# Chemoproteomic identification of a dipeptidyl peptidase 4 (DPP4) homolog in *Bacteroides thetaiotaomicron* important for envelope integrity and fitness

**DOI:** 10.1101/2022.07.25.501481

**Authors:** Laura J. Keller, Taylor H. Nguyen, Lawrence Liu, Markus Lakemeyer, Danielle J. Gelsinger, Rachael Chanin, Nhi Ngo, Kenneth M. Lum, Franco Faucher, Phillip Ipock, Micah J. Niphakis, Ami S. Bhatt, Anthony J. O’Donoghue, Kerwyn Casey Huang, Matthew Bogyo

**Affiliations:** Department of Chemical & Systems Biology, Stanford University School of Medicine, Stanford, CA 94305; Department of Bioengineering, Stanford University, Stanford, CA 94305; Skaggs School of Pharmacy and Pharmaceutical Sciences, University of California, San Diego, La Jolla, CA 92093; Department of Pathology, Stanford University School of Medicine, Stanford, CA 94305; Institute of Organic Chemistry and Macromolecular Chemistry, Friedrich-Schiller-Universität, Jena, Germany; Department of Genetics, Stanford University School of Medicine, Stanford, CA 94305; Lundbeck La Jolla Research Center, Inc., San Diego, CA 92121; Department of Chemistry, Stanford University, CA 94305; Divisions of Hematology and Blood & Marrow Transplantation, Department of Medicine, Stanford University, Stanford, CA 94305; Department of Microbiology and Immunology, Stanford University School of Medicine, Stanford, CA 94305; Chan Zuckerberg Biohub, San Francisco, CA 94158

**Keywords:** activity-based protein profiling, vancomycin, gut bacterial communities, synthetic communities, resource competition

## Abstract

Serine hydrolases play important roles in signaling and human metabolism, yet little is known about the functions of these enzymes in gut commensal bacteria. Using bioinformatics and chemoproteomics, we identify serine hydrolases in the gut commensal *Bacteroides thetaiotaomicron* that are specific to the Bacteroidetes phylum. Two are predicted homologs of the human protease dipeptidyl peptidase 4 (hDPP4), a key enzyme that regulates insulin signaling. Functional studies reveal that BT4193 is a true homolog of hDPP4 while the other is misannotated and is a proline-specific triaminopeptidase. We demonstrate that BT4193 is important for envelope integrity and is inhibited by FDA-approved type 2 diabetes drugs that target hDPP4. Loss of BT4193 reduces *B. thetaiotaomicron* fitness during *in vitro* growth within a diverse community. Taken together, our findings suggest that serine hydrolases contribute to gut microbiota dynamics and may be off-targets for existing drugs that could cause unintended impact on the microbiota.

## Introduction

The gut microbiota is a diverse microbial community whose dysbiosis has been implicated in a wide variety of conditions such as inflammatory bowel disease^1, 2^. The vast majority of studies aimed at understanding gut microbiota dynamics have focused on global analyses of genes, proteins, and metabolites using -omics methods^3^. While these methods have provided insight into the diversity of gut microbial communities and their metabolic outputs, they provide little information about the functional roles of specific enzymes and mechanisms that regulate these communities inside hosts. A better understanding of such regulatory mechanisms is a prerequisite for new avenues to develop therapeutics that modulate the gut microbiota. In addition, it is important to identify enzymes with homologous functions to human proteins, as they represent possible off-targets that need to be considered when developing new therapeutic agents. While existing approved drugs can have widespread impact on gut commensal bacteria^4–8^, the ability to predict these interactions and prioritize functionally relevant and druggable enzymes produced by commensal microbes in a high-throughput manner remains limited.

Activity-based protein profiling (ABPP) utilizes chemical probes to covalently modify enzymes in their endogenous environments. Integration with mass spectrometry-based proteomics allows class-wide identification of enzyme targets and measurement of enzyme activity across physiologically relevant conditions^9^. As a result, functionally relevant enzymes that are likely to be amenable to modulation by small molecules can be prioritized for characterization. In applying ABPP to study the gut microbiota, innovations such as the development of novel probes^10, 11^, integration of fluorescence-activated cell sorting^12^, and advances in metaproteomics analysis^13–15^ have improved efforts to identify functionally relevant enzymes in native microbiomes such as stool samples. However, the conservation of enzymes across closely related bacteria and limitations in metaproteomics techniques^16^ present obstacles to definitive determination of the function of specific enzymes using these top-down approaches.

Serine hydrolases are a broad class of proteases, lipases, and esterases that have a common catalytic serine residue. Studied extensively in mammalian systems, they play roles in proteolysis, metabolism, and cell signaling^17^, all of which likely affect the fitness of gut commensals in complex communities. Human serine hydrolases have been targeted with several FDA-approved therapeutics to treat diseases such as type 2 diabetes^18^, and recent efforts have identified serine hydrolases as novel drug targets for bacterial pathogens such as *Mycobacterium tuberculosis*^19–23^. However, little is known about the diversity of serine hydrolases in phylogenetically diverse gut commensal bacteria and their roles in interspecies communication and competition.

One of the most well-characterized human serine hydrolases is dipeptidyl peptidase 4 (hDPP4), which trims dipeptides from the N-terminus of oligopeptides with a preference for P1 proline or alanine residues. Ubiquitously expressed as a membrane protein that can also be shed to circulate in the bloodstream, this diaminopeptidase cleaves a number of peptide hormones, including glucagon-like peptide 1 (GLP1) and peptide YY, and leads to either degradation of the substrate or alteration of its receptor selectivity^24^. Because hDPP4-triggered degradation of GLP1, which induces the secretion of insulin, contributes to the regulation of glucose homeostasis^24^, multiple inhibitors of hDPP4 activity have been approved by the FDA for the treatment of type 2 diabetes^25^.

The bacterium *Bacteroides thetaiotaomicron* is prevalent and abundant in human gut microbiotas and has served as a model commensal due to its extensive capabilities for digesting complex polysaccharides^26, 27^. Early studies of fecal samples identified *Bacteroides* species as exhibiting high proteolytic activity^28, 29^, but the functions of these proteases have yet to be characterized. Recent work has implicated *Bacteroides*-derived proteases as key drivers of inflammatory bowel disease in patients with ulcerative colitis and in animal models of colitis^30^. In particular, dipeptidyl peptidases were identified to be abundant in *Bacteroides* species^30^, but the functions of these enzymes remains unclear.

Here, we utilize informatic predictions and a serine-reactive fluorophosphonate (FP) probe to profile serine hydrolases in *B. thetaiotaomicron* and identify six serine hydrolases annotated as dipeptidyl peptidases. Bioinformatic predictions indicated that these dipeptidyl peptidases are highly conserved within and restricted to the phylum Bacteroidetes. Two of these dipeptidyl peptidases, BT4193 and BT3254, have been annotated as putative homologs of hDPP4. We show biochemically that BT4193 is a bona fide functional homolog of hDPP4 that can be inhibited by hDPP4-targeted drugs, whereas BT3254 is a misannotated prolyl triaminopeptidase. We demonstrate that BT4193 plays a role in the maintenance of envelope integrity and in the establishment of *B. thetaiotaomicron* within multispecies synthetic communities *in vitro*. These findings highlight the potential for targeting serine hydrolases to regulate bacterial community dynamics and suggest that drugs designed for human enzymes may unintentionally alter gut microbiota composition.

## Results

### Identification of serine hydrolases in gut commensal bacterial species

Serine hydrolases have not been extensively studied in gut commensal bacteria, in part because they are difficult to identify bioinformatically due to the paucity of sequence homology and a large variety of protein folds^17^. Thus, we developed a pipeline to predict serine hydrolases by manually curating a list of protein family (Pfam) domains associated with serine hydrolases (Supplementary Fig. 1 and Supplementary Table 1), as defined by the MEROPS Peptidase Database^31^, ESTHER Database^32^, and known serine hydrolases identified in mammals^33^, plants^34^, bacteria^19, 21, 35, 36^, and archaea^37^ via ABPP with serine-reactive fluorophosphonate (FP)-based probes. Using these serine hydrolase-associated Pfam domains, we bioinformatically predicted the presence of serine hydrolases in 50 representative, phylogenetically diverse gut commensal bacterial strains (Fig. 1a, Supplementary Fig. 2, and Supplementary Tables 2-3). In clustering based on the number of serine hydrolases annotated with each Pfam domain, the human proteome appeared as an outgroup, having a distinct set of serine hydrolases primarily driven by the large number of trypsin-like (PF00089) serine hydrolases, and bacterial species clustered consistently by phylum. Furthermore, several types of serine hydrolases were predicted to be found exclusively in gut bacteria, highlighting the importance of characterizing serine hydrolases in these species. Similar to the human genome^17^, predicted serine hydrolases comprise 1-2.5% of each bacterial genome (Supplementary Fig. 3). Interestingly, bacteria from the Bacteroidetes phylum consistently are on the upper end of that range, in part driven by four serine hydrolase-associated Pfam domains that were markedly abundant in the Bacteroidetes phylum (Supplementary Fig. 4a). In a representative species, *Bacteroides thetaiotaomicron*, proteins annotated with two of these Bacteroidetes-specific Pfam domains, Sialic Acid-Specific Acetylesterase (PF03629) and GDSL-like Lipase (PF13472), are also predicted to be carbohydrate-processing enzymes (Supplementary Fig. 4b). As one of the hallmarks of bacteria in the Bacteroidetes phylum is their ability to digest complex polysaccharides, it is not surprising that these bacteria have a unique enrichment for serine hydrolases that are likely to process carbohydrates. The other two domains in this Bacteroidetes-specific signature are comprised of serine hydrolases with a Peptidase S9 (PF00326) or Peptidase S41 (PF03572) domain, suggesting that these peptidases may play unique and vital roles in these species.

**Figure 1:**
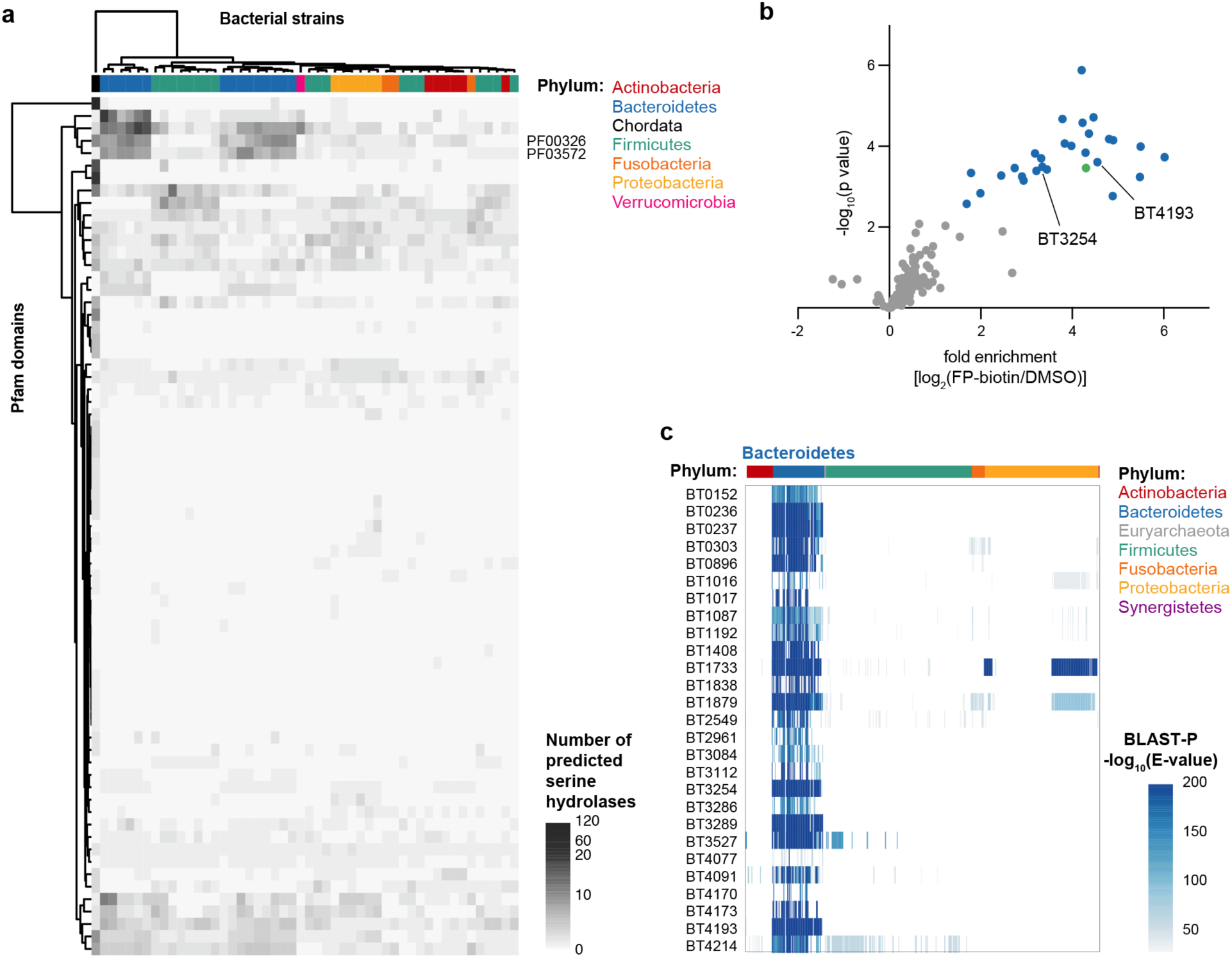
Bioinformatics and activity-based protein profiling of *B. thetaiotaomicron* identifies serine hydrolases specific to the Bacteroidetes phylum. **a,** Heatmap of number of proteins predicted to be serine hydrolases based on annotated Pfam domains in humans (Chordata) and 50 representative gut commensal bacterial species. Species are color-coded by phylum and hierarchically clustered by number of predicted serine hydrolases with each Pfam domain. Two Bacteroidetes-specific protease Pfam domains PF00326 (Peptidase S9) and PF03572 (Peptidase S41) are labeled on the right. **b,** Volcano plot of proteins labeled with fluorophosphonate (FP)- biotin relative to DMSO treatment in *B. thetaiotaomicron* VPI-5482. Significantly enriched hits are colored dark blue, an enriched ribosomal protein is colored light blue, and hDPP4 homologs BT4193 and BT3254 are labeled. c, Heatmap of homologs of *B. thetaiotaomicron* serine hydrolases across every bacterial strain in the Human Microbiome Project Reference Genomes for the Gastrointestinal Tract database via BLAST-P.

To determine whether the enzymes that make up the Bacteroidetes-specific peptidase signature are active and functionally relevant, we performed MS-based ABPP on *B. thetaiotaomicron* VPI-5482 by labeling intact bacteria with the fluorophosphonate probe, FP-biotin. We identified 27 FP-reactive enzymes active during *in vitro* monoculture growth (Fig. 1b and Supplementary Table 4), representing ∼1/3 of the total enzymes predicted by Pfam annotation. This incomplete coverage of the predicted serine hydrolases could be due to growth conditions impacting enzyme expression or activity, or the reactivity or permeability of the probe. A Protein-BLAST search against gastrointestinal tract reference genomes from the Human Microbiome Project^38^ confirmed that most *B. thetaiotaomicron* serine hydrolases identified by MS-ABPP are conserved and specific to the Bacteroidetes phylum (Fig. 1c). Six of the identified active serine hydrolases are in the Peptidase S9 or Peptidase S41 families, including BT3254 and BT4193, which are both annotated as putative homologs of hDPP4. Transposon insertions in BT3254 and BT4193 caused fitness defects in monocolonized gnotobiotic mice in a pooled transposon-site sequencing (Tn-Seq) assay^39^, supporting the relevance of these enzymes for growth *in vivo*. However, the function of hDPP4-like enzymes in a commensal bacterium residing in the gut lumen is unclear. Hence, we sought to confirm whether BT3254 and BT4193 are indeed hDPP4 homologs, determine their sensitivity to current FDA-approved hDPP4 inhibitors, and ascertain their physiological role in *B. thetaiotaomicron*.

### BT4193 is a functional homolog of hDPP4

By sequence identity, homologs of the hDPP4 enzyme are predicted to be highly conserved and exclusively found within the Bacteroidetes phylum (Fig. 2a and Supplementary Table 6). Crystallographic studies of hDPP4 have highlighted several sets of residues key to the activity of the protease: the catalytic triad (Ser630/Asp708/His740), the hydrophobic S1 binding pocket (Tyr631/Val656/Trp659/Tyr662/Try666/Val711) that confers P1 specificity, the double-Glu motif that recognizes the free N-terminus of the substrate (Glu205/Glu206), and the Arg125 residue that hydrogen bonds with Glu205 and the P2 carbonyl group^40–42^. Based on sequence alignments, both BT4193 and BT3254 have the conserved catalytic triad and Tyr662 and Tyr666 residues that ring- stack with the P1 Pro canonically found in hDPP4 substrates, as well as similar residues forming the rest of the S1 binding pocket (Fig. 2b and Supplementary Fig. 5-6). However, only BT4193 has the double-Glu motif and a corresponding Arg residue to recognize and stabilize the free N-terminus of a peptide substrate. Structural homology modeling confirmed the alignment of these residues and suggested that the orientation of the Gln-Glu motif in the predicted structure of BT3254 may not be capable of forming salt bridges with the N-terminus of a substrate (Fig. 2c).

**Figure 2:**
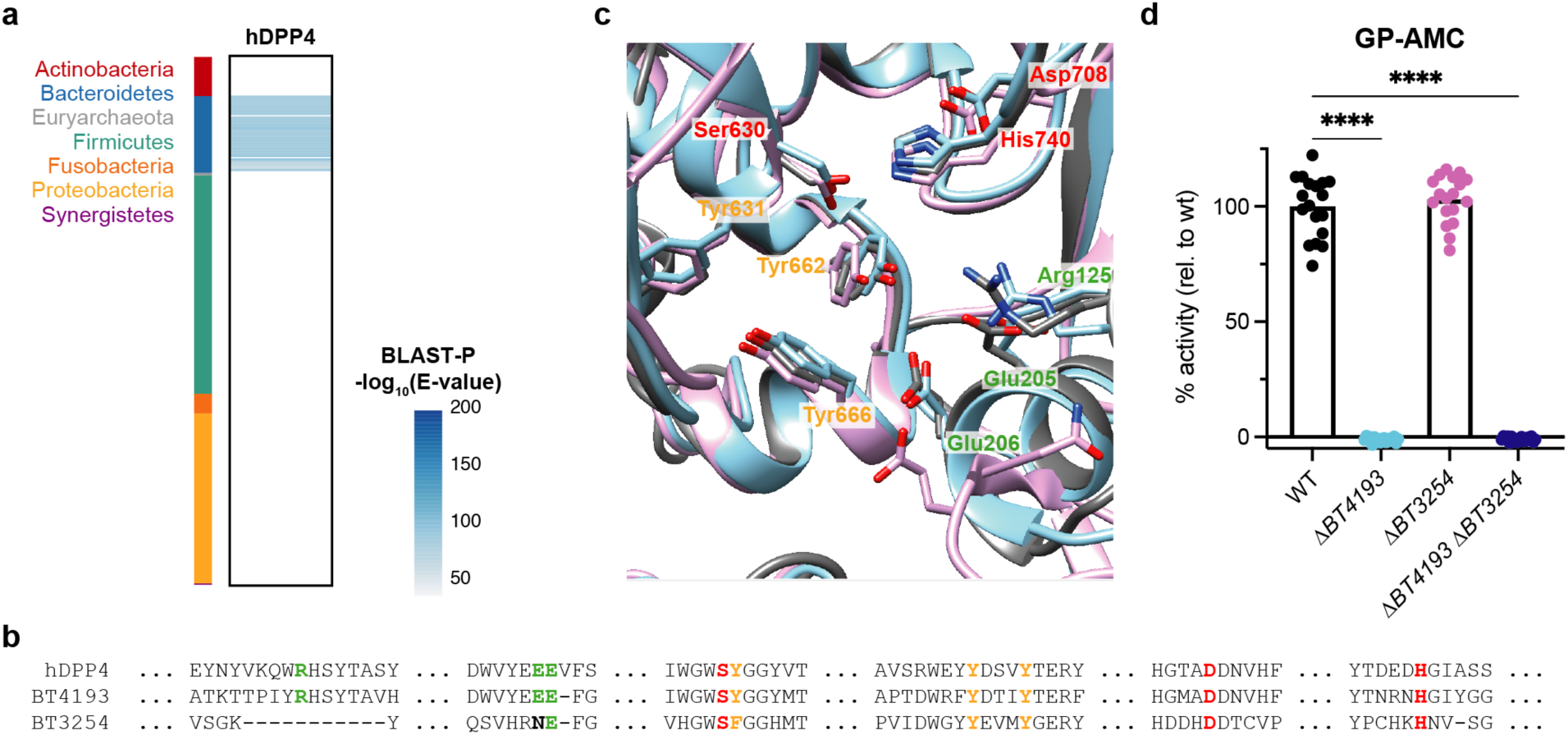
BT4193 is a functional homolog of human DPP4 (hDPP4). **a**, Heatmap of the phylogenetic distribution of hDPP4 homologs across every bacterial strain in the Human Microbiome Project Reference Genomes for the Gastrointestinal Tract database via BLAST-P. **b**, Multiple sequence alignment of hDPP4, BT4193, and BT3254. Residues relevant to hDPP4 function are colored red (catalytic triad, Ser630/Asp708/His740), gold (S1 binding pocket, Tyr631/Tyr662/Try666), or green (free amine-stabilizing residues, Arg125/Glu205/Glu206) along with their corresponding residues in BT4193 and BT3254. **c**, Structural homology modeling alignment of hDPP4 (grey; PDB: 1J2E, chain B) and AlphaFold^86^-predicted structures of BT4193 (blue) and BT3254 (pink). Side chains of residues relevant for hDPP4 function highlighted in (**a**) and their corresponding residues in BT4193 and BT3254 are shown, with numbering based on hDPP4. **d**, Quantification of cleavage of the canonical hDPP4 fluorogenic peptide substrate GP- AMC in *B. thetaiotaomicron* lysate (ex/em: 380/460 nm). Velocities of substrate cleavage are normalized to wild-type (WT) velocity. Statistical significance was determined using a one-way ANOVA test with posthoc Dunnett’s multiple comparisons tests compared to wild-type (****, p < 0.0001).

To determine whether this sequence and structural homology confers functional homology, we utilized the fluorogenic peptide substrate GlyPro-AMC, which is traditionally used to measure hDPP4-like activity via proteolytic release of fluorescent 7-amino-4-methylcoumarin (AMC)^43^.

Indeed, wild-type *B. thetaiotaomicron* lysate cleaved the substrate, suggesting that *B. thetaiotaomicron* has one or more proteases with hDPP4-like activity (Fig. 2d). These results corroborate previously reported evidence that the GlyPro-*p*-nitroanilide substrate is effectively cleaved by *B. thetaiotaomicron*, as well as other species in the Bacteroidetes phylum^44–46^ and the mouse cecum more generally^47^. To ascertain whether the putative hDPP4 homologs BT4193 and BT3254 contribute to this hDPP4-like activity, we generated single knockouts as well as the double knockout of these genes. Interestingly, GlyPro-AMC processing activity was completely eliminated in the Δ*BT4193* and Δ*BT4193* Δ*BT3254* strains but not in the Δ*BT3254* strain, suggesting that BT4193 is the only functional homolog of hDPP4 with proline dipeptidase activity (Fig. 2d). We also recombinantly expressed BT4193 and BT3254 and confirmed that only BT4193 can cleave the GlyPro-AMC substrate (Supplementary Fig. 7).

### BT4193 is targeted by hDPP4 inhibitors

As BT4193 is a functional homolog of hDPP4, we wanted to determine if it could be inhibited by compounds targeting the human enzyme, including talabostat and several members of the gliptin class of FDA-approved drugs for the treatment of type 2 diabetes (Fig. 3a). Pretreatment with two covalent inhibitors, the boronic acid-containing talabostat and the nitrile-containing saxagliptin, specifically and dose-dependently blocked labeling of BT4193 with FP-alkyne as measured by competitive gel-based ABPP (Fig. 3b). Additionally, talabostat and saxagliptin inhibited cleavage of the canonical hDPP4 substrate GP-AMC in wild-type *B. thetaiotaomicron* lysate, as did the non-covalent inhibitors sitagliptin and linagliptin, although to a lesser extent (Fig. 3c). The relative potency of these four inhibitors against BT4193 was confirmed using recombinantly expressed enzyme, with talabostat and saxagliptin having low nanomolar potency for both hDPP4 and BT4193 and sitagliptin and linagliptin being >100- and >1000-fold weaker inhibitors against BT4193 when compared with hDPP4 (Fig. 3d and Supplementary Fig. 8). None of these inhibitors were active against recombinant BT3254 (Supplementary Fig. 8).

**Figure 3:**
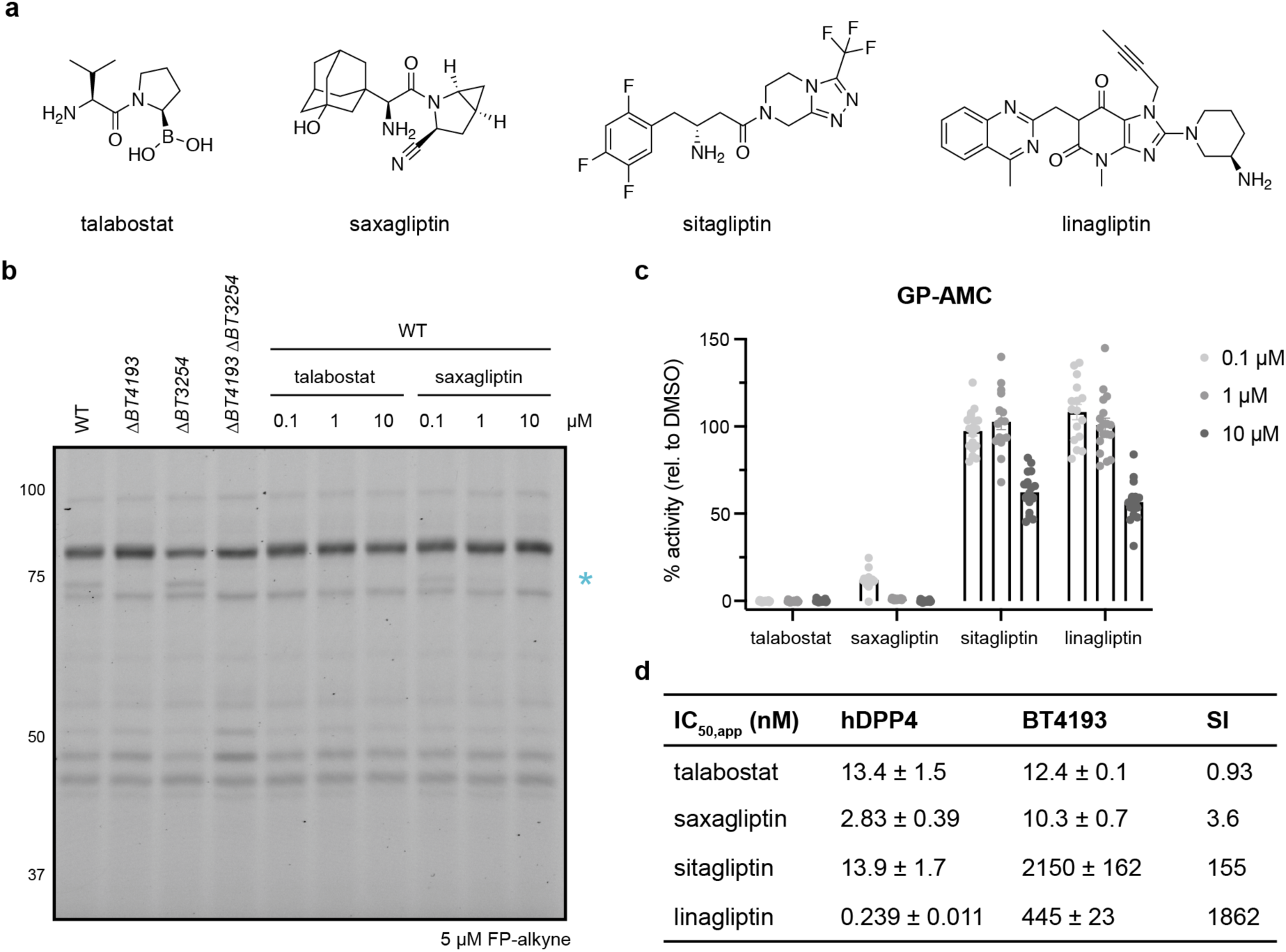
Human DPP4 inhibitors target BT4193. **a**, Structure of hDPP4 inhibitors. **b**, Representative in-gel fluorescence labeling of intact *B. thetaiotaomicron* strains. Pretreatment with talabostat or saxagliptin competes labeling of only BT4193 by FP-alkyne in wild-type (WT) *B. thetaiotaomicron*. Asterisk indicates the location of the band corresponding to BT4193. **c**, Quantification of percent inhibition of GP-AMC cleavage in WT *B. thetaiotaomicron* lysate (ex/em: 380/460 nm) after pre-treatment with inhibitor for 30 min at 37 °C. Activity was normalized to DMSO treatment. **d**, Apparent IC_50_ values of recombinant hDPP4 and BT4193 after treatment with inhibitor for 30 min prior to measuring activity via GP-AMC cleavage. Activity was normalized to DMSO treatment and fit with dose-dependent four parameter inhibition. Selectivity index (SI) is the ratio of BT4193 to hDPP4 apparent IC_50_ values.

### BT4193 and BT3254 are prolyl aminopeptidases

To comprehensively profile the substrate specificity of BT4193 and BT3254 in an unbiased manner, we applied both enzymes to multiplex substrate profiling by mass spectrometry (MSP-MS), in which recombinant enzyme is added to a library of synthetic tetradecapeptides whose cleavage is kinetically monitored by LC/MS^48, 49^. BT4193 primarily trimmed dipeptides from the N-terminus of the substrates, especially after Pro or Ala residues (Fig. 4a and Supplementary Fig. 9). Interestingly, BT3254 also exhibited aminopeptidase activity but trimmed tripeptides with P1 Pro or Ala, suggesting that it is a triaminopeptidase instead of a diaminopeptidase (Fig. 4a).

**Figure 4:**
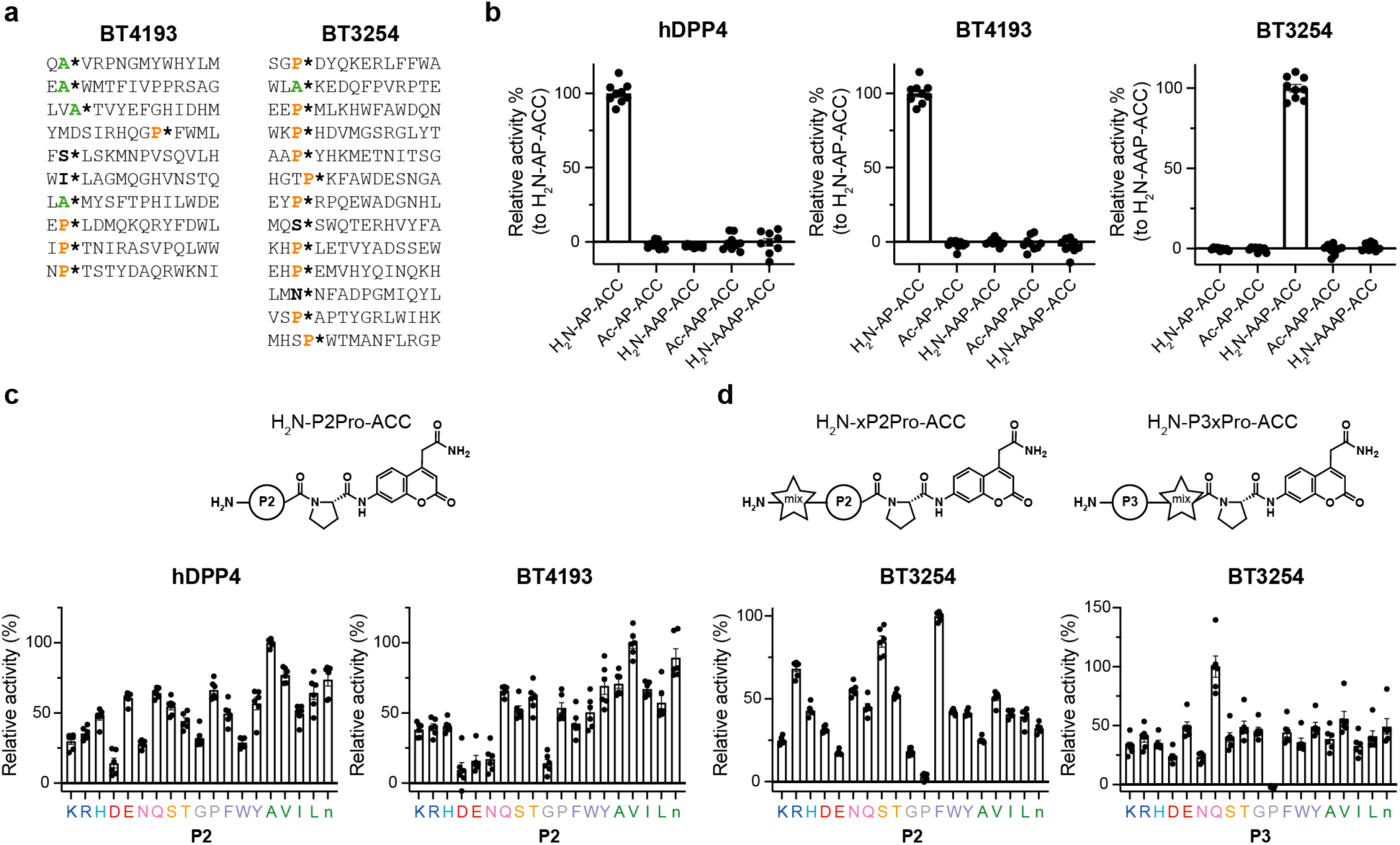
Substrate specificity of BT4193 and BT3254 is driven by P1 Pro. **a**, Substrates cleaved by recombinant BT4193 and BT3254 from multiplex substrate profiling by mass spectrometry (MSP-MS), defined as significant fold-change between 0 h and 4 h of incubation. Cleavage sites are denoted by an asterisk and P1 Pro and Ala residues are colored. **b**, Quantification of cleavage of fluorogenic peptide substrates by recombinant BT4193 and BT3254 (ex/em: 355/460 nm). Initial velocities were normalized to the best substrate (H_2_N-AP-ACC for BT4193, H_2_N-AAP-ACC for BT3254). BT4193 is a diaminopeptidase like hDPP4 and BT3254 is a triaminopeptidase. **c**, Quantification of cleavage velocity of a P2Pro-ACC positional scanning library relative to the peptide with the highest turnover for recombinant hDPP4 and BT4193. **d**, Quantification of cleavage velocity of xP2Pro-ACC and P3xPro-ACC positional scanning libraries by recombinant BT3254 relative to the peptide in the library with the highest turnover.

To confirm these activities, we synthesized a targeted library of fluorogenic peptide substrates that all contained proline in the P1 position and a 7-amino-4-carbamoylmethylcoumarin (ACC) reporter group. The substrates were various lengths and with or without an acetyl cap to determine the necessity of the free N-terminus. In agreement with the results from our MSP-MS screen, we found that BT4193, like hDPP4, is a diaminopeptidase while BT3254 is a triaminopeptidase (Fig. 4b). All three enzymes required substrates to have a free N-terminus. These substrate preferences were confirmed *in vitro* with *B. thetaiotaomicron* lysate (Supplementary Fig. 10). The preference for P1 Pro by both enzymes was confirmed with a small P1 library of fluorogenic peptide substrates (Supplementary Fig. 11), which matched the substrate specificity of hDPP4^50^.

To more comprehensively map substrate specificity at the P2 and P3 positions for each protease, we synthesized a series of positional scanning synthetic combinatorial libraries of P1 Pro fluorogenic peptides, in which each position is systematically scanned through each natural amino acid (excluding cysteine and methionine but including norleucine) and the remaining positions are composed of an isokinetic mixture of those amino acids^51^. Screening the dipeptide library against hDPP4 and BT4193 demonstrated a highly overlapping preference for bulky, hydrophobic P2 residues, although most residues could be accommodated by both the bacterial and human enzymes (Fig. 4c and Supplementary Fig. 12). Screening of two tripeptide libraries to scan the P2 and P3 position preference of BT3254 also confirmed that the majority of the residues could be tolerated in either position (Fig. 4d). Together, these results suggest that the specificities of BT4193 and BT3254 are driven primarily by the P1 proline residue and that both enzymes are prolyl aminopeptidases.

### BT4193 is important for cell envelope integrity

To characterize the functions of BT4193 and BT3254, we first sought to identify the localization of these proteases in *B. thetaiotaomicron* cells. Using SignalP^52^, both enzymes are predicted to have a type I Sec signal peptide cleavage site without a lipoprotein motif, and are predicted to be outer membrane-associated via CELLO^53^. Additionally, previous studies suggest that these enzymes are not surface-exposed^27^, and proteomic studies in the closely related *Bacteroides fragilis* suggest that homologs of BT4193 and BT3254 are not likely to be secreted^54^. Furthermore, we did not detect activity of either BT4193 or BT3254 (as indicated by turnover of their substrates AP-ACC and AAP-ACC, respectively) in *B. thetaiotaomicron* culture supernatants (Supplementary Fig. 13). Thus, we conclude that BT4193 and BT3254 are likely localized in the periplasm.

Based on the predicted periplasmic localization, we hypothesized that these enzymes may be important for envelope integrity. Previous fitness measurements of a pooled transposon library demonstrated that transposon insertions in *BT4193* have increased susceptibility to a number of cell envelope stressors including the antibiotic vancomycin and the bile salt deoxycholic acid^55^. We confirmed that *B. thetaiotaomicron* cannot grow in the presence of vancomycin when *BT4193* has been deleted (Fig. 5a-b), and this BT4193-dependent susceptibility to vancomycin is dose-dependent (Supplementary Fig. 14). Additionally, the Δ*BT4193* and Δ*BT4193* Δ*BT3254* strains exhibited increased susceptibility to deoxycholic acid (Fig. 5c-d and Supplementary Fig. 15). These results suggest that BT4193 plays a pivotal role in conferring resistance to cell envelope stressors and likely serves some function in maintaining outer membrane and/or cell wall integrity.

**Figure 5:**
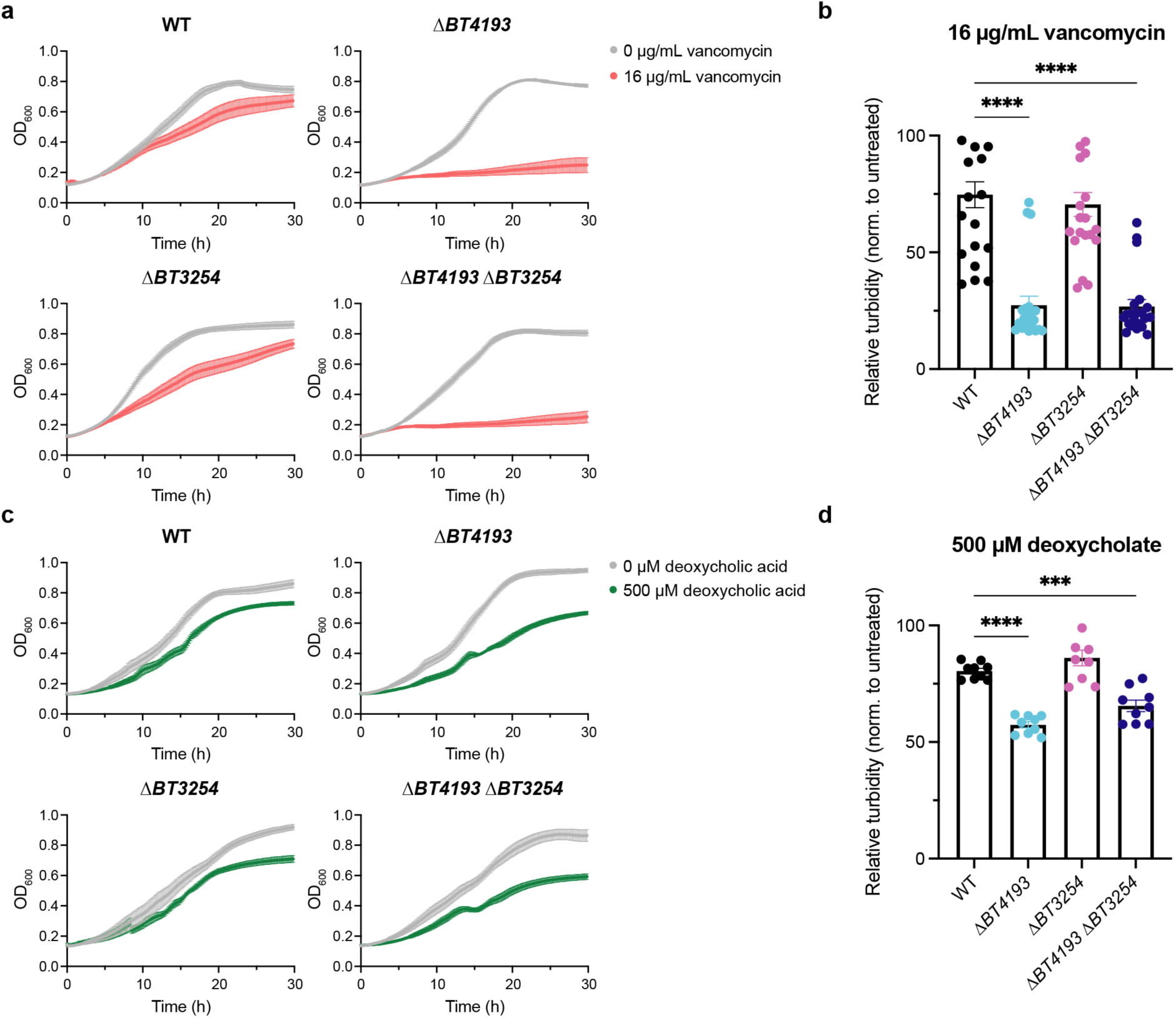
BT4193 confers resistance to envelope stressors vancomycin and deoxycholic acid. **a**, Growth of *B. thetaiotaomicron* strains during vancomycin treatment, as measured by OD_600_. Deletion of *BT4193* severely impacts fitness in the presence of vancomycin. **b**, Quantification of relative turbidity, as measured by the OD_600_ of vancomycin-treated bacteria normalized to vehicle- treated bacteria, at 20 h. **c**, Growth of *B. thetaiotaomicron* strains during deoxycholic acid treatment, as measured by OD_600_. Deletion of *BT4193* impacts fitness in the presence of deoxycholic acid. **d**, Quantification of relative turbidity, as measured by the OD_600_ of deoxycholic acid-treated bacteria normalized to vehicle-treated bacteria, at 20 h. Statistical significance was determined using a one-way ANOVA test with posthoc Dunnett’s multiple comparisons tests compared to wild-type (***, p < 0.001; ****, p < 0.0001).

### BT4193 is important for community fitness

Previous studies demonstrated that treatment of a diabetic mouse model with saxagliptin leads to depletion of bacteria from the Bacteroidetes phylum in the gut^56^. However, this model does not allow for deconvolution of the effect of inhibiting the host DPP4 enzyme versus bacterial DPP4 enzymes nor the distinction between host-bacteria and bacteria-bacteria interactions. Thus, to determine whether BT4193 plays a role in the establishment of *B. thetaiotaomicron* in a diverse complex bacterial community, we utilized a synthetic *in vitro* community of 14 stool-derived gut commensals that can be stably maintained in rich media and that reflects the diversity of bacterial families in the host from which the isolates were obtained^57^ (Fig. 6a). The 14 members comprise several *Bacteroides* species and a *Parabacteroides* species in the Bacteroidetes phylum, but not *B. thetaiotaomicron*. The single and double knockout strains did not exhibit any growth defects during monoculture growth in the rich Brain-Heart Infusion (BHI) medium (Fig. 6b). When any of our *B. thetaiotaomicron* strains were added individually to the full 14-member community, the overall community structure was not dramatically altered (Supplementary Fig. 16). However, the relative abundance of the Δ*BT4193* and Δ*BT4193* Δ*BT3254* strains was significantly lower than that of wild-type (Fig. 6c). To test the extent to which the lower fitness of the knockouts was sensitive to the nutrient environment, we passaged the same full communities in modified Gifu Anaerobic Medium (mGAM), another rich medium that supports rapid growth of *Bacteroides* species. Similar to BHI, the *ΔBT4193* and *ΔBT4193 ΔBT3254* strains had lower relative abundance than wild-type in mGAM (Fig. 6d), indicating that the hDPP4 homolog BT4193 generally plays an important role in the fitness of *B. thetaiotaomicron* within diverse bacterial communities.

**Figure 6:**
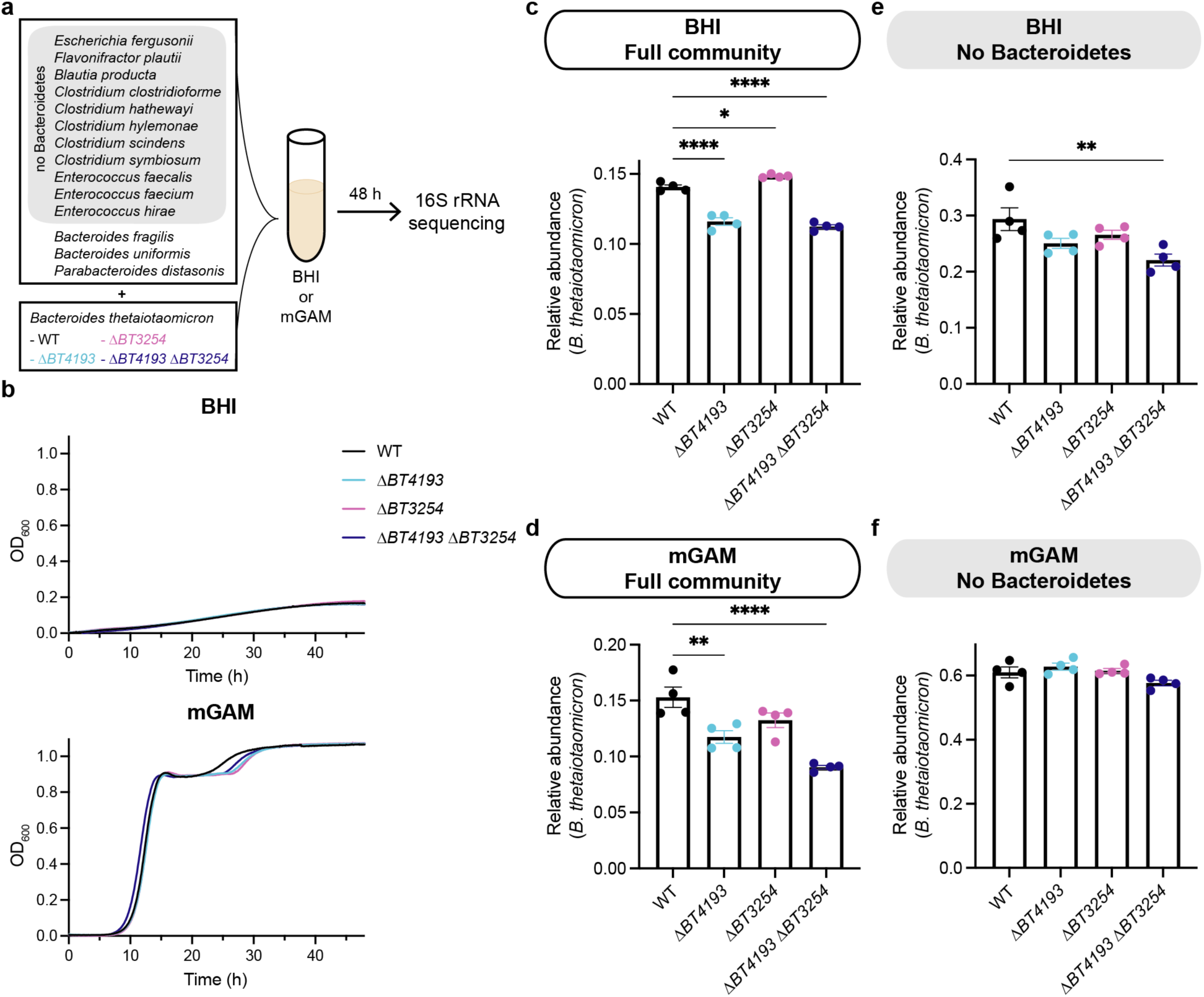
BT4193 impacts *B. thetaiotaomicron* during *in vitro* growth in a diverse community. **a**, Schematic of synthetic community experiments. Fourteen human gut commensal species were co-cultured with one of four *B. thetaiotaomicron* strains (wild-type (WT), Δ*BT4193*, Δ*BT3254*, Δ*BT4193* Δ*BT3254*) and co-cultured for 48 h in BHI or modified Gifu Anaerobic Medium (mGAM). The relative abundance of each species was quantified using 16S rRNA sequencing. To determine the contribution of other Bacteroidetes species, a community with only the 11 non-Bacteroidetes species along with one of the *B. thetaiotaomicron* strains (no Bacteroidetes) was also used. **b**, Growth of *B. thetaiotaomicron* strains in BHI and mGAM, as measured by OD_600_. **c**, Quantification of the relative abundance of *B. thetaiotaomicron* within the full community after 48 h of co-culturing in BHI. **d**, Quantification of the relative abundance of *B. thetaiotaomicron* within the full community after 48 h of co-culturing in mGAM. **e**, Quantification of the relative abundance of *B. thetaiotaomicron* within the no Bacteroidetes community after 48 h of co-culturing in BHI. **f**, Quantification of the relative abundance of *B. thetaiotaomicron* within the no Bacteroidetes community after 48 h of co-culturing in mGAM. Statistical significance was determined using a one-way ANOVA test with posthoc Dunnett’s multiple comparisons tests compared to wild-type (*, p < 0.05; **, p < 0.01; ****, p < 0.0001).

Since the mutants did not exhibit a growth defect in monoculture (Fig. 6b), we hypothesized that interspecies interactions such as nutrient competition were an important component of the fitness advantage conferred by BT4193. To test this hypothesis, we removed the other Bacteroidetes phylum members (*Bacteroides fragilis, Bacteroides uniformis,* and *Parabacteroides distasonis*), surmising that closely related species would have the largest niche overlap with *B. thetaiotaomicron*. In BHI, the fitness defect of the *ΔBT4193* strain was dependent on the other Bacteroidetes species (Fig. 6e), while in mGAM both the *ΔBT4193* and *ΔBT4193 ΔBT3254* mutants were able to grow to similar levels as wild-type in the absence of other Bacteroidetes (Fig. 6f). Thus, the microbial DPP4 protease can impact the assembly of diverse communities through modulation of interspecies nutrient competition.

## Discussion

Serine hydrolases play critical roles in mammalian pathophysiology including inflammation, blood clotting, and neurotransmitter signaling. As a result, they have been successfully targeted with small molecule drugs for treatment of a wide variety of human diseases^18^. However, little is known about the diversity and function of this enzyme superfamily in the phylogenetically divergent bacteria that comprise the human gut microbiota. Our bioinformatic approach should enable identification of predicted serine hydrolases more broadly in the gut microbiota, after which predictions can be functionally validated using chemoproteomics. Our analysis suggests that there are several types of bacterial serine hydrolases that are not found in humans (Fig. 1a), indicating that simple Protein BLAST analyses based on known serine hydrolases in humans will fail to identify many of these enzymes in bacteria.

Our chemoproteomic profiling of *B. thetaiotaomicron* led to our focus on BT4913 and BT3254, but we additionally identified several other putative dipeptidyl peptidases. A homologous network of aminopeptidases has been characterized in the phylogenetically related oral pathogen *Porphyromonas gingivalis*^58^. This asaccharolytic bacterium relies on these peptidases to generate di- and tripeptides from exogenous protein that can be more easily imported into the cytoplasm for protein catabolism^46, 58, 59^. Unlike *P. gingivalis*, the hallmark of gut commensal bacteria from the Bacteroidetes phylum are their carbohydrate utilization capabilities, so conservation of these aminopeptidases throughout the phylum and not just in asaccharolytic species suggests a role that extends beyond protein catabolism. Given that the other predicted dipeptidases did not exhibit fitness defects *in vivo* in monocolonized mice^39^, future investigation will be required to clarify how these enzymes interact with the larger network of dipeptidases in *B. thetaiotaomicron*.

The increased sensitivity of Δ*BT4193* cells to the antibiotic vancomycin (Fig. 5a-b) supports a role for BT4193 in regulating the overall integrity of the cell envelope. Vancomycin, which prevents cross-linking of nascent peptidoglycan chains in the cell wall, is traditionally thought to target Gram-positive bacteria. Gram-negative bacteria such as *B. thetaiotaomicron* have an outer membrane surrounding the peptidoglycan layer that can act as a barrier to vancomycin and hence confer resistance^60^. Mutants in *Escherichia coli*^61^ and *Caulobacter crescentus*^62^ that modulate outer membrane permeability have been shown to have increased sensitivity to both vancomycin and deoxycholic acid, similar to the phenotypes of strains lacking *BT4193*. Relative to other Gram-negative bacteria from the Proteobacteria phylum such as *E. coli*, little is known about the architecture of the cell envelope in *Bacteroides* species, although differences in the lipid A component of lipopolysaccharide molecules^63, 64^ impact resistance to the cationic antimicrobial peptide polymyxin B^65^. Our results suggest that aminopeptidases may regulate envelope integrity by trimming peptide substrates important in periplasmic homeostasis. These substrates could include oligopeptides serving analogous functions to Braun’s lipoprotein found in *E. coli*, which links the cell wall to the outer membrane^66^. Future efforts to map the endogenous substrates of BT4193 using proteomic methods may provide molecular insight into how BT4193 affects membrane permeability.

Within a diverse community, the effects of deleting *BT4193* and/or *BT3254* on *B. thetaiotaomicron* relative abundance (Fig. 6c-f) and overall community composition (Supplementary Fig. 16) are subtle, as one would expect since much of *B. thetaiotaomicron’s* niche is independent of the activity of these proteases. Nonetheless, BT4193 still has a significant impact on the establishment of *B. thetaiotaomicron* within a community. The dependence of the fitness of *ΔBT4193* and Δ*BT4193* Δ*BT3254* mutants on other community members (Fig. 6e,f) suggests that these proteases play key roles in competition for a niche that is shared by other Bacteroidetes members. The difference in fitness defects of the Δ*BT4193* Δ*BT3254* mutant between BHI (Fig. 6e) and mGAM (Fig. 6f) highlights the nutrient dependence of the competitive landscape and suggests that the more favorable growth of *Bacteroides* in mGAM may reflect greater competition, while the fitness defect of the Δ*BT4193* Δ*BT3254* mutant in BHI even in the absence of other Bacteroidetes likely reflects competition with members of other phyla^67^. The ability to modulate the levels of *B. thetaiotaomicron* through BT4193 has exciting implications for subtle community engineering. Our current tools for editing community composition such as antibiotics and bacteriophages typically have more drastic effects on community diversity and composition. More broadly, our results demonstrate that bacterial hDPP4 homologs are important targets whose disruption can affect community composition. As a result, our findings add to a growing body of evidence that human drugs can have unintentional side effects on the gut microbiota^4–6^.

The role of BT4193 in *B. thetaiotaomicron* physiology and overall fitness is particularly important considering 10-20% of patients with diabetes in the US take hDPP4 inhibitors^68, 69^. As a result, any cross-reactivity of these drugs with bacterial enzymes could impact microbial communities in the gut. We assessed the off-target effects of several FDA-approved inhibitors, saxagliptin (Onglyza), sitagliptin (Januvia), and linagliptin (Tradjenta) on BT4193 (Fig. 3). The potency of these inhibitors against BT4193 matched their known selectivity for hDPP4 over other human prolyl oligopeptidases^70^ and was similar to their efficacy against the *P. gingivalis* hDPP4 homolog^71^. The inhibitors sitagliptin and linagliptin gain their selectivity by occupying the S_2_ extensive and S_1_′/S_2_′ pockets on hDPP4, respectively^72^, which may also be the case for BT4193. While these inhibitors are primarily cleared renally, 20% of saxagliptin^73^ and 10% of sitagliptin^74^ are eliminated fecally, suggesting that *B. thetaiotaomicron* may be exposed to substantial concentrations of these drugs in the gut. We have shown that loss of BT4193 decreases *B. thetaiotaomicron* relative abundance within a diverse bacterial community (Fig. 6c-f), so the depletion of bacteria from the Bacteroidetes phylum upon saxagliptin treatment seen previously in a diabetic mouse model^56^ may be at least in part due to inhibition of bacterial DPP4 enzymes. While large cohort studies of diabetic patients characterizing the composition of their gut microbiota have often yielded conflicting results, a meta-analysis revealed that *Bacteroides* species are negatively associated with type 2 diabetes and they can be used as probiotics to improve glucose intolerance in mouse models of diabetes^75^. Thus, further loss of beneficial bacteria from the phylum Bacteroidetes upon hDPP4 inhibitor treatment may unintentionally exacerbate the dysbiosis associated with diabetes.

Previous studies identified DPP4 activity in the cecum of germ-free mice, likely from host DPP4^76^, and FP-reactive hDPP4 can be found in human fecal samples^15^. Interestingly, some of the canonical peptide hormone substrates of hDPP4 are secreted into the gut lumen—peptide YY is primarily secreted luminally^77^ and plays a role in regulating ion and water absorption^78^, and even GLP-1 has been shown to be secreted apically in *ex vivo* intestinal tissue.^79^ The physiological impact of these luminally secreted peptide hormones and hDPP4 is not yet understood, but will likely contribute to the multi-faceted role of bacterial DPP4 enzymes in the complex community within the gut lumen.

Overall, our study demonstrates how a combination of bioinformatics and chemoproteomics can be utilized to identify and prioritize physiologically relevant, druggable enzymes in gut commensal bacteria. This approach led to the successful characterization of two proteases and the discovery of a novel role for a hDPP4 homolog in regulating the integrity of the bacterial cell envelope. Moreover, our findings highlight the power of a bottom-up approach to discover molecular regulators of the dynamics of microbial communities. This approach can be similarly applied to other key gut commensal bacterial species. This work has identified gut bacterial serine hydrolases as potential drug targets that could be exploited to alter the composition of bacterial communities. Finally, it underscores the need to broadly understand the off-target effects of existing drugs on bacterial enzymes and how they impact the gut microbiota.

## Methods

### Chemicals

GlyPro-AMC was purchased from Santa Cruz Biotechnology. Z-GlyPro-AMC was purchased from Bachem. Recombinant hDPP4 was purchased from Sino Biological. Talabostat mesylate and sitagliptin were purchased from AChemBlock. Saxagliptin and linagliptin were purchased from Carbosynth. Fmoc-protected amino acids (Fmoc-Ala-OH, Fmoc-Asp(tBu)-OH, Fmoc-Glu(tBu)- OH, Fmoc-Phe-OH, Fmoc-Gly-OH, Fmoc-His(Trt)-OH, Fmoc-Ile-OH, Fmoc-Lys(Boc)-OH, Fmoc-Leu-OH, Fmoc-Nle-OH, Fmoc-Asn(Trt)-OH, Fmoc-Pro-OH, Fmoc-Gln(Trt)-OH, Fmoc-Arg(Pbf)-OH, Fmoc-Ser(But)-OH, Fmoc-Thr(But)-OH, Fmoc-Val-OH, Fmoc-Trp(Boc)-OH, Fmoc-Tyr(tBu)-OH) were purchased from Advanced ChemTech, Chem-Impex, AAPPTec, and Novabiochem. Vancomycin hydrochloride from *Streptomyces orientalis* was purchased from Sigma. Deoxycholic acid sodium salt was purchased from Fluka. FP-TMR and FP-biotin have been previously reported^36^.

### Bacterial strains and growth conditions

All bacterial strains and plasmids used in this study are listed in Supplementary Table 7. For mass spectrometry-based activity-based protein profiling and cloning of genomic DNA, *B. thetaiotaomicron* VPI-5482 was utilized, and for the remainder of the biological characterization assays, wild-type *B. thetaiotaomicron* refers to the parent strain *B. thetaiotaomicron tdk*. All *B. thetaiotaomicron* strains were grown anaerobically (90% N_2_, 5% CO_2_, 5% H_2_) at 37 °C in either Brain Heart Infusion (BHI) media or Varel-Bryant minimal media (Supplementary Table 8). All *E. coli* strains were grown aerobically at 37 °C in Luria Broth (LB) supplemented with 100 µg/mL carbenicillin as necessary for *B. thetaiotaomicron* mutagenesis and LB for protein expression.

Clean deletions were generated in *B. thetaiotaomicron* as previously described^80^, growing in BHIS. Briefly, suicide plasmids containing the flanking regions of the *B. thetaiotaomicron* genes *BT3254* and *BT4193* were amplified and assembled into pExchange-*tdk* using a Gibson Assembly Cloning Kit (New England Biolab). Plasmid inserts were verified by Sanger sequencing and the resulting plasmids (pRBC9 and pRBC8, respectively) were propagated in DH5α λ*pir*.

To generate *B. thetaiotaomicron* Δ*BT4193* and *B. thetaiotaomicron* Δ*BT3254*, the suicide plasmids pRBC8 and pRBC9, respectively were conjugated into *B. thetaiotaomicron tdk* using S17-1 λ*pir* as the conjugative donor strain. Exconjugants with chromosomally integrated plasmids were recovered on BHIS plates containing 200 μg/mL gentamycin and 25 μg/mL erythromycin. Second crossover events were selected using FudR plates (BHIS plates supplemented with 200 μg/mL 5- fluoro-2-deoxy-uridine). Deletion of the target gene was confirmed by PCR. *B. thetaiotaomicron tdk* Δ*BT4193* Δ*BT3254* was generated by conjugating pRBC8 into *B. thetaiotaomicron tdk* Δ*BT3254* and following the selection steps listed above.

### Bioinformatic analysis of serine hydrolases

For the prediction of serine hydrolases, a list of serine hydrolase-associated Pfam domains was generated by compiling the Pfam domains associated with each family of serine peptidases in the MEROPS database (https://www.ebi.ac.uk/merops/), each family of alpha/beta-hydrolases in the ESTHER database (https://bioweb.supagro.inrae.fr/ESTHER/general?what=index), and fluorophosphonate (FP)-reactive proteins identified by activity-based protein profiling in humans^33^, *Arabidopsis thaliana*^34^, *Mycobacterium tuberculosis*^19^, *Staphylococcus aureus*^21^, *Staphylococcus epidermidis*^36^, *Vibrio cholerae*^35^, *Sulfolobus solfataricus*, and *Haloferax volcanii*^37^. Each proteome of interest was downloaded from UniProt as a fasta file in March 2022. Proteomes were annotated with Pfam domains locally using pfam_scan.pl (http://ftp.ebi.ac.uk/pub/databases/Pfam/, November 2021). From each annotated proteome, proteins annotated with serine hydrolase-associated Pfam domains with an E-value < 0.0001 were considered serine hydrolases. Identified serine hydrolases were then tallied by Pfam domain, and the heatmap of the number of each type of serine hydrolase was generated using the *pheatmap* package with clustering in R. Predicted serine hydrolases from *B. thetaiotaomicron* were compared to its carbohydrate-active enzymes annotated on the CAZY database (http://www.cazy.org/) using the *eulerr* package in R.

To assess the conservation of active serine hydrolases identified in *B. thetaiotaomicron* throughout the gut microbiota, the pep.faa file for reference organisms isolated from the gastrointestinal tract as part of the NIH Human Microbiome Project (https://www.hmpdacc.org/hmp/HMRGD/, retrieved December 2019) was downloaded. This faa file was converted into a BLAST database locally using the *makeblastdb* application from NCBI BLAST+ v2.7.1. A protein fasta file of *B. thetaiotaomicron* active serine hydrolases was then queried against this database using the *blastp* application with a minimum e-value of 1e-30. These homologs were then sorted by bacterial strain of origin and query protein sequence, and -log_10_(E-value) was calculated for each homolog. For representation purposes, all BLASTp E-values of 0 were set equal to 1e-200. The heatmap of log_10_(E-values) was generated using the *pheatmap* package without clustering in R. This approach was also used to assess the conservation of hDPP4 in the gut microbiota.

To compare human DPP4 to the putative homologs in *B. thetaiotaomicron* BT4193 and BT3254 and other aminopeptidases, protein sequences were downloaded from UniProt and aligned using the web-based Clustal Omega tool (https://www.ebi.ac.uk/Tools/msa/clustalo/). Phylogenetic trees from multiple sequence alignments were visualized using FigTree v1.4.4. Predicted structures of BT4193 and BT3254 were generated using AlphaFold Colab (https://colab.research.google.com/github/deepmind/alphafold/blob/main/notebooks/AlphaFold.ipynb) and aligned to the crystal structure of hDPP4 (PDB 1J2E) using the *MatchMaker* function in UCSF Chimera v1.13.1.

### Mass spectrometry-based activity-based protein profiling sample preparation and analysis

*B. thetaiotaomicron* VPI-5482 cultures were grown in 30 mL of mBHI in triplicate for 72 h prior to being aliquoted into three separate tubes and pelleted. For each biological replicate, each pellet was resuspended in 200 µL of either 25 µM FP-TMR in PBS or 0.05% DMSO in PBS and incubated anaerobically for 30 min at 37 °C. Then, 2 µL of either 500 µM FP-biotin (final concentration, 5 µM) or DMSO was added to each sample and incubated anaerobically for 30 min at 37 °C. Bacterial pellets were stored at -20 °C prior to sample preparation.

Treated *B. thetaiotaomicron* cell pellets were thawed, diluted in PBS (200 μL) and lysed by probe sonication. The proteomes were denatured and precipitated using 4:1 MeOH/CHCl_3_, resuspended in 0.5 mL of 6 M urea in PBS, reduced using tris(2-carboxyethyl)phosphine (TCEP, 10 mM) for 30 min at 37 °C, and then alkylated using iodoacetamide (40 mM) for 30 min at room temperature in the dark. The biotinylated proteins were enriched with PBS-washed Pierce™ Streptavidin Agarose beads (100 µL per sample) by rotating at room temperature for 1.5 h in PBS with 0.2% SDS (6 mL). The beads were then washed sequentially with 5 mL 0.2% SDS in PBS (3x), 5 mL PBS (3x) and 5 mL H_2_O (3x). On-bead digestion was performed using sequencing-grade trypsin (2 μg; Promega) in 2 M urea in 100 mM triethylammonium bicarbonate (TEAB) buffer with 2 mM CaCl_2_ for 12–14 h at 37 °C (200 μL).

Tryptic digests were desalted using C18 solid phase extraction (SPE). C18 SPE plates (Thermo Scientific) were conditioned by the addition of acetonitrile (ACN) to each well and plates were centrifuged at 750 x *g* for 2 min. This process was repeated with Buffer A (200 μL, 0.1% formic acid in water). Samples were diluted with 560 μL Buffer A and triturated with a pipette to mix. A portion (560 μL) of each sample was added to separate wells of the SPE plate, the sample was allowed to slowly load over 5 min, then the plate was centrifuged for 5 min at 200 x *g*. The remaining sample volume was loaded into the same well of the SPE plate and the loading step was repeated. Samples were washed twice with Buffer A (200 μL, then 100 μL) followed by centrifugation of the SPE plate at 200 x *g* for 5 min after each wash. Samples were eluted into a clean 96-well plate by the addition of Buffer B (60 μL of 70% ACN and 0.1% formic acid in water) and centrifugation at 200 x *g* for 5 min. Samples were dried by centrifugal vacuum concentrator (50 °C, overnight) and resolubilized in TEAB (100 μL,100 mM, pH 8.5) containing 30% ACN. Peptides were labelled with the addition of TMT tags (3 μL/channel, 19.5 μg/μL) to each sample and incubated at room temperature for 30 min. Hydroxylamine (8 μL, 5% in water) was subsequently added to quench the labelling reaction (15 min) and the TMT-labelled samples were mixed and desalted on C18 using the same protocol as above.

Nanoflow LC-MS/MS measurements were performed on an Ultimate 3000 (Thermo Scientific) interfaced with an Orbitrap Fusion Lumos Tribrid mass spectrometer (Thermo Scientific) via an EASY-Spray source (Thermo Scientific). Peptides were separated on an EASY-Spray PepMap RSLC C18 column (2 µm particle size, 75 µm × 50 cm; Thermo Scientific, ES801) heated to 55 °C using a flow rate of 400 nL/min. The compositions of LC solvents were A: water and 0.1 % formic acid, and B: 95% acetonitrile, 5% water and 0.1% formic acid. Peptides were eluted over 4 hours using the linear gradient, 2.5-215 min 3-35% B, 215-230 min 25-40% B, 230-231 min 45-70% B, 231-233 min 70-90% B, 233-234 min 5-70% B, 234-236 min 70-90% B, 236-240 min 3% B.

MS data were acquired using an MS3 data-dependent acquisition method. MS1 profile scans were acquired in the Orbitrap (resolution: 120,000, scan range: 375–1500 m/z, AGC target: 4.0e5, maximum injection time: 50 ms). Monoisotopic peak determination was set to “peptide”. Only charge states 2-5 were included. Dynamic exclusion was enabled (repeat count, *n*: 1, exclusion duration: 60 s, mass tolerance: ppm, low: 10 and high: 10, excluding isotopes). An intensity threshold of 5e3 was set. Data-dependent MS2 spectra were acquired in centroid mode across a mass range of 400-1200 m/z. Precursor ions were isolated using the quadrupole (isolation window: 0.7 m/z), fragmented using CID (collision energy: 35%, activation time: 10 ms, activation Q: 0.25), and detected in the ion trap (scan range mode: auto m/z normal, scan rate: rapid, AGC target: 1.0e4, maximum injection time: 50 ms). Data-dependent MS3 spectra were acquired in the Orbitrap (resolution: 50000, scan range: 100-500 m/z, AGC target: 1.0e5, maximum injection time: 105 ms) following HCD activation (collision energy: 65%) using Synchronous Precursor Selection from up to 10 precursors.

Peptide and protein identifications were performed using MaxQuant v1.6.0.16^81^ using the search engine Andromeda^82^. Group-specific parameters were set to “Reporter ion MS3” with 10plex TMT isobaric labels for N-terminal and lysine residue modification selected. Reporter mass tolerance was set to 0.003 Da. Following parameters were used: Carbamidomethylation of cysteines as fixed modifications, oxidation of methionine and acetylation of N-terminus as dynamic modifications, trypsin/P as the proteolytic enzyme. Default settings were used for all other parameters. Searches were performed against the UniProt database for *B. thetaiotaomicron* VPI-5482 (Proteome ID: UP000001414, downloaded April 2019). Identification was performed with at least 2 unique peptides and quantification only with unique peptides.

Statistical analysis was performed with Perseus v1.6.0.7^83^. Putative contaminants, reverse hits, and proteins identified by side only were removed. LFQ intensities were log_2_-transformed. Missing values were imputed using a normal distribution (width = 0.3, down-shift = 1.8). *P*-values were calculated using a two-sided, two-sample t-test.

### Cloning, expression, and purification of recombinant BT4193 and BT3254

All enzymes were purchased from New England BioLabs. A single colony of *B. thetaiotaomicron* VPI-5482 was boiled in 10 µL of water at 95 °C for 5 min to isolate genomic DNA. The genomic sequences for *BT4193* and *BT3254* were amplified and then cloned into the pET28a plasmid with a C-terminal His_6_ tag using Gibson cloning following manufacturer’s instructions (for primer sequences, see Supplementary Table 7). Successful cloning was confirmed by selection on 50 µg/mL kanamycin followed by Sanger sequencing. The plasmids pET28a-BT4193-His and pET28a-BT3254-His (Supplementary Table 7) were then transformed into chemically competent *E. coli* Rosetta 2(DE3) cells with selection using 50 µg/mL kanamycin and 34 µg/mL chloramphenicol for expression.

For recombinant BT4193, an overnight culture of transformed cells was inoculated into 1 L of LB medium supplemented with 50 µg/mL kanamycin and 34 µg/mL chloramphenicol and grown with shaking until OD_600_ reached 0.6. After the culture was cold shocked on ice for 20 min, 0.1 mM isopropyl β-D-1-thiogalactopyranoside (IPTG) was added to the culture to induce protein expression, and the culture was incubated for 20 h at 19 °C. The bacterial pellet was flash frozen in liquid nitrogen (LN_2_) and stored at -80 °C. The pellet was lysed in 30 mL of lysis buffer (10 mM imidazole, 150 mM NaCl, pH 8) supplemented with 750 mM trehalose by probe sonication on ice (3 min at 30% power in 1 s bursts; 1 min at 60% power in 1 s bursts; 3 min at 30% power in 1 s bursts; 1 min at 60% power in 1 s bursts). Cell debris was pelleted at 38,400g for 20 min at 4 °C, and lysate was transferred to a new tube and centrifuged again at 12,000g for 15 min at 4 °C. The lysate was then clarified processively through a 5-µm and then 1-µm filter before purification. The clarified lysate was injected into an ÄKTApurifier FPLC system (GE Healthcare) to be separated with two in-parallel 1-mL HisTrap FF crude columns (GE Healthcare). The column was washed with lysis buffer supplemented with 10 mM imidazole, 20 mM imidazole, and then 40 mM imidazole. The protein was eluted with lysis buffer supplemented with 300 mM imidazole, and fractions containing protein (as determined by the UV trace and SDS-PAGE) were combined. The protein was further purified using a HiPrep 16/60 Sephacryl S-300 HR column (Cytiva Life Sciences) using lysis buffer, and fractions containing protein (as determined by the UV trace and SDS-PAGE) were combined.

For recombinant BT3254, an overnight culture of transformed cells was inoculated into 1 L of LB medium supplemented with 1% glucose (to suppress leaky expression), 50 µg/mL kanamycin, and 34 µg/mL chloramphenicol and grown with shaking until OD_600_ reached 0.8. After the culture was cold-shocked on ice for 20 min, 0.5 mM IPTG was added to the culture and incubated for 20 h at 19 °C. The bacterial pellet was flash frozen in LN_2_ and stored at -80 °C until purification. The pellet was lysed in lysis buffer (10 mM imidazole, 150 mM NaCl, pH 8) supplemented with 750 mM trehalose by probe sonication on ice (3 min at 30% power in 1 s bursts; 1 min at 60% power in 1 s bursts; 3 min at 30% power in 1 s bursts; 1 min at 60% power in 1 s bursts). Cell debris was pelleted at 15,000g for 30 min at 4 °C, but overexpressed protein remained in the insoluble fraction, as determined by SDS-PAGE. To purify the unfolded protein, the pellet was solubilized in denaturing buffer (100 mM Na_2_PO_4_, 10 mM Tris, 8 M urea) at pH 8 for 1 h at 25 °C with rotating and then centrifuged at 15,000g for 20 min at 25 °C. The lysate was incubated with 1 mL of Ni- NTA agarose (Qiagen) for 1 h at 25 °C with rotating. The resin was washed 3x in denaturing buffer at pH 6.3, and protein was eluted 3x in denaturing buffer at pH 5.9 and then 3x in denaturing buffer at pH 4.5. Elution fractions were combined and pH-adjusted to pH 7.5 prior to dialysis into 20 mM HEPES, 150 mM NaCl, 2mM reduced glutathione, 200 µM oxidized glutathione, pH 8 for 72 h, with the dialysis buffer being replaced once.

### Fluorogenic peptide substrate synthesis

7-Amino-4-carbamoylmethylcoumarin (ACC)-containing fluorogenic peptides were synthesized using solid-phase peptide synthesis as previously reported^84^. Each coupling step was performed at 25 °C with gentle shaking. The completion of each amide coupling was confirmed using the ninhydrin test in which several microliters of dried resin were incubated in 1 mL of ninhydrin solution (5 g ninhydrin in 100 mL ethanol) for 3 min at 120 °C. An incomplete coupling reaction with some remaining free amine groups was visualized by the resin changing to a dark blue color, whereas a complete coupling reaction leaves the resin colorless. Each synthesized peptide was purified by reverse-phase HPLC and lyophilized prior to use.

In brief, Fmoc-protected ACC was coupled to Rink amide AM resin (Chem Impex) using Fmoc-ACC (2.5 eq.) preactivated with hydroxybenzotriazole (HOBt, 2.5 eq.) and then diisopropylcarbodiimide (DIC, 2.5 eq.) in dimethylformamide (DMF) overnight. Resin was washed 3x in DMF and 6x in dichloromethane (DCM) and the reaction was checked with a ninhydrin test. Resin was swelled in DMF for 20 min and then subjected to an additional coupling with Fmoc-ACC (1.25 eq.) preactivated with HOBt (2.5 eq.) and then DIC (2.5 eq.) in DMF overnight. Resin was washed 3x in DMF and 6x in dichloromethane (DCM) and stored at -20 °C. For each peptide, Fmoc-ACC resin was swelled for 20 min in DMF prior to Fmoc deprotection with 20% piperidine in DMF 2x for 20 min. The Fmoc-P1 amino acid (5 eq.) was preactivated with N,N-diisopropylethylamine (DIPEA) or 2,4,6-collidine (5 eq.) and then coupled with HATU (5 eq.) in DMF 2-3x for 24 h until completion, washing with DMF between each coupling. P1 coupling was confirmed with a ninhydrin test and LC-MS after cleaving several microliters of resin in trifluoroacetic acid (TFA) for 20 min. Any remaining free ACC was capped with 1:1 acetic anhydride:DMF plus two drops of pyridine 2x for 1 h. P2, P3, and P4 amino acid couplings were performed similarly with alternating washes in DMF and DCM between each step: 2x Fmoc deprotection with 20% piperidine in DMF for 20 min, 1-2x coupling with Fmoc-Alanine (5 eq.) preactivated with DIPEA or 2,4,6-collidine (5 eq.) followed by coupling with HATU or HBTU (5 eq.) in DMF for 1 h. The synthesized peptides underwent Fmoc deprotection 2x with 20% piperidine in DMF for 20 min, and acetylated peptides were capped with 1:1 acetic anhydride:DMF plus two drops of pyridine for 2 h. Each peptide was cleaved from the resin in deprotection solution (95% TFA, 2.5% triisopropylsilane, 2.5% H_2_O) for 2.5 h and then precipitated in cold ether at -20 °C overnight.

### Enzyme kinetics

For each assay, 20 µL of lysate or recombinant enzyme was combined with 5 µL of 50 µM fluorogenic peptide substrate (final concentration, 10 µM, 0.2% DMSO) in a 384-well plate. Fluorescence (ex/em: 380/460 nm for AMC substrates; ex/em: 355/460 nm for ACC substrates) was measured every minute for 30 min at 37 °C on a Cytation 3 plate reader (BioTek). Initial velocities were quantified by calculating the slope of the linear phase of the reaction as relative fluorescent units (RFUs) per min.

To measure *in vitro* enzyme activity, *B. thetaiotaomicron* wild-type and Δ*BT4193*, Δ*BT3254*, and Δ*BT4193* Δ*BT3254* knockout strains were diluted into mBHIS and grown to late exponential phase in three parallel cultures. Bacteria were pelleted by centrifugation and lysed in 100 mM HEPES, pH 7.5 by probe sonication at 15% power (3x 20 s, with 2 min on ice in between). Cell debris was pelleted by centrifugation, and lysate concentration was normalized to 0.25 µg/µL with 100 mM HEPES, pH 7.5 (final concentration, 5 µg/well). To measure the activity of recombinant hDPP4 (Sino Biological), BT4193, and BT3254, enzyme was diluted to 1.25 nM in 100 mM HEPES, pH 7.5 (final concentration, 1 nM). All assays were performed in technical triplicates across 2-3 independent experiments.

For calculation of IC_50_ values, recombinant enzyme was diluted to 1.25 nM in 20 mM HEPES, pH 7.5 and incubated with inhibitor, diluted 1:100 from DMSO stocks, for 30min at 37 °C prior to monitoring activity. Initial velocities were normalized to DMSO-pretreated enzyme (100%) or buffer only (0%).

### Competitive gel-based activity-based protein profiling

*B. thetaiotaomicron* wild-type and Δ*BT4193*, Δ*BT3254*, and Δ*BT4193* Δ*BT3254* knockout strains were diluted into mBHIS and grown to late exponential phase. For each sample, 4 µL of DMSO or 100x stocks of covalent hDPP4 inhibitor (10 µM, 100 µM, or 1mM stocks of talabostat or saxagliptin in DMSO) were added to 400 µL of bacterial culture and incubated anerobically for 1 h at 37 °C. To each sample, 4 µL of 500 µM FP-alkyne (final concentration, 5 µM) were added and incubated anerobically for 30 min at 37 °C. Bacterial pellets were pelleted by centrifugation and lysed in PBS by probe sonication at 15% power (3 x 20 seconds, with 2 min on ice in between). Cell lysate was clarified by centrifugation, and lysate concentration was normalized. Tetramethylrhodamine (TMR)-azide was conjugated to FP-modified enzymes using Cu(I)- catalyzed azide-alkyne cycloaddition (5 mM BTTAA, 1 mM CuSO_4_, 15 mM sodium ascorbate, and 20 mM azide-TMR for 30 min at 37 °C), and labeled enzymes were separated via SDS-PAGE. In-gel fluorescence was visualized using the Cy3 channel on a Typhoon 9410 Imager (Amersham Biosciences).

### Multiplex substrate profiling by mass spectrometry

In a total volume of 150 µL, 10 nM of BT3254 and 10 nM of BT4193 were incubated with a mixture of 228 14-mer peptides (0.5 µM for each peptide) in 20 mM HEPES, pH 7.5 for 240 min at room temperature. Thirty microliters were removed and combined with 100 µL of 8 M urea. A control assay consisted of each enzyme mixed with 8 M urea prior to addition of the peptide library. Assays were conducted in quadruplicate. Samples were then acidified by addition of 40 µL of 2% TFA, desalted using spin tips consisting of octadecyl C18 (CDS Analytical), evaporated to dryness in a vacuum centrifuge, and placed at -80 °C. Samples were resuspended in 40 µL of 0.1% TFA and 4 µL were used for LC-MS/MS.

MS/MS data analysis was performed using PEAKS v8.5 software (Bioinformatics Solutions Inc.). MS2 data were searched with the 228-member 14-mer library sequence as a database, and a decoy search was conducted with peptide amino acid sequences in reverse order. A precursor tolerance of 20 ppm and 0.01 Da for MS2 fragments was defined. No protease digestion was specified. Data were filtered to 1% peptide- and protein-level false discovery rates with the target-decoy strategy. Peptides were quantified with label free quantification and data normalized by TIC. Outliers from replicates were removed by Dixon’s Q testing. Missing and zero values were imputed with random normally distributed numbers under the limit of detection, as determined by the lowest 5% of peptide intensities. ANOVA testing was performed for peptide data of the 0 (control) and 240 min incubation conditions; those with *p*<0.05 were considered for further analysis. Criteria for identification of cleaved peptide products included those with intensity scores of 8-fold or more above the quenched inactive enzymes (time 0 min), evaluated by log_2_(active/inactivated enzyme ratio) for each peptide product, with *p*<0.05 by two-tailed t-test.

LC-MS/MS data files can be obtained through <u>massive.ucsd.edu</u> under the dataset identifier numbers MSV000089969 and MSV000089970.

### Positional scanning library synthesis and screening

One dipeptide and two tripeptide positional scanning libraries with P1 proline residues were synthesized using solid-phase peptide synthesis and standard Fmoc coupling chemistry as reported previously^85^. Briefly, Fmoc-Pro-ACC resin was generated as described above. Unreacted ACC still containing the free amine was acetylated with 1:1 acetic anhydride:DMF plus two drops of pyridine 2x for 1 h. This resin was then moved forward into parallel synthesis on a Syro II parallel peptide synthesizer (Biotage) at a 5 mg scale. Before each coupling step, the resin was swelled in DMF for 10 min prior to Fmoc deprotection with 20% piperidine in DMF for 2x 5 min. For each coupling step, the appropriate Fmoc-protected amino acid (5 eq.) was preactivated with HOBt (5 eq.) and then added to the resin with DIC (5 eq.) to react for 2x 30 min. Between each deprotection and coupling step, the resin was washed in 5x DMF. Peptides were then cleaved from the resin in deprotection solution (95% TFA, 2.5% triisopropylsilane, 2.5% H_2_O) for 2 h, the TFA was evaporated, and the peptides were precipitated in cold ether at -20 °C overnight.

Each library scans through 19 amino acids: alanine (A), aspartate (D), glutamate (E), phenylalanine (F), glycine (G), histidine (H), isoleucine (I), lysine (K), leucine (L), norleucine (n), asparagine (N), proline (P), glutamine (Q), arginine (R), serine (S), threonine (T), valine (V), tryptophan (W), and tyrosine (Y). For the dipeptide library, each peptide was synthesized individually. The P2 tripeptide library consists of 19 members, each with one of the 19 residues in the P2 position and an isokinetic mixture of all 19 in the P3 position as described previously^85^. The P3 tripeptide library consists of 19 members, each with the isokinetic mixture in the P2 position and one of the 19 residues in the P3 position. The libraries were not HPLC-purified before screening.

Each library was screened at a final concentration of 10 µM for each substrate against recombinant enzyme at 1.25 nM in 100 mM HEPES, pH 7.5, as described above. For each library, to normalize across batches of synthesis, the absorbance at 355 nm for ACC for each member of the library was quantified and then used as a normalization factor for each initial velocity. They were then further normalized to the highest velocity for each replicate.

### Growth analyses

*B. thetaiotaomicron* wild-type and Δ*BT4193*, Δ*BT3254*, and Δ*BT4193* Δ*BT3254* knockout strains were grown in mBHIS until late exponential phase and then washed two times via centrifugation with Varel-Bryant medium supplemented with 0.5% glucose (VBG). Cultures were normalized to an OD_600_ of 1 and diluted 1:100 into VBG in three separate wells, supplemented with vancomycin hydrochloride (*n* = 6) or deoxycholic acid (*n* = 3). Anaerobic growth was monitored by measuring absorbance (OD_600_) every 10 min for 24 h at 37 °C using a Cytation 3 plate reader (BioTek).

### Community assembly experiments and 16S rRNA gene sequencing analysis

All bacterial isolates were struck on BHI-blood agar plates (5% defibrinated horse blood in 1.5% w/v agar). Resulting colonies were inoculated into 1mL or BHI, BHIhk, or mGAM and were grown in a 96-deep well plate at 37 °C for 48 h. Communities were assembled from stationary phase cultures of isolates mixed at equal volume, and 1 µL of the mixture was inoculated into 200 µL of BHI or mGAM growth medium. Experiments were performed in clear, flat-bottomed 96-well plates (Greiner Bio-One). Plates were sealed with breathable AeraSeals (Excel Scientific). Sealed plates were placed in a loosely closed bag to minimize evaporation and incubated at 37 °C without shaking for 48 h. Plates were then re-sealed with foil seals and stored at -80 °C for sequencing preparation.

Amplicon sequencing data were obtained and processed as previously described^57^. Relative abundances were determined to a minimum threshold of 10^-3^, reflecting the typical depth of sequencing.

### Software

Graphs and statistical analyses were generated using RStudio, GraphPad Prism 9, and Matlab (Mathworks, Inc.) Data are plotted as mean ± SEM. For the comparison of wild-type and knockout strains, one-way ANOVA tests were performed, with posthoc Dunnett’s multiple comparisons tests with wild-type as the control where appropriate (*, p < 0.05; **, p < 0.01; ***, p < 0.001; ****, p < 0.0001).

## Acknowledgements

Many thanks to the laboratories of Dr. Justin Sonnenburg and Dr. Michael Howitt for use of their equipment. L.J.K. was supported by the Stanford ChEM-H Chemistry/Biology Interface Predoctoral Training Program, a Stanford Molecular Pharmacology Training Grant, and a Stanford Graduate Fellowship. T.H.N. was supported by a National Science Foundation Graduate Research Fellowship. R.C. was supported by a National Institutes of Health Training grant T32 HG000044 and a Stand Up 2 Cancer grant (to A.S.B.). M.L. was supported by Deutsche Forschungsgemeinschaft (DFG) for funding under the Walter Benjamin Program. F.F. was supported by a National Science Foundation Graduate Research Fellowship (DGE-1656518), a Stanford ChEM-H O’Leary-Thiry Graduate Fellowship, and Stanford’s Enhancing Diversity in Graduate Education Doctoral Fellowship Program. This work was supported by National Institutes of Health grants R01 EB026332 and R01 EB026285 (to M.B.), RM GM135102 and R01 AI147023 (to K.C.H.), R01 AI158612 (to A.J.O.) and R01 AI14862302 and R01 AI14375702 (to A.S.B.), and National Science Foundation Grant EF-2125383 (to K.C.H.). K.C.H is a Chan Zuckerberg Biohub Investigator.

## Author Contributions

L.J.K. and M.B. conceived and designed the study. L.J.K., T.H.N., L.L., N.N., and K.M.L. acquired data. L.J.K., R.C., and F.F. generated reagents. L.J.K., T.H.N., L.L., M.L., D.J.G., P.I., A.J.O., K.C.H. and M.B. analyzed and interpreted data. L.J.K., T.H.N., K.C.H., and M.B. prepared the manuscript. All authors were involved in reviewing and revising the manuscript.

