## Supplemental Methods and Figures for "Chemoproteomic identification of a dipeptidyl peptidase 4 (DPP4) homolog in *Bacteroides thetaiotaomicron* important for envelope integrity and fitness"

**TABLE OF CONTENTS:**

**Supplementary Figure 1:** Schematic of workflow for bioinformatic prediction of serine hydrolases. **Page S3**

**Supplementary Figure 2:** Bioinformatically predicted serine hydrolases in representative gut commensal bacterial strains. **Page S4**

**Supplementary Figure 3:** Bacteria from the phylum Bacteroidetes have a larger fraction of their proteome predicted to be serine hydrolases. **Page S5**

**Supplementary Figure 4:** Serine hydrolases uniquely abundant in the Bacteroidetes phylum comprise two classes of proteases and two types of carbohydrate-active enzymes. **Page S6**

**Supplementary Figure 5:** Phylogenetic tree of BT4193 (BtDPP4) with other known bacterial DPP4 homologs and human prolyl peptidases. **Page S8**

**Supplementary Figure 6:** Sequence alignment of bacterial DPP4 homologs and human prolyl peptidases. **Page S9**

**Supplementary Figure 7:** hDPP4 substrate GP-AMC can only be cleaved by recombinant BT4193. **Page S10**

**Supplementary Figure 8:** hDPP4 inhibitors act on BT4193 but not BT3254. **Page S11**

**Supplementary Figure 9:** Ion intensities of peptides cleaved by BT4193 and BT3254. **Page S12**

**Supplementary Figure 10:** Peptide substrate cleavage in *B. thetaiotaomicron* lysate. **Page S12**

**Supplementary Figure 11:** Recombinant BT4193 and BT3254 prefer P1 Pro residues. **Page S13**

**Supplementary Figure 12:** Recombinant hDPP4 and BT4193 prefer similar P2 residues. **Page S13**

**Supplementary Figure 13:** Fluorogenic peptide substrates AP-ACC and AAP-ACC are not cleaved by *B. thetaiotaomicron* conditioned media. **Page S14**

**Supplementary Figure 14:** Dose-dependent inhibition of *B. thetaiotaomicron* growth with vancomycin treatment. **Page S14**

**Supplementary Figure 15:** Sensitivity to deoxycholic acid in the absence of *BT4193* occurs in stationary phase. **Page S15**

**Supplementary Figure 16:** Total community structure is not affected by the absence of *BT4193* or *BT3254*. **Page S16**

**Synthetic Methods and Characterization** **Page S17**

**References** **Page S21**

Supplementary Tables 1-8 are included in a separate Excel file.

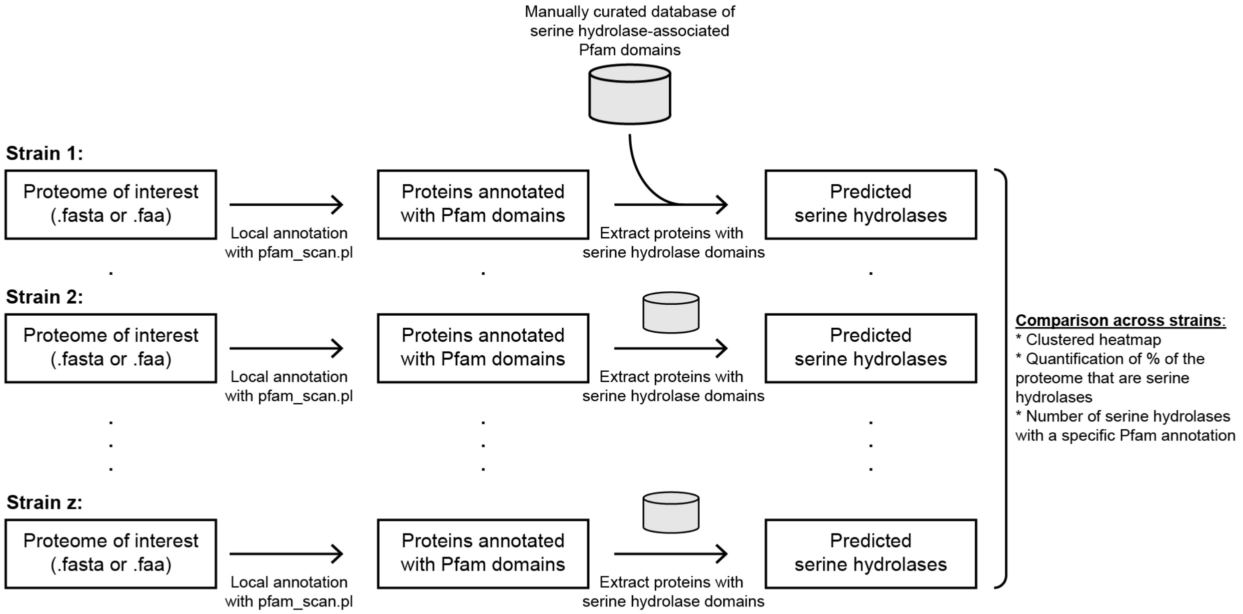


**Supplementary Figure 1: Schematic of workflow for bioinformatic prediction of serine hydrolases.** A list of serine hydrolase-associated Pfam domains was compiled based on ABPP-identified serine hydrolases and known serine hydrolase protein folds in the MEROPS and ESTHER databases. For each strain of interest, the proteome is annotated with Pfam domains with a .fasta file used as input. The annotated proteome is then filtered to select for proteins annotated with a serine hydrolase-associated Pfam domain. These predicted serine hydrolases can then be compared across bacterial strains.

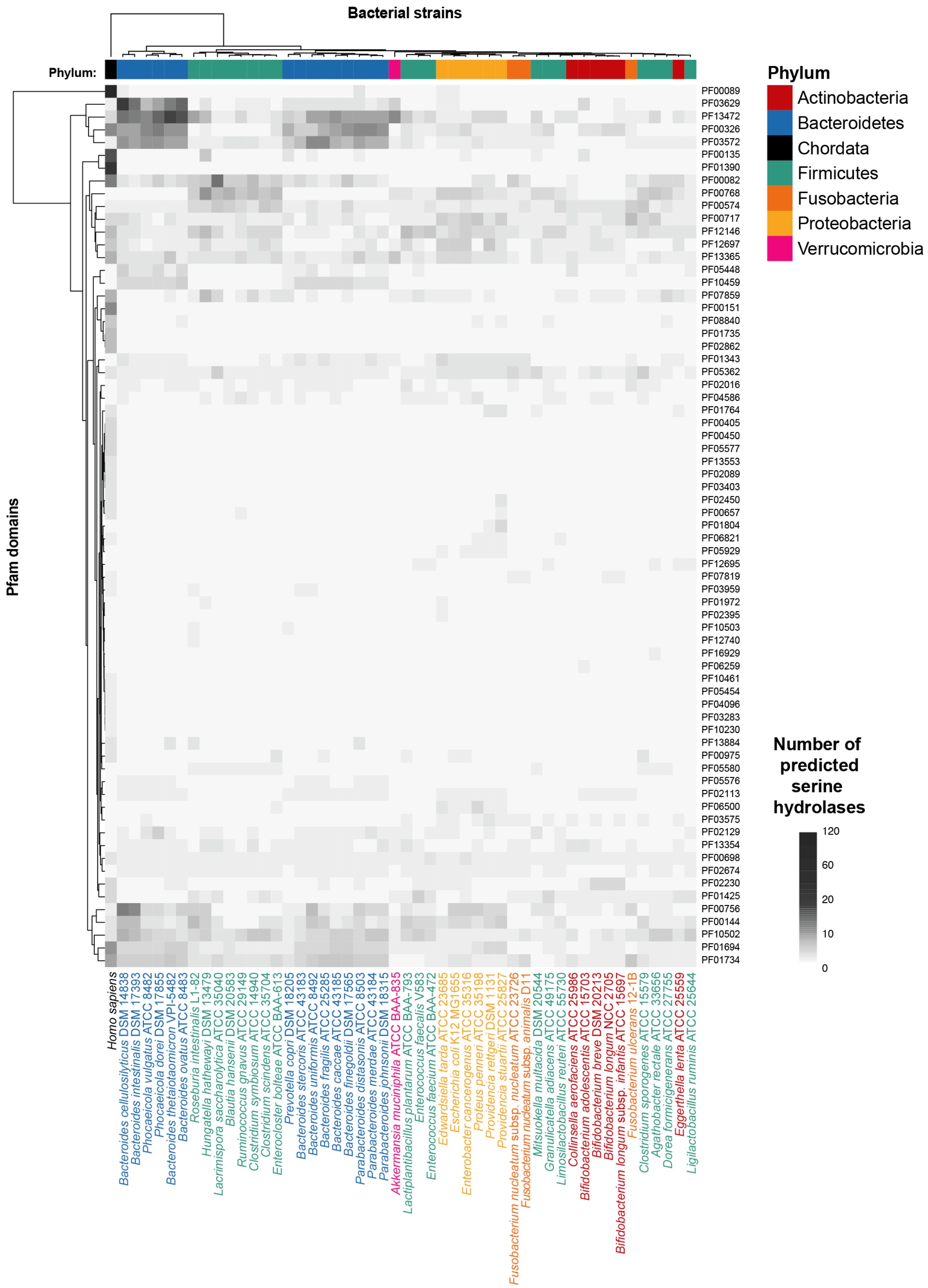


**Supplementary Figure 2: Bioinformatically predicted serine hydrolases in representative gut commensal bacterial strains.** Heatmap of number of proteins predicted to be serine hydrolases from Fig. 1a annotated with strain names and Pfam domains.

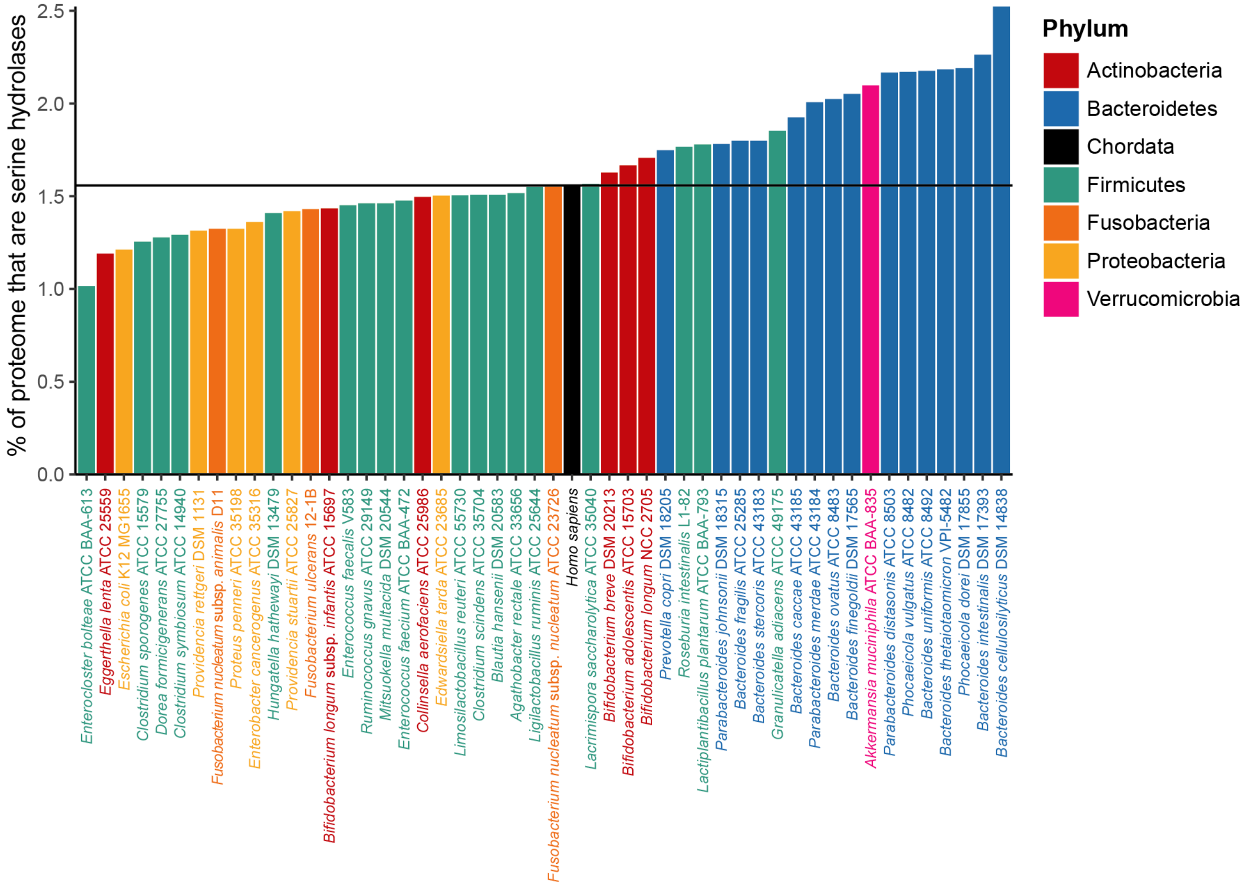


**Supplementary Figure 3: Bacteria from the phylum Bacteroidetes have a larger fraction of their proteome predicted to be serine hydrolases.** Percentage of the total proteome based on the number of proteins that are predicted serine hydrolases, colored by phylum. The horizontal line corresponds to the percentage of predicted human serine hydrolases as a reference.

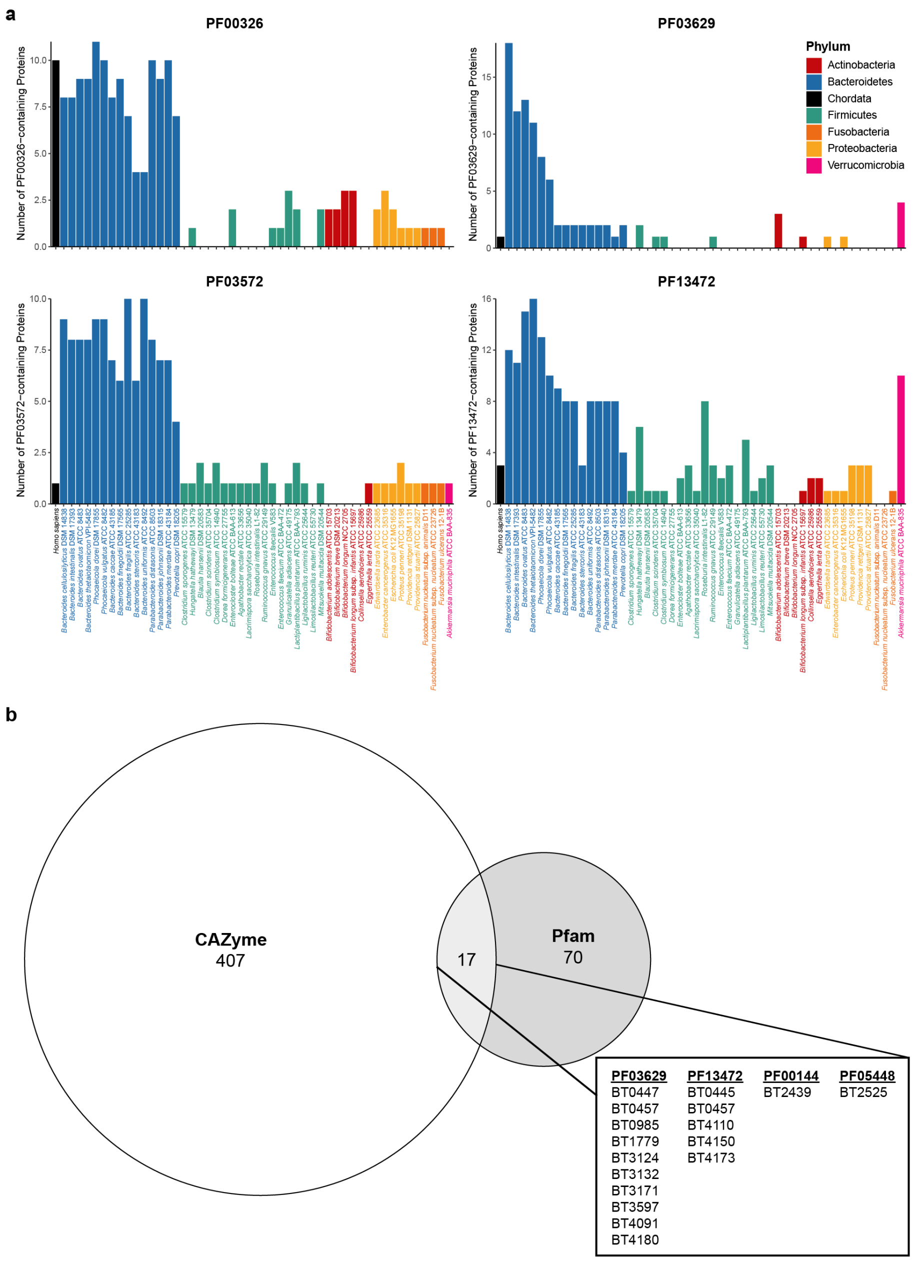


**Supplementary Figure 4: Serine hydrolases uniquely abundant in the Bacteroidetes phylum comprise two classes of proteases and two types of carbohydrate-active enzymes. a,** Number of proteins annotated with each of the four most abundant Pfam domains in the phylum Bacteroidetes for each of the 50 representative gut commensal species, colored by phylum. PF00326 corresponds to the Peptidase S9 domain, PF03572 corresponds to the Peptidase S41 domain, PF03629 corresponds to the Sialic Acid-Specific Acetylesterase domain, and PF13472 corresponds to the GDSL-like Lipase domain. **b,** Comparison of predicted serine hydrolases and proteins annotated as carbohydrate-active enzymes (https://www.cazy.org) in *Bacteroides thetaiotaomicron* VPI-5482. The shared proteins are shown with their serine hydrolase-associated domains in the inset, the majority of which are annotated with either PF03629 or PF13472 domains.

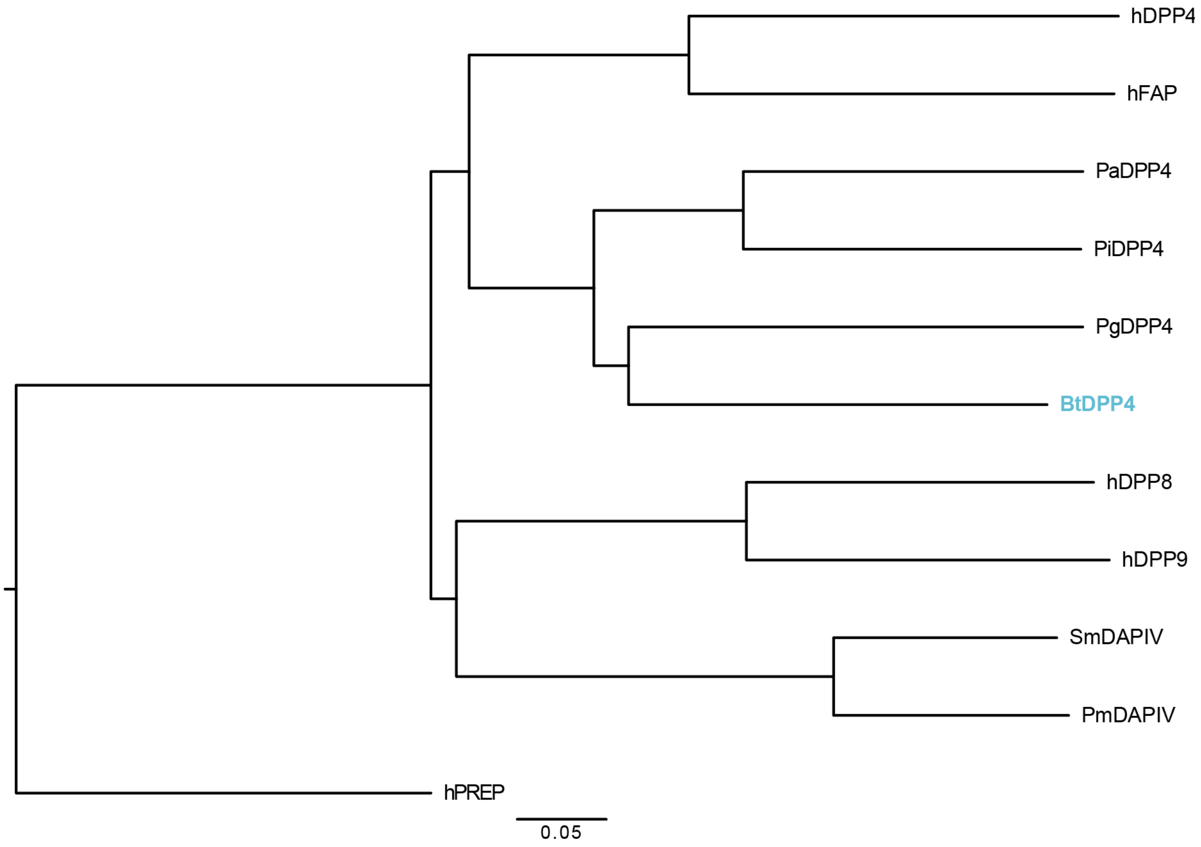


**Supplementary Figure 5: Phylogenetic tree of BT4193 (BtDPP4) with other known bacterial DPP4 homologs and human prolyl peptidases.** Phylogenetic tree of protein sequences of bacterial DPP4 homologs, bacterial DPP9 homologs, and human prolyl peptidases rooted on hPREP as an outgroup. Sequences include bacterial DPP4 homologs from *B. thetaiotaomicron* (BtDPP4; UniProt ID, Q8A028), *Prevotella albensis* (PaDPP4; UniProt ID Q93JY4), *Prevotella intermedia* (PiDPP4; UniProt ID, Q6L872), and *Porphyromonas gingivalis* (PgDPP4; UniProt ID, B2RKU3); bacterial DPP9 homologs from *Stenotrophomonas maltophilia* (SmDAPIV; UniProt ID, P95782) and *Pseudoxanthomonas mexicana* (PmDAPIV; UniProt ID, Q6F3I7); and human prolyl peptidases DPP4 (hDPP4; UniProt ID, P27487), fibroblast activation protein alpha (hFAP; UniProt ID, Q12884), dipeptidyl peptidase 8 (hDPP8; UniProt ID, Q6V1X1), dipeptidyl peptidase 9 (hDPP9; UniProt ID, Q86TI2), and prolyl endopeptidase (hPREP; Unirpto ID, P48147). BtDPP4 is highlighted in blue and clusters with hDPP4 rather than hDPP8/9.

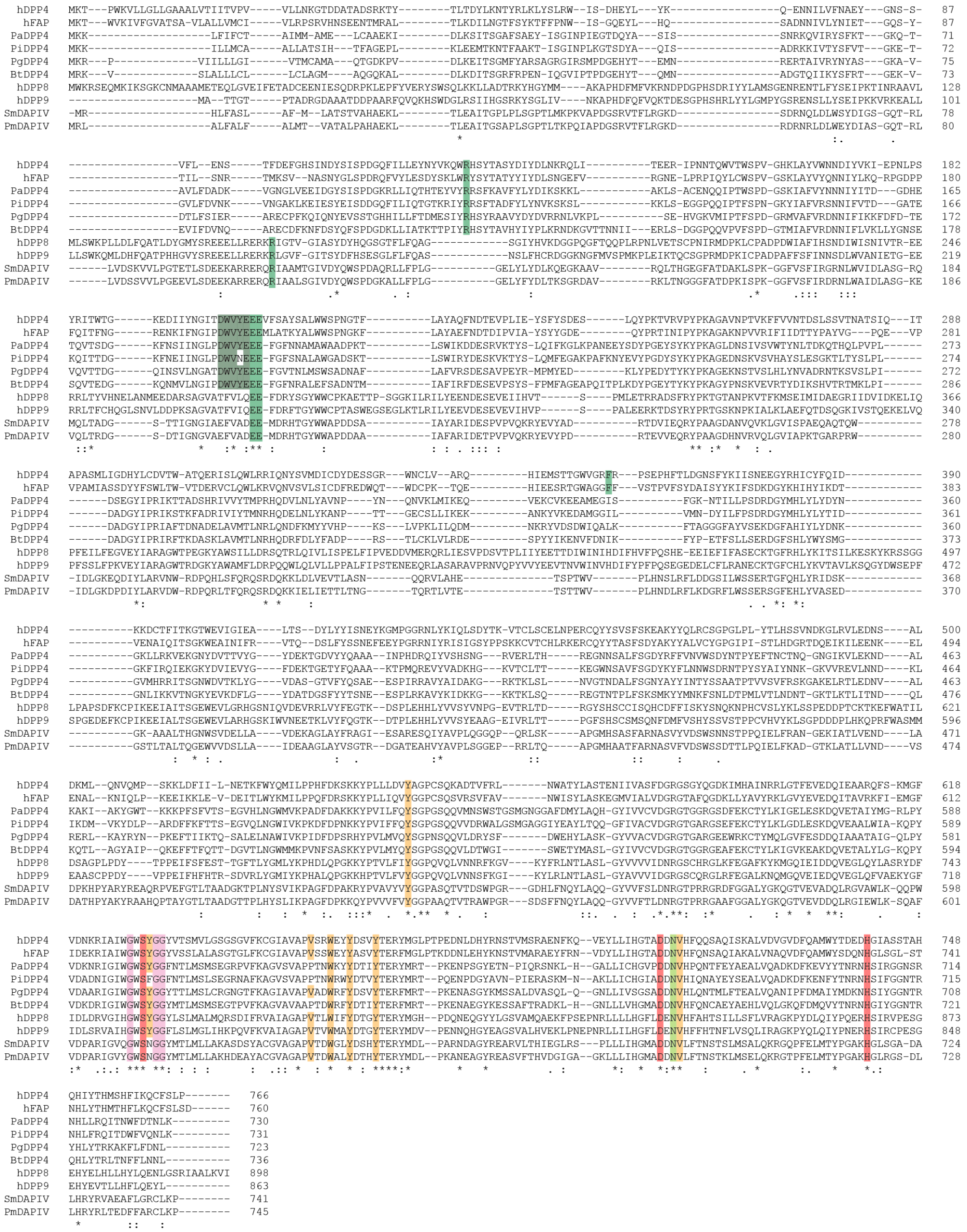

**Supplementary Figure 6: Sequence alignment of bacterial DPP4 homologs and human prolyl peptidases.** Multiple sequence alignment of bacterial DPP4 homologs, bacterial DPP9 homologs, and human prolyl peptidases. Sequences include bacterial DPP4 homologs from *B. thetaiotaomicron* (BtDPP4; UniProt ID, Q8A028), *Prevotella albensis* (PaDPP4; UniProt ID Q93JY4), *Prevotella intermedia* (PiDPP4; UniProt ID, Q6L872), and *Porphyromonas gingivalis* (PgDPP4; UniProt ID, B2RKU3), bacterial DPP9 homologs from *Stenotrophomonas maltophilia* (SmDAPIV; UniProt ID, P95782) and *Pseudoxanthomonas mexicana* (PmDAPIV; UniProt ID, Q6F3I7), and human prolyl peptidases DPP4 (hDPP4; UniProt ID, P27487), fibroblast activation protein alpha (hFAP; UniProt ID, Q12884), dipeptidyl peptidase 8 (hDPP8; UniProt ID, Q6V1X1), dipeptidyl peptidase 9 (hDPP9; UniProt ID, Q86TI2), and prolyl endopeptidase (hPREP; Unirpto ID, P48147). The conserved residues are colored as follows: catalytic triad, red; GxSxG domain, pink; S1 binding pocket, gold; free amine-stabilizing residues, green.

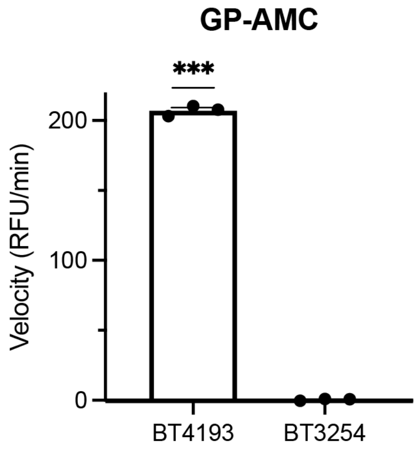


**Supplementary Figure 7: hDPP4 substrate GP-AMC can only be cleaved by recombinant BT4193.** Quantification of cleavage velocity of GP-AMC substrate (ex/em: 380/460 nm) by recombinantly expressed and purified BT4193 and BT3254. Data represent the mean ± SEM of one representative biological replicate with three technical replicates. Statistical significance was determined using a one-sample t-test compared to 0 (***, $p < 0.001$).

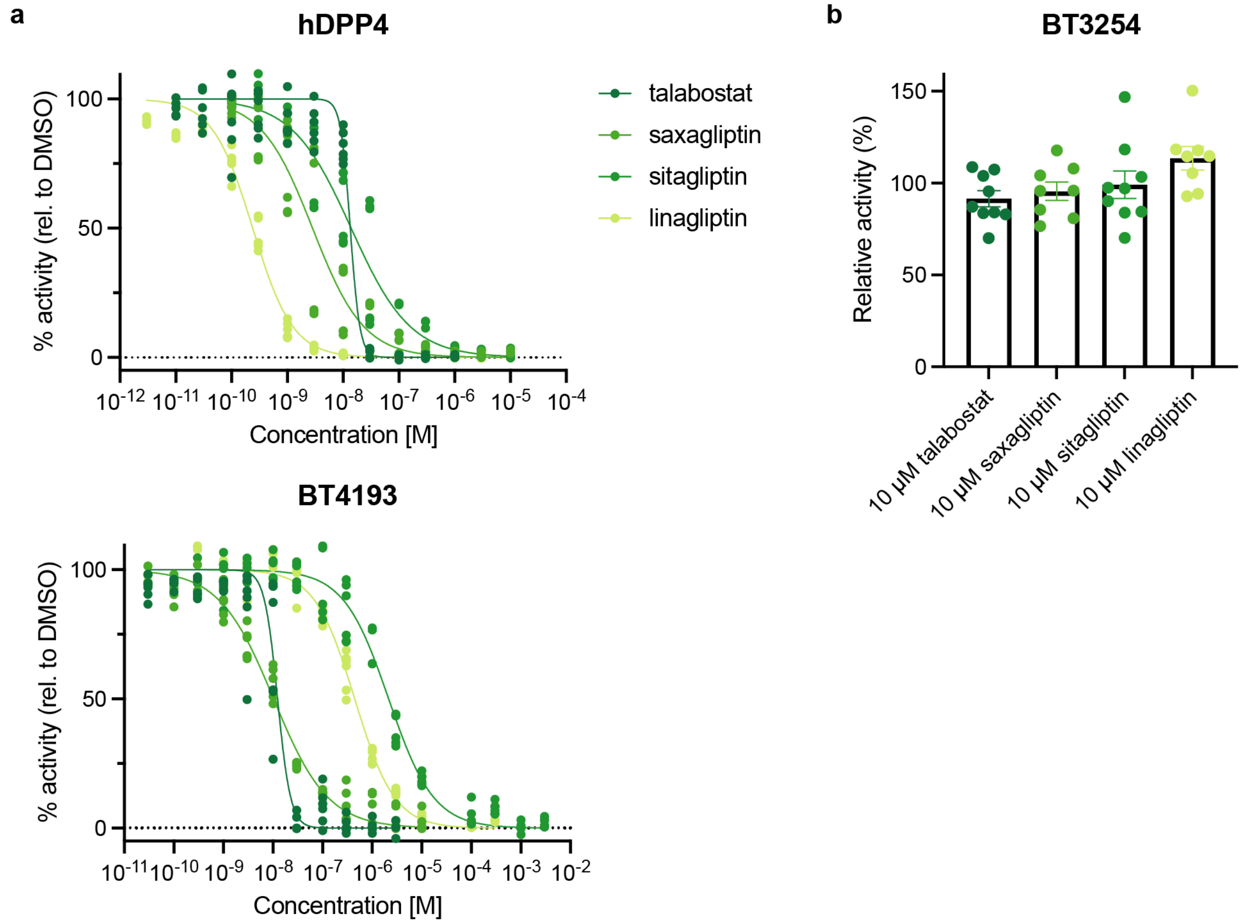


**Supplementary Figure 8: hDPP4 inhibitors act on BT4193 but not BT3254. a,** Dose-dependent inhibition of recombinant hDPP4 and BT4193 by hDPP4 inhibitors after 30 min of pretreatment used to generate apparent $IC_{50}$ values. Velocities were normalized to DMSO pretreatment (100%) and no enzyme (0%) controls. **b,** Quantification of inhibition of recombinant BT3254 by hDPP4 inhibitors after 30 min of pretreatment. Velocities were normalized to DMSO pretreatment.

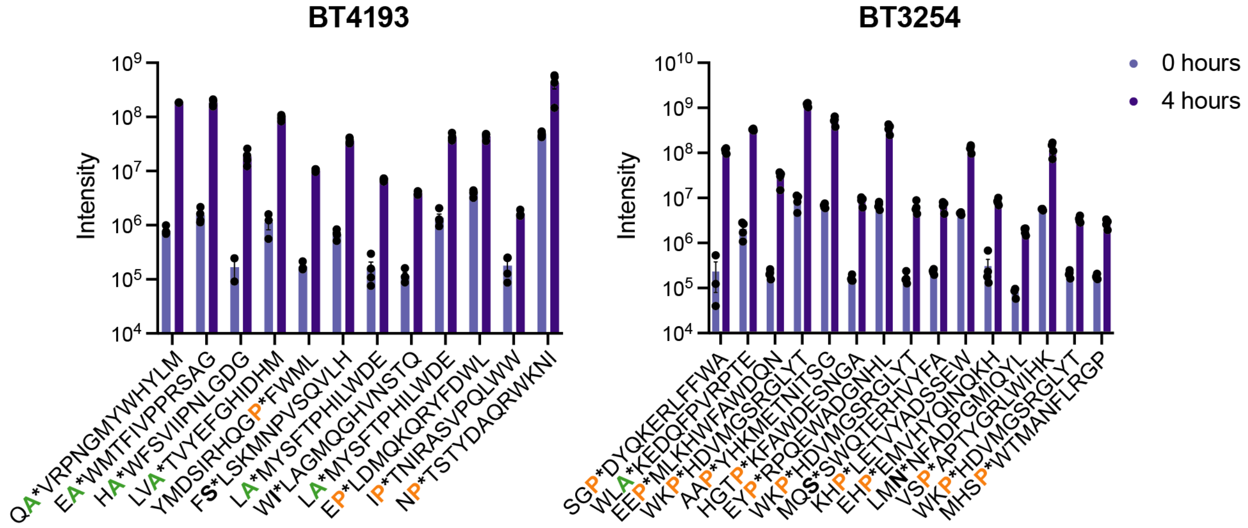


**Supplementary Figure 9: Ion intensities of peptides cleaved by BT4193 and BT3254.** Quantification of peak intensity of each cleaved peptide that was significantly upregulated before and after incubation with recombinant enzyme from multiplex substrate profiling. For each peptide, the cleavage site is denoted with an asterisk. P1 Ala residues are colored green and P1 Pro residues are colored orange.

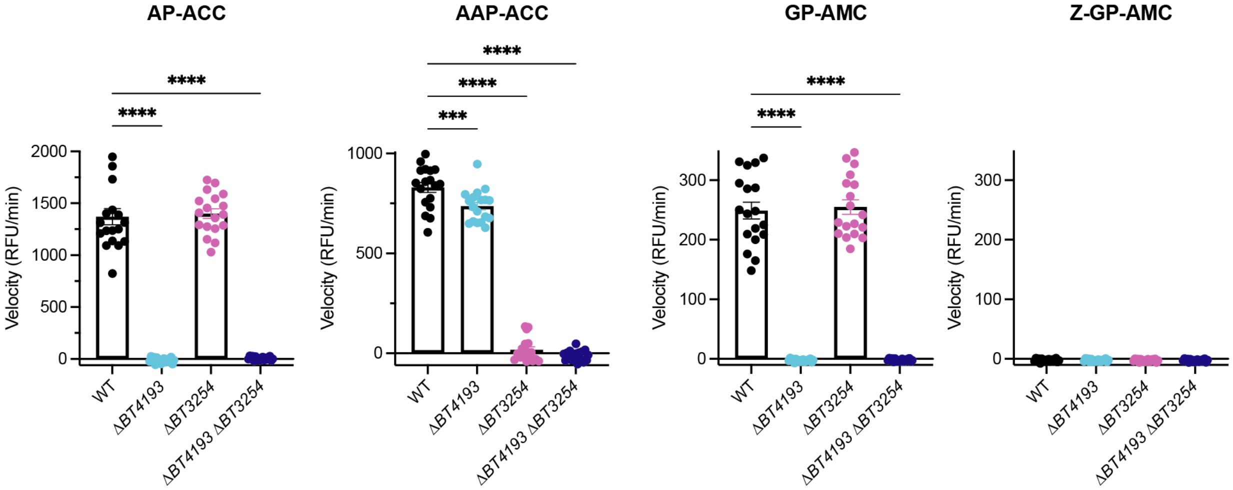


**Supplementary Figure 10: Peptide substrate cleavage in *B. thetaiotaomicron* lysate.** Quantification of initial velocities of fluorogenic peptide substrate cleavage (ex/em: 355/460 nm for ACC substrates; ex/em: 380/460 nm for AMC substrates) in lysate generated from wild-type (WT) and knockout *B. thetaiotaomicron* strains. Statistical significance was determined using a one-way ANOVA test with posthoc Dunnett's multiple comparisons tests compared to wild-type (***, p < 0.001; ****, p < 0.0001).

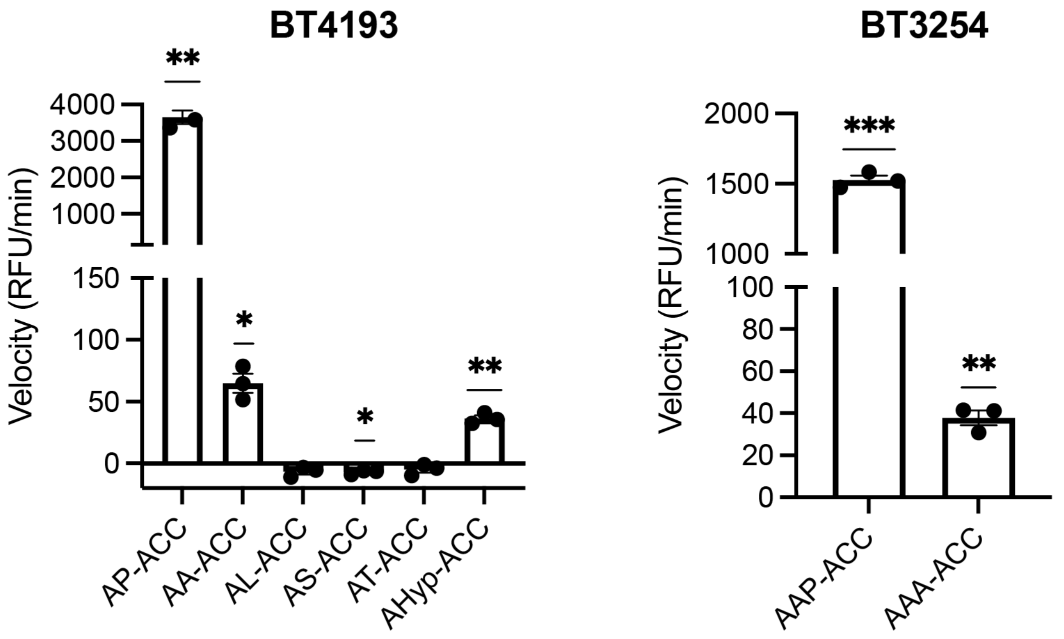


**Supplementary Figure 11: Recombinant BT4193 and BT3254 prefer P1 Pro residues.** Quantification of cleavage velocity of di- and tripeptide fluorogenic peptides with ACC-containing substrates (ex/em: 355/460 nm) by recombinantly expressed and purified BT4193 and BT3254. Data represent the mean ± SEM of one representative biological replicate with three technical replicates. Statistical significance was determined using a one-sample t-test compared to 0 (*, $p < 0.05$; **, $p < 0.01$; ***, $p < 0.001$).

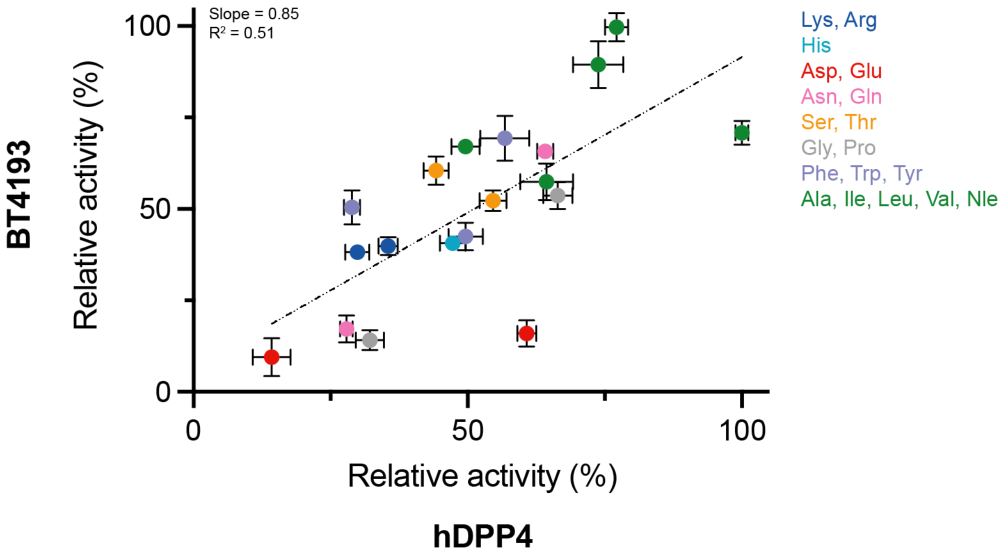


**Supplementary Figure 12: Recombinant hDPP4 and BT4193 prefer similar P2 residues.** Relative cleavage velocities of hDPP4 and BT4193 of each member of the P2Pro-ACC positional scanning library normalized to the best cleaved substrate. Each point is colored by chemical properties. Best-fit line shows linear regression.

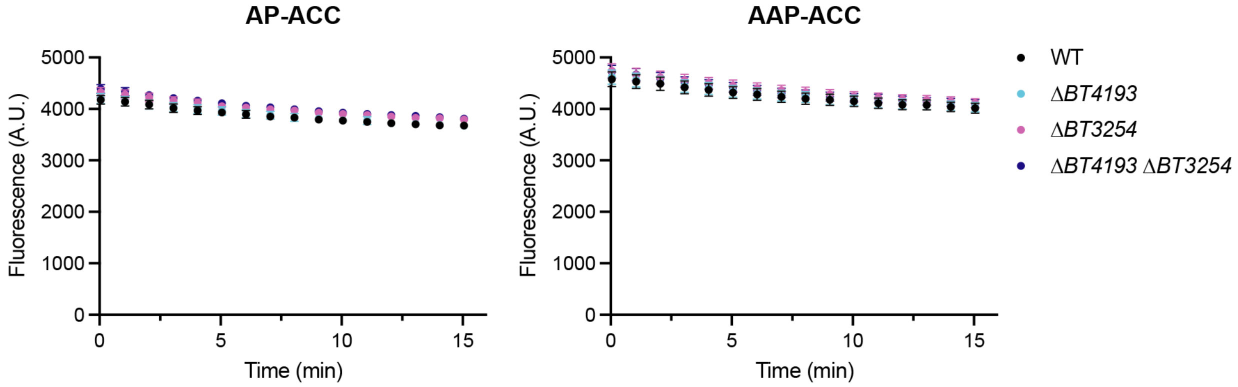


**Supplementary Figure 13: Fluorogenic peptide substrates AP-ACC and AAP-ACC are not cleaved by *B. thetaiotaomicron* conditioned media.** Quantification of fluorescence (ex/em: 355/460 nm) in 20 µL of conditioned mBHIS medium generated from wild-type (WT) and knockout *B. thetaiotaomicron* strains.

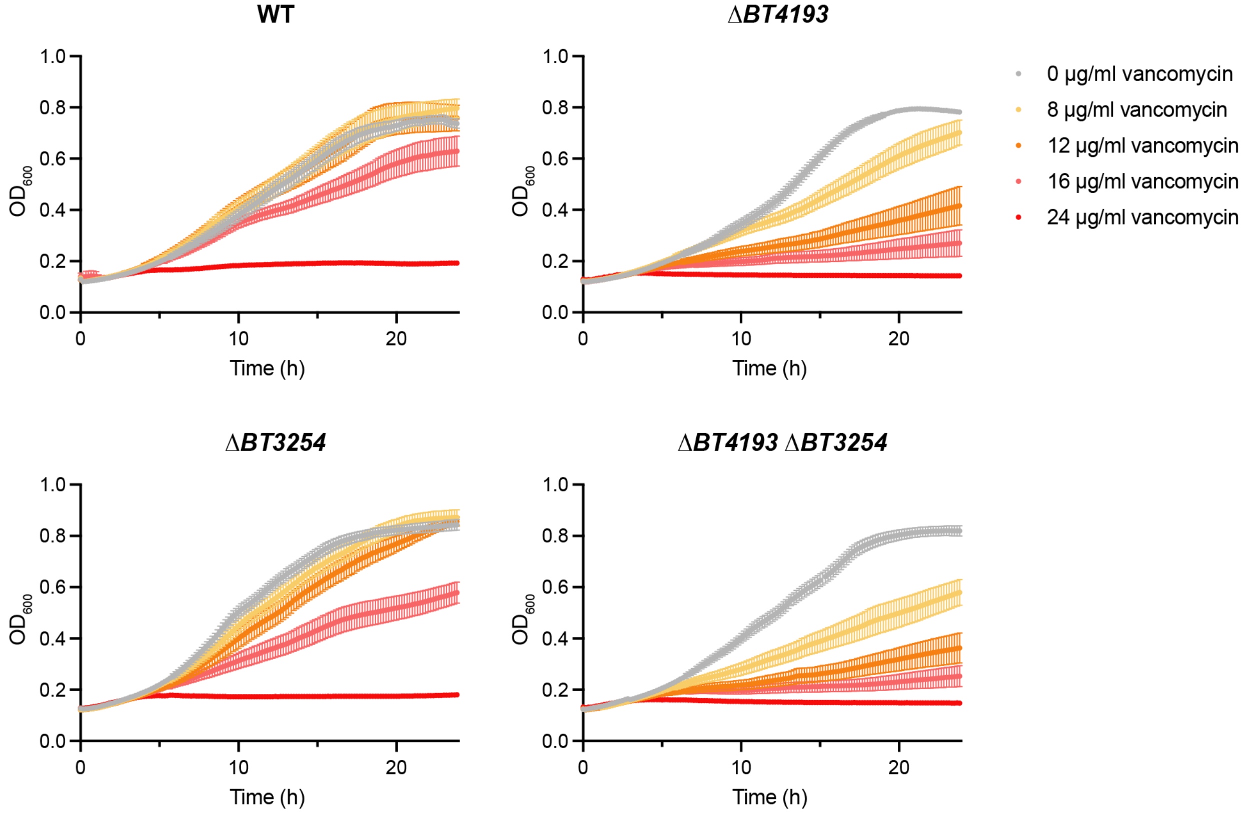


**Supplementary Figure 14: Dose-dependent inhibition of *B. thetaiotaomicron* growth with vancomycin treatment.** Growth of wild-type (WT) and knockout strains of *B. thetaiotaomicron* with increasing concentrations of vancomycin, as measured by $OD_{600}$.

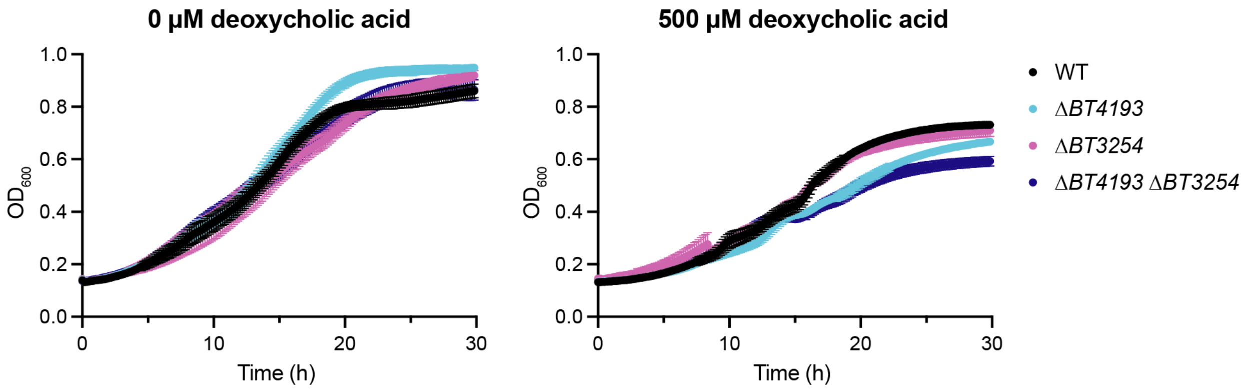


**Supplementary Figure 15: Sensitivity to deoxycholic acid in mutants lacking *BT4193* occurs in stationary phase.** Growth of *B. thetaiotaomicron* strains during deoxycholic acid treatment, as measured by $OD_{600}$.

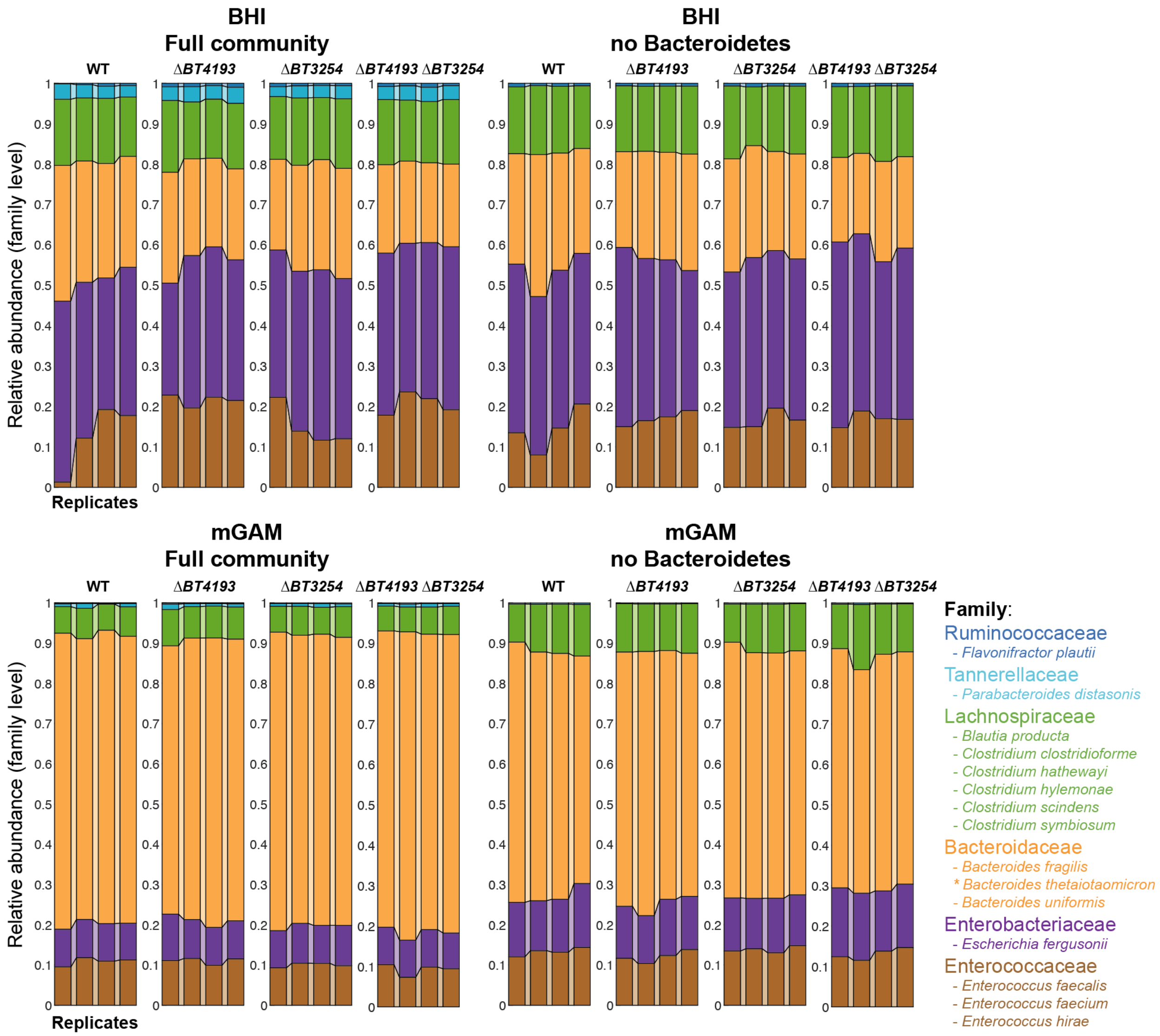


**Supplementary Figure 16: Overall community structure was not affected by the deletion of *BT4193* and/or *BT3254*.** Quantification of relative abundance of bacteria at the family level after 48 h of co-culturing in BHI or mGAM.

**Synthetic Methods and Characterization**

*Synthetic scheme*

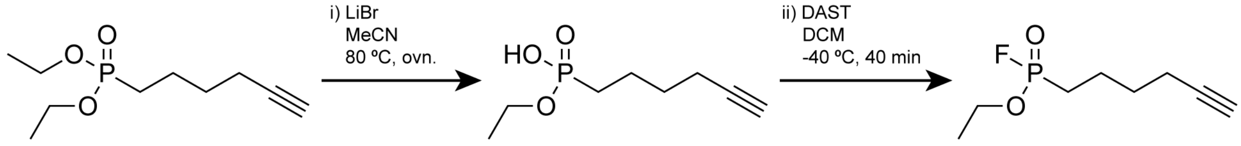


**1** **2** **FP-alkyne**

*Synthetic method*

Diethyl hex-5-ynylphosphonate (**1**) was synthesized as previously reported[1].

i) To a 20-dram vial, LiBr (79.6 mg, 0.92 mmol, 2.0 eq.) and a stir bar were added. This vial was then evacuated and back filled with argon 3x. Subsequently, MeCN (2 mL) and **1** (100 mg, 0.46 mmol, 1.0 eq.) were added. The vial was loosely capped and heated to 80°C in an oil bath. After stirring overnight, the reaction mixture was cooled to r.t. and concentrated in vacuum. The residue was dissolved in water and washed with DCM, the aqueous phase was acidified with 1 M HCl and extracted with DCM. The combined organic phases were dried over $Na_2SO_4$ and concentrated in vacuum without further purification to give the crude intermediate ethyl hydrogen hex-5-yn-1-ylphosphonate (**2**).

ii) To a 20-dram vial was added intermediate **2** (75 mg, 0.39 mmol, 1.0 eq.) and stir bar. The flask was evacuated and purged with argon 3x. Then intermediate **2** was dissolved in 2 mL of dry DCM. The solution was cooled in a cooling bath at -40°C for 15 min. Diethylaminosulfur trifluoride (DAST) (2 eq., 130 mg, 0.79 mmol) was added dropwise and allowed to stir at -40 °C for 40 min. 1 mL of water was added dropwise to quench the reaction and allowed to stir at -40 °C for 5 min. The solution was then allowed to cool to r.t. The solution was extracted 3x with DCM and dried with sodium sulfate, filtered, and evaporated to yield the product **FP-alkyne** as a white amorphous solid (71 mg, 0.37 mmol, 81% yield over two steps).

$^{1}H$ NMR (400 MHz, Chloroform-*d*) δ 4.31 – 4.19 (m, 2H), 2.23 (td, *J* = 6.9, 2.7 Hz, 2H), 1.98 – 1.71 (m, 4H), 1.64 (p, *J* = 6.9 Hz, 3H), 1.37 (t, *J* = 7.1 Hz, 3H). $^{19}F$ NMR (376 MHz, Chloroform-*d*) δ -62.93 (t, *J* = 4.1 Hz), -65.78 (d, *J* = 4.5 Hz). $^{31}P$ NMR (162 MHz, Chloroform-*d*) δ 34.35, 27.74.

*NMR characterization of FP-alkyne*

**$^1$H NMR**

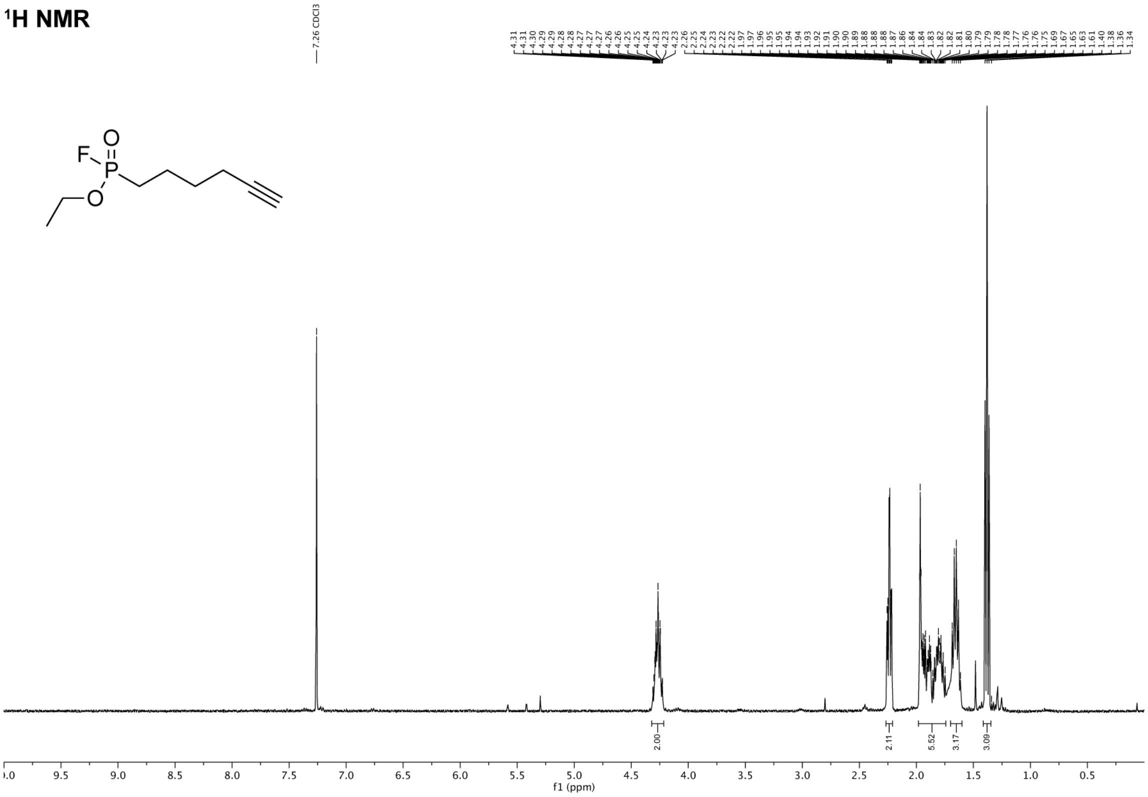

### $^{19}$F NMR

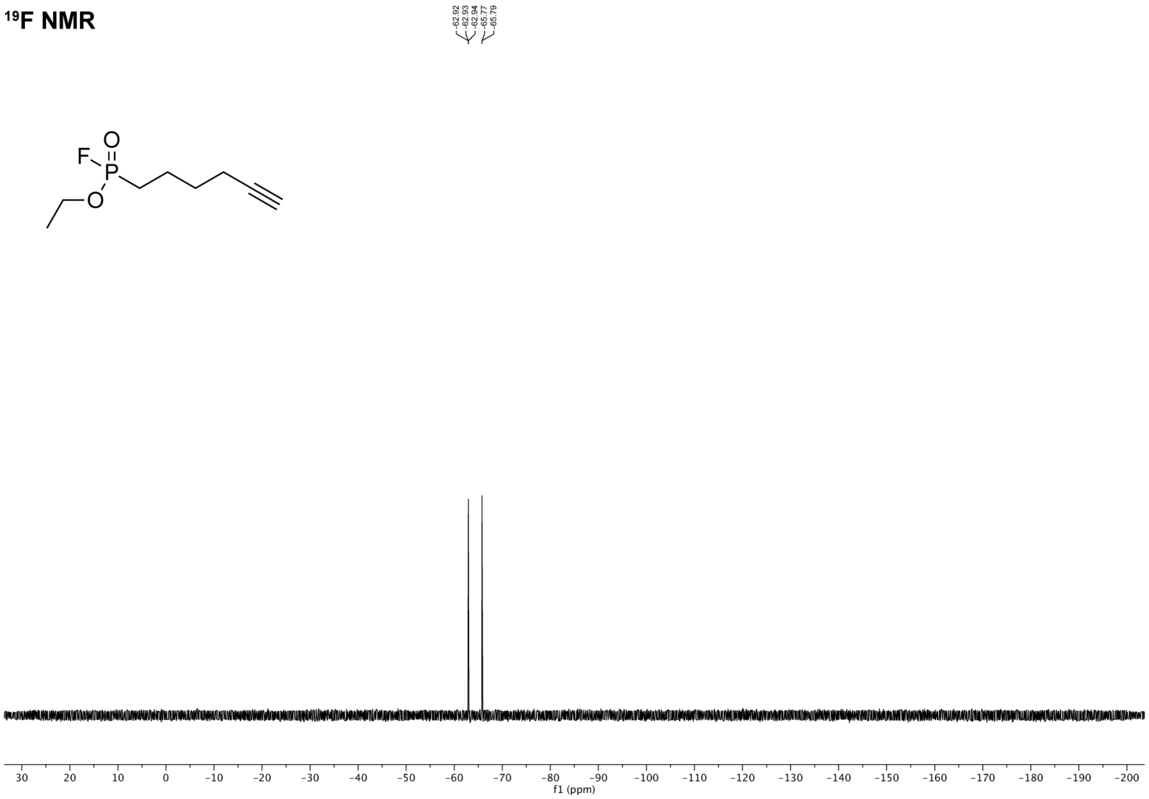


### $^{31}$P NMR

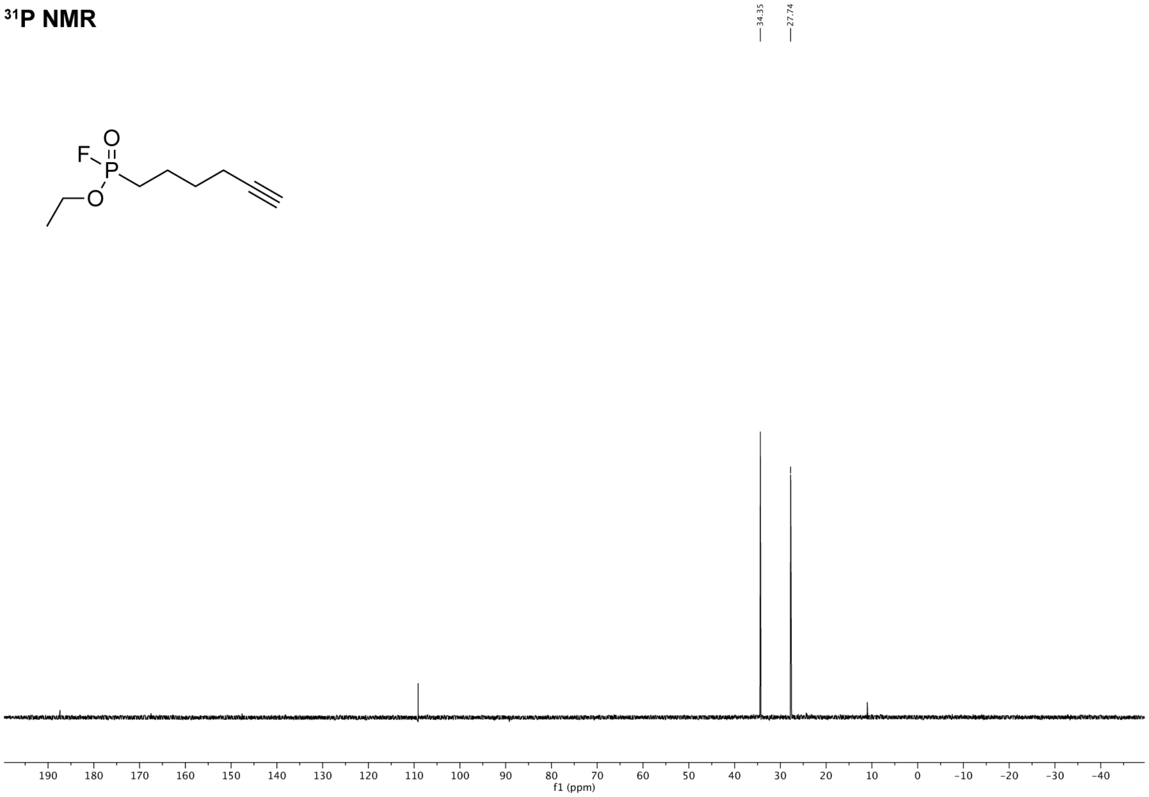
