## Supplemental Table S1 for "Chemoproteomic identification of a dipeptidyl peptidase 4 (DPP4) homolog in *Bacteroides thetaiotaomicron* important for envelope integrity and fitness"

Table S1: Serine hydrolase Pfam domains

| Pfam | Clan | Clan name | Pfam name | Likelihood of being a serine hydrolase |
|---|---|---|---|---|
| PF00135 | CL0028 | AB_hydrolase | Carboxylesterase family | Definite |
| PF00151 | CL0028 | AB_hydrolase | Lipase | Definite |
| PF00326 | CL0028 | AB_hydrolase | Prolyl oligopeptidase family | Definite |
| PF00450 | CL0028 | AB_hydrolase | Serine carboxypeptidase | Definite |
| PF00756 | CL0028 | AB_hydrolase | Putative esterase | Definite |
| PF00975 | CL0028 | AB_hydrolase | Thioesterase domain | Definite |
| PF01083 | CL0028 | AB_hydrolase | Cutinase | Definite |
| PF01674 | CL0028 | AB_hydrolase | Lipase (class 2) | Definite |
| PF01764 | CL0028 | AB_hydrolase | Lipase (class 3) | Definite |
| PF02089 | CL0028 | AB_hydrolase | Palmitoyl protein thioesterase | Definite |
| PF02129 | CL0028 | AB_hydrolase | X-Pro dipeptidyl-peptidase (S15 family) | Definite |
| PF02230 | CL0028 | AB_hydrolase | Phospholipase/Carboxylesterase | Definite |
| PF02273 | CL0028 | AB_hydrolase | Acyl transferase | Definite |
| PF02450 | CL0028 | AB_hydrolase | Lecithin:cholesterol acyltransferase | Definite |
| PF03283 | CL0028 | AB_hydrolase | Pectinacetylesterase | Definite |
| PF03403 | CL0028 | AB_hydrolase | Platelet-activating factor acetylhydrolase, isoform II | Definite |
| PF03583 | CL0028 | AB_hydrolase | Secretory lipase | Definite |
| PF03959 | CL0028 | AB_hydrolase | Serine hydrolase (FSH1) | Definite |
| PF05448 | CL0028 | AB_hydrolase | Acetyl xylan esterase (AXE1) | Definite |
| PF05576 | CL0028 | AB_hydrolase | PS-10 peptidase S37 | Definite |
| PF05577 | CL0028 | AB_hydrolase | Serine carboxypeptidase S28 | Definite |
| PF06259 | CL0028 | AB_hydrolase | Alpha/beta hydrolase | Definite |
| PF06500 | CL0028 | AB_hydrolase | Alpha/beta hydrolase of unknown function (DUF1100) | Definite |
| PF06821 | CL0028 | AB_hydrolase | Serine hydrolase | Definite |
| PF06850 | CL0028 | AB_hydrolase | PHB de-polymerase C-terminus | Definite |
| PF07224 | CL0028 | AB_hydrolase | Chlorophyllase | Definite |
| PF07519 | CL0028 | AB_hydrolase | Tannase and feruloyl esterase | Definite |
| PF07819 | CL0028 | AB_hydrolase | PGAP1-like protein | Definite |
| PF07859 | CL0028 | AB_hydrolase | alpha/beta hydrolase fold | Definite |
| PF08386 | CL0028 | AB_hydrolase | TAP-like protein | Definite |
| PF08840 | CL0028 | AB_hydrolase | BAAT / Acyl-CoA thioester hydrolase C terminal | Definite |
| PF10081 | CL0028 | AB_hydrolase | Alpha/beta-hydrolase family | Definite |
| PF10230 | CL0028 | AB_hydrolase | Lipid-droplet associated hydrolase | Definite |
| PF10340 | CL0028 | AB_hydrolase | Steryl acetyl hydrolase | Definite |
| PF10503 | CL0028 | AB_hydrolase | Esterase PHB depolymerase | Definite |
| PF12146 | CL0028 | AB_hydrolase | Serine aminopeptidase, S33 | Definite |
| PF12695 | CL0028 | AB_hydrolase | Alpha/beta hydrolase family | Definite |
| PF12697 | CL0028 | AB_hydrolase | Alpha/beta hydrolase family | Definite |
| PF12740 | CL0028 | AB_hydrolase | Chlorophyllase enzyme | Definite |
| PF16929 | CL0028 | AB_hydrolase | Accessory Sec system GspB-transporter | Definite |
| PF00144 | CL0013 | Beta-lactamase | Beta-lactamase | Definite |
| PF00768 | CL0013 | Beta-lactamase | D-alanyl-D-alanine carboxypeptidase | Definite |
| PF02113 | CL0013 | Beta-lactamase | D-Ala-D-Ala carboxypeptidase 3 (S13) family | Definite |
| PF13354 | CL0013 | Beta-lactamase | Beta-lactamase enzyme family | Definite |
| PF00574 | CL0127 | ClpP_crotonase | Clp protease | Definite |
| PF01343 | CL0127 | ClpP_crotonase | Peptidase family S49 | Definite |
| PF01972 | CL0127 | ClpP_crotonase | Serine dehydrogenase proteinase | Definite |
| PF03572 | CL0127 | ClpP_crotonase | Peptidase family S41 | Definite |
| PF04096 | CL0661 | Gain | Nucleoporin autopeptidase | Definite |
| PF02016 | CL0014 | Glutaminase_I | LD-carboxypeptidase N-terminal domain | Definite |
| PF03575 | CL0014 | Glutaminase_I | Peptidase family S51 | Definite |
| PF02674 | CL0292 | LysE | Colicin V production protein | Definite |
| PF05497 | CL0037 | Lysozyme | Destabilase | Definite |
| PF01804 | CL0052 | NTN | Penicillin amidase | Definite |
| PF00405 | CL0177 | PBP | Transferrin | Definite |
| PF00603 | CL0236 | PDDEXK | Influenza RNA-dependent RNA polymerase subunit PA | Definite |
| PF00698 | CL0323 | Patatin | Acyl transferase domain | Definite |
| PF01734 | CL0323 | Patatin | Patatin-like phospholipase | Definite |
| PF01735 | CL0323 | Patatin | Lysophospholipase catalytic domain | Definite |
| PF00089 | CL0124 | Peptidase_PA | Trypsin | Definite |
| PF00944 | CL0124 | Peptidase_PA | Alphavirus core protein | Definite |
| PF00949 | CL0124 | Peptidase_PA | Peptidase S7, Flavivirus NS3 serine protease | Definite |
| PF01577 | CL0124 | Peptidase_PA | Potyvirus P1 protease | Definite |
| PF02122 | CL0124 | Peptidase_PA | Peptidase S39 | Definite |
| PF02395 | CL0124 | Peptidase_PA | Immunoglobulin A1 protease | Definite |
| PF02907 | CL0124 | Peptidase_PA | Hepatitis C virus NS3 protease | Definite |
| PF03761 | CL0124 | Peptidase_PA | Nematode trypsin-6-like family | Definite |
| PF05578 | CL0124 | Peptidase_PA | Pestivirus NS3 polyprotein peptidase S31 | Definite |
| PF05579 | CL0124 | Peptidase_PA | Equine arteritis virus serine endopeptidase S32 | Definite |

| | | | | |
|---|---|---|---|---|
| PF05580 | CL0124 | Peptidase_PA | SpoIVB peptidase S55 | Definite |
| PF08192 | CL0124 | Peptidase_PA | Peptidase family S64 | Definite |
| PF10459 | CL0124 | Peptidase_PA | Peptidase S46 | Definite |
| PF13365 | CL0124 | Peptidase_PA | Trypsin-like peptidase domain | Definite |
| PF00717 | CL0299 | Peptidase_SF | Peptidase S24-like | Definite |
| PF10502 | CL0299 | Peptidase_SF | Signal peptidase, peptidase S26 | Definite |
| PF00716 | CL0201 | Peptidase_SH | Assemblin (Peptidase family S21) | Definite |
| PF03420 | CL0201 | Peptidase_SH | Prohead core protein serine protease | Definite |
| PF04586 | CL0201 | Peptidase_SH | Caudovirus prohead serine protease | Definite |
| PF01694 | CL0207 | Rhomboid-like | Rhomboid family | Definite |
| PF05362 | CL0329 | S5 | Lon protease (S16) C-terminal proteolytic domain | Definite |
| PF00657 | CL0264 | SGNH_hydrolas | GDSL-like Lipase/Acylhydrolase | Definite |
| PF03629 | CL0264 | SGNH_hydrolas | Carbohydrate esterase, sialic acid-specific acetylestera | Definite |
| PF13472 | CL0264 | SGNH_hydrolas | GDSL-like Lipase/Acylhydrolase family | Definite |
| PF00082 | | | Subtilase family | Definite |
| PF01390 | | | SEA domain | Definite |
| PF01425 | | | Amidase | Definite |
| PF01768 | | | Birnavirus VP4 protein | Definite |
| PF02862 | | | DDHD domain | Definite |
| PF03574 | | | Peptidase family S48 | Definite |
| PF05454 | | | Dystroglycan (Dystrophin-associated glycoprotein 1) | Definite |
| PF05929 | | | Phage capsid scaffolding protein (GPO) serine peptidas | Definite |
| PF10461 | | | Peptidase S68 | Definite |
| PF10605 | | | 3HB-oligomer hydrolase (3HBOH) | Definite |
| PF13553 | | | Function to find | Definite |
| PF13884 | | | Chaperone of endosialidase | Definite |
| PF00561 | CL0028 | AB_hydrolase | alpha/beta hydrolase fold | Likely |
| PF01738 | CL0028 | AB_hydrolase | Dienelactone hydrolase family | Likely |
| PF02551 | CL0050 | HotDog | Acyl-CoA thioesterase | Likely |
| PF03576 | CL0635 | DmpA_ArgJ | Peptidase_S58 | Likely |
| PF03633 | | | Glycosyl hydrolase family 65, C-terminal domain | Likely |
| PF06483 | | | Chitinase C | Likely |
| PF04301 | CL0028 | AB_hydrolase | Protein of unknown function (DUF452) | Probable |
| PF05057 | CL0028 | AB_hydrolase | Putative serine esterase (DUF676) | Probable |
| PF05277 | CL0028 | AB_hydrolase | Protein of unknown function (DUF726) | Probable |
| PF05677 | CL0028 | AB_hydrolase | Chlamydia CHLPS protein (DUF818) | Probable |
| PF05705 | CL0028 | AB_hydrolase | Eukaryotic protein of unknown function (DUF829) | Probable |
| PF05728 | CL0028 | AB_hydrolase | Uncharacterised protein family (UPF0227) | Probable |
| PF05990 | CL0028 | AB_hydrolase | Alpha/beta hydrolase of unknown function (DUF900) | Probable |
| PF06028 | CL0028 | AB_hydrolase | Alpha/beta hydrolase of unknown function (DUF915) | Probable |
| PF06057 | CL0028 | AB_hydrolase | Bacterial virulence protein (VirJ) | Probable |
| PF06342 | CL0028 | AB_hydrolase | Alpha/beta hydrolase of unknown function (DUF1057) | Probable |
| PF07082 | CL0028 | AB_hydrolase | Protein of unknown function (DUF1350) | Probable |
| PF07176 | CL0028 | AB_hydrolase | Alpha/beta hydrolase of unknown function (DUF1400) | Probable |
| PF08237 | CL0028 | AB_hydrolase | PE-PPE domain | Probable |
| PF08538 | CL0028 | AB_hydrolase | Protein of unknown function (DUF1749) | Probable |
| PF09752 | CL0028 | AB_hydrolase | Abhydrolase domain containing 18 | Probable |
| PF09994 | CL0028 | AB_hydrolase | Uncharacterized alpha/beta hydrolase domain (DUF223 | Probable |
| PF10142 | CL0028 | AB_hydrolase | PhoPQ-activated pathogenicity-related protein | Probable |
| PF11144 | CL0028 | AB_hydrolase | Protein of unknown function (DUF2920) | Probable |
| PF11187 | CL0028 | AB_hydrolase | Protein of unknown function (DUF2974) | Probable |
| PF11288 | CL0028 | AB_hydrolase | Protein of unknown function (DUF3089) | Probable |
| PF11339 | CL0028 | AB_hydrolase | Protein of unknown function (DUF3141) | Probable |
| PF12048 | CL0028 | AB_hydrolase | Protein of unknown function (DUF3530) | Probable |
| PF12715 | CL0028 | AB_hydrolase | Abhydrolase family | Probable |
| PF00472 | CL0337 | RF | RF-1 domain | Probable |
| PF04264 | | | YceI-like domain | Probable |
| PF18532 | | | Domain of unknown function (DUF5621) | Probable |
