## Supplemental Table S2 for "Chemoproteomic identification of a dipeptidyl peptidase 4 (DPP4) homolog in *Bacteroides thetaiotaomicron* important for envelope integrity and fitness"

Table S2: Gut commensal strains for bioinformatic predictions

| Strain | Phylum | UniProt Proteome ID |
|---|---|---|
| *Bifidobacterium adolescentis* ATCC 15703 | Actinobacteria | UP000008702 |
| *Bifidobacterium breve* DSM 20213 | Actinobacteria | UP000003191 |
| *Bifidobacterium longum* NCC 2705 | Actinobacteria | UP000000439 |
| *Bifidobacterium longum* subsp. *infantis* ATCC 15697 | Actinobacteria | UP000001360 |
| *Collinsella aerofaciens* ATCC 25986 | Actinobacteria | UP000464211 |
| *Eggerthella lenta* ATCC 25559 | Actinobacteria | UP000001377 |
| *Bacteroides caccae* ATCC 43185 | Bacteroidetes | UP000003325 |
| *Bacteroides cellulosilyticus* DSM 14838 | Bacteroidetes | UP000003711 |
| *Bacteroides finegoldii* DSM 17565 | Bacteroidetes | UP000003768 |
| *Bacteroides fragilis* ATCC 25285 | Bacteroidetes | UP000006731 |
| *Bacteroides intestinalis* DSM 17393 | Bacteroidetes | UP000004596 |
| *Bacteroides ovatus* ATCC 8483 | Bacteroidetes | UP000005475 |
| *Bacteroides stercoris* ATCC 43183 | Bacteroidetes | UP000004713 |
| *Bacteroides thetaiotaomicron* VPI-5482 | Bacteroidetes | UP000001414 |
| *Bacteroides uniformis* ATCC 8492 | Bacteroidetes | UP000004110 |
| *Parabacteroides distasonis* ATCC 8503 | Bacteroidetes | UP000000566 |
| *Parabacteroides johnsonii* DSM 18315 | Bacteroidetes | UP000005510 |
| *Parabacteroides merdae* ATCC 43184 | Bacteroidetes | UP000004276 |
| *Phocaeicola (Bacteroides) dorei* DSM 17855 | Bacteroidetes | UP000004849 |
| *Phocaeicola (Bacteroides) vulgatus* ATCC 8482 | Bacteroidetes | UP000002861 |
| *Prevotella copri* DSM 18205 | Bacteroidetes | UP000004477 |
| *Blautia hansenii* DSM 20583 | Firmicutes | UP000003755 |
| *Clostridium scindens* ATCC 35704 | Firmicutes | UP000003459 |
| *Clostridium sporogenes* ATCC 15579 | Firmicutes | UP000005747 |
| *Clostridium symbiosum* ATCC 14940 | Firmicutes | UP000016491 |
| *Dorea formicigenerans* ATCC 27755 | Firmicutes | UP000005359 |
| *Enterocloster (Clostridium) bolteae* ATCC BAA-613 | Firmicutes | UP000204173 |
| *Enterococcus faecalis* V583 | Firmicutes | UP000001415 |
| *Enterococcus faecium* ATCC BAA-472 | Firmicutes | UP000005269 |
| *Agathobacter (Eubacterium) rectale* ATCC 33656 | Firmicutes | UP000001477 |
| *Granulicatella adiacens* ATCC 49175 | Firmicutes | UP000005926 |
| *Hungatella (Clostridium) hathewayi* DSM 13479 | Firmicutes | UP000004968 |
| *Lacrimispora (Clostridium) saccharolytica* ATCC 35040 | Firmicutes | UP000001662 |
| *Lactiplantibacillus (Lactobacillus) plantarum* ATCC BAA-793 | Firmicutes | UP000000432 |
| *Ligilactobacillus (Lactobacillus) ruminis* ATCC 25644 | Firmicutes | UP000004099 |
| *Limosilactobacillus (Lactobacillus) reuteri* ATCC 55730 | Firmicutes | UP000001924 |
| *Mitsuokella multacida* DSM 20544 | Firmicutes | UP000003671 |
| *Roseburia intestinalis* L1-82 | Firmicutes | UP000004828 |
| *Ruminococcus gnavus* ATCC 29149 | Firmicutes | UP000004410 |
| *Fusobacterium nucleatum* subsp. *animalis* D11 | Fusobacteria | UP000004650 |
| *Fusobacterium nucleatum* subsp. *nucleatum* ATCC 23726 | Fusobacteria | UP000003643 |
| *Fusobacterium ulcerans* 12-1B | Fusobacteria | UP000003233 |
| *Edwardsiella tarda* ATCC 23685 | Proteobacteria | UP000003692 |
| *Enterobacter cancerogenus* ATCC 35316 | Proteobacteria | UP000003468 |
| *Escherichia coli* K12 MG1655 | Proteobacteria | UP000000625 |
| *Proteus penneri* ATCC 35198 | Proteobacteria | UP000006464 |
| *Providencia rettgeri* DSM 1131 | Proteobacteria | UP000003321 |
| *Providencia stuartii* ATCC 25827 | Proteobacteria | UP000004506 |
| *Akkermansia muciniphila* ATCC BAA-835 | Verrucomicrobia | UP000001031 |
| *Homo sapiens* | Chordata | UP000005640 |
