## Supplemental Table S3 for "Chemoproteomic identification of a dipeptidyl peptidase 4 (DPP4) homolog in *Bacteroides thetaiotaomicron* important for envelope integrity and fitness"

Table S3: Predicted serine hydrolases in gut commensal bacteria

| UniProt ID | Organism | Pfam | Pfam name | Evalue |
|---|---|---|---|---|
| Q88VI6 | LplantarumATCCBAA793 | PF00717.26 | Peptidase_S24 | 3.30E-29 |
| Q88XP7 | LplantarumATCCBAA793 | PF01425.24 | Amidase | 8.00E-154 |
| Q88YC0 | LplantarumATCCBAA793 | PF02129.21 | Peptidase_S15 | 2.10E-81 |
| Q88YH9 | LplantarumATCCBAA793 | PF00574.26 | CLP_protease | 2.30E-83 |
| F9UKV4 | LplantarumATCCBAA793 | PF13354.9 | Beta-lactamase2 | 1.20E-23 |
| F9UL51 | LplantarumATCCBAA793 | PF00975.23 | Thioesterase | 9.00E-12 |
| F9ULL8 | LplantarumATCCBAA793 | PF10502.12 | Peptidase_S26 | 8.00E-44 |
| F9ULR4 | LplantarumATCCBAA793 | PF13472.9 | Lipase_GDSL_2 | 6.90E-17 |
| F9UM18 | LplantarumATCCBAA793 | PF12146.11 | Hydrolase_4 | 8.60E-18 |
| F9UMG7 | LplantarumATCCBAA793 | PF07859.16 | Abhydrolase_3 | 1.10E-51 |
| F9UMT1 | LplantarumATCCBAA793 | PF12146.11 | Hydrolase_4 | 6.30E-09 |
| F9UMX0 | LplantarumATCCBAA793 | PF00144.27 | Beta-lactamase | 7.40E-36 |
| F9UN64 | LplantarumATCCBAA793 | PF00768.23 | Peptidase_S11 | 6.80E-58 |
| F9UNH0 | LplantarumATCCBAA793 | PF03572.21 | Peptidase_S41 | 1.80E-16 |
| F9UNV0 | LplantarumATCCBAA793 | PF01694.25 | Rhomboid | 4.90E-32 |
| F9UNV3 | LplantarumATCCBAA793 | PF03575.20 | Peptidase_S51 | 1.20E-18 |
| F9UP37 | LplantarumATCCBAA793 | PF00698.24 | Acyl_transf_1 | 1.40E-28 |
| F9UPB2 | LplantarumATCCBAA793 | PF00326.24 | Peptidase_S9 | 7.30E-11 |
| F9UPB5 | LplantarumATCCBAA793 | PF02016.18 | Peptidase_S66 | 6.80E-25 |
| F9UPB9 | LplantarumATCCBAA793 | PF02016.18 | Peptidase_S66 | 1.10E-24 |
| F9UPC3 | LplantarumATCCBAA793 | PF12697.10 | Abhydrolase_6 | 2.70E-10 |
| F9UPJ4 | LplantarumATCCBAA793 | PF03572.21 | Peptidase_S41 | 3.20E-48 |
| F9UPJ7 | LplantarumATCCBAA793 | PF13472.9 | Lipase_GDSL_2 | 3.40E-21 |
| F9UPY0 | LplantarumATCCBAA793 | PF00144.27 | Beta-lactamase | 4.10E-42 |
| F9UQ76 | LplantarumATCCBAA793 | PF05362.16 | Lon_C | 4.40E-09 |
| F9UQP5 | LplantarumATCCBAA793 | PF13354.9 | Beta-lactamase2 | 3.60E-48 |
| F9UQT0 | LplantarumATCCBAA793 | PF00144.27 | Beta-lactamase | 3.00E-54 |
| F9UQU5 | LplantarumATCCBAA793 | PF13472.9 | Lipase_GDSL_2 | 7.50E-12 |
| F9UQZ9 | LplantarumATCCBAA793 | PF04586.20 | Peptidase_S78 | 2.20E-49 |
| F9URC2 | LplantarumATCCBAA793 | PF12146.11 | Hydrolase_4 | 6.20E-08 |
| F9URD0 | LplantarumATCCBAA793 | PF00326.24 | Peptidase_S9 | 1.60E-07 |
| F9URF0 | LplantarumATCCBAA793 | PF00756.23 | Esterase | 3.20E-17 |
| F9URW3 | LplantarumATCCBAA793 | PF10502.12 | Peptidase_S26 | 4.50E-43 |
| F9URW4 | LplantarumATCCBAA793 | PF10502.12 | Peptidase_S26 | 2.30E-39 |
| F9US31 | LplantarumATCCBAA793 | PF12146.11 | Hydrolase_4 | 4.60E-17 |
| F9US66 | LplantarumATCCBAA793 | PF13365.9 | Trypsin_2 | 4.80E-28 |
| F9USU5 | LplantarumATCCBAA793 | PF12697.10 | Abhydrolase_6 | 4.30E-13 |
| F9UT98 | LplantarumATCCBAA793 | PF00768.23 | Peptidase_S11 | 3.30E-58 |
| F9UTF8 | LplantarumATCCBAA793 | PF12146.11 | Hydrolase_4 | 1.00E-12 |
| F9UTI8 | LplantarumATCCBAA793 | PF02674.19 | Colicin_V | 1.20E-26 |
| F9UTJ4 | LplantarumATCCBAA793 | PF00144.27 | Beta-lactamase | 6.90E-44 |
| F9UTK2 | LplantarumATCCBAA793 | PF13472.9 | Lipase_GDSL_2 | 1.50E-16 |
| F9UU44 | LplantarumATCCBAA793 | PF12697.10 | Abhydrolase_6 | 4.90E-08 |
| F9UU76 | LplantarumATCCBAA793 | PF12146.11 | Hydrolase_4 | 1.50E-10 |
| F9UUB1 | LplantarumATCCBAA793 | PF13472.9 | Lipase_GDSL_2 | 5.00E-11 |
| F9UUH1 | LplantarumATCCBAA793 | PF00756.23 | Esterase | 8.70E-24 |
| Q8G4R6 | BlongumNCC2705 | PF00717.26 | Peptidase_S24 | 1.60E-30 |
| Q8G5Q9 | BlongumNCC2705 | PF00574.26 | CLP_protease | 2.20E-70 |
| Q8G5R0 | BlongumNCC2705 | PF00574.26 | CLP_protease | 9.60E-75 |

| | | | | |
|---|---|---|---|---|
| Q8G768 | BlongumNCC2705 | PF01425.24 | Amidase | 1.80E-149 |
| Q8G3F7 | BlongumNCC2705 | PF07819.16 | PGAP1 | 5.70E-06 |
| Q8G3S5 | BlongumNCC2705 | PF02113.18 | Peptidase_S13 | 2.70E-43 |
| Q8G456 | BlongumNCC2705 | PF00698.24 | Acyl_transf_1 | 3.30E-35 |
| Q8G476 | BlongumNCC2705 | PF02230.19 | Abhydrolase_2 | 7.30E-17 |
| Q8G4Z5 | BlongumNCC2705 | PF12697.10 | Abhydrolase_6 | 5.70E-15 |
| Q8G4Z9 | BlongumNCC2705 | PF00326.24 | Peptidase_S9 | 3.70E-53 |
| Q8G5A7 | BlongumNCC2705 | PF07859.16 | Abhydrolase_3 | 2.00E-67 |
| Q8G5K7 | BlongumNCC2705 | PF00717.26 | Peptidase_S24 | 6.60E-17 |
| Q8G5V8 | BlongumNCC2705 | PF12697.10 | Abhydrolase_6 | 6.50E-18 |
| Q8G648 | BlongumNCC2705 | PF02230.19 | Abhydrolase_2 | 2.20E-12 |
| Q8G670 | BlongumNCC2705 | PF10502.12 | Peptidase_S26 | 4.60E-42 |
| Q8G6P4 | BlongumNCC2705 | PF01694.25 | Rhomboid | 3.40E-27 |
| Q8G6Q6 | BlongumNCC2705 | PF00326.24 | Peptidase_S9 | 6.20E-25 |
| Q8G6Q7 | BlongumNCC2705 | PF12146.11 | Hydrolase_4 | 1.70E-38 |
| Q8G6R7 | BlongumNCC2705 | PF00326.24 | Peptidase_S9 | 6.80E-07 |
| Q8G6T3 | BlongumNCC2705 | PF13365.9 | Trypsin_2 | 3.20E-31 |
| Q8G796 | BlongumNCC2705 | PF04586.20 | Peptidase_S78 | 5.60E-16 |
| Q8G7I8 | BlongumNCC2705 | PF10502.12 | Peptidase_S26 | 2.10E-40 |
| Q8G7Z0 | BlongumNCC2705 | PF05362.16 | Lon_C | 1.00E-12 |
| Q8G7Z9 | BlongumNCC2705 | PF01734.25 | Patatin | 1.90E-14 |
| Q8G810 | BlongumNCC2705 | PF02230.19 | Abhydrolase_2 | 8.80E-21 |
| A6LD45 | PdistasonisATCC8503 | PF05362.16 | Lon_C | 1.80E-83 |
| A6L837 | PdistasonisATCC8503 | PF03629.21 | SASA | 1.60E-20 |
| A6L880 | PdistasonisATCC8503 | PF13472.9 | Lipase_GDSL_2 | 2.90E-17 |
| A6L8H8 | PdistasonisATCC8503 | PF03572.21 | Peptidase_S41 | 6.40E-49 |
| A6L8K5 | PdistasonisATCC8503 | PF12146.11 | Hydrolase_4 | 1.80E-12 |
| A6L8L4 | PdistasonisATCC8503 | PF13472.9 | Lipase_GDSL_2 | 4.60E-15 |
| A6L8X8 | PdistasonisATCC8503 | PF13365.9 | Trypsin_2 | 5.80E-36 |
| A6L9E6 | PdistasonisATCC8503 | PF13472.9 | Lipase_GDSL_2 | 1.80E-15 |
| A6L9T6 | PdistasonisATCC8503 | PF01734.25 | Patatin | 8.80E-28 |
| A6L9W9 | PdistasonisATCC8503 | PF00326.24 | Peptidase_S9 | 3.90E-45 |
| A6LA14 | PdistasonisATCC8503 | PF13365.9 | Trypsin_2 | 1.70E-20 |
| A6LAA0 | PdistasonisATCC8503 | PF00717.26 | Peptidase_S24 | 2.60E-12 |
| A6LAL1 | PdistasonisATCC8503 | PF01694.25 | Rhomboid | 6.50E-26 |
| A6LAL3 | PdistasonisATCC8503 | PF13472.9 | Lipase_GDSL_2 | 3.80E-14 |
| A6LAN0 | PdistasonisATCC8503 | PF13472.9 | Lipase_GDSL_2 | 5.70E-28 |
| A6LAN4 | PdistasonisATCC8503 | PF02129.21 | Peptidase_S15 | 1.40E-10 |
| A6LAW0 | PdistasonisATCC8503 | PF01734.25 | Patatin | 4.90E-09 |
| A6LBC5 | PdistasonisATCC8503 | PF03572.21 | Peptidase_S41 | 1.70E-17 |
| A6LC23 | PdistasonisATCC8503 | PF00326.24 | Peptidase_S9 | 2.70E-54 |
| A6LC41 | PdistasonisATCC8503 | PF00144.27 | Beta-lactamase | 2.60E-31 |
| A6LC71 | PdistasonisATCC8503 | PF13365.9 | Trypsin_2 | 3.30E-07 |
| A6LCA7 | PdistasonisATCC8503 | PF12697.10 | Abhydrolase_6 | 3.10E-06 |
| A6LCU3 | PdistasonisATCC8503 | PF00698.24 | Acyl_transf_1 | 4.60E-43 |
| A6LCU4 | PdistasonisATCC8503 | PF00326.24 | Peptidase_S9 | 1.90E-10 |
| A6LDH8 | PdistasonisATCC8503 | PF00756.23 | Esterase | 4.50E-25 |
| A6LE10 | PdistasonisATCC8503 | PF00326.24 | Peptidase_S9 | 2.60E-55 |
| A6LE63 | PdistasonisATCC8503 | PF00326.24 | Peptidase_S9 | 5.20E-41 |
| A6LE97 | PdistasonisATCC8503 | PF00144.27 | Beta-lactamase | 7.30E-31 |
| A6LEE0 | PdistasonisATCC8503 | PF12146.11 | Hydrolase_4 | 2.90E-22 |
| A6LEE9 | PdistasonisATCC8503 | PF00144.27 | Beta-lactamase | 2.40E-46 |

| | | | | |
|---|---|---|---|---|
| A6LEK7 | PdistasonisATCC8503 | PF00574.26 | CLP_protease | 5.10E-79 |
| A6LEX8 | PdistasonisATCC8503 | PF00326.24 | Peptidase_S9 | 6.00E-43 |
| A6LFE2 | PdistasonisATCC8503 | PF00756.23 | Esterase | 9.80E-27 |
| A6LFF8 | PdistasonisATCC8503 | PF00082.25 | Peptidase_S8 | 5.90E-29 |
| A6LFH5 | PdistasonisATCC8503 | PF03572.21 | Peptidase_S41 | 3.70E-19 |
| A6LFI1 | PdistasonisATCC8503 | PF03572.21 | Peptidase_S41 | 6.30E-17 |
| A6LFJ2 | PdistasonisATCC8503 | PF00135.31 | COesterase | 3.40E-85 |
| A6LFL5 | PdistasonisATCC8503 | PF13365.9 | Trypsin_2 | 1.10E-06 |
| A6LFS8 | PdistasonisATCC8503 | PF00089.29 | Trypsin | 4.60E-05 |
| A6LFU7 | PdistasonisATCC8503 | PF10502.12 | Peptidase_S26 | 7.80E-29 |
| A6LFU8 | PdistasonisATCC8503 | PF10502.12 | Peptidase_S26 | 7.80E-32 |
| A6LFZ8 | PdistasonisATCC8503 | PF00089.29 | Trypsin | 3.70E-27 |
| A6LG07 | PdistasonisATCC8503 | PF10459.12 | Peptidase_S46 | 4.80E-242 |
| A6LG42 | PdistasonisATCC8503 | PF03629.21 | SASA | 8.40E-20 |
| A6LG42 | PdistasonisATCC8503 | PF13472.9 | Lipase_GDSL_2 | 4.00E-25 |
| A6LGD3 | PdistasonisATCC8503 | PF13884.9 | Peptidase_S74 | 6.90E-17 |
| A6LGD5 | PdistasonisATCC8503 | PF13884.9 | Peptidase_S74 | 1.20E-13 |
| A6LGG3 | PdistasonisATCC8503 | PF02230.19 | Abhydrolase_2 | 5.10E-13 |
| A6LGH2 | PdistasonisATCC8503 | PF05576.14 | Peptidase_S37 | 7.10E-51 |
| A6LGI6 | PdistasonisATCC8503 | PF00326.24 | Peptidase_S9 | 1.60E-47 |
| A6LGM2 | PdistasonisATCC8503 | PF13472.9 | Lipase_GDSL_2 | 1.10E-16 |
| A6LGN4 | PdistasonisATCC8503 | PF13472.9 | Lipase_GDSL_2 | 2.50E-15 |
| A6LGN7 | PdistasonisATCC8503 | PF00756.23 | Esterase | 2.70E-27 |
| A6LGP5 | PdistasonisATCC8503 | PF03572.21 | Peptidase_S41 | 7.50E-23 |
| A6LGT6 | PdistasonisATCC8503 | PF03572.21 | Peptidase_S41 | 3.30E-30 |
| A6LGV9 | PdistasonisATCC8503 | PF01734.25 | Patatin | 7.00E-29 |
| A6LH89 | PdistasonisATCC8503 | PF03572.21 | Peptidase_S41 | 9.20E-52 |
| A6LH98 | PdistasonisATCC8503 | PF02113.18 | Peptidase_S13 | 9.90E-70 |
| A6LHE1 | PdistasonisATCC8503 | PF00756.23 | Esterase | 4.70E-24 |
| A6LHG4 | PdistasonisATCC8503 | PF00082.25 | Peptidase_S8 | 2.40E-43 |
| A6LHH7 | PdistasonisATCC8503 | PF10502.12 | Peptidase_S26 | 5.80E-31 |
| A6LHK9 | PdistasonisATCC8503 | PF00326.24 | Peptidase_S9 | 2.40E-23 |
| A6LHS2 | PdistasonisATCC8503 | PF02674.19 | Colicin_V | 6.30E-18 |
| A6LHV3 | PdistasonisATCC8503 | PF00326.24 | Peptidase_S9 | 9.70E-22 |
| A6LHY7 | PdistasonisATCC8503 | PF01694.25 | Rhomboid | 1.20E-26 |
| A6LHY8 | PdistasonisATCC8503 | PF01694.25 | Rhomboid | 8.80E-22 |
| A6LI38 | PdistasonisATCC8503 | PF03572.21 | Peptidase_S41 | 7.20E-43 |
| A6LI91 | PdistasonisATCC8503 | PF00326.24 | Peptidase_S9 | 5.90E-34 |
| A6LID5 | PdistasonisATCC8503 | PF10459.12 | Peptidase_S46 | 9.50E-266 |
| A6LIE6 | PdistasonisATCC8503 | PF01734.25 | Patatin | 1.60E-27 |
| A6LIV8 | PdistasonisATCC8503 | PF01343.21 | Peptidase_S49 | 3.80E-45 |
| P00803 | EcoliK12MG1655 | PF10502.12 | Peptidase_S26 | 2.20E-58 |
| P00811 | EcoliK12MG1655 | PF00144.27 | Beta-lactamase | 1.60E-89 |
| P04335 | EcoliK12MG1655 | PF06500.14 | FrsA-like | 3.00E-235 |
| P07000 | EcoliK12MG1655 | PF12146.11 | Hydrolase_4 | 7.00E-59 |
| P08395 | EcoliK12MG1655 | PF01343.21 | Peptidase_S49 | 1.90E-53 |
| P08506 | EcoliK12MG1655 | PF00768.23 | Peptidase_S11 | 2.30E-105 |
| P08550 | EcoliK12MG1655 | PF02674.19 | Colicin_V | 4.60E-35 |
| P09391 | EcoliK12MG1655 | PF01694.25 | Rhomboid | 5.00E-29 |
| P0A6G7 | EcoliK12MG1655 | PF00574.26 | CLP_protease | 1.80E-88 |
| P0A7C2 | EcoliK12MG1655 | PF00717.26 | Peptidase_S24 | 2.30E-32 |
| P0A7C6 | EcoliK12MG1655 | PF03575.20 | Peptidase_S51 | 1.30E-63 |

| | | | | |
|---|---|---|---|---|
| P0A9M0 | EcoliK12MG1655 | PF05362.16 | Lon_C | 1.00E-97 |
| P0AAI9 | EcoliK12MG1655 | PF00698.24 | Acyl_transf_1 | 3.90E-39 |
| P0AD70 | EcoliK12MG1655 | PF00144.27 | Beta-lactamase | 2.20E-64 |
| P0ADA1 | EcoliK12MG1655 | PF13472.9 | Lipase_GDSL_2 | 6.50E-26 |
| P0AEB2 | EcoliK12MG1655 | PF00768.23 | Peptidase_S11 | 2.20E-104 |
| P0AEE3 | EcoliK12MG1655 | PF13365.9 | Trypsin_2 | 8.10E-29 |
| P0AFI5 | EcoliK12MG1655 | PF00768.23 | Peptidase_S11 | 2.80E-90 |
| P0AFR0 | EcoliK12MG1655 | PF01734.25 | Patatin | 1.90E-23 |
| P0AG11 | EcoliK12MG1655 | PF00717.26 | Peptidase_S24 | 1.80E-27 |
| P0AG14 | EcoliK12MG1655 | PF01343.21 | Peptidase_S49 | 6.60E-51 |
| P0C0V0 | EcoliK12MG1655 | PF13365.9 | Trypsin_2 | 3.90E-31 |
| P11454 | EcoliK12MG1655 | PF00975.23 | Thioesterase | 1.90E-51 |
| P13039 | EcoliK12MG1655 | PF00756.23 | Esterase | 1.10E-49 |
| P23865 | EcoliK12MG1655 | PF03572.21 | Peptidase_S41 | 2.00E-53 |
| P23872 | EcoliK12MG1655 | PF07859.16 | Abhydrolase_3 | 4.10E-53 |
| P24228 | EcoliK12MG1655 | PF02113.18 | Peptidase_S13 | 2.70E-178 |
| P24555 | EcoliK12MG1655 | PF00326.24 | Peptidase_S9 | 6.50E-67 |
| P31471 | EcoliK12MG1655 | PF00756.23 | Esterase | 2.80E-18 |
| P33013 | EcoliK12MG1655 | PF00768.23 | Peptidase_S11 | 1.80E-102 |
| P33018 | EcoliK12MG1655 | PF00756.23 | Esterase | 3.50E-72 |
| P37355 | EcoliK12MG1655 | PF12697.10 | Abhydrolase_6 | 8.00E-18 |
| P39099 | EcoliK12MG1655 | PF13365.9 | Trypsin_2 | 4.70E-29 |
| P39298 | EcoliK12MG1655 | PF00326.24 | Peptidase_S9 | 2.70E-11 |
| P39370 | EcoliK12MG1655 | PF03629.21 | SASA | 1.90E-12 |
| P39407 | EcoliK12MG1655 | PF01734.25 | Patatin | 1.70E-20 |
| P51025 | EcoliK12MG1655 | PF00756.23 | Esterase | 2.00E-76 |
| P75736 | EcoliK12MG1655 | PF12697.10 | Abhydrolase_6 | 2.60E-17 |
| P75867 | EcoliK12MG1655 | PF05362.16 | Lon_C | 4.00E-10 |
| P75895 | EcoliK12MG1655 | PF12697.10 | Abhydrolase_6 | 6.60E-17 |
| P75974 | EcoliK12MG1655 | PF00717.26 | Peptidase_S24 | 2.70E-27 |
| P76008 | EcoliK12MG1655 | PF02016.18 | Peptidase_S66 | 2.10E-19 |
| P76049 | EcoliK12MG1655 | PF02129.21 | Peptidase_S15 | 1.60E-06 |
| P76092 | EcoliK12MG1655 | PF12146.11 | Hydrolase_4 | 1.30E-63 |
| P76176 | EcoliK12MG1655 | PF00089.29 | Trypsin | 6.60E-11 |
| P76561 | EcoliK12MG1655 | PF02230.19 | Abhydrolase_2 | 1.50E-56 |
| P77044 | EcoliK12MG1655 | PF12697.10 | Abhydrolase_6 | 9.80E-23 |
| P77538 | EcoliK12MG1655 | PF12146.11 | Hydrolase_4 | 1.30E-12 |
| P77619 | EcoliK12MG1655 | PF00144.27 | Beta-lactamase | 5.40E-70 |
| B2UMP4 | AmuciniphilaATCCBAA835 | PF01425.24 | Amidase | 2.90E-151 |
| B2UKN7 | AmuciniphilaATCCBAA835 | PF01343.21 | Peptidase_S49 | 7.20E-13 |
| B2UL14 | AmuciniphilaATCCBAA835 | PF10502.12 | Peptidase_S26 | 8.20E-25 |
| B2UL29 | AmuciniphilaATCCBAA835 | PF00768.23 | Peptidase_S11 | 2.60E-33 |
| B2UL55 | AmuciniphilaATCCBAA835 | PF00698.24 | Acyl_transf_1 | 4.80E-36 |
| B2UL80 | AmuciniphilaATCCBAA835 | PF13472.9 | Lipase_GDSL_2 | 3.40E-17 |
| B2ULH8 | AmuciniphilaATCCBAA835 | PF13472.9 | Lipase_GDSL_2 | 5.10E-21 |
| B2ULI0 | AmuciniphilaATCCBAA835 | PF13472.9 | Lipase_GDSL_2 | 5.20E-20 |
| B2ULQ7 | AmuciniphilaATCCBAA835 | PF00144.27 | Beta-lactamase | 9.40E-35 |
| B2ULS4 | AmuciniphilaATCCBAA835 | PF03629.21 | SASA | 1.50E-09 |
| B2ULS5 | AmuciniphilaATCCBAA835 | PF13472.9 | Lipase_GDSL_2 | 7.90E-19 |
| B2ULX6 | AmuciniphilaATCCBAA835 | PF13365.9 | Trypsin_2 | 2.80E-23 |
| B2UM07 | AmuciniphilaATCCBAA835 | PF03572.21 | Peptidase_S41 | 7.60E-47 |
| B2UM46 | AmuciniphilaATCCBAA835 | PF03629.21 | SASA | 4.70E-28 |

| | | | | |
|---|---|---|---|---|
| B2UM78 | AmuciniphilaATCCBAA835 | PF13354.9 | Beta-lactamase2 | 3.00E-38 |
| B2UMQ3 | AmuciniphilaATCCBAA835 | PF13365.9 | Trypsin_2 | 1.20E-20 |
| B2UMQ6 | AmuciniphilaATCCBAA835 | PF00717.26 | Peptidase_S24 | 5.80E-12 |
| B2UMX7 | AmuciniphilaATCCBAA835 | PF13472.9 | Lipase_GDSL_2 | 3.70E-18 |
| B2UN08 | AmuciniphilaATCCBAA835 | PF02129.21 | Peptidase_S15 | 2.60E-10 |
| B2UNH0 | AmuciniphilaATCCBAA835 | PF00768.23 | Peptidase_S11 | 3.50E-53 |
| B2UNI0 | AmuciniphilaATCCBAA835 | PF03629.21 | SASA | 1.10E-11 |
| B2UNI1 | AmuciniphilaATCCBAA835 | PF13472.9 | Lipase_GDSL_2 | 7.30E-25 |
| B2UNY7 | AmuciniphilaATCCBAA835 | PF07859.16 | Abhydrolase_3 | 2.30E-10 |
| B2UP84 | AmuciniphilaATCCBAA835 | PF13472.9 | Lipase_GDSL_2 | 5.80E-14 |
| B2UPK1 | AmuciniphilaATCCBAA835 | PF13472.9 | Lipase_GDSL_2 | 2.50E-08 |
| B2UPX1 | AmuciniphilaATCCBAA835 | PF13365.9 | Trypsin_2 | 3.40E-22 |
| B2UPX2 | AmuciniphilaATCCBAA835 | PF13365.9 | Trypsin_2 | 1.30E-08 |
| B2UQ86 | AmuciniphilaATCCBAA835 | PF01694.25 | Rhomboid | 9.30E-36 |
| B2UQA8 | AmuciniphilaATCCBAA835 | PF13365.9 | Trypsin_2 | 7.40E-30 |
| B2UQD5 | AmuciniphilaATCCBAA835 | PF03629.21 | SASA | 1.00E-13 |
| B2UQV8 | AmuciniphilaATCCBAA835 | PF12697.10 | Abhydrolase_6 | 2.50E-10 |
| B2UQW3 | AmuciniphilaATCCBAA835 | PF13472.9 | Lipase_GDSL_2 | 1.40E-17 |
| B2UQZ2 | AmuciniphilaATCCBAA835 | PF00574.26 | CLP_protease | 6.10E-79 |
| B2URJ9 | AmuciniphilaATCCBAA835 | PF13472.9 | Lipase_GDSL_2 | 3.10E-24 |
| B7GQ64 | BinfantisATCC15697 | PF00717.26 | Peptidase_S24 | 1.60E-30 |
| B7GTU6 | BinfantisATCC15697 | PF01425.24 | Amidase | 2.20E-149 |
| B7GN36 | BinfantisATCC15697 | PF10502.12 | Peptidase_S26 | 6.00E-40 |
| B7GNI6 | BinfantisATCC15697 | PF00698.24 | Acyl_transf_1 | 3.40E-35 |
| B7GNX1 | BinfantisATCC15697 | PF00326.24 | Peptidase_S9 | 6.00E-15 |
| B7GP47 | BinfantisATCC15697 | PF02113.18 | Peptidase_S13 | 5.90E-43 |
| B7GPC2 | BinfantisATCC15697 | PF02230.19 | Abhydrolase_2 | 1.00E-16 |
| B7GPQ7 | BinfantisATCC15697 | PF07819.16 | PGAP1 | 5.10E-06 |
| B7GPW7 | BinfantisATCC15697 | PF02230.19 | Abhydrolase_2 | 4.80E-22 |
| B7GPZ0 | BinfantisATCC15697 | PF01734.25 | Patatin | 1.20E-14 |
| B7GPZ9 | BinfantisATCC15697 | PF05362.16 | Lon_C | 1.60E-12 |
| B7GQH2 | BinfantisATCC15697 | PF13472.9 | Lipase_GDSL_2 | 3.90E-21 |
| B7GRW1 | BinfantisATCC15697 | PF02230.19 | Abhydrolase_2 | 1.60E-13 |
| B7GSC2 | BinfantisATCC15697 | PF03575.20 | Peptidase_S51 | 9.00E-44 |
| B7GSM4 | BinfantisATCC15697 | PF01694.25 | Rhomboid | 3.00E-27 |
| B7GSN6 | BinfantisATCC15697 | PF00326.24 | Peptidase_S9 | 1.10E-37 |
| B7GSN7 | BinfantisATCC15697 | PF12146.11 | Hydrolase_4 | 1.50E-38 |
| B7GSN9 | BinfantisATCC15697 | PF00756.23 | Esterase | 4.20E-07 |
| B7GSQ3 | BinfantisATCC15697 | PF13365.9 | Trypsin_2 | 2.50E-31 |
| B7GSV3 | BinfantisATCC15697 | PF00574.26 | CLP_protease | 2.50E-74 |
| B7GSV4 | BinfantisATCC15697 | PF00574.26 | CLP_protease | 5.10E-70 |
| B7GT09 | BinfantisATCC15697 | PF00717.26 | Peptidase_S24 | 1.40E-17 |
| B7GU33 | BinfantisATCC15697 | PF03629.21 | SASA | 1.40E-05 |
| B7GU66 | BinfantisATCC15697 | PF07859.16 | Abhydrolase_3 | 1.10E-64 |
| B7GUI9 | BinfantisATCC15697 | PF00326.24 | Peptidase_S9 | 4.30E-53 |
| B7GUJ3 | BinfantisATCC15697 | PF12697.10 | Abhydrolase_6 | 1.00E-15 |
| C8WGC6 | ElentaATCC25559 | PF03575.20 | Peptidase_S51 | 2.80E-42 |
| C8WGN7 | ElentaATCC25559 | PF00768.23 | Peptidase_S11 | 1.20E-32 |
| C8WGW0 | ElentaATCC25559 | PF10502.12 | Peptidase_S26 | 4.80E-38 |
| C8WGW1 | ElentaATCC25559 | PF10502.12 | Peptidase_S26 | 1.50E-45 |
| C8WH27 | ElentaATCC25559 | PF01694.25 | Rhomboid | 2.90E-29 |
| C8WHF7 | ElentaATCC25559 | PF00768.23 | Peptidase_S11 | 1.30E-40 |

| | | | | |
|---|---|---|---|---|
| C8WHI8 | ElentaATCC25559 | PF00717.26 | Peptidase_S24 | 1.60E-29 |
| C8WHY8 | ElentaATCC25559 | PF00082.25 | Peptidase_S8 | 5.20E-39 |
| C8WI12 | ElentaATCC25559 | PF00144.27 | Beta-lactamase | 1.70E-51 |
| C8WI58 | ElentaATCC25559 | PF07859.16 | Abhydrolase_3 | 1.20E-50 |
| C8WIF2 | ElentaATCC25559 | PF00574.26 | CLP_protease | 3.80E-78 |
| C8WIL2 | ElentaATCC25559 | PF00698.24 | Acyl_transf_1 | 9.90E-38 |
| C8WIL8 | ElentaATCC25559 | PF03572.21 | Peptidase_S41 | 3.90E-42 |
| C8WJI0 | ElentaATCC25559 | PF13472.9 | Lipase_GDSL_2 | 7.00E-21 |
| C8WJL5 | ElentaATCC25559 | PF13472.9 | Lipase_GDSL_2 | 4.80E-08 |
| C8WK60 | ElentaATCC25559 | PF01734.25 | Patatin | 1.20E-13 |
| C8WKA0 | ElentaATCC25559 | PF07859.16 | Abhydrolase_3 | 3.60E-66 |
| C8WKG4 | ElentaATCC25559 | PF13354.9 | Beta-lactamase2 | 2.20E-36 |
| C8WL53 | ElentaATCC25559 | PF13365.9 | Trypsin_2 | 2.80E-30 |
| C8WMB7 | ElentaATCC25559 | PF01764.28 | Lipase_3 | 4.50E-12 |
| C8WMC1 | ElentaATCC25559 | PF12695.10 | Abhydrolase_5 | 2.50E-53 |
| C8WMC6 | ElentaATCC25559 | PF12146.11 | Hydrolase_4 | 4.80E-66 |
| C8WMD2 | ElentaATCC25559 | PF12697.10 | Abhydrolase_6 | 3.10E-17 |
| C8WMU6 | ElentaATCC25559 | PF00082.25 | Peptidase_S8 | 7.00E-13 |
| C8WMW0 | ElentaATCC25559 | PF01425.24 | Amidase | 3.00E-32 |
| C8WNU0 | ElentaATCC25559 | PF00082.25 | Peptidase_S8 | 1.30E-33 |
| C8WNX3 | ElentaATCC25559 | PF12697.10 | Abhydrolase_6 | 2.50E-22 |
| C8WPB6 | ElentaATCC25559 | PF12146.11 | Hydrolase_4 | 1.50E-51 |
| C8WPF6 | ElentaATCC25559 | PF01425.24 | Amidase | 1.60E-136 |
| C8WPH2 | ElentaATCC25559 | PF01343.21 | Peptidase_S49 | 1.20E-41 |
| Q8A129 | BthetaiotaomicronVPI5482 | PF00574.26 | CLP_protease | 1.30E-77 |
| Q89Z70 | BthetaiotaomicronVPI5482 | PF13354.9 | Beta-lactamase2 | 1.70E-31 |
| Q89Z75 | BthetaiotaomicronVPI5482 | PF13472.9 | Lipase_GDSL_2 | 2.60E-13 |
| Q89ZS4 | BthetaiotaomicronVPI5482 | PF01694.25 | Rhomboid | 6.40E-13 |
| Q89ZW2 | BthetaiotaomicronVPI5482 | PF03572.21 | Peptidase_S41 | 1.60E-52 |
| Q8A007 | BthetaiotaomicronVPI5482 | PF00326.24 | Peptidase_S9 | 3.80E-15 |
| Q8A028 | BthetaiotaomicronVPI5482 | PF00326.24 | Peptidase_S9 | 1.70E-59 |
| Q8A041 | BthetaiotaomicronVPI5482 | PF03629.21 | SASA | 6.60E-61 |
| Q8A042 | BthetaiotaomicronVPI5482 | PF03629.21 | SASA | 2.50E-14 |
| Q8A048 | BthetaiotaomicronVPI5482 | PF13472.9 | Lipase_GDSL_2 | 5.10E-13 |
| Q8A063 | BthetaiotaomicronVPI5482 | PF13472.9 | Lipase_GDSL_2 | 1.10E-13 |
| Q8A071 | BthetaiotaomicronVPI5482 | PF13472.9 | Lipase_GDSL_2 | 2.90E-13 |
| Q8A0B1 | BthetaiotaomicronVPI5482 | PF13472.9 | Lipase_GDSL_2 | 7.30E-11 |
| Q8A0C5 | BthetaiotaomicronVPI5482 | PF13472.9 | Lipase_GDSL_2 | 6.30E-16 |
| Q8A0D0 | BthetaiotaomicronVPI5482 | PF03629.21 | SASA | 1.10E-10 |
| Q8A0E3 | BthetaiotaomicronVPI5482 | PF13472.9 | Lipase_GDSL_2 | 8.60E-16 |
| Q8A0E4 | BthetaiotaomicronVPI5482 | PF00756.23 | Esterase | 6.20E-34 |
| Q8A0E6 | BthetaiotaomicronVPI5482 | PF13472.9 | Lipase_GDSL_2 | 1.80E-13 |
| Q8A0Y2 | BthetaiotaomicronVPI5482 | PF00082.25 | Peptidase_S8 | 1.10E-37 |
| Q8A140 | BthetaiotaomicronVPI5482 | PF13884.9 | Peptidase_S74 | 7.50E-14 |
| Q8A1Q1 | BthetaiotaomicronVPI5482 | PF00326.24 | Peptidase_S9 | 6.60E-06 |
| Q8A1R2 | BthetaiotaomicronVPI5482 | PF03629.21 | SASA | 1.60E-19 |
| Q8A1T1 | BthetaiotaomicronVPI5482 | PF03572.21 | Peptidase_S41 | 7.10E-33 |
| Q8A2A0 | BthetaiotaomicronVPI5482 | PF02674.19 | Colicin_V | 7.10E-32 |
| Q8A2I6 | BthetaiotaomicronVPI5482 | PF10502.12 | Peptidase_S26 | 4.60E-25 |
| Q8A2I7 | BthetaiotaomicronVPI5482 | PF10502.12 | Peptidase_S26 | 1.20E-17 |
| Q8A2L4 | BthetaiotaomicronVPI5482 | PF03572.21 | Peptidase_S41 | 7.50E-09 |
| Q8A2L6 | BthetaiotaomicronVPI5482 | PF10459.12 | Peptidase_S46 | 1.50E-266 |

| | | | | |
|---|---|---|---|---|
| Q8A2L9 | BthetaiotaomicronVPI5482 | PF02230.19 | Abhydrolase_2 | 6.50E-11 |
| Q8A2Q1 | BthetaiotaomicronVPI5482 | PF00326.24 | Peptidase_S9 | 5.00E-44 |
| Q8A2V9 | BthetaiotaomicronVPI5482 | PF00717.26 | Peptidase_S24 | 7.20E-18 |
| Q8A2W8 | BthetaiotaomicronVPI5482 | PF02113.18 | Peptidase_S13 | 6.60E-87 |
| Q8A2Y4 | BthetaiotaomicronVPI5482 | PF03629.21 | SASA | 3.90E-19 |
| Q8A323 | BthetaiotaomicronVPI5482 | PF03629.21 | SASA | 3.40E-12 |
| Q8A331 | BthetaiotaomicronVPI5482 | PF03629.21 | SASA | 7.80E-22 |
| Q8A343 | BthetaiotaomicronVPI5482 | PF12146.11 | Hydrolase_4 | 1.20E-16 |
| Q8A355 | BthetaiotaomicronVPI5482 | PF00326.24 | Peptidase_S9 | 5.20E-07 |
| Q8A3B9 | BthetaiotaomicronVPI5482 | PF03572.21 | Peptidase_S41 | 3.50E-52 |
| Q8A3J3 | BthetaiotaomicronVPI5482 | PF13472.9 | Lipase_GDSL_2 | 1.20E-16 |
| Q8A3M2 | BthetaiotaomicronVPI5482 | PF00717.26 | Peptidase_S24 | 1.00E-20 |
| Q8A3X2 | BthetaiotaomicronVPI5482 | PF01694.25 | Rhomboid | 6.40E-24 |
| Q8A3X3 | BthetaiotaomicronVPI5482 | PF01694.25 | Rhomboid | 1.40E-32 |
| Q8A406 | BthetaiotaomicronVPI5482 | PF03572.21 | Peptidase_S41 | 3.60E-50 |
| Q8A430 | BthetaiotaomicronVPI5482 | PF13365.9 | Trypsin_2 | 3.80E-15 |
| Q8A4C1 | BthetaiotaomicronVPI5482 | PF13472.9 | Lipase_GDSL_2 | 2.20E-13 |
| Q8A4H8 | BthetaiotaomicronVPI5482 | PF13472.9 | Lipase_GDSL_2 | 3.60E-17 |
| Q8A4M2 | BthetaiotaomicronVPI5482 | PF13365.9 | Trypsin_2 | 6.20E-09 |
| Q8A4P8 | BthetaiotaomicronVPI5482 | PF02016.18 | Peptidase_S66 | 2.50E-37 |
| Q8A4S2 | BthetaiotaomicronVPI5482 | PF05448.15 | AXE1 | 5.40E-51 |
| Q8A508 | BthetaiotaomicronVPI5482 | PF00144.27 | Beta-lactamase | 4.40E-48 |
| Q8A511 | BthetaiotaomicronVPI5482 | PF05576.14 | Peptidase_S37 | 5.70E-62 |
| Q8A5K0 | BthetaiotaomicronVPI5482 | PF03572.21 | Peptidase_S41 | 3.20E-27 |
| Q8A6K2 | BthetaiotaomicronVPI5482 | PF01343.21 | Peptidase_S49 | 2.90E-48 |
| Q8A6L4 | BthetaiotaomicronVPI5482 | PF13884.9 | Peptidase_S74 | 7.50E-14 |
| Q8A6N9 | BthetaiotaomicronVPI5482 | PF00326.24 | Peptidase_S9 | 1.40E-47 |
| Q8A6R9 | BthetaiotaomicronVPI5482 | PF01694.25 | Rhomboid | 2.00E-31 |
| Q8A6U4 | BthetaiotaomicronVPI5482 | PF01734.25 | Patatin | 9.60E-24 |
| Q8A6U7 | BthetaiotaomicronVPI5482 | PF03629.21 | SASA | 2.30E-10 |
| Q8A6Y2 | BthetaiotaomicronVPI5482 | PF13472.9 | Lipase_GDSL_2 | 4.90E-17 |
| Q8A7W6 | BthetaiotaomicronVPI5482 | PF03572.21 | Peptidase_S41 | 2.60E-19 |
| Q8A815 | BthetaiotaomicronVPI5482 | PF13365.9 | Trypsin_2 | 5.00E-06 |
| Q8A862 | BthetaiotaomicronVPI5482 | PF13365.9 | Trypsin_2 | 1.00E-33 |
| Q8A8A6 | BthetaiotaomicronVPI5482 | PF00326.24 | Peptidase_S9 | 1.20E-45 |
| Q8A8H5 | BthetaiotaomicronVPI5482 | PF13472.9 | Lipase_GDSL_2 | 1.70E-24 |
| Q8A8P6 | BthetaiotaomicronVPI5482 | PF02129.21 | Peptidase_S15 | 2.70E-07 |
| Q8A8Q3 | BthetaiotaomicronVPI5482 | PF12697.10 | Abhydrolase_6 | 2.70E-07 |
| Q8A8T0 | BthetaiotaomicronVPI5482 | PF07859.16 | Abhydrolase_3 | 1.60E-13 |
| Q8A901 | BthetaiotaomicronVPI5482 | PF01734.25 | Patatin | 4.20E-21 |
| Q8A909 | BthetaiotaomicronVPI5482 | PF00326.24 | Peptidase_S9 | 9.00E-14 |
| Q8A932 | BthetaiotaomicronVPI5482 | PF03629.21 | SASA | 1.80E-16 |
| Q8A9B9 | BthetaiotaomicronVPI5482 | PF01734.25 | Patatin | 2.40E-27 |
| Q8A9H9 | BthetaiotaomicronVPI5482 | PF05362.16 | Lon_C | 3.50E-58 |
| Q8A9M6 | BthetaiotaomicronVPI5482 | PF00698.24 | Acyl_transf_1 | 1.90E-36 |
| Q8A9P1 | BthetaiotaomicronVPI5482 | PF01734.25 | Patatin | 5.80E-21 |
| Q8A9Q0 | BthetaiotaomicronVPI5482 | PF13365.9 | Trypsin_2 | 2.30E-24 |
| Q8A9Z8 | BthetaiotaomicronVPI5482 | PF10502.12 | Peptidase_S26 | 1.00E-25 |
| Q8AA78 | BthetaiotaomicronVPI5482 | PF00326.24 | Peptidase_S9 | 4.00E-56 |
| Q8AA96 | BthetaiotaomicronVPI5482 | PF13472.9 | Lipase_GDSL_2 | 2.10E-21 |
| Q8AAK7 | BthetaiotaomicronVPI5482 | PF03629.21 | SASA | 2.60E-17 |
| Q8AAK7 | BthetaiotaomicronVPI5482 | PF13472.9 | Lipase_GDSL_2 | 2.50E-25 |

| | | | | |
|---|---|---|---|---|
| Q8AAL7 | BthetaiotaomicronVPI5482 | PF03629.21 | SASA | 3.50E-19 |
| Q8AAL9 | BthetaiotaomicronVPI5482 | PF13472.9 | Lipase_GDSL_2 | 3.00E-14 |
| Q8AB09 | BthetaiotaomicronVPI5482 | PF01734.25 | Patatin | 4.50E-26 |
| Q8AB73 | BthetaiotaomicronVPI5482 | PF10459.12 | Peptidase_S46 | 5.20E-256 |
| Q8AB74 | BthetaiotaomicronVPI5482 | PF10459.12 | Peptidase_S46 | 3.80E-253 |
| Q8AB98 | BthetaiotaomicronVPI5482 | PF00082.25 | Peptidase_S8 | 1.40E-42 |
| Q8ABF6 | BthetaiotaomicronVPI5482 | PF03572.21 | Peptidase_S41 | 4.60E-13 |
| Q8ABF8 | BthetaiotaomicronVPI5482 | PF00756.23 | Esterase | 3.50E-30 |
| Q834R0 | EfaecalisV583 | PF00717.26 | Peptidase_S24 | 1.80E-29 |
| Q837R0 | EfaecalisV583 | PF00574.26 | CLP_protease | 8.90E-87 |
| Q837V3 | EfaecalisV583 | PF01425.24 | Amidase | 3.60E-152 |
| H7C7B7 | EfaecalisV583 | PF00768.23 | Peptidase_S11 | 3.10E-51 |
| Q820U6 | EfaecalisV583 | PF13472.9 | Lipase_GDSL_2 | 2.70E-27 |
| Q82ZI3 | EfaecalisV583 | PF10502.12 | Peptidase_S26 | 8.20E-49 |
| Q82ZM6 | EfaecalisV583 | PF13365.9 | Trypsin_2 | 2.80E-31 |
| Q82ZR5 | EfaecalisV583 | PF01425.24 | Amidase | 8.50E-72 |
| Q82ZT4 | EfaecalisV583 | PF07859.16 | Abhydrolase_3 | 9.80E-24 |
| Q830B1 | EfaecalisV583 | PF00698.24 | Acyl_transf_1 | 1.00E-41 |
| Q830G5 | EfaecalisV583 | PF04586.20 | Peptidase_S78 | 4.20E-61 |
| Q830J0 | EfaecalisV583 | PF00326.24 | Peptidase_S9 | 5.90E-08 |
| Q830J2 | EfaecalisV583 | PF01694.25 | Rhomboid | 2.40E-35 |
| Q830Z6 | EfaecalisV583 | PF12146.11 | Hydrolase_4 | 9.20E-13 |
| Q831C1 | EfaecalisV583 | PF00144.27 | Beta-lactamase | 3.10E-50 |
| Q831Q0 | EfaecalisV583 | PF05362.16 | Lon_C | 9.50E-06 |
| Q833V8 | EfaecalisV583 | PF13365.9 | Trypsin_2 | 5.10E-15 |
| Q834H3 | EfaecalisV583 | PF03572.21 | Peptidase_S41 | 2.10E-51 |
| Q834H4 | EfaecalisV583 | PF10502.12 | Peptidase_S26 | 2.60E-49 |
| Q834I2 | EfaecalisV583 | PF02230.19 | Abhydrolase_2 | 1.70E-12 |
| Q834U6 | EfaecalisV583 | PF12146.11 | Hydrolase_4 | 7.80E-14 |
| Q834X2 | EfaecalisV583 | PF12146.11 | Hydrolase_4 | 1.10E-08 |
| Q834X5 | EfaecalisV583 | PF13354.9 | Beta-lactamase2 | 1.60E-43 |
| Q835H4 | EfaecalisV583 | PF02674.19 | Colicin_V | 1.50E-31 |
| Q835Y2 | EfaecalisV583 | PF05448.15 | AXE1 | 6.70E-53 |
| Q836K0 | EfaecalisV583 | PF10502.12 | Peptidase_S26 | 1.20E-50 |
| Q836S5 | EfaecalisV583 | PF01425.24 | Amidase | 1.50E-63 |
| Q837I5 | EfaecalisV583 | PF10502.12 | Peptidase_S26 | 8.30E-42 |
| Q837P7 | EfaecalisV583 | PF00756.23 | Esterase | 3.00E-26 |
| Q837T4 | EfaecalisV583 | PF00144.27 | Beta-lactamase | 7.60E-38 |
| Q837U3 | EfaecalisV583 | PF01425.24 | Amidase | 2.60E-62 |
| Q838I7 | EfaecalisV583 | PF00144.27 | Beta-lactamase | 3.70E-32 |
| Q838J8 | EfaecalisV583 | PF12697.10 | Abhydrolase_6 | 9.20E-17 |
| Q838Q5 | EfaecalisV583 | PF07859.16 | Abhydrolase_3 | 1.60E-12 |
| Q839A3 | EfaecalisV583 | PF12695.10 | Abhydrolase_5 | 2.10E-78 |
| Q839J6 | EfaecalisV583 | PF13472.9 | Lipase_GDSL_2 | 3.10E-27 |
| Q839Q7 | EfaecalisV583 | PF12146.11 | Hydrolase_4 | 2.30E-50 |
| C4ZGF4 | ArectaleATCC33656 | PF00574.26 | CLP_protease | 2.90E-87 |
| C4ZHB9 | ArectaleATCC33656 | PF01425.24 | Amidase | 3.20E-159 |
| C4Z801 | ArectaleATCC33656 | PF00717.26 | Peptidase_S24 | 2.20E-28 |
| C4Z845 | ArectaleATCC33656 | PF13472.9 | Lipase_GDSL_2 | 5.20E-14 |
| C4Z846 | ArectaleATCC33656 | PF13472.9 | Lipase_GDSL_2 | 2.20E-17 |
| C4Z8D0 | ArectaleATCC33656 | PF00768.23 | Peptidase_S11 | 8.10E-38 |
| C4Z8M3 | ArectaleATCC33656 | PF10502.12 | Peptidase_S26 | 5.00E-45 |

| | | | | |
|---|---|---|---|---|
| C4Z8P6 | ArectaleATCC33656 | PF00698.24 | Acyl_transf_1 | 6.60E-37 |
| C4Z934 | ArectaleATCC33656 | PF10502.12 | Peptidase_S26 | 5.10E-39 |
| C4Z9E7 | ArectaleATCC33656 | PF00768.23 | Peptidase_S11 | 9.20E-44 |
| C4Z9K8 | ArectaleATCC33656 | PF01694.25 | Rhomboid | 8.70E-34 |
| C4ZA04 | ArectaleATCC33656 | PF00717.26 | Peptidase_S24 | 3.40E-33 |
| C4ZA40 | ArectaleATCC33656 | PF00768.23 | Peptidase_S11 | 4.00E-49 |
| C4ZAT6 | ArectaleATCC33656 | PF10502.12 | Peptidase_S26 | 4.20E-06 |
| C4ZAV5 | ArectaleATCC33656 | PF10502.12 | Peptidase_S26 | 4.90E-34 |
| C4ZB65 | ArectaleATCC33656 | PF02674.19 | Colicin_V | 1.10E-22 |
| C4ZBE5 | ArectaleATCC33656 | PF00082.25 | Peptidase_S8 | 1.80E-40 |
| C4ZCV4 | ArectaleATCC33656 | PF00144.27 | Beta-lactamase | 4.60E-54 |
| C4ZCY0 | ArectaleATCC33656 | PF05580.15 | Peptidase_S55 | 1.40E-73 |
| C4ZD82 | ArectaleATCC33656 | PF02016.18 | Peptidase_S66 | 1.90E-21 |
| C4ZDN1 | ArectaleATCC33656 | PF01734.25 | Patatin | 5.70E-17 |
| C4ZDN2 | ArectaleATCC33656 | PF00768.23 | Peptidase_S11 | 3.20E-50 |
| C4ZE14 | ArectaleATCC33656 | PF00717.26 | Peptidase_S24 | 6.70E-27 |
| C4ZEI6 | ArectaleATCC33656 | PF03572.21 | Peptidase_S41 | 1.30E-49 |
| C4ZEW7 | ArectaleATCC33656 | PF12695.10 | Abhydrolase_5 | 1.10E-47 |
| C4ZF05 | ArectaleATCC33656 | PF00082.25 | Peptidase_S8 | 9.80E-19 |
| C4ZF28 | ArectaleATCC33656 | PF07859.16 | Abhydrolase_3 | 2.40E-48 |
| C4ZFJ7 | ArectaleATCC33656 | PF00717.26 | Peptidase_S24 | 2.50E-22 |
| C4ZFK7 | ArectaleATCC33656 | PF00768.23 | Peptidase_S11 | 1.80E-15 |
| C4ZG35 | ArectaleATCC33656 | PF16929.8 | Asp2 | 5.10E-213 |
| C4ZGF6 | ArectaleATCC33656 | PF05362.16 | Lon_C | 6.90E-90 |
| C4ZGJ1 | ArectaleATCC33656 | PF12146.11 | Hydrolase_4 | 9.80E-12 |
| C4ZGK4 | ArectaleATCC33656 | PF00756.23 | Esterase | 1.40E-16 |
| C4ZGK6 | ArectaleATCC33656 | PF13472.9 | Lipase_GDSL_2 | 5.60E-11 |
| C4ZGR9 | ArectaleATCC33656 | PF13365.9 | Trypsin_2 | 2.70E-30 |
| C4ZHC4 | ArectaleATCC33656 | PF00082.25 | Peptidase_S8 | 3.00E-18 |
| C4ZHP3 | ArectaleATCC33656 | PF12146.11 | Hydrolase_4 | 1.30E-50 |
| C4ZI06 | ArectaleATCC33656 | PF13365.9 | Trypsin_2 | 1.50E-10 |
| D9QZ71 | LsaccharolyticaATCC35040 | PF01734.25 | Patatin | 2.50E-12 |
| D9QZ95 | LsaccharolyticaATCC35040 | PF00574.26 | CLP_protease | 1.20E-31 |
| D9QZI2 | LsaccharolyticaATCC35040 | PF05362.16 | Lon_C | 1.20E-09 |
| D9QZR7 | LsaccharolyticaATCC35040 | PF00082.25 | Peptidase_S8 | 6.40E-49 |
| D9QZW7 | LsaccharolyticaATCC35040 | PF00082.25 | Peptidase_S8 | 1.00E-11 |
| D9R0H1 | LsaccharolyticaATCC35040 | PF10502.12 | Peptidase_S26 | 1.40E-43 |
| D9R0K6 | LsaccharolyticaATCC35040 | PF07859.16 | Abhydrolase_3 | 1.10E-41 |
| D9R0M7 | LsaccharolyticaATCC35040 | PF00698.24 | Acyl_transf_1 | 1.50E-36 |
| D9R0V8 | LsaccharolyticaATCC35040 | PF00082.25 | Peptidase_S8 | 1.00E-15 |
| D9R0Z1 | LsaccharolyticaATCC35040 | PF01694.25 | Rhomboid | 6.60E-37 |
| D9R140 | LsaccharolyticaATCC35040 | PF00717.26 | Peptidase_S24 | 1.20E-31 |
| D9R155 | LsaccharolyticaATCC35040 | PF01425.24 | Amidase | 8.30E-09 |
| D9R168 | LsaccharolyticaATCC35040 | PF00768.23 | Peptidase_S11 | 2.10E-50 |
| D9R1F8 | LsaccharolyticaATCC35040 | PF00144.27 | Beta-lactamase | 1.50E-60 |
| D9R1T1 | LsaccharolyticaATCC35040 | PF02674.19 | Colicin_V | 4.50E-11 |
| D9R233 | LsaccharolyticaATCC35040 | PF10502.12 | Peptidase_S26 | 7.30E-34 |
| D9R241 | LsaccharolyticaATCC35040 | PF12146.11 | Hydrolase_4 | 7.70E-15 |
| D9R261 | LsaccharolyticaATCC35040 | PF00574.26 | CLP_protease | 1.20E-21 |
| D9R2C2 | LsaccharolyticaATCC35040 | PF00082.25 | Peptidase_S8 | 3.40E-18 |
| D9R2P9 | LsaccharolyticaATCC35040 | PF12146.11 | Hydrolase_4 | 5.40E-59 |
| D9R2X4 | LsaccharolyticaATCC35040 | PF07859.16 | Abhydrolase_3 | 2.80E-20 |

| | | | | |
|---|---|---|---|---|
| D9R306 | LsaccharolyticaATCC35040 | PF13365.9 | Trypsin_2 | 6.80E-08 |
| D9R385 | LsaccharolyticaATCC35040 | PF00082.25 | Peptidase_S8 | 1.60E-13 |
| D9R386 | LsaccharolyticaATCC35040 | PF00082.25 | Peptidase_S8 | 3.60E-25 |
| D9R3K8 | LsaccharolyticaATCC35040 | PF00768.23 | Peptidase_S11 | 9.20E-56 |
| D9R3R7 | LsaccharolyticaATCC35040 | PF12697.10 | Abhydrolase_6 | 1.40E-07 |
| D9R429 | LsaccharolyticaATCC35040 | PF00574.26 | CLP_protease | 2.70E-73 |
| D9R466 | LsaccharolyticaATCC35040 | PF05362.16 | Lon_C | 5.80E-88 |
| D9R468 | LsaccharolyticaATCC35040 | PF00574.26 | CLP_protease | 6.40E-87 |
| D9R4C8 | LsaccharolyticaATCC35040 | PF00768.23 | Peptidase_S11 | 4.30E-71 |
| D9R4M4 | LsaccharolyticaATCC35040 | PF13365.9 | Trypsin_2 | 8.90E-29 |
| D9R4S7 | LsaccharolyticaATCC35040 | PF13354.9 | Beta-lactamase2 | 6.30E-56 |
| D9R4T6 | LsaccharolyticaATCC35040 | PF13354.9 | Beta-lactamase2 | 2.50E-52 |
| D9R4V6 | LsaccharolyticaATCC35040 | PF01734.25 | Patatin | 4.80E-20 |
| D9R4X5 | LsaccharolyticaATCC35040 | PF00768.23 | Peptidase_S11 | 2.20E-53 |
| D9R519 | LsaccharolyticaATCC35040 | PF05362.16 | Lon_C | 3.40E-82 |
| D9R595 | LsaccharolyticaATCC35040 | PF00082.25 | Peptidase_S8 | 1.90E-48 |
| D9R599 | LsaccharolyticaATCC35040 | PF00768.23 | Peptidase_S11 | 1.40E-50 |
| D9R5B9 | LsaccharolyticaATCC35040 | PF00082.25 | Peptidase_S8 | 5.80E-13 |
| D9R5G2 | LsaccharolyticaATCC35040 | PF00574.26 | CLP_protease | 3.30E-25 |
| D9R5R7 | LsaccharolyticaATCC35040 | PF12146.11 | Hydrolase_4 | 4.40E-14 |
| D9R5Y7 | LsaccharolyticaATCC35040 | PF03572.21 | Peptidase_S41 | 1.70E-47 |
| D9R6C0 | LsaccharolyticaATCC35040 | PF05362.16 | Lon_C | 2.30E-07 |
| D9R7K8 | LsaccharolyticaATCC35040 | PF02129.21 | Peptidase_S15 | 4.00E-09 |
| D9R871 | LsaccharolyticaATCC35040 | PF00144.27 | Beta-lactamase | 8.20E-44 |
| D9R8M8 | LsaccharolyticaATCC35040 | PF05580.15 | Peptidase_S55 | 1.40E-83 |
| D9R8R1 | LsaccharolyticaATCC35040 | PF07859.16 | Abhydrolase_3 | 2.80E-47 |
| D9R8U9 | LsaccharolyticaATCC35040 | PF00768.23 | Peptidase_S11 | 3.90E-47 |
| D9R951 | LsaccharolyticaATCC35040 | PF00082.25 | Peptidase_S8 | 5.90E-19 |
| D9R954 | LsaccharolyticaATCC35040 | PF00082.25 | Peptidase_S8 | 7.40E-25 |
| D9R956 | LsaccharolyticaATCC35040 | PF00082.25 | Peptidase_S8 | 1.70E-14 |
| D9R969 | LsaccharolyticaATCC35040 | PF01734.25 | Patatin | 2.30E-23 |
| D9R9D0 | LsaccharolyticaATCC35040 | PF01425.24 | Amidase | 7.70E-158 |
| D9R9L3 | LsaccharolyticaATCC35040 | PF13472.9 | Lipase_GDSL_2 | 1.20E-19 |
| D9R9U3 | LsaccharolyticaATCC35040 | PF00082.25 | Peptidase_S8 | 3.10E-13 |
| F8DL55 | LreuteriATCC55730 | PF01694.25 | Rhomboid | 9.60E-32 |
| F8DM65 | LreuteriATCC55730 | PF00574.26 | CLP_protease | 1.50E-81 |
| F8DM73 | LreuteriATCC55730 | PF12146.11 | Hydrolase_4 | 1.20E-15 |
| F8DMC2 | LreuteriATCC55730 | PF00144.27 | Beta-lactamase | 3.40E-44 |
| F8DMJ6 | LreuteriATCC55730 | PF00574.26 | CLP_protease | 5.60E-35 |
| F8DMX2 | LreuteriATCC55730 | PF00574.26 | CLP_protease | 4.70E-33 |
| F8DMY6 | LreuteriATCC55730 | PF01425.24 | Amidase | 5.60E-149 |
| F8DN38 | LreuteriATCC55730 | PF02674.19 | Colicin_V | 2.10E-31 |
| F8DND4 | LreuteriATCC55730 | PF07859.16 | Abhydrolase_3 | 1.20E-29 |
| F8DND9 | LreuteriATCC55730 | PF12697.10 | Abhydrolase_6 | 8.30E-15 |
| F8DNN8 | LreuteriATCC55730 | PF00768.23 | Peptidase_S11 | 7.80E-48 |
| F8DNN9 | LreuteriATCC55730 | PF13354.9 | Beta-lactamase2 | 1.80E-13 |
| F8DNW1 | LreuteriATCC55730 | PF05362.16 | Lon_C | 9.40E-07 |
| F8DP03 | LreuteriATCC55730 | PF00717.26 | Peptidase_S24 | 1.80E-29 |
| F8DPH9 | LreuteriATCC55730 | PF13472.9 | Lipase_GDSL_2 | 5.70E-25 |
| F8DPJ4 | LreuteriATCC55730 | PF02129.21 | Peptidase_S15 | 6.90E-77 |
| F8DPL7 | LreuteriATCC55730 | PF12146.11 | Hydrolase_4 | 1.20E-11 |
| F8DPN6 | LreuteriATCC55730 | PF12146.11 | Hydrolase_4 | 6.90E-16 |

| | | | | |
|---|---|---|---|---|
| F8DPY1 | LreuteriATCC55730 | PF04586.20 | Peptidase_S78 | 9.60E-46 |
| F8DQ72 | LreuteriATCC55730 | PF13472.9 | Lipase_GDSL_2 | 1.10E-10 |
| F8DQN7 | LreuteriATCC55730 | PF00768.23 | Peptidase_S11 | 4.40E-54 |
| F8DR11 | LreuteriATCC55730 | PF00144.27 | Beta-lactamase | 8.10E-36 |
| F8DR29 | LreuteriATCC55730 | PF10502.12 | Peptidase_S26 | 4.00E-43 |
| F8DR64 | LreuteriATCC55730 | PF13365.9 | Trypsin_2 | 4.30E-30 |
| F8DRB2 | LreuteriATCC55730 | PF00698.24 | Acyl_transf_1 | 3.90E-28 |
| F8DRC2 | LreuteriATCC55730 | PF01734.25 | Patatin | 1.40E-10 |
| F8DRZ7 | LreuteriATCC55730 | PF06821.16 | Ser_hydrolase | 7.60E-23 |
| A6L2J8 | PvulgatusATCC8482 | PF10459.12 | Peptidase_S46 | 1.70E-270 |
| A6KWF2 | PvulgatusATCC8482 | PF03629.21 | SASA | 2.50E-13 |
| A6KWF6 | PvulgatusATCC8482 | PF00756.23 | Esterase | 4.70E-20 |
| A6KWI0 | PvulgatusATCC8482 | PF10502.12 | Peptidase_S26 | 1.10E-33 |
| A6KWN5 | PvulgatusATCC8482 | PF00144.27 | Beta-lactamase | 3.00E-42 |
| A6KWQ9 | PvulgatusATCC8482 | PF00144.27 | Beta-lactamase | 6.50E-45 |
| A6KWS4 | PvulgatusATCC8482 | PF13472.9 | Lipase_GDSL_2 | 9.20E-12 |
| A6KWU6 | PvulgatusATCC8482 | PF13472.9 | Lipase_GDSL_2 | 2.60E-13 |
| A6KWX4 | PvulgatusATCC8482 | PF03572.21 | Peptidase_S41 | 5.80E-33 |
| A6KWZ2 | PvulgatusATCC8482 | PF13472.9 | Lipase_GDSL_2 | 2.10E-18 |
| A6KX28 | PvulgatusATCC8482 | PF00082.25 | Peptidase_S8 | 4.60E-37 |
| A6KX77 | PvulgatusATCC8482 | PF10502.12 | Peptidase_S26 | 2.20E-11 |
| A6KXJ5 | PvulgatusATCC8482 | PF00574.26 | CLP_protease | 1.30E-76 |
| A6KXT7 | PvulgatusATCC8482 | PF13472.9 | Lipase_GDSL_2 | 1.20E-26 |
| A6KYC3 | PvulgatusATCC8482 | PF00326.24 | Peptidase_S9 | 1.00E-32 |
| A6KYD0 | PvulgatusATCC8482 | PF03629.21 | SASA | 7.80E-12 |
| A6KYM0 | PvulgatusATCC8482 | PF03572.21 | Peptidase_S41 | 1.50E-45 |
| A6KZ23 | PvulgatusATCC8482 | PF01694.25 | Rhomboid | 1.80E-25 |
| A6KZ24 | PvulgatusATCC8482 | PF01694.25 | Rhomboid | 3.90E-35 |
| A6KZC1 | PvulgatusATCC8482 | PF00326.24 | Peptidase_S9 | 7.60E-46 |
| A6KZD5 | PvulgatusATCC8482 | PF02113.18 | Peptidase_S13 | 4.30E-92 |
| A6KZE8 | PvulgatusATCC8482 | PF00326.24 | Peptidase_S9 | 6.00E-45 |
| A6KZI1 | PvulgatusATCC8482 | PF03572.21 | Peptidase_S41 | 1.20E-19 |
| A6KZL9 | PvulgatusATCC8482 | PF03572.21 | Peptidase_S41 | 6.70E-17 |
| A6KZR3 | PvulgatusATCC8482 | PF03572.21 | Peptidase_S41 | 5.30E-51 |
| A6KZT4 | PvulgatusATCC8482 | PF10502.12 | Peptidase_S26 | 9.00E-23 |
| A6KZT5 | PvulgatusATCC8482 | PF10502.12 | Peptidase_S26 | 5.90E-26 |
| A6KZW8 | PvulgatusATCC8482 | PF10459.12 | Peptidase_S46 | 6.70E-267 |
| A6L018 | PvulgatusATCC8482 | PF05448.15 | AXE1 | 3.50E-49 |
| A6L031 | PvulgatusATCC8482 | PF02674.19 | Colicin_V | 3.10E-26 |
| A6L0D9 | PvulgatusATCC8482 | PF05576.14 | Peptidase_S37 | 4.30E-50 |
| A6L0H6 | PvulgatusATCC8482 | PF00082.25 | Peptidase_S8 | 5.20E-33 |
| A6L0S6 | PvulgatusATCC8482 | PF01343.21 | Peptidase_S49 | 1.80E-43 |
| A6L1C3 | PvulgatusATCC8482 | PF05448.15 | AXE1 | 7.40E-11 |
| A6L1C7 | PvulgatusATCC8482 | PF13472.9 | Lipase_GDSL_2 | 1.90E-13 |
| A6L1D0 | PvulgatusATCC8482 | PF00326.24 | Peptidase_S9 | 2.30E-36 |
| A6L1G9 | PvulgatusATCC8482 | PF02129.21 | Peptidase_S15 | 5.10E-51 |
| A6L1I6 | PvulgatusATCC8482 | PF03572.21 | Peptidase_S41 | 3.10E-20 |
| A6L1U9 | PvulgatusATCC8482 | PF00326.24 | Peptidase_S9 | 4.50E-54 |
| A6L1X5 | PvulgatusATCC8482 | PF13472.9 | Lipase_GDSL_2 | 3.20E-18 |
| A6L209 | PvulgatusATCC8482 | PF12146.11 | Hydrolase_4 | 2.90E-15 |
| A6L225 | PvulgatusATCC8482 | PF01734.25 | Patatin | 4.20E-20 |
| A6L245 | PvulgatusATCC8482 | PF13365.9 | Trypsin_2 | 1.70E-10 |

| | | | | |
|---|---|---|---|---|
| A6L2A7 | PvulgatusATCC8482 | PF13472.9 | Lipase_GDSL_2 | 1.40E-15 |
| A6L2J7 | PvulgatusATCC8482 | PF10459.12 | Peptidase_S46 | 5.20E-270 |
| A6L2K8 | PvulgatusATCC8482 | PF03572.21 | Peptidase_S41 | 2.00E-20 |
| A6L2S7 | PvulgatusATCC8482 | PF03629.21 | SASA | 1.70E-20 |
| A6L2S7 | PvulgatusATCC8482 | PF13472.9 | Lipase_GDSL_2 | 2.30E-13 |
| A6L378 | PvulgatusATCC8482 | PF00756.23 | Esterase | 1.60E-30 |
| A6L3D3 | PvulgatusATCC8482 | PF00326.24 | Peptidase_S9 | 2.00E-12 |
| A6L3L3 | PvulgatusATCC8482 | PF03572.21 | Peptidase_S41 | 4.90E-52 |
| A6L3T5 | PvulgatusATCC8482 | PF03629.21 | SASA | 5.50E-17 |
| A6L3X3 | PvulgatusATCC8482 | PF13365.9 | Trypsin_2 | 1.80E-34 |
| A6L409 | PvulgatusATCC8482 | PF01694.25 | Rhomboid | 1.00E-14 |
| A6L4P2 | PvulgatusATCC8482 | PF00326.24 | Peptidase_S9 | 6.50E-25 |
| A6L4Q9 | PvulgatusATCC8482 | PF01734.25 | Patatin | 4.60E-27 |
| A6L531 | PvulgatusATCC8482 | PF05362.16 | Lon_C | 3.80E-82 |
| A6L5C2 | PvulgatusATCC8482 | PF13354.9 | Beta-lactamase2 | 4.40E-33 |
| A6L5G6 | PvulgatusATCC8482 | PF01734.25 | Patatin | 3.10E-27 |
| A6L5L7 | PvulgatusATCC8482 | PF00326.24 | Peptidase_S9 | 1.50E-52 |
| A6L5T5 | PvulgatusATCC8482 | PF00717.26 | Peptidase_S24 | 5.70E-25 |
| A6L6M1 | PvulgatusATCC8482 | PF03575.20 | Peptidase_S51 | 3.80E-48 |
| A6L6Y2 | PvulgatusATCC8482 | PF03629.21 | SASA | 1.20E-12 |
| A6L715 | PvulgatusATCC8482 | PF00326.24 | Peptidase_S9 | 1.00E-40 |
| A6L7C8 | PvulgatusATCC8482 | PF00698.24 | Acyl_transf_1 | 1.20E-38 |
| A6L7C9 | PvulgatusATCC8482 | PF01734.25 | Patatin | 1.80E-23 |
| A6L7K9 | PvulgatusATCC8482 | PF03572.21 | Peptidase_S41 | 1.20E-16 |
| A6L7M1 | PvulgatusATCC8482 | PF00326.24 | Peptidase_S9 | 2.00E-42 |
| A6L7S8 | PvulgatusATCC8482 | PF03629.21 | SASA | 5.40E-20 |
| A6L7S8 | PvulgatusATCC8482 | PF13472.9 | Lipase_GDSL_2 | 2.10E-24 |
| A6L7S9 | PvulgatusATCC8482 | PF13472.9 | Lipase_GDSL_2 | 1.50E-19 |
| A6L7W1 | PvulgatusATCC8482 | PF00756.23 | Esterase | 1.00E-18 |
| A6L7W5 | PvulgatusATCC8482 | PF02129.21 | Peptidase_S15 | 1.50E-10 |
| D4BL85 | BbreveDSM20213 | PF02230.19 | Abhydrolase_2 | 5.00E-17 |
| D4BLK3 | BbreveDSM20213 | PF01694.25 | Rhomboid | 1.10E-26 |
| D4BLL6 | BbreveDSM20213 | PF00326.24 | Peptidase_S9 | 1.50E-25 |
| D4BLL8 | BbreveDSM20213 | PF12146.11 | Hydrolase_4 | 3.60E-38 |
| D4BLM2 | BbreveDSM20213 | PF00756.23 | Esterase | 1.70E-08 |
| D4BLN3 | BbreveDSM20213 | PF13365.9 | Trypsin_2 | 3.40E-31 |
| D4BLV6 | BbreveDSM20213 | PF12146.11 | Hydrolase_4 | 7.20E-16 |
| D4BM22 | BbreveDSM20213 | PF13354.9 | Beta-lactamase2 | 2.70E-29 |
| D4BM79 | BbreveDSM20213 | PF01425.24 | Amidase | 6.30E-150 |
| D4BMK9 | BbreveDSM20213 | PF10502.12 | Peptidase_S26 | 8.90E-39 |
| D4BMS7 | BbreveDSM20213 | PF12697.10 | Abhydrolase_6 | 9.00E-17 |
| D4BMT1 | BbreveDSM20213 | PF00326.24 | Peptidase_S9 | 5.70E-53 |
| D4BN64 | BbreveDSM20213 | PF07859.16 | Abhydrolase_3 | 2.40E-67 |
| D4BNC5 | BbreveDSM20213 | PF02230.19 | Abhydrolase_2 | 2.00E-20 |
| D4BND4 | BbreveDSM20213 | PF01734.25 | Patatin | 3.50E-14 |
| D4BNE3 | BbreveDSM20213 | PF05362.16 | Lon_C | 2.90E-12 |
| D4BNW9 | BbreveDSM20213 | PF00574.26 | CLP_protease | 1.70E-70 |
| D4BNX0 | BbreveDSM20213 | PF00574.26 | CLP_protease | 9.20E-75 |
| D4BP52 | BbreveDSM20213 | PF03575.20 | Peptidase_S51 | 1.50E-44 |
| D4BP56 | BbreveDSM20213 | PF12697.10 | Abhydrolase_6 | 3.70E-18 |
| D4BPE2 | BbreveDSM20213 | PF02230.19 | Abhydrolase_2 | 3.50E-12 |
| D4BQH0 | BbreveDSM20213 | PF00698.24 | Acyl_transf_1 | 2.20E-35 |

| | | | | |
|---|---|---|---|---|
| D4BQV0 | BbreveDSM20213 | PF05362.16 | Lon_C | 1.00E-05 |
| D4BR21 | BbreveDSM20213 | PF02113.18 | Peptidase_S13 | 2.00E-38 |
| D4BRQ8 | BbreveDSM20213 | PF00717.26 | Peptidase_S24 | 6.00E-31 |
| D4BRW8 | BbreveDSM20213 | PF04586.20 | Peptidase_S78 | 1.10E-27 |
| H1PPQ5 | Fulcerans121B | PF02016.18 | Peptidase_S66 | 3.90E-38 |
| H1PPX6 | Fulcerans121B | PF01734.25 | Patatin | 1.40E-14 |
| H1PQ10 | Fulcerans121B | PF01343.21 | Peptidase_S49 | 2.30E-39 |
| H1PQ57 | Fulcerans121B | PF00574.26 | CLP_protease | 2.00E-87 |
| H1PQ59 | Fulcerans121B | PF05362.16 | Lon_C | 1.00E-80 |
| H1PQK5 | Fulcerans121B | PF00717.26 | Peptidase_S24 | 2.70E-21 |
| H1PQN8 | Fulcerans121B | PF01734.25 | Patatin | 6.80E-25 |
| H1PQX4 | Fulcerans121B | PF00326.24 | Peptidase_S9 | 3.20E-56 |
| H1PR71 | Fulcerans121B | PF02674.19 | Colicin_V | 1.10E-20 |
| H1PRC3 | Fulcerans121B | PF12146.11 | Hydrolase_4 | 5.00E-54 |
| H1PRC6 | Fulcerans121B | PF01734.25 | Patatin | 1.80E-13 |
| H1PRG6 | Fulcerans121B | PF02016.18 | Peptidase_S66 | 2.10E-30 |
| H1PRI9 | Fulcerans121B | PF13354.9 | Beta-lactamase2 | 6.80E-46 |
| H1PS09 | Fulcerans121B | PF03572.21 | Peptidase_S41 | 5.70E-57 |
| H1PSI3 | Fulcerans121B | PF13472.9 | Lipase_GDSL_2 | 4.80E-10 |
| H1PTI9 | Fulcerans121B | PF00756.23 | Esterase | 1.80E-22 |
| H1PTR8 | Fulcerans121B | PF00574.26 | CLP_protease | 3.40E-28 |
| H1PTU9 | Fulcerans121B | PF00717.26 | Peptidase_S24 | 4.60E-26 |
| H1PUP4 | Fulcerans121B | PF01425.24 | Amidase | 1.20E-149 |
| H1PVI6 | Fulcerans121B | PF05362.16 | Lon_C | 4.70E-10 |
| H1PVJ3 | Fulcerans121B | PF00698.24 | Acyl_transf_1 | 6.90E-43 |
| H1PW52 | Fulcerans121B | PF00717.26 | Peptidase_S24 | 1.10E-27 |
| H1PWD3 | Fulcerans121B | PF00574.26 | CLP_protease | 3.70E-28 |
| H1PWE8 | Fulcerans121B | PF00717.26 | Peptidase_S24 | 6.30E-21 |
| H1PWE9 | Fulcerans121B | PF00717.26 | Peptidase_S24 | 4.20E-25 |
| H1PWH4 | Fulcerans121B | PF12146.11 | Hydrolase_4 | 1.10E-42 |
| H1PWJ2 | Fulcerans121B | PF00082.25 | Peptidase_S8 | 1.70E-30 |
| H1PWZ2 | Fulcerans121B | PF01734.25 | Patatin | 1.60E-23 |
| H1PX23 | Fulcerans121B | PF05448.15 | AXE1 | 1.00E-111 |
| H1PXA3 | Fulcerans121B | PF00756.23 | Esterase | 3.30E-14 |
| H1PXA4 | Fulcerans121B | PF00756.23 | Esterase | 1.30E-09 |
| H1PXU2 | Fulcerans121B | PF00768.23 | Peptidase_S11 | 5.40E-48 |
| H1PY48 | Fulcerans121B | PF10502.12 | Peptidase_S26 | 6.20E-39 |
| H1PY93 | Fulcerans121B | PF01734.25 | Patatin | 1.60E-24 |
| H1PY94 | Fulcerans121B | PF02113.18 | Peptidase_S13 | 8.80E-74 |
| H1PYB3 | Fulcerans121B | PF00717.26 | Peptidase_S24 | 6.20E-20 |
| H1PYJ9 | Fulcerans121B | PF10502.12 | Peptidase_S26 | 3.50E-23 |
| S2L1S0 | Fulcerans121B | PF00082.25 | Peptidase_S8 | 9.30E-29 |
| S2LQB3 | Fulcerans121B | PF00574.26 | CLP_protease | 6.20E-33 |
| D4BSR6 | PrettgeriDSM1131 | PF00698.24 | Acyl_transf_1 | 2.40E-36 |
| D4BT50 | PrettgeriDSM1131 | PF00326.24 | Peptidase_S9 | 6.80E-64 |
| D4BTE0 | PrettgeriDSM1131 | PF03572.21 | Peptidase_S41 | 4.30E-51 |
| D4BTG7 | PrettgeriDSM1131 | PF01343.21 | Peptidase_S49 | 1.10E-46 |
| D4BTL4 | PrettgeriDSM1131 | PF01343.21 | Peptidase_S49 | 2.20E-44 |
| D4BTV3 | PrettgeriDSM1131 | PF12146.11 | Hydrolase_4 | 1.20E-06 |
| D4BU14 | PrettgeriDSM1131 | PF05929.14 | Phage_GPO | 2.30E-62 |
| D4BUD9 | PrettgeriDSM1131 | PF12146.11 | Hydrolase_4 | 4.70E-63 |
| D4BUI1 | PrettgeriDSM1131 | PF00717.26 | Peptidase_S24 | 1.30E-18 |

| | | | | |
|---|---|---|---|---|
| D4BUY2 | PrettgeriDSM1131 | PF07859.16 | Abhydrolase_3 | 2.00E-46 |
| D4BVF7 | PrettgeriDSM1131 | PF05362.16 | Lon_C | 2.00E-09 |
| D4BVK3 | PrettgeriDSM1131 | PF12697.10 | Abhydrolase_6 | 7.10E-16 |
| D4BVS1 | PrettgeriDSM1131 | PF00768.23 | Peptidase_S11 | 5.40E-101 |
| D4BVW2 | PrettgeriDSM1131 | PF01694.25 | Rhomboid | 1.20E-24 |
| D4BW05 | PrettgeriDSM1131 | PF00768.23 | Peptidase_S11 | 1.60E-103 |
| D4BWC5 | PrettgeriDSM1131 | PF01764.28 | Lipase_3 | 2.90E-21 |
| D4BWC7 | PrettgeriDSM1131 | PF01764.28 | Lipase_3 | 6.70E-23 |
| D4BWI8 | PrettgeriDSM1131 | PF02674.19 | Colicin_V | 4.90E-38 |
| D4BWZ1 | PrettgeriDSM1131 | PF00082.25 | Peptidase_S8 | 3.20E-17 |
| D4BXA6 | PrettgeriDSM1131 | PF13365.9 | Trypsin_2 | 7.40E-30 |
| D4BXA7 | PrettgeriDSM1131 | PF13365.9 | Trypsin_2 | 2.20E-30 |
| D4BXD2 | PrettgeriDSM1131 | PF06821.16 | Ser_hydrolase | 3.90E-27 |
| D4BYL2 | PrettgeriDSM1131 | PF12697.10 | Abhydrolase_6 | 2.80E-16 |
| D4BYU8 | PrettgeriDSM1131 | PF13472.9 | Lipase_GDSL_2 | 7.90E-22 |
| D4BZB8 | PrettgeriDSM1131 | PF03575.20 | Peptidase_S51 | 9.10E-45 |
| D4BZP5 | PrettgeriDSM1131 | PF13472.9 | Lipase_GDSL_2 | 2.60E-08 |
| D4BZP6 | PrettgeriDSM1131 | PF13472.9 | Lipase_GDSL_2 | 2.00E-13 |
| D4BZT1 | PrettgeriDSM1131 | PF00717.26 | Peptidase_S24 | 4.00E-33 |
| D4C056 | PrettgeriDSM1131 | PF03575.20 | Peptidase_S51 | 3.80E-60 |
| D4C067 | PrettgeriDSM1131 | PF00756.23 | Esterase | 2.90E-60 |
| D4C0H3 | PrettgeriDSM1131 | PF12697.10 | Abhydrolase_6 | 1.30E-07 |
| D4C198 | PrettgeriDSM1131 | PF05362.16 | Lon_C | 4.20E-97 |
| D4C1A0 | PrettgeriDSM1131 | PF00574.26 | CLP_protease | 2.00E-89 |
| D4C1P4 | PrettgeriDSM1131 | PF01734.25 | Patatin | 1.80E-20 |
| D4C1R1 | PrettgeriDSM1131 | PF00756.23 | Esterase | 8.90E-21 |
| D4C1X4 | PrettgeriDSM1131 | PF00144.27 | Beta-lactamase | 2.10E-44 |
| D4C2D7 | PrettgeriDSM1131 | PF00144.27 | Beta-lactamase | 3.60E-75 |
| D4C2F8 | PrettgeriDSM1131 | PF12697.10 | Abhydrolase_6 | 7.20E-19 |
| D4C2G3 | PrettgeriDSM1131 | PF01694.25 | Rhomboid | 1.80E-26 |
| D4C2J6 | PrettgeriDSM1131 | PF00756.23 | Esterase | 2.20E-18 |
| D4C399 | PrettgeriDSM1131 | PF02129.21 | Peptidase_S15 | 3.80E-56 |
| D4C3G9 | PrettgeriDSM1131 | PF00082.25 | Peptidase_S8 | 1.30E-17 |
| D4C3K4 | PrettgeriDSM1131 | PF00144.27 | Beta-lactamase | 4.30E-56 |
| D4C3R6 | PrettgeriDSM1131 | PF02113.18 | Peptidase_S13 | 1.20E-153 |
| D4C3Y0 | PrettgeriDSM1131 | PF12146.11 | Hydrolase_4 | 4.20E-46 |
| D4C4L5 | PrettgeriDSM1131 | PF06500.14 | FrsA-like | 1.30E-162 |
| D4C514 | PrettgeriDSM1131 | PF10502.12 | Peptidase_S26 | 2.60E-56 |
| D4C542 | PrettgeriDSM1131 | PF12697.10 | Abhydrolase_6 | 7.60E-08 |
| D4C5T2 | PrettgeriDSM1131 | PF01694.25 | Rhomboid | 5.80E-16 |
| D4C5T3 | PrettgeriDSM1131 | PF01804.21 | Penicil_amidase | 1.70E-138 |
| A5ZB40 | BcaccaeATCC43185 | PF01694.25 | Rhomboid | 2.30E-24 |
| A5ZB41 | BcaccaeATCC43185 | PF01694.25 | Rhomboid | 8.50E-32 |
| A5ZB64 | BcaccaeATCC43185 | PF03572.21 | Peptidase_S41 | 5.30E-49 |
| A5ZBL2 | BcaccaeATCC43185 | PF13472.9 | Lipase_GDSL_2 | 9.50E-13 |
| A5ZBR6 | BcaccaeATCC43185 | PF02016.18 | Peptidase_S66 | 6.70E-36 |
| A5ZBW2 | BcaccaeATCC43185 | PF00144.27 | Beta-lactamase | 3.50E-46 |
| A5ZC96 | BcaccaeATCC43185 | PF01734.25 | Patatin | 3.60E-22 |
| A5ZCG6 | BcaccaeATCC43185 | PF01734.25 | Patatin | 2.20E-28 |
| A5ZCL8 | BcaccaeATCC43185 | PF05362.16 | Lon_C | 5.40E-82 |
| A5ZCQ4 | BcaccaeATCC43185 | PF00698.24 | Acyl_transf_1 | 1.80E-36 |
| A5ZCR7 | BcaccaeATCC43185 | PF01734.25 | Patatin | 1.40E-21 |

| | | | | |
|---|---|---|---|---|
| A5ZCY2 | BcaccaeATCC43185 | PF04586.20 | Peptidase_S78 | 6.60E-05 |
| A5ZD81 | BcaccaeATCC43185 | PF10502.12 | Peptidase_S26 | 8.60E-31 |
| A5ZDH6 | BcaccaeATCC43185 | PF00326.24 | Peptidase_S9 | 2.60E-56 |
| A5ZDS2 | BcaccaeATCC43185 | PF13472.9 | Lipase_GDSL_2 | 1.40E-15 |
| A5ZDX7 | BcaccaeATCC43185 | PF03629.21 | SASA | 2.20E-11 |
| A5ZDX7 | BcaccaeATCC43185 | PF13472.9 | Lipase_GDSL_2 | 5.00E-25 |
| A5ZDX8 | BcaccaeATCC43185 | PF13472.9 | Lipase_GDSL_2 | 5.90E-20 |
| A5ZEB2 | BcaccaeATCC43185 | PF00717.26 | Peptidase_S24 | 1.10E-16 |
| A5ZEC4 | BcaccaeATCC43185 | PF13472.9 | Lipase_GDSL_2 | 1.40E-18 |
| A5ZEG7 | BcaccaeATCC43185 | PF03572.21 | Peptidase_S41 | 2.70E-51 |
| A5ZEK1 | BcaccaeATCC43185 | PF02129.21 | Peptidase_S15 | 1.00E-16 |
| A5ZET3 | BcaccaeATCC43185 | PF02113.18 | Peptidase_S13 | 2.90E-79 |
| A5ZEY9 | BcaccaeATCC43185 | PF00326.24 | Peptidase_S9 | 5.80E-44 |
| A5ZF11 | BcaccaeATCC43185 | PF00326.24 | Peptidase_S9 | 2.80E-37 |
| A5ZF12 | BcaccaeATCC43185 | PF00326.24 | Peptidase_S9 | 7.70E-31 |
| A5ZF13 | BcaccaeATCC43185 | PF00326.24 | Peptidase_S9 | 2.10E-22 |
| A5ZF21 | BcaccaeATCC43185 | PF10459.12 | Peptidase_S46 | 5.80E-265 |
| A5ZF37 | BcaccaeATCC43185 | PF10502.12 | Peptidase_S26 | 1.90E-09 |
| A5ZF38 | BcaccaeATCC43185 | PF10502.12 | Peptidase_S26 | 1.90E-24 |
| A5ZFT4 | BcaccaeATCC43185 | PF13472.9 | Lipase_GDSL_2 | 1.40E-12 |
| A5ZFX9 | BcaccaeATCC43185 | PF00326.24 | Peptidase_S9 | 2.40E-58 |
| A5ZFZ9 | BcaccaeATCC43185 | PF00326.24 | Peptidase_S9 | 1.80E-14 |
| A5ZG53 | BcaccaeATCC43185 | PF03572.21 | Peptidase_S41 | 6.00E-52 |
| A5ZG69 | BcaccaeATCC43185 | PF13472.9 | Lipase_GDSL_2 | 3.70E-15 |
| A5ZG72 | BcaccaeATCC43185 | PF01694.25 | Rhomboid | 1.70E-12 |
| A5ZGC2 | BcaccaeATCC43185 | PF00082.25 | Peptidase_S8 | 5.80E-25 |
| A5ZGG1 | BcaccaeATCC43185 | PF13472.9 | Lipase_GDSL_2 | 1.80E-11 |
| A5ZGG6 | BcaccaeATCC43185 | PF13354.9 | Beta-lactamase2 | 5.60E-31 |
| A5ZGX3 | BcaccaeATCC43185 | PF05448.15 | AXE1 | 2.30E-14 |
| A5ZGZ6 | BcaccaeATCC43185 | PF00756.23 | Esterase | 9.50E-29 |
| A5ZGZ7 | BcaccaeATCC43185 | PF03572.21 | Peptidase_S41 | 3.40E-13 |
| A5ZH64 | BcaccaeATCC43185 | PF10459.12 | Peptidase_S46 | 9.70E-262 |
| A5ZH65 | BcaccaeATCC43185 | PF10459.12 | Peptidase_S46 | 4.40E-257 |
| A5ZH99 | BcaccaeATCC43185 | PF01734.25 | Patatin | 1.50E-26 |
| A5ZHN2 | BcaccaeATCC43185 | PF03629.21 | SASA | 2.60E-21 |
| A5ZHV0 | BcaccaeATCC43185 | PF03572.21 | Peptidase_S41 | 1.80E-19 |
| A5ZI60 | BcaccaeATCC43185 | PF03572.21 | Peptidase_S41 | 4.80E-15 |
| A5ZI65 | BcaccaeATCC43185 | PF13365.9 | Trypsin_2 | 1.40E-34 |
| A5ZID0 | BcaccaeATCC43185 | PF10502.12 | Peptidase_S26 | 2.40E-35 |
| A5ZID9 | BcaccaeATCC43185 | PF13472.9 | Lipase_GDSL_2 | 2.00E-12 |
| A5ZIH6 | BcaccaeATCC43185 | PF01694.25 | Rhomboid | 3.00E-30 |
| A5ZIJ5 | BcaccaeATCC43185 | PF01734.25 | Patatin | 2.60E-21 |
| A5ZKD4 | BcaccaeATCC43185 | PF00574.26 | CLP_protease | 1.00E-77 |
| A5ZKH2 | BcaccaeATCC43185 | PF00082.25 | Peptidase_S8 | 4.20E-37 |
| A5ZKL6 | BcaccaeATCC43185 | PF02674.19 | Colicin_V | 2.50E-31 |
| A5ZKW2 | BcaccaeATCC43185 | PF03572.21 | Peptidase_S41 | 3.30E-32 |
| A5ZLG4 | BcaccaeATCC43185 | PF01343.21 | Peptidase_S49 | 2.00E-48 |
| A5ZLN4 | BcaccaeATCC43185 | PF00326.24 | Peptidase_S9 | 4.50E-48 |
| B0N9E4 | CscindensATCC35704 | PF13365.9 | Trypsin_2 | 1.50E-27 |
| B0N9U3 | CscindensATCC35704 | PF00574.26 | CLP_protease | 4.40E-30 |
| B0N9Y2 | CscindensATCC35704 | PF00144.27 | Beta-lactamase | 1.70E-18 |
| B0NA73 | CscindensATCC35704 | PF00082.25 | Peptidase_S8 | 6.20E-17 |

| | | | | |
|---|---|---|---|---|
| B0NAG5 | CscindensATCC35704 | PF05580.15 | Peptidase_S55 | 1.60E-84 |
| B0NAM2 | CscindensATCC35704 | PF00717.26 | Peptidase_S24 | 4.00E-31 |
| B0NAP1 | CscindensATCC35704 | PF00768.23 | Peptidase_S11 | 1.50E-45 |
| B0NAT0 | CscindensATCC35704 | PF00574.26 | CLP_protease | 2.80E-28 |
| B0NAX6 | CscindensATCC35704 | PF02016.18 | Peptidase_S66 | 2.80E-28 |
| B0NAY6 | CscindensATCC35704 | PF00574.26 | CLP_protease | 1.40E-25 |
| B0NBK8 | CscindensATCC35704 | PF01734.25 | Patatin | 2.80E-20 |
| B0NC41 | CscindensATCC35704 | PF10502.12 | Peptidase_S26 | 4.30E-16 |
| B0NCW7 | CscindensATCC35704 | PF01734.25 | Patatin | 9.30E-07 |
| B0NCZ1 | CscindensATCC35704 | PF00082.25 | Peptidase_S8 | 2.50E-49 |
| B0NCZ9 | CscindensATCC35704 | PF10502.12 | Peptidase_S26 | 4.50E-37 |
| B0ND01 | CscindensATCC35704 | PF10502.12 | Peptidase_S26 | 1.30E-39 |
| B0ND57 | CscindensATCC35704 | PF00768.23 | Peptidase_S11 | 1.40E-74 |
| B0ND61 | CscindensATCC35704 | PF00768.23 | Peptidase_S11 | 9.30E-47 |
| B0NDA4 | CscindensATCC35704 | PF03629.21 | SASA | 2.00E-13 |
| B0NDG2 | CscindensATCC35704 | PF00574.26 | CLP_protease | 6.40E-24 |
| B0NDS8 | CscindensATCC35704 | PF10502.12 | Peptidase_S26 | 4.40E-23 |
| B0NEN7 | CscindensATCC35704 | PF12697.10 | Abhydrolase_6 | 8.10E-10 |
| B0NEX4 | CscindensATCC35704 | PF02674.19 | Colicin_V | 9.30E-09 |
| B0NFQ9 | CscindensATCC35704 | PF00698.24 | Acyl_transf_1 | 2.90E-40 |
| B0NGF3 | CscindensATCC35704 | PF01694.25 | Rhomboid | 7.90E-36 |
| B0NGQ9 | CscindensATCC35704 | PF10502.12 | Peptidase_S26 | 7.20E-30 |
| B0NGR0 | CscindensATCC35704 | PF00768.23 | Peptidase_S11 | 3.10E-42 |
| B0NHB7 | CscindensATCC35704 | PF07859.16 | Abhydrolase_3 | 4.20E-69 |
| B0NHR7 | CscindensATCC35704 | PF12146.11 | Hydrolase_4 | 7.10E-47 |
| B0NHT9 | CscindensATCC35704 | PF00768.23 | Peptidase_S11 | 2.10E-47 |
| B0NI42 | CscindensATCC35704 | PF12697.10 | Abhydrolase_6 | 1.10E-14 |
| B0NI49 | CscindensATCC35704 | PF01425.24 | Amidase | 1.70E-49 |
| B0NI56 | CscindensATCC35704 | PF13365.9 | Trypsin_2 | 9.00E-08 |
| B0NIQ3 | CscindensATCC35704 | PF12697.10 | Abhydrolase_6 | 4.40E-10 |
| B0NJ85 | CscindensATCC35704 | PF01343.21 | Peptidase_S49 | 1.50E-44 |
| B0NJH0 | CscindensATCC35704 | PF00144.27 | Beta-lactamase | 2.10E-06 |
| B0NJL6 | CscindensATCC35704 | PF00574.26 | CLP_protease | 7.30E-86 |
| B0NJL8 | CscindensATCC35704 | PF05362.16 | Lon_C | 3.60E-90 |
| B0NJQ7 | CscindensATCC35704 | PF00082.25 | Peptidase_S8 | 1.60E-09 |
| B0NJR5 | CscindensATCC35704 | PF13472.9 | Lipase_GDSL_2 | 1.10E-16 |
| B0NJT4 | CscindensATCC35704 | PF01425.24 | Amidase | 7.50E-160 |
| B0NKA4 | CscindensATCC35704 | PF12146.11 | Hydrolase_4 | 7.60E-12 |
| B0NKB4 | CscindensATCC35704 | PF12146.11 | Hydrolase_4 | 1.10E-57 |
| B0NKL2 | CscindensATCC35704 | PF03572.21 | Peptidase_S41 | 2.80E-55 |
| D2Z9B7 | EcancerogenusATCC35316 | PF13365.9 | Trypsin_2 | 9.90E-32 |
| D2Z9W8 | EcancerogenusATCC35316 | PF01734.25 | Patatin | 5.30E-19 |
| D2ZA34 | EcancerogenusATCC35316 | PF00574.26 | CLP_protease | 1.90E-64 |
| D2ZA93 | EcancerogenusATCC35316 | PF00768.23 | Peptidase_S11 | 3.40E-103 |
| D2ZAP5 | EcancerogenusATCC35316 | PF05362.16 | Lon_C | 5.00E-09 |
| D2ZB05 | EcancerogenusATCC35316 | PF00698.24 | Acyl_transf_1 | 1.40E-38 |
| D2ZB95 | EcancerogenusATCC35316 | PF01343.21 | Peptidase_S49 | 4.20E-51 |
| D2ZC78 | EcancerogenusATCC35316 | PF00135.31 | COesterase | 5.20E-89 |
| D2ZCG6 | EcancerogenusATCC35316 | PF12146.11 | Hydrolase_4 | 1.40E-13 |
| D2ZCS5 | EcancerogenusATCC35316 | PF00717.26 | Peptidase_S24 | 2.90E-27 |
| D2ZD03 | EcancerogenusATCC35316 | PF01343.21 | Peptidase_S49 | 2.30E-46 |
| D2ZD34 | EcancerogenusATCC35316 | PF12146.11 | Hydrolase_4 | 1.50E-07 |

| | | | | |
|---|---|---|---|---|
| D2ZD75 | EcancerogenusATCC35316 | PF05929.14 | Phage_GPO | 4.30E-103 |
| D2ZDA3 | EcancerogenusATCC35316 | PF01734.25 | Patatin | 6.70E-22 |
| D2ZDG0 | EcancerogenusATCC35316 | PF02016.18 | Peptidase_S66 | 1.10E-20 |
| D2ZDK2 | EcancerogenusATCC35316 | PF03572.21 | Peptidase_S41 | 1.70E-52 |
| D2ZDM0 | EcancerogenusATCC35316 | PF00326.24 | Peptidase_S9 | 2.10E-66 |
| D2ZDV1 | EcancerogenusATCC35316 | PF12146.11 | Hydrolase_4 | 2.20E-22 |
| D2ZE23 | EcancerogenusATCC35316 | PF00717.26 | Peptidase_S24 | 6.50E-27 |
| D2ZE79 | EcancerogenusATCC35316 | PF00144.27 | Beta-lactamase | 1.50E-42 |
| D2ZEA7 | EcancerogenusATCC35316 | PF00768.23 | Peptidase_S11 | 1.30E-94 |
| D2ZEL8 | EcancerogenusATCC35316 | PF00768.23 | Peptidase_S11 | 6.80E-87 |
| D2ZEP2 | EcancerogenusATCC35316 | PF00756.23 | Esterase | 2.40E-70 |
| D2ZEW2 | EcancerogenusATCC35316 | PF12697.10 | Abhydrolase_6 | 9.60E-16 |
| D2ZF13 | EcancerogenusATCC35316 | PF02674.19 | Colicin_V | 9.50E-37 |
| D2ZF95 | EcancerogenusATCC35316 | PF02230.19 | Abhydrolase_2 | 8.70E-48 |
| D2ZFC3 | EcancerogenusATCC35316 | PF12697.10 | Abhydrolase_6 | 4.00E-19 |
| D2ZFJ0 | EcancerogenusATCC35316 | PF10502.12 | Peptidase_S26 | 6.70E-58 |
| D2ZFR8 | EcancerogenusATCC35316 | PF06500.14 | FrsA-like | 1.20E-216 |
| D2ZFU8 | EcancerogenusATCC35316 | PF00144.27 | Beta-lactamase | 3.30E-62 |
| D2ZG22 | EcancerogenusATCC35316 | PF00574.26 | CLP_protease | 1.40E-89 |
| D2ZG24 | EcancerogenusATCC35316 | PF05362.16 | Lon_C | 4.60E-98 |
| D2ZG90 | EcancerogenusATCC35316 | PF13472.9 | Lipase_GDSL_2 | 1.10E-24 |
| D2ZGI5 | EcancerogenusATCC35316 | PF00756.23 | Esterase | 7.20E-39 |
| D2ZGI7 | EcancerogenusATCC35316 | PF00975.23 | Thioesterase | 5.70E-45 |
| D2ZGQ2 | EcancerogenusATCC35316 | PF00768.23 | Peptidase_S11 | 2.00E-103 |
| D2ZGT7 | EcancerogenusATCC35316 | PF12697.10 | Abhydrolase_6 | 8.30E-19 |
| D2ZH71 | EcancerogenusATCC35316 | PF00756.23 | Esterase | 1.80E-08 |
| D2ZH72 | EcancerogenusATCC35316 | PF00756.23 | Esterase | 8.80E-40 |
| D2ZHL3 | EcancerogenusATCC35316 | PF00717.26 | Peptidase_S24 | 5.90E-32 |
| D2ZIL2 | EcancerogenusATCC35316 | PF00698.24 | Acyl_transf_1 | 1.40E-25 |
| D2ZIT0 | EcancerogenusATCC35316 | PF01694.25 | Rhomboid | 7.90E-28 |
| D2ZJ78 | EcancerogenusATCC35316 | PF01425.24 | Amidase | 5.50E-72 |
| D2ZJ88 | EcancerogenusATCC35316 | PF12697.10 | Abhydrolase_6 | 7.90E-14 |
| D2ZJW9 | EcancerogenusATCC35316 | PF00326.24 | Peptidase_S9 | 2.00E-57 |
| D2ZKY0 | EcancerogenusATCC35316 | PF02113.18 | Peptidase_S13 | 2.10E-167 |
| D2ZL42 | EcancerogenusATCC35316 | PF13365.9 | Trypsin_2 | 1.70E-28 |
| D2ZL43 | EcancerogenusATCC35316 | PF13365.9 | Trypsin_2 | 4.10E-29 |
| D2ZL87 | EcancerogenusATCC35316 | PF04586.20 | Peptidase_S78 | 3.90E-48 |
| D2ZLH5 | EcancerogenusATCC35316 | PF00144.27 | Beta-lactamase | 9.30E-86 |
| D2ZLK8 | EcancerogenusATCC35316 | PF00326.24 | Peptidase_S9 | 2.90E-14 |
| D2ZLX7 | EcancerogenusATCC35316 | PF07859.16 | Abhydrolase_3 | 1.50E-62 |
| D2ZM60 | EcancerogenusATCC35316 | PF12146.11 | Hydrolase_4 | 9.20E-57 |
| D2ZME4 | EcancerogenusATCC35316 | PF00717.26 | Peptidase_S24 | 1.50E-30 |
| D5RA65 | FnucleatumATCC23726 | PF00082.25 | Peptidase_S8 | 6.60E-20 |
| D5RAD2 | FnucleatumATCC23726 | PF02674.19 | Colicin_V | 1.60E-17 |
| D5RB77 | FnucleatumATCC23726 | PF13354.9 | Beta-lactamase2 | 4.80E-61 |
| D5RB82 | FnucleatumATCC23726 | PF00717.26 | Peptidase_S24 | 3.90E-28 |
| D5RB84 | FnucleatumATCC23726 | PF00082.25 | Peptidase_S8 | 8.60E-20 |
| D5RBR0 | FnucleatumATCC23726 | PF12146.11 | Hydrolase_4 | 4.50E-51 |
| D5RBX7 | FnucleatumATCC23726 | PF10502.12 | Peptidase_S26 | 2.50E-40 |
| D5RC68 | FnucleatumATCC23726 | PF05362.16 | Lon_C | 1.10E-08 |
| D5RC76 | FnucleatumATCC23726 | PF00698.24 | Acyl_transf_1 | 6.30E-45 |
| D5RCA6 | FnucleatumATCC23726 | PF12146.11 | Hydrolase_4 | 3.30E-12 |

| | | | | |
|---|---|---|---|---|
| D5RCG9 | FnucleatumATCC23726 | PF00326.24 | Peptidase_S9 | 2.40E-56 |
| D5RCI4 | FnucleatumATCC23726 | PF03575.20 | Peptidase_S51 | 5.20E-16 |
| D5RCM5 | FnucleatumATCC23726 | PF07819.16 | PGAP1 | 3.90E-06 |
| D5RCZ0 | FnucleatumATCC23726 | PF00082.25 | Peptidase_S8 | 3.10E-23 |
| D5RD53 | FnucleatumATCC23726 | PF03572.21 | Peptidase_S41 | 1.60E-56 |
| D5RDI5 | FnucleatumATCC23726 | PF05362.16 | Lon_C | 1.40E-80 |
| D5RDI7 | FnucleatumATCC23726 | PF00574.26 | CLP_protease | 1.30E-84 |
| D5RDP1 | FnucleatumATCC23726 | PF05362.16 | Lon_C | 2.40E-05 |
| D5RDX7 | FnucleatumATCC23726 | PF01425.24 | Amidase | 1.10E-154 |
| D5RE18 | FnucleatumATCC23726 | PF00768.23 | Peptidase_S11 | 4.90E-45 |
| D5REA6 | FnucleatumATCC23726 | PF01734.25 | Patatin | 2.30E-16 |
| D5REB4 | FnucleatumATCC23726 | PF01343.21 | Peptidase_S49 | 5.10E-38 |
| D5RES6 | FnucleatumATCC23726 | PF02016.18 | Peptidase_S66 | 1.40E-26 |
| D5REV2 | FnucleatumATCC23726 | PF13365.9 | Trypsin_2 | 9.50E-21 |
| D5REW1 | FnucleatumATCC23726 | PF01343.21 | Peptidase_S49 | 9.10E-39 |
| D5RFI3 | FnucleatumATCC23726 | PF01734.25 | Patatin | 5.90E-30 |
| D5RFJ8 | FnucleatumATCC23726 | PF00082.25 | Peptidase_S8 | 1.00E-22 |
| C9KIM2 | MmultacidaDSM20544 | PF01425.24 | Amidase | 2.20E-168 |
| C9KIR6 | MmultacidaDSM20544 | PF00768.23 | Peptidase_S11 | 1.20E-64 |
| C9KJ11 | MmultacidaDSM20544 | PF00326.24 | Peptidase_S9 | 3.40E-14 |
| C9KJV3 | MmultacidaDSM20544 | PF00326.24 | Peptidase_S9 | 3.20E-08 |
| C9KK96 | MmultacidaDSM20544 | PF13472.9 | Lipase_GDSL_2 | 4.80E-15 |
| C9KKA6 | MmultacidaDSM20544 | PF00574.26 | CLP_protease | 2.80E-88 |
| C9KKA8 | MmultacidaDSM20544 | PF05362.16 | Lon_C | 5.10E-97 |
| C9KKD3 | MmultacidaDSM20544 | PF13472.9 | Lipase_GDSL_2 | 4.10E-14 |
| C9KKJ5 | MmultacidaDSM20544 | PF00717.26 | Peptidase_S24 | 8.40E-12 |
| C9KKJ6 | MmultacidaDSM20544 | PF00768.23 | Peptidase_S11 | 2.30E-54 |
| C9KL17 | MmultacidaDSM20544 | PF00756.23 | Esterase | 1.50E-12 |
| C9KLD4 | MmultacidaDSM20544 | PF01764.28 | Lipase_3 | 9.20E-22 |
| C9KLG2 | MmultacidaDSM20544 | PF00756.23 | Esterase | 1.70E-05 |
| C9KLH5 | MmultacidaDSM20544 | PF13472.9 | Lipase_GDSL_2 | 1.60E-15 |
| C9KLK5 | MmultacidaDSM20544 | PF13365.9 | Trypsin_2 | 8.70E-34 |
| C9KM28 | MmultacidaDSM20544 | PF05362.16 | Lon_C | 1.30E-33 |
| C9KNB0 | MmultacidaDSM20544 | PF12146.11 | Hydrolase_4 | 1.40E-17 |
| C9KNK3 | MmultacidaDSM20544 | PF03572.21 | Peptidase_S41 | 2.70E-57 |
| C9KP04 | MmultacidaDSM20544 | PF05362.16 | Lon_C | 2.00E-07 |
| C9KP05 | MmultacidaDSM20544 | PF00768.23 | Peptidase_S11 | 1.10E-62 |
| C9KP26 | MmultacidaDSM20544 | PF02129.21 | Peptidase_S15 | 6.50E-09 |
| C9KP32 | MmultacidaDSM20544 | PF12697.10 | Abhydrolase_6 | 9.70E-08 |
| C9KP39 | MmultacidaDSM20544 | PF02129.21 | Peptidase_S15 | 3.40E-12 |
| C9KP53 | MmultacidaDSM20544 | PF00135.31 | COesterase | 1.20E-18 |
| C9KP86 | MmultacidaDSM20544 | PF12697.10 | Abhydrolase_6 | 2.90E-07 |
| C9KPS2 | MmultacidaDSM20544 | PF10502.12 | Peptidase_S26 | 2.90E-50 |
| C9KQ81 | MmultacidaDSM20544 | PF00698.24 | Acyl_transf_1 | 1.80E-40 |
| C9KQS6 | MmultacidaDSM20544 | PF13365.9 | Trypsin_2 | 6.10E-35 |
| C9KQW1 | MmultacidaDSM20544 | PF05580.15 | Peptidase_S55 | 8.70E-12 |
| D4F0A3 | EtardaATCC23685 | PF00717.26 | Peptidase_S24 | 5.20E-32 |
| D4F0N3 | EtardaATCC23685 | PF00326.24 | Peptidase_S9 | 2.80E-11 |
| D4F0R9 | EtardaATCC23685 | PF02113.18 | Peptidase_S13 | 3.40E-158 |
| D4F100 | EtardaATCC23685 | PF05929.14 | Phage_GPO | 3.30E-104 |
| D4F154 | EtardaATCC23685 | PF01734.25 | Patatin | 6.80E-09 |
| D4F183 | EtardaATCC23685 | PF13365.9 | Trypsin_2 | 7.40E-30 |

| | | | | |
|---|---|---|---|---|
| D4F184 | EtardaATCC23685 | PF13365.9 | Trypsin_2 | 1.40E-30 |
| D4F2M5 | EtardaATCC23685 | PF06500.14 | FrsA-like | 3.80E-123 |
| D4F2M6 | EtardaATCC23685 | PF06500.14 | FrsA-like | 9.90E-68 |
| D4F2N7 | EtardaATCC23685 | PF02395.19 | Peptidase_S6 | 1.50E-129 |
| D4F2S3 | EtardaATCC23685 | PF00768.23 | Peptidase_S11 | 2.90E-87 |
| D4F2T5 | EtardaATCC23685 | PF03629.21 | SASA | 1.10E-08 |
| D4F301 | EtardaATCC23685 | PF00574.26 | CLP_protease | 2.00E-89 |
| D4F303 | EtardaATCC23685 | PF05362.16 | Lon_C | 2.80E-98 |
| D4F3A9 | EtardaATCC23685 | PF02230.19 | Abhydrolase_2 | 5.40E-27 |
| D4F3U8 | EtardaATCC23685 | PF05362.16 | Lon_C | 5.50E-10 |
| D4F482 | EtardaATCC23685 | PF12697.10 | Abhydrolase_6 | 1.40E-14 |
| D4F4I9 | EtardaATCC23685 | PF01343.21 | Peptidase_S49 | 1.30E-46 |
| D4F4R3 | EtardaATCC23685 | PF01343.21 | Peptidase_S49 | 4.60E-47 |
| D4F535 | EtardaATCC23685 | PF00717.26 | Peptidase_S24 | 2.40E-19 |
| D4F561 | EtardaATCC23685 | PF01343.21 | Peptidase_S49 | 2.30E-31 |
| D4F5B0 | EtardaATCC23685 | PF00326.24 | Peptidase_S9 | 3.60E-33 |
| D4F5B1 | EtardaATCC23685 | PF12697.10 | Abhydrolase_6 | 5.90E-18 |
| D4F5H3 | EtardaATCC23685 | PF00089.29 | Trypsin | 3.90E-08 |
| D4F5L2 | EtardaATCC23685 | PF02129.21 | Peptidase_S15 | 2.70E-13 |
| D4F5U5 | EtardaATCC23685 | PF03572.21 | Peptidase_S41 | 9.60E-53 |
| D4F5Z5 | EtardaATCC23685 | PF01972.19 | SDH_sah | 1.60E-06 |
| D4F607 | EtardaATCC23685 | PF00717.26 | Peptidase_S24 | 1.70E-23 |
| D4F6U4 | EtardaATCC23685 | PF00698.24 | Acyl_transf_1 | 6.30E-39 |
| D4F7D4 | EtardaATCC23685 | PF00768.23 | Peptidase_S11 | 6.90E-98 |
| D4F7P0 | EtardaATCC23685 | PF12697.10 | Abhydrolase_6 | 2.90E-14 |
| D4F7U4 | EtardaATCC23685 | PF02674.19 | Colicin_V | 1.30E-37 |
| D4F812 | EtardaATCC23685 | PF04586.20 | Peptidase_S78 | 5.60E-46 |
| D4F839 | EtardaATCC23685 | PF00717.26 | Peptidase_S24 | 2.60E-26 |
| D4F8A6 | EtardaATCC23685 | PF00768.23 | Peptidase_S11 | 2.00E-71 |
| D4F8I2 | EtardaATCC23685 | PF12697.10 | Abhydrolase_6 | 1.50E-16 |
| D4F8M8 | EtardaATCC23685 | PF00768.23 | Peptidase_S11 | 6.40E-101 |
| D4F8Q4 | EtardaATCC23685 | PF07859.16 | Abhydrolase_3 | 4.40E-50 |
| D4F8V1 | EtardaATCC23685 | PF13472.9 | Lipase_GDSL_2 | 2.40E-22 |
| D4F8X2 | EtardaATCC23685 | PF10502.12 | Peptidase_S26 | 3.20E-57 |
| D4F916 | EtardaATCC23685 | PF13365.9 | Trypsin_2 | 8.70E-31 |
| D4F978 | EtardaATCC23685 | PF00144.27 | Beta-lactamase | 1.10E-76 |
| D4F9H5 | EtardaATCC23685 | PF00144.27 | Beta-lactamase | 5.70E-61 |
| D4F9Z2 | EtardaATCC23685 | PF03575.20 | Peptidase_S51 | 2.80E-46 |
| D4FAC9 | EtardaATCC23685 | PF01694.25 | Rhomboid | 6.40E-27 |
| D4FAG0 | EtardaATCC23685 | PF12146.11 | Hydrolase_4 | 3.10E-40 |
| D4FAG1 | EtardaATCC23685 | PF12146.11 | Hydrolase_4 | 5.60E-10 |
| E2N6Z1 | BcellulosilyticusDSM14838 | PF01343.21 | Peptidase_S49 | 1.90E-16 |
| E2N712 | BcellulosilyticusDSM14838 | PF03629.21 | SASA | 9.30E-16 |
| E2N746 | BcellulosilyticusDSM14838 | PF13472.9 | Lipase_GDSL_2 | 4.30E-13 |
| E2N764 | BcellulosilyticusDSM14838 | PF03629.21 | SASA | 4.90E-22 |
| E2N767 | BcellulosilyticusDSM14838 | PF03629.21 | SASA | 4.70E-19 |
| E2N787 | BcellulosilyticusDSM14838 | PF03629.21 | SASA | 1.50E-16 |
| E2N788 | BcellulosilyticusDSM14838 | PF03629.21 | SASA | 1.70E-11 |
| E2N7R4 | BcellulosilyticusDSM14838 | PF13365.9 | Trypsin_2 | 3.40E-34 |
| E2N7Z8 | BcellulosilyticusDSM14838 | PF04586.20 | Peptidase_S78 | 3.30E-34 |
| E2N830 | BcellulosilyticusDSM14838 | PF00717.26 | Peptidase_S24 | 1.60E-09 |
| E2N868 | BcellulosilyticusDSM14838 | PF00717.26 | Peptidase_S24 | 2.00E-10 |

| | | | | |
|---|---|---|---|---|
| E2N913 | BcellulosilyticusDSM14838 | PF10502.12 | Peptidase_S26 | 5.30E-27 |
| E2N980 | BcellulosilyticusDSM14838 | PF00574.26 | CLP_protease | 1.60E-76 |
| E2N9D1 | BcellulosilyticusDSM14838 | PF13472.9 | Lipase_GDSL_2 | 4.60E-12 |
| E2N9G4 | BcellulosilyticusDSM14838 | PF03572.21 | Peptidase_S41 | 2.30E-11 |
| E2N9X4 | BcellulosilyticusDSM14838 | PF01343.21 | Peptidase_S49 | 1.50E-51 |
| E2NA18 | BcellulosilyticusDSM14838 | PF00756.23 | Esterase | 5.80E-23 |
| E2NA23 | BcellulosilyticusDSM14838 | PF00082.25 | Peptidase_S8 | 2.60E-35 |
| E2NA77 | BcellulosilyticusDSM14838 | PF03629.21 | SASA | 6.20E-20 |
| E2NAG8 | BcellulosilyticusDSM14838 | PF00144.27 | Beta-lactamase | 1.20E-25 |
| E2NAP6 | BcellulosilyticusDSM14838 | PF00144.27 | Beta-lactamase | 1.00E-62 |
| E2NAP6 | BcellulosilyticusDSM14838 | PF13472.9 | Lipase_GDSL_2 | 9.50E-16 |
| E2NAQ2 | BcellulosilyticusDSM14838 | PF03572.21 | Peptidase_S41 | 4.20E-34 |
| E2NB10 | BcellulosilyticusDSM14838 | PF03629.21 | SASA | 1.50E-09 |
| E2NB81 | BcellulosilyticusDSM14838 | PF02674.19 | Colicin_V | 3.40E-31 |
| E2NB89 | BcellulosilyticusDSM14838 | PF00756.23 | Esterase | 8.60E-26 |
| E2NBC6 | BcellulosilyticusDSM14838 | PF00082.25 | Peptidase_S8 | 4.40E-33 |
| E2NBJ1 | BcellulosilyticusDSM14838 | PF10502.12 | Peptidase_S26 | 4.60E-28 |
| E2NBJ2 | BcellulosilyticusDSM14838 | PF10502.12 | Peptidase_S26 | 1.10E-17 |
| E2NBN0 | BcellulosilyticusDSM14838 | PF10502.12 | Peptidase_S26 | 3.20E-30 |
| E2NBR9 | BcellulosilyticusDSM14838 | PF10459.12 | Peptidase_S46 | 3.00E-258 |
| E2NBU1 | BcellulosilyticusDSM14838 | PF01734.25 | Patatin | 5.30E-23 |
| E2NBV5 | BcellulosilyticusDSM14838 | PF10502.12 | Peptidase_S26 | 4.10E-32 |
| E2NBV7 | BcellulosilyticusDSM14838 | PF03629.21 | SASA | 1.70E-09 |
| E2NC38 | BcellulosilyticusDSM14838 | PF13472.9 | Lipase_GDSL_2 | 7.60E-23 |
| E2NC76 | BcellulosilyticusDSM14838 | PF03572.21 | Peptidase_S41 | 8.30E-20 |
| E2NCB5 | BcellulosilyticusDSM14838 | PF10502.12 | Peptidase_S26 | 3.30E-29 |
| E2NCE0 | BcellulosilyticusDSM14838 | PF03572.21 | Peptidase_S41 | 1.50E-19 |
| E2NCF1 | BcellulosilyticusDSM14838 | PF00326.24 | Peptidase_S9 | 8.60E-46 |
| E2NCK0 | BcellulosilyticusDSM14838 | PF13365.9 | Trypsin_2 | 2.20E-26 |
| E2NCM7 | BcellulosilyticusDSM14838 | PF00756.23 | Esterase | 2.60E-12 |
| E2NCQ4 | BcellulosilyticusDSM14838 | PF00756.23 | Esterase | 4.40E-25 |
| E2NCT5 | BcellulosilyticusDSM14838 | PF02113.18 | Peptidase_S13 | 4.70E-83 |
| E2NCW7 | BcellulosilyticusDSM14838 | PF13472.9 | Lipase_GDSL_2 | 1.80E-11 |
| E2NCY6 | BcellulosilyticusDSM14838 | PF00756.23 | Esterase | 1.80E-29 |
| E2NCY7 | BcellulosilyticusDSM14838 | PF00756.23 | Esterase | 1.20E-21 |
| E2NCZ6 | BcellulosilyticusDSM14838 | PF00756.23 | Esterase | 3.00E-19 |
| E2ND19 | BcellulosilyticusDSM14838 | PF00326.24 | Peptidase_S9 | 8.90E-46 |
| E2ND86 | BcellulosilyticusDSM14838 | PF03572.21 | Peptidase_S41 | 3.00E-53 |
| E2NDM1 | BcellulosilyticusDSM14838 | PF05448.15 | AXE1 | 5.10E-38 |
| E2NDM8 | BcellulosilyticusDSM14838 | PF00144.27 | Beta-lactamase | 3.40E-47 |
| E2NDM9 | BcellulosilyticusDSM14838 | PF00144.27 | Beta-lactamase | 1.70E-45 |
| E2NDV9 | BcellulosilyticusDSM14838 | PF01694.25 | Rhomboid | 1.00E-23 |
| E2NDW0 | BcellulosilyticusDSM14838 | PF01694.25 | Rhomboid | 3.60E-32 |
| E2NDY7 | BcellulosilyticusDSM14838 | PF03572.21 | Peptidase_S41 | 7.20E-52 |
| E2NE15 | BcellulosilyticusDSM14838 | PF03629.21 | SASA | 1.10E-22 |
| E2NE21 | BcellulosilyticusDSM14838 | PF00326.24 | Peptidase_S9 | 5.30E-10 |
| E2NE21 | BcellulosilyticusDSM14838 | PF13472.9 | Lipase_GDSL_2 | 5.20E-23 |
| E2NE50 | BcellulosilyticusDSM14838 | PF02016.18 | Peptidase_S66 | 1.60E-35 |
| E2NE92 | BcellulosilyticusDSM14838 | PF03629.21 | SASA | 1.90E-10 |
| E2NEB0 | BcellulosilyticusDSM14838 | PF00326.24 | Peptidase_S9 | 8.10E-10 |
| E2NEB0 | BcellulosilyticusDSM14838 | PF13472.9 | Lipase_GDSL_2 | 3.70E-29 |
| E2NED9 | BcellulosilyticusDSM14838 | PF00144.27 | Beta-lactamase | 5.10E-33 |

| | | | | |
|---|---|---|---|---|
| E2NEN5 | BcellulosilyticusDSM14838 | PF13472.9 | Lipase_GDSL_2 | 2.20E-14 |
| E2NES6 | BcellulosilyticusDSM14838 | PF12697.10 | Abhydrolase_6 | 5.60E-19 |
| E2NET9 | BcellulosilyticusDSM14838 | PF05448.15 | AXE1 | 1.30E-07 |
| E2NEZ7 | BcellulosilyticusDSM14838 | PF03629.21 | SASA | 5.40E-17 |
| E2NF64 | BcellulosilyticusDSM14838 | PF03572.21 | Peptidase_S41 | 2.40E-19 |
| E2NFL0 | BcellulosilyticusDSM14838 | PF10502.12 | Peptidase_S26 | 6.20E-34 |
| E2NFN9 | BcellulosilyticusDSM14838 | PF00717.26 | Peptidase_S24 | 7.30E-19 |
| E2NFQ1 | BcellulosilyticusDSM14838 | PF13472.9 | Lipase_GDSL_2 | 7.30E-14 |
| E2NFQ2 | BcellulosilyticusDSM14838 | PF05448.15 | AXE1 | 4.90E-49 |
| E2NFS6 | BcellulosilyticusDSM14838 | PF00326.24 | Peptidase_S9 | 4.80E-58 |
| E2NFW0 | BcellulosilyticusDSM14838 | PF12146.11 | Hydrolase_4 | 1.70E-12 |
| E2NFY8 | BcellulosilyticusDSM14838 | PF00326.24 | Peptidase_S9 | 4.20E-10 |
| E2NG83 | BcellulosilyticusDSM14838 | PF03572.21 | Peptidase_S41 | 3.30E-52 |
| E2NGB8 | BcellulosilyticusDSM14838 | PF13472.9 | Lipase_GDSL_2 | 1.00E-10 |
| E2NGC3 | BcellulosilyticusDSM14838 | PF01694.25 | Rhomboid | 1.20E-11 |
| E2NGH8 | BcellulosilyticusDSM14838 | PF03629.21 | SASA | 1.50E-12 |
| E2NGJ2 | BcellulosilyticusDSM14838 | PF00756.23 | Esterase | 8.10E-30 |
| E2NGJ3 | BcellulosilyticusDSM14838 | PF03629.21 | SASA | 1.90E-33 |
| E2NGJ6 | BcellulosilyticusDSM14838 | PF12146.11 | Hydrolase_4 | 4.00E-09 |
| E2NGL2 | BcellulosilyticusDSM14838 | PF03629.21 | SASA | 3.00E-14 |
| E2NHC4 | BcellulosilyticusDSM14838 | PF12697.10 | Abhydrolase_6 | 4.80E-12 |
| E2NI36 | BcellulosilyticusDSM14838 | PF13354.9 | Beta-lactamase2 | 2.40E-36 |
| E2NIA0 | BcellulosilyticusDSM14838 | PF00144.27 | Beta-lactamase | 4.70E-44 |
| E2NIJ2 | BcellulosilyticusDSM14838 | PF13472.9 | Lipase_GDSL_2 | 3.80E-16 |
| E2NJ61 | BcellulosilyticusDSM14838 | PF00326.24 | Peptidase_S9 | 3.40E-56 |
| E2NJM0 | BcellulosilyticusDSM14838 | PF00756.23 | Esterase | 2.40E-18 |
| E2NJP9 | BcellulosilyticusDSM14838 | PF13472.9 | Lipase_GDSL_2 | 2.10E-10 |
| E2NK70 | BcellulosilyticusDSM14838 | PF01734.25 | Patatin | 1.40E-21 |
| E2NKA8 | BcellulosilyticusDSM14838 | PF02129.21 | Peptidase_S15 | 1.70E-07 |
| E2NKB5 | BcellulosilyticusDSM14838 | PF00698.24 | Acyl_transf_1 | 8.50E-38 |
| E2NKI7 | BcellulosilyticusDSM14838 | PF05362.16 | Lon_C | 1.20E-81 |
| E2NKQ5 | BcellulosilyticusDSM14838 | PF01734.25 | Patatin | 2.90E-29 |
| E2NL22 | BcellulosilyticusDSM14838 | PF05576.14 | Peptidase_S37 | 6.30E-59 |
| E2NLB4 | BcellulosilyticusDSM14838 | PF03629.21 | SASA | 3.20E-21 |
| E2NLE8 | BcellulosilyticusDSM14838 | PF10459.12 | Peptidase_S46 | 9.60E-261 |
| E2NLE9 | BcellulosilyticusDSM14838 | PF10459.12 | Peptidase_S46 | 3.10E-258 |
| E2NLM2 | BcellulosilyticusDSM14838 | PF03629.21 | SASA | 4.10E-32 |
| E2NLM3 | BcellulosilyticusDSM14838 | PF03629.21 | SASA | 2.90E-12 |
| E2NLN2 | BcellulosilyticusDSM14838 | PF03629.21 | SASA | 3.70E-51 |
| E2NLU2 | BcellulosilyticusDSM14838 | PF03572.21 | Peptidase_S41 | 3.40E-22 |
| E2NLU4 | BcellulosilyticusDSM14838 | PF00756.23 | Esterase | 1.00E-28 |
| E2NLU8 | BcellulosilyticusDSM14838 | PF05448.15 | AXE1 | 1.20E-16 |
| E2NM93 | BcellulosilyticusDSM14838 | PF00089.29 | Trypsin | 2.90E-09 |
| E2NM95 | BcellulosilyticusDSM14838 | PF00756.23 | Esterase | 4.40E-19 |
| E2NMH0 | BcellulosilyticusDSM14838 | PF01734.25 | Patatin | 3.30E-21 |
| E2NMP1 | BcellulosilyticusDSM14838 | PF00326.24 | Peptidase_S9 | 1.00E-67 |
| C9L3S6 | BhanseniiDSM20583 | PF01694.25 | Rhomboid | 1.60E-19 |
| C9L3X6 | BhanseniiDSM20583 | PF01694.25 | Rhomboid | 1.20E-36 |
| C9L436 | BhanseniiDSM20583 | PF13472.9 | Lipase_GDSL_2 | 7.10E-21 |
| C9L4D7 | BhanseniiDSM20583 | PF00574.26 | CLP_protease | 2.10E-25 |
| C9L519 | BhanseniiDSM20583 | PF13365.9 | Trypsin_2 | 5.70E-15 |
| C9L570 | BhanseniiDSM20583 | PF00082.25 | Peptidase_S8 | 1.60E-19 |

| | | | | |
|---|---|---|---|---|
| C9L5M9 | BhanseniiDSM20583 | PF00082.25 | Peptidase_S8 | 1.80E-46 |
| C9L5N9 | BhanseniiDSM20583 | PF00574.26 | CLP_protease | 5.70E-27 |
| C9L5U7 | BhanseniiDSM20583 | PF00768.23 | Peptidase_S11 | 2.90E-28 |
| C9L5Y1 | BhanseniiDSM20583 | PF00768.23 | Peptidase_S11 | 5.10E-46 |
| C9L631 | BhanseniiDSM20583 | PF10502.12 | Peptidase_S26 | 9.90E-47 |
| C9L693 | BhanseniiDSM20583 | PF00768.23 | Peptidase_S11 | 2.90E-44 |
| C9L697 | BhanseniiDSM20583 | PF00768.23 | Peptidase_S11 | 8.20E-67 |
| C9L701 | BhanseniiDSM20583 | PF00698.24 | Acyl_transf_1 | 2.10E-37 |
| C9L734 | BhanseniiDSM20583 | PF05580.15 | Peptidase_S55 | 4.20E-88 |
| C9L7D4 | BhanseniiDSM20583 | PF00574.26 | CLP_protease | 1.10E-25 |
| C9L845 | BhanseniiDSM20583 | PF13354.9 | Beta-lactamase2 | 1.70E-34 |
| C9L8E8 | BhanseniiDSM20583 | PF00717.26 | Peptidase_S24 | 1.70E-32 |
| C9L8N1 | BhanseniiDSM20583 | PF10502.12 | Peptidase_S26 | 2.70E-38 |
| C9L8T4 | BhanseniiDSM20583 | PF00768.23 | Peptidase_S11 | 2.60E-48 |
| C9L8V0 | BhanseniiDSM20583 | PF00082.25 | Peptidase_S8 | 7.10E-13 |
| C9L932 | BhanseniiDSM20583 | PF12697.10 | Abhydrolase_6 | 8.40E-09 |
| C9L989 | BhanseniiDSM20583 | PF00768.23 | Peptidase_S11 | 1.60E-48 |
| C9L9B9 | BhanseniiDSM20583 | PF01734.25 | Patatin | 1.00E-24 |
| C9L9R0 | BhanseniiDSM20583 | PF02674.19 | Colicin_V | 5.80E-11 |
| C9LA07 | BhanseniiDSM20583 | PF05362.16 | Lon_C | 7.00E-92 |
| C9LA09 | BhanseniiDSM20583 | PF00574.26 | CLP_protease | 3.20E-86 |
| C9LA89 | BhanseniiDSM20583 | PF03572.21 | Peptidase_S41 | 3.80E-55 |
| C9LBB4 | BhanseniiDSM20583 | PF01734.25 | Patatin | 5.80E-15 |
| C9LBR1 | BhanseniiDSM20583 | PF02016.18 | Peptidase_S66 | 1.20E-27 |
| C9LBS8 | BhanseniiDSM20583 | PF12146.11 | Hydrolase_4 | 1.30E-46 |
| C9LC09 | BhanseniiDSM20583 | PF03572.21 | Peptidase_S41 | 8.30E-26 |
| C9LC66 | BhanseniiDSM20583 | PF00082.25 | Peptidase_S8 | 7.60E-10 |
| C9LCJ0 | BhanseniiDSM20583 | PF13365.9 | Trypsin_2 | 1.30E-27 |
| C9LCJ1 | BhanseniiDSM20583 | PF00768.23 | Peptidase_S11 | 1.30E-48 |
| C9LCN4 | BhanseniiDSM20583 | PF04586.20 | Peptidase_S78 | 1.40E-26 |
| C9KQZ5 | BfinegoldiiDSM17565 | PF03572.21 | Peptidase_S41 | 2.40E-51 |
| C9KR99 | BfinegoldiiDSM17565 | PF05448.15 | AXE1 | 2.00E-39 |
| C9KRB0 | BfinegoldiiDSM17565 | PF01694.25 | Rhomboid | 1.00E-23 |
| C9KRB1 | BfinegoldiiDSM17565 | PF01694.25 | Rhomboid | 4.30E-33 |
| C9KRC8 | BfinegoldiiDSM17565 | PF03572.21 | Peptidase_S41 | 4.60E-49 |
| C9KRL6 | BfinegoldiiDSM17565 | PF03572.21 | Peptidase_S41 | 7.10E-18 |
| C9KRN1 | BfinegoldiiDSM17565 | PF13365.9 | Trypsin_2 | 3.50E-06 |
| C9KS09 | BfinegoldiiDSM17565 | PF00698.24 | Acyl_transf_1 | 9.60E-37 |
| C9KS16 | BfinegoldiiDSM17565 | PF01734.25 | Patatin | 5.50E-22 |
| C9KS33 | BfinegoldiiDSM17565 | PF01694.25 | Rhomboid | 2.30E-31 |
| C9KS91 | BfinegoldiiDSM17565 | PF10459.12 | Peptidase_S46 | 4.90E-262 |
| C9KT13 | BfinegoldiiDSM17565 | PF00082.25 | Peptidase_S8 | 1.40E-39 |
| C9KT66 | BfinegoldiiDSM17565 | PF00574.26 | CLP_protease | 1.30E-77 |
| C9KTG4 | BfinegoldiiDSM17565 | PF00082.25 | Peptidase_S8 | 2.50E-42 |
| C9KTT0 | BfinegoldiiDSM17565 | PF10502.12 | Peptidase_S26 | 5.70E-25 |
| C9KTT1 | BfinegoldiiDSM17565 | PF10502.12 | Peptidase_S26 | 4.30E-18 |
| C9KTT4 | BfinegoldiiDSM17565 | PF13472.9 | Lipase_GDSL_2 | 4.20E-09 |
| C9KU58 | BfinegoldiiDSM17565 | PF13354.9 | Beta-lactamase2 | 4.20E-30 |
| C9KUE6 | BfinegoldiiDSM17565 | PF00574.26 | CLP_protease | 1.40E-08 |
| C9KUT4 | BfinegoldiiDSM17565 | PF03629.21 | SASA | 4.50E-20 |
| C9KUV3 | BfinegoldiiDSM17565 | PF13472.9 | Lipase_GDSL_2 | 3.50E-09 |
| C9KUW0 | BfinegoldiiDSM17565 | PF13472.9 | Lipase_GDSL_2 | 2.50E-13 |

| | | | | |
|---|---|---|---|---|
| C9KUX1 | BfinegoldiiDSM17565 | PF13472.9 | Lipase_GDSL_2 | 3.90E-13 |
| C9KUY9 | BfinegoldiiDSM17565 | PF13472.9 | Lipase_GDSL_2 | 2.90E-14 |
| C9KV34 | BfinegoldiiDSM17565 | PF01343.21 | Peptidase_S49 | 2.20E-49 |
| C9KV87 | BfinegoldiiDSM17565 | PF00326.24 | Peptidase_S9 | 1.90E-68 |
| C9KVC1 | BfinegoldiiDSM17565 | PF00717.26 | Peptidase_S24 | 6.80E-15 |
| C9KVE6 | BfinegoldiiDSM17565 | PF00574.26 | CLP_protease | 7.50E-34 |
| C9KW66 | BfinegoldiiDSM17565 | PF13472.9 | Lipase_GDSL_2 | 3.40E-23 |
| C9KW84 | BfinegoldiiDSM17565 | PF00326.24 | Peptidase_S9 | 2.70E-56 |
| C9KWT4 | BfinegoldiiDSM17565 | PF03629.21 | SASA | 1.00E-17 |
| C9KWV6 | BfinegoldiiDSM17565 | PF01734.25 | Patatin | 2.40E-21 |
| C9KXA5 | BfinegoldiiDSM17565 | PF05362.16 | Lon_C | 1.70E-81 |
| C9KXQ0 | BfinegoldiiDSM17565 | PF07859.16 | Abhydrolase_3 | 1.20E-51 |
| C9KXY5 | BfinegoldiiDSM17565 | PF13472.9 | Lipase_GDSL_2 | 1.00E-25 |
| C9KYA4 | BfinegoldiiDSM17565 | PF00144.27 | Beta-lactamase | 2.30E-44 |
| C9KYA4 | BfinegoldiiDSM17565 | PF12697.10 | Abhydrolase_6 | 1.50E-17 |
| C9KYB8 | BfinegoldiiDSM17565 | PF00756.23 | Esterase | 6.50E-28 |
| C9KYC0 | BfinegoldiiDSM17565 | PF03572.21 | Peptidase_S41 | 1.30E-10 |
| C9KYH7 | BfinegoldiiDSM17565 | PF00082.25 | Peptidase_S8 | 4.20E-44 |
| C9KYI6 | BfinegoldiiDSM17565 | PF10459.12 | Peptidase_S46 | 1.70E-259 |
| C9KYI7 | BfinegoldiiDSM17565 | PF10459.12 | Peptidase_S46 | 1.30E-256 |
| C9KZ41 | BfinegoldiiDSM17565 | PF10502.12 | Peptidase_S26 | 5.70E-12 |
| C9KZ42 | BfinegoldiiDSM17565 | PF10502.12 | Peptidase_S26 | 7.20E-07 |
| C9KZF6 | BfinegoldiiDSM17565 | PF01734.25 | Patatin | 2.60E-21 |
| C9KZK4 | BfinegoldiiDSM17565 | PF03572.21 | Peptidase_S41 | 4.70E-30 |
| C9L0K6 | BfinegoldiiDSM17565 | PF02674.19 | Colicin_V | 4.70E-30 |
| C9L0Y8 | BfinegoldiiDSM17565 | PF02016.18 | Peptidase_S66 | 1.70E-37 |
| C9L140 | BfinegoldiiDSM17565 | PF00144.27 | Beta-lactamase | 5.20E-46 |
| C9L1G3 | BfinegoldiiDSM17565 | PF00082.25 | Peptidase_S8 | 1.30E-45 |
| C9L1J3 | BfinegoldiiDSM17565 | PF00326.24 | Peptidase_S9 | 9.40E-46 |
| C9L1L6 | BfinegoldiiDSM17565 | PF00326.24 | Peptidase_S9 | 4.20E-37 |
| C9L1L7 | BfinegoldiiDSM17565 | PF00326.24 | Peptidase_S9 | 2.00E-30 |
| C9L1L8 | BfinegoldiiDSM17565 | PF00326.24 | Peptidase_S9 | 4.10E-23 |
| C9L1P8 | BfinegoldiiDSM17565 | PF00326.24 | Peptidase_S9 | 2.60E-34 |
| C9L1P9 | BfinegoldiiDSM17565 | PF00082.25 | Peptidase_S8 | 2.30E-54 |
| C9L1Q3 | BfinegoldiiDSM17565 | PF02113.18 | Peptidase_S13 | 7.40E-82 |
| C9L1Q8 | BfinegoldiiDSM17565 | PF00717.26 | Peptidase_S24 | 3.00E-07 |
| C9L1T5 | BfinegoldiiDSM17565 | PF03572.21 | Peptidase_S41 | 5.20E-53 |
| C9L1V2 | BfinegoldiiDSM17565 | PF01694.25 | Rhomboid | 1.50E-12 |
| C9L252 | BfinegoldiiDSM17565 | PF01734.25 | Patatin | 2.50E-25 |
| C9L2K9 | BfinegoldiiDSM17565 | PF00326.24 | Peptidase_S9 | 1.00E-14 |
| C9L2M1 | BfinegoldiiDSM17565 | PF00326.24 | Peptidase_S9 | 3.40E-59 |
| C9L379 | BfinegoldiiDSM17565 | PF13472.9 | Lipase_GDSL_2 | 9.60E-14 |
| C9L3C3 | BfinegoldiiDSM17565 | PF02129.21 | Peptidase_S15 | 2.50E-09 |
| C9L3D1 | BfinegoldiiDSM17565 | PF01734.25 | Patatin | 2.90E-28 |
| C9L3G5 | BfinegoldiiDSM17565 | PF13365.9 | Trypsin_2 | 3.50E-34 |
| E7FM85 | LruminisATCC25644 | PF10502.12 | Peptidase_S26 | 5.50E-38 |
| E7FMP2 | LruminisATCC25644 | PF02129.21 | Peptidase_S15 | 8.50E-80 |
| E7FMS7 | LruminisATCC25644 | PF12146.11 | Hydrolase_4 | 1.10E-10 |
| E7FMU6 | LruminisATCC25644 | PF05448.15 | AXE1 | 2.70E-13 |
| E7FNC9 | LruminisATCC25644 | PF00144.27 | Beta-lactamase | 1.20E-31 |
| E7FNT8 | LruminisATCC25644 | PF10502.12 | Peptidase_S26 | 1.70E-40 |
| E7FNW0 | LruminisATCC25644 | PF10502.12 | Peptidase_S26 | 1.40E-44 |

| | | | | |
|---|---|---|---|---|
| E7FNW1 | LruminisATCC25644 | PF03572.21 | Peptidase_S41 | 7.60E-49 |
| E7FNW4 | LruminisATCC25644 | PF13472.9 | Lipase_GDSL_2 | 1.60E-26 |
| E7FP26 | LruminisATCC25644 | PF00144.27 | Beta-lactamase | 1.50E-49 |
| E7FPD0 | LruminisATCC25644 | PF01734.25 | Patatin | 6.00E-07 |
| E7FPR6 | LruminisATCC25644 | PF00574.26 | CLP_protease | 8.50E-83 |
| E7FQC1 | LruminisATCC25644 | PF12146.11 | Hydrolase_4 | 2.30E-48 |
| E7FQL4 | LruminisATCC25644 | PF12695.10 | Abhydrolase_5 | 2.90E-53 |
| E7FR08 | LruminisATCC25644 | PF05362.16 | Lon_C | 9.10E-08 |
| E7FR32 | LruminisATCC25644 | PF01694.25 | Rhomboid | 4.40E-36 |
| E7FRD3 | LruminisATCC25644 | PF07859.16 | Abhydrolase_3 | 9.90E-48 |
| E7FRI4 | LruminisATCC25644 | PF00717.26 | Peptidase_S24 | 5.10E-29 |
| E7FRR7 | LruminisATCC25644 | PF13365.9 | Trypsin_2 | 9.50E-29 |
| E7FRY4 | LruminisATCC25644 | PF10502.12 | Peptidase_S26 | 4.80E-25 |
| E7FS97 | LruminisATCC25644 | PF00082.25 | Peptidase_S8 | 5.00E-53 |
| E7FSC6 | LruminisATCC25644 | PF01425.24 | Amidase | 2.20E-72 |
| E7FSU5 | LruminisATCC25644 | PF01425.24 | Amidase | 1.00E-150 |
| E7FTC8 | LruminisATCC25644 | PF00698.24 | Acyl_transf_1 | 3.30E-29 |
| E7FTE6 | LruminisATCC25644 | PF02674.19 | Colicin_V | 1.00E-22 |
| E7FTI6 | LruminisATCC25644 | PF00768.23 | Peptidase_S11 | 3.50E-69 |
| A7UXU9 | BuniformisATCC8492 | PF13472.9 | Lipase_GDSL_2 | 1.00E-10 |
| A7UY07 | BuniformisATCC8492 | PF03572.21 | Peptidase_S41 | 1.40E-24 |
| A7UY40 | BuniformisATCC8492 | PF00144.27 | Beta-lactamase | 3.70E-31 |
| A7UY93 | BuniformisATCC8492 | PF00144.27 | Beta-lactamase | 1.20E-61 |
| A7UY93 | BuniformisATCC8492 | PF13472.9 | Lipase_GDSL_2 | 9.60E-18 |
| A7UY98 | BuniformisATCC8492 | PF00756.23 | Esterase | 7.30E-32 |
| A7UYA4 | BuniformisATCC8492 | PF02674.19 | Colicin_V | 3.40E-29 |
| A7UYF6 | BuniformisATCC8492 | PF00144.27 | Beta-lactamase | 9.20E-32 |
| A7UYK8 | BuniformisATCC8492 | PF05448.15 | AXE1 | 1.60E-37 |
| A7UYM0 | BuniformisATCC8492 | PF00144.27 | Beta-lactamase | 1.40E-48 |
| A7UYM1 | BuniformisATCC8492 | PF00144.27 | Beta-lactamase | 2.70E-51 |
| A7UYX9 | BuniformisATCC8492 | PF02113.18 | Peptidase_S13 | 5.30E-83 |
| A7UZ00 | BuniformisATCC8492 | PF00717.26 | Peptidase_S24 | 1.50E-16 |
| A7UZB7 | BuniformisATCC8492 | PF05362.16 | Lon_C | 1.40E-81 |
| A7UZG8 | BuniformisATCC8492 | PF00698.24 | Acyl_transf_1 | 8.70E-38 |
| A7UZI8 | BuniformisATCC8492 | PF01734.25 | Patatin | 6.00E-21 |
| A7UZM5 | BuniformisATCC8492 | PF13472.9 | Lipase_GDSL_2 | 3.80E-16 |
| A7V035 | BuniformisATCC8492 | PF00756.23 | Esterase | 2.50E-24 |
| A7V036 | BuniformisATCC8492 | PF00756.23 | Esterase | 2.00E-21 |
| A7V037 | BuniformisATCC8492 | PF00756.23 | Esterase | 5.50E-28 |
| A7V041 | BuniformisATCC8492 | PF00135.31 | COesterase | 6.50E-102 |
| A7V0B6 | BuniformisATCC8492 | PF01734.25 | Patatin | 1.30E-24 |
| A7V0F0 | BuniformisATCC8492 | PF10502.12 | Peptidase_S26 | 1.80E-26 |
| A7V0F1 | BuniformisATCC8492 | PF10502.12 | Peptidase_S26 | 3.10E-25 |
| A7V0H7 | BuniformisATCC8492 | PF10459.12 | Peptidase_S46 | 2.40E-264 |
| A7V0R6 | BuniformisATCC8492 | PF01734.25 | Patatin | 1.10E-06 |
| A7V149 | BuniformisATCC8492 | PF13472.9 | Lipase_GDSL_2 | 1.30E-23 |
| A7V184 | BuniformisATCC8492 | PF00756.23 | Esterase | 3.30E-32 |
| A7V186 | BuniformisATCC8492 | PF03572.21 | Peptidase_S41 | 1.20E-18 |
| A7V1I5 | BuniformisATCC8492 | PF10459.12 | Peptidase_S46 | 2.00E-265 |
| A7V1I6 | BuniformisATCC8492 | PF10459.12 | Peptidase_S46 | 7.00E-261 |
| A7V2B9 | BuniformisATCC8492 | PF00756.23 | Esterase | 2.50E-18 |
| A7V2T5 | BuniformisATCC8492 | PF05576.14 | Peptidase_S37 | 7.60E-62 |

| | | | | |
|---|---|---|---|---|
| A7V358 | BuniformisATCC8492 | PF01734.25 | Patatin | 3.30E-29 |
| A7V369 | BuniformisATCC8492 | PF13354.9 | Beta-lactamase2 | 2.50E-35 |
| A7V4D7 | BuniformisATCC8492 | PF00326.24 | Peptidase_S9 | 2.20E-42 |
| A7V4H2 | BuniformisATCC8492 | PF01343.21 | Peptidase_S49 | 3.20E-50 |
| A7V4Z8 | BuniformisATCC8492 | PF00717.26 | Peptidase_S24 | 1.20E-09 |
| A7V517 | BuniformisATCC8492 | PF00574.26 | CLP_protease | 1.30E-13 |
| A7V5C1 | BuniformisATCC8492 | PF00326.24 | Peptidase_S9 | 1.00E-55 |
| A7V5D8 | BuniformisATCC8492 | PF13472.9 | Lipase_GDSL_2 | 1.40E-17 |
| A7V5T5 | BuniformisATCC8492 | PF07859.16 | Abhydrolase_3 | 1.00E-13 |
| A7V604 | BuniformisATCC8492 | PF00574.26 | CLP_protease | 8.30E-77 |
| A7V699 | BuniformisATCC8492 | PF00082.25 | Peptidase_S8 | 1.60E-38 |
| A7V6C0 | BuniformisATCC8492 | PF03572.21 | Peptidase_S41 | 3.40E-15 |
| A7V6D1 | BuniformisATCC8492 | PF05448.15 | AXE1 | 7.80E-46 |
| A7V732 | BuniformisATCC8492 | PF03572.21 | Peptidase_S41 | 2.70E-08 |
| A7V741 | BuniformisATCC8492 | PF01694.25 | Rhomboid | 4.10E-12 |
| A7V743 | BuniformisATCC8492 | PF13472.9 | Lipase_GDSL_2 | 9.80E-24 |
| A7V753 | BuniformisATCC8492 | PF03572.21 | Peptidase_S41 | 4.60E-53 |
| A7V7A4 | BuniformisATCC8492 | PF03959.16 | FSH1 | 5.30E-06 |
| A7V7E4 | BuniformisATCC8492 | PF00082.25 | Peptidase_S8 | 1.50E-45 |
| A7V7G7 | BuniformisATCC8492 | PF12146.11 | Hydrolase_4 | 1.60E-09 |
| A7V7K4 | BuniformisATCC8492 | PF10502.12 | Peptidase_S26 | 6.30E-30 |
| A7V7M2 | BuniformisATCC8492 | PF00326.24 | Peptidase_S9 | 1.30E-57 |
| A7V7Z2 | BuniformisATCC8492 | PF13472.9 | Lipase_GDSL_2 | 4.60E-25 |
| A7V806 | BuniformisATCC8492 | PF03572.21 | Peptidase_S41 | 1.60E-16 |
| A7V829 | BuniformisATCC8492 | PF03572.21 | Peptidase_S41 | 1.50E-22 |
| A7V893 | BuniformisATCC8492 | PF03629.21 | SASA | 1.30E-19 |
| A7V899 | BuniformisATCC8492 | PF03572.21 | Peptidase_S41 | 3.10E-32 |
| A7V8K0 | BuniformisATCC8492 | PF02016.18 | Peptidase_S66 | 4.30E-36 |
| A7V8Q8 | BuniformisATCC8492 | PF03572.21 | Peptidase_S41 | 4.40E-49 |
| A7V8S9 | BuniformisATCC8492 | PF01694.25 | Rhomboid | 8.50E-34 |
| A7V8T0 | BuniformisATCC8492 | PF01694.25 | Rhomboid | 5.90E-24 |
| A7V966 | BuniformisATCC8492 | PF03572.21 | Peptidase_S41 | 1.30E-52 |
| A7V993 | BuniformisATCC8492 | PF00326.24 | Peptidase_S9 | 1.40E-45 |
| A7V9B4 | BuniformisATCC8492 | PF03629.21 | SASA | 4.40E-22 |
| A7V9P8 | BuniformisATCC8492 | PF13365.9 | Trypsin_2 | 4.20E-34 |
| A7VA83 | BuniformisATCC8492 | PF13472.9 | Lipase_GDSL_2 | 1.40E-15 |
| A7VAB3 | BuniformisATCC8492 | PF02129.21 | Peptidase_S15 | 1.70E-10 |
| A7A9P6 | PmerdaeATCC43184 | PF01343.21 | Peptidase_S49 | 1.50E-50 |
| A7A9P9 | PmerdaeATCC43184 | PF01734.25 | Patatin | 2.00E-24 |
| A7A9V7 | PmerdaeATCC43184 | PF10502.12 | Peptidase_S26 | 1.90E-28 |
| A7A9V8 | PmerdaeATCC43184 | PF10502.12 | Peptidase_S26 | 1.20E-29 |
| A7AAA4 | PmerdaeATCC43184 | PF10459.12 | Peptidase_S46 | 1.40E-245 |
| A7AAC8 | PmerdaeATCC43184 | PF03629.21 | SASA | 5.60E-18 |
| A7AAC8 | PmerdaeATCC43184 | PF13472.9 | Lipase_GDSL_2 | 4.70E-23 |
| A7AAI3 | PmerdaeATCC43184 | PF13472.9 | Lipase_GDSL_2 | 1.00E-15 |
| A7AAT3 | PmerdaeATCC43184 | PF10502.12 | Peptidase_S26 | 7.30E-30 |
| A7AAY6 | PmerdaeATCC43184 | PF02113.18 | Peptidase_S13 | 3.10E-71 |
| A7AAZ9 | PmerdaeATCC43184 | PF03572.21 | Peptidase_S41 | 4.30E-51 |
| A7AB54 | PmerdaeATCC43184 | PF13365.9 | Trypsin_2 | 2.20E-12 |
| A7AB83 | PmerdaeATCC43184 | PF00326.24 | Peptidase_S9 | 3.20E-06 |
| A7AB83 | PmerdaeATCC43184 | PF13472.9 | Lipase_GDSL_2 | 7.10E-09 |
| A7ABD4 | PmerdaeATCC43184 | PF13472.9 | Lipase_GDSL_2 | 5.30E-20 |

| | | | | |
|---|---|---|---|---|
| A7ABE7 | PmerdaeATCC43184 | PF13472.9 | Lipase_GDSL_2 | 4.10E-15 |
| A7ABK1 | PmerdaeATCC43184 | PF00756.23 | Esterase | 5.10E-21 |
| A7ABY6 | PmerdaeATCC43184 | PF13472.9 | Lipase_GDSL_2 | 4.10E-17 |
| A7AC51 | PmerdaeATCC43184 | PF03572.21 | Peptidase_S41 | 8.20E-50 |
| A7ACQ0 | PmerdaeATCC43184 | PF00082.25 | Peptidase_S8 | 5.20E-33 |
| A7AD16 | PmerdaeATCC43184 | PF00326.24 | Peptidase_S9 | 1.40E-41 |
| A7AD57 | PmerdaeATCC43184 | PF12146.11 | Hydrolase_4 | 4.30E-22 |
| A7AD82 | PmerdaeATCC43184 | PF01734.25 | Patatin | 4.40E-29 |
| A7ADM3 | PmerdaeATCC43184 | PF03572.21 | Peptidase_S41 | 3.50E-45 |
| A7ADN7 | PmerdaeATCC43184 | PF01694.25 | Rhomboid | 2.80E-21 |
| A7ADN8 | PmerdaeATCC43184 | PF01694.25 | Rhomboid | 2.20E-27 |
| A7ADU7 | PmerdaeATCC43184 | PF02674.19 | Colicin_V | 3.00E-15 |
| A7AE05 | PmerdaeATCC43184 | PF00326.24 | Peptidase_S9 | 2.90E-48 |
| A7AE10 | PmerdaeATCC43184 | PF00326.24 | Peptidase_S9 | 8.60E-24 |
| A7AE59 | PmerdaeATCC43184 | PF01343.21 | Peptidase_S49 | 4.50E-09 |
| A7AEH7 | PmerdaeATCC43184 | PF01734.25 | Patatin | 3.40E-27 |
| A7AEL1 | PmerdaeATCC43184 | PF00326.24 | Peptidase_S9 | 3.00E-47 |
| A7AET9 | PmerdaeATCC43184 | PF01734.25 | Patatin | 1.30E-18 |
| A7AFN0 | PmerdaeATCC43184 | PF00326.24 | Peptidase_S9 | 3.40E-38 |
| A7AFU6 | PmerdaeATCC43184 | PF00756.23 | Esterase | 6.00E-39 |
| A7AFV9 | PmerdaeATCC43184 | PF12697.10 | Abhydrolase_6 | 5.90E-07 |
| A7AFX0 | PmerdaeATCC43184 | PF10502.12 | Peptidase_S26 | 2.40E-28 |
| A7AFX9 | PmerdaeATCC43184 | PF02129.21 | Peptidase_S15 | 2.30E-08 |
| A7AGF8 | PmerdaeATCC43184 | PF00698.24 | Acyl_transf_1 | 4.80E-41 |
| A7AGN1 | PmerdaeATCC43184 | PF00756.23 | Esterase | 1.40E-28 |
| A7AH47 | PmerdaeATCC43184 | PF00326.24 | Peptidase_S9 | 9.30E-49 |
| A7AH55 | PmerdaeATCC43184 | PF05576.14 | Peptidase_S37 | 6.80E-52 |
| A7AH72 | PmerdaeATCC43184 | PF03572.21 | Peptidase_S41 | 5.60E-30 |
| A7AHP4 | PmerdaeATCC43184 | PF10459.12 | Peptidase_S46 | 6.50E-263 |
| A7AHU8 | PmerdaeATCC43184 | PF00326.24 | Peptidase_S9 | 7.00E-55 |
| A7AI23 | PmerdaeATCC43184 | PF13472.9 | Lipase_GDSL_2 | 6.00E-16 |
| A7AI74 | PmerdaeATCC43184 | PF03572.21 | Peptidase_S41 | 1.70E-15 |
| A7AI83 | PmerdaeATCC43184 | PF03572.21 | Peptidase_S41 | 4.90E-20 |
| A7AI93 | PmerdaeATCC43184 | PF00082.25 | Peptidase_S8 | 2.00E-32 |
| A7AIE3 | PmerdaeATCC43184 | PF10502.12 | Peptidase_S26 | 4.60E-35 |
| A7AIG9 | PmerdaeATCC43184 | PF01694.25 | Rhomboid | 1.90E-26 |
| A7AII8 | PmerdaeATCC43184 | PF00326.24 | Peptidase_S9 | 7.00E-44 |
| A7AJV0 | PmerdaeATCC43184 | PF00574.26 | CLP_protease | 6.10E-80 |
| A7AK11 | PmerdaeATCC43184 | PF00326.24 | Peptidase_S9 | 1.60E-54 |
| A7AK61 | PmerdaeATCC43184 | PF00144.27 | Beta-lactamase | 2.00E-45 |
| A7AKC8 | PmerdaeATCC43184 | PF05362.16 | Lon_C | 1.80E-82 |
| A7AL82 | PmerdaeATCC43184 | PF05448.15 | AXE1 | 2.50E-47 |
| A7AL88 | PmerdaeATCC43184 | PF03572.21 | Peptidase_S41 | 7.50E-20 |
| A7ALD2 | PmerdaeATCC43184 | PF13365.9 | Trypsin_2 | 3.80E-35 |
| A7ALL9 | PmerdaeATCC43184 | PF13472.9 | Lipase_GDSL_2 | 3.40E-14 |
| A7ALW2 | PmerdaeATCC43184 | PF12697.10 | Abhydrolase_6 | 2.90E-07 |
| A7ALX2 | PmerdaeATCC43184 | PF02129.21 | Peptidase_S15 | 2.50E-07 |
| A7AXU6 | RgnavusATCC29149 | PF13365.9 | Trypsin_2 | 4.90E-08 |
| A7AXY5 | RgnavusATCC29149 | PF03572.21 | Peptidase_S41 | 2.50E-56 |
| A7AXZ5 | RgnavusATCC29149 | PF13472.9 | Lipase_GDSL_2 | 8.50E-15 |
| A7AY03 | RgnavusATCC29149 | PF00082.25 | Peptidase_S8 | 1.40E-09 |
| A7AY63 | RgnavusATCC29149 | PF04586.20 | Peptidase_S78 | 1.90E-11 |

| | | | | |
|---|---|---|---|---|
| A7AYB3 | RgnavusATCC29149 | PF05580.15 | Peptidase_S55 | 2.80E-83 |
| A7AYH0 | RgnavusATCC29149 | PF00768.23 | Peptidase_S11 | 2.40E-73 |
| A7AYH4 | RgnavusATCC29149 | PF00768.23 | Peptidase_S11 | 5.40E-47 |
| A7AZ52 | RgnavusATCC29149 | PF00574.26 | CLP_protease | 1.70E-23 |
| A7AZB1 | RgnavusATCC29149 | PF00768.23 | Peptidase_S11 | 2.40E-38 |
| A7B007 | RgnavusATCC29149 | PF12697.10 | Abhydrolase_6 | 4.50E-09 |
| A7B0N6 | RgnavusATCC29149 | PF10502.12 | Peptidase_S26 | 3.80E-42 |
| A7B0N8 | RgnavusATCC29149 | PF10502.12 | Peptidase_S26 | 4.70E-37 |
| A7B0P6 | RgnavusATCC29149 | PF00082.25 | Peptidase_S8 | 1.10E-44 |
| A7B169 | RgnavusATCC29149 | PF13365.9 | Trypsin_2 | 1.90E-08 |
| A7B1H5 | RgnavusATCC29149 | PF00082.25 | Peptidase_S8 | 7.60E-10 |
| A7B1N3 | RgnavusATCC29149 | PF03572.21 | Peptidase_S41 | 8.30E-26 |
| A7B234 | RgnavusATCC29149 | PF00717.26 | Peptidase_S24 | 1.90E-07 |
| A7B252 | RgnavusATCC29149 | PF00657.25 | Lipase_GDSL | 6.20E-11 |
| A7B271 | RgnavusATCC29149 | PF00698.24 | Acyl_transf_1 | 1.20E-37 |
| A7B2F2 | RgnavusATCC29149 | PF01734.25 | Patatin | 1.70E-15 |
| A7B2X2 | RgnavusATCC29149 | PF00768.23 | Peptidase_S11 | 6.70E-51 |
| A7B2Z6 | RgnavusATCC29149 | PF00717.26 | Peptidase_S24 | 8.10E-32 |
| A7B332 | RgnavusATCC29149 | PF00768.23 | Peptidase_S11 | 2.00E-42 |
| A7B3D5 | RgnavusATCC29149 | PF04586.20 | Peptidase_S78 | 1.40E-26 |
| A7B441 | RgnavusATCC29149 | PF00574.26 | CLP_protease | 5.70E-36 |
| A7B4Z5 | RgnavusATCC29149 | PF03629.21 | SASA | 4.40E-46 |
| A7B564 | RgnavusATCC29149 | PF13472.9 | Lipase_GDSL_2 | 1.20E-22 |
| A7B574 | RgnavusATCC29149 | PF13472.9 | Lipase_GDSL_2 | 1.30E-12 |
| A7B5F0 | RgnavusATCC29149 | PF01734.25 | Patatin | 2.30E-20 |
| A7B5Q3 | RgnavusATCC29149 | PF00082.25 | Peptidase_S8 | 5.90E-15 |
| A7B5T8 | RgnavusATCC29149 | PF10502.12 | Peptidase_S26 | 9.20E-23 |
| A7B686 | RgnavusATCC29149 | PF02674.19 | Colicin_V | 6.20E-12 |
| A7B6D7 | RgnavusATCC29149 | PF12146.11 | Hydrolase_4 | 1.90E-46 |
| A7B6G2 | RgnavusATCC29149 | PF01694.25 | Rhomboid | 4.50E-35 |
| A7B6I3 | RgnavusATCC29149 | PF12146.11 | Hydrolase_4 | 4.50E-14 |
| A7B6U5 | RgnavusATCC29149 | PF07859.16 | Abhydrolase_3 | 1.10E-55 |
| A7B7E4 | RgnavusATCC29149 | PF13365.9 | Trypsin_2 | 1.30E-29 |
| A7B7K4 | RgnavusATCC29149 | PF05362.16 | Lon_C | 1.00E-90 |
| A7B7K6 | RgnavusATCC29149 | PF00574.26 | CLP_protease | 1.50E-87 |
| D1P8K7 | PcopriDSM18205 | PF01343.21 | Peptidase_S49 | 1.70E-43 |
| D1P8K9 | PcopriDSM18205 | PF07859.16 | Abhydrolase_3 | 3.80E-12 |
| D1P8T9 | PcopriDSM18205 | PF03572.21 | Peptidase_S41 | 6.90E-52 |
| D1P990 | PcopriDSM18205 | PF03572.21 | Peptidase_S41 | 1.20E-13 |
| D1P9K2 | PcopriDSM18205 | PF00717.26 | Peptidase_S24 | 3.00E-14 |
| D1PAQ4 | PcopriDSM18205 | PF00144.27 | Beta-lactamase | 1.80E-44 |
| D1PAT0 | PcopriDSM18205 | PF00326.24 | Peptidase_S9 | 3.00E-52 |
| D1PBU3 | PcopriDSM18205 | PF01734.25 | Patatin | 2.20E-19 |
| D1PBV8 | PcopriDSM18205 | PF00698.24 | Acyl_transf_1 | 2.30E-42 |
| D1PBX1 | PcopriDSM18205 | PF00326.24 | Peptidase_S9 | 5.80E-10 |
| D1PC07 | PcopriDSM18205 | PF00326.24 | Peptidase_S9 | 5.20E-05 |
| D1PC11 | PcopriDSM18205 | PF00717.26 | Peptidase_S24 | 1.10E-25 |
| D1PC16 | PcopriDSM18205 | PF00326.24 | Peptidase_S9 | 1.70E-43 |
| D1PC25 | PcopriDSM18205 | PF00326.24 | Peptidase_S9 | 5.30E-53 |
| D1PC28 | PcopriDSM18205 | PF13365.9 | Trypsin_2 | 6.20E-34 |
| D1PCE6 | PcopriDSM18205 | PF00082.25 | Peptidase_S8 | 2.30E-39 |
| D1PCM0 | PcopriDSM18205 | PF13472.9 | Lipase_GDSL_2 | 9.40E-25 |

| | | | | |
|---|---|---|---|---|
| D1PCP7 | PcopriDSM18205 | PF13472.9 | Lipase_GDSL_2 | 6.80E-21 |
| D1PCU2 | PcopriDSM18205 | PF00326.24 | Peptidase_S9 | 7.20E-50 |
| D1PCW9 | PcopriDSM18205 | PF00082.25 | Peptidase_S8 | 1.20E-39 |
| D1PDU3 | PcopriDSM18205 | PF01694.25 | Rhomboid | 4.00E-33 |
| D1PED1 | PcopriDSM18205 | PF00135.31 | COesterase | 4.10E-108 |
| D1PEK5 | PcopriDSM18205 | PF00574.26 | CLP_protease | 4.00E-77 |
| D1PF02 | PcopriDSM18205 | PF13472.9 | Lipase_GDSL_2 | 9.60E-16 |
| D1PFI3 | PcopriDSM18205 | PF13472.9 | Lipase_GDSL_2 | 6.70E-16 |
| D1PFS4 | PcopriDSM18205 | PF10503.12 | Esterase_PHB | 2.40E-10 |
| D1PG85 | PcopriDSM18205 | PF00135.31 | COesterase | 4.50E-106 |
| D1PG87 | PcopriDSM18205 | PF05448.15 | AXE1 | 1.10E-51 |
| D1PG90 | PcopriDSM18205 | PF03572.21 | Peptidase_S41 | 2.20E-47 |
| D1PGA3 | PcopriDSM18205 | PF10502.12 | Peptidase_S26 | 5.70E-25 |
| D1PGE0 | PcopriDSM18205 | PF01734.25 | Patatin | 2.60E-15 |
| D1PGS6 | PcopriDSM18205 | PF05362.16 | Lon_C | 2.10E-81 |
| D1PGT4 | PcopriDSM18205 | PF00756.23 | Esterase | 4.60E-21 |
| D1PGT4 | PcopriDSM18205 | PF03629.21 | SASA | 5.10E-31 |
| D1PH04 | PcopriDSM18205 | PF03572.21 | Peptidase_S41 | 2.00E-46 |
| D1PH08 | PcopriDSM18205 | PF03629.21 | SASA | 4.70E-22 |
| D1PH69 | PcopriDSM18205 | PF10459.12 | Peptidase_S46 | 2.40E-212 |
| D1PHB6 | PcopriDSM18205 | PF00326.24 | Peptidase_S9 | 2.10E-09 |
| D1PHG0 | PcopriDSM18205 | PF07859.16 | Abhydrolase_3 | 1.60E-36 |
| D1PHG0 | PcopriDSM18205 | PF13354.9 | Beta-lactamase2 | 2.00E-31 |
| D1PHI4 | PcopriDSM18205 | PF02113.18 | Peptidase_S13 | 1.40E-43 |
| B2PU03 | PstuartiiATCC25827 | PF06500.14 | FrsA-like | 5.70E-164 |
| B2PUJ8 | PstuartiiATCC25827 | PF01694.25 | Rhomboid | 2.60E-25 |
| B2PUM5 | PstuartiiATCC25827 | PF00144.27 | Beta-lactamase | 1.10E-71 |
| B2PUS0 | PstuartiiATCC25827 | PF00144.27 | Beta-lactamase | 1.10E-45 |
| B2PV17 | PstuartiiATCC25827 | PF01804.21 | Penicil_amidase | 7.70E-139 |
| B2PV18 | PstuartiiATCC25827 | PF01694.25 | Rhomboid | 9.10E-20 |
| B2PV27 | PstuartiiATCC25827 | PF00657.25 | Lipase_GDSL | 4.60E-23 |
| B2PVR9 | PstuartiiATCC25827 | PF03575.20 | Peptidase_S51 | 1.40E-42 |
| B2PWR3 | PstuartiiATCC25827 | PF01804.21 | Penicil_amidase | 8.20E-170 |
| B2PXI9 | PstuartiiATCC25827 | PF13365.9 | Trypsin_2 | 2.10E-29 |
| B2PXJ0 | PstuartiiATCC25827 | PF13365.9 | Trypsin_2 | 1.20E-29 |
| B2PXL6 | PstuartiiATCC25827 | PF06821.16 | Ser_hydrolase | 5.80E-27 |
| B2PXQ4 | PstuartiiATCC25827 | PF01764.28 | Lipase_3 | 8.60E-23 |
| B2PY44 | PstuartiiATCC25827 | PF00144.27 | Beta-lactamase | 1.10E-39 |
| B2PY66 | PstuartiiATCC25827 | PF00326.24 | Peptidase_S9 | 5.00E-65 |
| B2PZE4 | PstuartiiATCC25827 | PF01804.21 | Penicil_amidase | 1.20E-117 |
| B2PZE6 | PstuartiiATCC25827 | PF02450.18 | LCAT | 4.50E-08 |
| B2PZG0 | PstuartiiATCC25827 | PF00698.24 | Acyl_transf_1 | 1.00E-38 |
| B2PZK2 | PstuartiiATCC25827 | PF12697.10 | Abhydrolase_6 | 1.60E-19 |
| B2Q016 | PstuartiiATCC25827 | PF00717.26 | Peptidase_S24 | 7.00E-21 |
| B2Q0E0 | PstuartiiATCC25827 | PF02674.19 | Colicin_V | 1.30E-38 |
| B2Q0R1 | PstuartiiATCC25827 | PF01764.28 | Lipase_3 | 8.80E-24 |
| B2Q0U0 | PstuartiiATCC25827 | PF12146.11 | Hydrolase_4 | 2.50E-62 |
| B2Q163 | PstuartiiATCC25827 | PF10502.12 | Peptidase_S26 | 3.00E-56 |
| B2Q1N5 | PstuartiiATCC25827 | PF05929.14 | Phage_GPO | 1.90E-61 |
| B2Q208 | PstuartiiATCC25827 | PF05362.16 | Lon_C | 5.30E-10 |
| B2Q236 | PstuartiiATCC25827 | PF08840.14 | BAAT_C | 5.50E-06 |
| B2Q241 | PstuartiiATCC25827 | PF00756.23 | Esterase | 4.90E-61 |

| | | | | |
|---|---|---|---|---|
| B2Q2B8 | PstuartiiATCC25827 | PF12697.10 | Abhydrolase_6 | 1.40E-06 |
| B2Q2H1 | PstuartiiATCC25827 | PF00768.23 | Peptidase_S11 | 1.10E-102 |
| B2Q2K6 | PstuartiiATCC25827 | PF02450.18 | LCAT | 2.30E-14 |
| B2Q3F8 | PstuartiiATCC25827 | PF00717.26 | Peptidase_S24 | 5.60E-17 |
| B2Q3M0 | PstuartiiATCC25827 | PF01343.21 | Peptidase_S49 | 3.20E-45 |
| B2Q3T9 | PstuartiiATCC25827 | PF13472.9 | Lipase_GDSL_2 | 2.20E-22 |
| B2Q3W8 | PstuartiiATCC25827 | PF01694.25 | Rhomboid | 4.30E-25 |
| B2Q410 | PstuartiiATCC25827 | PF00768.23 | Peptidase_S11 | 1.50E-101 |
| B2Q4B3 | PstuartiiATCC25827 | PF01343.21 | Peptidase_S49 | 2.20E-48 |
| B2Q4D8 | PstuartiiATCC25827 | PF03572.21 | Peptidase_S41 | 3.60E-52 |
| B2Q4P6 | PstuartiiATCC25827 | PF12146.11 | Hydrolase_4 | 6.00E-47 |
| B2Q4V5 | PstuartiiATCC25827 | PF02113.18 | Peptidase_S13 | 3.60E-153 |
| B2Q538 | PstuartiiATCC25827 | PF00144.27 | Beta-lactamase | 8.90E-58 |
| B2Q563 | PstuartiiATCC25827 | PF00717.26 | Peptidase_S24 | 1.50E-32 |
| B2Q586 | PstuartiiATCC25827 | PF13472.9 | Lipase_GDSL_2 | 1.60E-12 |
| B2Q587 | PstuartiiATCC25827 | PF13472.9 | Lipase_GDSL_2 | 4.60E-09 |
| B2Q5M2 | PstuartiiATCC25827 | PF00756.23 | Esterase | 2.80E-20 |
| B2Q6B8 | PstuartiiATCC25827 | PF00717.26 | Peptidase_S24 | 7.80E-22 |
| B2Q6J1 | PstuartiiATCC25827 | PF00756.23 | Esterase | 2.80E-22 |
| B2Q6L1 | PstuartiiATCC25827 | PF01734.25 | Patatin | 4.90E-19 |
| B2Q703 | PstuartiiATCC25827 | PF00574.26 | CLP_protease | 2.50E-89 |
| B2Q705 | PstuartiiATCC25827 | PF05362.16 | Lon_C | 1.00E-96 |
| B3C536 | BintestinalisDSM17393 | PF00574.26 | CLP_protease | 8.20E-77 |
| B3C594 | BintestinalisDSM17393 | PF00756.23 | Esterase | 5.80E-23 |
| B3C596 | BintestinalisDSM17393 | PF00082.25 | Peptidase_S8 | 5.80E-36 |
| B3C5Z4 | BintestinalisDSM17393 | PF13472.9 | Lipase_GDSL_2 | 5.50E-15 |
| B3C638 | BintestinalisDSM17393 | PF03629.21 | SASA | 1.30E-20 |
| B3C639 | BintestinalisDSM17393 | PF03629.21 | SASA | 6.50E-17 |
| B3C6E9 | BintestinalisDSM17393 | PF01734.25 | Patatin | 1.10E-23 |
| B3C6G4 | BintestinalisDSM17393 | PF10502.12 | Peptidase_S26 | 1.60E-32 |
| B3C6H5 | BintestinalisDSM17393 | PF03572.21 | Peptidase_S41 | 2.40E-19 |
| B3C6J1 | BintestinalisDSM17393 | PF00326.24 | Peptidase_S9 | 1.60E-45 |
| B3C6M6 | BintestinalisDSM17393 | PF00144.27 | Beta-lactamase | 9.30E-46 |
| B3C6M7 | BintestinalisDSM17393 | PF00144.27 | Beta-lactamase | 4.10E-44 |
| B3C6V6 | BintestinalisDSM17393 | PF00326.24 | Peptidase_S9 | 1.10E-07 |
| B3C6V6 | BintestinalisDSM17393 | PF13472.9 | Lipase_GDSL_2 | 5.50E-09 |
| B3C6Z6 | BintestinalisDSM17393 | PF00144.27 | Beta-lactamase | 3.60E-29 |
| B3C759 | BintestinalisDSM17393 | PF00144.27 | Beta-lactamase | 4.00E-63 |
| B3C759 | BintestinalisDSM17393 | PF13472.9 | Lipase_GDSL_2 | 4.30E-16 |
| B3C767 | BintestinalisDSM17393 | PF03572.21 | Peptidase_S41 | 5.40E-33 |
| B3C7M6 | BintestinalisDSM17393 | PF13472.9 | Lipase_GDSL_2 | 9.00E-12 |
| B3C7P2 | BintestinalisDSM17393 | PF05448.15 | AXE1 | 9.20E-41 |
| B3C7S3 | BintestinalisDSM17393 | PF01694.25 | Rhomboid | 7.40E-25 |
| B3C7S4 | BintestinalisDSM17393 | PF01694.25 | Rhomboid | 1.70E-33 |
| B3C7T8 | BintestinalisDSM17393 | PF03572.21 | Peptidase_S41 | 7.40E-52 |
| B3C7Y7 | BintestinalisDSM17393 | PF02016.18 | Peptidase_S66 | 2.10E-35 |
| B3C818 | BintestinalisDSM17393 | PF00326.24 | Peptidase_S9 | 5.80E-07 |
| B3C818 | BintestinalisDSM17393 | PF13472.9 | Lipase_GDSL_2 | 2.60E-29 |
| B3C828 | BintestinalisDSM17393 | PF00144.27 | Beta-lactamase | 1.70E-31 |
| B3C8B7 | BintestinalisDSM17393 | PF13472.9 | Lipase_GDSL_2 | 3.00E-13 |
| B3C8J8 | BintestinalisDSM17393 | PF05576.14 | Peptidase_S37 | 3.30E-57 |
| B3C8T8 | BintestinalisDSM17393 | PF01734.25 | Patatin | 2.90E-29 |

| | | | | |
|---|---|---|---|---|
| B3C900 | BintestinalisDSM17393 | PF03572.21 | Peptidase_S41 | 8.20E-54 |
| B3C934 | BintestinalisDSM17393 | PF00717.26 | Peptidase_S24 | 7.60E-17 |
| B3C948 | BintestinalisDSM17393 | PF00326.24 | Peptidase_S9 | 1.60E-44 |
| B3C969 | BintestinalisDSM17393 | PF00756.23 | Esterase | 1.90E-19 |
| B3C974 | BintestinalisDSM17393 | PF00756.23 | Esterase | 6.70E-20 |
| B3C974 | BintestinalisDSM17393 | PF03629.21 | SASA | 1.40E-35 |
| B3C975 | BintestinalisDSM17393 | PF00756.23 | Esterase | 4.40E-21 |
| B3C9A2 | BintestinalisDSM17393 | PF02113.18 | Peptidase_S13 | 3.20E-80 |
| B3C9D0 | BintestinalisDSM17393 | PF00756.23 | Esterase | 1.80E-24 |
| B3C9E5 | BintestinalisDSM17393 | PF10502.12 | Peptidase_S26 | 3.20E-18 |
| B3C9J5 | BintestinalisDSM17393 | PF13365.9 | Trypsin_2 | 2.20E-26 |
| B3C9M4 | BintestinalisDSM17393 | PF10502.12 | Peptidase_S26 | 3.50E-29 |
| B3C9W9 | BintestinalisDSM17393 | PF03629.21 | SASA | 9.60E-10 |
| B3C9Y2 | BintestinalisDSM17393 | PF10459.12 | Peptidase_S46 | 2.60E-258 |
| B3CA27 | BintestinalisDSM17393 | PF10502.12 | Peptidase_S26 | 4.00E-17 |
| B3CA28 | BintestinalisDSM17393 | PF10502.12 | Peptidase_S26 | 2.70E-28 |
| B3CAB3 | BintestinalisDSM17393 | PF02674.19 | Colicin_V | 3.80E-31 |
| B3CAD2 | BintestinalisDSM17393 | PF03629.21 | SASA | 3.20E-55 |
| B3CB37 | BintestinalisDSM17393 | PF13365.9 | Trypsin_2 | 1.20E-33 |
| B3CB90 | BintestinalisDSM17393 | PF12146.11 | Hydrolase_4 | 9.10E-14 |
| B3CB92 | BintestinalisDSM17393 | PF00756.23 | Esterase | 2.70E-15 |
| B3CBJ7 | BintestinalisDSM17393 | PF00144.27 | Beta-lactamase | 2.60E-39 |
| B3CBQ6 | BintestinalisDSM17393 | PF13354.9 | Beta-lactamase2 | 1.40E-35 |
| B3CC11 | BintestinalisDSM17393 | PF05362.16 | Lon_C | 5.30E-82 |
| B3CCP2 | BintestinalisDSM17393 | PF00698.24 | Acyl_transf_1 | 1.60E-36 |
| B3CCS4 | BintestinalisDSM17393 | PF01734.25 | Patatin | 4.80E-21 |
| B3CE43 | BintestinalisDSM17393 | PF12697.10 | Abhydrolase_6 | 3.20E-10 |
| B3CEI1 | BintestinalisDSM17393 | PF00756.23 | Esterase | 3.80E-05 |
| B3CEK6 | BintestinalisDSM17393 | PF00326.24 | Peptidase_S9 | 8.50E-56 |
| B3CER9 | BintestinalisDSM17393 | PF03629.21 | SASA | 2.40E-14 |
| B3CET0 | BintestinalisDSM17393 | PF03629.21 | SASA | 1.70E-34 |
| B3CET1 | BintestinalisDSM17393 | PF00756.23 | Esterase | 3.10E-29 |
| B3CEU1 | BintestinalisDSM17393 | PF03629.21 | SASA | 1.30E-12 |
| B3CF45 | BintestinalisDSM17393 | PF03572.21 | Peptidase_S41 | 7.20E-20 |
| B3CF93 | BintestinalisDSM17393 | PF01694.25 | Rhomboid | 1.50E-12 |
| B3CF98 | BintestinalisDSM17393 | PF13472.9 | Lipase_GDSL_2 | 1.30E-11 |
| B3CFA3 | BintestinalisDSM17393 | PF13472.9 | Lipase_GDSL_2 | 2.90E-13 |
| B3CFC8 | BintestinalisDSM17393 | PF03572.21 | Peptidase_S41 | 3.30E-52 |
| B3CFP4 | BintestinalisDSM17393 | PF13472.9 | Lipase_GDSL_2 | 2.00E-15 |
| B3CFV2 | BintestinalisDSM17393 | PF00326.24 | Peptidase_S9 | 4.20E-10 |
| B3CFY2 | BintestinalisDSM17393 | PF12146.11 | Hydrolase_4 | 1.80E-11 |
| B3CG11 | BintestinalisDSM17393 | PF00326.24 | Peptidase_S9 | 1.40E-58 |
| B3CG29 | BintestinalisDSM17393 | PF05448.15 | AXE1 | 2.40E-46 |
| B3CG30 | BintestinalisDSM17393 | PF13472.9 | Lipase_GDSL_2 | 6.20E-14 |
| B3CG40 | BintestinalisDSM17393 | PF00717.26 | Peptidase_S24 | 1.10E-18 |
| B3CGC9 | BintestinalisDSM17393 | PF03629.21 | SASA | 3.40E-21 |
| B3CGS2 | BintestinalisDSM17393 | PF10502.12 | Peptidase_S26 | 6.70E-31 |
| B3CGT4 | BintestinalisDSM17393 | PF10459.12 | Peptidase_S46 | 3.50E-263 |
| B3CGT5 | BintestinalisDSM17393 | PF10459.12 | Peptidase_S46 | 1.00E-260 |
| B3CGY5 | BintestinalisDSM17393 | PF03629.21 | SASA | 4.20E-49 |
| B3CH21 | BintestinalisDSM17393 | PF03572.21 | Peptidase_S41 | 5.80E-19 |
| B3CHD9 | BintestinalisDSM17393 | PF03572.21 | Peptidase_S41 | 1.40E-19 |

| | | | | |
|---|---|---|---|---|
| B3CHE1 | BintestinalisDSM17393 | PF00756.23 | Esterase | 4.30E-28 |
| B3CHU5 | BintestinalisDSM17393 | PF03629.21 | SASA | 7.80E-19 |
| B3CHU8 | BintestinalisDSM17393 | PF03629.21 | SASA | 4.00E-21 |
| B3CHW3 | BintestinalisDSM17393 | PF13472.9 | Lipase_GDSL_2 | 2.60E-13 |
| B3CI38 | BintestinalisDSM17393 | PF00756.23 | Esterase | 1.30E-28 |
| B3CIL7 | BintestinalisDSM17393 | PF01734.25 | Patatin | 2.70E-21 |
| B3CIT2 | BintestinalisDSM17393 | PF01343.21 | Peptidase_S49 | 8.20E-51 |
| B3CIY1 | BintestinalisDSM17393 | PF00326.24 | Peptidase_S9 | 1.80E-67 |
| A0A0K9CLL2 | FanimalisD11 | PF01734.25 | Patatin | 5.20E-16 |
| A0A0K9CMT | FanimalisD11 | PF01734.25 | Patatin | 1.30E-09 |
| A0A0K9CP24 | FanimalisD11 | PF02674.19 | Colicin_V | 1.30E-17 |
| A0A0K9CP82 | FanimalisD11 | PF12146.11 | Hydrolase_4 | 1.30E-50 |
| D6BCT4 | FanimalisD11 | PF01343.21 | Peptidase_S49 | 7.60E-39 |
| D6BD79 | FanimalisD11 | PF00326.24 | Peptidase_S9 | 8.10E-56 |
| D6BD96 | FanimalisD11 | PF03575.20 | Peptidase_S51 | 1.20E-51 |
| D6BDC7 | FanimalisD11 | PF07819.16 | PGAP1 | 2.90E-06 |
| D6BDF5 | FanimalisD11 | PF00082.25 | Peptidase_S8 | 3.50E-18 |
| D6BDS5 | FanimalisD11 | PF01734.25 | Patatin | 2.60E-14 |
| D6BE02 | FanimalisD11 | PF00574.26 | CLP_protease | 1.20E-28 |
| D6BE69 | FanimalisD11 | PF10502.12 | Peptidase_S26 | 9.10E-20 |
| D6BEK9 | FanimalisD11 | PF00698.24 | Acyl_transf_1 | 6.40E-44 |
| D6BEL6 | FanimalisD11 | PF05362.16 | Lon_C | 1.30E-08 |
| D6BG23 | FanimalisD11 | PF00082.25 | Peptidase_S8 | 5.40E-23 |
| D6BG51 | FanimalisD11 | PF00768.23 | Peptidase_S11 | 4.50E-44 |
| D6BGC0 | FanimalisD11 | PF01343.21 | Peptidase_S49 | 1.90E-40 |
| D6BGK9 | FanimalisD11 | PF01425.24 | Amidase | 1.20E-154 |
| D6BGY9 | FanimalisD11 | PF03572.21 | Peptidase_S41 | 3.40E-56 |
| D6BHN0 | FanimalisD11 | PF13354.9 | Beta-lactamase2 | 3.80E-58 |
| D6BHN3 | FanimalisD11 | PF00717.26 | Peptidase_S24 | 7.40E-27 |
| D6BIV3 | FanimalisD11 | PF00574.26 | CLP_protease | 1.30E-84 |
| D6BIV5 | FanimalisD11 | PF05362.16 | Lon_C | 4.90E-80 |
| D6BJ99 | FanimalisD11 | PF10502.12 | Peptidase_S26 | 9.00E-30 |
| B0NKS9 | BstercorisATCC43183 | PF01343.21 | Peptidase_S49 | 5.40E-21 |
| B0NKW3 | BstercorisATCC43183 | PF00326.24 | Peptidase_S9 | 3.50E-43 |
| B0NL30 | BstercorisATCC43183 | PF00144.27 | Beta-lactamase | 7.40E-52 |
| B0NLQ2 | BstercorisATCC43183 | PF03629.21 | SASA | 1.20E-18 |
| B0NM79 | BstercorisATCC43183 | PF04586.20 | Peptidase_S78 | 3.00E-26 |
| B0NMM2 | BstercorisATCC43183 | PF13365.9 | Trypsin_2 | 2.50E-34 |
| B0NPA5 | BstercorisATCC43183 | PF00326.24 | Peptidase_S9 | 1.10E-55 |
| B0NPF1 | BstercorisATCC43183 | PF01734.25 | Patatin | 8.90E-22 |
| B0NPG9 | BstercorisATCC43183 | PF00698.24 | Acyl_transf_1 | 2.00E-37 |
| B0NPL3 | BstercorisATCC43183 | PF05362.16 | Lon_C | 6.70E-82 |
| B0NPN2 | BstercorisATCC43183 | PF01694.25 | Rhomboid | 2.00E-30 |
| B0NPP4 | BstercorisATCC43183 | PF13354.9 | Beta-lactamase2 | 8.50E-36 |
| B0NPR8 | BstercorisATCC43183 | PF01734.25 | Patatin | 1.20E-28 |
| B0NPW7 | BstercorisATCC43183 | PF03629.21 | SASA | 8.50E-21 |
| B0NQC8 | BstercorisATCC43183 | PF01734.25 | Patatin | 2.10E-20 |
| B0NQQ7 | BstercorisATCC43183 | PF00756.23 | Esterase | 1.50E-28 |
| B0NQQ9 | BstercorisATCC43183 | PF03572.21 | Peptidase_S41 | 4.10E-18 |
| B0NQY3 | BstercorisATCC43183 | PF10459.12 | Peptidase_S46 | 3.10E-257 |
| B0NQY4 | BstercorisATCC43183 | PF10459.12 | Peptidase_S46 | 3.00E-258 |
| B0NRI5 | BstercorisATCC43183 | PF02016.18 | Peptidase_S66 | 6.60E-36 |

| | | | | |
|---|---|---|---|---|
| B0NRK7 | BstercorisATCC43183 | PF03572.21 | Peptidase_S41 | 7.60E-51 |
| B0NRM9 | BstercorisATCC43183 | PF01694.25 | Rhomboid | 5.10E-33 |
| B0NRN0 | BstercorisATCC43183 | PF01694.25 | Rhomboid | 1.50E-23 |
| B0NS26 | BstercorisATCC43183 | PF03572.21 | Peptidase_S41 | 3.10E-53 |
| B0NS50 | BstercorisATCC43183 | PF00326.24 | Peptidase_S9 | 2.20E-45 |
| B0NS93 | BstercorisATCC43183 | PF02113.18 | Peptidase_S13 | 1.30E-79 |
| B0NSC7 | BstercorisATCC43183 | PF13365.9 | Trypsin_2 | 9.60E-26 |
| B0NSG5 | BstercorisATCC43183 | PF10459.12 | Peptidase_S46 | 1.40E-261 |
| B0NSI2 | BstercorisATCC43183 | PF10502.12 | Peptidase_S26 | 4.40E-09 |
| B0NSI3 | BstercorisATCC43183 | PF10502.12 | Peptidase_S26 | 1.40E-26 |
| B0NSP4 | BstercorisATCC43183 | PF02674.19 | Colicin_V | 8.00E-30 |
| B0NSY1 | BstercorisATCC43183 | PF03572.21 | Peptidase_S41 | 4.60E-34 |
| B0NSY7 | BstercorisATCC43183 | PF00144.27 | Beta-lactamase | 2.60E-62 |
| B0NSY7 | BstercorisATCC43183 | PF13472.9 | Lipase_GDSL_2 | 4.10E-17 |
| B0NT27 | BstercorisATCC43183 | PF02129.21 | Peptidase_S15 | 9.50E-44 |
| B0NT93 | BstercorisATCC43183 | PF13472.9 | Lipase_GDSL_2 | 2.00E-14 |
| B0NTN3 | BstercorisATCC43183 | PF05576.14 | Peptidase_S37 | 2.10E-61 |
| B0NTT7 | BstercorisATCC43183 | PF01343.21 | Peptidase_S49 | 8.00E-50 |
| B0NU47 | BstercorisATCC43183 | PF00082.25 | Peptidase_S8 | 4.90E-33 |
| B0NU93 | BstercorisATCC43183 | PF12146.11 | Hydrolase_4 | 1.10E-15 |
| B0NUT4 | BstercorisATCC43183 | PF00326.24 | Peptidase_S9 | 1.30E-57 |
| B0NUW3 | BstercorisATCC43183 | PF05448.15 | AXE1 | 4.60E-50 |
| B0NUW4 | BstercorisATCC43183 | PF13472.9 | Lipase_GDSL_2 | 3.50E-12 |
| B0NV56 | BstercorisATCC43183 | PF03572.21 | Peptidase_S41 | 2.30E-18 |
| B0NVP5 | BstercorisATCC43183 | PF03572.21 | Peptidase_S41 | 1.40E-51 |
| B0NVS2 | BstercorisATCC43183 | PF03959.16 | FSH1 | 9.40E-06 |
| B0NW45 | BstercorisATCC43183 | PF00082.25 | Peptidase_S8 | 6.70E-41 |
| B0NW66 | BstercorisATCC43183 | PF00082.25 | Peptidase_S8 | 4.60E-34 |
| B0NWE3 | BstercorisATCC43183 | PF00574.26 | CLP_protease | 3.40E-77 |
| C7G5A1 | RintestinalisL182 | PF03572.21 | Peptidase_S41 | 4.90E-55 |
| C7G5F7 | RintestinalisL182 | PF13472.9 | Lipase_GDSL_2 | 1.20E-10 |
| C7G5L1 | RintestinalisL182 | PF00698.24 | Acyl_transf_1 | 2.30E-34 |
| C7G5Q7 | RintestinalisL182 | PF01734.25 | Patatin | 3.80E-16 |
| C7G5R0 | RintestinalisL182 | PF02016.18 | Peptidase_S66 | 5.50E-20 |
| C7G5S3 | RintestinalisL182 | PF01694.25 | Rhomboid | 4.30E-30 |
| C7G624 | RintestinalisL182 | PF00574.26 | CLP_protease | 4.10E-17 |
| C7G659 | RintestinalisL182 | PF00082.25 | Peptidase_S8 | 5.10E-13 |
| C7G6E7 | RintestinalisL182 | PF01734.25 | Patatin | 2.20E-19 |
| C7G6F8 | RintestinalisL182 | PF13472.9 | Lipase_GDSL_2 | 3.20E-13 |
| C7G6H9 | RintestinalisL182 | PF12146.11 | Hydrolase_4 | 8.80E-45 |
| C7G715 | RintestinalisL182 | PF13472.9 | Lipase_GDSL_2 | 1.70E-10 |
| C7G770 | RintestinalisL182 | PF13365.9 | Trypsin_2 | 2.20E-28 |
| C7G7L2 | RintestinalisL182 | PF00144.27 | Beta-lactamase | 5.80E-30 |
| C7G7L7 | RintestinalisL182 | PF13472.9 | Lipase_GDSL_2 | 4.90E-10 |
| C7G7T7 | RintestinalisL182 | PF00717.26 | Peptidase_S24 | 2.90E-33 |
| C7G889 | RintestinalisL182 | PF01694.25 | Rhomboid | 1.10E-21 |
| C7G8T8 | RintestinalisL182 | PF07859.16 | Abhydrolase_3 | 4.10E-49 |
| C7G8W6 | RintestinalisL182 | PF00756.23 | Esterase | 2.60E-18 |
| C7G903 | RintestinalisL182 | PF10503.12 | Esterase_PHB | 4.30E-13 |
| C7G908 | RintestinalisL182 | PF00756.23 | Esterase | 3.60E-16 |
| C7G911 | RintestinalisL182 | PF00756.23 | Esterase | 2.10E-12 |
| C7G912 | RintestinalisL182 | PF00756.23 | Esterase | 3.90E-21 |

| | | | | |
|---|---|---|---|---|
| C7G966 | RintestinalisL182 | PF00768.23 | Peptidase_S11 | 2.20E-55 |
| C7G993 | RintestinalisL182 | PF01425.24 | Amidase | 1.40E-67 |
| C7GA96 | RintestinalisL182 | PF00144.27 | Beta-lactamase | 3.70E-32 |
| C7GAT3 | RintestinalisL182 | PF13472.9 | Lipase_GDSL_2 | 1.90E-21 |
| C7GAV2 | RintestinalisL182 | PF12146.11 | Hydrolase_4 | 7.80E-45 |
| C7GB00 | RintestinalisL182 | PF10502.12 | Peptidase_S26 | 1.50E-40 |
| C7GB12 | RintestinalisL182 | PF12146.11 | Hydrolase_4 | 1.20E-06 |
| C7GBZ1 | RintestinalisL182 | PF13884.9 | Peptidase_S74 | 2.20E-15 |
| C7GBZ4 | RintestinalisL182 | PF13884.9 | Peptidase_S74 | 1.70E-08 |
| C7GCD7 | RintestinalisL182 | PF05580.15 | Peptidase_S55 | 2.70E-80 |
| C7GCE9 | RintestinalisL182 | PF13472.9 | Lipase_GDSL_2 | 1.30E-15 |
| C7GCL5 | RintestinalisL182 | PF12697.10 | Abhydrolase_6 | 1.80E-08 |
| C7GCT1 | RintestinalisL182 | PF03959.16 | FSH1 | 3.90E-06 |
| C7GDJ0 | RintestinalisL182 | PF00768.23 | Peptidase_S11 | 3.50E-48 |
| C7GDJ3 | RintestinalisL182 | PF00768.23 | Peptidase_S11 | 7.50E-74 |
| C7GDJ5 | RintestinalisL182 | PF00082.25 | Peptidase_S8 | 2.20E-48 |
| C7GDS4 | RintestinalisL182 | PF13472.9 | Lipase_GDSL_2 | 2.40E-11 |
| C7GDZ9 | RintestinalisL182 | PF05362.16 | Lon_C | 1.50E-93 |
| C7GE01 | RintestinalisL182 | PF00574.26 | CLP_protease | 4.60E-85 |
| C7GE42 | RintestinalisL182 | PF13472.9 | Lipase_GDSL_2 | 3.90E-13 |
| C7GEA5 | RintestinalisL182 | PF00768.23 | Peptidase_S11 | 1.70E-49 |
| C7GEG1 | RintestinalisL182 | PF00082.25 | Peptidase_S8 | 1.50E-21 |
| C7GEK6 | RintestinalisL182 | PF12146.11 | Hydrolase_4 | 3.90E-12 |
| C7GF27 | RintestinalisL182 | PF00144.27 | Beta-lactamase | 4.30E-56 |
| C7GF76 | RintestinalisL182 | PF12740.10 | Chlorophyllase2 | 6.60E-09 |
| C7GF99 | RintestinalisL182 | PF02674.19 | Colicin_V | 3.50E-12 |
| C7GFQ1 | RintestinalisL182 | PF01425.24 | Amidase | 6.50E-161 |
| C7GG56 | RintestinalisL182 | PF13365.9 | Trypsin_2 | 4.10E-11 |
| C7GG63 | RintestinalisL182 | PF00144.27 | Beta-lactamase | 1.30E-33 |
| C7GGG1 | RintestinalisL182 | PF10502.12 | Peptidase_S26 | 8.00E-40 |
| C7GGG3 | RintestinalisL182 | PF10502.12 | Peptidase_S26 | 1.20E-40 |
| C7GH70 | RintestinalisL182 | PF00574.26 | CLP_protease | 7.80E-24 |
| C7GHL5 | RintestinalisL182 | PF00082.25 | Peptidase_S8 | 1.80E-12 |
| C7GHZ3 | RintestinalisL182 | PF00144.27 | Beta-lactamase | 4.70E-49 |
| C7GHZ4 | RintestinalisL182 | PF00144.27 | Beta-lactamase | 5.60E-39 |
| B6VRX5 | PdoreiDSM17855 | PF00574.26 | CLP_protease | 1.30E-76 |
| B6VSF9 | PdoreiDSM17855 | PF00082.25 | Peptidase_S8 | 4.60E-37 |
| B6VSH9 | PdoreiDSM17855 | PF13472.9 | Lipase_GDSL_2 | 5.60E-19 |
| B6VST6 | PdoreiDSM17855 | PF03572.21 | Peptidase_S41 | 6.10E-33 |
| B6VSW6 | PdoreiDSM17855 | PF13472.9 | Lipase_GDSL_2 | 2.60E-13 |
| B6VSZ4 | PdoreiDSM17855 | PF13472.9 | Lipase_GDSL_2 | 6.70E-12 |
| B6VT10 | PdoreiDSM17855 | PF05448.15 | AXE1 | 1.70E-42 |
| B6VT19 | PdoreiDSM17855 | PF03629.21 | SASA | 2.70E-19 |
| B6VT52 | PdoreiDSM17855 | PF02129.21 | Peptidase_S15 | 4.60E-11 |
| B6VT64 | PdoreiDSM17855 | PF00144.27 | Beta-lactamase | 2.10E-45 |
| B6VT82 | PdoreiDSM17855 | PF00756.23 | Esterase | 3.60E-18 |
| B6VTD5 | PdoreiDSM17855 | PF00144.27 | Beta-lactamase | 4.40E-48 |
| B6VTE7 | PdoreiDSM17855 | PF13472.9 | Lipase_GDSL_2 | 1.00E-18 |
| B6VTE8 | PdoreiDSM17855 | PF03629.21 | SASA | 3.10E-20 |
| B6VTE8 | PdoreiDSM17855 | PF13472.9 | Lipase_GDSL_2 | 2.10E-24 |
| B6VTR2 | PdoreiDSM17855 | PF10502.12 | Peptidase_S26 | 5.40E-31 |
| B6VTU8 | PdoreiDSM17855 | PF03629.21 | SASA | 1.20E-13 |

| | | | | |
|---|---|---|---|---|
| B6VTW8 | PdoreiDSM17855 | PF00326.24 | Peptidase_S9 | 1.90E-42 |
| B6VTX6 | PdoreiDSM17855 | PF03572.21 | Peptidase_S41 | 1.20E-16 |
| B6VU72 | PdoreiDSM17855 | PF01734.25 | Patatin | 7.00E-23 |
| B6VU73 | PdoreiDSM17855 | PF00698.24 | Acyl_transf_1 | 1.60E-38 |
| B6VUK3 | PdoreiDSM17855 | PF00326.24 | Peptidase_S9 | 4.90E-41 |
| B6VVB8 | PdoreiDSM17855 | PF13472.9 | Lipase_GDSL_2 | 7.80E-26 |
| B6VVK7 | PdoreiDSM17855 | PF03629.21 | SASA | 7.60E-12 |
| B6VVS2 | PdoreiDSM17855 | PF13472.9 | Lipase_GDSL_2 | 3.10E-12 |
| B6VW01 | PdoreiDSM17855 | PF02129.21 | Peptidase_S15 | 2.40E-11 |
| B6VWP0 | PdoreiDSM17855 | PF03629.21 | SASA | 2.90E-26 |
| B6VWX7 | PdoreiDSM17855 | PF13365.9 | Trypsin_2 | 1.70E-21 |
| B6VX11 | PdoreiDSM17855 | PF01694.25 | Rhomboid | 1.80E-25 |
| B6VX12 | PdoreiDSM17855 | PF01694.25 | Rhomboid | 3.50E-35 |
| B6VX93 | PdoreiDSM17855 | PF00326.24 | Peptidase_S9 | 7.60E-46 |
| B6VXB3 | PdoreiDSM17855 | PF02113.18 | Peptidase_S13 | 2.70E-92 |
| B6VXC2 | PdoreiDSM17855 | PF00326.24 | Peptidase_S9 | 3.30E-45 |
| B6VXH5 | PdoreiDSM17855 | PF03572.21 | Peptidase_S41 | 1.20E-19 |
| B6VXL4 | PdoreiDSM17855 | PF03572.21 | Peptidase_S41 | 6.70E-17 |
| B6VXM4 | PdoreiDSM17855 | PF10502.12 | Peptidase_S26 | 1.00E-22 |
| B6VXM5 | PdoreiDSM17855 | PF10502.12 | Peptidase_S26 | 1.10E-25 |
| B6VXV3 | PdoreiDSM17855 | PF03572.21 | Peptidase_S41 | 8.40E-52 |
| B6VXZ2 | PdoreiDSM17855 | PF10459.12 | Peptidase_S46 | 2.90E-265 |
| B6VY01 | PdoreiDSM17855 | PF00326.24 | Peptidase_S9 | 5.90E-51 |
| B6VY72 | PdoreiDSM17855 | PF05448.15 | AXE1 | 1.90E-49 |
| B6VY86 | PdoreiDSM17855 | PF02674.19 | Colicin_V | 5.90E-27 |
| B6VYB0 | PdoreiDSM17855 | PF00082.25 | Peptidase_S8 | 1.60E-47 |
| B6VYN8 | PdoreiDSM17855 | PF05576.14 | Peptidase_S37 | 1.10E-50 |
| B6VYT6 | PdoreiDSM17855 | PF01343.21 | Peptidase_S49 | 1.80E-43 |
| B6VZ50 | PdoreiDSM17855 | PF00082.25 | Peptidase_S8 | 5.60E-34 |
| B6VZC7 | PdoreiDSM17855 | PF00326.24 | Peptidase_S9 | 1.00E-52 |
| B6VZH2 | PdoreiDSM17855 | PF01734.25 | Patatin | 3.10E-27 |
| B6VZN3 | PdoreiDSM17855 | PF13354.9 | Beta-lactamase2 | 1.40E-33 |
| B6VZW8 | PdoreiDSM17855 | PF05362.16 | Lon_C | 3.80E-82 |
| B6W063 | PdoreiDSM17855 | PF01734.25 | Patatin | 3.60E-26 |
| B6W0K6 | PdoreiDSM17855 | PF00326.24 | Peptidase_S9 | 1.00E-54 |
| B6W0W6 | PdoreiDSM17855 | PF03572.21 | Peptidase_S41 | 3.20E-19 |
| B6W0Y3 | PdoreiDSM17855 | PF02129.21 | Peptidase_S15 | 1.10E-50 |
| B6W163 | PdoreiDSM17855 | PF00326.24 | Peptidase_S9 | 2.40E-36 |
| B6W166 | PdoreiDSM17855 | PF13472.9 | Lipase_GDSL_2 | 2.20E-13 |
| B6W170 | PdoreiDSM17855 | PF05448.15 | AXE1 | 2.10E-09 |
| B6W1I8 | PdoreiDSM17855 | PF10459.12 | Peptidase_S46 | 1.50E-270 |
| B6W1J1 | PdoreiDSM17855 | PF13472.9 | Lipase_GDSL_2 | 3.20E-18 |
| B6W1J2 | PdoreiDSM17855 | PF13472.9 | Lipase_GDSL_2 | 3.10E-14 |
| B6W1N2 | PdoreiDSM17855 | PF12146.11 | Hydrolase_4 | 7.10E-15 |
| B6W1N8 | PdoreiDSM17855 | PF10459.12 | Peptidase_S46 | 1.00E-269 |
| B6W1P8 | PdoreiDSM17855 | PF03572.21 | Peptidase_S41 | 2.20E-20 |
| B6W1U5 | PdoreiDSM17855 | PF01734.25 | Patatin | 2.90E-19 |
| B6W293 | PdoreiDSM17855 | PF03629.21 | SASA | 1.60E-20 |
| B6W293 | PdoreiDSM17855 | PF13472.9 | Lipase_GDSL_2 | 2.30E-13 |
| B6W2J6 | PdoreiDSM17855 | PF00756.23 | Esterase | 4.10E-30 |
| B6W2S3 | PdoreiDSM17855 | PF02129.21 | Peptidase_S15 | 2.70E-07 |
| B6W2U2 | PdoreiDSM17855 | PF12697.10 | Abhydrolase_6 | 2.30E-06 |

| | | | | |
|---|---|---|---|---|
| B6W2X3 | PdoreiDSM17855 | PF03629.21 | SASA | 6.00E-17 |
| B6W320 | PdoreiDSM17855 | PF03572.21 | Peptidase_S41 | 4.80E-52 |
| B6W3P8 | PdoreiDSM17855 | PF00326.24 | Peptidase_S9 | 1.90E-11 |
| B6W429 | PdoreiDSM17855 | PF13365.9 | Trypsin_2 | 1.80E-34 |
| B6W485 | PdoreiDSM17855 | PF01694.25 | Rhomboid | 1.30E-14 |
| B6W4G9 | PdoreiDSM17855 | PF00082.25 | Peptidase_S8 | 8.90E-52 |
| B6W4Q5 | PdoreiDSM17855 | PF13472.9 | Lipase_GDSL_2 | 2.20E-16 |
| B6W4Q6 | PdoreiDSM17855 | PF13472.9 | Lipase_GDSL_2 | 1.70E-16 |
| B6W4W7 | PdoreiDSM17855 | PF00756.23 | Esterase | 3.30E-34 |
| B6W4Z4 | PdoreiDSM17855 | PF00326.24 | Peptidase_S9 | 5.30E-25 |
| B6W5B6 | PdoreiDSM17855 | PF03572.21 | Peptidase_S41 | 5.80E-46 |
| B6W5E8 | PdoreiDSM17855 | PF00326.24 | Peptidase_S9 | 3.50E-32 |
| B6W5F6 | PdoreiDSM17855 | PF03629.21 | SASA | 6.60E-12 |
| D3A9C2 | HhathewayiDSM13479 | PF13354.9 | Beta-lactamase2 | 1.00E-63 |
| D3A9D1 | HhathewayiDSM13479 | PF13354.9 | Beta-lactamase2 | 3.10E-56 |
| D3A9I0 | HhathewayiDSM13479 | PF00082.25 | Peptidase_S8 | 1.40E-50 |
| D3A9J6 | HhathewayiDSM13479 | PF00135.31 | COesterase | 6.60E-65 |
| D3A9K1 | HhathewayiDSM13479 | PF00756.23 | Esterase | 1.80E-18 |
| D3A9K3 | HhathewayiDSM13479 | PF00756.23 | Esterase | 2.50E-27 |
| D3AA25 | HhathewayiDSM13479 | PF00756.23 | Esterase | 1.10E-26 |
| D3AA29 | HhathewayiDSM13479 | PF10502.12 | Peptidase_S26 | 3.70E-44 |
| D3AAC6 | HhathewayiDSM13479 | PF13365.9 | Trypsin_2 | 1.20E-07 |
| D3AAF7 | HhathewayiDSM13479 | PF07859.16 | Abhydrolase_3 | 3.50E-24 |
| D3AAS1 | HhathewayiDSM13479 | PF00135.31 | COesterase | 1.70E-70 |
| D3AAU0 | HhathewayiDSM13479 | PF07859.16 | Abhydrolase_3 | 3.60E-57 |
| D3AB41 | HhathewayiDSM13479 | PF00144.27 | Beta-lactamase | 2.00E-48 |
| D3AB92 | HhathewayiDSM13479 | PF02129.21 | Peptidase_S15 | 1.60E-56 |
| D3ABU1 | HhathewayiDSM13479 | PF12146.11 | Hydrolase_4 | 7.30E-17 |
| D3ABU7 | HhathewayiDSM13479 | PF03629.21 | SASA | 1.40E-23 |
| D3ACN9 | HhathewayiDSM13479 | PF13472.9 | Lipase_GDSL_2 | 2.80E-19 |
| D3ACP0 | HhathewayiDSM13479 | PF00082.25 | Peptidase_S8 | 1.10E-50 |
| D3AD37 | HhathewayiDSM13479 | PF12146.11 | Hydrolase_4 | 1.00E-49 |
| D3ADN2 | HhathewayiDSM13479 | PF00144.27 | Beta-lactamase | 5.50E-27 |
| D3AE08 | HhathewayiDSM13479 | PF07859.16 | Abhydrolase_3 | 8.90E-26 |
| D3AE49 | HhathewayiDSM13479 | PF13472.9 | Lipase_GDSL_2 | 2.10E-18 |
| D3AEG4 | HhathewayiDSM13479 | PF05580.15 | Peptidase_S55 | 3.00E-82 |
| D3AEJ3 | HhathewayiDSM13479 | PF00698.24 | Acyl_transf_1 | 1.10E-39 |
| D3AES2 | HhathewayiDSM13479 | PF12697.10 | Abhydrolase_6 | 6.40E-09 |
| D3AF45 | HhathewayiDSM13479 | PF01694.25 | Rhomboid | 6.20E-38 |
| D3AF81 | HhathewayiDSM13479 | PF00144.27 | Beta-lactamase | 1.80E-46 |
| D3AFJ0 | HhathewayiDSM13479 | PF00326.24 | Peptidase_S9 | 7.80E-06 |
| D3AGS0 | HhathewayiDSM13479 | PF00768.23 | Peptidase_S11 | 1.50E-57 |
| D3AH68 | HhathewayiDSM13479 | PF07859.16 | Abhydrolase_3 | 6.00E-51 |
| D3AH92 | HhathewayiDSM13479 | PF02674.19 | Colicin_V | 4.00E-12 |
| D3AHV0 | HhathewayiDSM13479 | PF00082.25 | Peptidase_S8 | 3.60E-11 |
| D3AHV3 | HhathewayiDSM13479 | PF13365.9 | Trypsin_2 | 2.10E-28 |
| D3AI05 | HhathewayiDSM13479 | PF13472.9 | Lipase_GDSL_2 | 5.00E-18 |
| D3AIN4 | HhathewayiDSM13479 | PF01972.19 | SDH_sah | 1.50E-18 |
| D3AIY2 | HhathewayiDSM13479 | PF12697.10 | Abhydrolase_6 | 3.40E-07 |
| D3AJE4 | HhathewayiDSM13479 | PF01425.24 | Amidase | 5.30E-154 |
| D3AJW1 | HhathewayiDSM13479 | PF00768.23 | Peptidase_S11 | 3.80E-55 |
| D3AK71 | HhathewayiDSM13479 | PF12146.11 | Hydrolase_4 | 3.00E-13 |

| | | | | |
|---|---|---|---|---|
| D3AKX6 | HhathewayiDSM13479 | PF12146.11 | Hydrolase_4 | 2.10E-43 |
| D3AL20 | HhathewayiDSM13479 | PF00768.23 | Peptidase_S11 | 2.10E-47 |
| D3AL31 | HhathewayiDSM13479 | PF00135.31 | COesterase | 8.00E-78 |
| D3ALB7 | HhathewayiDSM13479 | PF03572.21 | Peptidase_S41 | 2.00E-49 |
| D3ALK8 | HhathewayiDSM13479 | PF00768.23 | Peptidase_S11 | 6.60E-13 |
| D3ALK9 | HhathewayiDSM13479 | PF00768.23 | Peptidase_S11 | 2.60E-12 |
| D3ALS5 | HhathewayiDSM13479 | PF04586.20 | Peptidase_S78 | 3.20E-24 |
| D3AM11 | HhathewayiDSM13479 | PF03629.21 | SASA | 4.80E-12 |
| D3AMN3 | HhathewayiDSM13479 | PF00574.26 | CLP_protease | 2.50E-79 |
| D3ANG2 | HhathewayiDSM13479 | PF00082.25 | Peptidase_S8 | 4.50E-15 |
| D3ANK4 | HhathewayiDSM13479 | PF00768.23 | Peptidase_S11 | 2.50E-24 |
| D3ANV5 | HhathewayiDSM13479 | PF00082.25 | Peptidase_S8 | 2.50E-17 |
| D3AP13 | HhathewayiDSM13479 | PF00144.27 | Beta-lactamase | 9.50E-30 |
| D3AP25 | HhathewayiDSM13479 | PF12146.11 | Hydrolase_4 | 3.10E-15 |
| D3APJ2 | HhathewayiDSM13479 | PF00768.23 | Peptidase_S11 | 7.40E-57 |
| D3APV1 | HhathewayiDSM13479 | PF01734.25 | Patatin | 1.50E-19 |
| D3APY8 | HhathewayiDSM13479 | PF07859.16 | Abhydrolase_3 | 1.90E-45 |
| D3AQ56 | HhathewayiDSM13479 | PF10502.12 | Peptidase_S26 | 1.10E-24 |
| D3AQ57 | HhathewayiDSM13479 | PF10502.12 | Peptidase_S26 | 1.30E-10 |
| D3AQ81 | HhathewayiDSM13479 | PF00574.26 | CLP_protease | 2.00E-87 |
| D3AQ83 | HhathewayiDSM13479 | PF05362.16 | Lon_C | 7.60E-88 |
| D3ARC4 | HhathewayiDSM13479 | PF00768.23 | Peptidase_S11 | 6.70E-72 |
| D3AS06 | HhathewayiDSM13479 | PF13472.9 | Lipase_GDSL_2 | 1.00E-14 |
| D3ASI3 | HhathewayiDSM13479 | PF07859.16 | Abhydrolase_3 | 9.20E-21 |
| D3ATT1 | HhathewayiDSM13479 | PF12146.11 | Hydrolase_4 | 1.70E-08 |
| D3ATT2 | HhathewayiDSM13479 | PF13472.9 | Lipase_GDSL_2 | 1.40E-13 |
| D3ATW1 | HhathewayiDSM13479 | PF00717.26 | Peptidase_S24 | 1.90E-30 |
| D3AU11 | HhathewayiDSM13479 | PF12697.10 | Abhydrolase_6 | 4.20E-08 |
| D3AU19 | HhathewayiDSM13479 | PF00135.31 | COesterase | 4.60E-09 |
| D3AU79 | HhathewayiDSM13479 | PF00135.31 | COesterase | 2.70E-73 |
| D3AUG4 | HhathewayiDSM13479 | PF00768.23 | Peptidase_S11 | 1.30E-23 |
| D3AUK8 | HhathewayiDSM13479 | PF10502.12 | Peptidase_S26 | 1.90E-34 |
| D3AUN6 | HhathewayiDSM13479 | PF00756.23 | Esterase | 1.00E-05 |
| D3AUZ9 | HhathewayiDSM13479 | PF13472.9 | Lipase_GDSL_2 | 2.50E-13 |
| D3AVC2 | HhathewayiDSM13479 | PF01734.25 | Patatin | 3.50E-08 |
| D3AVC8 | HhathewayiDSM13479 | PF00574.26 | CLP_protease | 3.50E-11 |
| I3TYY8 | EfaeciumATCCBAA472 | PF00698.24 | Acyl_transf_1 | 1.60E-18 |
| I3U046 | EfaeciumATCCBAA472 | PF10502.12 | Peptidase_S26 | 6.20E-05 |
| I3U1H7 | EfaeciumATCCBAA472 | PF05362.16 | Lon_C | 1.20E-08 |
| I3U1P5 | EfaeciumATCCBAA472 | PF00717.26 | Peptidase_S24 | 2.60E-28 |
| I3U216 | EfaeciumATCCBAA472 | PF01425.24 | Amidase | 3.10E-68 |
| I3U2A3 | EfaeciumATCCBAA472 | PF12146.11 | Hydrolase_4 | 2.20E-06 |
| I3U2K8 | EfaeciumATCCBAA472 | PF01343.21 | Peptidase_S49 | 3.50E-50 |
| I3U2U5 | EfaeciumATCCBAA472 | PF01694.25 | Rhomboid | 9.30E-34 |
| I3U3E0 | EfaeciumATCCBAA472 | PF00768.23 | Peptidase_S11 | 4.90E-66 |
| I3U4H3 | EfaeciumATCCBAA472 | PF07859.16 | Abhydrolase_3 | 1.30E-64 |
| I3U4I5 | EfaeciumATCCBAA472 | PF10502.12 | Peptidase_S26 | 7.00E-47 |
| I3U4K3 | EfaeciumATCCBAA472 | PF00144.27 | Beta-lactamase | 1.40E-48 |
| I3U515 | EfaeciumATCCBAA472 | PF00326.24 | Peptidase_S9 | 2.80E-06 |
| I3U574 | EfaeciumATCCBAA472 | PF00756.23 | Esterase | 2.60E-21 |
| Q3XWL4 | EfaeciumATCCBAA472 | PF12146.11 | Hydrolase_4 | 9.90E-13 |
| Q3XWX3 | EfaeciumATCCBAA472 | PF02016.18 | Peptidase_S66 | 1.30E-29 |

| | | | | |
|---|---|---|---|---|
| Q3XX12 | EfaeciumATCCBAA472 | PF12146.11 | Hydrolase_4 | 1.80E-10 |
| Q3XX76 | EfaeciumATCCBAA472 | PF00574.26 | CLP_protease | 6.70E-88 |
| Q3XXW9 | EfaeciumATCCBAA472 | PF12695.10 | Abhydrolase_5 | 1.50E-67 |
| Q3XXY1 | EfaeciumATCCBAA472 | PF00144.27 | Beta-lactamase | 1.10E-39 |
| Q3XY33 | EfaeciumATCCBAA472 | PF00144.27 | Beta-lactamase | 1.20E-16 |
| Q3XY80 | EfaeciumATCCBAA472 | PF12146.11 | Hydrolase_4 | 5.30E-12 |
| Q3XYR5 | EfaeciumATCCBAA472 | PF13472.9 | Lipase_GDSL_2 | 4.10E-22 |
| Q3XZ82 | EfaeciumATCCBAA472 | PF01425.24 | Amidase | 8.10E-152 |
| Q3XZB9 | EfaeciumATCCBAA472 | PF10502.12 | Peptidase_S26 | 7.70E-45 |
| Q3XZC3 | EfaeciumATCCBAA472 | PF13472.9 | Lipase_GDSL_2 | 6.90E-25 |
| Q3XZI0 | EfaeciumATCCBAA472 | PF02129.21 | Peptidase_S15 | 2.30E-09 |
| Q3XZI8 | EfaeciumATCCBAA472 | PF02230.19 | Abhydrolase_2 | 1.50E-08 |
| Q3XZZ6 | EfaeciumATCCBAA472 | PF12146.11 | Hydrolase_4 | 1.50E-08 |
| Q3Y091 | EfaeciumATCCBAA472 | PF13365.9 | Trypsin_2 | 1.40E-32 |
| Q3Y0S5 | EfaeciumATCCBAA472 | PF00698.24 | Acyl_transf_1 | 1.30E-41 |
| Q3Y1A6 | EfaeciumATCCBAA472 | PF01734.25 | Patatin | 6.00E-27 |
| Q3Y1P7 | EfaeciumATCCBAA472 | PF04586.20 | Peptidase_S78 | 3.20E-47 |
| Q3Y1R9 | EfaeciumATCCBAA472 | PF02674.19 | Colicin_V | 4.50E-29 |
| Q3Y2N2 | EfaeciumATCCBAA472 | PF00574.26 | CLP_protease | 1.50E-32 |
| Q3Y3G0 | EfaeciumATCCBAA472 | PF13472.9 | Lipase_GDSL_2 | 7.10E-15 |
| B0G1G6 | DformicigeneransATCC27755 | PF05580.15 | Peptidase_S55 | 9.60E-82 |
| B0G1L5 | DformicigeneransATCC27755 | PF00768.23 | Peptidase_S11 | 2.60E-71 |
| B0G1Y3 | DformicigeneransATCC27755 | PF01425.24 | Amidase | 1.10E-159 |
| B0G220 | DformicigeneransATCC27755 | PF12146.11 | Hydrolase_4 | 2.80E-49 |
| B0G243 | DformicigeneransATCC27755 | PF00082.25 | Peptidase_S8 | 6.30E-13 |
| B0G2E1 | DformicigeneransATCC27755 | PF00144.27 | Beta-lactamase | 3.90E-28 |
| B0G2G8 | DformicigeneransATCC27755 | PF12146.11 | Hydrolase_4 | 4.00E-34 |
| B0G2Q2 | DformicigeneransATCC27755 | PF07859.16 | Abhydrolase_3 | 4.00E-24 |
| B0G2Q4 | DformicigeneransATCC27755 | PF07859.16 | Abhydrolase_3 | 2.10E-26 |
| B0G2Y7 | DformicigeneransATCC27755 | PF07859.16 | Abhydrolase_3 | 6.70E-67 |
| B0G367 | DformicigeneransATCC27755 | PF05362.16 | Lon_C | 2.00E-92 |
| B0G369 | DformicigeneransATCC27755 | PF00574.26 | CLP_protease | 6.80E-86 |
| B0G3R6 | DformicigeneransATCC27755 | PF13365.9 | Trypsin_2 | 7.10E-30 |
| B0G576 | DformicigeneransATCC27755 | PF00768.23 | Peptidase_S11 | 2.40E-48 |
| B0G5E9 | DformicigeneransATCC27755 | PF00574.26 | CLP_protease | 5.00E-25 |
| B0G5K4 | DformicigeneransATCC27755 | PF00768.23 | Peptidase_S11 | 4.10E-44 |
| B0G5L9 | DformicigeneransATCC27755 | PF00717.26 | Peptidase_S24 | 4.60E-30 |
| B0G5Y3 | DformicigeneransATCC27755 | PF10502.12 | Peptidase_S26 | 1.80E-46 |
| B0G5Z1 | DformicigeneransATCC27755 | PF10502.12 | Peptidase_S26 | 1.30E-41 |
| B0G5Z3 | DformicigeneransATCC27755 | PF10502.12 | Peptidase_S26 | 7.40E-37 |
| B0G602 | DformicigeneransATCC27755 | PF00082.25 | Peptidase_S8 | 3.50E-46 |
| B0G614 | DformicigeneransATCC27755 | PF01734.25 | Patatin | 4.70E-14 |
| B0G6P0 | DformicigeneransATCC27755 | PF00756.23 | Esterase | 3.70E-09 |
| B0G6T5 | DformicigeneransATCC27755 | PF00082.25 | Peptidase_S8 | 1.50E-16 |
| B0G748 | DformicigeneransATCC27755 | PF00698.24 | Acyl_transf_1 | 1.50E-37 |
| B0G7E2 | DformicigeneransATCC27755 | PF08840.14 | BAAT_C | 3.40E-22 |
| B0G7R2 | DformicigeneransATCC27755 | PF01343.21 | Peptidase_S49 | 1.80E-43 |
| B0G7Z9 | DformicigeneransATCC27755 | PF10502.12 | Peptidase_S26 | 5.30E-16 |
| B0G8A0 | DformicigeneransATCC27755 | PF03572.21 | Peptidase_S41 | 8.70E-54 |
| B0G8Z9 | DformicigeneransATCC27755 | PF00768.23 | Peptidase_S11 | 4.60E-42 |
| B0G939 | DformicigeneransATCC27755 | PF01694.25 | Rhomboid | 2.60E-35 |
| B0G9F8 | DformicigeneransATCC27755 | PF02674.19 | Colicin_V | 2.20E-11 |

| | | | | |
|---|---|---|---|---|
| A7LQU0 | BovatusATCC8483 | PF03629.21 | SASA | 1.00E-17 |
| A7LR33 | BovatusATCC8483 | PF03629.21 | SASA | 1.50E-26 |
| A7LR75 | BovatusATCC8483 | PF03572.21 | Peptidase_S41 | 2.90E-14 |
| A7LRA9 | BovatusATCC8483 | PF00574.26 | CLP_protease | 1.30E-77 |
| A7LRE5 | BovatusATCC8483 | PF00082.25 | Peptidase_S8 | 8.80E-44 |
| A7LRF7 | BovatusATCC8483 | PF10459.12 | Peptidase_S46 | 4.70E-260 |
| A7LRF8 | BovatusATCC8483 | PF10459.12 | Peptidase_S46 | 6.30E-257 |
| A7LRJ2 | BovatusATCC8483 | PF01734.25 | Patatin | 7.40E-26 |
| A7LS28 | BovatusATCC8483 | PF03572.21 | Peptidase_S41 | 7.30E-14 |
| A7LS33 | BovatusATCC8483 | PF13365.9 | Trypsin_2 | 3.50E-34 |
| A7LSB6 | BovatusATCC8483 | PF00326.24 | Peptidase_S9 | 2.70E-67 |
| A7LSD1 | BovatusATCC8483 | PF01734.25 | Patatin | 3.50E-28 |
| A7LSM6 | BovatusATCC8483 | PF03572.21 | Peptidase_S41 | 2.30E-19 |
| A7LSN4 | BovatusATCC8483 | PF08840.14 | BAAT_C | 9.90E-05 |
| A7LSV0 | BovatusATCC8483 | PF00326.24 | Peptidase_S9 | 2.60E-36 |
| A7LSV1 | BovatusATCC8483 | PF00326.24 | Peptidase_S9 | 9.60E-31 |
| A7LSV2 | BovatusATCC8483 | PF00326.24 | Peptidase_S9 | 3.50E-23 |
| A7LSV7 | BovatusATCC8483 | PF02230.19 | Abhydrolase_2 | 9.20E-12 |
| A7LSW0 | BovatusATCC8483 | PF10459.12 | Peptidase_S46 | 8.20E-262 |
| A7LT18 | BovatusATCC8483 | PF10502.12 | Peptidase_S26 | 8.40E-17 |
| A7LT19 | BovatusATCC8483 | PF10502.12 | Peptidase_S26 | 5.50E-25 |
| A7LT40 | BovatusATCC8483 | PF13472.9 | Lipase_GDSL_2 | 6.10E-15 |
| A7LT44 | BovatusATCC8483 | PF13472.9 | Lipase_GDSL_2 | 5.70E-13 |
| A7LTG4 | BovatusATCC8483 | PF00082.25 | Peptidase_S8 | 2.40E-39 |
| A7LTY7 | BovatusATCC8483 | PF03572.21 | Peptidase_S41 | 1.20E-12 |
| A7LTY9 | BovatusATCC8483 | PF00756.23 | Esterase | 1.50E-28 |
| A7LU17 | BovatusATCC8483 | PF00144.27 | Beta-lactamase | 3.80E-45 |
| A7LU17 | BovatusATCC8483 | PF12697.10 | Abhydrolase_6 | 8.50E-18 |
| A7LU53 | BovatusATCC8483 | PF00326.24 | Peptidase_S9 | 1.10E-45 |
| A7LUS8 | BovatusATCC8483 | PF00144.27 | Beta-lactamase | 2.40E-48 |
| A7LUX0 | BovatusATCC8483 | PF05448.15 | AXE1 | 1.80E-50 |
| A7LUY3 | BovatusATCC8483 | PF02016.18 | Peptidase_S66 | 8.50E-37 |
| A7LVF2 | BovatusATCC8483 | PF03629.21 | SASA | 9.20E-20 |
| A7LVF8 | BovatusATCC8483 | PF03629.21 | SASA | 1.20E-19 |
| A7LVQ0 | BovatusATCC8483 | PF02674.19 | Colicin_V | 8.30E-30 |
| A7LW22 | BovatusATCC8483 | PF03572.21 | Peptidase_S41 | 9.60E-49 |
| A7LW53 | BovatusATCC8483 | PF00756.23 | Esterase | 1.30E-30 |
| A7LW54 | BovatusATCC8483 | PF00756.23 | Esterase | 4.10E-37 |
| A7LW76 | BovatusATCC8483 | PF01694.25 | Rhomboid | 2.60E-33 |
| A7LW77 | BovatusATCC8483 | PF01694.25 | Rhomboid | 5.30E-24 |
| A7LWA1 | BovatusATCC8483 | PF05448.15 | AXE1 | 2.40E-39 |
| A7LWI9 | BovatusATCC8483 | PF13472.9 | Lipase_GDSL_2 | 3.90E-18 |
| A7LXE4 | BovatusATCC8483 | PF01734.25 | Patatin | 6.50E-21 |
| A7LXG7 | BovatusATCC8483 | PF00698.24 | Acyl_transf_1 | 2.20E-36 |
| A7LXK7 | BovatusATCC8483 | PF05362.16 | Lon_C | 5.80E-82 |
| A7LXS2 | BovatusATCC8483 | PF03629.21 | SASA | 8.10E-09 |
| A7LXS7 | BovatusATCC8483 | PF03572.21 | Peptidase_S41 | 1.20E-50 |
| A7LXX7 | BovatusATCC8483 | PF13472.9 | Lipase_GDSL_2 | 4.40E-14 |
| A7LXZ7 | BovatusATCC8483 | PF13472.9 | Lipase_GDSL_2 | 9.20E-14 |
| A7LY32 | BovatusATCC8483 | PF03629.21 | SASA | 1.40E-23 |
| A7LY96 | BovatusATCC8483 | PF12146.11 | Hydrolase_4 | 1.30E-15 |
| A7LYE1 | BovatusATCC8483 | PF03629.21 | SASA | 3.30E-20 |

| | | | | |
|---|---|---|---|---|
| A7LYH5 | BovatusATCC8483 | PF02113.18 | Peptidase_S13 | 1.20E-82 |
| A7LYH9 | BovatusATCC8483 | PF00717.26 | Peptidase_S24 | 1.10E-17 |
| A7LYM1 | BovatusATCC8483 | PF01343.21 | Peptidase_S49 | 4.00E-51 |
| A7LZV9 | BovatusATCC8483 | PF03572.21 | Peptidase_S41 | 1.90E-29 |
| A7M008 | BovatusATCC8483 | PF00756.23 | Esterase | 1.60E-21 |
| A7M008 | BovatusATCC8483 | PF03629.21 | SASA | 1.10E-32 |
| A7M023 | BovatusATCC8483 | PF03629.21 | SASA | 3.80E-18 |
| A7M0E0 | BovatusATCC8483 | PF05448.15 | AXE1 | 2.00E-41 |
| A7M0I9 | BovatusATCC8483 | PF03629.21 | SASA | 6.00E-59 |
| A7M0R4 | BovatusATCC8483 | PF00326.24 | Peptidase_S9 | 2.70E-56 |
| A7M0T5 | BovatusATCC8483 | PF13472.9 | Lipase_GDSL_2 | 6.80E-22 |
| A7M1K1 | BovatusATCC8483 | PF13472.9 | Lipase_GDSL_2 | 1.70E-26 |
| A7M2Q2 | BovatusATCC8483 | PF03629.21 | SASA | 3.60E-14 |
| A7M2R8 | BovatusATCC8483 | PF10459.12 | Peptidase_S46 | 3.50E-172 |
| A7M321 | BovatusATCC8483 | PF01734.25 | Patatin | 5.30E-23 |
| A7M352 | BovatusATCC8483 | PF01694.25 | Rhomboid | 3.30E-29 |
| A7M430 | BovatusATCC8483 | PF13472.9 | Lipase_GDSL_2 | 1.10E-13 |
| A7M445 | BovatusATCC8483 | PF13472.9 | Lipase_GDSL_2 | 1.30E-17 |
| A7M462 | BovatusATCC8483 | PF13472.9 | Lipase_GDSL_2 | 8.80E-14 |
| A7M496 | BovatusATCC8483 | PF00326.24 | Peptidase_S9 | 2.90E-06 |
| A7M496 | BovatusATCC8483 | PF13472.9 | Lipase_GDSL_2 | 7.30E-22 |
| A7M4A8 | BovatusATCC8483 | PF13472.9 | Lipase_GDSL_2 | 9.00E-13 |
| A7M4D1 | BovatusATCC8483 | PF13472.9 | Lipase_GDSL_2 | 3.40E-13 |
| A7M4E2 | BovatusATCC8483 | PF13472.9 | Lipase_GDSL_2 | 1.60E-10 |
| A7M4F0 | BovatusATCC8483 | PF03629.21 | SASA | 6.30E-19 |
| A7M4G3 | BovatusATCC8483 | PF03629.21 | SASA | 3.40E-20 |
| A7M4K5 | BovatusATCC8483 | PF00326.24 | Peptidase_S9 | 9.30E-60 |
| A7M4M3 | BovatusATCC8483 | PF00326.24 | Peptidase_S9 | 1.90E-15 |
| A7M4P9 | BovatusATCC8483 | PF03572.21 | Peptidase_S41 | 1.70E-52 |
| A7M4Q9 | BovatusATCC8483 | PF01694.25 | Rhomboid | 2.00E-13 |
| A7M4V2 | BovatusATCC8483 | PF00082.25 | Peptidase_S8 | 5.60E-24 |
| A7M5B2 | BovatusATCC8483 | PF13472.9 | Lipase_GDSL_2 | 2.80E-11 |
| A7M5B8 | BovatusATCC8483 | PF13354.9 | Beta-lactamase2 | 9.60E-29 |
| B7B4T6 | PjohnsoniiDSM18315 | PF00756.23 | Esterase | 5.80E-32 |
| B7B4Y3 | PjohnsoniiDSM18315 | PF13365.9 | Trypsin_2 | 1.90E-35 |
| B7B504 | PjohnsoniiDSM18315 | PF01694.25 | Rhomboid | 6.20E-20 |
| B7B5N8 | PjohnsoniiDSM18315 | PF03629.21 | SASA | 6.30E-11 |
| B7B5S1 | PjohnsoniiDSM18315 | PF01343.21 | Peptidase_S49 | 1.50E-49 |
| B7B5S6 | PjohnsoniiDSM18315 | PF01734.25 | Patatin | 3.10E-25 |
| B7B5Y0 | PjohnsoniiDSM18315 | PF10502.12 | Peptidase_S26 | 6.90E-27 |
| B7B5Y1 | PjohnsoniiDSM18315 | PF10502.12 | Peptidase_S26 | 6.60E-30 |
| B7B691 | PjohnsoniiDSM18315 | PF10459.12 | Peptidase_S46 | 7.00E-245 |
| B7B6A8 | PjohnsoniiDSM18315 | PF03629.21 | SASA | 6.40E-18 |
| B7B6A8 | PjohnsoniiDSM18315 | PF13472.9 | Lipase_GDSL_2 | 2.60E-23 |
| B7B6Q0 | PjohnsoniiDSM18315 | PF00326.24 | Peptidase_S9 | 2.50E-49 |
| B7B6Q5 | PjohnsoniiDSM18315 | PF05576.14 | Peptidase_S37 | 2.30E-51 |
| B7B6S7 | PjohnsoniiDSM18315 | PF03572.21 | Peptidase_S41 | 8.50E-32 |
| B7B730 | PjohnsoniiDSM18315 | PF02113.18 | Peptidase_S13 | 1.20E-69 |
| B7B739 | PjohnsoniiDSM18315 | PF03572.21 | Peptidase_S41 | 3.90E-51 |
| B7B788 | PjohnsoniiDSM18315 | PF13365.9 | Trypsin_2 | 2.10E-12 |
| B7B7B7 | PjohnsoniiDSM18315 | PF00326.24 | Peptidase_S9 | 2.60E-06 |
| B7B7B7 | PjohnsoniiDSM18315 | PF13472.9 | Lipase_GDSL_2 | 1.80E-09 |

| | | | | |
|---|---|---|---|---|
| B7B7T9 | PjohnsoniiDSM18315 | PF13472.9 | Lipase_GDSL_2 | 5.90E-15 |
| B7B865 | PjohnsoniiDSM18315 | PF02129.21 | Peptidase_S15 | 1.10E-07 |
| B7B935 | PjohnsoniiDSM18315 | PF01734.25 | Patatin | 7.50E-19 |
| B7B9I1 | PjohnsoniiDSM18315 | PF00326.24 | Peptidase_S9 | 5.60E-39 |
| B7B9Z6 | PjohnsoniiDSM18315 | PF03572.21 | Peptidase_S41 | 8.20E-50 |
| B7BA65 | PjohnsoniiDSM18315 | PF13472.9 | Lipase_GDSL_2 | 3.70E-17 |
| B7BAM0 | PjohnsoniiDSM18315 | PF00698.24 | Acyl_transf_1 | 4.50E-41 |
| B7BB12 | PjohnsoniiDSM18315 | PF00326.24 | Peptidase_S9 | 1.60E-47 |
| B7BB87 | PjohnsoniiDSM18315 | PF13472.9 | Lipase_GDSL_2 | 1.30E-13 |
| B7BBA1 | PjohnsoniiDSM18315 | PF01734.25 | Patatin | 1.30E-27 |
| B7BBD2 | PjohnsoniiDSM18315 | PF13354.9 | Beta-lactamase2 | 2.00E-34 |
| B7BBM7 | PjohnsoniiDSM18315 | PF00326.24 | Peptidase_S9 | 5.20E-54 |
| B7BBT8 | PjohnsoniiDSM18315 | PF00574.26 | CLP_protease | 4.60E-80 |
| B7BBZ8 | PjohnsoniiDSM18315 | PF13472.9 | Lipase_GDSL_2 | 3.80E-15 |
| B7BC10 | PjohnsoniiDSM18315 | PF13472.9 | Lipase_GDSL_2 | 4.50E-21 |
| B7BC75 | PjohnsoniiDSM18315 | PF00326.24 | Peptidase_S9 | 5.30E-55 |
| B7BCD6 | PjohnsoniiDSM18315 | PF10459.12 | Peptidase_S46 | 1.20E-262 |
| B7BCH1 | PjohnsoniiDSM18315 | PF05362.16 | Lon_C | 5.40E-83 |
| B7BDP9 | PjohnsoniiDSM18315 | PF00144.27 | Beta-lactamase | 5.30E-44 |
| B7BE54 | PjohnsoniiDSM18315 | PF00144.27 | Beta-lactamase | 1.20E-28 |
| B7BE70 | PjohnsoniiDSM18315 | PF05448.15 | AXE1 | 3.60E-46 |
| B7BE76 | PjohnsoniiDSM18315 | PF03572.21 | Peptidase_S41 | 6.60E-20 |
| B7BED0 | PjohnsoniiDSM18315 | PF12146.11 | Hydrolase_4 | 1.00E-12 |
| B7BEE4 | PjohnsoniiDSM18315 | PF13354.9 | Beta-lactamase2 | 7.00E-27 |
| B7BFK8 | PjohnsoniiDSM18315 | PF00326.24 | Peptidase_S9 | 7.50E-42 |
| B7BFR4 | PjohnsoniiDSM18315 | PF12146.11 | Hydrolase_4 | 4.30E-22 |
| B7BFT8 | PjohnsoniiDSM18315 | PF13472.9 | Lipase_GDSL_2 | 3.40E-22 |
| B7BFU3 | PjohnsoniiDSM18315 | PF01734.25 | Patatin | 1.70E-29 |
| B7BG80 | PjohnsoniiDSM18315 | PF03572.21 | Peptidase_S41 | 1.20E-44 |
| B7BG99 | PjohnsoniiDSM18315 | PF01694.25 | Rhomboid | 1.60E-21 |
| B7BGA0 | PjohnsoniiDSM18315 | PF01694.25 | Rhomboid | 2.90E-27 |
| B7BGH8 | PjohnsoniiDSM18315 | PF02674.19 | Colicin_V | 5.30E-15 |
| B7BGN8 | PjohnsoniiDSM18315 | PF00326.24 | Peptidase_S9 | 1.60E-47 |
| B7BGP4 | PjohnsoniiDSM18315 | PF00326.24 | Peptidase_S9 | 1.20E-23 |
| B7BGU5 | PjohnsoniiDSM18315 | PF10502.12 | Peptidase_S26 | 9.40E-22 |
| B7BGU6 | PjohnsoniiDSM18315 | PF10502.12 | Peptidase_S26 | 3.40E-07 |
| B7BH11 | PjohnsoniiDSM18315 | PF10502.12 | Peptidase_S26 | 8.70E-34 |
| B7BH50 | PjohnsoniiDSM18315 | PF00082.25 | Peptidase_S8 | 2.60E-32 |
| B7BH62 | PjohnsoniiDSM18315 | PF03572.21 | Peptidase_S41 | 7.30E-20 |
| B7BH68 | PjohnsoniiDSM18315 | PF03572.21 | Peptidase_S41 | 4.80E-14 |
| B7BHB3 | PjohnsoniiDSM18315 | PF00756.23 | Esterase | 2.00E-05 |
| B7BHK4 | PjohnsoniiDSM18315 | PF00756.23 | Esterase | 2.30E-28 |
| A0A1B0GVH | Hsapiens | PF00089.29 | Trypsin | 1.30E-13 |
| A1L453 | Hsapiens | PF00089.29 | Trypsin | 6.50E-58 |
| A4D1T9 | Hsapiens | PF00089.29 | Trypsin | 1.40E-26 |
| A6NIE9 | Hsapiens | PF00089.29 | Trypsin | 2.90E-59 |
| A8MTI9 | Hsapiens | PF00089.29 | Trypsin | 8.10E-46 |
| A8MY62 | Hsapiens | PF00144.27 | Beta-lactamase | 7.70E-40 |
| E5RG02 | Hsapiens | PF00089.29 | Trypsin | 2.80E-31 |
| O00187 | Hsapiens | PF00089.29 | Trypsin | 3.20E-56 |
| O00468 | Hsapiens | PF01390.23 | SEA | 4.00E-15 |
| O00519 | Hsapiens | PF01425.24 | Amidase | 9.70E-126 |

| | | | | |
|---|---|---|---|---|
| O00562 | Hsapiens | PF02862.20 | DDHD | 2.30E-30 |
| O00748 | Hsapiens | PF00135.31 | COesterase | 1.10E-169 |
| O14773 | Hsapiens | PF00082.25 | Peptidase_S8 | 1.00E-05 |
| O15393 | Hsapiens | PF00089.29 | Trypsin | 1.10E-66 |
| O43240 | Hsapiens | PF00089.29 | Trypsin | 4.00E-58 |
| O43464 | Hsapiens | PF13365.9 | Trypsin_2 | 8.60E-30 |
| O60235 | Hsapiens | PF00089.29 | Trypsin | 2.20E-72 |
| O60235 | Hsapiens | PF01390.23 | SEA | 8.30E-28 |
| O60259 | Hsapiens | PF00089.29 | Trypsin | 8.40E-74 |
| O60733 | Hsapiens | PF01734.25 | Patatin | 2.90E-15 |
| O75608 | Hsapiens | PF02230.19 | Abhydrolase_2 | 3.70E-87 |
| O75783 | Hsapiens | PF01694.25 | Rhomboid | 9.00E-36 |
| O75884 | Hsapiens | PF06821.16 | Ser_hydrolase | 4.10E-13 |
| O94830 | Hsapiens | PF02862.20 | DDHD | 3.80E-53 |
| O95084 | Hsapiens | PF00089.29 | Trypsin | 8.90E-10 |
| O95372 | Hsapiens | PF02230.19 | Abhydrolase_2 | 3.50E-89 |
| P00734 | Hsapiens | PF00089.29 | Trypsin | 8.50E-68 |
| P00736 | Hsapiens | PF00089.29 | Trypsin | 4.70E-49 |
| P00738 | Hsapiens | PF00089.29 | Trypsin | 2.10E-58 |
| P00739 | Hsapiens | PF00089.29 | Trypsin | 1.80E-54 |
| P00740 | Hsapiens | PF00089.29 | Trypsin | 1.30E-69 |
| P00742 | Hsapiens | PF00089.29 | Trypsin | 7.20E-70 |
| P00746 | Hsapiens | PF00089.29 | Trypsin | 4.60E-68 |
| P00747 | Hsapiens | PF00089.29 | Trypsin | 1.00E-65 |
| P00748 | Hsapiens | PF00089.29 | Trypsin | 3.70E-63 |
| P00749 | Hsapiens | PF00089.29 | Trypsin | 1.40E-69 |
| P00750 | Hsapiens | PF00089.29 | Trypsin | 4.50E-70 |
| P00751 | Hsapiens | PF00089.29 | Trypsin | 2.30E-47 |
| P01266 | Hsapiens | PF00135.31 | COesterase | 3.70E-131 |
| P02787 | Hsapiens | PF00405.20 | Transferrin | 1.30E-174 |
| P02788 | Hsapiens | PF00405.20 | Transferrin | 9.60E-179 |
| P03951 | Hsapiens | PF00089.29 | Trypsin | 4.10E-73 |
| P03952 | Hsapiens | PF00089.29 | Trypsin | 3.00E-75 |
| P04070 | Hsapiens | PF00089.29 | Trypsin | 7.90E-70 |
| P04180 | Hsapiens | PF02450.18 | LCAT | 3.30E-116 |
| P05156 | Hsapiens | PF00089.29 | Trypsin | 5.70E-62 |
| P05981 | Hsapiens | PF00089.29 | Trypsin | 1.20E-67 |
| P06276 | Hsapiens | PF00135.31 | COesterase | 1.50E-179 |
| P06681 | Hsapiens | PF00089.29 | Trypsin | 2.10E-29 |
| P06858 | Hsapiens | PF00151.22 | Lipase | 1.30E-130 |
| P06870 | Hsapiens | PF00089.29 | Trypsin | 9.10E-69 |
| P07288 | Hsapiens | PF00089.29 | Trypsin | 9.10E-69 |
| P07477 | Hsapiens | PF00089.29 | Trypsin | 6.50E-82 |
| P07478 | Hsapiens | PF00089.29 | Trypsin | 4.70E-80 |
| P08217 | Hsapiens | PF00089.29 | Trypsin | 1.40E-70 |
| P08218 | Hsapiens | PF00089.29 | Trypsin | 1.90E-67 |
| P08246 | Hsapiens | PF00089.29 | Trypsin | 4.00E-57 |
| P08311 | Hsapiens | PF00089.29 | Trypsin | 9.70E-65 |
| P08519 | Hsapiens | PF00089.29 | Trypsin | 2.40E-62 |
| P08582 | Hsapiens | PF00405.20 | Transferrin | 3.80E-145 |
| P08709 | Hsapiens | PF00089.29 | Trypsin | 4.80E-59 |
| P08861 | Hsapiens | PF00089.29 | Trypsin | 3.80E-68 |

| | | | | |
|---|---|---|---|---|
| P09093 | Hsapiens | PF00089.29 | Trypsin | 5.50E-66 |
| P09871 | Hsapiens | PF00089.29 | Trypsin | 4.20E-54 |
| P09958 | Hsapiens | PF00082.25 | Peptidase_S8 | 1.00E-47 |
| P0C7V7 | Hsapiens | PF00717.26 | Peptidase_S24 | 1.40E-05 |
| P0C869 | Hsapiens | PF01735.21 | PLA2_B | 6.60E-28 |
| P0CW18 | Hsapiens | PF00089.29 | Trypsin | 5.30E-66 |
| P10144 | Hsapiens | PF00089.29 | Trypsin | 1.10E-62 |
| P10323 | Hsapiens | PF00089.29 | Trypsin | 3.00E-69 |
| P10619 | Hsapiens | PF00450.25 | Peptidase_S10 | 2.50E-139 |
| P10745 | Hsapiens | PF03572.21 | Peptidase_S41 | 7.10E-25 |
| P10768 | Hsapiens | PF00756.23 | Esterase | 3.70E-73 |
| P11150 | Hsapiens | PF00151.22 | Lipase | 3.10E-130 |
| P12544 | Hsapiens | PF00089.29 | Trypsin | 2.40E-61 |
| P13798 | Hsapiens | PF00326.24 | Peptidase_S9 | 1.00E-33 |
| P14210 | Hsapiens | PF00089.29 | Trypsin | 1.10E-50 |
| P15941 | Hsapiens | PF01390.23 | SEA | 6.00E-12 |
| P16233 | Hsapiens | PF00151.22 | Lipase | 6.20E-159 |
| P16519 | Hsapiens | PF00082.25 | Peptidase_S8 | 4.40E-42 |
| P17538 | Hsapiens | PF00089.29 | Trypsin | 5.40E-73 |
| P19835 | Hsapiens | PF00135.31 | COesterase | 2.40E-164 |
| P20151 | Hsapiens | PF00089.29 | Trypsin | 9.40E-71 |
| P20160 | Hsapiens | PF00089.29 | Trypsin | 1.40E-49 |
| P20231 | Hsapiens | PF00089.29 | Trypsin | 2.50E-69 |
| P20718 | Hsapiens | PF00089.29 | Trypsin | 3.60E-63 |
| P22303 | Hsapiens | PF00135.31 | COesterase | 1.20E-172 |
| P22760 | Hsapiens | PF07859.16 | Abhydrolase_3 | 1.40E-37 |
| P22891 | Hsapiens | PF00089.29 | Trypsin | 4.70E-24 |
| P23141 | Hsapiens | PF00135.31 | COesterase | 7.50E-165 |
| P23946 | Hsapiens | PF00089.29 | Trypsin | 1.50E-60 |
| P24158 | Hsapiens | PF00089.29 | Trypsin | 5.40E-61 |
| P26927 | Hsapiens | PF00089.29 | Trypsin | 9.20E-47 |
| P27487 | Hsapiens | PF00326.24 | Peptidase_S9 | 3.80E-56 |
| P28039 | Hsapiens | PF00657.25 | Lipase_GDSL | 5.60E-28 |
| P29120 | Hsapiens | PF00082.25 | Peptidase_S8 | 2.70E-46 |
| P29122 | Hsapiens | PF00082.25 | Peptidase_S8 | 6.60E-47 |
| P29144 | Hsapiens | PF00082.25 | Peptidase_S8 | 1.00E-67 |
| P35030 | Hsapiens | PF00089.29 | Trypsin | 1.30E-79 |
| P36776 | Hsapiens | PF05362.16 | Lon_C | 7.70E-69 |
| P40313 | Hsapiens | PF00089.29 | Trypsin | 5.70E-71 |
| P41247 | Hsapiens | PF01734.25 | Patatin | 8.20E-15 |
| P42658 | Hsapiens | PF00326.24 | Peptidase_S9 | 1.20E-39 |
| P42785 | Hsapiens | PF05577.15 | Peptidase_S28 | 4.80E-97 |
| P47712 | Hsapiens | PF01735.21 | PLA2_B | 5.80E-149 |
| P48147 | Hsapiens | PF00326.24 | Peptidase_S9 | 2.90E-68 |
| P48740 | Hsapiens | PF00089.29 | Trypsin | 6.90E-74 |
| P49327 | Hsapiens | PF00698.24 | Acyl_transf_1 | 5.20E-116 |
| P49327 | Hsapiens | PF00975.23 | Thioesterase | 1.40E-66 |
| P49753 | Hsapiens | PF08840.14 | BAAT_C | 1.10E-90 |
| P49862 | Hsapiens | PF00089.29 | Trypsin | 1.30E-68 |
| P49863 | Hsapiens | PF00089.29 | Trypsin | 1.80E-67 |
| P50897 | Hsapiens | PF02089.18 | Palm_thioest | 2.80E-88 |
| P51124 | Hsapiens | PF00089.29 | Trypsin | 4.40E-59 |

| | | | | |
|---|---|---|---|---|
| P52948 | Hsapiens | PF04096.17 | Nucleoporin2 | 2.90E-45 |
| P54315 | Hsapiens | PF00151.22 | Lipase | 7.30E-154 |
| P54317 | Hsapiens | PF00151.22 | Lipase | 8.40E-161 |
| P56730 | Hsapiens | PF00089.29 | Trypsin | 4.80E-65 |
| P57727 | Hsapiens | PF00089.29 | Trypsin | 2.70E-68 |
| P58872 | Hsapiens | PF01694.25 | Rhomboid | 3.90E-35 |
| P67812 | Hsapiens | PF00717.26 | Peptidase_S24 | 1.70E-09 |
| P68402 | Hsapiens | PF13472.9 | Lipase_GDSL_2 | 4.00E-15 |
| P83105 | Hsapiens | PF13365.9 | Trypsin_2 | 2.20E-28 |
| P83110 | Hsapiens | PF13365.9 | Trypsin_2 | 7.30E-29 |
| P83111 | Hsapiens | PF00144.27 | Beta-lactamase | 4.10E-29 |
| P98073 | Hsapiens | PF00089.29 | Trypsin | 3.80E-71 |
| P98073 | Hsapiens | PF01390.23 | SEA | 3.80E-18 |
| Q04756 | Hsapiens | PF00089.29 | Trypsin | 6.00E-68 |
| Q05469 | Hsapiens | PF07859.16 | Abhydrolase_3 | 7.20E-34 |
| Q12884 | Hsapiens | PF00326.24 | Peptidase_S9 | 4.80E-54 |
| Q13093 | Hsapiens | PF03403.16 | PAF-AH_p_II | 7.50E-183 |
| Q14032 | Hsapiens | PF08840.14 | BAAT_C | 1.40E-72 |
| Q14118 | Hsapiens | PF05454.14 | DAG1 | 2.60E-162 |
| Q14520 | Hsapiens | PF00089.29 | Trypsin | 2.30E-62 |
| Q14703 | Hsapiens | PF00082.25 | Peptidase_S8 | 1.20E-42 |
| Q15102 | Hsapiens | PF13472.9 | Lipase_GDSL_2 | 6.00E-17 |
| Q15661 | Hsapiens | PF00089.29 | Trypsin | 2.50E-69 |
| Q16549 | Hsapiens | PF00082.25 | Peptidase_S8 | 5.70E-42 |
| Q16651 | Hsapiens | PF00089.29 | Trypsin | 1.30E-74 |
| Q16740 | Hsapiens | PF00574.26 | CLP_protease | 3.00E-80 |
| Q17R60 | Hsapiens | PF01390.23 | SEA | 1.40E-14 |
| Q17RR3 | Hsapiens | PF00151.22 | Lipase | 8.70E-115 |
| Q2L4Q9 | Hsapiens | PF00089.29 | Trypsin | 1.90E-43 |
| Q2T9J0 | Hsapiens | PF13365.9 | Trypsin_2 | 2.40E-19 |
| Q2TAA2 | Hsapiens | PF13472.9 | Lipase_GDSL_2 | 1.60E-31 |
| Q2TV78 | Hsapiens | PF00089.29 | Trypsin | 1.90E-44 |
| Q3I5F7 | Hsapiens | PF08840.14 | BAAT_C | 7.20E-84 |
| Q3MJ16 | Hsapiens | PF01735.21 | PLA2_B | 3.80E-34 |
| Q4J6C6 | Hsapiens | PF00326.24 | Peptidase_S9 | 7.20E-30 |
| Q53H76 | Hsapiens | PF00151.22 | Lipase | 6.80E-104 |
| Q5DID0 | Hsapiens | PF01390.23 | SEA | 5.90E-07 |
| Q5K4E3 | Hsapiens | PF00089.29 | Trypsin | 4.90E-63 |
| Q5T601 | Hsapiens | PF01390.23 | SEA | 1.40E-08 |
| Q5VST6 | Hsapiens | PF00326.24 | Peptidase_S9 | 2.60E-05 |
| Q5VUY0 | Hsapiens | PF07859.16 | Abhydrolase_3 | 2.60E-34 |
| Q5VUY2 | Hsapiens | PF07859.16 | Abhydrolase_3 | 4.00E-27 |
| Q5VWZ2 | Hsapiens | PF02230.19 | Abhydrolase_2 | 3.60E-42 |
| Q5XG92 | Hsapiens | PF00135.31 | COesterase | 3.30E-158 |
| Q63HM1 | Hsapiens | PF07859.16 | Abhydrolase_3 | 4.70E-15 |
| Q685J3 | Hsapiens | PF01390.23 | SEA | 5.30E-13 |
| Q68DD2 | Hsapiens | PF01735.21 | PLA2_B | 1.00E-33 |
| Q6GMR7 | Hsapiens | PF01425.24 | Amidase | 1.80E-75 |
| Q6GPI1 | Hsapiens | PF00089.29 | Trypsin | 1.30E-72 |
| Q6NT32 | Hsapiens | PF00135.31 | COesterase | 1.40E-153 |
| Q6NTF9 | Hsapiens | PF01694.25 | Rhomboid | 1.10E-22 |
| Q6P093 | Hsapiens | PF07859.16 | Abhydrolase_3 | 2.80E-38 |

| | | | | |
|---|---|---|---|---|
| Q6P1J6 | Hsapiens | PF00657.25 | Lipase_GDSL | 2.30E-31 |
| Q6P988 | Hsapiens | PF03283.16 | PAE | 3.20E-89 |
| Q6PCB6 | Hsapiens | PF12146.11 | Hydrolase_4 | 1.50E-06 |
| Q6PEW0 | Hsapiens | PF00089.29 | Trypsin | 3.30E-26 |
| Q6PIU2 | Hsapiens | PF07859.16 | Abhydrolase_3 | 1.40E-34 |
| Q6PJF5 | Hsapiens | PF01694.25 | Rhomboid | 2.60E-27 |
| Q6SJ93 | Hsapiens | PF13365.9 | Trypsin_2 | 2.70E-21 |
| Q6UW60 | Hsapiens | PF00082.25 | Peptidase_S8 | 3.90E-45 |
| Q6UWB4 | Hsapiens | PF00089.29 | Trypsin | 1.80E-62 |
| Q6UWW8 | Hsapiens | PF00135.31 | COesterase | 3.30E-156 |
| Q6UWY2 | Hsapiens | PF00089.29 | Trypsin | 2.40E-56 |
| Q6UXH9 | Hsapiens | PF00089.29 | Trypsin | 8.70E-39 |
| Q6V1X1 | Hsapiens | PF00326.24 | Peptidase_S9 | 2.20E-61 |
| Q6XZB0 | Hsapiens | PF00151.22 | Lipase | 1.70E-85 |
| Q6ZMR5 | Hsapiens | PF00089.29 | Trypsin | 9.00E-65 |
| Q6ZMR5 | Hsapiens | PF01390.23 | SEA | 2.40E-28 |
| Q6ZV29 | Hsapiens | PF01734.25 | Patatin | 2.90E-20 |
| Q6ZWK6 | Hsapiens | PF00089.29 | Trypsin | 6.50E-67 |
| Q6ZWK6 | Hsapiens | PF01390.23 | SEA | 1.50E-27 |
| Q75T13 | Hsapiens | PF07819.16 | PGAP1 | 2.00E-81 |
| Q7L211 | Hsapiens | PF12146.11 | Hydrolase_4 | 8.60E-10 |
| Q7RTY3 | Hsapiens | PF00089.29 | Trypsin | 1.90E-60 |
| Q7RTY5 | Hsapiens | PF00089.29 | Trypsin | 1.00E-63 |
| Q7RTY7 | Hsapiens | PF00089.29 | Trypsin | 1.90E-65 |
| Q7RTY8 | Hsapiens | PF00089.29 | Trypsin | 8.50E-62 |
| Q7RTY8 | Hsapiens | PF01390.23 | SEA | 1.30E-21 |
| Q7RTY9 | Hsapiens | PF00089.29 | Trypsin | 7.80E-61 |
| Q7RTZ1 | Hsapiens | PF00089.29 | Trypsin | 1.90E-65 |
| Q7Z410 | Hsapiens | PF00089.29 | Trypsin | 1.70E-69 |
| Q7Z5A4 | Hsapiens | PF00089.29 | Trypsin | 1.90E-54 |
| Q7Z5M8 | Hsapiens | PF12146.11 | Hydrolase_4 | 2.00E-16 |
| Q7Z6Z6 | Hsapiens | PF01734.25 | Patatin | 2.20E-14 |
| Q86T26 | Hsapiens | PF00089.29 | Trypsin | 1.90E-67 |
| Q86T26 | Hsapiens | PF01390.23 | SEA | 8.90E-20 |
| Q86TI2 | Hsapiens | PF00326.24 | Peptidase_S9 | 3.90E-56 |
| Q86TX2 | Hsapiens | PF08840.14 | BAAT_C | 1.00E-90 |
| Q86WA8 | Hsapiens | PF05362.16 | Lon_C | 3.80E-79 |
| Q86WS5 | Hsapiens | PF00089.29 | Trypsin | 1.00E-62 |
| Q86XP0 | Hsapiens | PF01735.21 | PLA2_B | 7.50E-26 |
| Q8IU80 | Hsapiens | PF00089.29 | Trypsin | 8.60E-69 |
| Q8IU80 | Hsapiens | PF01390.23 | SEA | 3.80E-09 |
| Q8IVS2 | Hsapiens | PF00698.24 | Acyl_transf_1 | 6.00E-24 |
| Q8IY17 | Hsapiens | PF01734.25 | Patatin | 6.00E-23 |
| Q8IYP2 | Hsapiens | PF00089.29 | Trypsin | 3.30E-41 |
| Q8IZF2 | Hsapiens | PF01390.23 | SEA | 2.90E-08 |
| Q8N0W4 | Hsapiens | PF00135.31 | COesterase | 2.30E-206 |
| Q8N2K0 | Hsapiens | PF12146.11 | Hydrolase_4 | 1.20E-13 |
| Q8N2Q7 | Hsapiens | PF00135.31 | COesterase | 8.90E-198 |
| Q8N3Z0 | Hsapiens | PF00089.29 | Trypsin | 3.40E-09 |
| Q8N608 | Hsapiens | PF00326.24 | Peptidase_S9 | 2.10E-47 |
| Q8N9L9 | Hsapiens | PF08840.14 | BAAT_C | 8.40E-86 |
| Q8NBP7 | Hsapiens | PF00082.25 | Peptidase_S8 | 4.80E-29 |

| | | | | |
|---|---|---|---|---|
| Q8NCC3 | Hsapiens | PF02450.18 | LCAT | 3.20E-73 |
| Q8NCG7 | Hsapiens | PF01764.28 | Lipase_3 | 1.90E-18 |
| Q8NEL9 | Hsapiens | PF02862.20 | DDHD | 7.00E-57 |
| Q8NF86 | Hsapiens | PF00089.29 | Trypsin | 4.60E-67 |
| Q8NFV4 | Hsapiens | PF12697.10 | Abhydrolase_6 | 1.20E-24 |
| Q8NFZ3 | Hsapiens | PF00135.31 | COesterase | 5.80E-205 |
| Q8NFZ4 | Hsapiens | PF00135.31 | COesterase | 4.30E-198 |
| Q8NHM4 | Hsapiens | PF00089.29 | Trypsin | 5.00E-80 |
| Q8TEB9 | Hsapiens | PF01694.25 | Rhomboid | 6.70E-22 |
| Q8WWY8 | Hsapiens | PF00151.22 | Lipase | 1.80E-81 |
| Q8WXI7 | Hsapiens | PF01390.23 | SEA | 4.70E-26 |
| Q8WZ82 | Hsapiens | PF03959.16 | FSH1 | 1.80E-54 |
| Q92743 | Hsapiens | PF13365.9 | Trypsin_2 | 5.90E-31 |
| Q92824 | Hsapiens | PF00082.25 | Peptidase_S8 | 7.70E-47 |
| Q92876 | Hsapiens | PF00089.29 | Trypsin | 6.70E-73 |
| Q96AD5 | Hsapiens | PF01734.25 | Patatin | 1.00E-14 |
| Q96CC6 | Hsapiens | PF01694.25 | Rhomboid | 7.60E-28 |
| Q96GS6 | Hsapiens | PF12146.11 | Hydrolase_4 | 8.10E-07 |
| Q96IU4 | Hsapiens | PF12697.10 | Abhydrolase_6 | 1.80E-08 |
| Q96LU5 | Hsapiens | PF10502.12 | Peptidase_S26 | 3.40E-14 |
| Q96LU7 | Hsapiens | PF13884.9 | Peptidase_S74 | 9.40E-15 |
| Q96PZ2 | Hsapiens | PF13365.9 | Trypsin_2 | 4.60E-16 |
| Q96T52 | Hsapiens | PF10502.12 | Peptidase_S26 | 6.00E-13 |
| Q99487 | Hsapiens | PF03403.16 | PAF-AH_p_II | 9.90E-172 |
| Q99685 | Hsapiens | PF12146.11 | Hydrolase_4 | 7.90E-74 |
| Q99895 | Hsapiens | PF00089.29 | Trypsin | 1.60E-68 |
| Q9BQR3 | Hsapiens | PF00089.29 | Trypsin | 1.60E-68 |
| Q9BUJ0 | Hsapiens | PF12697.10 | Abhydrolase_6 | 2.10E-07 |
| Q9BY50 | Hsapiens | PF00717.26 | Peptidase_S24 | 7.60E-10 |
| Q9BYE2 | Hsapiens | PF00089.29 | Trypsin | 9.60E-68 |
| Q9BZ71 | Hsapiens | PF02862.20 | DDHD | 1.20E-41 |
| Q9BZ72 | Hsapiens | PF02862.20 | DDHD | 4.30E-55 |
| Q9BZJ3 | Hsapiens | PF00089.29 | Trypsin | 8.40E-55 |
| Q9BZV3 | Hsapiens | PF01390.23 | SEA | 5.50E-15 |
| Q9C000 | Hsapiens | PF13553.9 | FIIND | 4.80E-106 |
| Q9GZN4 | Hsapiens | PF00089.29 | Trypsin | 2.60E-65 |
| Q9H0R6 | Hsapiens | PF01425.24 | Amidase | 2.40E-125 |
| Q9H195 | Hsapiens | PF01390.23 | SEA | 5.60E-12 |
| Q9H2R5 | Hsapiens | PF00089.29 | Trypsin | 6.60E-70 |
| Q9H300 | Hsapiens | PF01694.25 | Rhomboid | 2.30E-29 |
| Q9H3G5 | Hsapiens | PF00450.25 | Peptidase_S10 | 5.40E-104 |
| Q9H3R2 | Hsapiens | PF01390.23 | SEA | 2.30E-11 |
| Q9H3S3 | Hsapiens | PF00089.29 | Trypsin | 6.10E-72 |
| Q9H6B9 | Hsapiens | PF12697.10 | Abhydrolase_6 | 1.60E-18 |
| Q9H6V9 | Hsapiens | PF10230.12 | LIDHydrolase | 1.10E-80 |
| Q9HAT2 | Hsapiens | PF03629.21 | SASA | 7.50E-14 |
| Q9HB40 | Hsapiens | PF00450.25 | Peptidase_S10 | 4.10E-93 |
| Q9HB75 | Hsapiens | PF10461.12 | Peptidase_S68 | 3.80E-16 |
| Q9NP80 | Hsapiens | PF01734.25 | Patatin | 2.50E-23 |
| Q9NQE7 | Hsapiens | PF05577.15 | Peptidase_S28 | 3.10E-153 |
| Q9NRR2 | Hsapiens | PF00089.29 | Trypsin | 1.10E-65 |
| Q9NRS4 | Hsapiens | PF00089.29 | Trypsin | 1.00E-65 |

| | | | | |
|---|---|---|---|---|
| Q9NST1 | Hsapiens | PF01734.25 | Patatin | 2.70E-13 |
| Q9NV23 | Hsapiens | PF00975.23 | Thioesterase | 1.40E-39 |
| Q9NX52 | Hsapiens | PF01694.25 | Rhomboid | 3.90E-40 |
| Q9NZ94 | Hsapiens | PF00135.31 | COesterase | 5.80E-204 |
| Q9NZP8 | Hsapiens | PF00089.29 | Trypsin | 7.10E-45 |
| Q9P0G3 | Hsapiens | PF00089.29 | Trypsin | 9.10E-73 |
| Q9UBX7 | Hsapiens | PF00089.29 | Trypsin | 2.80E-70 |
| Q9UHL4 | Hsapiens | PF05577.15 | Peptidase_S28 | 1.50E-85 |
| Q9UI38 | Hsapiens | PF00089.29 | Trypsin | 1.10E-48 |
| Q9UKN1 | Hsapiens | PF01390.23 | SEA | 2.90E-08 |
| Q9UKQ9 | Hsapiens | PF00089.29 | Trypsin | 1.80E-63 |
| Q9UKR0 | Hsapiens | PF00089.29 | Trypsin | 6.30E-64 |
| Q9UKR3 | Hsapiens | PF00089.29 | Trypsin | 1.90E-74 |
| Q9UKY3 | Hsapiens | PF00135.31 | COesterase | 1.40E-99 |
| Q9UL52 | Hsapiens | PF00089.29 | Trypsin | 4.50E-67 |
| Q9UL52 | Hsapiens | PF01390.23 | SEA | 1.20E-20 |
| Q9UMR5 | Hsapiens | PF02089.18 | Palm_thioest | 3.50E-63 |
| Q9UNI1 | Hsapiens | PF00089.29 | Trypsin | 3.80E-69 |
| Q9UP65 | Hsapiens | PF01735.21 | PLA2_B | 5.00E-25 |
| Q9Y2G1 | Hsapiens | PF13884.9 | Peptidase_S74 | 5.80E-13 |
| Q9Y2G2 | Hsapiens | PF13553.9 | FIIND | 6.60E-96 |
| Q9Y337 | Hsapiens | PF00089.29 | Trypsin | 1.90E-68 |
| Q9Y3P4 | Hsapiens | PF01694.25 | Rhomboid | 2.60E-07 |
| Q9Y4D2 | Hsapiens | PF01764.28 | Lipase_3 | 5.30E-10 |
| Q9Y570 | Hsapiens | PF12697.10 | Abhydrolase_6 | 3.60E-30 |
| Q9Y5K2 | Hsapiens | PF00089.29 | Trypsin | 7.00E-63 |
| Q9Y5Q5 | Hsapiens | PF00089.29 | Trypsin | 2.70E-60 |
| Q9Y5X9 | Hsapiens | PF00151.22 | Lipase | 1.10E-107 |
| Q9Y5Y6 | Hsapiens | PF00089.29 | Trypsin | 4.30E-65 |
| Q9Y5Y6 | Hsapiens | PF01390.23 | SEA | 6.10E-13 |
| Q9Y6M0 | Hsapiens | PF00089.29 | Trypsin | 5.80E-59 |
| Q9Y6Y8 | Hsapiens | PF02862.20 | DDHD | 6.00E-58 |
| A0A7U9C4B | CsporogenesATCC15579 | PF04586.20 | Peptidase_S78 | 1.40E-31 |
| A0A7U9C5A | CsporogenesATCC15579 | PF00574.26 | CLP_protease | 3.70E-33 |
| A0A7U9C5U | CsporogenesATCC15579 | PF00768.23 | Peptidase_S11 | 1.20E-55 |
| A0A7U9C5V | CsporogenesATCC15579 | PF05362.16 | Lon_C | 8.60E-35 |
| A0A7U9C5Y | CsporogenesATCC15579 | PF00698.24 | Acyl_transf_1 | 6.30E-36 |
| A0A7U9C61 | CsporogenesATCC15579 | PF13365.9 | Trypsin_2 | 1.70E-31 |
| A0A7U9C7F | CsporogenesATCC15579 | PF00768.23 | Peptidase_S11 | 1.70E-76 |
| A0A7U9C7P | CsporogenesATCC15579 | PF00144.27 | Beta-lactamase | 4.10E-59 |
| A0A7U9C9G | CsporogenesATCC15579 | PF12146.11 | Hydrolase_4 | 6.80E-07 |
| A0A7U9C9K | CsporogenesATCC15579 | PF05580.15 | Peptidase_S55 | 1.80E-89 |
| A0A7U9C9P | CsporogenesATCC15579 | PF00574.26 | CLP_protease | 4.70E-29 |
| A0A7U9CAF | CsporogenesATCC15579 | PF01734.25 | Patatin | 5.50E-22 |
| A0A7U9CAI7 | CsporogenesATCC15579 | PF00574.26 | CLP_protease | 2.90E-67 |
| A0A7U9CBT | CsporogenesATCC15579 | PF01734.25 | Patatin | 4.60E-27 |
| A0A7U9CCB | CsporogenesATCC15579 | PF00717.26 | Peptidase_S24 | 3.60E-25 |
| A0A7U9CCC | CsporogenesATCC15579 | PF05362.16 | Lon_C | 1.50E-16 |
| A0A7U9CD0 | CsporogenesATCC15579 | PF12146.11 | Hydrolase_4 | 2.00E-62 |
| A0A7U9CFB | CsporogenesATCC15579 | PF00574.26 | CLP_protease | 9.20E-89 |
| A0A7U9GI3 | CsporogenesATCC15579 | PF00717.26 | Peptidase_S24 | 1.60E-07 |
| A0A7U9GIF | CsporogenesATCC15579 | PF03572.21 | Peptidase_S41 | 7.20E-51 |

| | | | | |
|---|---|---|---|---|
| A0A7U9GIQ | CsporogenesATCC15579 | PF00975.23 | Thioesterase | 1.20E-37 |
| A0A7U9GIR | CsporogenesATCC15579 | PF00082.25 | Peptidase_S8 | 1.60E-26 |
| A0A7U9GJ0 | CsporogenesATCC15579 | PF00768.23 | Peptidase_S11 | 1.70E-63 |
| A0A7U9GJJ | CsporogenesATCC15579 | PF10502.12 | Peptidase_S26 | 1.10E-40 |
| A0A7U9GJJ | CsporogenesATCC15579 | PF00144.27 | Beta-lactamase | 1.00E-56 |
| A0A7U9GJK | CsporogenesATCC15579 | PF00975.23 | Thioesterase | 8.30E-36 |
| A0A7U9GJR | CsporogenesATCC15579 | PF00717.26 | Peptidase_S24 | 1.40E-30 |
| A0A7U9GKA | CsporogenesATCC15579 | PF02016.18 | Peptidase_S66 | 3.60E-37 |
| A0A7U9GKC | CsporogenesATCC15579 | PF12146.11 | Hydrolase_4 | 3.10E-08 |
| A0A7U9GKF | CsporogenesATCC15579 | PF01425.24 | Amidase | 1.50E-159 |
| A0A7U9GKK | CsporogenesATCC15579 | PF13472.9 | Lipase_GDSL_2 | 4.80E-19 |
| A0A7U9GKT | CsporogenesATCC15579 | PF07859.16 | Abhydrolase_3 | 1.50E-68 |
| A0A7U9GL0 | CsporogenesATCC15579 | PF03575.20 | Peptidase_S51 | 8.80E-24 |
| A0A7U9GL0 | CsporogenesATCC15579 | PF01694.25 | Rhomboid | 8.80E-37 |
| A0A7U9GL7 | CsporogenesATCC15579 | PF05362.16 | Lon_C | 9.90E-35 |
| A0A7U9GL8 | CsporogenesATCC15579 | PF00144.27 | Beta-lactamase | 1.30E-60 |
| A0A7U9GLG | CsporogenesATCC15579 | PF05362.16 | Lon_C | 1.90E-95 |
| A0A7U9GLQ | CsporogenesATCC15579 | PF00574.26 | CLP_protease | 1.40E-27 |
| A0A7U9GM3 | CsporogenesATCC15579 | PF10502.12 | Peptidase_S26 | 3.30E-52 |
| A0A7U9GM9 | CsporogenesATCC15579 | PF00768.23 | Peptidase_S11 | 1.00E-50 |
| C8NDN5 | GadiacensATCC49175 | PF00326.24 | Peptidase_S9 | 4.90E-05 |
| C8NDN5 | GadiacensATCC49175 | PF12146.11 | Hydrolase_4 | 1.90E-08 |
| C8NDY2 | GadiacensATCC49175 | PF01694.25 | Rhomboid | 8.80E-35 |
| C8NE35 | GadiacensATCC49175 | PF00717.26 | Peptidase_S24 | 3.40E-30 |
| C8NE84 | GadiacensATCC49175 | PF00326.24 | Peptidase_S9 | 2.40E-12 |
| C8NED3 | GadiacensATCC49175 | PF00698.24 | Acyl_transf_1 | 3.30E-34 |
| C8NEG0 | GadiacensATCC49175 | PF00768.23 | Peptidase_S11 | 7.00E-55 |
| C8NEM7 | GadiacensATCC49175 | PF03572.21 | Peptidase_S41 | 6.00E-47 |
| C8NEM8 | GadiacensATCC49175 | PF10502.12 | Peptidase_S26 | 3.70E-51 |
| C8NF65 | GadiacensATCC49175 | PF00756.23 | Esterase | 1.70E-11 |
| C8NFF9 | GadiacensATCC49175 | PF00326.24 | Peptidase_S9 | 3.40E-53 |
| C8NFG2 | GadiacensATCC49175 | PF00768.23 | Peptidase_S11 | 4.90E-47 |
| C8NFM5 | GadiacensATCC49175 | PF12695.10 | Abhydrolase_5 | 3.40E-49 |
| C8NFW3 | GadiacensATCC49175 | PF02674.19 | Colicin_V | 2.70E-24 |
| C8NG30 | GadiacensATCC49175 | PF01425.24 | Amidase | 7.00E-74 |
| C8NG95 | GadiacensATCC49175 | PF00574.26 | CLP_protease | 5.40E-87 |
| C8NGA9 | GadiacensATCC49175 | PF00756.23 | Esterase | 7.30E-20 |
| C8NGS7 | GadiacensATCC49175 | PF05362.16 | Lon_C | 4.80E-08 |
| C8NH78 | GadiacensATCC49175 | PF02016.18 | Peptidase_S66 | 5.30E-27 |
| C8NHC9 | GadiacensATCC49175 | PF12697.10 | Abhydrolase_6 | 2.50E-11 |
| C8NHF7 | GadiacensATCC49175 | PF01425.24 | Amidase | 1.10E-155 |
| C8NHK1 | GadiacensATCC49175 | PF12146.11 | Hydrolase_4 | 1.20E-10 |
| C8NHP5 | GadiacensATCC49175 | PF08840.14 | BAAT_C | 1.20E-15 |
| C8NHQ1 | GadiacensATCC49175 | PF03575.20 | Peptidase_S51 | 3.60E-44 |
| C8NHW2 | GadiacensATCC49175 | PF04586.20 | Peptidase_S78 | 7.10E-43 |
| C8NHZ8 | GadiacensATCC49175 | PF13365.9 | Trypsin_2 | 1.30E-29 |
| C8NIC3 | GadiacensATCC49175 | PF00144.27 | Beta-lactamase | 3.80E-65 |
| C8NIE3 | GadiacensATCC49175 | PF01425.24 | Amidase | 1.60E-83 |
| C8NIH9 | GadiacensATCC49175 | PF12146.11 | Hydrolase_4 | 7.50E-09 |
| C8NIM7 | GadiacensATCC49175 | PF00768.23 | Peptidase_S11 | 6.90E-62 |
| C0AQJ0 | PpenneriATCC35198 | PF04586.20 | Peptidase_S78 | 6.70E-18 |
| C0AQM6 | PpenneriATCC35198 | PF01343.21 | Peptidase_S49 | 7.20E-43 |

| | | | | |
|---|---|---|---|---|
| C0AQU8 | PpenneriATCC35198 | PF00717.26 | Peptidase_S24 | 1.60E-12 |
| C0AQW6 | PpenneriATCC35198 | PF00326.24 | Peptidase_S9 | 5.80E-66 |
| C0AR23 | PpenneriATCC35198 | PF13472.9 | Lipase_GDSL_2 | 6.40E-10 |
| C0AR24 | PpenneriATCC35198 | PF13472.9 | Lipase_GDSL_2 | 3.20E-15 |
| C0ARL0 | PpenneriATCC35198 | PF00082.25 | Peptidase_S8 | 1.90E-19 |
| C0ARN1 | PpenneriATCC35198 | PF00717.26 | Peptidase_S24 | 2.80E-18 |
| C0ARV4 | PpenneriATCC35198 | PF12146.11 | Hydrolase_4 | 3.40E-09 |
| C0ARV5 | PpenneriATCC35198 | PF12146.11 | Hydrolase_4 | 7.40E-34 |
| C0ASP9 | PpenneriATCC35198 | PF01694.25 | Rhomboid | 6.00E-19 |
| C0AT67 | PpenneriATCC35198 | PF00082.25 | Peptidase_S8 | 1.00E-19 |
| C0ATJ6 | PpenneriATCC35198 | PF01734.25 | Patatin | 1.70E-12 |
| C0ATL1 | PpenneriATCC35198 | PF00756.23 | Esterase | 7.70E-25 |
| C0AUC4 | PpenneriATCC35198 | PF01694.25 | Rhomboid | 3.20E-27 |
| C0AUL4 | PpenneriATCC35198 | PF00144.27 | Beta-lactamase | 1.40E-45 |
| C0AVE5 | PpenneriATCC35198 | PF02674.19 | Colicin_V | 2.30E-27 |
| C0AW03 | PpenneriATCC35198 | PF00717.26 | Peptidase_S24 | 2.40E-20 |
| C0AW22 | PpenneriATCC35198 | PF10502.12 | Peptidase_S26 | 7.20E-55 |
| C0AWX8 | PpenneriATCC35198 | PF04586.20 | Peptidase_S78 | 2.90E-45 |
| C0AX09 | PpenneriATCC35198 | PF00717.26 | Peptidase_S24 | 2.50E-17 |
| C0AX49 | PpenneriATCC35198 | PF13472.9 | Lipase_GDSL_2 | 2.60E-22 |
| C0AXH2 | PpenneriATCC35198 | PF00756.23 | Esterase | 8.10E-10 |
| C0AXT5 | PpenneriATCC35198 | PF06500.14 | FrsA-like | 1.30E-12 |
| C0AXT6 | PpenneriATCC35198 | PF06500.14 | FrsA-like | 9.10E-84 |
| C0AXT7 | PpenneriATCC35198 | PF06500.14 | FrsA-like | 2.10E-45 |
| C0AYP4 | PpenneriATCC35198 | PF05362.16 | Lon_C | 2.80E-74 |
| C0AYP7 | PpenneriATCC35198 | PF00574.26 | CLP_protease | 2.70E-89 |
| C0AZE4 | PpenneriATCC35198 | PF13365.9 | Trypsin_2 | 1.80E-30 |
| C0AZE5 | PpenneriATCC35198 | PF13365.9 | Trypsin_2 | 7.70E-29 |
| C0B017 | PpenneriATCC35198 | PF03575.20 | Peptidase_S51 | 4.50E-42 |
| C0B0A5 | PpenneriATCC35198 | PF00144.27 | Beta-lactamase | 7.20E-55 |
| C0B0D0 | PpenneriATCC35198 | PF06821.16 | Ser_hydrolase | 1.70E-54 |
| C0B0H4 | PpenneriATCC35198 | PF02113.18 | Peptidase_S13 | 6.10E-146 |
| C0B0P7 | PpenneriATCC35198 | PF03572.21 | Peptidase_S41 | 2.40E-17 |
| C0B0P8 | PpenneriATCC35198 | PF03572.21 | Peptidase_S41 | 6.10E-24 |
| C0B0V8 | PpenneriATCC35198 | PF00717.26 | Peptidase_S24 | 3.30E-27 |
| C0B104 | PpenneriATCC35198 | PF00698.24 | Acyl_transf_1 | 3.10E-39 |
| C0B1C2 | PpenneriATCC35198 | PF05362.16 | Lon_C | 2.30E-10 |
| C0B1Q2 | PpenneriATCC35198 | PF00768.23 | Peptidase_S11 | 1.70E-13 |
| C0B1Q3 | PpenneriATCC35198 | PF00768.23 | Peptidase_S11 | 2.90E-79 |
| C0B2B6 | PpenneriATCC35198 | PF00756.23 | Esterase | 1.80E-21 |
| C0B2B7 | PpenneriATCC35198 | PF00756.23 | Esterase | 1.20E-35 |
| C0B2E5 | PpenneriATCC35198 | PF00768.23 | Peptidase_S11 | 5.70E-100 |
| C0B2G9 | PpenneriATCC35198 | PF12146.11 | Hydrolase_4 | 2.60E-61 |
| C0B2Q3 | PpenneriATCC35198 | PF13354.9 | Beta-lactamase2 | 1.20E-39 |
| C0B323 | PpenneriATCC35198 | PF08840.14 | BAAT_C | 5.90E-06 |
| C0B388 | PpenneriATCC35198 | PF05929.14 | Phage_GPO | 1.80E-105 |
| C0B3Z6 | PpenneriATCC35198 | PF01343.21 | Peptidase_S49 | 1.50E-44 |
| Q5L8L6 | BfragilisATCC25285 | PF00574.26 | CLP_protease | 1.10E-78 |
| Q5LB17 | BfragilisATCC25285 | PF10459.12 | Peptidase_S46 | 1.80E-260 |
| A0A0I9RSQ7 | BfragilisATCC25285 | PF01343.21 | Peptidase_S49 | 2.70E-17 |
| A0A0K6BNH | BfragilisATCC25285 | PF10502.12 | Peptidase_S26 | 7.40E-25 |
| A0A380YP31 | BfragilisATCC25285 | PF01734.25 | Patatin | 8.60E-21 |

| | | | | |
|---|---|---|---|---|
| A0A380YRJ2 | BfragilisATCC25285 | PF00144.27 | Beta-lactamase | 9.10E-40 |
| A0A380YS74 | BfragilisATCC25285 | PF02230.19 | Abhydrolase_2 | 3.40E-11 |
| A0A380YSS8 | BfragilisATCC25285 | PF03572.21 | Peptidase_S41 | 7.50E-17 |
| A0A380YT33 | BfragilisATCC25285 | PF13472.9 | Lipase_GDSL_2 | 3.90E-15 |
| A0A380YT44 | BfragilisATCC25285 | PF00082.25 | Peptidase_S8 | 3.90E-38 |
| A0A380YU49 | BfragilisATCC25285 | PF02016.18 | Peptidase_S66 | 8.70E-36 |
| A0A380YVD1 | BfragilisATCC25285 | PF02674.19 | Colicin_V | 2.40E-31 |
| A0A380YY03 | BfragilisATCC25285 | PF03572.21 | Peptidase_S41 | 5.80E-19 |
| A0A380YZM5 | BfragilisATCC25285 | PF13472.9 | Lipase_GDSL_2 | 3.30E-18 |
| Q5L7F0 | BfragilisATCC25285 | PF03572.21 | Peptidase_S41 | 1.80E-52 |
| Q5L7Q5 | BfragilisATCC25285 | PF01694.25 | Rhomboid | 3.40E-24 |
| Q5L7Q6 | BfragilisATCC25285 | PF01694.25 | Rhomboid | 3.00E-33 |
| Q5L7V4 | BfragilisATCC25285 | PF03572.21 | Peptidase_S41 | 4.60E-48 |
| Q5L832 | BfragilisATCC25285 | PF00326.24 | Peptidase_S9 | 2.20E-07 |
| Q5L832 | BfragilisATCC25285 | PF13472.9 | Lipase_GDSL_2 | 7.20E-27 |
| Q5L8F3 | BfragilisATCC25285 | PF13472.9 | Lipase_GDSL_2 | 1.60E-12 |
| Q5L8P8 | BfragilisATCC25285 | PF00756.23 | Esterase | 1.60E-28 |
| Q5L8Y8 | BfragilisATCC25285 | PF13472.9 | Lipase_GDSL_2 | 2.10E-21 |
| Q5L9E7 | BfragilisATCC25285 | PF01734.25 | Patatin | 2.70E-26 |
| Q5LA76 | BfragilisATCC25285 | PF03629.21 | SASA | 2.10E-09 |
| Q5LAA5 | BfragilisATCC25285 | PF01343.21 | Peptidase_S49 | 3.70E-49 |
| Q5LAE1 | BfragilisATCC25285 | PF00326.24 | Peptidase_S9 | 6.20E-46 |
| Q5LB16 | BfragilisATCC25285 | PF10459.12 | Peptidase_S46 | 2.30E-266 |
| Q5LB38 | BfragilisATCC25285 | PF10502.12 | Peptidase_S26 | 4.20E-32 |
| Q5LB79 | BfragilisATCC25285 | PF01734.25 | Patatin | 2.50E-25 |
| Q5LBI9 | BfragilisATCC25285 | PF00975.23 | Thioesterase | 2.80E-07 |
| Q5LBR2 | BfragilisATCC25285 | PF13365.9 | Trypsin_2 | 1.50E-32 |
| Q5LC39 | BfragilisATCC25285 | PF00326.24 | Peptidase_S9 | 7.30E-58 |
| Q5LCE8 | BfragilisATCC25285 | PF03572.21 | Peptidase_S41 | 7.10E-20 |
| Q5LCH5 | BfragilisATCC25285 | PF01734.25 | Patatin | 6.80E-27 |
| Q5LCS2 | BfragilisATCC25285 | PF05362.16 | Lon_C | 2.70E-81 |
| Q5LCV0 | BfragilisATCC25285 | PF03572.21 | Peptidase_S41 | 1.20E-17 |
| Q5LCW3 | BfragilisATCC25285 | PF00698.24 | Acyl_transf_1 | 1.60E-37 |
| Q5LCX6 | BfragilisATCC25285 | PF01734.25 | Patatin | 2.80E-21 |
| Q5LD72 | BfragilisATCC25285 | PF13472.9 | Lipase_GDSL_2 | 6.60E-19 |
| Q5LD97 | BfragilisATCC25285 | PF00326.24 | Peptidase_S9 | 7.40E-12 |
| Q5LDF9 | BfragilisATCC25285 | PF03572.21 | Peptidase_S41 | 9.90E-15 |
| Q5LDG9 | BfragilisATCC25285 | PF02129.21 | Peptidase_S15 | 8.80E-10 |
| Q5LE26 | BfragilisATCC25285 | PF00717.26 | Peptidase_S24 | 1.00E-24 |
| Q5LEE3 | BfragilisATCC25285 | PF03629.21 | SASA | 6.60E-22 |
| Q5LEE3 | BfragilisATCC25285 | PF13472.9 | Lipase_GDSL_2 | 1.30E-24 |
| Q5LFN9 | BfragilisATCC25285 | PF00756.23 | Esterase | 3.00E-28 |
| Q5LG20 | BfragilisATCC25285 | PF13354.9 | Beta-lactamase2 | 2.40E-30 |
| Q5LGS7 | BfragilisATCC25285 | PF01694.25 | Rhomboid | 5.00E-13 |
| Q5LGU5 | BfragilisATCC25285 | PF00326.24 | Peptidase_S9 | 1.20E-56 |
| Q5LGW2 | BfragilisATCC25285 | PF03572.21 | Peptidase_S41 | 8.80E-51 |
| Q5LH06 | BfragilisATCC25285 | PF00326.24 | Peptidase_S9 | 2.00E-16 |
| Q5LH43 | BfragilisATCC25285 | PF12146.11 | Hydrolase_4 | 2.30E-14 |
| Q5LHV2 | BfragilisATCC25285 | PF05576.14 | Peptidase_S37 | 5.80E-61 |
| Q5LHV8 | BfragilisATCC25285 | PF00144.27 | Beta-lactamase | 7.10E-50 |
| Q5LHV9 | BfragilisATCC25285 | PF00144.27 | Beta-lactamase | 4.60E-53 |
| Q5LI90 | BfragilisATCC25285 | PF00144.27 | Beta-lactamase | 2.00E-60 |

| | | | | |
|---|---|---|---|---|
| Q5LI90 | BfragilisATCC25285 | PF13472.9 | Lipase_GDSL_2 | 1.20E-15 |
| Q5LIA5 | BfragilisATCC25285 | PF03572.21 | Peptidase_S41 | 1.20E-33 |
| Q5LIV3 | BfragilisATCC25285 | PF10502.12 | Peptidase_S26 | 2.40E-27 |
| Q5LIW7 | BfragilisATCC25285 | PF10459.12 | Peptidase_S46 | 3.30E-268 |
| Q5LJ01 | BfragilisATCC25285 | PF00326.24 | Peptidase_S9 | 2.00E-44 |
| Q5LJ18 | BfragilisATCC25285 | PF03572.21 | Peptidase_S41 | 1.40E-18 |
| Q5LJ73 | BfragilisATCC25285 | PF02113.18 | Peptidase_S13 | 2.80E-80 |
| A1A2G3 | BadolescentisATCC15703 | PF00717.26 | Peptidase_S24 | 1.90E-31 |
| A0ZZE3 | BadolescentisATCC15703 | PF00326.24 | Peptidase_S9 | 3.50E-40 |
| A0ZZE6 | BadolescentisATCC15703 | PF12146.11 | Hydrolase_4 | 4.30E-39 |
| A0ZZF5 | BadolescentisATCC15703 | PF13365.9 | Trypsin_2 | 2.00E-31 |
| A0ZZI1 | BadolescentisATCC15703 | PF03629.21 | SASA | 2.10E-08 |
| A1A004 | BadolescentisATCC15703 | PF00698.24 | Acyl_transf_1 | 4.20E-36 |
| A1A050 | BadolescentisATCC15703 | PF00135.31 | COesterase | 1.10E-86 |
| A1A0F5 | BadolescentisATCC15703 | PF02113.18 | Peptidase_S13 | 4.20E-38 |
| A1A0H7 | BadolescentisATCC15703 | PF03629.21 | SASA | 9.50E-13 |
| A1A0H8 | BadolescentisATCC15703 | PF00135.31 | COesterase | 8.00E-79 |
| A1A0V3 | BadolescentisATCC15703 | PF06259.15 | Abhydrolase_8 | 6.70E-05 |
| A1A0Y5 | BadolescentisATCC15703 | PF00574.26 | CLP_protease | 1.30E-70 |
| A1A0Y6 | BadolescentisATCC15703 | PF00574.26 | CLP_protease | 7.50E-75 |
| A1A124 | BadolescentisATCC15703 | PF00756.23 | Esterase | 2.40E-15 |
| A1A2K9 | BadolescentisATCC15703 | PF05362.16 | Lon_C | 4.90E-12 |
| A1A2L7 | BadolescentisATCC15703 | PF01734.25 | Patatin | 1.00E-15 |
| A1A306 | BadolescentisATCC15703 | PF13354.9 | Beta-lactamase2 | 1.70E-32 |
| A1A307 | BadolescentisATCC15703 | PF13354.9 | Beta-lactamase2 | 1.00E-05 |
| A1A340 | BadolescentisATCC15703 | PF00326.24 | Peptidase_S9 | 1.70E-52 |
| A1A345 | BadolescentisATCC15703 | PF12697.10 | Abhydrolase_6 | 1.60E-19 |
| A1A3F6 | BadolescentisATCC15703 | PF01425.24 | Amidase | 4.50E-149 |
| A1A3L1 | BadolescentisATCC15703 | PF03629.21 | SASA | 4.90E-28 |
| A1A3S7 | BadolescentisATCC15703 | PF02230.19 | Abhydrolase_2 | 4.20E-18 |
| U2BDQ5 | CsymbiosumATCC14940 | PF13354.9 | Beta-lactamase2 | 5.40E-64 |
| U2BIR6 | CsymbiosumATCC14940 | PF05580.15 | Peptidase_S55 | 7.50E-84 |
| U2BJR4 | CsymbiosumATCC14940 | PF03572.21 | Peptidase_S41 | 1.70E-22 |
| U2BND4 | CsymbiosumATCC14940 | PF05362.16 | Lon_C | 1.10E-83 |
| U2BNP7 | CsymbiosumATCC14940 | PF13472.9 | Lipase_GDSL_2 | 8.20E-17 |
| U2BNU7 | CsymbiosumATCC14940 | PF12146.11 | Hydrolase_4 | 1.20E-46 |
| U2BPV6 | CsymbiosumATCC14940 | PF10502.12 | Peptidase_S26 | 3.80E-20 |
| U2BQ68 | CsymbiosumATCC14940 | PF00768.23 | Peptidase_S11 | 3.60E-43 |
| U2BQF5 | CsymbiosumATCC14940 | PF00768.23 | Peptidase_S11 | 6.60E-52 |
| U2BS34 | CsymbiosumATCC14940 | PF12146.11 | Hydrolase_4 | 1.30E-15 |
| U2BS82 | CsymbiosumATCC14940 | PF02674.19 | Colicin_V | 6.10E-13 |
| U2BT47 | CsymbiosumATCC14940 | PF07859.16 | Abhydrolase_3 | 5.60E-56 |
| U2BTA7 | CsymbiosumATCC14940 | PF02016.18 | Peptidase_S66 | 8.10E-26 |
| U2BYP6 | CsymbiosumATCC14940 | PF02129.21 | Peptidase_S15 | 2.40E-10 |
| U2BZK6 | CsymbiosumATCC14940 | PF00768.23 | Peptidase_S11 | 2.00E-54 |
| U2C153 | CsymbiosumATCC14940 | PF07859.16 | Abhydrolase_3 | 9.30E-11 |
| U2C1H1 | CsymbiosumATCC14940 | PF01734.25 | Patatin | 1.60E-10 |
| U2C2X1 | CsymbiosumATCC14940 | PF00082.25 | Peptidase_S8 | 5.00E-18 |
| U2C336 | CsymbiosumATCC14940 | PF00082.25 | Peptidase_S8 | 3.20E-11 |
| U2C3Z0 | CsymbiosumATCC14940 | PF10502.12 | Peptidase_S26 | 8.90E-31 |
| U2C414 | CsymbiosumATCC14940 | PF00768.23 | Peptidase_S11 | 4.20E-48 |
| U2C5E0 | CsymbiosumATCC14940 | PF13354.9 | Beta-lactamase2 | 1.20E-63 |

| | | | | |
|---|---|---|---|---|
| U2C5P4 | CsymbiosumATCC14940 | PF03572.21 | Peptidase_S41 | 7.10E-52 |
| U2C665 | CsymbiosumATCC14940 | PF10502.12 | Peptidase_S26 | 2.10E-43 |
| U2C6W1 | CsymbiosumATCC14940 | PF12697.10 | Abhydrolase_6 | 2.70E-06 |
| U2C7Z6 | CsymbiosumATCC14940 | PF01734.25 | Patatin | 1.30E-20 |
| U2C8N6 | CsymbiosumATCC14940 | PF12146.11 | Hydrolase_4 | 6.70E-27 |
| U2C989 | CsymbiosumATCC14940 | PF10502.12 | Peptidase_S26 | 3.70E-41 |
| U2CYP7 | CsymbiosumATCC14940 | PF00717.26 | Peptidase_S24 | 5.40E-32 |
| U2D1C7 | CsymbiosumATCC14940 | PF00574.26 | CLP_protease | 1.40E-86 |
| U2D1I3 | CsymbiosumATCC14940 | PF00574.26 | CLP_protease | 6.00E-19 |
| U2D7B4 | CsymbiosumATCC14940 | PF00082.25 | Peptidase_S8 | 7.70E-55 |
| U2D968 | CsymbiosumATCC14940 | PF03629.21 | SASA | 5.90E-52 |
| U2D9D4 | CsymbiosumATCC14940 | PF07859.16 | Abhydrolase_3 | 2.50E-50 |
| U2DAD2 | CsymbiosumATCC14940 | PF00698.24 | Acyl_transf_1 | 2.60E-37 |
| U2DAZ5 | CsymbiosumATCC14940 | PF12697.10 | Abhydrolase_6 | 4.30E-08 |
| U2DBF5 | CsymbiosumATCC14940 | PF00768.23 | Peptidase_S11 | 1.10E-69 |
| U2DBK6 | CsymbiosumATCC14940 | PF00082.25 | Peptidase_S8 | 1.10E-53 |
| U2DD62 | CsymbiosumATCC14940 | PF04586.20 | Peptidase_S78 | 1.00E-10 |
| U2DDJ1 | CsymbiosumATCC14940 | PF01694.25 | Rhomboid | 6.80E-35 |
| U2DDQ9 | CsymbiosumATCC14940 | PF00082.25 | Peptidase_S8 | 8.30E-14 |
| U2DDR4 | CsymbiosumATCC14940 | PF13365.9 | Trypsin_2 | 1.10E-27 |
| U2DGI3 | CsymbiosumATCC14940 | PF13365.9 | Trypsin_2 | 4.20E-10 |
| U2DGK0 | CsymbiosumATCC14940 | PF00082.25 | Peptidase_S8 | 2.50E-16 |
| U2DHH1 | CsymbiosumATCC14940 | PF10502.12 | Peptidase_S26 | 8.00E-29 |
| U2DHN2 | CsymbiosumATCC14940 | PF01425.24 | Amidase | 3.00E-153 |
| U2DJU3 | CsymbiosumATCC14940 | PF00768.23 | Peptidase_S11 | 9.00E-52 |
| A0A412YUG | EbolteaeATCCBAA613 | PF03572.21 | Peptidase_S41 | 4.20E-49 |
| A0A412YXW | EbolteaeATCCBAA613 | PF02674.19 | Colicin_V | 2.40E-11 |
| A0A412YYF | EbolteaeATCCBAA613 | PF00082.25 | Peptidase_S8 | 2.10E-50 |
| A0A412Z5S | EbolteaeATCCBAA613 | PF00768.23 | Peptidase_S11 | 1.70E-48 |
| A0A412Z5S | EbolteaeATCCBAA613 | PF00768.23 | Peptidase_S11 | 3.80E-51 |
| A0A412Z6C | EbolteaeATCCBAA613 | PF10502.12 | Peptidase_S26 | 1.30E-33 |
| A0A412Z6F | EbolteaeATCCBAA613 | PF10502.12 | Peptidase_S26 | 1.20E-44 |
| A0A412Z7U | EbolteaeATCCBAA613 | PF10502.12 | Peptidase_S26 | 1.80E-42 |
| A0A412Z7X | EbolteaeATCCBAA613 | PF05580.15 | Peptidase_S55 | 1.20E-83 |
| A0A412Z827 | EbolteaeATCCBAA613 | PF07859.16 | Abhydrolase_3 | 5.50E-52 |
| A0A412ZFE | EbolteaeATCCBAA613 | PF13472.9 | Lipase_GDSL_2 | 5.10E-18 |
| A0A414AEZ | EbolteaeATCCBAA613 | PF13365.9 | Trypsin_2 | 1.50E-10 |
| A0A414AL79 | EbolteaeATCCBAA613 | PF00082.25 | Peptidase_S8 | 5.10E-13 |
| A0A414ANF | EbolteaeATCCBAA613 | PF00975.23 | Thioesterase | 4.10E-29 |
| A0A414ARK | EbolteaeATCCBAA613 | PF01694.25 | Rhomboid | 4.10E-34 |
| A0A414AWB | EbolteaeATCCBAA613 | PF07859.16 | Abhydrolase_3 | 1.30E-56 |
| A0A414AXV | EbolteaeATCCBAA613 | PF00574.26 | CLP_protease | 6.10E-21 |
| A0A414AYY | EbolteaeATCCBAA613 | PF05362.16 | Lon_C | 1.60E-88 |
| A0A414AYZ | EbolteaeATCCBAA613 | PF00574.26 | CLP_protease | 2.30E-85 |
| A0A414AZM | EbolteaeATCCBAA613 | PF01734.25 | Patatin | 5.80E-23 |
| A0A414AZT | EbolteaeATCCBAA613 | PF12697.10 | Abhydrolase_6 | 6.80E-15 |
| A0A414B0N | EbolteaeATCCBAA613 | PF02016.18 | Peptidase_S66 | 3.60E-35 |
| A0A6M5G6 | EbolteaeATCCBAA613 | PF00717.26 | Peptidase_S24 | 6.20E-31 |
| A0A6N2XPI1 | EbolteaeATCCBAA613 | PF13354.9 | Beta-lactamase2 | 1.90E-34 |
| A0A855Q3M | EbolteaeATCCBAA613 | PF01425.24 | Amidase | 5.30E-155 |
| A0A855Q98 | EbolteaeATCCBAA613 | PF00768.23 | Peptidase_S11 | 1.50E-43 |
| A0A8D4EIZ1 | EbolteaeATCCBAA613 | PF00326.24 | Peptidase_S9 | 4.50E-33 |

| | | | | |
|---|---|---|---|---|
| A0A8D4EJE | EbolteaeATCCBAA613 | PF00326.24 | Peptidase_S9 | 6.70E-44 |
| A0A8D4EKD | EbolteaeATCCBAA613 | PF00082.25 | Peptidase_S8 | 1.00E-53 |
| A0A8D4EKE | EbolteaeATCCBAA613 | PF12146.11 | Hydrolase_4 | 2.40E-09 |
| A0A8D4ELI5 | EbolteaeATCCBAA613 | PF00975.23 | Thioesterase | 5.60E-08 |
| A0A8D4ELM | EbolteaeATCCBAA613 | PF06500.14 | FrsA-like | 5.90E-17 |
| A0A8D4ELM | EbolteaeATCCBAA613 | PF00768.23 | Peptidase_S11 | 4.20E-57 |
| A0A8D4ELY | EbolteaeATCCBAA613 | PF13472.9 | Lipase_GDSL_2 | 3.20E-14 |
| A0A8D4EP4 | EbolteaeATCCBAA613 | PF00574.26 | CLP_protease | 1.80E-23 |
| A0A8D4EPH | EbolteaeATCCBAA613 | PF00082.25 | Peptidase_S8 | 2.10E-17 |
| A0A8D4EQ1 | EbolteaeATCCBAA613 | PF01734.25 | Patatin | 3.10E-19 |
| A0A8D4EQ | EbolteaeATCCBAA613 | PF12146.11 | Hydrolase_4 | 7.40E-16 |
| A0A8D4ERA | EbolteaeATCCBAA613 | PF01734.25 | Patatin | 1.90E-17 |
| A0A8D4ES7 | EbolteaeATCCBAA613 | PF00144.27 | Beta-lactamase | 4.90E-35 |
| A0A8D4ESC | EbolteaeATCCBAA613 | PF04586.20 | Peptidase_S78 | 1.90E-23 |
| A0A8D4ESF | EbolteaeATCCBAA613 | PF13365.9 | Trypsin_2 | 7.50E-29 |
| A0A8D4JF7 | EbolteaeATCCBAA613 | PF00768.23 | Peptidase_S11 | 1.10E-55 |
| A0A8D4JFP | EbolteaeATCCBAA613 | PF00768.23 | Peptidase_S11 | 1.90E-72 |
| A0A8D4JFV | EbolteaeATCCBAA613 | PF00698.24 | Acyl_transf_1 | 2.60E-39 |
| A0A8D4JGH | EbolteaeATCCBAA613 | PF12146.11 | Hydrolase_4 | 5.60E-12 |
| A0A8D4JGQ | EbolteaeATCCBAA613 | PF13365.9 | Trypsin_2 | 9.30E-29 |
| A0A8D4JGT | EbolteaeATCCBAA613 | PF00574.26 | CLP_protease | 4.80E-79 |
| A0A8D4JJ64 | EbolteaeATCCBAA613 | PF12697.10 | Abhydrolase_6 | 3.00E-08 |
| A0A8D4JJ7 | EbolteaeATCCBAA613 | PF00144.27 | Beta-lactamase | 1.90E-25 |
| A0A858B376 | CaerofaciensATCC25986 | PF00717.26 | Peptidase_S24 | 5.60E-08 |
| A0A858B3J6 | CaerofaciensATCC25986 | PF00574.26 | CLP_protease | 2.60E-80 |
| A0A858B4K | CaerofaciensATCC25986 | PF12697.10 | Abhydrolase_6 | 2.40E-16 |
| A0A858B4M | CaerofaciensATCC25986 | PF13354.9 | Beta-lactamase2 | 4.60E-19 |
| A0A858B4T | CaerofaciensATCC25986 | PF13354.9 | Beta-lactamase2 | 1.90E-64 |
| A0A858B5Z | CaerofaciensATCC25986 | PF00717.26 | Peptidase_S24 | 8.60E-29 |
| A0A858B6C | CaerofaciensATCC25986 | PF10502.12 | Peptidase_S26 | 1.20E-14 |
| A0A858B6R | CaerofaciensATCC25986 | PF00768.23 | Peptidase_S11 | 9.20E-43 |
| A0A858B766 | CaerofaciensATCC25986 | PF00082.25 | Peptidase_S8 | 5.80E-15 |
| A0A858B7G | CaerofaciensATCC25986 | PF13472.9 | Lipase_GDSL_2 | 1.70E-24 |
| A0A858B7J2 | CaerofaciensATCC25986 | PF00756.23 | Esterase | 2.50E-07 |
| A4E6R7 | CaerofaciensATCC25986 | PF03575.20 | Peptidase_S51 | 2.80E-44 |
| A4E7D0 | CaerofaciensATCC25986 | PF13472.9 | Lipase_GDSL_2 | 8.90E-17 |
| A4E7Q8 | CaerofaciensATCC25986 | PF00574.26 | CLP_protease | 5.30E-44 |
| A4E845 | CaerofaciensATCC25986 | PF01734.25 | Patatin | 4.90E-14 |
| A4E8L2 | CaerofaciensATCC25986 | PF01734.25 | Patatin | 1.60E-08 |
| A4E8M4 | CaerofaciensATCC25986 | PF01425.24 | Amidase | 1.50E-139 |
| A4E8P7 | CaerofaciensATCC25986 | PF13365.9 | Trypsin_2 | 1.40E-33 |
| A4E9I4 | CaerofaciensATCC25986 | PF00698.24 | Acyl_transf_1 | 4.60E-36 |
| A4EB38 | CaerofaciensATCC25986 | PF13365.9 | Trypsin_2 | 4.00E-09 |
| A4EBJ9 | CaerofaciensATCC25986 | PF12146.11 | Hydrolase_4 | 7.80E-07 |
| A4EBS2 | CaerofaciensATCC25986 | PF07859.16 | Abhydrolase_3 | 8.50E-44 |
| A4ECI5 | CaerofaciensATCC25986 | PF10502.12 | Peptidase_S26 | 6.00E-45 |
| A4ECP1 | CaerofaciensATCC25986 | PF05448.15 | AXE1 | 1.50E-11 |
| A4ECP2 | CaerofaciensATCC25986 | PF05448.15 | AXE1 | 4.80E-81 |
