## Supplemental Table S4 for "Chemoproteomic identification of a dipeptidyl peptidase 4 (DPP4) homolog in *Bacteroides thetaiotaomicron* important for envelope integrity and fitness"

Table S4: FP-reactive enzymes in B. thetaiotaomicron VPI-5482

| UniProt ID | Probe/DMSO Fold Change | Probe/DMSO t-test significance | Probe/Competition t-test significance | Gene name | Protein name | Pfam domain(s) | Interpro domains | Gene ontology (molecular function) |
|---|---|---|---|---|---|---|---|---|
| Q8A051_BACTN | 4.80738 | + | + | BT4170 | Putative pectate lyase L | PF16380; | IPR032153;IPR012334;IPR011050; | lyase activity [GO:0016829] |
| Q8A0E4_BACTN | 3.44965 | + | + | BT4077 | Endo-1,4-beta-xylanase Z | PF00756; | IPR029058;IPR000801; | hydrolase activity, acting on glycosyl bonds [GO:0016798]; transferase activity, transferring acyl groups other than amino-acyl groups [GO:0016747] |
| Q8A343_BACTN | 4.29238 | + | + | BT3112 | Putative lipoprotein | PF12146; | IPR029058;IPR022742; | carboxylic ester hydrolase activity [GO:0052689] |
| Q8A900_BACTN | 3.98469 | + | + | BT1017 | Uncharacterized protein | PF12875; | IPR029058;IPR024284; | |
| Q8A901_BACTN | 3.78534 | + | + | BT1016 | Conserved protein with a conserved patatin-like phospholipase domain | PF01734; | IPR016035;IPR002641; | hydrolase activity [GO:0016787] |
| Q8A9B9_BACTN | 4.46885 | + | + | BT0896 | Putative patatin-like phospholipase | PF01734; | IPR016035;IPR002641; | hydrolase activity [GO:0016787] |
| Q8A007_BACTN | 4.36943 | + | | BT4214 | Hydrolase of the alpha/beta superfamily | PF00326; | IPR029058;IPR001375; | serine-type peptidase activity [GO:0008236] |
| Q8A028_BACTN | 4.55252 | + | | BT4193 | Dipeptidyl peptidase IV | PF00930;PF00326; | IPR029058;IPR001375;IPR002469;IPR038554; | serine-type peptidase activity [GO:0008236] |
| Q8A048_BACTN | 1.78861 | + | | BT4173 | Putative rhamnogalacturonan acetylesterase | PF13472; | IPR037459;IPR013830;IPR036514; | carboxylic ester hydrolase activity [GO:0052689] |
| Q8A0D0_BACTN | 4.22505 | + | | BT4091 | Sialic acid-specific 9-O-acetylesterase | PF02837;PF03629; | IPR008979;IPR006104;IPR013783;IPR005181;IPR036514;IPR039329; | hydrolase activity, hydrolyzing O-glycosyl compounds [GO:0004553]; sialate O-acetylesterase activity [GO:0001681] |
| Q8A1Y1_BACTN | 4.88806 | + | | BT3527 | Alpha-1,2-mannosidase | PF07971;PF17678; | IPR008928;IPR005887;IPR014718;IPR041371;IPR012939; | carbohydrate binding [GO:0030246]; peptide-N4-(N-acetyl-beta-glucosaminyl)asparagine amidase activity [GO:0000224] |
| Q8A2L6_BACTN | 2.9393 | + | | BT3289 | Dipeptidyl-peptidase | PF10459; | IPR019500;IPR009003; | dipeptidyl-peptidase activity [GO:0008239]; serine-type aminopeptidase activity [GO:0070009] |
| Q8A2L9_BACTN | 6.02334 | + | | BT3286 | Phospholipase/Carboxylesterase | PF02230; | IPR029058;IPR003140; | hydrolase activity [GO:0016787] |
| Q8A2Q1_BACTN | 3.34744 | + | | BT3254 | Dipeptidyl peptidase IV | PF00930;PF00326; | IPR029058;IPR001375;IPR002469;IPR038554; | serine-type peptidase activity [GO:0008236] |
| Q8A371_BACTN | 3.21804 | + | | BT3084 | Lipase, putative | PF07859; | IPR029058;IPR013094; | hydrolase activity [GO:0016787] |
| Q8A3J3_BACTN | 5.48393 | + | | BT2961 | Putative lipase | PF13472; | IPR013830;IPR036514; | |
| Q8A4P8_BACTN | 3.18768 | + | | BT2549 | Putative carboxypeptidase | PF02016;PF17676; | IPR027461;IPR029062;IPR027478;IPR040449;IPR040921;IPR003507; | carboxypeptidase activity [GO:0004180] |
| Q8A6K2_BACTN | 4.20797 | + | | BT1879 | Protease IV | PF01343; | IPR029045;IPR004634;IPR004635;IPR002142; | peptidase activity [GO:0008233] |
| Q8A6N9_BACTN | 3.83853 | + | | BT1838 | Putative alanyl dipeptidyl peptidase | PF00326; | IPR011042;IPR029058;IPR001375; | serine-type peptidase activity [GO:0008236] |
| Q8A6Z2_BACTN | 1.99256 | + | | BT1733 | Phosphoribosylformylglycinamidine synthase | PF02769;PF18072;PF18076; | IPR029062;IPR040707;IPR017926;IPR010073;IPR041609;IPR010918;IPR036676;IPR036921;IPR036604; | ATP binding [GO:0005524]; metal ion binding [GO:0046872]; phosphoribosylformylglycinamidine synthase activity [GO:0004642] |
| Q8A7W6_BACTN | 2.90411 | + | | BT1408 | Tricorn protease homolog | PF07676;PF03572;PF14684; | IPR011042;IPR029045;IPR011659;IPR036034;IPR005151;IPR028204;IPR012393; | serine-type peptidase activity [GO:0008236] |
| Q8A8H5_BACTN | 2.44566 | + | | BT1192 | Putative xylanase | PF13472;PF00326; | IPR029058;IPR001375;IPR013830;IPR036514; | hydrolase activity, acting on glycosyl bonds [GO:0016798]; serine-type peptidase activity [GO:0008236] |
| Q8A8T0_BACTN | 1.69331 | + | | BT1087 | Putative xylanase | PF07859;PF07313; | IPR029058;IPR013094;IPR010846;IPR038765; | hydrolase activity, acting on glycosyl bonds [GO:0016798] |
| Q8AB09_BACTN | 3.3229 | + | | BT0303 | Putative patatin-like phospholipase | PF01734; | IPR016035;IPR002641; | hydrolase activity [GO:0016787] |
| Q8AB73_BACTN | 5.49573 | + | | BT0237 | Dipeptidyl-peptidase | PF10459; | IPR019500;IPR009003; | dipeptidyl-peptidase activity [GO:0008239]; serine-type aminopeptidase activity [GO:0070009] |
| Q8AB74_BACTN | 2.74372 | + | | BT0236 | Dipeptidyl-peptidase | PF10459; | IPR019500;IPR009003; | dipeptidyl-peptidase activity [GO:0008239]; serine-type aminopeptidase activity [GO:0070009] |
| Q8ABF8_BACTN | 4.89907 | + | | BT0152 | Acetyl esterase (Acetylxylosidase) | PF00756; | IPR029058;IPR000801; | transferase activity, transferring acyl groups other than amino-acyl groups |

**Key for Pfam domains**
Definite serine hydrolase
Found in serine hydrolases but not catalytic
Not a serine hydrolase
