## Supplemental Table S5 for "Chemoproteomic identification of a dipeptidyl peptidase 4 (DPP4) homolog in *Bacteroides thetaiotaomicron* important for envelope integrity and fitness"

Table S5: Protein-BLAST results for B. thetaiotaomicron serine hydrolases in HMP gut reference isolates

| Query sequence | Target sequence | % identity | % coverage | E-value | Bit score | Target sequence description |
|---|---|---|---|---|---|---|
| Q8AB73 | BSIG_0060 | 100 | 100 | 0 | 1442 | hypothetical protein [Bacteroides sp. 1_1_6] |
| Q8AB73 | HMPREF9007_01172 | 99.568 | 100 | 0 | 1437 | hypothetical protein [Bacteroides sp. 1_1_14] |
| Q8AB73 | BACFIN_07388 | 91.655 | 100 | 0 | 1344 | hypothetical protein [Bacteroides finegoldii DSM 17565] |
| Q8AB73 | BSGG_0735 | 91.079 | 100 | 0 | 1332 | hypothetical protein [Bacteroides sp. D2] |
| Q8AB73 | HMPREF1017_03509 | 91.079 | 100 | 0 | 1332 | hypothetical protein [Bacteroides ovatus 3_8_47FAA] |
| Q8AB73 | BACCAC_02237 | 90.647 | 100 | 0 | 1332 | hypothetical protein [Bacteroides caccae ATCC 43185] |
| Q8AB73 | CUY_4106 | 91.079 | 100 | 0 | 1332 | conserved hypothetical protein [Bacteroides ovatus SD CMC 3f] |
| Q8AB73 | HMPREF0106_04191 | 90.791 | 100 | 0 | 1331 | hypothetical protein [Bacteroides sp. D22] |
| Q8AB73 | HMPREF9010_01792 | 90.935 | 100 | 0 | 1331 | hypothetical protein [Bacteroides sp. 3_1_23] |
| Q8AB73 | BXY_42310 | 90.647 | 100 | 0 | 1331 | dipeptidyl-peptidase 7. Serine peptidase. MEROPS family S46 [Bacteroides xylanisolvens XB1A] |
| Q8AB73 | HMPREF0127_02920 | 90.504 | 100 | 0 | 1330 | hypothetical protein [Bacteroides sp. 1_1_30] |
| Q8AB73 | HMPREF0102_02504 | 90.647 | 100 | 0 | 1330 | hypothetical protein [Bacteroides sp. 2_1_22] |
| Q8AB73 | BSAG_01080 | 90.647 | 100 | 0 | 1330 | hypothetical protein [Bacteroides sp. D1] |
| Q8AB73 | CW3_4226 | 90.647 | 100 | 0 | 1330 | conserved hypothetical protein [Bacteroides xylanisolvens SD CC 1b] |
| Q8AB73 | CW1_0171 | 90.647 | 100 | 0 | 1330 | conserved hypothetical protein [Bacteroides xylanisolvens SD CC 2a] |
| Q8AB73 | BSCG_00818 | 90.935 | 100 | 0 | 1328 | hypothetical protein [Bacteroides sp. 2_2_4] |
| Q8AB73 | BACOVA_00386 | 90.791 | 100 | 0 | 1327 | hypothetical protein [Bacteroides ovatus ATCC 8483] |
| Q8AB73 | BSHG_1934 | 85.94 | 100 | 0 | 1264 | hypothetical protein [Bacteroides sp. 3_2_5] |
| Q8AB73 | HMPREF1018_03521 | 85.94 | 100 | 0 | 1264 | hypothetical protein [Bacteroides sp. 2_1_56FAA] |
| Q8AB73 | HMPREF0101_03127 | 85.94 | 100 | 0 | 1264 | hypothetical protein [Bacteroides sp. 2_1_16] |
| Q8AB73 | BFAG_03217 | 84.792 | 100 | 0 | 1251 | hypothetical protein [Bacteroides fragilis 3_1_12] |
| Q8AB73 | HMPREF1214_03868 | 83.501 | 100 | 0 | 1219 | hypothetical protein [Bacteroides sp. HPS0048] |
| Q8AB73 | HMPREF1532_00292 | 83.214 | 100 | 0 | 1215 | hypothetical protein [Bacteroides salyersiae WAL 10018 = DSM 18765 = JCM 12988] |
| Q8AB73 | BACINT_02882 | 79.627 | 100 | 0 | 1154 | hypothetical protein [Bacteroides intestinalis DSM 17393] |
| Q8AB73 | HMPREF9447_03715 | 79.053 | 100 | 0 | 1152 | hypothetical protein [Bacteroides oleiciplenus YIT 12058] |
| Q8AB73 | BACCELL_05142 | 79.627 | 100 | 0 | 1145 | hypothetical protein [Bacteroides cellulosilyticus DSM 14838] |
| Q8AB73 | BACEGG_02070 | 74.175 | 100 | 0 | 1073 | hypothetical protein [Bacteroides eggerthii DSM 20697] |
| Q8AB73 | BACSTE_01784 | 74.032 | 100 | 0 | 1071 | hypothetical protein [Bacteroides stercoris ATCC 43183] |
| Q8AB73 | HMPREF1016_00219 | 73.601 | 100 | 0 | 1062 | peptidase S46 [Bacteroides eggerthii 1_2_48FAA] |
| Q8AB73 | HMPREF9445_00327 | 73.171 | 100 | 0 | 1058 | hypothetical protein [Bacteroides clarus YIT 12056] |
| Q8AB73 | HMPREF0969_00399 | 71.306 | 100 | 0 | 1031 | hypothetical protein [Bacteroides sp. D20] |
| Q8AB73 | BACUNI_01423 | 71.306 | 100 | 0 | 1031 | hypothetical protein [Bacteroides uniformis ATCC 8492] |
| Q8AB73 | HMPREF1007_01538 | 71.306 | 100 | 0 | 1031 | peptidase S46 [Bacteroides sp. 4_1_36] |
| Q8AB73 | HMPREF9446_00758 | 71.593 | 100 | 0 | 1031 | hypothetical protein [Bacteroides fluxus YIT 12057] |
| Q8AB73 | BSFG_01936 | 71.306 | 100 | 0 | 1019 | hypothetical protein [Bacteroides sp. 4_3_47FAA] |
| Q8AB73 | HMPREF9011_02238 | 71.306 | 100 | 0 | 1019 | hypothetical protein [Bacteroides sp. 3_1_40A] |
| Q8AB73 | HMPREF0105_0857 | 71.019 | 100 | 0 | 1017 | hypothetical protein [Bacteroides sp. 3_1_33FAA] |
| Q8AB73 | BSEG_00914 | 71.019 | 100 | 0 | 1017 | hypothetical protein [Bacteroides dorei 5_1_36/D4] |
| Q8AB73 | BSBG_02476 | 71.019 | 100 | 0 | 1017 | hypothetical protein [Bacteroides sp. 9_1_42FAA] |
| Q8AB73 | CUU_0388 | 71.162 | 100 | 0 | 1016 | conserved hypothetical protein [Bacteroides vulgatus PC510] |
| Q8AB73 | BACDOR_03540 | 70.732 | 100 | 0 | 1013 | hypothetical protein [Bacteroides dorei DSM 17855] |
| Q8AB73 | BACCOP_03276 | 66.858 | 100 | 0 | 989 | hypothetical protein [Bacteroides coprocola DSM 17136] |
| Q8AB73 | BACPLE_01878 | 66.952 | 100 | 0 | 983 | hypothetical protein [Bacteroides plebeius DSM 17135] |
| Q8AB73 | BACCOPRO_02783 | 64.849 | 100 | 0 | 964 | hypothetical protein [Bacteroides coprophilus DSM 18228 = JCM 13818] |
| Q8AB73 | HMPREF1033_00928 | 53.228 | 100 | 0 | 770 | hypothetical protein [Tannerella sp. 6_1_58FAA_CT1] |
| Q8AB73 | BACCOPRO_02782 | 48.997 | 99 | 0 | 706 | hypothetical protein [Bacteroides coprophilus DSM 18228 = JCM 13818] |
| Q8AB73 | HMPREF0105_0858 | 50.212 | 100 | 0 | 705 | hypothetical protein [Bacteroides sp. 3_1_33FAA] |
| Q8AB73 | BACDOR_03539 | 50.212 | 100 | 0 | 705 | hypothetical protein [Bacteroides dorei DSM 17855] |
| Q8AB73 | BSEG_00915 | 50.071 | 100 | 0 | 704 | hypothetical protein [Bacteroides dorei 5_1_36/D4] |
| Q8AB73 | BSFG_01937 | 50.212 | 100 | 0 | 704 | hypothetical protein [Bacteroides sp. 4_3_47FAA] |
| Q8AB73 | HMPREF9011_02237 | 50.212 | 100 | 0 | 704 | hypothetical protein [Bacteroides sp. 3_1_40A] |
| Q8AB73 | BSBG_02477 | 50.212 | 100 | 0 | 704 | hypothetical protein [Bacteroides sp. 9_1_42FAA] |
| Q8AB73 | CUU_0387 | 50.071 | 100 | 0 | 703 | conserved hypothetical protein [Bacteroides vulgatus PC510] |
| Q8AB73 | BACPLE_01877 | 49.14 | 99 | 0 | 694 | hypothetical protein [Bacteroides plebeius DSM 17135] |
| Q8AB73 | BACCOP_03275 | 48.854 | 99 | 0 | 689 | hypothetical protein [Bacteroides coprocola DSM 17136] |
| Q8AB73 | BACINT_02883 | 47.949 | 100 | 0 | 683 | hypothetical protein [Bacteroides intestinalis DSM 17393] |
| Q8AB73 | PRABACTJOHN_02704 | 48.355 | 100 | 0 | 678 | hypothetical protein [Parabacteroides johnsonii DSM 18315] |
| Q8AB73 | HMPREF9447_03716 | 47.383 | 100 | 0 | 676 | hypothetical protein [Bacteroides oleiciplenus YIT 12058] |

| | | | | | | |
|---|---|---|---|---|---|---|
| Q8AB73 | BFAG_03218 | 47.653 | 100 | 0 | 676 | hypothetical protein [Bacteroides fragilis 3_1_12] |
| Q8AB73 | BACCAC_02236 | 48.212 | 100 | 0 | 676 | hypothetical protein [Bacteroides caccae ATCC 43185] |
| Q8AB73 | HMPREF1033_02741 | 47.757 | 98 | 0 | 675 | hypothetical protein [Tannerella sp. 6_1_58FAA_CT1] |
| Q8AB73 | PARMER_02946 | 47.708 | 100 | 0 | 674 | hypothetical protein [Parabacteroides merdae ATCC 43184] |
| Q8AB73 | HMPREF1532_00293 | 47.489 | 99 | 0 | 674 | hypothetical protein [Bacteroides salyersiae WAL 10018 = DSM 18765 = JCM 12988] |
| Q8AB73 | HMPREF1214_03869 | 48.069 | 100 | 0 | 673 | hypothetical protein [Bacteroides sp. HPS0048] |
| Q8AB73 | BACCELL_05143 | 47.242 | 100 | 0 | 672 | hypothetical protein [Bacteroides cellulosilyticus DSM 14838] |
| Q8AB73 | HMPREF0969_00398 | 46.714 | 99 | 0 | 670 | hypothetical protein [Bacteroides sp. D20] |
| Q8AB73 | BACUNI_01424 | 46.571 | 99 | 0 | 669 | hypothetical protein [Bacteroides uniformis ATCC 8492] |
| Q8AB73 | HMPREF1007_01539 | 46.571 | 99 | 0 | 669 | peptidase S46 [Bacteroides sp. 4_1_36] |
| Q8AB73 | HMPREF9446_00759 | 47.143 | 99 | 0 | 669 | hypothetical protein [Bacteroides fluxus YIT 12057] |
| Q8AB73 | HMPREF1016_00218 | 46.714 | 99 | 0 | 669 | peptidase S46 [Bacteroides eggerthii 1_2_48FAA] |
| Q8AB73 | BACEGG_02069 | 46.714 | 99 | 0 | 669 | hypothetical protein [Bacteroides eggerthii DSM 20697] |
| Q8AB73 | HMPREF0619_00990 | 48.276 | 99 | 0 | 668 | hypothetical protein [Parabacteroides sp. D13] |
| Q8AB73 | HMPREF9008_01598 | 48.132 | 99 | 0 | 668 | hypothetical protein [Parabacteroides sp. 20_3] |
| Q8AB73 | HMPREF0104_00878 | 48.276 | 99 | 0 | 668 | hypothetical protein [Bacteroides sp. 3_1_19] |
| Q8AB73 | BSHG_1935 | 46.924 | 100 | 0 | 666 | hypothetical protein [Bacteroides sp. 3_2_5] |
| Q8AB73 | HMPREF0103_1442 | 47.994 | 99 | 0 | 665 | hypothetical protein [Bacteroides sp. 2_1_33B] |
| Q8AB73 | BACSTE_01783 | 46.581 | 100 | 0 | 665 | hypothetical protein [Bacteroides stercoris ATCC 43183] |
| Q8AB73 | HMPREF0101_03126 | 46.924 | 100 | 0 | 665 | hypothetical protein [Bacteroides sp. 2_1_16] |
| Q8AB73 | HMPREF1018_03522 | 46.924 | 100 | 0 | 665 | hypothetical protein [Bacteroides sp. 2_1_56FAA] |
| Q8AB73 | HMPREF9445_00326 | 46.571 | 99 | 0 | 664 | hypothetical protein [Bacteroides clarus YIT 12056] |
| Q8AB73 | BSIG_0059 | 47.059 | 99 | 0 | 663 | hypothetical protein [Bacteroides sp. 1_1_6] |
| Q8AB73 | BACFIN_07387 | 46.79 | 100 | 0 | 654 | hypothetical protein [Bacteroides finegoldii DSM 17565] |
| Q8AB73 | HMPREF9010_01793 | 46.857 | 99 | 0 | 654 | hypothetical protein [Bacteroides sp. 3_1_23] |
| Q8AB73 | BACOVA_00385 | 46.857 | 99 | 0 | 654 | hypothetical protein [Bacteroides ovatus ATCC 8483] |
| Q8AB73 | BSGG_0736 | 46.857 | 99 | 0 | 653 | hypothetical protein [Bacteroides sp. D2] |
| Q8AB73 | HMPREF0127_02919 | 46.714 | 99 | 0 | 653 | hypothetical protein [Bacteroides sp. 1_1_30] |
| Q8AB73 | BXY_42300 | 46.714 | 99 | 0 | 653 | dipeptidyl-peptidase 7. Serine peptidase. MEROPS family S46 [Bacteroides xylanisolvens XB1A] |
| Q8AB73 | HMPREF0106_04190 | 46.714 | 99 | 0 | 653 | hypothetical protein [Bacteroides sp. D22] |
| Q8AB73 | BSCG_00819 | 46.857 | 99 | 0 | 653 | hypothetical protein [Bacteroides sp. 2_2_4] |
| Q8AB73 | HMPREF1017_03508 | 46.857 | 99 | 0 | 653 | hypothetical protein [Bacteroides ovatus 3_8_47FAA] |
| Q8AB73 | CUY_4107 | 46.857 | 99 | 0 | 652 | conserved hypothetical protein [Bacteroides ovatus SD CMC 3f] |
| Q8AB73 | HMPREF0102_02503 | 46.571 | 99 | 0 | 650 | hypothetical protein [Bacteroides sp. 2_1_22] |
| Q8AB73 | BSAG_01079 | 46.571 | 99 | 0 | 650 | hypothetical protein [Bacteroides sp. D1] |
| Q8AB73 | CW3_4225 | 46.571 | 99 | 0 | 650 | conserved hypothetical protein [Bacteroides xylanisolvens SD CC 1b] |
| Q8AB73 | CW1_0172 | 46.571 | 99 | 0 | 650 | conserved hypothetical protein [Bacteroides xylanisolvens SD CC 2a] |
| Q8AB73 | HMPREF9450_00755 | 46.993 | 100 | 0 | 631 | hypothetical protein [Alistipes indistinctus YIT 12060] |
| Q8AB73 | ALIPUT_02112 | 46.439 | 100 | 0 | 618 | hypothetical protein [Alistipes putredinis DSM 17216] |
| Q8AB73 | AL1_16030 | 47.202 | 99 | 0 | 612 | Peptidase S46. [Alistipes shahii WAL 8301] |
| Q8AB73 | HMPREF9449_02133 | 44.444 | 100 | 0 | 595 | hypothetical protein [Odoribacter laneus YIT 12061] |
| Q8AB73 | HMPREF9456_01457 | 45.739 | 99 | 0 | 590 | hypothetical protein [Dysgonomonas mossii DSM 22836] |
| Q8AB73 | HMPREF9455_01975 | 44.193 | 100 | 0 | 586 | hypothetical protein [Dysgonomonas gadei ATCC BAA-286] |
| Q8AB73 | PARMER_00304 | 43.198 | 100 | 0 | 580 | hypothetical protein [Parabacteroides merdae ATCC 43184] |
| Q8AB73 | PRABACTJOHN_00535 | 42.676 | 100 | 0 | 568 | hypothetical protein [Parabacteroides johnsonii DSM 18315] |
| Q8AB73 | HMPREF0103_2247 | 43 | 99 | 0 | 568 | hypothetical protein [Bacteroides sp. 2_1_33B] |
| Q8AB73 | HMPREF0619_00047 | 42.181 | 99 | 0 | 561 | hypothetical protein [Parabacteroides sp. D13] |
| Q8AB73 | HMPREF9008_04196 | 42.181 | 99 | 0 | 559 | hypothetical protein [Parabacteroides sp. 20_3] |
| Q8AB73 | HMPREF0104_00053 | 42.037 | 99 | 0 | 557 | hypothetical protein [Bacteroides sp. 3_1_19] |
| Q8AB73 | CUU_0716 | 41.218 | 100 | 0 | 541 | conserved hypothetical protein [Bacteroides vulgatus PC510] |
| Q8AB73 | BSEG_03017 | 41.193 | 99 | 0 | 540 | hypothetical protein [Bacteroides dorei 5_1_36/D4] |
| Q8AB73 | BACDOR_02152 | 41.193 | 99 | 0 | 540 | hypothetical protein [Bacteroides dorei DSM 17855] |
| Q8AB73 | HMPREF0105_3274 | 41.193 | 99 | 0 | 540 | hypothetical protein [Bacteroides sp. 3_1_33FAA] |
| Q8AB73 | BSBG_03122 | 41.193 | 99 | 0 | 540 | hypothetical protein [Bacteroides sp. 9_1_42FAA] |
| Q8AB73 | BSFG_00163 | 41.076 | 100 | 0 | 538 | hypothetical protein [Bacteroides sp. 4_3_47FAA] |
| Q8AB73 | HMPREF9011_00248 | 41.076 | 100 | 0 | 538 | hypothetical protein [Bacteroides sp. 3_1_40A] |
| Q8AB73 | BXY_25380 | 41.453 | 99 | 1.83E-176 | 523 | dipeptidyl-peptidase 7. Serine peptidase. MEROPS family S46 [Bacteroides xylanisolvens XB1A] |
| Q8AB73 | HMPREF0127_01618 | 41.512 | 99 | 2.22E-176 | 524 | hypothetical protein [Bacteroides sp. 1_1_30] |
| Q8AB73 | HMPREF0106_04280 | 41.512 | 99 | 2.22E-176 | 524 | hypothetical protein [Bacteroides sp. D22] |
| Q8AB73 | HMPREF9441_00315 | 40.704 | 99 | 7.71E-176 | 523 | hypothetical protein [Paraprevotella clara YIT 11840] |

| | | | | | | |
|---|---|---|---|---|---|---|
| Q8AB73 | HMPREF0102_00568 | 41.369 | 99 | 8.03E-176 | 523 | hypothetical protein [Bacteroides sp. 2_1_22] |
| Q8AB73 | BSAG_00318 | 41.369 | 99 | 8.03E-176 | 523 | hypothetical protein [Bacteroides sp. D1] |
| Q8AB73 | CW3_2069 | 41.369 | 99 | 8.03E-176 | 523 | conserved hypothetical protein [Bacteroides xylanisolvens SD CC 1b] |
| Q8AB73 | CW1_0771 | 41.369 | 99 | 8.03E-176 | 523 | conserved hypothetical protein [Bacteroides xylanisolvens SD CC 2a] |
| Q8AB73 | HMPREF1532_01775 | 40.483 | 99 | 2.56E-175 | 521 | hypothetical protein [Bacteroides salyersiae WAL 10018 = DSM 18765 = JCM 12988] |
| Q8AB73 | BACCAC_01485 | 40.967 | 99 | 6.91E-175 | 520 | hypothetical protein [Bacteroides caccae ATCC 43185] |
| Q8AB73 | BSCG_03059 | 41.084 | 99 | 1.55E-174 | 519 | hypothetical protein [Bacteroides sp. 2_2_4] |
| Q8AB73 | CUY_4404 | 41.084 | 99 | 2.47E-174 | 519 | conserved hypothetical protein [Bacteroides ovatus SD CMC 3f] |
| Q8AB73 | HMPREF9010_00752 | 41.084 | 99 | 2.47E-174 | 519 | hypothetical protein [Bacteroides sp. 3_1_23] |
| Q8AB73 | HMPREF9442_02956 | 41.152 | 99 | 2.79E-174 | 519 | hypothetical protein [Paraprevotella xylaniphila YIT 11841] |
| Q8AB73 | HMPREF1017_00565 | 41.084 | 99 | 3.03E-174 | 518 | hypothetical protein [Bacteroides ovatus 3_8_47FAA] |
| Q8AB73 | HMPREF9447_02263 | 40.96 | 99 | 8.68E-174 | 517 | hypothetical protein [Bacteroides oleiciplenus YIT 12058] |
| Q8AB73 | BSGG_2332 | 40.942 | 99 | 9.34E-174 | 517 | hypothetical protein [Bacteroides sp. D2] |
| Q8AB73 | BACOVA_00899 | 40.942 | 99 | 9.34E-174 | 517 | hypothetical protein [Bacteroides ovatus ATCC 8483] |
| Q8AB73 | BACINT_01296 | 41.044 | 99 | 1.54E-173 | 516 | hypothetical protein [Bacteroides intestinalis DSM 17393] |
| Q8AB73 | BACFIN_05173 | 40.625 | 100 | 4.77E-172 | 513 | hypothetical protein [Bacteroides finegoldii DSM 17565] |
| Q8AB73 | BACCELL_01726 | 40.788 | 99 | 7.94E-172 | 512 | hypothetical protein [Bacteroides cellulosilyticus DSM 14838] |
| Q8AB73 | HMPREF1214_00608 | 39.773 | 99 | 8.49E-172 | 512 | hypothetical protein [Bacteroides sp. HPS0048] |
| Q8AB73 | HMPREF9445_01804 | 41.408 | 99 | 1.87E-171 | 512 | hypothetical protein [Bacteroides clarus YIT 12056] |
| Q8AB73 | HMPREF9450_02265 | 40.288 | 99 | 2.08E-170 | 508 | hypothetical protein [Alistipes indistinctus YIT 12060] |
| Q8AB73 | BSIG_1382 | 40.426 | 99 | 1.28E-169 | 506 | hypothetical protein [Bacteroides sp. 1_1_6] |
| Q8AB73 | HMPREF9007_03829 | 40.426 | 99 | 1.41E-169 | 506 | hypothetical protein [Bacteroides sp. 1_1_14] |
| Q8AB73 | HMPREF1016_00890 | 40.17 | 99 | 1.17E-168 | 504 | peptidase S46 [Bacteroides eggerthii 1_2_48FAA] |
| Q8AB73 | BACEGG_03368 | 40.17 | 99 | 1.17E-168 | 504 | hypothetical protein [Bacteroides eggerthii DSM 20697] |
| Q8AB73 | BACSTE_02429 | 40.845 | 99 | 1.52E-168 | 504 | hypothetical protein [Bacteroides stercoris ATCC 43183] |
| Q8AB73 | BFAG_02873 | 39.459 | 99 | 4.89E-167 | 500 | hypothetical protein [Bacteroides fragilis 3_1_12] |
| Q8AB73 | HMPREF0101_02131 | 39.316 | 99 | 1.64E-166 | 499 | hypothetical protein [Bacteroides sp. 2_1_16] |
| Q8AB73 | HMPREF1018_00169 | 39.316 | 99 | 7.45E-166 | 497 | hypothetical protein [Bacteroides sp. 2_1_56FAA] |
| Q8AB73 | HMPREF9446_00563 | 40.426 | 99 | 8.74E-166 | 497 | hypothetical protein [Bacteroides fluxus YIT 12057] |
| Q8AB73 | BSHG_1458 | 39.316 | 99 | 9.15E-166 | 497 | hypothetical protein [Bacteroides sp. 3_2_5] |
| Q8AB73 | HMPREF0969_01517 | 40.114 | 99 | 1.77E-165 | 496 | hypothetical protein [Bacteroides sp. D20] |
| Q8AB73 | HMPREF1007_00282 | 40.114 | 99 | 1.99E-165 | 496 | peptidase S46 [Bacteroides sp. 4_1_36] |
| Q8AB73 | BACUNI_01062 | 39.972 | 99 | 6.74E-165 | 494 | hypothetical protein [Bacteroides uniformis ATCC 8492] |
| Q8AB73 | HMPREF9449_02174 | 38.764 | 100 | 7.61E-163 | 489 | hypothetical protein [Odoribacter laneus YIT 12061] |
| Q8AB73 | HMPREF9449_00904 | 39.548 | 99 | 3.27E-161 | 485 | hypothetical protein [Odoribacter laneus YIT 12061] |
| Q8AB73 | ALIPUT_00511 | 39.056 | 100 | 6.26E-157 | 472 | hypothetical protein [Alistipes putredinis DSM 17216] |
| Q8AB73 | PREVCOP_06589 | 37.571 | 98 | 7.53E-155 | 469 | hypothetical protein [Prevotella copri DSM 18205] |
| Q8AB73 | AL1_16890 | 38.494 | 99 | 4.31E-152 | 460 | Peptidase S46. [Alistipes shahii WAL 8301] |
| Q8AB73 | HMPREF9420_1755 | 37.907 | 99 | 1.29E-147 | 449 | hypothetical protein [Prevotella salivae DSM 15606] |
| Q8AB73 | HMPREF1475_01683 | 38.102 | 99 | 1.47E-144 | 442 | hypothetical protein [Prevotella oralis HGA0225] |
| Q8AB73 | HMPREF0673_00993 | 36.975 | 99 | 3.33E-144 | 441 | hypothetical protein [Prevotella stercorea DSM 18206] |
| Q8AB73 | AL1_16630 | 36.25 | 100 | 4.82E-133 | 412 | Peptidase S46. [Alistipes shahii WAL 8301] |
| Q8AB73 | BSGG_3154 | 29.42 | 99 | 1.76E-90 | 301 | hypothetical protein [Bacteroides sp. D2] |
| Q8AB73 | BACOVA_04406 | 29.144 | 99 | 1.25E-88 | 296 | hypothetical protein [Bacteroides ovatus ATCC 8483] |
| Q8AB09 | BSIG_0127 | 100 | 100 | 0 | 1529 | hypothetical protein [Bacteroides sp. 1_1_6] |
| Q8AB09 | BACCAC_02271 | 90.257 | 100 | 0 | 1411 | phospholipase, patatin family [Bacteroides caccae ATCC 43185] |
| Q8AB09 | HMPREF0106_03968 | 87.398 | 100 | 0 | 1379 | phospholipase, patatin family [Bacteroides sp. D22] |
| Q8AB09 | HMPREF9010_01762 | 87.534 | 100 | 0 | 1377 | phospholipase, patatin family [Bacteroides sp. 3_1_23] |
| Q8AB09 | BSCG_00789 | 87.534 | 100 | 0 | 1377 | phospholipase, patatin family [Bacteroides sp. 2_2_4] |
| Q8AB09 | BACOVA_00420 | 87.534 | 100 | 0 | 1377 | phospholipase, patatin family [Bacteroides ovatus ATCC 8483] |
| Q8AB09 | CUY_4079 | 87.398 | 100 | 0 | 1377 | phospholipase, patatin family [Bacteroides ovatus SD CMC 3f] |
| Q8AB09 | HMPREF1017_05053 | 87.398 | 100 | 0 | 1376 | hypothetical protein [Bacteroides ovatus 3_8_47FAA] |
| Q8AB09 | BACFIN_08682 | 87.551 | 100 | 0 | 1374 | phospholipase, patatin family [Bacteroides finegoldii DSM 17565] |
| Q8AB09 | BXY_15130 | 87.127 | 100 | 0 | 1374 | Predicted esterase of the alpha-beta hydrolase superfamily [Bacteroides xylanisolvens XB1A] |
| Q8AB09 | HMPREF0127_03476 | 87.127 | 100 | 0 | 1373 | hypothetical protein [Bacteroides sp. 1_1_30] |
| Q8AB09 | HMPREF0102_02535 | 87.127 | 100 | 0 | 1373 | phospholipase, patatin family [Bacteroides sp. 2_1_22] |
| Q8AB09 | BSAG_01109 | 87.127 | 100 | 0 | 1373 | phospholipase, patatin family [Bacteroides sp. D1] |
| Q8AB09 | BSGG_0662 | 87.263 | 100 | 0 | 1373 | hypothetical protein [Bacteroides sp. D2] |
| Q8AB09 | CW3_2075 | 87.535 | 98 | 0 | 1352 | phospholipase, patatin family [Bacteroides xylanisolvens SD CC 1b] |
| Q8AB09 | HMPREF9007_01235 | 99.08 | 88 | 0 | 1344 | phospholipase, patatin family [Bacteroides sp. 1_1_14] |

| | | | | | | |
|---|---|---|---|---|---|---|
| Q8AB09 | HMPREF0101_03192 | 73.713 | 100 | 0 | 1162 | phospholipase, patatin family [Bacteroides sp. 2_1_16] |
| Q8AB09 | BFAG_03154 | 75.35 | 97 | 0 | 1161 | putative phage tail component domain protein [Bacteroides fragilis 3_1_12] |
| Q8AB09 | HMPREF1018_04237 | 73.577 | 100 | 0 | 1161 | hypothetical protein [Bacteroides sp. 2_1_56FAA] |
| Q8AB09 | BSHG_1871 | 73.442 | 100 | 0 | 1159 | hypothetical protein [Bacteroides sp. 3_2_5] |
| Q8AB09 | HMPREF1214_03820 | 68.606 | 100 | 0 | 1087 | NTE family protein [Bacteroides sp. HPS0048] |
| Q8AB09 | HMPREF1532_00269 | 67.659 | 100 | 0 | 1068 | hypothetical protein [Bacteroides salyersiae WAL 10018 = DSM 18765 = JCM 12988] |
| Q8AB09 | HMPREF1018_03072 | 63.158 | 100 | 0 | 1004 | hypothetical protein [Bacteroides sp. 2_1_56FAA] |
| Q8AB09 | HMPREF0101_03770 | 63.158 | 100 | 0 | 1004 | phospholipase, patatin family [Bacteroides sp. 2_1_16] |
| Q8AB09 | BSHG_3165 | 63.158 | 100 | 0 | 1003 | hypothetical protein [Bacteroides sp. 3_2_5] |
| Q8AB09 | BFAG_03874 | 65.126 | 97 | 0 | 999 | phospholipase, patatin family [Bacteroides fragilis 3_1_12] |
| Q8AB09 | HMPREF9011_01083 | 62.803 | 100 | 0 | 967 | patatin-like phospholipase [Bacteroides sp. 3_1_40A] |
| Q8AB09 | CUU_1427 | 62.822 | 99 | 0 | 962 | phospholipase, patatin family [Bacteroides vulgatus PC510] |
| Q8AB09 | BSFG_03784 | 62.822 | 99 | 0 | 962 | phospholipase, patatin family [Bacteroides sp. 4_3_47FAA] |
| Q8AB09 | BACDOR_02941 | 62.264 | 100 | 0 | 956 | phospholipase, patatin family [Bacteroides dorei DSM 17855] |
| Q8AB09 | HMPREF0105_4378 | 62.129 | 100 | 0 | 955 | phospholipase, patatin family [Bacteroides sp. 3_1_33FAA] |
| Q8AB09 | BSEG_00063 | 62.144 | 99 | 0 | 951 | phospholipase, patatin family [Bacteroides dorei 5_1_36/D4] |
| Q8AB09 | BSBG_01896 | 62.144 | 99 | 0 | 951 | phospholipase, patatin family [Bacteroides sp. 9_1_42FAA] |
| Q8AB09 | CW1_1829 | 86.907 | 60 | 0 | 830 | conserved hypothetical protein [Bacteroides xylanisolvens SD CC 2a] |
| Q8AB09 | HMPREF9449_00398 | 51.24 | 98 | 0 | 769 | hypothetical protein [Odoribacter laneus YIT 12061] |
| Q8AB09 | PARMER_00099 | 49.446 | 97 | 0 | 724 | phospholipase, patatin family [Parabacteroides merdae ATCC 43184] |
| Q8AB09 | PRABACTJOHN_00370 | 49.307 | 97 | 0 | 724 | phospholipase, patatin family [Parabacteroides johnsonii DSM 18315] |
| Q8AB09 | HMPREF9008_04488 | 50.143 | 94 | 0 | 713 | surface antigen variable number repeat protein [Parabacteroides sp. 20_3] |
| Q8AB09 | HMPREF0103_1929 | 50.143 | 94 | 0 | 712 | surface antigen variable number repeat protein [Bacteroides sp. 2_1_33B] |
| Q8AB09 | HMPREF0104_00348 | 49.427 | 94 | 0 | 703 | hypothetical protein [Bacteroides sp. 3_1_19] |
| Q8AB09 | HMPREF0619_00262 | 49.427 | 94 | 0 | 702 | surface antigen variable number repeat protein [Parabacteroides sp. D13] |
| Q8AB09 | HMPREF9449_01091 | 47.632 | 97 | 0 | 682 | hypothetical protein [Odoribacter laneus YIT 12061] |
| Q8AB09 | HMPREF1475_00755 | 42.876 | 99 | 0 | 593 | hypothetical protein [Prevotella oralis HGA0225] |
| Q8AB09 | HMPREF1007_00232 | 41.473 | 99 | 0 | 559 | patatin-like phospholipase [Bacteroides sp. 4_1_36] |
| Q8AB09 | HMPREF0969_01456 | 41.633 | 99 | 0 | 557 | phospholipase, patatin family [Bacteroides sp. D20] |
| Q8AB09 | BACUNI_01001 | 41.848 | 99 | 0 | 557 | phospholipase, patatin family [Bacteroides uniformis ATCC 8492] |
| Q8AB09 | HMPREF9445_00032 | 42.996 | 97 | 0 | 554 | phospholipase, patatin family [Bacteroides clarus YIT 12056] |
| Q8AB09 | BACINT_00484 | 42.259 | 97 | 0 | 554 | phospholipase, patatin family [Bacteroides intestinalis DSM 17393] |
| Q8AB09 | HMPREF9446_03118 | 42.003 | 97 | 0 | 547 | phospholipase, patatin family [Bacteroides fluxus YIT 12057] |
| Q8AB09 | CW1_1828 | 87.747 | 34 | 7.16E-160 | 467 | phospholipase, patatin family [Bacteroides xylanisolvens SD CC 2a] |
| Q8AB09 | PREVCOP_06304 | 35.839 | 99 | 1.19E-153 | 469 | phospholipase, patatin family [Prevotella copri DSM 18205] |
| Q8AB09 | HMPREF9447_01746 | 61.538 | 88 | 2.12E-132 | 411 | hypothetical protein [Bacteroides oleiciplenus YIT 12058] |
| Q8AB09 | HMPREF9447_01746 | 37.117 | 88 | 8.85E-62 | 223 | hypothetical protein [Bacteroides oleiciplenus YIT 12058] |
| Q8AB09 | BACCELL_01748 | 57.975 | 89 | 1.32E-129 | 404 | phospholipase, patatin family [Bacteroides cellulosilyticus DSM 14838] |
| Q8AB09 | BACCELL_01748 | 36.937 | 89 | 2.74E-65 | 233 | phospholipase, patatin family [Bacteroides cellulosilyticus DSM 14838] |
| Q8AB09 | HMPREF9441_01694 | 55.882 | 87 | 2.31E-112 | 358 | phospholipase, patatin family [Paraprevotella clara YIT 11840] |
| Q8AB09 | HMPREF9441_01694 | 32.558 | 87 | 3.92E-51 | 193 | phospholipase, patatin family [Paraprevotella clara YIT 11840] |
| Q8AB09 | HMPREF9442_02648 | 55.882 | 87 | 3.01E-112 | 358 | phospholipase, patatin family [Paraprevotella xylaniphila YIT 11841] |
| Q8AB09 | HMPREF9442_02648 | 32.849 | 87 | 1.75E-52 | 197 | phospholipase, patatin family [Paraprevotella xylaniphila YIT 11841] |
| Q8AB09 | BFAG_02511 | 50.599 | 89 | 3.00E-103 | 334 | phospholipase, patatin family [Bacteroides fragilis 3_1_12] |
| Q8AB09 | BFAG_02511 | 29.878 | 89 | 1.78E-38 | 156 | phospholipase, patatin family [Bacteroides fragilis 3_1_12] |
| Q8AB09 | HMPREF0102_01311 | 50.489 | 86 | 3.60E-99 | 323 | phospholipase, patatin family [Bacteroides sp. 2_1_22] |
| Q8AB09 | HMPREF0102_01311 | 32.515 | 86 | 2.93E-49 | 188 | phospholipase, patatin family [Bacteroides sp. 2_1_22] |
| Q8AB09 | BSAG_03782 | 50.489 | 86 | 3.60E-99 | 323 | phospholipase, patatin family [Bacteroides sp. D1] |
| Q8AB09 | BSAG_03782 | 32.515 | 86 | 2.93E-49 | 188 | phospholipase, patatin family [Bacteroides sp. D1] |
| Q8AB09 | CW3_2182 | 50.489 | 86 | 3.60E-99 | 323 | phospholipase, patatin family [Bacteroides xylanisolvens SD CC 1b] |
| Q8AB09 | CW3_2182 | 32.515 | 86 | 2.93E-49 | 188 | phospholipase, patatin family [Bacteroides xylanisolvens SD CC 1b] |
| Q8AB09 | CW1_0310 | 50.489 | 86 | 3.60E-99 | 323 | phospholipase, patatin family [Bacteroides xylanisolvens SD CC 2a] |
| Q8AB09 | CW1_0310 | 32.515 | 86 | 2.93E-49 | 188 | phospholipase, patatin family [Bacteroides xylanisolvens SD CC 2a] |
| Q8AB09 | HMPREF9007_04140 | 50.489 | 86 | 3.71E-99 | 323 | phospholipase, patatin family [Bacteroides sp. 1_1_14] |
| Q8AB09 | HMPREF9007_04140 | 32.209 | 86 | 8.11E-48 | 184 | phospholipase, patatin family [Bacteroides sp. 1_1_14] |
| Q8AB09 | HMPREF0105_1378 | 50.492 | 86 | 1.69E-98 | 322 | phospholipase, patatin family [Bacteroides sp. 3_1_33FAA] |
| Q8AB09 | HMPREF0105_1378 | 31.288 | 86 | 1.03E-45 | 177 | phospholipase, patatin family [Bacteroides sp. 3_1_33FAA] |
| Q8AB09 | HMPREF9447_00151 | 50.331 | 85 | 2.27E-97 | 318 | hypothetical protein [Bacteroides oleiciplenus YIT 12058] |
| Q8AB09 | HMPREF9447_00151 | 32.515 | 85 | 3.04E-49 | 188 | hypothetical protein [Bacteroides oleiciplenus YIT 12058] |
| Q8AB09 | HMPREF9466_00771 | 37.543 | 40 | 1.36E-53 | 193 | hypothetical protein [Fusobacterium necrophorum subsp. funduliforme 1_1_36S] |

| | | | | | | |
|---|---|---|---|---|---|---|
| Q8AB09 | HMPREF1030_00884 | 28.571 | 75 | 3.98E-53 | 200 | hypothetical protein [Pseudomonas sp. 2_1_26] |
| Q8AB09 | FMAG_00818 | 26.064 | 96 | 1.76E-51 | 196 | phospholipase, patatin family [Fusobacterium mortiferum ATCC 9817] |
| Q8AB09 | FSBG_01353 | 25.035 | 97 | 6.10E-51 | 194 | surface antigen variable number repeat protein [Fusobacterium gonidiaformans 3-1-5R] |
| Q8AB09 | HMPREF9440_02002 | 31.25 | 54 | 4.28E-50 | 191 | phospholipase, patatin family [Sutterella parvirubra YIT 11816] |
| Q8AB09 | FVAG_01248 | 27.07 | 97 | 7.81E-50 | 191 | surface antigen variable number repeat protein [Fusobacterium varium ATCC 27725] |
| Q8AB09 | FGAG_01009 | 24.62 | 97 | 9.45E-49 | 188 | surface antigen variable number repeat protein [Fusobacterium gonidiaformans ATCC 25563] |
| Q8AB09 | HMPREF0402_02935 | 35.461 | 38 | 9.71E-49 | 188 | hypothetical protein [Fusobacterium ulcerans 12-1B] |
| Q8AB09 | FUAG_02317 | 36.014 | 38 | 1.92E-48 | 187 | hypothetical protein [Fusobacterium ulcerans ATCC 49185] |
| Q8AB09 | FMAG_00016 | 26.316 | 79 | 3.01E-48 | 186 | surface antigen variable number repeat protein [Fusobacterium mortiferum ATCC 9817] |
| Q8AB09 | HMPREF9465_00024 | 27.373 | 80 | 4.58E-48 | 186 | hypothetical protein [Sutterella wadsworthensis 2_1_59BFAA] |
| Q8AB09 | HMPREF1705_01578 | 28.13 | 77 | 4.81E-47 | 182 | phospholipase, patatin family [Anaerobaculum hydrogeniformans ATCC BAA-1850] |
| Q8AB09 | HMPREF1004_02453 | 32.402 | 48 | 5.57E-47 | 183 | surface antigen variable number repeat protein [Ralstonia sp. 5_7_47FAA] |
| Q8AB09 | HMPREF9464_00407 | 30.923 | 53 | 5.84E-47 | 182 | hypothetical protein [Sutterella wadsworthensis 3_1_45B] |
| Q8AB09 | HMPREF0989_02691 | 32.402 | 48 | 6.20E-47 | 183 | hypothetical protein [Ralstonia sp. 5_2_56FAA] |
| Q8AB09 | HMPREF1476_02169 | 30.673 | 53 | 1.92E-46 | 181 | hypothetical protein [Sutterella wadsworthensis HGA0223] |
| Q8AB09 | BACCOP_02003 | 28.106 | 63 | 7.57E-43 | 171 | phospholipase, patatin family [Bacteroides coprocola DSM 17136] |
| Q8AB09 | HMPREF1033_00307 | 29.314 | 56 | 1.91E-42 | 169 | hypothetical protein [Tannerella sp. 6_1_58FAA_CT1] |
| Q8AB09 | FUAG_00538 | 29.363 | 48 | 2.57E-42 | 169 | hypothetical protein [Fusobacterium ulcerans ATCC 49185] |
| Q8AB09 | HMPREF0402_00731 | 29.363 | 48 | 7.56E-42 | 167 | hypothetical protein [Fusobacterium ulcerans 12-1B] |
| Q8AB09 | FVAG_00457 | 23.777 | 96 | 7.77E-42 | 167 | phospholipase, patatin family [Fusobacterium varium ATCC 27725] |
| Q8AB09 | HMPREF1214_02219 | 27.14 | 62 | 3.99E-41 | 165 | NTE family protein [Bacteroides sp. HPS0048] |
| Q8AB09 | BACPLE_00812 | 27.893 | 62 | 4.08E-41 | 165 | phospholipase, patatin family [Bacteroides plebeius DSM 17135] |
| Q8AB09 | BFAG_01977 | 26.778 | 62 | 1.15E-40 | 164 | phospholipase, patatin family [Bacteroides fragilis 3_1_12] |
| Q8AB09 | HMPREF9011_01286 | 27.391 | 59 | 2.77E-40 | 163 | patatin-like phospholipase [Bacteroides sp. 3_1_40A] |
| Q8AB09 | CUU_3874 | 27.391 | 59 | 2.98E-40 | 163 | phospholipase, patatin family [Bacteroides vulgatus PC510] |
| Q8AB09 | BSFG_01260 | 27.391 | 59 | 3.87E-40 | 162 | phospholipase, patatin family [Bacteroides sp. 4_3_47FAA] |
| Q8AB09 | BSEG_02311 | 27.609 | 59 | 5.12E-40 | 162 | phospholipase, patatin family [Bacteroides dorei 5_1_36/D4] |
| Q8AB09 | HMPREF0105_2599 | 27.609 | 59 | 5.22E-40 | 162 | phospholipase, patatin family [Bacteroides sp. 3_1_33FAA] |
| Q8AB09 | BSBG_02713 | 27.609 | 59 | 5.22E-40 | 162 | phospholipase, patatin family [Bacteroides sp. 9_1_42FAA] |
| Q8AB09 | FSBG_00285 | 31.757 | 40 | 5.36E-40 | 162 | phospholipase, patatin family [Fusobacterium gonidiaformans 3-1-5R] |
| Q8AB09 | HMPREF1006_01762 | 32.055 | 47 | 5.62E-40 | 162 | hypothetical protein [Synergistes sp. 3_1_syn1] |
| Q8AB09 | BACDOR_02695 | 27.609 | 59 | 6.56E-40 | 162 | phospholipase, patatin family [Bacteroides dorei DSM 17855] |
| Q8AB09 | BSIG_0700 | 27.368 | 62 | 7.99E-40 | 161 | hypothetical protein [Bacteroides sp. 1_1_6] |
| Q8AB09 | HMPREF9450_00746 | 33.227 | 41 | 1.32E-39 | 160 | hypothetical protein [Alistipes indistinctus YIT 12060] |
| Q8AB09 | BSHG_2622 | 26.84 | 59 | 6.23E-39 | 159 | hypothetical protein [Bacteroides sp. 3_2_5] |
| Q8AB09 | HMPREF1018_02557 | 26.84 | 59 | 6.23E-39 | 159 | hypothetical protein [Bacteroides sp. 2_1_56FAA] |
| Q8AB09 | HMPREF0101_01591 | 26.84 | 59 | 6.81E-39 | 159 | phospholipase, patatin family [Bacteroides sp. 2_1_16] |
| Q8AB09 | FGAG_01187 | 31.419 | 40 | 7.93E-39 | 158 | phospholipase, patatin family [Fusobacterium gonidiaformans ATCC 25563] |
| Q8AB09 | HMPREF9010_01070 | 27.31 | 62 | 8.06E-39 | 158 | phospholipase, patatin family [Bacteroides sp. 3_1_23] |
| Q8AB09 | HMPREF9447_00200 | 37.86 | 32 | 8.97E-39 | 158 | hypothetical protein [Bacteroides oleiciplenus YIT 12058] |
| Q8AB09 | HMPREF1017_03552 | 27.31 | 62 | 1.07E-38 | 158 | hypothetical protein [Bacteroides ovatus 3_8_47FAA] |
| Q8AB09 | BACFIN_09123 | 26.694 | 63 | 1.10E-38 | 158 | phospholipase, patatin family [Bacteroides finegoldii DSM 17565] |
| Q8AB09 | BSCG_02216 | 27.31 | 62 | 1.18E-38 | 158 | phospholipase, patatin family [Bacteroides sp. 2_2_4] |
| Q8AB09 | BACOVA_00719 | 27.31 | 62 | 1.18E-38 | 158 | phospholipase, patatin family [Bacteroides ovatus ATCC 8483] |
| Q8AB09 | CUY_5128 | 27.31 | 62 | 1.27E-38 | 158 | phospholipase, patatin family [Bacteroides ovatus SD CMC 3f] |
| Q8AB09 | BSGG_0001 | 27.31 | 62 | 1.27E-38 | 158 | hypothetical protein [Bacteroides sp. D2] |
| Q8AB09 | HMPREF0102_03076 | 26.749 | 62 | 1.48E-38 | 157 | phospholipase, patatin family [Bacteroides sp. 2_1_22] |
| Q8AB09 | BSAG_01665 | 26.749 | 62 | 1.48E-38 | 157 | phospholipase, patatin family [Bacteroides sp. D1] |
| Q8AB09 | BACCELL_04898 | 27.887 | 59 | 1.49E-38 | 157 | phospholipase, patatin family [Bacteroides cellulosilyticus DSM 14838] |
| Q8AB09 | CW3_4571 | 26.749 | 62 | 1.58E-38 | 157 | phospholipase, patatin family [Bacteroides xylanisolvens SD CC 1b] |
| Q8AB09 | CW1_4352 | 26.749 | 62 | 1.58E-38 | 157 | phospholipase, patatin family [Bacteroides xylanisolvens SD CC 2a] |
| Q8AB09 | BACCAC_00558 | 27.158 | 62 | 2.63E-38 | 157 | phospholipase, patatin family [Bacteroides caccae ATCC 43185] |
| Q8AB09 | HMPREF9446_02666 | 37.398 | 32 | 3.51E-38 | 157 | phospholipase, patatin family [Bacteroides fluxus YIT 12057] |
| Q8AB09 | BACINT_01992 | 27.887 | 59 | 4.13E-38 | 156 | phospholipase, patatin family [Bacteroides intestinalis DSM 17393] |
| Q8AB09 | HMPREF1532_03279 | 28.78 | 53 | 5.50E-38 | 156 | hypothetical protein [Bacteroides salyersiae WAL 10018 = DSM 18765 = JCM 12988] |
| Q8AB09 | BXY_04710 | 26.543 | 62 | 9.79E-38 | 155 | Predicted esterase of the alpha-beta hydrolase superfamily [Bacteroides xylanisolvens XB1A] |
| Q8AB09 | HMPREF0127_00278 | 26.543 | 62 | 1.06E-37 | 155 | hypothetical protein [Bacteroides sp. 1_1_30] |
| Q8AB09 | HMPREF0106_04656 | 26.543 | 62 | 1.06E-37 | 155 | phospholipase, patatin family [Bacteroides sp. D22] |
| Q8AB09 | BACCOPRO_02612 | 35.223 | 32 | 1.07E-37 | 155 | phospholipase, patatin family [Bacteroides coprophilus DSM 18228 = JCM 13818] |
| Q8AB09 | HMPREF0969_03217 | 29.056 | 53 | 1.26E-37 | 155 | phospholipase, patatin family [Bacteroides sp. D20] |

| | | | | | | |
|---|---|---|---|---|---|---|
| Q8AB09 | HMPREF1007_02295 | 29.056 | 53 | 1.26E-37 | 155 | patatin-like phospholipase [Bacteroides sp. 4_1_36] |
| Q8AB09 | BACUNI_02000 | 29.056 | 53 | 1.46E-37 | 154 | phospholipase, patatin family [Bacteroides uniformis ATCC 8492] |
| Q8AB09 | HMPREF9445_00702 | 33.878 | 32 | 1.41E-36 | 151 | phospholipase, patatin family [Bacteroides clarus YIT 12056] |
| Q8AB09 | BACSTE_01473 | 27.451 | 59 | 2.21E-36 | 151 | phospholipase, patatin family [Bacteroides stercoris ATCC 43183] |
| Q8AB09 | HMPREF1016_03234 | 34.286 | 32 | 3.20E-36 | 150 | patatin-like phospholipase [Bacteroides eggerthii 1_2_48FAA] |
| Q8AB09 | BACEGG_02714 | 34.286 | 32 | 3.26E-36 | 150 | phospholipase, patatin family [Bacteroides eggerthii DSM 20697] |
| Q8AB09 | PRABACTJOHN_02312 | 30.295 | 48 | 3.87E-36 | 148 | phospholipase, patatin family [Parabacteroides johnsonii DSM 18315] |
| Q8AB09 | FSEG_01208 | 29.352 | 40 | 5.09E-36 | 150 | phospholipase, patatin family [Fusobacterium necrophorum D12] |
| Q8AB09 | PARMER_01815 | 30.203 | 52 | 9.78E-36 | 149 | phospholipase, patatin family [Parabacteroides merdae ATCC 43184] |
| Q8AB09 | HMPREF9448_00383 | 34.201 | 35 | 2.38E-35 | 148 | hypothetical protein [Barnesiella intestinihominis YIT 11860] |
| Q8AB09 | HMPREF1014_00937 | 33.684 | 39 | 3.04E-35 | 139 | hypothetical protein [Bacillus sp. 7_6_55CFAA_CT2] |
| Q8AB09 | ALIPUT_02121 | 36.327 | 31 | 6.08E-35 | 146 | phospholipase, patatin family [Alistipes putredinis DSM 17216] |
| Q8AB09 | HMPREF9008_01022 | 29.041 | 48 | 9.79E-35 | 146 | phospholipase, patatin family [Parabacteroides sp. 20_3] |
| Q8AB09 | HMPREF0619_01433 | 29.041 | 48 | 1.20E-34 | 145 | phospholipase, patatin family [Parabacteroides sp. D13] |
| Q8AB09 | HMPREF0103_1148 | 29.041 | 48 | 1.90E-34 | 145 | phospholipase, patatin family [Bacteroides sp. 2_1_33B] |
| Q8AB09 | HMPREF0104_01483 | 29.041 | 48 | 1.90E-34 | 145 | phospholipase, patatin family [Bacteroides sp. 3_1_19] |
| Q8AB09 | HMPREF1015_00726 | 34.495 | 39 | 2.04E-34 | 136 | hypothetical protein [Bacillus smithii 7_3_47FAA] |
| Q8AB09 | FSAG_001224 | 30.592 | 40 | 4.95E-34 | 144 | NTE family protein [Fusobacterium periodonticum 2_1_31] |
| Q8AB09 | FSCG_01360 | 30.07 | 39 | 2.64E-33 | 141 | phospholipase, patatin family [Fusobacterium nucleatum subsp. vincentii 4_1_13] |
| Q8AB09 | HMPREF0404_01566 | 28.009 | 57 | 9.37E-33 | 140 | hypothetical protein [Fusobacterium nucleatum subsp. animalis 21_1A] |
| Q8AB09 | HMPREF0400_01371 | 30.389 | 38 | 1.30E-32 | 139 | phospholipase, patatin family [Fusobacterium periodonticum 1_1_41FAA] |
| Q8AB09 | FSDG_01861 | 28.009 | 57 | 1.73E-32 | 139 | hypothetical protein [Fusobacterium nucleatum subsp. animalis 7_1] |
| Q8AB09 | HMPREF0406_01169 | 30.769 | 39 | 2.15E-32 | 139 | hypothetical protein [Fusobacterium nucleatum subsp. animalis 3_1_33] |
| Q8AB09 | PSAG_02178 | 28.009 | 57 | 2.66E-32 | 138 | phospholipase, patatin family [Fusobacterium nucleatum subsp. animalis D11] |
| Q8AB09 | AL1_31780 | 35.426 | 30 | 2.77E-32 | 130 | Predicted esterase of the alpha-beta hydrolase superfamily [Alistipes shahii WAL 8301] |
| Q8AB09 | HMPREF0401_02248 | 27.778 | 57 | 4.30E-32 | 137 | serine protease [Fusobacterium nucleatum subsp. animalis 11_3_2] |
| Q8AB09 | AL1_15940 | 38.356 | 28 | 9.76E-32 | 136 | Predicted esterase of the alpha-beta hydrolase superfamily [Alistipes shahii WAL 8301] |
| Q8AB09 | ANASTE_00165 | 34.4 | 34 | 7.87E-31 | 126 | phospholipase, patatin family [Anaerofustis stercorihominis DSM 17244] |
| Q8A9B9 | BSIG_0700 | 100 | 100 | 0 | 1560 | hypothetical protein [Bacteroides sp. 1_1_6] |
| Q8A9B9 | BACCAC_00558 | 92.819 | 100 | 0 | 1464 | phospholipase, patatin family [Bacteroides caccae ATCC 43185] |
| Q8A9B9 | HMPREF0102_03076 | 91.755 | 100 | 0 | 1457 | phospholipase, patatin family [Bacteroides sp. 2_1_22] |
| Q8A9B9 | BSAG_01665 | 91.755 | 100 | 0 | 1457 | phospholipase, patatin family [Bacteroides sp. D1] |
| Q8A9B9 | CW3_4571 | 91.755 | 100 | 0 | 1456 | phospholipase, patatin family [Bacteroides xylanisolvens SD CC 1b] |
| Q8A9B9 | CW1_4352 | 91.755 | 100 | 0 | 1456 | phospholipase, patatin family [Bacteroides xylanisolvens SD CC 2a] |
| Q8A9B9 | CUY_5128 | 91.888 | 100 | 0 | 1455 | phospholipase, patatin family [Bacteroides ovatus SD CMC 3f] |
| Q8A9B9 | BSGG_0001 | 91.888 | 100 | 0 | 1455 | hypothetical protein [Bacteroides sp. D2] |
| Q8A9B9 | BXY_04710 | 91.755 | 100 | 0 | 1455 | Predicted esterase of the alpha-beta hydrolase superfamily [Bacteroides xylanisolvens XB1A] |
| Q8A9B9 | BSCG_02216 | 91.888 | 100 | 0 | 1454 | phospholipase, patatin family [Bacteroides sp. 2_2_4] |
| Q8A9B9 | HMPREF1017_03552 | 91.755 | 100 | 0 | 1454 | hypothetical protein [Bacteroides ovatus 3_8_47FAA] |
| Q8A9B9 | BACOVA_00719 | 91.888 | 100 | 0 | 1454 | phospholipase, patatin family [Bacteroides ovatus ATCC 8483] |
| Q8A9B9 | HMPREF0127_00278 | 91.489 | 100 | 0 | 1453 | hypothetical protein [Bacteroides sp. 1_1_30] |
| Q8A9B9 | HMPREF0106_04656 | 91.489 | 100 | 0 | 1453 | phospholipase, patatin family [Bacteroides sp. D22] |
| Q8A9B9 | HMPREF9010_01070 | 91.622 | 100 | 0 | 1452 | phospholipase, patatin family [Bacteroides sp. 3_1_23] |
| Q8A9B9 | BACFIN_09123 | 90.957 | 100 | 0 | 1448 | phospholipase, patatin family [Bacteroides finegoldii DSM 17565] |
| Q8A9B9 | HMPREF1532_03279 | 83.311 | 99 | 0 | 1345 | hypothetical protein [Bacteroides salyersiae WAL 10018 = DSM 18765 = JCM 12988] |
| Q8A9B9 | HMPREF0101_01591 | 83.979 | 99 | 0 | 1342 | phospholipase, patatin family [Bacteroides sp. 2_1_16] |
| Q8A9B9 | BSHG_2622 | 83.979 | 99 | 0 | 1342 | hypothetical protein [Bacteroides sp. 3_2_5] |
| Q8A9B9 | HMPREF1018_02557 | 83.979 | 99 | 0 | 1342 | hypothetical protein [Bacteroides sp. 2_1_56FAA] |
| Q8A9B9 | HMPREF1214_02219 | 82.911 | 99 | 0 | 1341 | NTE family protein [Bacteroides sp. HPS0048] |
| Q8A9B9 | BFAG_01977 | 83.178 | 99 | 0 | 1335 | phospholipase, patatin family [Bacteroides fragilis 3_1_12] |
| Q8A9B9 | BACINT_01992 | 78.046 | 99 | 0 | 1239 | phospholipase, patatin family [Bacteroides intestinalis DSM 17393] |
| Q8A9B9 | HMPREF9447_00200 | 77.63 | 99 | 0 | 1239 | hypothetical protein [Bacteroides oleiciplenus YIT 12058] |
| Q8A9B9 | BACCELL_04898 | 77.644 | 99 | 0 | 1233 | phospholipase, patatin family [Bacteroides cellulosilyticus DSM 14838] |
| Q8A9B9 | HMPREF9446_02666 | 75.532 | 99 | 0 | 1219 | phospholipase, patatin family [Bacteroides fluxus YIT 12057] |
| Q8A9B9 | BACSTE_01473 | 75.233 | 99 | 0 | 1218 | phospholipase, patatin family [Bacteroides stercoris ATCC 43183] |
| Q8A9B9 | HMPREF9445_00702 | 75.368 | 99 | 0 | 1217 | phospholipase, patatin family [Bacteroides clarus YIT 12056] |
| Q8A9B9 | HMPREF0969_03217 | 75.668 | 99 | 0 | 1216 | phospholipase, patatin family [Bacteroides sp. D20] |
| Q8A9B9 | HMPREF1007_02295 | 75.668 | 99 | 0 | 1216 | patatin-like phospholipase [Bacteroides sp. 4_1_36] |
| Q8A9B9 | BACUNI_02000 | 75.668 | 99 | 0 | 1216 | phospholipase, patatin family [Bacteroides uniformis ATCC 8492] |
| Q8A9B9 | HMPREF1016_03234 | 75.1 | 99 | 0 | 1212 | patatin-like phospholipase [Bacteroides eggerthii 1_2_48FAA] |

| | | | | | | |
|---|---|---|---|---|---|---|
| Q8A9B9 | BACEGG_02714 | 75.1 | 99 | 0 | 1212 | phospholipase, patatin family [Bacteroides eggerthii DSM 20697] |
| Q8A9B9 | BSFG_01260 | 69.415 | 99 | 0 | 1122 | phospholipase, patatin family [Bacteroides sp. 4_3_47FAA] |
| Q8A9B9 | HMPREF9011_01286 | 69.415 | 99 | 0 | 1121 | patatin-like phospholipase [Bacteroides sp. 3_1_40A] |
| Q8A9B9 | CUU_3874 | 69.6 | 99 | 0 | 1121 | phospholipase, patatin family [Bacteroides vulgatus PC510] |
| Q8A9B9 | BACDOR_02695 | 69.467 | 99 | 0 | 1116 | phospholipase, patatin family [Bacteroides dorei DSM 17855] |
| Q8A9B9 | HMPREF0105_2599 | 69.467 | 99 | 0 | 1115 | phospholipase, patatin family [Bacteroides sp. 3_1_33FAA] |
| Q8A9B9 | BSBG_02713 | 69.467 | 99 | 0 | 1115 | phospholipase, patatin family [Bacteroides sp. 9_1_42FAA] |
| Q8A9B9 | BSEG_02311 | 69.333 | 99 | 0 | 1113 | phospholipase, patatin family [Bacteroides dorei 5_1_36/D4] |
| Q8A9B9 | BACCOP_02003 | 65.2 | 99 | 0 | 1061 | phospholipase, patatin family [Bacteroides coprocola DSM 17136] |
| Q8A9B9 | BACPLE_00812 | 63.165 | 99 | 0 | 1032 | phospholipase, patatin family [Bacteroides plebeius DSM 17135] |
| Q8A9B9 | BACCOPRO_02612 | 63.467 | 99 | 0 | 1025 | phospholipase, patatin family [Bacteroides coprophilus DSM 18228 = JCM 13818] |
| Q8A9B9 | HMPREF1033_00307 | 50.469 | 99 | 0 | 786 | hypothetical protein [Tannerella sp. 6_1_58FAA_CT1] |
| Q8A9B9 | HMPREF9448_00383 | 46.941 | 100 | 0 | 731 | hypothetical protein [Barnesiella intestinihominis YIT 11860] |
| Q8A9B9 | HMPREF9008_01022 | 47.16 | 99 | 0 | 728 | phospholipase, patatin family [Parabacteroides sp. 20_3] |
| Q8A9B9 | HMPREF0103_1148 | 47.16 | 99 | 0 | 728 | phospholipase, patatin family [Bacteroides sp. 2_1_33B] |
| Q8A9B9 | HMPREF0619_01433 | 47.16 | 99 | 0 | 728 | phospholipase, patatin family [Parabacteroides sp. D13] |
| Q8A9B9 | HMPREF0104_01483 | 47.16 | 99 | 0 | 728 | phospholipase, patatin family [Bacteroides sp. 3_1_19] |
| Q8A9B9 | PARMER_01815 | 46.021 | 99 | 0 | 712 | phospholipase, patatin family [Parabacteroides merdae ATCC 43184] |
| Q8A9B9 | PRABACTJOHN_02312 | 48.638 | 68 | 2.28E-175 | 517 | phospholipase, patatin family [Parabacteroides johnsonii DSM 18315] |
| Q8A9B9 | HMPREF9450_00746 | 35.333 | 99 | 2.66E-158 | 481 | hypothetical protein [Alistipes indistinctus YIT 12060] |
| Q8A9B9 | ALIPUT_02121 | 30.135 | 96 | 9.61E-122 | 386 | phospholipase, patatin family [Alistipes putredinis DSM 17216] |
| Q8A9B9 | AL1_15940 | 31.174 | 96 | 2.51E-114 | 367 | Predicted esterase of the alpha-beta hydrolase superfamily [Alistipes shahii WAL 8301] |
| Q8A9B9 | PRABACTJOHN_02311 | 41.35 | 31 | 2.59E-56 | 196 | hypothetical protein [Parabacteroides johnsonii DSM 18315] |
| Q8A9B9 | BACCAC_02271 | 27.912 | 59 | 2.66E-42 | 169 | phospholipase, patatin family [Bacteroides caccae ATCC 43185] |
| Q8A9B9 | HMPREF1030_00884 | 35.862 | 37 | 4.08E-41 | 165 | hypothetical protein [Pseudomonas sp. 2_1_26] |
| Q8A9B9 | PRABACTJOHN_00370 | 27.495 | 62 | 4.24E-41 | 165 | phospholipase, patatin family [Parabacteroides johnsonii DSM 18315] |
| Q8A9B9 | HMPREF9449_00398 | 26.609 | 59 | 1.71E-40 | 163 | hypothetical protein [Odoribacter laneus YIT 12061] |
| Q8A9B9 | PARMER_00099 | 27.126 | 62 | 2.96E-40 | 162 | phospholipase, patatin family [Parabacteroides merdae ATCC 43184] |
| Q8A9B9 | CW1_1828 | 36.626 | 32 | 2.97E-40 | 154 | phospholipase, patatin family [Bacteroides xylanisolvens SD CC 2a] |
| Q8A9B9 | CUY_4079 | 27.473 | 59 | 8.04E-40 | 161 | phospholipase, patatin family [Bacteroides ovatus SD CMC 3f] |
| Q8A9B9 | HMPREF9010_01762 | 27.473 | 59 | 8.26E-40 | 161 | phospholipase, patatin family [Bacteroides sp. 3_1_23] |
| Q8A9B9 | BSCG_00789 | 27.473 | 59 | 8.26E-40 | 161 | phospholipase, patatin family [Bacteroides sp. 2_2_4] |
| Q8A9B9 | BACOVA_00420 | 27.473 | 59 | 8.26E-40 | 161 | phospholipase, patatin family [Bacteroides ovatus ATCC 8483] |
| Q8A9B9 | HMPREF1017_05053 | 27.473 | 59 | 1.11E-39 | 161 | hypothetical protein [Bacteroides ovatus 3_8_47FAA] |
| Q8A9B9 | BSGG_0662 | 27.253 | 59 | 1.42E-39 | 160 | hypothetical protein [Bacteroides sp. D2] |
| Q8A9B9 | HMPREF9007_01235 | 27.193 | 59 | 1.43E-39 | 160 | phospholipase, patatin family [Bacteroides sp. 1_1_14] |
| Q8A9B9 | BSIG_0127 | 27.193 | 59 | 1.67E-39 | 160 | hypothetical protein [Bacteroides sp. 1_1_6] |
| Q8A9B9 | HMPREF9008_04488 | 26.489 | 63 | 2.85E-39 | 160 | surface antigen variable number repeat protein [Parabacteroides sp. 20_3] |
| Q8A9B9 | CUU_1427 | 27.983 | 59 | 5.38E-39 | 159 | phospholipase, patatin family [Bacteroides vulgatus PC510] |
| Q8A9B9 | BSFG_03784 | 27.983 | 59 | 5.63E-39 | 159 | phospholipase, patatin family [Bacteroides sp. 4_3_47FAA] |
| Q8A9B9 | BFAG_03154 | 27.154 | 70 | 6.06E-39 | 159 | putative phage tail component domain protein [Bacteroides fragilis 3_1_12] |
| Q8A9B9 | HMPREF9011_01083 | 27.983 | 59 | 6.52E-39 | 159 | patatin-like phospholipase [Bacteroides sp. 3_1_40A] |
| Q8A9B9 | HMPREF1018_04237 | 27.505 | 70 | 7.41E-39 | 158 | hypothetical protein [Bacteroides sp. 2_1_56FAA] |
| Q8A9B9 | HMPREF0101_03192 | 27.505 | 70 | 7.48E-39 | 158 | phospholipase, patatin family [Bacteroides sp. 2_1_16] |
| Q8A9B9 | BSHG_1871 | 27.505 | 70 | 7.97E-39 | 158 | hypothetical protein [Bacteroides sp. 3_2_5] |
| Q8A9B9 | HMPREF0103_1929 | 26.283 | 63 | 1.43E-38 | 157 | surface antigen variable number repeat protein [Bacteroides sp. 2_1_33B] |
| Q8A9B9 | FMAG_00818 | 28.629 | 60 | 1.93E-38 | 157 | phospholipase, patatin family [Fusobacterium mortiferum ATCC 9817] |
| Q8A9B9 | BFAG_03874 | 27.22 | 67 | 2.00E-38 | 157 | phospholipase, patatin family [Bacteroides fragilis 3_1_12] |
| Q8A9B9 | HMPREF0106_03968 | 26.258 | 59 | 2.20E-38 | 157 | phospholipase, patatin family [Bacteroides sp. D22] |
| Q8A9B9 | CW3_2075 | 27.253 | 59 | 2.22E-38 | 157 | phospholipase, patatin family [Bacteroides xylanisolvens SD CC 1b] |
| Q8A9B9 | HMPREF0127_03476 | 27.253 | 59 | 2.22E-38 | 157 | hypothetical protein [Bacteroides sp. 1_1_30] |
| Q8A9B9 | HMPREF0102_02535 | 27.253 | 59 | 2.22E-38 | 157 | phospholipase, patatin family [Bacteroides sp. 2_1_22] |
| Q8A9B9 | BSAG_01109 | 27.253 | 59 | 2.22E-38 | 157 | phospholipase, patatin family [Bacteroides sp. D1] |
| Q8A9B9 | BXY_15130 | 27.253 | 59 | 2.28E-38 | 157 | Predicted esterase of the alpha-beta hydrolase superfamily [Bacteroides xylanisolvens XB1A] |
| Q8A9B9 | BACFIN_08682 | 27.033 | 59 | 2.47E-38 | 157 | phospholipase, patatin family [Bacteroides finegoldii DSM 17565] |
| Q8A9B9 | HMPREF1214_03820 | 36.111 | 33 | 3.01E-38 | 157 | NTE family protein [Bacteroides sp. HPS0048] |
| Q8A9B9 | HMPREF0105_1378 | 38.843 | 32 | 7.96E-38 | 154 | phospholipase, patatin family [Bacteroides sp. 3_1_33FAA] |
| Q8A9B9 | HMPREF1532_00269 | 37.154 | 33 | 1.78E-37 | 154 | hypothetical protein [Bacteroides salyersiae WAL 10018 = DSM 18765 = JCM 12988] |
| Q8A9B9 | HMPREF0619_00262 | 26.744 | 55 | 2.13E-37 | 154 | surface antigen variable number repeat protein [Parabacteroides sp. D13] |
| Q8A9B9 | HMPREF0104_00348 | 26.744 | 55 | 2.17E-37 | 154 | hypothetical protein [Bacteroides sp. 3_1_19] |

| | | | | | | |
|---|---|---|---|---|---|---|
| Q8A9B9 | FVAG_00457 | 28.44 | 57 | 2.18E-37 | 154 | phospholipase, patatin family [Fusobacterium varium ATCC 27725] |
| Q8A9B9 | HMPREF0105_4378 | 26.477 | 59 | 2.80E-37 | 154 | phospholipase, patatin family [Bacteroides sp. 3_1_33FAA] |
| Q8A9B9 | BACDOR_02941 | 26.477 | 59 | 2.96E-37 | 154 | phospholipase, patatin family [Bacteroides dorei DSM 17855] |
| Q8A9B9 | BSEG_00063 | 26.477 | 59 | 3.46E-37 | 153 | phospholipase, patatin family [Bacteroides dorei 5_1_36/D4] |
| Q8A9B9 | BSBG_01896 | 26.477 | 59 | 3.46E-37 | 153 | phospholipase, patatin family [Bacteroides sp. 9_1_42FAA] |
| Q8A9B9 | FUAG_00538 | 27.865 | 55 | 1.29E-36 | 152 | hypothetical protein [Fusobacterium ulcerans ATCC 49185] |
| Q8A9B9 | HMPREF0402_00731 | 27.865 | 55 | 1.75E-36 | 151 | hypothetical protein [Fusobacterium ulcerans 12-1B] |
| Q8A9B9 | HMPREF1018_03072 | 26.013 | 59 | 2.32E-36 | 151 | hypothetical protein [Bacteroides sp. 2_1_56FAA] |
| Q8A9B9 | HMPREF0101_03770 | 26.013 | 59 | 2.32E-36 | 151 | phospholipase, patatin family [Bacteroides sp. 2_1_16] |
| Q8A9B9 | HMPREF9447_00151 | 38.017 | 32 | 3.36E-36 | 149 | hypothetical protein [Bacteroides oleiciplenus YIT 12058] |
| Q8A9B9 | HMPREF9449_01091 | 37.218 | 35 | 6.18E-36 | 149 | hypothetical protein [Odoribacter laneus YIT 12061] |
| Q8A9B9 | HMPREF9466_00771 | 34.956 | 30 | 6.32E-36 | 143 | hypothetical protein [Fusobacterium necrophorum subsp. funduliforme 1_1_36S] |
| Q8A9B9 | BSHG_3165 | 25.8 | 59 | 2.22E-35 | 148 | hypothetical protein [Bacteroides sp. 3_2_5] |
| Q8A9B9 | HMPREF9007_04140 | 37.19 | 32 | 3.10E-35 | 147 | phospholipase, patatin family [Bacteroides sp. 1_1_14] |
| Q8A9B9 | HMPREF0102_01311 | 37.19 | 32 | 3.13E-35 | 147 | phospholipase, patatin family [Bacteroides sp. 2_1_22] |
| Q8A9B9 | BSAG_03782 | 37.19 | 32 | 3.13E-35 | 147 | phospholipase, patatin family [Bacteroides sp. D1] |
| Q8A9B9 | CW3_2182 | 37.19 | 32 | 3.13E-35 | 147 | phospholipase, patatin family [Bacteroides xylanisolvens SD CC 1b] |
| Q8A9B9 | CW1_0310 | 37.19 | 32 | 3.13E-35 | 147 | phospholipase, patatin family [Bacteroides xylanisolvens SD CC 2a] |
| Q8A9B9 | HMPREF1007_00232 | 27.273 | 64 | 5.08E-35 | 146 | patatin-like phospholipase [Bacteroides sp. 4_1_36] |
| Q8A9B9 | HMPREF0969_01456 | 27.273 | 64 | 1.00E-34 | 145 | phospholipase, patatin family [Bacteroides sp. D20] |
| Q8A9B9 | BACUNI_01001 | 27.273 | 64 | 1.02E-34 | 145 | phospholipase, patatin family [Bacteroides uniformis ATCC 8492] |
| Q8A9B9 | FVAG_01248 | 24.66 | 76 | 4.88E-32 | 137 | surface antigen variable number repeat protein [Fusobacterium varium ATCC 27725] |
| Q8A901 | HMPREF9007_02462 | 98.846 | 100 | 0 | 533 | phospholipase, patatin family [Bacteroides sp. 1_1_14] |
| Q8A901 | BSIG_0599 | 98.846 | 100 | 0 | 533 | hypothetical protein [Bacteroides sp. 1_1_6] |
| Q8A901 | BACCAC_00487 | 92.52 | 98 | 3.01E-178 | 494 | phospholipase, patatin family [Bacteroides caccae ATCC 43185] |
| Q8A901 | HMPREF9010_00947 | 90.909 | 97 | 6.36E-174 | 483 | phospholipase, patatin family [Bacteroides sp. 3_1_23] |
| Q8A901 | HMPREF0106_01781 | 89.883 | 99 | 1.33E-173 | 482 | serine protease [Bacteroides sp. D22] |
| Q8A901 | HMPREF0127_00560 | 89.494 | 99 | 1.05E-172 | 480 | hypothetical protein [Bacteroides sp. 1_1_30] |
| Q8A901 | CUY_4619 | 90.514 | 97 | 2.68E-172 | 479 | phospholipase, patatin family [Bacteroides ovatus SD CMC 3f] |
| Q8A901 | BSCG_02360 | 90.514 | 97 | 2.68E-172 | 479 | phospholipase, patatin family [Bacteroides sp. 2_2_4] |
| Q8A901 | HMPREF1017_03705 | 90.514 | 97 | 2.68E-172 | 479 | hypothetical protein [Bacteroides ovatus 3_8_47FAA] |
| Q8A901 | BSGG_0149 | 89.105 | 99 | 5.90E-172 | 478 | hypothetical protein [Bacteroides sp. D2] |
| Q8A901 | BACFIN_06798 | 89.37 | 98 | 9.48E-172 | 478 | phospholipase, patatin family [Bacteroides finegoldii DSM 17565] |
| Q8A901 | CW3_0809 | 89.723 | 97 | 2.91E-171 | 476 | phospholipase, patatin family [Bacteroides xylanisolvens SD CC 1b] |
| Q8A901 | HMPREF0102_02966 | 89.723 | 97 | 4.39E-171 | 476 | phospholipase, patatin family [Bacteroides sp. 2_1_22] |
| Q8A901 | BSAG_01552 | 89.723 | 97 | 4.39E-171 | 476 | phospholipase, patatin family [Bacteroides sp. D1] |
| Q8A901 | CW1_4083 | 89.723 | 97 | 4.39E-171 | 476 | phospholipase, patatin family [Bacteroides xylanisolvens SD CC 2a] |
| Q8A901 | AL1_31780 | 50 | 98 | 3.68E-91 | 274 | Predicted esterase of the alpha-beta hydrolase superfamily [Alistipes shahii WAL 8301] |
| Q8A901 | HMPREF9420_2337 | 44.599 | 99 | 5.82E-77 | 239 | phospholipase, patatin family [Prevotella salivae DSM 15606] |
| Q8A901 | HMPREF9456_02925 | 49.718 | 68 | 6.03E-59 | 193 | hypothetical protein [Dysgonomonas mossii DSM 22836] |
| Q8A901 | HMPREF1030_02876 | 49.451 | 70 | 8.74E-52 | 176 | hypothetical protein [Pseudomonas sp. 2_1_26] |
| Q8A901 | HMPREF0880_03779 | 30.496 | 94 | 3.20E-41 | 147 | NTE family protein RssA [Yokenella regensburgei ATCC 43003] |
| Q8A901 | HMPREF0485_00764 | 30.249 | 94 | 1.04E-40 | 145 | NTE family protein RssA [Klebsiella sp. 1_1_55] |
| Q8A901 | HMPREF1308_02348 | 30.605 | 94 | 1.42E-40 | 145 | NTE family protein rssA [Klebsiella pneumoniae subsp. pneumoniae WGLW5] |
| Q8A901 | HMPREF9538_04284 | 30.605 | 94 | 1.42E-40 | 145 | NTE family protein RssA [Klebsiella sp. MS 92-3] |
| Q8A901 | HMPREF1024_00512 | 30.605 | 94 | 1.42E-40 | 145 | NTE family protein rssA [Klebsiella sp. 4_1_44FAA] |
| Q8A901 | HMPREF1307_02299 | 30.605 | 94 | 1.42E-40 | 145 | NTE family protein rssA [Klebsiella pneumoniae subsp. pneumoniae WGLW3] |
| Q8A901 | ENTCAN_06436 | 30.961 | 94 | 2.96E-40 | 144 | NTE family protein RssA [Enterobacter cancerogenus ATCC 35316] |
| Q8A901 | HMPREF9543_03507 | 30.851 | 94 | 7.65E-40 | 144 | NTE family protein RssA [Escherichia coli MS 146-1] |
| Q8A901 | HMPREF9532_04293 | 30.496 | 94 | 1.09E-39 | 143 | NTE family protein RssA [Escherichia coli MS 57-2] |
| Q8A901 | HMPREF9348_01512 | 30.496 | 94 | 1.21E-39 | 143 | NTE family protein RssA [Escherichia coli MS 145-7] |
| Q8A901 | HMPREF9347_03424 | 30.496 | 94 | 1.21E-39 | 143 | NTE family protein RssA [Escherichia coli MS 124-1] |
| Q8A901 | HMPREF9553_04763 | 30.496 | 94 | 1.21E-39 | 143 | NTE family protein RssA [Escherichia coli MS 200-1] |
| Q8A901 | HMPREF9530_04333 | 30.496 | 94 | 1.21E-39 | 143 | NTE family protein RssA [Escherichia coli MS 21-1] |
| Q8A901 | HMPREF9350_03303 | 30.496 | 94 | 1.21E-39 | 143 | NTE family protein RssA [Escherichia coli MS 85-1] |
| Q8A901 | HMPREF9542_02344 | 30.496 | 94 | 1.21E-39 | 143 | NTE family protein RssA [Escherichia coli MS 117-3] |
| Q8A901 | HMPREF9539_01412 | 30.496 | 94 | 1.21E-39 | 143 | NTE family protein RssA [Escherichia coli MS 110-3] |
| Q8A901 | HMPREF9545_03662 | 30.496 | 94 | 1.21E-39 | 143 | NTE family protein RssA [Escherichia coli MS 16-3] |
| Q8A901 | HMPREF9544_04880 | 30.496 | 94 | 1.21E-39 | 143 | NTE family protein RssA [Escherichia coli MS 153-1] |
| Q8A901 | HMPREF9346_00813 | 30.496 | 94 | 1.21E-39 | 143 | NTE family protein RssA [Escherichia coli MS 119-7] |

| | | | | | | |
|---|---|---|---|---|---|---|
| Q8A901 | HMPREF9549_00985 | 30.496 | 94 | 1.21E-39 | 143 | NTE family protein RssA [Escherichia coli MS 185-1] |
| Q8A901 | HMPREF9531_00208 | 30.496 | 94 | 1.21E-39 | 143 | NTE family protein RssA [Escherichia coli MS 45-1] |
| Q8A901 | ESAG_01925 | 30.496 | 94 | 1.21E-39 | 143 | NTE family protein RssA [Escherichia sp. 3_2_53FAA] |
| Q8A901 | HMPREF9533_00200 | 30.496 | 94 | 1.21E-39 | 143 | NTE family protein RssA [Escherichia coli MS 60-1] |
| Q8A901 | HMPREF9536_02261 | 30.496 | 94 | 1.21E-39 | 143 | NTE family protein RssA [Escherichia coli MS 84-1] |
| Q8A901 | HMPREF9349_05315 | 30.496 | 94 | 1.25E-39 | 143 | NTE family protein RssA [Escherichia coli MS 79-10] |
| Q8A901 | SSJG_01324 | 30.496 | 94 | 1.25E-39 | 143 | NTE family protein RssA [Escherichia coli D9] |
| Q8A901 | HMPREF9345_04781 | 30.496 | 94 | 1.25E-39 | 143 | NTE family protein RssA [Escherichia coli MS 107-1] |
| Q8A901 | HMPREF9541_05257 | 30.496 | 94 | 1.25E-39 | 143 | NTE family protein RssA [Escherichia coli MS 116-1] |
| Q8A901 | HMPREF9540_02921 | 30.496 | 94 | 1.25E-39 | 143 | NTE family protein RssA [Escherichia coli MS 115-1] |
| Q8A901 | HMPREF9547_01760 | 30.496 | 94 | 1.25E-39 | 143 | NTE family protein RssA [Escherichia coli MS 175-1] |
| Q8A901 | HMPREF9548_04614 | 30.496 | 94 | 1.25E-39 | 143 | NTE family protein RssA [Escherichia coli MS 182-1] |
| Q8A901 | ESCG_03269 | 30.496 | 94 | 1.25E-39 | 143 | NTE family protein rssA [Escherichia sp. 1_1_43] |
| Q8A901 | ESBG_01451 | 30.496 | 94 | 1.25E-39 | 143 | NTE family protein rssA [Escherichia sp. 4_1_40B] |
| Q8A901 | HMPREF9535_01952 | 30.496 | 94 | 1.25E-39 | 143 | NTE family protein RssA [Escherichia coli MS 78-1] |
| Q8A901 | HMPREF9552_03846 | 30.496 | 94 | 1.28E-39 | 143 | NTE family protein RssA [Escherichia coli MS 198-1] |
| Q8A901 | ECSE_1284 | 30.496 | 94 | 1.28E-39 | 143 | conserved hypothetical protein [Escherichia coli SE11] |
| Q8A901 | ECSF_1215 | 30.496 | 94 | 1.28E-39 | 143 | conserved hypothetical protein [Escherichia coli SE15] |
| Q8A901 | HMPREF9534_04406 | 30.496 | 94 | 1.28E-39 | 143 | NTE family protein RssA [Escherichia coli MS 69-1] |
| Q8A901 | HMPREF9551_04057 | 30.496 | 94 | 1.33E-39 | 143 | NTE family protein RssA [Escherichia coli MS 196-1] |
| Q8A901 | HMPREF0986_04085 | 30.496 | 94 | 1.67E-39 | 143 | NTE family protein rssA [Escherichia coli 4_1_47FAA] |
| Q8A901 | HMPREF9550_00115 | 30.496 | 94 | 1.72E-39 | 143 | NTE family protein RssA [Escherichia coli MS 187-1] |
| Q8A901 | HMPREF9428_01548 | 30.282 | 94 | 5.29E-39 | 141 | NTE family protein rssA [Citrobacter freundii 4_7_47CFAA] |
| Q8A901 | HMPREF0201_02441 | 30.035 | 94 | 6.73E-39 | 141 | NTE family protein RssA [Cedecea davisae DSM 4568] |
| Q8A901 | CIT292_06864 | 29.93 | 94 | 2.47E-38 | 139 | NTE family protein RssA [Citrobacter youngae ATCC 29220] |
| Q8A901 | HMPREF1014_00937 | 35.102 | 94 | 3.07E-38 | 138 | hypothetical protein [Bacillus sp. 7_6_55CFAA_CT2] |
| Q8A901 | HMPREF1705_01953 | 31.559 | 93 | 4.64E-37 | 135 | phospholipase, patatin family [Anaerobaculum hydrogeniformans ATCC BAA-1850] |
| Q8A901 | HMPREF9418_2280 | 37.5 | 69 | 2.71E-34 | 129 | patatin family phospholipase [Neisseria macacae ATCC 33926] |
| Q8A901 | HMPREF0322_03075 | 30.233 | 98 | 6.29E-34 | 127 | phospholipase, patatin family [Desulfitobacterium hafniense DP7] |
| Q8A900 | HMPREF9007_02463 | 99.387 | 100 | 0 | 1353 | conserved hypothetical protein [Bacteroides sp. 1_1_14] |
| Q8A900 | BSIG_0598 | 99.387 | 100 | 0 | 1352 | hypothetical protein [Bacteroides sp. 1_1_6] |
| Q8A900 | CW3_0810 | 89.74 | 100 | 0 | 1221 | conserved hypothetical protein [Bacteroides xylanisolvens SD CC 1b] |
| Q8A900 | CW1_4082 | 89.74 | 100 | 0 | 1221 | conserved hypothetical protein [Bacteroides xylanisolvens SD CC 2a] |
| Q8A900 | BACOVA_04123 | 89.587 | 100 | 0 | 1221 | hypothetical protein [Bacteroides ovatus ATCC 8483] |
| Q8A900 | HMPREF0102_02965 | 89.433 | 100 | 0 | 1219 | hypothetical protein [Bacteroides sp. 2_1_22] |
| Q8A900 | BSAG_01551 | 89.433 | 100 | 0 | 1219 | hypothetical protein [Bacteroides sp. D1] |
| Q8A900 | HMPREF9010_00946 | 89.433 | 100 | 0 | 1219 | conserved hypothetical protein [Bacteroides sp. 3_1_23] |
| Q8A900 | HMPREF0127_00561 | 89.587 | 100 | 0 | 1218 | hypothetical protein [Bacteroides sp. 1_1_30] |
| Q8A900 | BSGG_0151 | 89.587 | 100 | 0 | 1217 | hypothetical protein [Bacteroides sp. D2] |
| Q8A900 | HMPREF1017_03706 | 89.28 | 100 | 0 | 1216 | hypothetical protein [Bacteroides ovatus 3_8_47FAA] |
| Q8A900 | BACCELL_02887 | 88.361 | 100 | 0 | 1215 | hypothetical protein [Bacteroides cellulosilyticus DSM 14838] |
| Q8A900 | BACFIN_06799 | 88.974 | 100 | 0 | 1214 | hypothetical protein [Bacteroides finegoldii DSM 17565] |
| Q8A900 | BXY_06400 | 89.628 | 99 | 0 | 1206 | hypothetical protein [Bacteroides xylanisolvens XB1A] |
| Q8A900 | CUY_4620 | 89.319 | 99 | 0 | 1204 | conserved hypothetical protein [Bacteroides ovatus SD CMC 3f] |
| Q8A900 | BSCG_02361 | 89.474 | 99 | 0 | 1202 | hypothetical protein [Bacteroides sp. 2_2_4] |
| Q8A900 | HMPREF1532_00979 | 74.178 | 98 | 0 | 986 | hypothetical protein [Bacteroides salyersiae WAL 10018 = DSM 18765 = JCM 12988] |
| Q8A900 | BSEG_00316 | 77.027 | 68 | 0 | 701 | hypothetical protein [Bacteroides dorei 5_1_36/D4] |
| Q8A900 | BSBG_01478 | 77.027 | 68 | 0 | 700 | hypothetical protein [Bacteroides sp. 9_1_42FAA] |
| Q8A900 | BACDOR_04321 | 76.802 | 68 | 0 | 699 | hypothetical protein [Bacteroides dorei DSM 17855] |
| Q8A900 | HMPREF0105_0259 | 76.351 | 68 | 0 | 697 | hypothetical protein [Bacteroides sp. 3_1_33FAA] |
| Q8A900 | CUU_1073 | 78.538 | 65 | 0 | 694 | conserved hypothetical protein [Bacteroides vulgatus PC510] |
| Q8A900 | BSFG_02978 | 78.538 | 65 | 0 | 694 | hypothetical protein [Bacteroides sp. 4_3_47FAA] |
| Q8A900 | HMPREF9011_03678 | 78.538 | 65 | 0 | 694 | hypothetical protein [Bacteroides sp. 3_1_40A] |
| Q8A900 | HMPREF1016_01117 | 69.124 | 66 | 0 | 620 | hypothetical protein [Bacteroides eggerthii 1_2_48FAA] |
| Q8A900 | BACEGG_00660 | 69.124 | 66 | 0 | 620 | hypothetical protein [Bacteroides eggerthii DSM 20697] |
| Q8A6N9 | BSIG_4403 | 99.524 | 100 | 0 | 1742 | hypothetical protein [Bacteroides sp. 1_1_6] |
| Q8A6N9 | BACCAC_03838 | 89.881 | 100 | 0 | 1607 | peptidase, S9A/B/C family, catalytic domain protein [Bacteroides caccae ATCC 43185] |
| Q8A6N9 | BFAG_03532 | 82.976 | 100 | 0 | 1503 | peptidase, S9A/B/C family, catalytic domain protein [Bacteroides fragilis 3_1_12] |
| Q8A6N9 | HMPREF0101_02878 | 82.143 | 100 | 0 | 1493 | peptidase, S9A/B/C family, catalytic domain protein [Bacteroides sp. 2_1_16] |
| Q8A6N9 | HMPREF1018_03843 | 82.143 | 100 | 0 | 1492 | hypothetical protein [Bacteroides sp. 2_1_56FAA] |

| | | | | | | |
|---|---|---|---|---|---|---|
| Q8A6N9 | BSHG_2181 | 82.024 | 100 | 0 | 1492 | hypothetical protein [Bacteroides sp. 3_2_5] |
| Q8A6N9 | HMPREF1007_03537 | 67.922 | 98 | 0 | 1211 | prolyl oligopeptidase [Bacteroides sp. 4_1_36] |
| Q8A6N9 | HMPREF0969_01197 | 67.801 | 98 | 0 | 1209 | putative phage prohead protease, HK97 family [Bacteroides sp. D20] |
| Q8A6N9 | BACUNI_02439 | 67.922 | 98 | 0 | 1207 | peptidase, S9A/B/C family, catalytic domain protein [Bacteroides uniformis ATCC 8492] |
| Q8A6N9 | HMPREF9446_01327 | 67.801 | 98 | 0 | 1207 | peptidase, S9A/B/C family, catalytic domain protein [Bacteroides fluxus YIT 12057] |
| Q8A6N9 | HMPREF9445_02770 | 65.632 | 99 | 0 | 1181 | peptidase, S9A/B/C family, catalytic domain protein [Bacteroides clarus YIT 12056] |
| Q8A6N9 | BACEGG_00665 | 66.264 | 98 | 0 | 1175 | peptidase, S9A/B/C family, catalytic domain protein [Bacteroides eggerthii DSM 20697] |
| Q8A6N9 | HMPREF1016_01121 | 66.264 | 98 | 0 | 1174 | prolyl oligopeptidase [Bacteroides eggerthii 1_2_48FAA] |
| Q8A6N9 | BACSTE_00111 | 65.463 | 99 | 0 | 1174 | peptidase, S9A/B/C family, catalytic domain protein [Bacteroides stercoris ATCC 43183] |
| Q8A6N9 | HMPREF9008_00988 | 48.727 | 98 | 0 | 804 | peptidase, S9A/B/C family, catalytic domain protein [Parabacteroides sp. 20_3] |
| Q8A6N9 | HMPREF0619_01467 | 48.606 | 98 | 0 | 803 | peptidase, S9A/B/C family, catalytic domain protein [Parabacteroides sp. D13] |
| Q8A6N9 | HMPREF9456_01244 | 48.104 | 99 | 0 | 801 | hypothetical protein [Dysgonomonas mossii DSM 22836] |
| Q8A6N9 | HMPREF0103_1113 | 48.606 | 98 | 0 | 800 | peptidase, S9A/B/C family, catalytic domain protein [Bacteroides sp. 2_1_33B] |
| Q8A6N9 | HMPREF0104_01518 | 48.485 | 98 | 0 | 800 | alanyl dipeptidyl peptidase [Bacteroides sp. 3_1_19] |
| Q8A6N9 | PRABACTJOHN_02222 | 46.437 | 100 | 0 | 799 | peptidase, S9A/B/C family, catalytic domain protein [Parabacteroides johnsonii DSM 18315] |
| Q8A6N9 | HMPREF9455_02957 | 46.968 | 99 | 0 | 793 | hypothetical protein [Dysgonomonas gadei ATCC BAA-286] |
| Q8A6N9 | PARMER_01851 | 45.724 | 100 | 0 | 787 | peptidase, S9A/B/C family, catalytic domain protein [Parabacteroides merdae ATCC 43184] |
| Q8A6N9 | HMPREF0103_3070 | 46.229 | 98 | 0 | 760 | peptidase, S9A/B/C family, catalytic domain protein [Bacteroides sp. 2_1_33B] |
| Q8A6N9 | HMPREF9008_01869 | 46.22 | 98 | 0 | 759 | peptidase, S9A/B/C family, catalytic domain protein [Parabacteroides sp. 20_3] |
| Q8A6N9 | PARMER_03244 | 46.228 | 99 | 0 | 759 | peptidase, S9A/B/C family, catalytic domain protein [Parabacteroides merdae ATCC 43184] |
| Q8A6N9 | HMPREF0619_03491 | 46.107 | 98 | 0 | 755 | peptidase, S9A/B/C family, catalytic domain protein [Parabacteroides sp. D13] |
| Q8A6N9 | HMPREF0104_03313 | 45.864 | 98 | 0 | 750 | alanyl dipeptidyl peptidase [Bacteroides sp. 3_1_19] |
| Q8A6N9 | BSFG_00784 | 44.471 | 100 | 0 | 739 | peptidase, S9A/B/C family, catalytic domain protein [Bacteroides sp. 4_3_47FAA] |
| Q8A6N9 | HMPREF9011_03158 | 44.471 | 100 | 0 | 739 | alanyl dipeptidyl peptidase [Bacteroides sp. 3_1_40A] |
| Q8A6N9 | CUU_2819 | 44.352 | 100 | 0 | 738 | peptidase, S9A/B/C familie, catalytic domain protein [Bacteroides vulgatus PC510] |
| Q8A6N9 | HMPREF0105_1477 | 44.891 | 98 | 0 | 736 | peptidase, S9A/B/C family, catalytic domain protein [Bacteroides sp. 3_1_33FAA] |
| Q8A6N9 | BACDOR_00752 | 44.891 | 98 | 0 | 736 | peptidase, S9A/B/C family, catalytic domain protein [Bacteroides dorei DSM 17855] |
| Q8A6N9 | BSEG_01192 | 44.471 | 100 | 0 | 735 | peptidase, S9A/B/C family, catalytic domain protein [Bacteroides dorei 5_1_36/D4] |
| Q8A6N9 | BSBG_00318 | 44.471 | 100 | 0 | 735 | peptidase, S9A/B/C family, catalytic domain protein [Bacteroides sp. 9_1_42FAA] |
| Q8A6N9 | BACPLE_01436 | 41.232 | 100 | 0 | 692 | peptidase, S9A/B/C family, catalytic domain protein [Bacteroides plebeius DSM 17135] |
| Q8A6N9 | PREVCOP_04746 | 40.767 | 99 | 0 | 650 | peptidase, S9A/B/C family, catalytic domain protein [Prevotella copri DSM 18205] |
| Q8A6N9 | HMPREF9449_00768 | 36.364 | 100 | 0 | 565 | hypothetical protein [Odoribacter laneus YIT 12061] |
| Q8A6N9 | HMPREF9448_01354 | 27.824 | 55 | 3.06E-36 | 150 | hypothetical protein [Barnesiella intestinihominis YIT 11860] |
| Q8A6N9 | HMPREF1033_00381 | 26.261 | 55 | 6.64E-35 | 147 | hypothetical protein [Tannerella sp. 6_1_58FAA_CT1] |
| Q8A6N9 | HMPREF9449_00790 | 26.263 | 45 | 3.35E-34 | 144 | hypothetical protein [Odoribacter laneus YIT 12061] |
| Q8A6N9 | HMPREF1016_02497 | 26.337 | 55 | 9.91E-34 | 143 | prolyl oligopeptidase [Bacteroides eggerthii 1_2_48FAA] |
| Q8A6N9 | BACEGG_02554 | 26.337 | 55 | 1.25E-33 | 143 | peptidase, S9A/B/C family, catalytic domain protein [Bacteroides eggerthii DSM 20697] |
| Q8A6N9 | HMPREF1210_02618 | 26.441 | 58 | 1.36E-33 | 142 | hypothetical protein [Paenisporosarcina sp. HGH0030] |
| Q8A6N9 | CUY_1699 | 24.371 | 59 | 5.98E-33 | 140 | peptidase, S9A/B/C familie, catalytic domain protein [Bacteroides ovatus SD CMC 3f] |
| Q8A6N9 | BSCG_04041 | 24.371 | 59 | 6.57E-33 | 140 | peptidase, S9A/B/C family, catalytic domain protein [Bacteroides sp. 2_2_4] |
| Q8A6N9 | BSGG_4123 | 25.275 | 53 | 6.63E-33 | 140 | hypothetical protein [Bacteroides sp. D2] |
| Q8A6N9 | HMPREF9010_01642 | 25.275 | 53 | 7.71E-33 | 140 | prolyl oligopeptidase family protein [Bacteroides sp. 3_1_23] |
| Q8A6N9 | BACFIN_06575 | 25.275 | 53 | 7.71E-33 | 140 | peptidase, S9A/B/C family, catalytic domain protein [Bacteroides finegoldii DSM 17565] |
| Q8A6N9 | BACOVA_03695 | 25.275 | 53 | 7.92E-33 | 140 | peptidase, S9A/B/C family, catalytic domain protein [Bacteroides ovatus ATCC 8483] |
| Q8A6N9 | HMPREF1017_01228 | 25.275 | 53 | 8.81E-33 | 140 | hypothetical protein [Bacteroides ovatus 3_8_47FAA] |
| Q8A6N9 | HMPREF0106_03341 | 25.275 | 53 | 8.81E-33 | 140 | prolyl oligopeptidase family protein [Bacteroides sp. D22] |
| Q8A6N9 | HMPREF0102_01335 | 25.275 | 53 | 9.81E-33 | 140 | peptidase, S9A/B/C family, catalytic domain protein [Bacteroides sp. 2_1_22] |
| Q8A6N9 | BSAG_02778 | 25.275 | 53 | 9.81E-33 | 140 | peptidase, S9A/B/C family, catalytic domain protein [Bacteroides sp. D1] |
| Q8A6N9 | CW3_1188 | 25.275 | 53 | 9.81E-33 | 140 | peptidase, S9A/B/C familie, catalytic domain protein [Bacteroides xylanisolvens SD CC 1b] |
| Q8A6N9 | CW1_0332 | 25.275 | 53 | 9.81E-33 | 140 | peptidase, S9A/B/C familie, catalytic domain protein [Bacteroides xylanisolvens SD CC 2a] |
| Q8A6N9 | BXY_01510 | 25.055 | 53 | 1.03E-32 | 140 | Dipeptidyl aminopeptidases/acylaminoacyl-peptidases [Bacteroides xylanisolvens XB1A] |
| Q8A6N9 | BACCELL_04347 | 25.572 | 55 | 1.56E-32 | 139 | peptidase, S9A/B/C family, catalytic domain protein [Bacteroides cellulosilyticus DSM 14838] |
| Q8A6N9 | BACINT_02525 | 25.259 | 55 | 2.15E-32 | 139 | peptidase, S9A/B/C family, catalytic domain protein [Bacteroides intestinalis DSM 17393] |
| Q8A6N9 | HMPREF9447_00898 | 25.155 | 55 | 4.01E-32 | 138 | hypothetical protein [Bacteroides oleiciplenus YIT 12058] |
| Q8A6N9 | HMPREF9007_04165 | 25.442 | 52 | 4.97E-32 | 137 | prolyl oligopeptidase family protein [Bacteroides sp. 1_1_14] |
| Q8A6N9 | BSIG_1028 | 25.442 | 52 | 4.97E-32 | 137 | hypothetical protein [Bacteroides sp. 1_1_6] |
| Q8A6N9 | HMPREF9456_01334 | 25.852 | 55 | 8.74E-32 | 137 | hypothetical protein [Dysgonomonas mossii DSM 22836] |
| Q8A6N9 | HMPREF0673_02840 | 26.632 | 45 | 9.16E-32 | 137 | peptidase, S9A/B/C family, catalytic domain protein [Prevotella stercorea DSM 18206] |
| Q8A6N9 | HMPREF0969_03426 | 26.132 | 55 | 2.02E-31 | 136 | peptidase, S9A/B/C family, catalytic domain protein [Bacteroides sp. D20] |
| Q8A6N9 | HMPREF1007_01374 | 26.132 | 55 | 2.64E-31 | 135 | prolyl oligopeptidase [Bacteroides sp. 4_1_36] |

| | | | | | | |
|---|---|---|---|---|---|---|
| Q8A6N9 | BACCOPRO_00063 | 25.553 | 55 | 4.38E-31 | 135 | peptidase, S9A/B/C family, catalytic domain protein [Bacteroides coprophilus DSM 18228 = JCM 13818] |
| Q8A6N9 | HMPREF9455_01886 | 25.455 | 55 | 6.69E-31 | 134 | hypothetical protein [Dysgonomonas gadei ATCC BAA-286] |
| Q8A6N9 | BACUNI_02780 | 25.926 | 55 | 7.42E-31 | 134 | peptidase, S9A/B/C family, catalytic domain protein [Bacteroides uniformis ATCC 8492] |
| Q8A6K2 | BSIG_5237 | 100 | 100 | 0 | 1206 | signal peptide peptidase SppA, 67K type [Bacteroides sp. 1_1_6] |
| Q8A6K2 | HMPREF9007_01978 | 99.663 | 100 | 0 | 1199 | signal peptide peptidase SppA, 67K type [Bacteroides sp. 1_1_14] |
| Q8A6K2 | BXY_17770 | 88.176 | 100 | 0 | 1090 | signal peptide peptidase A. Serine peptidase. MEROPS family S49 [Bacteroides xylanisolvens XB1A] |
| Q8A6K2 | HMPREF0102_04197 | 88.176 | 100 | 0 | 1090 | signal peptide peptidase SppA, 67K type [Bacteroides sp. 2_1_22] |
| Q8A6K2 | BSAG_03511 | 88.176 | 100 | 0 | 1090 | signal peptide peptidase SppA, 67K type [Bacteroides sp. D1] |
| Q8A6K2 | HMPREF0106_02790 | 88.176 | 100 | 0 | 1090 | signal peptide peptidase SppA, 67K type [Bacteroides sp. D22] |
| Q8A6K2 | CW3_4739 | 88.176 | 100 | 0 | 1090 | signal peptide peptidase SppA, 67K type [Bacteroides xylanisolvens SD CC 1b] |
| Q8A6K2 | CW1_4635 | 88.176 | 100 | 0 | 1090 | signal peptide peptidase SppA, 67K type [Bacteroides xylanisolvens SD CC 2a] |
| Q8A6K2 | HMPREF0127_04404 | 87.669 | 100 | 0 | 1085 | signal peptide peptidase SppA, 67K type [Bacteroides sp. 1_1_30] |
| Q8A6K2 | HMPREF9010_04722 | 88.176 | 100 | 0 | 1084 | signal peptide peptidase SppA, 67K type [Bacteroides sp. 3_1_23] |
| Q8A6K2 | BSGG_4715 | 88.176 | 100 | 0 | 1084 | signal peptide peptidase SppA, 67K type [Bacteroides sp. D2] |
| Q8A6K2 | BACOVA_02940 | 87.838 | 100 | 0 | 1082 | signal peptide peptidase SppA, 67K type [Bacteroides ovatus ATCC 8483] |
| Q8A6K2 | HMPREF1017_04443 | 88.007 | 100 | 0 | 1082 | signal peptide peptidase SppA, 67K type [Bacteroides ovatus 3_8_47FAA] |
| Q8A6K2 | CUY_2234 | 87.838 | 100 | 0 | 1080 | signal peptide peptidase SppA, 67K type [Bacteroides ovatus SD CMC 3f] |
| Q8A6K2 | BSCG_00026 | 87.838 | 100 | 0 | 1080 | signal peptide peptidase SppA, 67K type [Bacteroides sp. 2_2_4] |
| Q8A6K2 | BACCAC_03765 | 87.5 | 100 | 0 | 1074 | signal peptide peptidase SppA, 67K type [Bacteroides caccae ATCC 43185] |
| Q8A6K2 | BACFIN_06171 | 86.318 | 100 | 0 | 1065 | signal peptide peptidase SppA, 67K type [Bacteroides finegoldii DSM 17565] |
| Q8A6K2 | HMPREF1214_04296 | 82.77 | 100 | 0 | 1029 | signal peptide peptidase SppA, 67K type [Bacteroides sp. HPS0048] |
| Q8A6K2 | HMPREF1532_03953 | 84.628 | 100 | 0 | 1027 | signal peptide peptidase SppA, 67K type [Bacteroides salyersiae WAL 10018 = DSM 18765 = JCM 12988] |
| Q8A6K2 | HMPREF1018_03441 | 77.534 | 100 | 0 | 982 | signal peptide peptidase SppA, 67K type [Bacteroides sp. 2_1_56FAA] |
| Q8A6K2 | BFAG_04481 | 77.365 | 100 | 0 | 979 | signal peptide peptidase SppA, 67K type [Bacteroides fragilis 3_1_12] |
| Q8A6K2 | HMPREF0101_03398 | 77.196 | 100 | 0 | 979 | signal peptide peptidase SppA, 67K type [Bacteroides sp. 2_1_16] |
| Q8A6K2 | BSHG_2803 | 77.365 | 100 | 0 | 978 | signal peptide peptidase SppA, 67K type [Bacteroides sp. 3_2_5] |
| Q8A6K2 | HMPREF9446_01284 | 68.919 | 100 | 0 | 873 | signal peptide peptidase SppA [Bacteroides fluxus YIT 12057] |
| Q8A6K2 | BACUNI_02479 | 70.101 | 100 | 0 | 866 | signal peptide peptidase SppA, 67K type [Bacteroides uniformis ATCC 8492] |
| Q8A6K2 | HMPREF1007_03691 | 70.101 | 100 | 0 | 866 | signal peptide peptidase SppA [Bacteroides sp. 4_1_36] |
| Q8A6K2 | BACEGG_00527 | 68.581 | 100 | 0 | 862 | signal peptide peptidase SppA, 67K type [Bacteroides eggerthii DSM 20697] |
| Q8A6K2 | HMPREF1016_03692 | 68.412 | 100 | 0 | 861 | signal peptide peptidase SppA [Bacteroides eggerthii 1_2_48FAA] |
| Q8A6K2 | HMPREF9445_03245 | 68.243 | 100 | 0 | 857 | signal peptide peptidase SppA [Bacteroides clarus YIT 12056] |
| Q8A6K2 | HMPREF0969_00725 | 70.101 | 100 | 0 | 852 | signal peptide peptidase SppA, 67K type [Bacteroides sp. D20] |
| Q8A6K2 | HMPREF9447_03349 | 69.036 | 99 | 0 | 845 | signal peptide peptidase SppA, 67K type [Bacteroides oleiciplenus YIT 12058] |
| Q8A6K2 | BACCELL_01072 | 67.682 | 99 | 0 | 832 | signal peptide peptidase SppA, 67K type [Bacteroides cellulosilyticus DSM 14838] |
| Q8A6K2 | BACINT_03271 | 67.005 | 99 | 0 | 828 | signal peptide peptidase SppA, 67K type [Bacteroides intestinalis DSM 17393] |
| Q8A6K2 | BACSTE_02920 | 67.831 | 92 | 0 | 791 | signal peptide peptidase SppA, 67K type [Bacteroides stercoris ATCC 43183] |
| Q8A6K2 | HMPREF0105_4531 | 60.702 | 100 | 0 | 741 | signal peptide peptidase SppA, 67K type [Bacteroides sp. 3_1_33FAA] |
| Q8A6K2 | CUU_3737 | 60.368 | 100 | 0 | 737 | signal peptide peptidase SppA, 67K type [Bacteroides vulgatus PC510] |
| Q8A6K2 | BSEG_04329 | 60.535 | 100 | 0 | 736 | signal peptide peptidase SppA, 67K type [Bacteroides dorei 5_1_36/D4] |
| Q8A6K2 | BACDOR_02450 | 60.535 | 100 | 0 | 736 | signal peptide peptidase SppA, 67K type [Bacteroides dorei DSM 17855] |
| Q8A6K2 | BSFG_02121 | 60.201 | 100 | 0 | 735 | signal peptide peptidase SppA, 67K type [Bacteroides sp. 4_3_47FAA] |
| Q8A6K2 | HMPREF9011_00004 | 60.201 | 100 | 0 | 735 | protease IV [Bacteroides sp. 3_1_40A] |
| Q8A6K2 | BACCOP_00445 | 60.606 | 100 | 0 | 732 | signal peptide peptidase SppA, 67K type [Bacteroides coprocola DSM 17136] |
| Q8A6K2 | BSBG_04376 | 60.201 | 100 | 0 | 731 | signal peptide peptidase SppA, 67K type [Bacteroides sp. 9_1_42FAA] |
| Q8A6K2 | BACCOPRO_01364 | 56.998 | 100 | 0 | 694 | signal peptide peptidase SppA, 67K type [Bacteroides coprophilus DSM 18228 = JCM 13818] |
| Q8A6K2 | BACPLE_01534 | 56.83 | 100 | 0 | 686 | signal peptide peptidase SppA, 67K type [Bacteroides plebeius DSM 17135] |
| Q8A6K2 | HMPREF9455_03155 | 51.007 | 100 | 0 | 595 | signal peptide peptidase SppA, 67K type [Dysgonomonas gadei ATCC BAA-286] |
| Q8A6K2 | HMPREF9456_00216 | 49.155 | 100 | 0 | 578 | signal peptide peptidase SppA, 67K type [Dysgonomonas mossii DSM 22836] |
| Q8A6K2 | HMPREF9442_02060 | 49.241 | 99 | 0 | 566 | signal peptide peptidase SppA [Paraprevotella xylaniphila YIT 11841] |
| Q8A6K2 | HMPREF9441_00749 | 49.241 | 99 | 0 | 563 | signal peptide peptidase SppA [Paraprevotella clara YIT 11840] |
| Q8A6K2 | HMPREF1475_00617 | 48.074 | 100 | 0 | 561 | signal peptide peptidase SppA, 67K type [Prevotella oralis HGA0225] |
| Q8A6K2 | HMPREF0673_00209 | 46.411 | 100 | 0 | 548 | signal peptide peptidase SppA [Prevotella stercorea DSM 18206] |
| Q8A6K2 | HMPREF1033_03183 | 46.791 | 99 | 1.30E-176 | 516 | signal peptide peptidase SppA, 67K type [Tannerella sp. 6_1_58FAA_CT1] |
| Q8A6K2 | PARMER_00097 | 46.231 | 99 | 1.81E-175 | 513 | signal peptide peptidase SppA, 67K type [Parabacteroides merdae ATCC 43184] |
| Q8A6K2 | HMPREF9008_02733 | 45.638 | 99 | 3.11E-175 | 513 | signal peptide peptidase SppA, 67K type [Parabacteroides sp. 20_3] |
| Q8A6K2 | HMPREF0103_1292 | 46.988 | 97 | 2.97E-174 | 510 | signal peptide peptidase SppA, 67K type [Bacteroides sp. 2_1_33B] |
| Q8A6K2 | HMPREF0619_01105 | 46.988 | 97 | 1.42E-173 | 508 | signal peptide peptidase SppA, 67K type [Parabacteroides sp. D13] |
| Q8A6K2 | HMPREF9448_00100 | 46.167 | 99 | 1.94E-173 | 508 | signal peptide peptidase SppA, 67K type [Barnesiella intestinihominis YIT 11860] |
| Q8A6K2 | HMPREF0104_01029 | 46.644 | 97 | 1.05E-172 | 506 | signal peptide peptidase SppA, 67K type [Bacteroides sp. 3_1_19] |

| | | | | | | |
|---|---|---|---|---|---|---|
| Q8A6K2 | PREVCOP_03542 | 45.274 | 100 | 1.45E-172 | 506 | signal peptide peptidase SppA, 67K type [Prevotella copri DSM 18205] |
| Q8A6K2 | PRABACTJOHN_00365 | 48.767 | 88 | 2.03E-168 | 493 | signal peptide peptidase SppA, 67K type [Parabacteroides johnsonii DSM 18315] |
| Q8A6K2 | HMPREF9449_02554 | 43.493 | 99 | 2.23E-165 | 488 | signal peptide peptidase SppA, 67K type [Odoribacter laneus YIT 12061] |
| Q8A6K2 | HMPREF9420_0821 | 42.14 | 100 | 4.33E-154 | 459 | signal peptide peptidase SppA, 67K type [Prevotella salivae DSM 15606] |
| Q8A6K2 | HMPREF9450_00758 | 38.5 | 100 | 4.63E-130 | 397 | signal peptide peptidase SppA, 67K type [Alistipes indistinctus YIT 12060] |
| Q8A6K2 | AL1_16080 | 37.248 | 99 | 4.82E-128 | 392 | signal peptide peptidase SppA, 67K type [Alistipes shahii WAL 8301] |
| Q8A6K2 | ALIPUT_02108 | 41.098 | 88 | 8.06E-125 | 384 | signal peptide peptidase SppA, 67K type [Alistipes putredinis DSM 17216] |
| Q8A6K2 | HMPREF9538_03670 | 38.241 | 92 | 8.77E-96 | 309 | signal peptide peptidase SppA [Klebsiella sp. MS 92-3] |
| Q8A6K2 | HMPREF0485_04543 | 38.241 | 92 | 2.38E-95 | 308 | signal peptide peptidase SppA, 67K type [Klebsiella sp. 1_1_55] |
| Q8A6K2 | HMPREF1308_01348 | 38.061 | 92 | 4.54E-95 | 307 | protease 4 [Klebsiella pneumoniae subsp. pneumoniae WGLW5] |
| Q8A6K2 | HMPREF1024_03570 | 38.061 | 92 | 4.54E-95 | 307 | protease 4 [Klebsiella sp. 4_1_44FAA] |
| Q8A6K2 | ENTCAN_05736 | 37.276 | 92 | 8.35E-95 | 306 | signal peptide peptidase SppA, 67K type [Enterobacter cancerogenus ATCC 35316] |
| Q8A6K2 | HMPREF1307_03247 | 38.241 | 92 | 1.66E-94 | 306 | protease 4 [Klebsiella pneumoniae subsp. pneumoniae WGLW3] |
| Q8A6K2 | ENC_14690 | 37.226 | 90 | 1.68E-94 | 306 | signal peptide peptidase SppA, 67K type [Enterobacter cloacae subsp. cloacae NCTC 9394] |
| Q8A6K2 | HMPREF9552_03126 | 36.528 | 90 | 6.84E-93 | 303 | signal peptide peptidase SppA, 67K type [Escherichia coli MS 198-1] |
| Q8A6K2 | ESBG_00912 | 36.528 | 90 | 1.58E-92 | 301 | protease 4 [Escherichia sp. 4_1_40B] |
| Q8A6K2 | HMPREF9553_00668 | 36.528 | 90 | 1.62E-92 | 301 | signal peptide peptidase SppA, 67K type [Escherichia coli MS 200-1] |
| Q8A6K2 | HMPREF9532_03193 | 36.528 | 90 | 1.62E-92 | 301 | signal peptide peptidase SppA, 67K type [Escherichia coli MS 57-2] |
| Q8A6K2 | HMPREF9530_03735 | 36.528 | 90 | 1.62E-92 | 301 | signal peptide peptidase SppA, 67K type [Escherichia coli MS 21-1] |
| Q8A6K2 | HMPREF9545_01074 | 36.528 | 90 | 1.62E-92 | 301 | signal peptide peptidase SppA, 67K type [Escherichia coli MS 16-3] |
| Q8A6K2 | HMPREF9544_05148 | 36.528 | 90 | 1.62E-92 | 301 | signal peptide peptidase SppA, 67K type [Escherichia coli MS 153-1] |
| Q8A6K2 | HMPREF9549_04795 | 36.528 | 90 | 1.62E-92 | 301 | signal peptide peptidase SppA, 67K type [Escherichia coli MS 185-1] |
| Q8A6K2 | HMPREF9531_05162 | 36.528 | 90 | 1.62E-92 | 301 | signal peptide peptidase SppA, 67K type [Escherichia coli MS 45-1] |
| Q8A6K2 | HMPREF9533_04455 | 36.528 | 90 | 1.62E-92 | 301 | signal peptide peptidase SppA, 67K type [Escherichia coli MS 60-1] |
| Q8A6K2 | HMPREF9539_03584 | 36.528 | 90 | 1.78E-92 | 300 | signal peptide peptidase SppA, 67K type [Escherichia coli MS 110-3] |
| Q8A6K2 | ESAG_02445 | 36.528 | 90 | 1.78E-92 | 300 | signal peptide peptidase SppA, 67K type [Escherichia sp. 3_2_53FAA] |
| Q8A6K2 | HMPREF9428_01022 | 37.996 | 87 | 1.86E-92 | 300 | protease 4 [Citrobacter freundii 4_7_47CFAA] |
| Q8A6K2 | HMPREF9551_01037 | 36.528 | 90 | 1.91E-92 | 300 | signal peptide peptidase SppA, 67K type [Escherichia coli MS 196-1] |
| Q8A6K2 | CSAG_01034 | 37.341 | 90 | 2.50E-92 | 300 | protease 4 [Citrobacter sp. 30_2] |
| Q8A6K2 | ECSE_1937 | 36.528 | 90 | 2.78E-92 | 300 | protease IV [Escherichia coli SE11] |
| Q8A6K2 | ESCG_03780 | 36.528 | 90 | 2.80E-92 | 300 | protease 4 [Escherichia sp. 1_1_43] |
| Q8A6K2 | HMPREF9534_04964 | 36.528 | 90 | 2.96E-92 | 300 | signal peptide peptidase SppA, 67K type [Escherichia coli MS 69-1] |
| Q8A6K2 | CIT292_07480 | 37.341 | 90 | 3.18E-92 | 300 | signal peptide peptidase SppA, 67K type [Citrobacter youngae ATCC 29220] |
| Q8A6K2 | ECSF_1626 | 36.528 | 90 | 3.43E-92 | 300 | protease IV [Escherichia coli SE15] |
| Q8A6K2 | HMPREF9347_02540 | 36.528 | 90 | 4.48E-92 | 301 | signal peptide peptidase SppA, 67K type [Escherichia coli MS 124-1] |
| Q8A6K2 | HMPREF9350_00701 | 36.528 | 90 | 4.48E-92 | 301 | signal peptide peptidase SppA, 67K type [Escherichia coli MS 85-1] |
| Q8A6K2 | HMPREF9536_02699 | 36.528 | 90 | 4.48E-92 | 301 | signal peptide peptidase SppA, 67K type [Escherichia coli MS 84-1] |
| Q8A6K2 | HMPREF9541_04686 | 36.528 | 90 | 4.72E-92 | 301 | signal peptide peptidase SppA, 67K type [Escherichia coli MS 116-1] |
| Q8A6K2 | HMPREF9547_04363 | 36.528 | 90 | 4.72E-92 | 301 | signal peptide peptidase SppA, 67K type [Escherichia coli MS 175-1] |
| Q8A6K2 | HMPREF9543_04464 | 36.528 | 90 | 4.72E-92 | 301 | signal peptide peptidase SppA, 67K type [Escherichia coli MS 146-1] |
| Q8A6K2 | HMPREF9542_04947 | 36.528 | 90 | 5.71E-92 | 300 | signal peptide peptidase SppA, 67K type [Escherichia coli MS 117-3] |
| Q8A6K2 | HMPREF9346_03034 | 36.528 | 90 | 5.71E-92 | 300 | signal peptide peptidase SppA, 67K type [Escherichia coli MS 119-7] |
| Q8A6K2 | HMPREF9348_00699 | 36.528 | 90 | 6.15E-92 | 300 | signal peptide peptidase SppA, 67K type [Escherichia coli MS 145-7] |
| Q8A6K2 | HMPREF9548_02937 | 36.528 | 90 | 6.15E-92 | 300 | signal peptide peptidase SppA, 67K type [Escherichia coli MS 182-1] |
| Q8A6K2 | HMPREF9535_02368 | 36.528 | 90 | 6.15E-92 | 300 | signal peptide peptidase SppA, 67K type [Escherichia coli MS 78-1] |
| Q8A6K2 | HMPREF9349_00484 | 36.347 | 90 | 1.06E-91 | 300 | signal peptide peptidase SppA [Escherichia coli MS 79-10] |
| Q8A6K2 | HMPREF9345_03487 | 36.347 | 90 | 1.06E-91 | 300 | signal peptide peptidase SppA, 67K type [Escherichia coli MS 107-1] |
| Q8A6K2 | HMPREF9550_03013 | 36.528 | 90 | 1.06E-91 | 300 | signal peptide peptidase SppA, 67K type [Escherichia coli MS 187-1] |
| Q8A6K2 | HMPREF9540_01493 | 36.528 | 90 | 1.18E-91 | 300 | signal peptide peptidase SppA, 67K type [Escherichia coli MS 115-1] |
| Q8A6K2 | SSJG_00825 | 36.347 | 90 | 1.40E-91 | 298 | signal peptide peptidase SppA, 67K type [Escherichia coli D9] |
| Q8A6K2 | EDWATA_01660 | 37.376 | 83 | 1.90E-90 | 295 | signal peptide peptidase SppA, 67K type [Edwardsiella tarda ATCC 23685] |
| Q8A6K2 | HMPREF0880_03126 | 37.549 | 84 | 2.33E-90 | 295 | signal peptide peptidase SppA [Yokenella regensburgei ATCC 43003] |
| Q8A6K2 | HMPREF0864_04762 | 36.312 | 86 | 2.37E-87 | 287 | signal peptide peptidase SppA, 67K type [Enterobacteriaceae bacterium 9_2_54FAA] |
| Q8A6K2 | HMPREF0454_02659 | 36.053 | 86 | 4.70E-87 | 286 | signal peptide peptidase SppA [Hafnia alvei ATCC 51873] |
| Q8A6K2 | HMPREF0201_01655 | 37.6 | 82 | 2.25E-85 | 282 | signal peptide peptidase SppA [Cedecea davisae DSM 4568] |
| Q8A6K2 | PROVALCAL_01811 | 34.981 | 85 | 2.98E-83 | 276 | signal peptide peptidase SppA, 67K type [Providencia alcalifaciens DSM 30120] |
| Q8A6K2 | PROVRETT_05588 | 36.154 | 85 | 3.39E-83 | 276 | signal peptide peptidase SppA, 67K type [Providencia rettgeri DSM 1131] |
| Q8A6K2 | PROSTU_02769 | 32.671 | 88 | 3.04E-81 | 271 | signal peptide peptidase SppA, 67K type [Providencia stuartii ATCC 25827] |
| Q8A6K2 | PROVRUST_04795 | 36.842 | 84 | 3.88E-81 | 271 | signal peptide peptidase SppA, 67K type [Providencia rustigianii DSM 4541] |
| Q8A6K2 | HMPREF1311_01475 | 34.275 | 87 | 1.01E-77 | 261 | signal peptide peptidase SppA, 67K type [Proteus mirabilis WGLW6] |

| | | | | | | |
|---|---|---|---|---|---|---|
| Q8A6K2 | HMPREF9440_02189 | 32.177 | 85 | 1.33E-72 | 246 | signal peptide peptidase SppA [Sutterella parvirubra YIT 11816] |
| Q8A6K2 | HMPREF9444_01298 | 32.806 | 81 | 1.23E-66 | 233 | signal peptide peptidase SppA, 67K type [Succinatimonas hippei YIT 12066] |
| Q8A6K2 | FMAG_00843 | 28.159 | 91 | 2.76E-66 | 230 | signal peptide peptidase SppA, 67K type [Fusobacterium mortiferum ATCC 9817] |
| Q8A6K2 | HMPREF9465_00588 | 32.669 | 83 | 1.39E-65 | 229 | signal peptide peptidase SppA, 67K type [Sutterella wadsworthensis 2_1_59BFAA] |
| Q8A6K2 | HMPREF1476_02235 | 30.755 | 90 | 1.31E-62 | 221 | signal peptide peptidase SppA, 67K type [Sutterella wadsworthensis HGA0223] |
| Q8A6K2 | HMPREF9464_00344 | 30.755 | 90 | 6.16E-62 | 219 | hypothetical protein [Sutterella wadsworthensis 3_1_45B] |
| Q8A6K2 | FVAG_01920 | 27.82 | 88 | 1.36E-58 | 209 | signal peptide peptidase SppA, 36K type [Fusobacterium varium ATCC 27725] |
| Q8A6K2 | FSBG_01203 | 27.898 | 84 | 4.21E-57 | 205 | signal peptide peptidase SppA, 36K type [Fusobacterium gonidiaformans 3-1-5R] |
| Q8A6K2 | FUAG_00685 | 27.032 | 87 | 2.33E-56 | 204 | signal peptide peptidase SppA, 67K type [Fusobacterium ulcerans ATCC 49185] |
| Q8A6K2 | PSAG_01252 | 30.387 | 87 | 4.85E-56 | 202 | signal peptide peptidase SppA, 36K type [Fusobacterium nucleatum subsp. animalis D11] |
| Q8A6K2 | HMPREF0406_00328 | 30.387 | 87 | 5.65E-56 | 202 | signal peptide peptidase SppA, 67K type [Fusobacterium nucleatum subsp. animalis 3_1_33] |
| Q8A6K2 | HMPREF0401_00940 | 30.387 | 87 | 7.48E-56 | 202 | protease IV [Fusobacterium nucleatum subsp. animalis 11_3_2] |
| Q8A6K2 | HMPREF0404_01303 | 30.387 | 87 | 7.90E-56 | 202 | signal peptide peptidase SppA, 67K type [Fusobacterium nucleatum subsp. animalis 21_1A] |
| Q8A6K2 | FSDG_00193 | 30.387 | 87 | 7.90E-56 | 202 | signal peptide peptidase SppA, 67K type [Fusobacterium nucleatum subsp. animalis 7_1] |
| Q8A6K2 | HMPREF0402_00503 | 27.032 | 87 | 9.18E-56 | 202 | signal peptide peptidase SppA, 67K type [Fusobacterium ulcerans 12-1B] |
| Q8A6K2 | FGAG_00701 | 27.701 | 84 | 1.09E-55 | 201 | signal peptide peptidase SppA, 36K type [Fusobacterium gonidiaformans ATCC 25563] |
| Q8A6K2 | FSCG_00839 | 31.471 | 88 | 8.95E-55 | 199 | signal peptide peptidase SppA, 36K type [Fusobacterium nucleatum subsp. vincentii 4_1_13] |
| Q8A6K2 | HMPREF0404_00901 | 28.807 | 82 | 9.18E-53 | 193 | signal peptide peptidase SppA, 67K type [Fusobacterium nucleatum subsp. animalis 21_1A] |
| Q8A6K2 | HMPREF0406_00596 | 30.415 | 73 | 5.40E-52 | 191 | signal peptide peptidase SppA, 67K type [Fusobacterium nucleatum subsp. animalis 3_1_33] |
| Q8A6K2 | FSDG_00633 | 28.395 | 82 | 1.04E-51 | 190 | signal peptide peptidase SppA, 67K type [Fusobacterium nucleatum subsp. animalis 7_1] |
| Q8A6K2 | FSCG_00389 | 30.485 | 73 | 1.69E-51 | 189 | signal peptide peptidase SppA, 36K type [Fusobacterium nucleatum subsp. vincentii 4_1_13] |
| Q8A6K2 | HMPREF0400_00082 | 29.253 | 81 | 6.05E-51 | 188 | signal peptide peptidase SppA, 36K type [Fusobacterium periodonticum 1_1_41FAA] |
| Q8A6K2 | FSEG_00901 | 26.892 | 83 | 1.73E-49 | 184 | signal peptide peptidase SppA, 36K type [Fusobacterium necrophorum D12] |
| Q8A6K2 | HMPREF0401_00466 | 29.33 | 73 | 1.77E-49 | 184 | protease IV [Fusobacterium nucleatum subsp. animalis 11_3_2] |
| Q8A6K2 | HMPREF9466_00192 | 27.191 | 73 | 4.19E-48 | 179 | signal peptide peptidase SppA, 67K type [Fusobacterium necrophorum subsp. funduliforme 1_1_36S] |
| Q8A6K2 | FSAG_001173 | 31.507 | 83 | 5.07E-48 | 181 | signal peptide peptidase SppA, 67K type [Fusobacterium periodonticum 2_1_31] |
| Q8A6K2 | HMPREF1015_00545 | 39.916 | 40 | 1.37E-46 | 171 | signal peptide peptidase SppA, 36K type [Bacillus smithii 7_3_47FAA] |
| Q8A6K2 | HMPREF0400_01423 | 30.624 | 87 | 3.09E-46 | 176 | signal peptide peptidase SppA, 36K type [Fusobacterium periodonticum 1_1_41FAA] |
| Q8A6K2 | HMPREF0556_12201 | 40.26 | 38 | 1.90E-44 | 165 | signal peptide peptidase SppA, 36K type [Listeria grayi DSM 20601] |
| Q8A6K2 | HMPREF0986_02546 | 46.734 | 33 | 3.28E-44 | 162 | protease 4 [Escherichia coli 4_1_47FAA] |
| Q8A6K2 | HMPREF0557_02255 | 36.782 | 43 | 4.90E-43 | 161 | signal peptide peptidase SppA [Listeria innocua ATCC 33091] |
| Q8A6K2 | PROPEN_00089 | 34.891 | 53 | 9.81E-43 | 162 | signal peptide peptidase SppA, 67K type [Proteus penneri ATCC 35198] |
| Q8A6K2 | HMPREF1210_00129 | 37.594 | 43 | 6.49E-42 | 158 | signal peptide peptidase SppA, 36K type [Paenisporosarcina sp. HGH0030] |
| Q8A6K2 | HMPREF9453_00285 | 36.232 | 35 | 4.33E-40 | 152 | signal peptide peptidase SppA, 36K type [Dialister succinatiphilus YIT 11850] |
| Q8A6K2 | PSAG_00016 | 34.363 | 44 | 4.45E-40 | 152 | signal peptide peptidase SppA, 36K type [Fusobacterium nucleatum subsp. animalis D11] |
| Q8A6K2 | FSAG_000806 | 34.363 | 44 | 1.05E-39 | 150 | signal peptide peptidase SppA, 36K type [Fusobacterium periodonticum 2_1_31] |
| Q8A6K2 | HMPREF0986_02545 | 34.432 | 44 | 1.31E-39 | 152 | protease 4 [Escherichia coli 4_1_47FAA] |
| Q8A6K2 | HMPREF9443_01679 | 33.878 | 40 | 5.59E-39 | 149 | signal peptide peptidase SppA, 36K type [Phascolarctobacterium succinatutens YIT 12067] |
| Q8A6K2 | CUO_0563 | 38 | 42 | 1.18E-37 | 146 | signal peptide peptidase SppA, 36K type [Enterococcus faecium PC4.1] |
| Q8A6K2 | HMPREF0352_1624 | 38 | 42 | 1.18E-37 | 146 | signal peptide peptidase SppA, 36K type [Enterococcus faecium TX1330] |
| Q8A6K2 | HMPREF0326_00798 | 35.242 | 38 | 1.73E-35 | 141 | signal peptide peptidase SppA, 36K type [Desulfovibrio sp. 3_1_syn3] |
| Q8A6K2 | HMPREF1022_02534 | 35.242 | 38 | 1.87E-35 | 140 | hypothetical protein [Desulfovibrio sp. 6_1_46AFAA] |
| Q8A6K2 | DESPIG_01689 | 39.444 | 30 | 5.41E-35 | 139 | signal peptide peptidase SppA, 36K type [Desulfovibrio piger ATCC 29098] |
| Q8A6K2 | HMPREF9478_01319 | 37.104 | 37 | 3.53E-33 | 134 | hypothetical protein [Enterococcus saccharolyticus 30_1] |
| Q8A6K2 | HMPREF1208_01081 | 32.243 | 36 | 1.80E-32 | 132 | signal peptide peptidase SppA, 36K type [Staphylococcus sp. HGB0015] |
| Q8A6K2 | HMPREF9477_00951 | 34.862 | 36 | 3.11E-31 | 128 | signal peptide peptidase SppA, 36K type [Lachnospiraceae bacterium 2_1_46FAA] |
| Q8A343 | BSIG_1537 | 99.786 | 100 | 0 | 969 | hypothetical protein [Bacteroides sp. 1_1_6] |
| Q8A343 | HMPREF9007_02503 | 93.59 | 100 | 0 | 907 | lipoprotein [Bacteroides sp. 1_1_14] |
| Q8A343 | HMPREF0102_00749 | 79.915 | 100 | 0 | 802 | hydrolase, alpha/beta domain protein [Bacteroides sp. 2_1_22] |
| Q8A343 | CW3_3541 | 79.915 | 100 | 0 | 802 | hydrolase, alpha/beta domain protein [Bacteroides xylanisolvens SD CC 1b] |
| Q8A343 | CW1_0407 | 79.915 | 100 | 0 | 802 | hydrolase, alpha/beta domain protein [Bacteroides xylanisolvens SD CC 2a] |
| Q8A343 | BSAG_00501 | 80.525 | 98 | 0 | 790 | hydrolase, alpha/beta domain protein [Bacteroides sp. D1] |
| Q8A343 | CUY_1592 | 80.574 | 97 | 0 | 785 | hydrolase, alpha/beta domain protein [Bacteroides ovatus SD CMC 3f] |
| Q8A343 | HMPREF1017_00377 | 80.574 | 97 | 0 | 785 | hypothetical protein [Bacteroides ovatus 3_8_47FAA] |
| Q8A343 | HMPREF0106_03002 | 80.574 | 97 | 0 | 785 | lipoprotein [Bacteroides sp. D22] |
| Q8A343 | BACCAC_01314 | 80.795 | 97 | 0 | 785 | hydrolase, alpha/beta domain protein [Bacteroides caccae ATCC 43185] |
| Q8A343 | HMPREF0127_01382 | 80.574 | 97 | 0 | 784 | hypothetical protein [Bacteroides sp. 1_1_30] |
| Q8A343 | BXY_22330 | 80.574 | 97 | 0 | 783 | Dienelactone hydrolase and related enzymes [Bacteroides xylanisolvens XB1A] |
| Q8A343 | BACOVA_02813 | 80.574 | 97 | 0 | 783 | hydrolase, alpha/beta domain protein [Bacteroides ovatus ATCC 8483] |
| Q8A343 | BSCG_01598 | 80.353 | 97 | 0 | 781 | hydrolase, alpha/beta domain protein [Bacteroides sp. 2_2_4] |

| | | | | | | |
|---|---|---|---|---|---|---|
| Q8A343 | HMPREF9010_00493 | 79.303 | 98 | 0 | 778 | putative lipoprotein [Bacteroides sp. 3_1_23] |
| Q8A343 | BSGG_2510 | 78.065 | 98 | 0 | 773 | hypothetical protein [Bacteroides sp. D2] |
| Q8A343 | BSCG_01593 | 79.773 | 94 | 0 | 764 | hydrolase, alpha/beta domain protein [Bacteroides sp. 2_2_4] |
| Q8A343 | HMPREF9447_04915 | 68.722 | 97 | 0 | 662 | hypothetical protein [Bacteroides oleiciplenus YIT 12058] |
| Q8A343 | BACINT_03477 | 64.967 | 96 | 0 | 623 | hypothetical protein [Bacteroides intestinalis DSM 17393] |
| Q8A343 | HMPREF9420_1395 | 42.273 | 93 | 1.39E-110 | 336 | hypothetical protein [Prevotella salivae DSM 15606] |
| Q8A343 | HMPREF9441_02648 | 43.524 | 94 | 2.86E-107 | 330 | hydrolase, alpha/beta domain protein [Paraprevotella clara YIT 11840] |
| Q8A343 | HMPREF0619_00319 | 40.129 | 98 | 3.47E-105 | 324 | hydrolase, alpha/beta domain protein [Parabacteroides sp. D13] |
| Q8A343 | HMPREF9442_03095 | 42.275 | 94 | 7.71E-105 | 324 | hydrolase, alpha/beta domain protein [Paraprevotella xylaniphila YIT 11841] |
| Q8A343 | BACCOPRO_03807 | 39.479 | 98 | 1.46E-102 | 316 | hypothetical protein [Bacteroides coprophilus DSM 18228 = JCM 13818] |
| Q8A343 | PRABACTJOHN_03409 | 40.43 | 97 | 1.33E-100 | 312 | hydrolase, alpha/beta domain protein [Parabacteroides johnsonii DSM 18315] |
| Q8A343 | HMPREF0673_02827 | 43.077 | 95 | 5.84E-100 | 310 | hydrolase, alpha/beta domain protein [Prevotella stercorea DSM 18206] |
| Q8A343 | HMPREF1475_01274 | 40.389 | 86 | 1.47E-88 | 279 | hypothetical protein [Prevotella oralis HGA0225] |
| Q8A343 | HMPREF0101_02774 | 31.977 | 70 | 6.56E-36 | 142 | hypothetical protein [Bacteroides sp. 2_1_16] |
| Q8A343 | HMPREF1018_00895 | 31.977 | 70 | 6.69E-36 | 142 | hypothetical protein [Bacteroides sp. 2_1_56FAA] |
| Q8A343 | BSHG_0771 | 31.977 | 70 | 7.62E-36 | 142 | hypothetical protein [Bacteroides sp. 3_2_5] |
| Q8A2L9 | HMPREF9007_03826 | 99.608 | 100 | 0 | 529 | conserved hypothetical protein [Bacteroides sp. 1_1_14] |
| Q8A2L9 | BSIG_1385 | 99.216 | 100 | 0 | 528 | hypothetical protein [Bacteroides sp. 1_1_6] |
| Q8A2L9 | HMPREF0127_01615 | 82.745 | 100 | 1.42E-164 | 459 | hypothetical protein [Bacteroides sp. 1_1_30] |
| Q8A2L9 | HMPREF0106_04277 | 82.745 | 100 | 1.42E-164 | 459 | conserved hypothetical protein [Bacteroides sp. D22] |
| Q8A2L9 | BXY_25350 | 82.353 | 100 | 9.49E-164 | 457 | Predicted peptidase [Bacteroides xylanisolvens XB1A] |
| Q8A2L9 | HMPREF0102_00571 | 82.745 | 100 | 2.23E-163 | 456 | phospholipase/carboxylesterase [Bacteroides sp. 2_1_22] |
| Q8A2L9 | BSAG_00321 | 82.745 | 100 | 2.23E-163 | 456 | phospholipase/carboxylesterase [Bacteroides sp. D1] |
| Q8A2L9 | CW3_2066 | 82.745 | 100 | 2.23E-163 | 456 | phospholipase/carboxylesterase [Bacteroides xylanisolvens SD CC 1b] |
| Q8A2L9 | CW1_0768 | 82.745 | 100 | 2.23E-163 | 456 | phospholipase/carboxylesterase [Bacteroides xylanisolvens SD CC 2a] |
| Q8A2L9 | BSCG_03062 | 83.137 | 100 | 1.09E-154 | 434 | phospholipase/carboxylesterase [Bacteroides sp. 2_2_4] |
| Q8A2L9 | CUY_4407 | 83.137 | 100 | 1.27E-154 | 434 | phospholipase/carboxylesterase [Bacteroides ovatus SD CMC 3f] |
| Q8A2L9 | HMPREF9010_00749 | 83.137 | 100 | 1.27E-154 | 434 | phospholipase/carboxylesterase superfamily [Bacteroides sp. 3_1_23] |
| Q8A2L9 | BSGG_2335 | 83.137 | 100 | 1.27E-154 | 434 | hypothetical protein [Bacteroides sp. D2] |
| Q8A2L9 | HMPREF1017_00562 | 83.137 | 100 | 1.27E-154 | 434 | hypothetical protein [Bacteroides ovatus 3_8_47FAA] |
| Q8A2L9 | BACOVA_00896 | 83.137 | 100 | 1.27E-154 | 434 | phospholipase/carboxylesterase [Bacteroides ovatus ATCC 8483] |
| Q8A2L9 | HMPREF9447_05110 | 76.096 | 98 | 1.36E-147 | 416 | hypothetical protein [Bacteroides oleiciplenus YIT 12058] |
| Q8A2L9 | HMPREF1214_00184 | 72.835 | 99 | 3.49E-144 | 407 | hypothetical protein [Bacteroides sp. HPS0048] |
| Q8A2L9 | BACCOP_00633 | 76.151 | 94 | 5.15E-140 | 397 | phospholipase/carboxylesterase [Bacteroides coprocola DSM 17136] |
| Q8A2L9 | HMPREF1532_02134 | 71.654 | 99 | 2.36E-139 | 395 | hypothetical protein [Bacteroides salyersiae WAL 10018 = DSM 18765 = JCM 12988] |
| Q8A2L9 | BSHG_3515 | 70.312 | 100 | 2.49E-138 | 392 | hypothetical protein [Bacteroides sp. 3_2_5] |
| Q8A2L9 | BFAG_04177 | 69.38 | 100 | 3.49E-137 | 390 | phospholipase/carboxylesterase [Bacteroides fragilis 3_1_12] |
| Q8A2L9 | HMPREF1018_04045 | 69.922 | 100 | 5.07E-137 | 389 | hypothetical protein [Bacteroides sp. 2_1_56FAA] |
| Q8A2L9 | HMPREF0101_04075 | 69.531 | 100 | 8.77E-137 | 389 | phospholipase/carboxylesterase [Bacteroides sp. 2_1_16] |
| Q8A2L9 | BACINT_01644 | 74.262 | 93 | 8.91E-137 | 389 | phospholipase/carboxylesterase [Bacteroides intestinalis DSM 17393] |
| Q8A2L9 | HMPREF0969_01919 | 70.98 | 100 | 4.62E-136 | 387 | phospholipase/carboxylesterase [Bacteroides sp. D20] |
| Q8A2L9 | BACUNI_03954 | 70.155 | 100 | 1.55E-133 | 380 | phospholipase/carboxylesterase [Bacteroides uniformis ATCC 8492] |
| Q8A2L9 | HMPREF1007_00666 | 70.155 | 100 | 1.55E-133 | 380 | hypothetical protein [Bacteroides sp. 4_1_36] |
| Q8A2L9 | BACCELL_02512 | 77.073 | 80 | 1.54E-120 | 345 | phospholipase/carboxylesterase [Bacteroides cellulosilyticus DSM 14838] |
| Q8A2L9 | HMPREF1214_02993 | 57.708 | 98 | 9.12E-105 | 308 | hypothetical protein [Bacteroides sp. HPS0048] |
| Q8A2L9 | HMPREF9448_00395 | 50 | 91 | 2.64E-82 | 258 | por secretion system C-terminal sorting domain-containing protein [Barnesiella intestinihominis YIT 11860] |
| Q8A2L9 | HMPREF0103_2108 | 46.457 | 97 | 9.11E-78 | 239 | phospholipase/carboxylesterase [Bacteroides sp. 2_1_33B] |
| Q8A2L9 | HMPREF0104_00201 | 48.729 | 90 | 8.71E-77 | 237 | phospholipase/carboxylesterase superfamily [Bacteroides sp. 3_1_19] |
| Q8A2L9 | HMPREF0619_00485 | 48.729 | 90 | 1.07E-76 | 237 | phospholipase/carboxylesterase [Parabacteroides sp. D13] |
| Q8A2L9 | HMPREF9008_03720 | 48.729 | 90 | 1.33E-76 | 236 | phospholipase/carboxylesterase [Parabacteroides sp. 20_3] |
| Q8A2L9 | HMPREF9456_01149 | 44.186 | 100 | 1.10E-67 | 213 | hypothetical protein [Dysgonomonas mossii DSM 22836] |
| Q8A2L9 | HMPREF9455_02119 | 47.577 | 88 | 2.63E-65 | 207 | hypothetical protein [Dysgonomonas gadei ATCC BAA-286] |
| Q8A2L9 | BACCELL_01985 | 49.405 | 64 | 2.79E-55 | 179 | hypothetical protein [Bacteroides cellulosilyticus DSM 14838] |
| Q8A1Y1 | HMPREF9007_03406 | 100 | 100 | 0 | 1596 | alpha-1,2-mannosidase family protein [Bacteroides sp. 1_1_14] |
| Q8A1Y1 | BSIG_3494 | 99.869 | 100 | 0 | 1593 | putative alpha-1,2-mannosidase [Bacteroides sp. 1_1_6] |
| Q8A1Y1 | BACOVA_03346 | 94.364 | 100 | 0 | 1527 | putative alpha-1,2-mannosidase [Bacteroides ovatus ATCC 8483] |
| Q8A1Y1 | BSGG_2052 | 94.102 | 100 | 0 | 1526 | putative alpha-1,2-mannosidase [Bacteroides sp. D2] |
| Q8A1Y1 | HMPREF0102_00338 | 94.233 | 100 | 0 | 1526 | putative alpha-1,2-mannosidase [Bacteroides sp. 2_1_22] |
| Q8A1Y1 | BSAG_00082 | 94.233 | 100 | 0 | 1526 | putative alpha-1,2-mannosidase [Bacteroides sp. D1] |
| Q8A1Y1 | CW3_3351 | 94.233 | 100 | 0 | 1526 | putative alpha-1,2-mannosidase [Bacteroides xylanisolvens SD CC 1b] |

| | | | | | | |
|---|---|---|---|---|---|---|
| Q8A1Y1 | CW1_3018 | 94.233 | 100 | 0 | 1526 | putative alpha-1,2-mannosidase [Bacteroides xylanisolvens SD CC 2a] |
| Q8A1Y1 | CUY_0237 | 94.233 | 100 | 0 | 1525 | putative alpha-1,2-mannosidase [Bacteroides ovatus SD CMC 3f] |
| Q8A1Y1 | BSCG_02805 | 94.233 | 100 | 0 | 1525 | putative alpha-1,2-mannosidase [Bacteroides sp. 2_2_4] |
| Q8A1Y1 | HMPREF0106_01040 | 94.233 | 100 | 0 | 1525 | alpha-1,2-mannosidase family protein [Bacteroides sp. D22] |
| Q8A1Y1 | HMPREF0127_02546 | 93.971 | 100 | 0 | 1524 | hypothetical protein [Bacteroides sp. 1_1_30] |
| Q8A1Y1 | HMPREF1017_02873 | 93.84 | 100 | 0 | 1523 | hypothetical protein [Bacteroides ovatus 3_8_47FAA] |
| Q8A1Y1 | HMPREF9010_03872 | 93.84 | 100 | 0 | 1522 | alpha-1,2-mannosidase family protein [Bacteroides sp. 3_1_23] |
| Q8A1Y1 | BXY_28180 | 93.84 | 100 | 0 | 1519 | alpha-1,2-mannosidase, putative [Bacteroides xylanisolvens XB1A] |
| Q8A1Y1 | BSHG_1278 | 86.898 | 98 | 0 | 1395 | putative alpha-1,2-mannosidase [Bacteroides sp. 3_2_5] |
| Q8A1Y1 | HMPREF1018_00339 | 86.898 | 98 | 0 | 1395 | hypothetical protein [Bacteroides sp. 2_1_56FAA] |
| Q8A1Y1 | HMPREF0101_02300 | 86.898 | 98 | 0 | 1395 | putative alpha-1,2-mannosidase [Bacteroides sp. 2_1_16] |
| Q8A1Y1 | BFAG_02665 | 87.332 | 97 | 0 | 1394 | putative alpha-1,2-mannosidase [Bacteroides fragilis 3_1_12] |
| Q8A1Y1 | HMPREF9455_03280 | 80.105 | 99 | 0 | 1317 | hypothetical protein [Dysgonomonas gadei ATCC BAA-286] |
| Q8A1Y1 | HMPREF9456_00301 | 79.578 | 99 | 0 | 1306 | hypothetical protein [Dysgonomonas mossii DSM 22836] |
| Q8A1Y1 | HMPREF9455_03989 | 79.974 | 99 | 0 | 1305 | hypothetical protein [Dysgonomonas gadei ATCC BAA-286] |
| Q8A1Y1 | BACSTE_03757 | 78.516 | 100 | 0 | 1305 | putative alpha-1,2-mannosidase [Bacteroides stercoris ATCC 43183] |
| Q8A1Y1 | HMPREF0969_02272 | 78.125 | 100 | 0 | 1301 | putative alpha-1,2-mannosidase [Bacteroides sp. D20] |
| Q8A1Y1 | BACUNI_03096 | 78.125 | 100 | 0 | 1301 | putative alpha-1,2-mannosidase [Bacteroides uniformis ATCC 8492] |
| Q8A1Y1 | PARMER_03861 | 79.134 | 99 | 0 | 1299 | putative alpha-1,2-mannosidase [Parabacteroides merdae ATCC 43184] |
| Q8A1Y1 | HMPREF1007_02049 | 80.379 | 97 | 0 | 1299 | glycosyl hydrolase family 92 [Bacteroides sp. 4_1_36] |
| Q8A1Y1 | PRABACTJOHN_00066 | 78.74 | 99 | 0 | 1293 | putative alpha-1,2-mannosidase [Parabacteroides johnsonii DSM 18315] |
| Q8A1Y1 | HMPREF1475_00939 | 77.177 | 99 | 0 | 1256 | hypothetical protein [Prevotella oralis HGA0225] |
| Q8A1Y1 | BACOVA_04809 | 74.67 | 99 | 0 | 1234 | putative alpha-1,2-mannosidase [Bacteroides ovatus ATCC 8483] |
| Q8A1Y1 | HMPREF0106_00730 | 75.198 | 99 | 0 | 1233 | alpha-1,2-mannosidase family protein [Bacteroides sp. D22] |
| Q8A1Y1 | BXY_31140 | 75.066 | 99 | 0 | 1232 | alpha-1,2-mannosidase, putative [Bacteroides xylanisolvens XB1A] |
| Q8A1Y1 | HMPREF0127_01951 | 74.67 | 99 | 0 | 1231 | hypothetical protein [Bacteroides sp. 1_1_30] |
| Q8A1Y1 | HMPREF9007_01571 | 74.274 | 99 | 0 | 1231 | alpha-1,2-mannosidase family protein [Bacteroides sp. 1_1_14] |
| Q8A1Y1 | HMPREF9010_02577 | 74.67 | 99 | 0 | 1231 | alpha-1,2-mannosidase family protein [Bacteroides sp. 3_1_23] |
| Q8A1Y1 | HMPREF1017_03049 | 74.67 | 99 | 0 | 1230 | hypothetical protein [Bacteroides ovatus 3_8_47FAA] |
| Q8A1Y1 | CUY_1101 | 74.538 | 99 | 0 | 1229 | putative alpha-1,2-mannosidase [Bacteroides ovatus SD CMC 3f] |
| Q8A1Y1 | BSCG_04343 | 74.406 | 99 | 0 | 1228 | putative alpha-1,2-mannosidase [Bacteroides sp. 2_2_4] |
| Q8A1Y1 | BSGG_1570 | 74.406 | 99 | 0 | 1228 | putative alpha-1,2-mannosidase [Bacteroides sp. D2] |
| Q8A1Y1 | BSIG_2675 | 74.011 | 99 | 0 | 1226 | putative alpha-1,2-mannosidase [Bacteroides sp. 1_1_6] |
| Q8A1Y1 | HMPREF0102_03255 | 74.538 | 99 | 0 | 1224 | putative alpha-1,2-mannosidase [Bacteroides sp. 2_1_22] |
| Q8A1Y1 | BSAG_03128 | 74.538 | 99 | 0 | 1224 | putative alpha-1,2-mannosidase [Bacteroides sp. D1] |
| Q8A1Y1 | CW3_3115 | 74.538 | 99 | 0 | 1224 | putative alpha-1,2-mannosidase [Bacteroides xylanisolvens SD CC 1b] |
| Q8A1Y1 | CW1_2676 | 74.538 | 99 | 0 | 1224 | putative alpha-1,2-mannosidase [Bacteroides xylanisolvens SD CC 2a] |
| Q8A1Y1 | BACCAC_01682 | 73.483 | 99 | 0 | 1219 | putative alpha-1,2-mannosidase [Bacteroides caccae ATCC 43185] |
| Q8A1Y1 | BACFIN_05353 | 73.483 | 99 | 0 | 1216 | putative alpha-1,2-mannosidase [Bacteroides finegoldii DSM 17565] |
| Q8A1Y1 | BACUNI_02645 | 72.405 | 99 | 0 | 1213 | putative alpha-1,2-mannosidase [Bacteroides uniformis ATCC 8492] |
| Q8A1Y1 | HMPREF1007_03652 | 72.405 | 99 | 0 | 1213 | glycosyl hydrolase family 92 [Bacteroides sp. 4_1_36] |
| Q8A1Y1 | HMPREF0673_02685 | 73.289 | 99 | 0 | 1212 | putative alpha-1,2-mannosidase [Prevotella stercorea DSM 18206] |
| Q8A1Y1 | HMPREF0969_00775 | 71.879 | 99 | 0 | 1207 | putative alpha-1,2-mannosidase [Bacteroides sp. D20] |
| Q8A1Y1 | BACOVA_01636 | 72.215 | 100 | 0 | 1202 | putative alpha-1,2-mannosidase [Bacteroides ovatus ATCC 8483] |
| Q8A1Y1 | HMPREF1214_01295 | 71.711 | 99 | 0 | 1201 | hypothetical protein [Bacteroides sp. HPS0048] |
| Q8A1Y1 | HMPREF9010_04618 | 72.084 | 100 | 0 | 1201 | alpha-1,2-mannosidase family protein [Bacteroides sp. 3_1_23] |
| Q8A1Y1 | HMPREF1017_04368 | 72.215 | 100 | 0 | 1200 | hypothetical protein [Bacteroides ovatus 3_8_47FAA] |
| Q8A1Y1 | CUY_2712 | 72.084 | 100 | 0 | 1199 | putative alpha-1,2-mannosidase [Bacteroides ovatus SD CMC 3f] |
| Q8A1Y1 | BSCG_00124 | 72.084 | 100 | 0 | 1199 | putative alpha-1,2-mannosidase [Bacteroides sp. 2_2_4] |
| Q8A1Y1 | BSGG_4850 | 72.084 | 100 | 0 | 1198 | putative alpha-1,2-mannosidase [Bacteroides sp. D2] |
| Q8A1Y1 | HMPREF1532_01086 | 72.354 | 99 | 0 | 1197 | hypothetical protein [Bacteroides salyersiae WAL 10018 = DSM 18765 = JCM 12988] |
| Q8A1Y1 | BFAG_00091 | 72.559 | 99 | 0 | 1196 | putative alpha-1,2-mannosidase [Bacteroides fragilis 3_1_12] |
| Q8A1Y1 | BACCELL_03369 | 71.916 | 99 | 0 | 1195 | putative alpha-1,2-mannosidase [Bacteroides cellulosilyticus DSM 14838] |
| Q8A1Y1 | HMPREF9447_01541 | 72.608 | 99 | 0 | 1193 | hypothetical protein [Bacteroides oleiciplenus YIT 12058] |
| Q8A1Y1 | BACSTE_03544 | 71.391 | 99 | 0 | 1193 | putative alpha-1,2-mannosidase [Bacteroides stercoris ATCC 43183] |
| Q8A1Y1 | HMPREF1018_00776 | 72.222 | 99 | 0 | 1192 | hypothetical protein [Bacteroides sp. 2_1_56FAA] |
| Q8A1Y1 | HMPREF0101_02657 | 72.354 | 99 | 0 | 1192 | putative alpha-1,2-mannosidase [Bacteroides sp. 2_1_16] |
| Q8A1Y1 | PRABACTJOHN_01667 | 71.782 | 100 | 0 | 1191 | putative alpha-1,2-mannosidase [Parabacteroides johnsonii DSM 18315] |
| Q8A1Y1 | BACINT_04249 | 73.605 | 96 | 0 | 1189 | putative alpha-1,2-mannosidase [Bacteroides intestinalis DSM 17393] |
| Q8A1Y1 | BSHG_0892 | 72.09 | 99 | 0 | 1189 | putative alpha-1,2-mannosidase [Bacteroides sp. 3_2_5] |

| | | | | | | |
|---|---|---|---|---|---|---|
| Q8A1Y1 | BACDOR_01141 | 71.818 | 99 | 0 | 1184 | putative alpha-1,2-mannosidase [Bacteroides dorei DSM 17855] |
| Q8A1Y1 | BSEG_01825 | 71.688 | 99 | 0 | 1182 | putative alpha-1,2-mannosidase [Bacteroides dorei 5_1_36/D4] |
| Q8A1Y1 | BSBG_01328 | 71.688 | 99 | 0 | 1182 | putative alpha-1,2-mannosidase [Bacteroides sp. 9_1_42FAA] |
| Q8A1Y1 | CUU_0910 | 71.447 | 99 | 0 | 1181 | putative alpha-1,2-mannosidase [Bacteroides vulgatus PC510] |
| Q8A1Y1 | BSFG_01739 | 71.447 | 99 | 0 | 1181 | putative alpha-1,2-mannosidase [Bacteroides sp. 4_3_47FAA] |
| Q8A1Y1 | HMPREF9011_00727 | 71.447 | 99 | 0 | 1181 | glycoside hydrolase family 92 [Bacteroides sp. 3_1_40A] |
| Q8A1Y1 | HMPREF0104_01337 | 73.224 | 96 | 0 | 1181 | alpha-1,2-mannosidase family protein [Bacteroides sp. 3_1_19] |
| Q8A1Y1 | BACCOP_04375 | 71.222 | 99 | 0 | 1181 | putative alpha-1,2-mannosidase [Bacteroides coprocola DSM 17136] |
| Q8A1Y1 | HMPREF0619_01279 | 73.087 | 96 | 0 | 1180 | putative alpha-1,2-mannosidase [Parabacteroides sp. D13] |
| Q8A1Y1 | PARMER_02236 | 70.741 | 100 | 0 | 1180 | putative alpha-1,2-mannosidase [Parabacteroides merdae ATCC 43184] |
| Q8A1Y1 | HMPREF9008_03225 | 73.087 | 96 | 0 | 1180 | putative alpha-1,2-mannosidase [Parabacteroides sp. 20_3] |
| Q8A1Y1 | HMPREF0105_2413 | 71.558 | 99 | 0 | 1179 | putative alpha-1,2-mannosidase [Bacteroides sp. 3_1_33FAA] |
| Q8A1Y1 | HMPREF0127_01911 | 70.773 | 99 | 0 | 1179 | hypothetical protein [Bacteroides sp. 1_1_30] |
| Q8A1Y1 | HMPREF9455_02175 | 72.919 | 99 | 0 | 1179 | hypothetical protein [Dysgonomonas gadei ATCC BAA-286] |
| Q8A1Y1 | BSGG_1592 | 72.951 | 96 | 0 | 1179 | putative alpha-1,2-mannosidase [Bacteroides sp. D2] |
| Q8A1Y1 | HMPREF0106_00701 | 70.904 | 99 | 0 | 1179 | alpha-1,2-mannosidase family protein [Bacteroides sp. D22] |
| Q8A1Y1 | CUY_1145 | 70.565 | 99 | 0 | 1177 | putative alpha-1,2-mannosidase [Bacteroides ovatus SD CMC 3f] |
| Q8A1Y1 | BSCG_04297 | 70.565 | 99 | 0 | 1177 | putative alpha-1,2-mannosidase [Bacteroides sp. 2_2_4] |
| Q8A1Y1 | BXY_30750 | 70.696 | 99 | 0 | 1177 | alpha-1,2-mannosidase, putative [Bacteroides xylanisolvens XB1A] |
| Q8A1Y1 | HMPREF0102_03218 | 70.565 | 99 | 0 | 1177 | putative alpha-1,2-mannosidase [Bacteroides sp. 2_1_22] |
| Q8A1Y1 | BSAG_03093 | 70.565 | 99 | 0 | 1177 | putative alpha-1,2-mannosidase [Bacteroides sp. D1] |
| Q8A1Y1 | CW3_1776 | 70.565 | 99 | 0 | 1177 | putative alpha-1,2-mannosidase [Bacteroides xylanisolvens SD CC 1b] |
| Q8A1Y1 | HMPREF0103_1289 | 72.814 | 96 | 0 | 1176 | putative alpha-1,2-mannosidase [Bacteroides sp. 2_1_33B] |
| Q8A1Y1 | HMPREF9007_03646 | 73.522 | 98 | 0 | 1176 | alpha-1,2-mannosidase family protein [Bacteroides sp. 1_1_14] |
| Q8A1Y1 | BACPLE_00637 | 71.693 | 99 | 0 | 1176 | putative alpha-1,2-mannosidase [Bacteroides plebeius DSM 17135] |
| Q8A1Y1 | BACOVA_01222 | 72.343 | 96 | 0 | 1176 | putative alpha-1,2-mannosidase [Bacteroides ovatus ATCC 8483] |
| Q8A1Y1 | HMPREF9010_02546 | 70.642 | 99 | 0 | 1176 | alpha-1,2-mannosidase family protein [Bacteroides sp. 3_1_23] |
| Q8A1Y1 | HMPREF1017_03076 | 70.642 | 99 | 0 | 1176 | hypothetical protein [Bacteroides ovatus 3_8_47FAA] |
| Q8A1Y1 | BSIG_3777 | 73.387 | 98 | 0 | 1175 | putative alpha-1,2-mannosidase [Bacteroides sp. 1_1_6] |
| Q8A1Y1 | BACCOP_03489 | 71.108 | 99 | 0 | 1175 | putative alpha-1,2-mannosidase [Bacteroides coprocola DSM 17136] |
| Q8A1Y1 | BXY_48410 | 73.118 | 98 | 0 | 1174 | alpha-1,2-mannosidase, putative [Bacteroides xylanisolvens XB1A] |
| Q8A1Y1 | HMPREF9010_04161 | 73.118 | 98 | 0 | 1174 | alpha-1,2-mannosidase family protein [Bacteroides sp. 3_1_23] |
| Q8A1Y1 | BSGG_1778 | 73.118 | 98 | 0 | 1173 | putative alpha-1,2-mannosidase [Bacteroides sp. D2] |
| Q8A1Y1 | BACOVA_03629 | 73.118 | 98 | 0 | 1173 | putative alpha-1,2-mannosidase [Bacteroides ovatus ATCC 8483] |
| Q8A1Y1 | BSCG_03589 | 72.984 | 98 | 0 | 1172 | putative alpha-1,2-mannosidase [Bacteroides sp. 2_2_4] |
| Q8A1Y1 | HMPREF0102_00038 | 72.984 | 98 | 0 | 1172 | putative alpha-1,2-mannosidase [Bacteroides sp. 2_1_22] |
| Q8A1Y1 | BSAG_02041 | 72.984 | 98 | 0 | 1172 | putative alpha-1,2-mannosidase [Bacteroides sp. D1] |
| Q8A1Y1 | CW3_1133 | 72.984 | 98 | 0 | 1172 | putative alpha-1,2-mannosidase [Bacteroides xylanisolvens SD CC 1b] |
| Q8A1Y1 | CW1_1393 | 72.984 | 98 | 0 | 1172 | putative alpha-1,2-mannosidase [Bacteroides xylanisolvens SD CC 2a] |
| Q8A1Y1 | HMPREF0127_04531 | 73.118 | 98 | 0 | 1172 | hypothetical protein [Bacteroides sp. 1_1_30] |
| Q8A1Y1 | HMPREF9007_01544 | 70.434 | 99 | 0 | 1171 | alpha-1,2-mannosidase family protein [Bacteroides sp. 1_1_14] |
| Q8A1Y1 | BSIG_2707 | 70.434 | 99 | 0 | 1171 | putative alpha-1,2-mannosidase [Bacteroides sp. 1_1_6] |
| Q8A1Y1 | HMPREF1017_02632 | 72.984 | 98 | 0 | 1171 | hypothetical protein [Bacteroides ovatus 3_8_47FAA] |
| Q8A1Y1 | CUY_0573 | 72.984 | 98 | 0 | 1170 | putative alpha-1,2-mannosidase [Bacteroides ovatus SD CMC 3f] |
| Q8A1Y1 | HMPREF9447_05316 | 68.997 | 99 | 0 | 1147 | hypothetical protein [Bacteroides oleiciplenus YIT 12058] |
| Q8A1Y1 | HMPREF1532_00850 | 68.146 | 99 | 0 | 1138 | hypothetical protein [Bacteroides salyersiae WAL 10018 = DSM 18765 = JCM 12988] |
| Q8A1Y1 | BACCOPRO_02261 | 69.083 | 96 | 0 | 1135 | putative alpha-1,2-mannosidase [Bacteroides coprophilus DSM 18228 = JCM 13818] |
| Q8A1Y1 | HMPREF1214_04464 | 67.984 | 99 | 0 | 1134 | hypothetical protein [Bacteroides sp. HPS0048] |
| Q8A1Y1 | HMPREF0104_02111 | 69.353 | 99 | 0 | 1132 | alpha-1,2-mannosidase family protein [Bacteroides sp. 3_1_19] |
| Q8A1Y1 | HMPREF0103_2758 | 69.221 | 99 | 0 | 1131 | putative alpha-1,2-mannosidase [Bacteroides sp. 2_1_33B] |
| Q8A1Y1 | HMPREF1214_01521 | 68.016 | 99 | 0 | 1129 | hypothetical protein [Bacteroides sp. HPS0048] |
| Q8A1Y1 | HMPREF0619_02982 | 68.956 | 99 | 0 | 1129 | putative alpha-1,2-mannosidase [Parabacteroides sp. D13] |
| Q8A1Y1 | HMPREF9008_02500 | 68.692 | 99 | 0 | 1125 | putative alpha-1,2-mannosidase [Parabacteroides sp. 20_3] |
| Q8A1Y1 | HMPREF9448_02300 | 69.054 | 97 | 0 | 1123 | hypothetical protein [Barnesiella intestinihominis YIT 11860] |
| Q8A1Y1 | HMPREF9448_00581 | 69.293 | 96 | 0 | 1120 | hypothetical protein [Barnesiella intestinihominis YIT 11860] |
| Q8A1Y1 | HMPREF1532_04078 | 66.008 | 99 | 0 | 1111 | hypothetical protein [Bacteroides salyersiae WAL 10018 = DSM 18765 = JCM 12988] |
| Q8A1Y1 | HMPREF9455_03359 | 68.594 | 99 | 0 | 1103 | hypothetical protein [Dysgonomonas gadei ATCC BAA-286] |
| Q8A1Y1 | HMPREF1033_02449 | 67.152 | 99 | 0 | 1101 | hypothetical protein [Tannerella sp. 6_1_58FAA_CT1] |
| Q8A1Y1 | HMPREF9449_01284 | 68.169 | 96 | 0 | 1094 | putative alpha-1,2-mannosidase [Odoribacter laneus YIT 12061] |
| Q8A1Y1 | BACCELL_01862 | 64.954 | 99 | 0 | 1080 | putative alpha-1,2-mannosidase [Bacteroides cellulosilyticus DSM 14838] |

| | | | | | | |
|---|---|---|---|---|---|---|
| Q8A1Y1 | HMPREF9420_1354 | 64.875 | 99 | 0 | 1061 | putative alpha-1,2-mannosidase [Prevotella salivae DSM 15606] |
| Q8A1Y1 | AL1_30810 | 65.314 | 99 | 0 | 1059 | alpha-1,2-mannosidase, putative [Alistipes shahii WAL 8301] |
| Q8A1Y1 | HMPREF9420_1435 | 52.917 | 96 | 0 | 840 | putative alpha-1,2-mannosidase [Prevotella salivae DSM 15606] |
| Q8A1Y1 | BSIG_4488 | 50.87 | 98 | 0 | 815 | putative alpha-1,2-mannosidase [Bacteroides sp. 1_1_6] |
| Q8A1Y1 | HMPREF9007_00258 | 50.531 | 99 | 0 | 812 | alpha-1,2-mannosidase family protein [Bacteroides sp. 1_1_14] |
| Q8A1Y1 | BFAG_00064 | 49.801 | 99 | 0 | 805 | putative alpha-1,2-mannosidase [Bacteroides fragilis 3_1_12] |
| Q8A1Y1 | CW1_2595 | 69.346 | 70 | 0 | 804 | putative alpha-1,2-mannosidase [Bacteroides xylanisolvens SD CC 2a] |
| Q8A1Y1 | HMPREF1018_00753 | 49.404 | 99 | 0 | 799 | hypothetical protein [Bacteroides sp. 2_1_56FAA] |
| Q8A1Y1 | BSHG_0923 | 49.404 | 99 | 0 | 798 | putative alpha-1,2-mannosidase [Bacteroides sp. 3_2_5] |
| Q8A1Y1 | HMPREF0101_02632 | 49.139 | 99 | 0 | 795 | putative alpha-1,2-mannosidase [Bacteroides sp. 2_1_16] |
| Q8A1Y1 | HMPREF1033_02712 | 50.41 | 96 | 0 | 783 | hypothetical protein [Tannerella sp. 6_1_58FAA_CT1] |
| Q8A1Y1 | BACCELL_00834 | 49.009 | 99 | 0 | 775 | putative alpha-1,2-mannosidase [Bacteroides cellulosilyticus DSM 14838] |
| Q8A1Y1 | HMPREF9446_00568 | 49.655 | 95 | 0 | 764 | putative alpha-1,2-mannosidase [Bacteroides fluxus YIT 12057] |
| Q8A1Y1 | BXY_31760 | 46.789 | 99 | 0 | 720 | alpha-1,2-mannosidase, putative [Bacteroides xylanisolvens XB1A] |
| Q8A1Y1 | HMPREF9007_01647 | 46.789 | 99 | 0 | 718 | alpha-1,2-mannosidase family protein [Bacteroides sp. 1_1_14] |
| Q8A1Y1 | BSIG_2554 | 46.789 | 99 | 0 | 718 | putative alpha-1,2-mannosidase [Bacteroides sp. 1_1_6] |
| Q8A1Y1 | HMPREF9455_02450 | 46.447 | 99 | 0 | 712 | hypothetical protein [Dysgonomonas gadei ATCC BAA-286] |
| Q8A1Y1 | CUY_2886 | 46.396 | 99 | 0 | 709 | putative alpha-1,2-mannosidase [Bacteroides ovatus SD CMC 3f] |
| Q8A1Y1 | BACOVA_04883 | 46.396 | 99 | 0 | 708 | putative alpha-1,2-mannosidase [Bacteroides ovatus ATCC 8483] |
| Q8A1Y1 | CW3_2373 | 46.396 | 99 | 0 | 708 | putative alpha-1,2-mannosidase [Bacteroides xylanisolvens SD CC 1b] |
| Q8A1Y1 | BSGG_1511 | 45.872 | 99 | 0 | 707 | putative alpha-1,2-mannosidase [Bacteroides sp. D2] |
| Q8A1Y1 | HMPREF1017_02978 | 46.396 | 99 | 0 | 707 | hypothetical protein [Bacteroides ovatus 3_8_47FAA] |
| Q8A1Y1 | HMPREF0106_00798 | 46.396 | 99 | 0 | 707 | alpha-1,2-mannosidase family protein [Bacteroides sp. D22] |
| Q8A1Y1 | CW1_1107 | 46.396 | 99 | 0 | 707 | putative alpha-1,2-mannosidase [Bacteroides xylanisolvens SD CC 2a] |
| Q8A1Y1 | BSCG_05454 | 46.265 | 99 | 0 | 707 | putative alpha-1,2-mannosidase [Bacteroides sp. 2_2_4] |
| Q8A1Y1 | HMPREF0127_02020 | 46.134 | 99 | 0 | 706 | hypothetical protein [Bacteroides sp. 1_1_30] |
| Q8A1Y1 | HMPREF9456_02215 | 46.06 | 96 | 0 | 705 | hypothetical protein [Dysgonomonas mossii DSM 22836] |
| Q8A1Y1 | HMPREF0103_2009 | 45.986 | 96 | 0 | 684 | putative alpha-1,2-mannosidase [Bacteroides sp. 2_1_33B] |
| Q8A1Y1 | HMPREF9412_2222 | 36.07 | 96 | 5.43E-157 | 477 | putative alpha-1,2-mannosidase [Paenibacillus sp. HGF5] |
| Q8A1Y1 | HMPREF0556_10520 | 34.454 | 93 | 2.93E-149 | 457 | putative alpha-1,2-mannosidase [Listeria grayi DSM 20601] |
| Q8A1Y1 | HMPREF1486_03614 | 35.097 | 97 | 9.73E-145 | 457 | hypothetical protein [Streptomyces sp. HPH0547] |
| Q8A1Y1 | HMPREF9447_02745 | 34.76 | 96 | 2.38E-144 | 446 | hypothetical protein [Bacteroides oleiciplenus YIT 12058] |
| Q8A1Y1 | HMPREF1033_02160 | 34.478 | 97 | 1.01E-143 | 444 | hypothetical protein [Tannerella sp. 6_1_58FAA_CT1] |
| Q8A1Y1 | CUO_1869 | 33.607 | 95 | 1.67E-143 | 442 | putative alpha-1,2-mannosidase [Enterococcus faecium PC4.1] |
| Q8A1Y1 | HMPREF0352_0231 | 33.607 | 95 | 1.67E-143 | 442 | putative alpha-1,2-mannosidase [Enterococcus faecium TX1330] |
| Q8A1Y1 | BACPLE_03760 | 32.875 | 99 | 5.13E-143 | 442 | putative alpha-1,2-mannosidase [Bacteroides plebeius DSM 17135] |
| Q8A1Y1 | BACCELL_02347 | 35.397 | 93 | 6.70E-142 | 438 | putative alpha-1,2-mannosidase [Bacteroides cellulosilyticus DSM 14838] |
| Q8A1Y1 | HMPREF0348_3255 | 32.834 | 95 | 2.50E-140 | 433 | putative alpha-1,2-mannosidase [Enterococcus faecalis TX0104] |
| Q8A1Y1 | HMPREF0349_0483 | 32.834 | 95 | 2.50E-140 | 433 | putative alpha-1,2-mannosidase [Enterococcus faecalis TX1322] |
| Q8A1Y1 | HMPREF9517_03053 | 32.834 | 95 | 2.50E-140 | 433 | putative alpha-1,2-mannosidase [Enterococcus faecalis TX1341] |
| Q8A1Y1 | HMPREF9494_02405 | 32.834 | 95 | 2.73E-140 | 433 | putative alpha-1,2-mannosidase [Enterococcus faecalis TX2137] |
| Q8A1Y1 | ENT_15020 | 32.834 | 95 | 3.17E-140 | 433 | alpha-1,2-mannosidase, putative [Enterococcus sp. 7L76] |
| Q8A1Y1 | HMPREF9521_02858 | 32.834 | 95 | 3.17E-140 | 433 | putative alpha-1,2-mannosidase [Enterococcus faecalis TX2134] |
| Q8A1Y1 | CUI_1538 | 32.698 | 95 | 1.66E-139 | 431 | putative alpha-1,2-mannosidase [Enterococcus faecalis PC1.1] |
| Q8A1Y1 | HMPREF9516_02045 | 32.698 | 95 | 1.66E-139 | 431 | putative alpha-1,2-mannosidase [Enterococcus faecalis TX1302] |
| Q8A1Y1 | HMPREF9497_02798 | 32.698 | 95 | 1.66E-139 | 431 | putative alpha-1,2-mannosidase [Enterococcus faecalis TX4244] |
| Q8A1Y1 | HMPREF9518_00653 | 32.834 | 95 | 3.33E-139 | 431 | putative alpha-1,2-mannosidase [Enterococcus faecalis TX1342] |
| Q8A1Y1 | HMPREF9478_00919 | 33.471 | 95 | 3.52E-139 | 431 | hypothetical protein [Enterococcus saccharolyticus 30_1] |
| Q8A1Y1 | HMPREF9456_02581 | 34.531 | 99 | 6.84E-139 | 437 | hypothetical protein [Dysgonomonas mossii DSM 22836] |
| Q8A1Y1 | HMPREF9519_01315 | 32.698 | 95 | 7.92E-139 | 429 | putative alpha-1,2-mannosidase [Enterococcus faecalis TX1346] |
| Q8A1Y1 | HMPREF9447_02712 | 33.243 | 94 | 1.92E-138 | 429 | hypothetical protein [Bacteroides oleiciplenus YIT 12058] |
| Q8A1Y1 | PRABACTJOHN_00262 | 34.538 | 94 | 3.18E-137 | 426 | putative alpha-1,2-mannosidase [Parabacteroides johnsonii DSM 18315] |
| Q8A1Y1 | HMPREF9449_00242 | 34.815 | 99 | 3.55E-137 | 427 | putative alpha-1,2-mannosidase [Odoribacter laneus YIT 12061] |
| Q8A1Y1 | PARMER_03390 | 34.631 | 94 | 6.37E-137 | 425 | putative alpha-1,2-mannosidase [Parabacteroides merdae ATCC 43184] |
| Q8A1Y1 | BACINT_01519 | 33.636 | 96 | 1.00E-136 | 425 | putative alpha-1,2-mannosidase [Bacteroides intestinalis DSM 17393] |
| Q8A1Y1 | HMPREF9446_03563 | 35 | 94 | 1.14E-136 | 426 | putative alpha-1,2-mannosidase [Bacteroides fluxus YIT 12057] |
| Q8A1Y1 | BSHG_0888 | 36.544 | 96 | 1.20E-136 | 425 | putative alpha-1,2-mannosidase [Bacteroides sp. 3_2_5] |
| Q8A1Y1 | HMPREF9007_01545 | 34.491 | 99 | 1.34E-136 | 426 | alpha-1,2-mannosidase family protein [Bacteroides sp. 1_1_14] |
| Q8A1Y1 | HMPREF0101_02661 | 36.544 | 96 | 1.39E-136 | 424 | putative alpha-1,2-mannosidase [Bacteroides sp. 2_1_16] |
| Q8A1Y1 | BFAG_00095 | 36.828 | 96 | 1.57E-136 | 424 | putative alpha-1,2-mannosidase [Bacteroides fragilis 3_1_12] |

| | | | | | | |
|---|---|---|---|---|---|---|
| Q8A1Y1 | BSIG_2706 | 34.491 | 99 | 2.54E-136 | 425 | putative alpha-1,2-mannosidase [Bacteroides sp. 1_1_6] |
| Q8A1Y1 | HMPREF1018_00780 | 36.412 | 96 | 4.30E-136 | 423 | hypothetical protein [Bacteroides sp. 2_1_56FAA] |
| Q8A1Y1 | HMPREF0101_00696 | 34.533 | 96 | 4.68E-136 | 430 | putative alpha-1,2-mannosidase [Bacteroides sp. 2_1_16] |
| Q8A1Y1 | BACCELL_02295 | 33.333 | 94 | 4.76E-136 | 423 | putative alpha-1,2-mannosidase [Bacteroides cellulosilyticus DSM 14838] |
| Q8A1Y1 | HMPREF1018_01858 | 34.533 | 96 | 6.84E-136 | 430 | hypothetical protein [Bacteroides sp. 2_1_56FAA] |
| Q8A1Y1 | HMPREF9447_02948 | 33.938 | 99 | 1.25E-135 | 429 | hypothetical protein [Bacteroides oleiciplenus YIT 12058] |
| Q8A1Y1 | BFAG_03931 | 34.371 | 99 | 2.44E-135 | 422 | putative alpha-1,2-mannosidase [Bacteroides fragilis 3_1_12] |
| Q8A1Y1 | BACCELL_03366 | 36.471 | 97 | 3.12E-135 | 421 | putative alpha-1,2-mannosidase [Bacteroides cellulosilyticus DSM 14838] |
| Q8A1Y1 | HMPREF1532_01618 | 32.992 | 97 | 3.85E-135 | 421 | hypothetical protein [Bacteroides salyersiae WAL 10018 = DSM 18765 = JCM 12988] |
| Q8A1Y1 | HMPREF0106_00702 | 34.271 | 97 | 1.14E-134 | 421 | alpha-1,2-mannosidase family protein [Bacteroides sp. D22] |
| Q8A1Y1 | CUY_1144 | 34.177 | 97 | 1.51E-134 | 421 | putative alpha-1,2-mannosidase [Bacteroides ovatus SD CMC 3f] |
| Q8A1Y1 | HMPREF1214_01298 | 36.184 | 96 | 2.48E-134 | 419 | hypothetical protein [Bacteroides sp. HPS0048] |
| Q8A1Y1 | BSCG_04298 | 34.576 | 96 | 3.39E-134 | 420 | putative alpha-1,2-mannosidase [Bacteroides sp. 2_2_4] |
| Q8A1Y1 | BXY_30760 | 34.576 | 96 | 3.39E-134 | 420 | alpha-1,2-mannosidase, putative [Bacteroides xylanisolvens XB1A] |
| Q8A1Y1 | HMPREF1211_02120 | 34.238 | 96 | 3.69E-134 | 432 | hypothetical protein [Streptomyces sp. HGB0020] |
| Q8A1Y1 | BACINT_04252 | 36.315 | 95 | 1.01E-133 | 417 | putative alpha-1,2-mannosidase [Bacteroides intestinalis DSM 17393] |
| Q8A1Y1 | HMPREF0102_03219 | 34.051 | 97 | 1.11E-133 | 418 | putative alpha-1,2-mannosidase [Bacteroides sp. 2_1_22] |
| Q8A1Y1 | BSAG_03094 | 34.051 | 97 | 1.11E-133 | 418 | putative alpha-1,2-mannosidase [Bacteroides sp. D1] |
| Q8A1Y1 | CW3_1777 | 34.051 | 97 | 1.11E-133 | 418 | putative alpha-1,2-mannosidase [Bacteroides xylanisolvens SD CC 1b] |
| Q8A1Y1 | CW1_2593 | 34.051 | 97 | 1.11E-133 | 418 | putative alpha-1,2-mannosidase [Bacteroides xylanisolvens SD CC 2a] |
| Q8A1Y1 | HMPREF1018_02985 | 33.873 | 99 | 1.39E-133 | 418 | hypothetical protein [Bacteroides sp. 2_1_56FAA] |
| Q8A1Y1 | HMPREF0127_01912 | 34.177 | 97 | 1.79E-133 | 418 | hypothetical protein [Bacteroides sp. 1_1_30] |
| Q8A1Y1 | BSGG_1591 | 34.271 | 97 | 1.93E-133 | 418 | putative alpha-1,2-mannosidase [Bacteroides sp. D2] |
| Q8A1Y1 | HMPREF1214_00022 | 33.553 | 94 | 2.11E-133 | 416 | hypothetical protein [Bacteroides sp. HPS0048] |
| Q8A1Y1 | BSEG_01944 | 33.29 | 96 | 2.52E-133 | 417 | putative alpha-1,2-mannosidase [Bacteroides dorei 5_1_36/D4] |
| Q8A1Y1 | BSBG_01024 | 33.29 | 96 | 2.52E-133 | 417 | putative alpha-1,2-mannosidase [Bacteroides sp. 9_1_42FAA] |
| Q8A1Y1 | HMPREF1017_03075 | 34.131 | 98 | 2.82E-133 | 417 | hypothetical protein [Bacteroides ovatus 3_8_47FAA] |
| Q8A1Y1 | BACUNI_02648 | 36.17 | 95 | 4.68E-133 | 415 | putative alpha-1,2-mannosidase [Bacteroides uniformis ATCC 8492] |
| Q8A1Y1 | HMPREF0105_2294 | 33.29 | 96 | 4.92E-133 | 416 | putative alpha-1,2-mannosidase [Bacteroides sp. 3_1_33FAA] |
| Q8A1Y1 | HMPREF9447_01544 | 35.964 | 97 | 7.38E-133 | 415 | hypothetical protein [Bacteroides oleiciplenus YIT 12058] |
| Q8A1Y1 | HMPREF0969_00778 | 36.085 | 95 | 7.57E-133 | 414 | putative alpha-1,2-mannosidase [Bacteroides sp. D20] |
| Q8A1Y1 | CLOSPI_00338 | 34.702 | 94 | 7.71E-133 | 433 | putative alpha-1,2-mannosidase [Erysipelatoclostridium [Clostridium] spiroforme DSM 1552] |
| Q8A1Y1 | BACDOR_04886 | 33.29 | 96 | 9.12E-133 | 415 | putative alpha-1,2-mannosidase [Bacteroides dorei DSM 17855] |
| Q8A1Y1 | HMPREF1007_03655 | 35.64 | 97 | 1.12E-132 | 414 | glycosyl hydrolase family 92 [Bacteroides sp. 4_1_36] |
| Q8A1Y1 | HMPREF9456_00934 | 33.584 | 98 | 1.20E-132 | 416 | hypothetical protein [Dysgonomonas mossii DSM 22836] |
| Q8A1Y1 | PARMER_01576 | 33.81 | 95 | 1.56E-132 | 415 | putative alpha-1,2-mannosidase [Parabacteroides merdae ATCC 43184] |
| Q8A1Y1 | HMPREF9455_00268 | 33.634 | 99 | 2.15E-132 | 421 | hypothetical protein [Dysgonomonas gadei ATCC BAA-286] |
| Q8A1Y1 | HMPREF1017_04454 | 35.405 | 93 | 2.85E-132 | 413 | hypothetical protein [Bacteroides ovatus 3_8_47FAA] |
| Q8A1Y1 | HMPREF9455_03238 | 33.806 | 96 | 5.03E-132 | 414 | hypothetical protein [Dysgonomonas gadei ATCC BAA-286] |
| Q8A1Y1 | HMPREF9010_02547 | 34.005 | 98 | 5.67E-132 | 414 | alpha-1,2-mannosidase family protein [Bacteroides sp. 3_1_23] |
| Q8A1Y1 | BACSTE_03551 | 36.559 | 94 | 6.64E-132 | 412 | putative alpha-1,2-mannosidase [Bacteroides stercoris ATCC 43183] |
| Q8A1Y1 | HMPREF0101_03828 | 33.624 | 99 | 9.21E-132 | 413 | putative alpha-1,2-mannosidase [Bacteroides sp. 2_1_16] |
| Q8A1Y1 | BSHG_3280 | 33.624 | 99 | 9.41E-132 | 413 | putative alpha-1,2-mannosidase [Bacteroides sp. 3_2_5] |
| Q8A1Y1 | HMPREF9446_02424 | 33.511 | 96 | 1.11E-131 | 419 | putative alpha-1,2-mannosidase [Bacteroides fluxus YIT 12057] |
| Q8A1Y1 | HMPREF9455_03240 | 35.117 | 96 | 2.70E-131 | 412 | hypothetical protein [Dysgonomonas gadei ATCC BAA-286] |
| Q8A1Y1 | BSEG_01829 | 35.248 | 97 | 2.71E-131 | 411 | putative alpha-1,2-mannosidase [Bacteroides dorei 5_1_36/D4] |
| Q8A1Y1 | HMPREF1532_01083 | 36.376 | 93 | 3.37E-131 | 410 | hypothetical protein [Bacteroides salyersiae WAL 10018 = DSM 18765 = JCM 12988] |
| Q8A1Y1 | BACPLE_00636 | 35.088 | 93 | 3.67E-131 | 411 | putative alpha-1,2-mannosidase [Bacteroides plebeius DSM 17135] |
| Q8A1Y1 | BACCOPRO_02654 | 33.464 | 96 | 6.09E-131 | 410 | putative alpha-1,2-mannosidase [Bacteroides coprophilus DSM 18228 = JCM 13818] |
| Q8A1Y1 | HMPREF1214_04023 | 33.812 | 96 | 6.66E-131 | 410 | hypothetical protein [Bacteroides sp. HPS0048] |
| Q8A1Y1 | BACCOPRO_01389 | 33.892 | 96 | 7.06E-131 | 411 | putative alpha-1,2-mannosidase [Bacteroides coprophilus DSM 18228 = JCM 13818] |
| Q8A1Y1 | BACDOR_01137 | 35.248 | 97 | 1.23E-130 | 409 | putative alpha-1,2-mannosidase [Bacteroides dorei DSM 17855] |
| Q8A1Y1 | HMPREF0105_2409 | 35.248 | 97 | 1.59E-130 | 409 | putative alpha-1,2-mannosidase [Bacteroides sp. 3_1_33FAA] |
| Q8A1Y1 | BSBG_01324 | 35.248 | 97 | 1.59E-130 | 409 | putative alpha-1,2-mannosidase [Bacteroides sp. 9_1_42FAA] |
| Q8A1Y1 | HMPREF0127_01955 | 35.348 | 96 | 2.11E-130 | 409 | hypothetical protein [Bacteroides sp. 1_1_30] |
| Q8A1Y1 | HMPREF9011_00723 | 35.379 | 97 | 2.17E-130 | 409 | glycoside hydrolase family 92 [Bacteroides sp. 3_1_40A] |
| Q8A1Y1 | HMPREF0101_02658 | 34.231 | 97 | 2.37E-130 | 409 | putative alpha-1,2-mannosidase [Bacteroides sp. 2_1_16] |
| Q8A1Y1 | BSGG_2469 | 33.64 | 96 | 3.39E-130 | 408 | putative alpha-1,2-mannosidase [Bacteroides sp. D2] |
| Q8A1Y1 | CUU_0906 | 35.379 | 97 | 3.92E-130 | 408 | putative alpha-1,2-mannosidase [Bacteroides vulgatus PC510] |
| Q8A1Y1 | BSFG_01743 | 35.379 | 97 | 3.92E-130 | 408 | putative alpha-1,2-mannosidase [Bacteroides sp. 4_3_47FAA] |

| | | | | | | |
|---|---|---|---|---|---|---|
| Q8A1Y1 | BACSTE_03661 | 34.153 | 99 | 4.46E-130 | 414 | putative alpha-1,2-mannosidase [Bacteroides stercoris ATCC 43183] |
| Q8A1Y1 | HMPREF0969_02361 | 32.513 | 99 | 4.86E-130 | 414 | putative alpha-1,2-mannosidase [Bacteroides sp. D20] |
| Q8A1Y1 | BSGG_1788 | 32.078 | 97 | 5.43E-130 | 407 | putative alpha-1,2-mannosidase [Bacteroides sp. D2] |
| Q8A1Y1 | HMPREF0969_00769 | 34.759 | 94 | 5.82E-130 | 408 | putative alpha-1,2-mannosidase [Bacteroides sp. D20] |
| Q8A1Y1 | BACUNI_02639 | 34.759 | 94 | 5.82E-130 | 408 | putative alpha-1,2-mannosidase [Bacteroides uniformis ATCC 8492] |
| Q8A1Y1 | HMPREF1007_03646 | 34.759 | 94 | 5.82E-130 | 408 | glycosyl hydrolase family 92 [Bacteroides sp. 4_1_36] |
| Q8A1Y1 | BACCOP_04376 | 34.225 | 94 | 6.58E-130 | 408 | putative alpha-1,2-mannosidase [Bacteroides coprocola DSM 17136] |
| Q8A1Y1 | BSFG_00538 | 32.474 | 96 | 9.80E-130 | 408 | putative alpha-1,2-mannosidase [Bacteroides sp. 4_3_47FAA] |
| Q8A1Y1 | HMPREF9011_00613 | 32.474 | 96 | 9.80E-130 | 408 | hypothetical protein [Bacteroides sp. 3_1_40A] |
| Q8A1Y1 | BACSTE_03542 | 34.94 | 93 | 1.11E-129 | 407 | putative alpha-1,2-mannosidase [Bacteroides stercoris ATCC 43183] |
| Q8A1Y1 | CUU_0909 | 34.026 | 96 | 1.19E-129 | 407 | putative alpha-1,2-mannosidase [Bacteroides vulgatus PC510] |
| Q8A1Y1 | BACCOP_04391 | 34.536 | 97 | 1.24E-129 | 406 | putative alpha-1,2-mannosidase [Bacteroides coprocola DSM 17136] |
| Q8A1Y1 | BSHG_2180 | 34.225 | 94 | 1.54E-129 | 406 | putative alpha-1,2-mannosidase [Bacteroides sp. 3_2_5] |
| Q8A1Y1 | HMPREF1018_03842 | 34.225 | 94 | 1.54E-129 | 406 | hypothetical protein [Bacteroides sp. 2_1_56FAA] |
| Q8A1Y1 | HMPREF0101_02879 | 34.225 | 94 | 1.54E-129 | 406 | putative alpha-1,2-mannosidase [Bacteroides sp. 2_1_16] |
| Q8A1Y1 | CUY_1097 | 35.348 | 96 | 1.75E-129 | 406 | putative alpha-1,2-mannosidase [Bacteroides ovatus SD CMC 3f] |
| Q8A1Y1 | BSAG_03132 | 35.425 | 96 | 1.79E-129 | 406 | putative alpha-1,2-mannosidase [Bacteroides sp. D1] |
| Q8A1Y1 | CW3_3119 | 35.425 | 96 | 1.79E-129 | 406 | putative alpha-1,2-mannosidase [Bacteroides xylanisolvens SD CC 1b] |
| Q8A1Y1 | HMPREF1017_03045 | 35.294 | 96 | 1.89E-129 | 406 | hypothetical protein [Bacteroides ovatus 3_8_47FAA] |
| Q8A1Y1 | CW1_2680 | 35.425 | 96 | 1.92E-129 | 406 | putative alpha-1,2-mannosidase [Bacteroides xylanisolvens SD CC 2a] |
| Q8A1Y1 | HMPREF9441_01695 | 33.735 | 94 | 2.19E-129 | 405 | putative alpha-1,2-mannosidase [Paraprevotella clara YIT 11840] |
| Q8A1Y1 | BXY_31180 | 35.724 | 93 | 2.41E-129 | 405 | alpha-1,2-mannosidase, putative [Bacteroides xylanisolvens XB1A] |
| Q8A1Y1 | HMPREF0106_00734 | 35.724 | 93 | 2.59E-129 | 405 | alpha-1,2-mannosidase family protein [Bacteroides sp. D22] |
| Q8A1Y1 | HMPREF1033_02713 | 33.205 | 97 | 3.69E-129 | 406 | hypothetical protein [Tannerella sp. 6_1_58FAA_CT1] |
| Q8A1Y1 | BFAG_03531 | 34.057 | 93 | 3.89E-129 | 405 | putative alpha-1,2-mannosidase [Bacteroides fragilis 3_1_12] |
| Q8A1Y1 | BSFG_01740 | 34.026 | 96 | 3.95E-129 | 406 | putative alpha-1,2-mannosidase [Bacteroides sp. 4_3_47FAA] |
| Q8A1Y1 | HMPREF9011_00726 | 34.026 | 96 | 3.95E-129 | 406 | glycoside hydrolase family 92 [Bacteroides sp. 3_1_40A] |
| Q8A1Y1 | BSGG_1566 | 35.332 | 97 | 4.73E-129 | 405 | putative alpha-1,2-mannosidase [Bacteroides sp. D2] |
| Q8A1Y1 | BACCAC_01687 | 35.995 | 93 | 5.58E-129 | 404 | putative alpha-1,2-mannosidase [Bacteroides caccae ATCC 43185] |
| Q8A1Y1 | HMPREF9010_02581 | 35.666 | 93 | 5.95E-129 | 404 | alpha-1,2-mannosidase family protein [Bacteroides sp. 3_1_23] |
| Q8A1Y1 | BSHG_0891 | 34.103 | 97 | 6.68E-129 | 405 | putative alpha-1,2-mannosidase [Bacteroides sp. 3_2_5] |
| Q8A1Y1 | HMPREF0102_03259 | 35.801 | 93 | 7.76E-129 | 404 | putative alpha-1,2-mannosidase [Bacteroides sp. 2_1_22] |
| Q8A1Y1 | HMPREF1033_02199 | 35.171 | 96 | 9.99E-129 | 404 | hypothetical protein [Tannerella sp. 6_1_58FAA_CT1] |
| Q8A1Y1 | CUU_0351 | 32.216 | 96 | 1.02E-128 | 405 | putative alpha-1,2-mannosidase [Bacteroides vulgatus PC510] |
| Q8A1Y1 | HMPREF0847_01777 | 32.398 | 93 | 1.11E-128 | 403 | putative alpha-1,2-mannosidase [Streptococcus sp. 2_1_36FAA] |
| Q8A1Y1 | BACCELL_03377 | 33.914 | 93 | 1.23E-128 | 404 | putative alpha-1,2-mannosidase [Bacteroides cellulosilyticus DSM 14838] |
| Q8A1Y1 | BFAG_00092 | 34.487 | 97 | 1.36E-128 | 404 | putative alpha-1,2-mannosidase [Bacteroides fragilis 3_1_12] |
| Q8A1Y1 | BSEG_01826 | 33.987 | 96 | 1.39E-128 | 404 | putative alpha-1,2-mannosidase [Bacteroides dorei 5_1_36/D4] |
| Q8A1Y1 | BSBG_01327 | 33.987 | 96 | 1.56E-128 | 404 | putative alpha-1,2-mannosidase [Bacteroides sp. 9_1_42FAA] |
| Q8A1Y1 | BSIG_2671 | 34.993 | 96 | 1.60E-128 | 404 | putative alpha-1,2-mannosidase [Bacteroides sp. 1_1_6] |
| Q8A1Y1 | HMPREF1007_01972 | 32.383 | 99 | 1.64E-128 | 410 | glycosyl hydrolase family 92 [Bacteroides sp. 4_1_36] |
| Q8A1Y1 | BACUNI_03279 | 32.383 | 99 | 1.95E-128 | 411 | putative alpha-1,2-mannosidase [Bacteroides uniformis ATCC 8492] |
| Q8A1Y1 | HMPREF0105_2412 | 33.987 | 96 | 2.24E-128 | 404 | putative alpha-1,2-mannosidase [Bacteroides sp. 3_1_33FAA] |
| Q8A1Y1 | HMPREF9007_01575 | 34.993 | 96 | 2.28E-128 | 403 | alpha-1,2-mannosidase family protein [Bacteroides sp. 1_1_14] |
| Q8A1Y1 | BSIG_1521 | 32.941 | 97 | 2.78E-128 | 403 | putative alpha-1,2-mannosidase [Bacteroides sp. 1_1_6] |
| Q8A1Y1 | HMPREF0103_3527 | 34.902 | 97 | 3.13E-128 | 403 | putative alpha-1,2-mannosidase [Bacteroides sp. 2_1_33B] |
| Q8A1Y1 | HMPREF1018_00777 | 33.974 | 97 | 3.16E-128 | 403 | hypothetical protein [Bacteroides sp. 2_1_56FAA] |
| Q8A1Y1 | HMPREF1017_05023 | 33.777 | 96 | 3.85E-128 | 419 | hypothetical protein [Bacteroides ovatus 3_8_47FAA] |
| Q8A1Y1 | PRABACTJOHN_04168 | 32.702 | 99 | 4.54E-128 | 403 | putative alpha-1,2-mannosidase [Parabacteroides johnsonii DSM 18315] |
| Q8A1Y1 | BACFIN_05349 | 35.724 | 93 | 4.89E-128 | 402 | putative alpha-1,2-mannosidase [Bacteroides finegoldii DSM 17565] |
| Q8A1Y1 | CUU_3832 | 34.474 | 96 | 4.94E-128 | 402 | putative alpha-1,2-mannosidase [Bacteroides vulgatus PC510] |
| Q8A1Y1 | HMPREF0104_00314 | 33.03 | 98 | 6.14E-128 | 409 | alpha-1,2-mannosidase family protein [Bacteroides sp. 3_1_19] |
| Q8A1Y1 | HMPREF0103_1974 | 33.246 | 98 | 6.17E-128 | 409 | putative alpha-1,2-mannosidase [Bacteroides sp. 2_1_33B] |
| Q8A1Y1 | BSCG_04347 | 35.589 | 93 | 7.98E-128 | 402 | putative alpha-1,2-mannosidase [Bacteroides sp. 2_2_4] |
| Q8A1Y1 | BACOVA_04813 | 35.318 | 93 | 8.69E-128 | 401 | putative alpha-1,2-mannosidase [Bacteroides ovatus ATCC 8483] |
| Q8A1Y1 | BACDOR_01140 | 33.856 | 96 | 1.01E-127 | 402 | putative alpha-1,2-mannosidase [Bacteroides dorei DSM 17855] |
| Q8A1Y1 | HMPREF9447_01531 | 33.914 | 94 | 1.04E-127 | 402 | hypothetical protein [Bacteroides oleiciplenus YIT 12058] |
| Q8A1Y1 | HMPREF0127_03506 | 33.696 | 94 | 1.08E-127 | 418 | hypothetical protein [Bacteroides sp. 1_1_30] |
| Q8A1Y1 | HMPREF9449_00243 | 33.924 | 99 | 1.29E-127 | 402 | putative alpha-1,2-mannosidase [Odoribacter laneus YIT 12061] |
| Q8A1Y1 | HMPREF0619_00369 | 33.116 | 98 | 1.53E-127 | 407 | putative alpha-1,2-mannosidase [Parabacteroides sp. D13] |

| | | | | | | |
|---|---|---|---|---|---|---|
| Q8A1Y1 | BSIG_5236 | 34.328 | 93 | 1.72E-127 | 400 | putative alpha-1,2-mannosidase [Bacteroides sp. 1_1_6] |
| Q8A1Y1 | HMPREF0104_01159 | 34.771 | 97 | 1.98E-127 | 402 | alpha-1,2-mannosidase family protein [Bacteroides sp. 3_1_19] |
| Q8A1Y1 | HMPREF9450_00989 | 33.873 | 99 | 2.31E-127 | 402 | hypothetical protein [Alistipes indistinctus YIT 12060] |
| Q8A1Y1 | BSFG_01673 | 34.342 | 96 | 2.31E-127 | 400 | putative alpha-1,2-mannosidase [Bacteroides sp. 4_3_47FAA] |
| Q8A1Y1 | HMPREF9011_00793 | 34.342 | 96 | 2.31E-127 | 400 | glycoside hydrolase family 92 [Bacteroides sp. 3_1_40A] |
| Q8A1Y1 | BSGG_1569 | 33.912 | 94 | 2.85E-127 | 401 | putative alpha-1,2-mannosidase [Bacteroides sp. D2] |
| Q8A1Y1 | HMPREF9456_00933 | 34.817 | 96 | 2.90E-127 | 402 | hypothetical protein [Dysgonomonas mossii DSM 22836] |
| Q8A1Y1 | HMPREF9456_02505 | 34.051 | 94 | 2.99E-127 | 401 | hypothetical protein [Dysgonomonas mossii DSM 22836] |
| Q8A1Y1 | HMPREF9008_03082 | 34.641 | 97 | 3.09E-127 | 401 | putative alpha-1,2-mannosidase [Parabacteroides sp. 20_3] |
| Q8A1Y1 | BACCOPRO_02262 | 33.163 | 98 | 3.32E-127 | 400 | putative alpha-1,2-mannosidase [Bacteroides coprophilus DSM 18228 = JCM 13818] |
| Q8A1Y1 | BSGG_4713 | 33.152 | 93 | 3.70E-127 | 400 | putative alpha-1,2-mannosidase [Bacteroides sp. D2] |
| Q8A1Y1 | BACOVA_01687 | 33.644 | 96 | 4.27E-127 | 416 | putative alpha-1,2-mannosidase [Bacteroides ovatus ATCC 8483] |
| Q8A1Y1 | HMPREF9007_03696 | 32.81 | 97 | 5.29E-127 | 399 | alpha-1,2-mannosidase family protein [Bacteroides sp. 1_1_14] |
| Q8A1Y1 | HMPREF9007_01979 | 34.057 | 93 | 5.94E-127 | 399 | alpha-1,2-mannosidase family protein [Bacteroides sp. 1_1_14] |
| Q8A1Y1 | BACINT_04242 | 33.118 | 97 | 7.11E-127 | 400 | putative alpha-1,2-mannosidase [Bacteroides intestinalis DSM 17393] |
| Q8A1Y1 | BACPLE_02380 | 33.376 | 96 | 7.66E-127 | 400 | putative alpha-1,2-mannosidase [Bacteroides plebeius DSM 17135] |
| Q8A1Y1 | PARMER_02507 | 35.589 | 93 | 7.82E-127 | 399 | putative alpha-1,2-mannosidase [Parabacteroides merdae ATCC 43184] |
| Q8A1Y1 | HMPREF9448_01740 | 32.524 | 94 | 9.11E-127 | 399 | hypothetical protein [Barnesiella intestinihominis YIT 11860] |
| Q8A1Y1 | HMPREF9447_03114 | 33.921 | 93 | 9.98E-127 | 399 | hypothetical protein [Bacteroides oleiciplenus YIT 12058] |
| Q8A1Y1 | HMPREF9008_04130 | 32.855 | 98 | 1.03E-126 | 405 | putative alpha-1,2-mannosidase [Parabacteroides sp. 20_3] |
| Q8A1Y1 | HMPREF0619_01141 | 34.641 | 97 | 1.07E-126 | 400 | putative alpha-1,2-mannosidase [Parabacteroides sp. D13] |
| Q8A1Y1 | HMPREF0103_1774 | 32.789 | 98 | 1.14E-126 | 400 | putative alpha-1,2-mannosidase [Bacteroides sp. 2_1_33B] |
| Q8A1Y1 | HMPREF1033_00968 | 33.646 | 94 | 1.21E-126 | 399 | hypothetical protein [Tannerella sp. 6_1_58FAA_CT1] |
| Q8A1Y1 | HMPREF9008_03595 | 32.789 | 98 | 1.50E-126 | 400 | putative alpha-1,2-mannosidase [Parabacteroides sp. 20_3] |
| Q8A1Y1 | HMPREF1481_01621 | 32.823 | 94 | 3.74E-126 | 396 | hypothetical protein [Streptococcus sp. HPH0090] |
| Q8A1Y1 | HMPREF1214_00154 | 32.353 | 95 | 4.36E-126 | 397 | hypothetical protein [Bacteroides sp. HPS0048] |
| Q8A1Y1 | HMPREF9520_03180 | 32.985 | 87 | 4.42E-126 | 394 | putative alpha-1,2-mannosidase [Enterococcus faecalis TX1467] |
| Q8A1Y1 | BSCG_04344 | 33.867 | 94 | 4.74E-126 | 398 | putative alpha-1,2-mannosidase [Bacteroides sp. 2_2_4] |
| Q8A1Y1 | HMPREF0531_12430 | 30.78 | 94 | 7.06E-126 | 396 | putative alpha-1,2-mannosidase [Lactobacillus plantarum subsp. plantarum ATCC 14917 = JCM 1149 =] |
| Q8A1Y1 | CUY_2236 | 33.152 | 93 | 7.91E-126 | 396 | putative alpha-1,2-mannosidase [Bacteroides ovatus SD CMC 3f] |
| Q8A1Y1 | HMPREF0127_04405 | 33.152 | 93 | 1.06E-125 | 396 | hypothetical protein [Bacteroides sp. 1_1_30] |
| Q8A1Y1 | HMPREF0105_3796 | 33.816 | 96 | 1.42E-125 | 395 | putative alpha-1,2-mannosidase [Bacteroides sp. 3_1_33FAA] |
| Q8A1Y1 | HMPREF0102_03256 | 33.867 | 94 | 1.61E-125 | 397 | putative alpha-1,2-mannosidase [Bacteroides sp. 2_1_22] |
| Q8A1Y1 | BSAG_03129 | 33.867 | 94 | 1.61E-125 | 397 | putative alpha-1,2-mannosidase [Bacteroides sp. D1] |
| Q8A1Y1 | CW3_3116 | 33.867 | 94 | 1.61E-125 | 397 | putative alpha-1,2-mannosidase [Bacteroides xylanisolvens SD CC 1b] |
| Q8A1Y1 | CW1_2677 | 33.867 | 94 | 1.61E-125 | 397 | putative alpha-1,2-mannosidase [Bacteroides xylanisolvens SD CC 2a] |
| Q8A1Y1 | HMPREF0619_00578 | 32.538 | 98 | 2.07E-125 | 397 | putative alpha-1,2-mannosidase [Parabacteroides sp. D13] |
| Q8A1Y1 | BACDOR_01210 | 33.947 | 96 | 2.49E-125 | 395 | putative alpha-1,2-mannosidase [Bacteroides dorei DSM 17855] |
| Q8A1Y1 | BSEG_03714 | 33.816 | 96 | 3.18E-125 | 395 | putative alpha-1,2-mannosidase [Bacteroides dorei 5_1_36/D4] |
| Q8A1Y1 | BSBG_04080 | 33.816 | 96 | 3.18E-125 | 395 | putative alpha-1,2-mannosidase [Bacteroides sp. 9_1_42FAA] |
| Q8A1Y1 | HMPREF1017_04445 | 33.016 | 93 | 3.66E-125 | 394 | hypothetical protein [Bacteroides ovatus 3_8_47FAA] |
| Q8A1Y1 | HMPREF9448_00369 | 33.633 | 97 | 5.22E-125 | 395 | hypothetical protein [Barnesiella intestinihominis YIT 11860] |
| Q8A1Y1 | HMPREF1017_03048 | 33.733 | 94 | 5.27E-125 | 395 | hypothetical protein [Bacteroides ovatus 3_8_47FAA] |
| Q8A1Y1 | BACCOP_04378 | 35.405 | 93 | 5.86E-125 | 394 | putative alpha-1,2-mannosidase [Bacteroides coprocola DSM 17136] |
| Q8A1Y1 | BXY_31150 | 33.867 | 94 | 6.45E-125 | 395 | alpha-1,2-mannosidase, putative [Bacteroides xylanisolvens XB1A] |
| Q8A1Y1 | HMPREF0106_00731 | 33.867 | 94 | 6.45E-125 | 395 | alpha-1,2-mannosidase family protein [Bacteroides sp. D22] |
| Q8A1Y1 | HMPREF9455_03816 | 33.119 | 98 | 8.11E-125 | 394 | hypothetical protein [Dysgonomonas gadei ATCC BAA-286] |
| Q8A1Y1 | BSCG_00024 | 32.88 | 93 | 9.11E-125 | 394 | putative alpha-1,2-mannosidase [Bacteroides sp. 2_2_4] |
| Q8A1Y1 | BXY_17780 | 33.152 | 93 | 9.11E-125 | 394 | alpha-1,2-mannosidase, putative [Bacteroides xylanisolvens XB1A] |
| Q8A1Y1 | HMPREF0102_04198 | 33.152 | 93 | 9.11E-125 | 394 | putative alpha-1,2-mannosidase [Bacteroides sp. 2_1_22] |
| Q8A1Y1 | BSAG_03510 | 33.152 | 93 | 9.11E-125 | 394 | putative alpha-1,2-mannosidase [Bacteroides sp. D1] |
| Q8A1Y1 | HMPREF0106_02791 | 33.152 | 93 | 9.11E-125 | 394 | alpha-1,2-mannosidase family protein [Bacteroides sp. D22] |
| Q8A1Y1 | PRABACTJOHN_02081 | 35.589 | 93 | 9.84E-125 | 394 | putative alpha-1,2-mannosidase [Parabacteroides johnsonii DSM 18315] |
| Q8A1Y1 | CW3_4738 | 33.152 | 93 | 1.00E-124 | 393 | putative alpha-1,2-mannosidase [Bacteroides xylanisolvens SD CC 1b] |
| Q8A1Y1 | CW1_4636 | 33.152 | 93 | 1.00E-124 | 393 | putative alpha-1,2-mannosidase [Bacteroides xylanisolvens SD CC 2a] |
| Q8A1Y1 | HMPREF9442_02649 | 33.199 | 94 | 1.20E-124 | 393 | putative alpha-1,2-mannosidase [Paraprevotella xylaniphila YIT 11841] |
| Q8A1Y1 | HMPREF0127_01952 | 33.733 | 94 | 1.66E-124 | 394 | hypothetical protein [Bacteroides sp. 1_1_30] |
| Q8A1Y1 | HMPREF9010_04724 | 32.88 | 93 | 1.90E-124 | 393 | alpha-1,2-mannosidase family protein [Bacteroides sp. 3_1_23] |
| Q8A1Y1 | BSBG_00732 | 33.917 | 99 | 2.02E-124 | 394 | putative alpha-1,2-mannosidase [Bacteroides sp. 9_1_42FAA] |
| Q8A1Y1 | HMPREF0104_00477 | 32.412 | 98 | 2.11E-124 | 394 | alpha-1,2-mannosidase family protein [Bacteroides sp. 3_1_19] |

| | | | | | | |
|---|---|---|---|---|---|---|
| Q8A1Y1 | BACPLE_02378 | 32.666 | 99 | 2.22E-124 | 394 | putative alpha-1,2-mannosidase [Bacteroides plebeius DSM 17135] |
| Q8A1Y1 | CUY_1100 | 33.733 | 94 | 3.37E-124 | 393 | putative alpha-1,2-mannosidase [Bacteroides ovatus SD CMC 3f] |
| Q8A1Y1 | BACOVA_04810 | 33.6 | 94 | 3.56E-124 | 393 | putative alpha-1,2-mannosidase [Bacteroides ovatus ATCC 8483] |
| Q8A1Y1 | PARMER_03860 | 33.42 | 96 | 3.63E-124 | 393 | putative alpha-1,2-mannosidase [Parabacteroides merdae ATCC 43184] |
| Q8A1Y1 | BACFIN_06170 | 31.884 | 96 | 1.67E-123 | 390 | putative alpha-1,2-mannosidase [Bacteroides finegoldii DSM 17565] |
| Q8A1Y1 | BSCG_04300 | 32.866 | 98 | 2.52E-123 | 391 | putative alpha-1,2-mannosidase [Bacteroides sp. 2_2_4] |
| Q8A1Y1 | HMPREF9007_01572 | 32.005 | 97 | 3.22E-123 | 390 | alpha-1,2-mannosidase family protein [Bacteroides sp. 1_1_14] |
| Q8A1Y1 | CUY_1142 | 33.333 | 96 | 6.75E-123 | 390 | putative alpha-1,2-mannosidase [Bacteroides ovatus SD CMC 3f] |
| Q8A1Y1 | HMPREF1017_03073 | 33.246 | 96 | 7.27E-123 | 390 | hypothetical protein [Bacteroides ovatus 3_8_47FAA] |
| Q8A1Y1 | BACOVA_02942 | 32.745 | 93 | 8.32E-123 | 389 | putative alpha-1,2-mannosidase [Bacteroides ovatus ATCC 8483] |
| Q8A1Y1 | BACFIN_05352 | 33.422 | 94 | 8.73E-123 | 389 | putative alpha-1,2-mannosidase [Bacteroides finegoldii DSM 17565] |
| Q8A1Y1 | HMPREF9010_02549 | 33.246 | 96 | 9.66E-123 | 390 | alpha-1,2-mannosidase family protein [Bacteroides sp. 3_1_23] |
| Q8A1Y1 | HMPREF9010_02578 | 33.467 | 94 | 1.23E-122 | 389 | alpha-1,2-mannosidase family protein [Bacteroides sp. 3_1_23] |
| Q8A1Y1 | HMPREF9450_00814 | 33.814 | 96 | 1.27E-122 | 389 | hypothetical protein [Alistipes indistinctus YIT 12060] |
| Q8A1Y1 | BSBG_00734 | 33.775 | 94 | 1.83E-122 | 388 | putative alpha-1,2-mannosidase [Bacteroides sp. 9_1_42FAA] |
| Q8A1Y1 | BSIG_2674 | 31.877 | 97 | 1.83E-122 | 389 | putative alpha-1,2-mannosidase [Bacteroides sp. 1_1_6] |
| Q8A1Y1 | BACCAC_03767 | 33.105 | 93 | 2.15E-122 | 387 | putative alpha-1,2-mannosidase [Bacteroides caccae ATCC 43185] |
| Q8A1Y1 | HMPREF0673_01916 | 33.071 | 96 | 3.02E-122 | 387 | putative alpha-1,2-mannosidase [Prevotella stercorea DSM 18206] |
| Q8A1Y1 | HMPREF1532_00417 | 33.984 | 96 | 5.14E-122 | 387 | hypothetical protein [Bacteroides salyersiae WAL 10018 = DSM 18765 = JCM 12988] |
| Q8A1Y1 | BACCAC_01684 | 32.756 | 94 | 5.49E-122 | 387 | putative alpha-1,2-mannosidase [Bacteroides caccae ATCC 43185] |
| Q8A1Y1 | BXY_30780 | 33.115 | 96 | 1.31E-121 | 387 | alpha-1,2-mannosidase, putative [Bacteroides xylanisolvens XB1A] |
| Q8A1Y1 | HMPREF0102_03221 | 32.984 | 96 | 1.41E-121 | 387 | putative alpha-1,2-mannosidase [Bacteroides sp. 2_1_22] |
| Q8A1Y1 | HMPREF0106_00704 | 33.115 | 96 | 1.41E-121 | 387 | alpha-1,2-mannosidase family protein [Bacteroides sp. D22] |
| Q8A1Y1 | CW3_1779 | 32.984 | 96 | 1.47E-121 | 387 | putative alpha-1,2-mannosidase [Bacteroides xylanisolvens SD CC 1b] |
| Q8A1Y1 | CW1_2591 | 32.984 | 96 | 1.47E-121 | 387 | putative alpha-1,2-mannosidase [Bacteroides xylanisolvens SD CC 2a] |
| Q8A1Y1 | BSAG_03096 | 32.984 | 96 | 1.84E-121 | 386 | putative alpha-1,2-mannosidase [Bacteroides sp. D1] |
| Q8A1Y1 | PARMER_03862 | 33.209 | 99 | 2.15E-121 | 386 | putative alpha-1,2-mannosidase [Parabacteroides merdae ATCC 43184] |
| Q8A1Y1 | PARMER_00605 | 33.001 | 99 | 3.77E-121 | 385 | putative alpha-1,2-mannosidase [Parabacteroides merdae ATCC 43184] |
| Q8A1Y1 | HMPREF0127_01914 | 32.984 | 96 | 4.35E-121 | 385 | hypothetical protein [Bacteroides sp. 1_1_30] |
| Q8A1Y1 | BSIG_2704 | 33.595 | 96 | 6.39E-121 | 385 | putative alpha-1,2-mannosidase [Bacteroides sp. 1_1_6] |
| Q8A1Y1 | BSEG_01863 | 32.068 | 97 | 1.22E-120 | 383 | putative alpha-1,2-mannosidase [Bacteroides dorei 5_1_36/D4] |
| Q8A1Y1 | PRABACTJOHN_00883 | 32.83 | 98 | 2.43E-120 | 383 | putative alpha-1,2-mannosidase [Parabacteroides johnsonii DSM 18315] |
| Q8A1Y1 | BACOVA_04608 | 32.866 | 98 | 4.30E-120 | 390 | putative alpha-1,2-mannosidase [Bacteroides ovatus ATCC 8483] |
| Q8A1Y1 | BACDOR_04794 | 31.937 | 97 | 4.94E-120 | 381 | putative alpha-1,2-mannosidase [Bacteroides dorei DSM 17855] |
| Q8A1Y1 | HMPREF0105_2377 | 31.937 | 97 | 6.79E-120 | 381 | putative alpha-1,2-mannosidase [Bacteroides sp. 3_1_33FAA] |
| Q8A1Y1 | BSBG_01109 | 31.806 | 97 | 7.60E-119 | 378 | putative alpha-1,2-mannosidase [Bacteroides sp. 9_1_42FAA] |
| Q8A1Y1 | HMPREF0556_10574 | 32.513 | 96 | 1.51E-118 | 390 | putative alpha-1,2-mannosidase [Listeria grayi DSM 20601] |
| Q8A1Y1 | HMPREF1475_02052 | 33.208 | 99 | 1.69E-118 | 378 | hypothetical protein [Prevotella oralis HGA0225] |
| Q8A1Y1 | BACCELL_02612 | 31.759 | 96 | 7.38E-118 | 375 | putative alpha-1,2-mannosidase [Bacteroides cellulosilyticus DSM 14838] |
| Q8A1Y1 | BSGG_1589 | 32.549 | 96 | 1.18E-117 | 376 | putative alpha-1,2-mannosidase [Bacteroides sp. D2] |
| Q8A1Y1 | HMPREF1214_04117 | 32.337 | 93 | 1.74E-117 | 374 | hypothetical protein [Bacteroides sp. HPS0048] |
| Q8A1Y1 | HMPREF1532_00849 | 32.51 | 98 | 9.89E-117 | 373 | hypothetical protein [Bacteroides salyersiae WAL 10018 = DSM 18765 = JCM 12988] |
| Q8A1Y1 | HMPREF0673_00375 | 33.246 | 96 | 1.02E-116 | 372 | putative alpha-1,2-mannosidase [Prevotella stercorea DSM 18206] |
| Q8A1Y1 | HMPREF9449_01412 | 32.21 | 94 | 6.37E-116 | 371 | putative alpha-1,2-mannosidase [Odoribacter laneus YIT 12061] |
| Q8A1Y1 | HMPREF9420_2426 | 31.736 | 98 | 1.04E-115 | 370 | putative alpha-1,2-mannosidase [Prevotella salivae DSM 15606] |
| Q8A1Y1 | BACINT_04747 | 32.129 | 93 | 1.56E-115 | 370 | putative alpha-1,2-mannosidase [Bacteroides intestinalis DSM 17393] |
| Q8A1Y1 | BSHG_3694 | 35.556 | 76 | 2.82E-115 | 371 | putative alpha-1,2-mannosidase [Bacteroides sp. 3_2_5] |
| Q8A1Y1 | BACCELL_00110 | 31.995 | 93 | 8.53E-115 | 368 | putative alpha-1,2-mannosidase [Bacteroides cellulosilyticus DSM 14838] |
| Q8A1Y1 | HMPREF9448_02298 | 31.76 | 99 | 8.98E-115 | 368 | hypothetical protein [Barnesiella intestinihominis YIT 11860] |
| Q8A1Y1 | EUBVEN_00090 | 32.824 | 97 | 9.13E-115 | 387 | putative alpha-1,2-mannosidase [Eubacterium ventriosum ATCC 27560] |
| Q8A1Y1 | HMPREF1214_01526 | 31.73 | 98 | 1.26E-114 | 368 | hypothetical protein [Bacteroides sp. HPS0048] |
| Q8A1Y1 | HMPREF1475_02050 | 31.886 | 99 | 2.51E-114 | 368 | hypothetical protein [Prevotella oralis HGA0225] |
| Q8A1Y1 | CLOSPI_00329 | 33.29 | 95 | 2.83E-114 | 385 | putative alpha-1,2-mannosidase [Erysipelatoclostridium [Clostridium] spiroforme DSM 1552] |
| Q8A1Y1 | HMPREF9447_04444 | 32.262 | 93 | 3.06E-114 | 367 | hypothetical protein [Bacteroides oleiciplenus YIT 12058] |
| Q8A1Y1 | HMPREF0103_2021 | 31.701 | 96 | 4.08E-114 | 366 | putative alpha-1,2-mannosidase [Bacteroides sp. 2_1_33B] |
| Q8A1Y1 | BSIG_3766 | 31.484 | 99 | 6.98E-114 | 366 | putative alpha-1,2-mannosidase [Bacteroides sp. 1_1_6] |
| Q8A1Y1 | HMPREF9007_03635 | 31.443 | 99 | 1.06E-113 | 365 | alpha-1,2-mannosidase family protein [Bacteroides sp. 1_1_14] |
| Q8A1Y1 | HMPREF9455_00116 | 31.226 | 97 | 3.91E-111 | 358 | hypothetical protein [Dysgonomonas gadei ATCC BAA-286] |
| Q8A1Y1 | HMPREF1033_02448 | 31.378 | 98 | 7.28E-111 | 358 | hypothetical protein [Tannerella sp. 6_1_58FAA_CT1] |
| Q8A1Y1 | BSGG_1783 | 31.491 | 98 | 7.93E-111 | 358 | putative alpha-1,2-mannosidase [Bacteroides sp. D2] |

| | | | | | | |
|---|---|---|---|---|---|---|
| Q8A1Y1 | HMPREF1016_00475 | 32.129 | 93 | 9.99E-111 | 357 | glycosyl hydrolase family 92 [Bacteroides eggerthii 1_2_48FAA] |
| Q8A1Y1 | BXY_48360 | 31.491 | 98 | 1.16E-110 | 357 | alpha-1,2-mannosidase, putative [Bacteroides xylanisolvens XB1A] |
| Q8A1Y1 | CUY_0578 | 31.491 | 98 | 1.26E-110 | 357 | putative alpha-1,2-mannosidase [Bacteroides ovatus SD CMC 3f] |
| Q8A1Y1 | HMPREF9010_04156 | 31.362 | 98 | 3.77E-110 | 356 | alpha-1,2-mannosidase family protein [Bacteroides sp. 3_1_23] |
| Q8A1Y1 | BACEGG_02280 | 32.129 | 93 | 4.10E-110 | 356 | putative alpha-1,2-mannosidase [Bacteroides eggerthii DSM 20697] |
| Q8A1Y1 | BACOVA_03623 | 31.362 | 98 | 1.16E-109 | 355 | putative alpha-1,2-mannosidase [Bacteroides ovatus ATCC 8483] |
| Q8A1Y1 | HMPREF9456_02549 | 31.25 | 97 | 1.24E-109 | 355 | hypothetical protein [Dysgonomonas mossii DSM 22836] |
| Q8A1Y1 | HMPREF1017_02637 | 31.362 | 98 | 1.39E-109 | 355 | hypothetical protein [Bacteroides ovatus 3_8_47FAA] |
| Q8A1Y1 | HMPREF0127_04526 | 31.362 | 98 | 1.39E-109 | 355 | hypothetical protein [Bacteroides sp. 1_1_30] |
| Q8A1Y1 | HMPREF9441_00876 | 31.143 | 97 | 5.17E-109 | 353 | putative alpha-1,2-mannosidase [Paraprevotella clara YIT 11840] |
| Q8A1Y1 | BSCG_03594 | 31.234 | 98 | 1.60E-108 | 352 | putative alpha-1,2-mannosidase [Bacteroides sp. 2_2_4] |
| Q8A1Y1 | HMPREF0102_00043 | 31.234 | 98 | 1.60E-108 | 352 | putative alpha-1,2-mannosidase [Bacteroides sp. 2_1_22] |
| Q8A1Y1 | BSAG_02046 | 31.234 | 98 | 1.60E-108 | 352 | putative alpha-1,2-mannosidase [Bacteroides sp. D1] |
| Q8A1Y1 | CW3_1128 | 31.234 | 98 | 1.60E-108 | 352 | putative alpha-1,2-mannosidase [Bacteroides xylanisolvens SD CC 1b] |
| Q8A1Y1 | CW1_1388 | 31.234 | 98 | 1.60E-108 | 352 | putative alpha-1,2-mannosidase [Bacteroides xylanisolvens SD CC 2a] |
| Q8A1Y1 | HMPREF9442_01432 | 31.143 | 97 | 3.59E-108 | 350 | putative alpha-1,2-mannosidase [Paraprevotella xylaniphila YIT 11841] |
| Q8A1Y1 | HMPREF1033_01716 | 30.14 | 99 | 4.29E-108 | 350 | hypothetical protein [Tannerella sp. 6_1_58FAA_CT1] |
| Q8A1Y1 | HMPREF1532_00842 | 29.51 | 99 | 1.94E-107 | 348 | hypothetical protein [Bacteroides salyersiae WAL 10018 = DSM 18765 = JCM 12988] |
| Q8A1Y1 | HMPREF9441_01613 | 29.346 | 98 | 2.20E-106 | 347 | putative alpha-1,2-mannosidase [Paraprevotella clara YIT 11840] |
| Q8A1Y1 | HMPREF0102_03321 | 43.564 | 52 | 4.45E-106 | 334 | hypothetical protein [Bacteroides sp. 2_1_22] |
| Q8A1Y1 | HMPREF0103_2019 | 32.179 | 99 | 7.84E-106 | 344 | putative alpha-1,2-mannosidase [Bacteroides sp. 2_1_33B] |
| Q8A1Y1 | BACPLE_00638 | 29.685 | 93 | 8.87E-106 | 344 | putative alpha-1,2-mannosidase [Bacteroides plebeius DSM 17135] |
| Q8A1Y1 | BACUNI_04064 | 31.907 | 99 | 1.45E-105 | 345 | putative alpha-1,2-mannosidase [Bacteroides uniformis ATCC 8492] |
| Q8A1Y1 | HMPREF1007_00591 | 31.907 | 99 | 1.63E-105 | 345 | glycosyl hydrolase family 92 [Bacteroides sp. 4_1_36] |
| Q8A1Y1 | HMPREF0104_00272 | 32.051 | 99 | 2.74E-105 | 342 | alpha-1,2-mannosidase family protein [Bacteroides sp. 3_1_19] |
| Q8A1Y1 | HMPREF9008_03793 | 31.923 | 99 | 1.74E-104 | 340 | putative alpha-1,2-mannosidase [Parabacteroides sp. 20_3] |
| Q8A1Y1 | CW1_2594 | 69.608 | 27 | 2.48E-104 | 322 | conserved domain protein [Bacteroides xylanisolvens SD CC 2a] |
| Q8A1Y1 | HMPREF9456_02222 | 31.392 | 95 | 3.34E-104 | 341 | hypothetical protein [Dysgonomonas mossii DSM 22836] |
| Q8A1Y1 | HMPREF9442_00572 | 28.993 | 98 | 7.45E-104 | 340 | putative alpha-1,2-mannosidase [Paraprevotella xylaniphila YIT 11841] |
| Q8A1Y1 | HMPREF1017_00409 | 30.617 | 97 | 2.19E-103 | 338 | hypothetical protein [Bacteroides ovatus 3_8_47FAA] |
| Q8A1Y1 | HMPREF0673_01384 | 31.259 | 91 | 1.45E-102 | 333 | putative alpha-1,2-mannosidase [Prevotella stercorea DSM 18206] |
| Q8A1Y1 | HMPREF0619_03709 | 31.861 | 94 | 3.69E-102 | 335 | putative alpha-1,2-mannosidase [Parabacteroides sp. D13] |
| Q8A1Y1 | HMPREF9008_02102 | 31.861 | 94 | 6.16E-102 | 334 | putative alpha-1,2-mannosidase [Parabacteroides sp. 20_3] |
| Q8A1Y1 | HMPREF0104_03069 | 31.861 | 94 | 9.17E-102 | 334 | alpha-1,2-mannosidase family protein [Bacteroides sp. 3_1_19] |
| Q8A1Y1 | HMPREF0103_3265 | 31.861 | 94 | 9.46E-102 | 334 | putative alpha-1,2-mannosidase [Bacteroides sp. 2_1_33B] |
| Q8A1Y1 | BACINT_00091 | 30.653 | 96 | 3.30E-101 | 333 | putative alpha-1,2-mannosidase [Bacteroides intestinalis DSM 17393] |
| Q8A1Y1 | BACCELL_01131 | 30.137 | 96 | 1.00E-100 | 331 | putative alpha-1,2-mannosidase [Bacteroides cellulosilyticus DSM 14838] |
| Q8A1Y1 | HMPREF0619_01280 | 30.72 | 98 | 2.86E-100 | 330 | putative alpha-1,2-mannosidase [Parabacteroides sp. D13] |
| Q8A1Y1 | HMPREF0103_1288 | 30.591 | 98 | 4.26E-100 | 329 | putative alpha-1,2-mannosidase [Bacteroides sp. 2_1_33B] |
| Q8A1Y1 | HMPREF9008_03226 | 31.034 | 95 | 4.49E-100 | 329 | putative alpha-1,2-mannosidase [Parabacteroides sp. 20_3] |
| Q8A1Y1 | HMPREF0104_01338 | 30.902 | 95 | 1.62E-99 | 328 | alpha-1,2-mannosidase family protein [Bacteroides sp. 3_1_19] |
| Q8A1Y1 | HMPREF9008_03799 | 29.238 | 98 | 2.58E-99 | 328 | putative alpha-1,2-mannosidase [Parabacteroides sp. 20_3] |
| Q8A1Y1 | HMPREF0103_2010 | 29.449 | 94 | 3.11E-99 | 328 | putative alpha-1,2-mannosidase [Bacteroides sp. 2_1_33B] |
| Q8A1Y1 | HMPREF9007_01443 | 30.633 | 98 | 5.33E-99 | 327 | alpha-1,2-mannosidase family protein [Bacteroides sp. 1_1_14] |
| Q8A1Y1 | BSIG_2812 | 30.672 | 98 | 5.36E-99 | 327 | putative alpha-1,2-mannosidase [Bacteroides sp. 1_1_6] |
| Q8A1Y1 | PARMER_03122 | 30.68 | 97 | 1.09E-98 | 326 | putative alpha-1,2-mannosidase [Parabacteroides merdae ATCC 43184] |
| Q8A1Y1 | HMPREF9447_02878 | 29.68 | 97 | 1.24E-98 | 326 | hypothetical protein [Bacteroides oleiciplenus YIT 12058] |
| Q8A1Y1 | HMPREF0104_00279 | 29.028 | 94 | 1.05E-97 | 323 | alpha-1,2-mannosidase family protein [Bacteroides sp. 3_1_19] |
| Q8A1Y1 | HMPREF0619_00406 | 29.156 | 94 | 1.89E-97 | 323 | putative alpha-1,2-mannosidase [Parabacteroides sp. D13] |
| Q8A1Y1 | BSIG_1698 | 36.346 | 67 | 2.23E-97 | 326 | putative alpha-1,2-mannosidase [Bacteroides sp. 1_1_6] |
| Q8A1Y1 | BACPLE_00486 | 30.74 | 96 | 7.38E-97 | 320 | putative alpha-1,2-mannosidase [Bacteroides plebeius DSM 17135] |
| Q8A1Y1 | HMPREF9007_04278 | 32.376 | 96 | 9.44E-97 | 320 | alpha-1,2-mannosidase family protein [Bacteroides sp. 1_1_14] |
| Q8A1Y1 | PRABACTJOHN_04421 | 30.882 | 94 | 1.87E-96 | 320 | putative alpha-1,2-mannosidase [Parabacteroides johnsonii DSM 18315] |
| Q8A1Y1 | HMPREF9455_02174 | 30.337 | 99 | 5.96E-96 | 318 | hypothetical protein [Dysgonomonas gadei ATCC BAA-286] |
| Q8A1Y1 | BSGG_1519 | 29.747 | 94 | 9.77E-96 | 318 | putative alpha-1,2-mannosidase [Bacteroides sp. D2] |
| Q8A1Y1 | BSGG_4847 | 28.837 | 98 | 1.42E-95 | 318 | putative alpha-1,2-mannosidase [Bacteroides sp. D2] |
| Q8A1Y1 | BSIG_4336 | 31.95 | 99 | 1.68E-95 | 317 | putative alpha-1,2-mannosidase [Bacteroides sp. 1_1_6] |
| Q8A1Y1 | HMPREF9010_00933 | 30.462 | 96 | 1.71E-95 | 317 | alpha-1,2-mannosidase family protein [Bacteroides sp. 3_1_23] |
| Q8A1Y1 | HMPREF1211_01156 | 30.093 | 94 | 1.76E-95 | 318 | hypothetical protein [Streptomyces sp. HGB0020] |
| Q8A1Y1 | HMPREF1214_04622 | 30.098 | 98 | 2.37E-95 | 317 | hypothetical protein [Bacteroides sp. HPS0048] |

| | | | | | | |
|---|---|---|---|---|---|---|
| Q8A1Y1 | HMPREF9455_01365 | 31.579 | 95 | 2.61E-95 | 317 | hypothetical protein [Dysgonomonas gadei ATCC BAA-286] |
| Q8A1Y1 | BACFIN_06815 | 30.818 | 96 | 2.81E-95 | 317 | putative alpha-1,2-mannosidase [Bacteroides finegoldii DSM 17565] |
| Q8A1Y1 | HMPREF9007_02477 | 29.787 | 96 | 5.96E-95 | 316 | alpha-1,2-mannosidase family protein [Bacteroides sp. 1_1_14] |
| Q8A1Y1 | BACINT_00710 | 30.564 | 85 | 6.43E-95 | 314 | putative alpha-1,2-mannosidase [Bacteroides intestinalis DSM 17393] |
| Q8A1Y1 | BSGG_0162 | 30.262 | 96 | 1.07E-94 | 315 | putative alpha-1,2-mannosidase [Bacteroides sp. D2] |
| Q8A1Y1 | BSIG_0581 | 29.873 | 96 | 1.18E-94 | 315 | putative alpha-1,2-mannosidase [Bacteroides sp. 1_1_6] |
| Q8A1Y1 | HMPREF1033_02421 | 29.733 | 99 | 1.22E-94 | 315 | hypothetical protein [Tannerella sp. 6_1_58FAA_CT1] |
| Q8A1Y1 | BACCOP_04372 | 30.877 | 96 | 1.80E-94 | 314 | putative alpha-1,2-mannosidase [Bacteroides coprocola DSM 17136] |
| Q8A1Y1 | HMPREF1214_04005 | 30.38 | 96 | 2.43E-94 | 314 | hypothetical protein [Bacteroides sp. HPS0048] |
| Q8A1Y1 | HMPREF9447_01528 | 29.887 | 96 | 2.90E-94 | 314 | hypothetical protein [Bacteroides oleiciplenus YIT 12058] |
| Q8A1Y1 | BSHG_4425 | 30.303 | 96 | 3.42E-94 | 313 | putative alpha-1,2-mannosidase [Bacteroides sp. 3_2_5] |
| Q8A1Y1 | HMPREF1018_01841 | 30.303 | 96 | 3.42E-94 | 313 | hypothetical protein [Bacteroides sp. 2_1_56FAA] |
| Q8A1Y1 | HMPREF1033_01752 | 29.412 | 94 | 3.68E-94 | 314 | hypothetical protein [Tannerella sp. 6_1_58FAA_CT1] |
| Q8A0E4 | BSIG_2569 | 100 | 100 | 0 | 604 | hypothetical protein [Bacteroides sp. 1_1_6] |
| Q8A0E4 | PARMER_02286 | 45.763 | 99 | 3.83E-76 | 237 | putative esterase [Parabacteroides merdae ATCC 43184] |
| Q8A0E4 | AL1_14240 | 40.074 | 91 | 4.33E-66 | 211 | Enterochelin esterase and related enzymes [Alistipes shahii WAL 8301] |
| Q8A0E4 | PRABACTJOHN_00027 | 44.094 | 85 | 2.82E-60 | 195 | putative esterase [Parabacteroides johnsonii DSM 18315] |
| Q8A0E4 | BACUNI_00922 | 37.374 | 99 | 1.23E-58 | 192 | putative esterase [Bacteroides uniformis ATCC 8492] |
| Q8A0E4 | HMPREF1007_00163 | 37.374 | 99 | 1.88E-58 | 192 | hypothetical protein [Bacteroides sp. 4_1_36] |
| Q8A0E4 | HMPREF0969_01394 | 37.037 | 99 | 4.21E-58 | 191 | putative esterase [Bacteroides sp. D20] |
| Q8A0E4 | HMPREF9447_01465 | 35.548 | 99 | 2.77E-56 | 187 | hypothetical protein [Bacteroides oleiciplenus YIT 12058] |
| Q8A0E4 | BACINT_04212 | 35.135 | 99 | 1.48E-54 | 182 | putative esterase [Bacteroides intestinalis DSM 17393] |
| Q8A0E4 | BACCELL_03417 | 34.884 | 99 | 5.34E-54 | 181 | putative esterase [Bacteroides cellulosilyticus DSM 14838] |
| Q8A0E4 | CUY_0321 | 36.992 | 85 | 4.39E-53 | 177 | conserved hypothetical protein [Bacteroides ovatus SD CMC 3f] |
| Q8A0E4 | BSGG_1946 | 33.445 | 99 | 2.29E-51 | 174 | hypothetical protein [Bacteroides sp. D2] |
| Q8A0E4 | BACPLE_01969 | 36.12 | 99 | 5.82E-47 | 162 | putative esterase [Bacteroides plebeius DSM 17135] |
| Q8A0E4 | CUY_3721 | 33.445 | 99 | 1.13E-40 | 151 | conserved domain protein [Bacteroides ovatus SD CMC 3f] |
| Q8A0E4 | HMPREF0106_00538 | 33.11 | 99 | 5.07E-40 | 149 | esterase [Bacteroides sp. D22] |
| Q8A0E4 | BACOVA_02057 | 33.11 | 99 | 5.34E-40 | 149 | putative esterase [Bacteroides ovatus ATCC 8483] |
| Q8A0E4 | HMPREF0106_00539 | 34.328 | 88 | 6.61E-40 | 149 | endo-1,4-beta-xylanase Z [Bacteroides sp. D22] |
| Q8A0E4 | BACOVA_02056 | 33.708 | 88 | 8.11E-40 | 149 | putative esterase [Bacteroides ovatus ATCC 8483] |
| Q8A0E4 | CUY_3722 | 34.328 | 88 | 1.06E-39 | 149 | conserved domain protein [Bacteroides ovatus SD CMC 3f] |
| Q8A0E4 | HMPREF0127_02350 | 32.558 | 99 | 2.85E-39 | 147 | hypothetical protein [Bacteroides sp. 1_1_30] |
| Q8A0E4 | BXY_19590 | 32.558 | 99 | 3.19E-39 | 147 | Enterochelin esterase and related enzymes [Bacteroides xylanisolvens XB1A] |
| Q8A0E4 | HMPREF0127_02351 | 33.955 | 88 | 2.06E-38 | 145 | hypothetical protein [Bacteroides sp. 1_1_30] |
| Q8A0E4 | BSGG_2893 | 34.432 | 88 | 7.88E-38 | 144 | hypothetical protein [Bacteroides sp. D2] |
| Q8A0E4 | HMPREF0969_02050 | 34.47 | 85 | 6.48E-36 | 136 | isoamylase N-terminal domain protein [Bacteroides sp. D20] |
| Q8A0E4 | BACUNI_00269 | 34.47 | 85 | 6.48E-36 | 136 | isoamylase N-terminal domain protein [Bacteroides uniformis ATCC 8492] |
| Q8A0E4 | BACCOP_00729 | 34.118 | 84 | 6.33E-34 | 131 | putative esterase [Bacteroides coprocola DSM 17136] |
| Q8A0E4 | BACINT_01094 | 33.708 | 90 | 1.92E-31 | 127 | isoamylase N-terminal domain protein [Bacteroides intestinalis DSM 17393] |
| Q8A0E4 | BACCELL_02062 | 34.231 | 88 | 2.57E-31 | 127 | isoamylase N-terminal domain protein [Bacteroides cellulosilyticus DSM 14838] |
| Q8A0E4 | HMPREF9420_1580 | 31.203 | 88 | 6.30E-31 | 123 | isoamylase N-terminal domain protein [Prevotella salivae DSM 15606] |
| Q8A051 | BSIG_2479 | 100 | 100 | 0 | 1103 | parallel beta-helix [Bacteroides sp. 1_1_6] |
| Q8A051 | HMPREF9007_00401 | 71.429 | 99 | 0 | 810 | pectate lyase [Bacteroides sp. 1_1_14] |
| Q8A051 | BSAG_03172 | 50.094 | 99 | 8.58E-163 | 476 | hypothetical protein [Bacteroides sp. D1] |
| Q8A051 | CW3_4502 | 50.094 | 99 | 8.58E-163 | 476 | conserved hypothetical protein [Bacteroides xylanisolvens SD CC 1b] |
| Q8A051 | CW1_4251 | 50.094 | 99 | 8.58E-163 | 476 | conserved hypothetical protein [Bacteroides xylanisolvens SD CC 2a] |
| Q8A051 | BACOVA_05000 | 49.906 | 99 | 8.96E-163 | 476 | hypothetical protein [Bacteroides ovatus ATCC 8483] |
| Q8A051 | BSGG_1409 | 49.153 | 99 | 1.66E-158 | 465 | hypothetical protein [Bacteroides sp. D2] |
| Q8A051 | HMPREF9445_01478 | 47.637 | 96 | 4.44E-155 | 457 | hypothetical protein [Bacteroides clarus YIT 12056] |
| Q8A051 | CUU_3793 | 47.538 | 98 | 2.81E-153 | 452 | conserved hypothetical protein [Bacteroides vulgatus PC510] |
| Q8A051 | HMPREF0106_02588 | 51.153 | 88 | 1.86E-149 | 440 | pectate lyase [Bacteroides sp. D22] |
| Q8A051 | HMPREF0102_03405 | 51.153 | 88 | 9.40E-149 | 438 | hypothetical protein [Bacteroides sp. 2_1_22] |
| Q8A051 | BSIG_2465 | 51.454 | 83 | 2.97E-142 | 421 | hypothetical protein [Bacteroides sp. 1_1_6] |
| Q8A051 | HMPREF9007_00416 | 51.454 | 83 | 3.82E-141 | 418 | pectate lyase [Bacteroides sp. 1_1_14] |
| Q8A028 | BSIG_2455 | 100 | 100 | 0 | 1537 | hypothetical protein [Bacteroides sp. 1_1_6] |
| Q8A028 | HMPREF9007_00426 | 99.592 | 100 | 0 | 1531 | dipeptidyl-peptidase IV [Bacteroides sp. 1_1_14] |
| Q8A028 | HMPREF0127_04782 | 94.429 | 100 | 0 | 1427 | hypothetical protein [Bacteroides sp. 1_1_30] |
| Q8A028 | BXY_33000 | 94.429 | 100 | 0 | 1427 | dipeptidyl-peptidase IV . Serine peptidase. MEROPS family S09B [Bacteroides xylanisolvens XB1A] |
| Q8A028 | HMPREF0102_03488 | 94.293 | 100 | 0 | 1424 | peptidase, S9A/B/C family, catalytic domain protein [Bacteroides sp. 2_1_22] |

| | | | | | | |
|---|---|---|---|---|---|---|
| Q8A028 | BSAG_01729 | 94.293 | 100 | 0 | 1424 | peptidase, S9A/B/C family, catalytic domain protein [Bacteroides sp. D1] |
| Q8A028 | CW3_2404 | 94.293 | 100 | 0 | 1424 | dipeptidyl peptidase IV N-terminal domain protein [Bacteroides xylanisolvens SD CC 1b] |
| Q8A028 | CW1_1547 | 94.293 | 100 | 0 | 1424 | dipeptidyl peptidase IV N-terminal domain protein [Bacteroides xylanisolvens SD CC 2a] |
| Q8A028 | HMPREF0106_02634 | 94.293 | 100 | 0 | 1423 | dipeptidyl-peptidase IV [Bacteroides sp. D22] |
| Q8A028 | HMPREF1017_02246 | 94.158 | 100 | 0 | 1422 | hypothetical protein [Bacteroides ovatus 3_8_47FAA] |
| Q8A028 | HMPREF9010_02811 | 94.022 | 100 | 0 | 1421 | dipeptidyl-peptidase IV [Bacteroides sp. 3_1_23] |
| Q8A028 | CUY_0743 | 94.158 | 100 | 0 | 1420 | dipeptidyl peptidase IV N-terminal domain protein [Bacteroides ovatus SD CMC 3f] |
| Q8A028 | BSGG_1381 | 93.886 | 100 | 0 | 1419 | hypothetical protein [Bacteroides sp. D2] |
| Q8A028 | BACOVA_05046 | 94.993 | 98 | 0 | 1418 | peptidase, S9A/B/C family, catalytic domain protein [Bacteroides ovatus ATCC 8483] |
| Q8A028 | BACFIN_08852 | 93.342 | 100 | 0 | 1414 | peptidase, S9A/B/C family, catalytic domain protein [Bacteroides finegoldii DSM 17565] |
| Q8A028 | BSCG_01817 | 94.576 | 98 | 0 | 1413 | peptidase, S9A/B/C family, catalytic domain protein [Bacteroides sp. 2_2_4] |
| Q8A028 | BACCAC_01795 | 93.602 | 98 | 0 | 1405 | peptidase, S9A/B/C family, catalytic domain protein [Bacteroides caccae ATCC 43185] |
| Q8A028 | HMPREF1018_00999 | 80.707 | 100 | 0 | 1292 | hypothetical protein [Bacteroides sp. 2_1_56FAA] |
| Q8A028 | BFAG_00311 | 81.655 | 99 | 0 | 1292 | peptidase, S9A/B/C family, catalytic domain protein [Bacteroides fragilis 3_1_12] |
| Q8A028 | HMPREF0101_00028 | 80.571 | 100 | 0 | 1291 | peptidase, S9A/B/C family, catalytic domain protein [Bacteroides sp. 2_1_16] |
| Q8A028 | BSHG_0669 | 80.435 | 100 | 0 | 1289 | hypothetical protein [Bacteroides sp. 3_2_5] |
| Q8A028 | HMPREF1532_00929 | 82.609 | 100 | 0 | 1286 | hypothetical protein [Bacteroides salyersiae WAL 10018 = DSM 18765 = JCM 12988] |
| Q8A028 | HMPREF1214_01399 | 82.88 | 100 | 0 | 1280 | dipeptidyl-peptidase 4 [Bacteroides sp. HPS0048] |
| Q8A028 | HMPREF9446_02000 | 79.647 | 100 | 0 | 1251 | peptidase, S9A/B/C family, catalytic domain protein [Bacteroides fluxus YIT 12057] |
| Q8A028 | BACCELL_03148 | 78.426 | 100 | 0 | 1238 | peptidase, S9A/B/C family, catalytic domain protein [Bacteroides cellulosilyticus DSM 14838] |
| Q8A028 | BACINT_04434 | 78.562 | 100 | 0 | 1229 | peptidase, S9A/B/C family, catalytic domain protein [Bacteroides intestinalis DSM 17393] |
| Q8A028 | BACUNI_03587 | 78.155 | 100 | 0 | 1215 | peptidase, S9A/B/C family, catalytic domain protein [Bacteroides uniformis ATCC 8492] |
| Q8A028 | HMPREF1007_03240 | 78.155 | 100 | 0 | 1215 | dipeptidyl peptidase IV [Bacteroides sp. 4_1_36] |
| Q8A028 | HMPREF0969_00958 | 78.155 | 100 | 0 | 1214 | peptidase, S9A/B/C family, catalytic domain protein [Bacteroides sp. D20] |
| Q8A028 | HMPREF9447_04168 | 78.155 | 100 | 0 | 1213 | hypothetical protein [Bacteroides oleiciplenus YIT 12058] |
| Q8A028 | HMPREF9445_02996 | 76.934 | 100 | 0 | 1211 | peptidase, S9A/B/C family, catalytic domain protein [Bacteroides clarus YIT 12056] |
| Q8A028 | BACSTE_03271 | 76.398 | 99 | 0 | 1207 | peptidase, S9A/B/C family, catalytic domain protein [Bacteroides stercoris ATCC 43183] |
| Q8A028 | BACEGG_00881 | 76.391 | 100 | 0 | 1201 | peptidase, S9A/B/C family, catalytic domain protein [Bacteroides eggerthii DSM 20697] |
| Q8A028 | HMPREF1016_01518 | 76.255 | 100 | 0 | 1198 | dipeptidyl peptidase IV [Bacteroides eggerthii 1_2_48FAA] |
| Q8A028 | BACCOP_03926 | 64.905 | 99 | 0 | 1022 | peptidase, S9A/B/C family, catalytic domain protein [Bacteroides coprocola DSM 17136] |
| Q8A028 | BSFG_01970 | 63.93 | 100 | 0 | 1014 | peptidase, S9A/B/C family, catalytic domain protein [Bacteroides sp. 4_3_47FAA] |
| Q8A028 | HMPREF9011_04151 | 63.93 | 100 | 0 | 1014 | dipeptidyl peptidase IV [Bacteroides sp. 3_1_40A] |
| Q8A028 | BSEG_03560 | 63.624 | 100 | 0 | 1011 | peptidase, S9A/B/C family, catalytic domain protein [Bacteroides dorei 5_1_36/D4] |
| Q8A028 | BACDOR_03064 | 63.624 | 100 | 0 | 1011 | peptidase, S9A/B/C family, catalytic domain protein [Bacteroides dorei DSM 17855] |
| Q8A028 | BSBG_03825 | 63.624 | 100 | 0 | 1011 | peptidase, S9A/B/C family, catalytic domain protein [Bacteroides sp. 9_1_42FAA] |
| Q8A028 | CUU_1533 | 63.661 | 100 | 0 | 1009 | putative prolyl dipeptidyl peptidase [Bacteroides vulgatus PC510] |
| Q8A028 | HMPREF0105_4156 | 63.49 | 100 | 0 | 1009 | peptidase, S9A/B/C family, catalytic domain protein [Bacteroides sp. 3_1_33FAA] |
| Q8A028 | BACCOPRO_03615 | 64.363 | 99 | 0 | 991 | peptidase, S9A/B/C family, catalytic domain protein [Bacteroides coprophilus DSM 18228 = JCM 13818] |
| Q8A028 | BACPLE_03112 | 62.584 | 100 | 0 | 959 | peptidase, S9A/B/C family, catalytic domain protein [Bacteroides plebeius DSM 17135] |
| Q8A028 | HMPREF9442_00881 | 58.008 | 100 | 0 | 886 | peptidase, S9A/B/C family, catalytic domain protein [Paraprevotella xylaniphila YIT 11841] |
| Q8A028 | HMPREF9441_02325 | 57.604 | 100 | 0 | 883 | peptidase, S9A/B/C family, catalytic domain protein [Paraprevotella clara YIT 11840] |
| Q8A028 | PARMER_03777 | 52.901 | 100 | 0 | 803 | peptidase, S9A/B/C family, catalytic domain protein [Parabacteroides merdae ATCC 43184] |
| Q8A028 | PRABACTJOHN_02440 | 52.497 | 100 | 0 | 799 | peptidase, S9A/B/C family, catalytic domain protein [Parabacteroides johnsonii DSM 18315] |
| Q8A028 | HMPREF9420_0548 | 51.413 | 100 | 0 | 794 | peptidase, S9A/B/C family, catalytic domain protein [Prevotella salivae DSM 15606] |
| Q8A028 | HMPREF9008_02510 | 52.092 | 100 | 0 | 785 | peptidase, S9A/B/C family, catalytic domain protein [Parabacteroides sp. 20_3] |
| Q8A028 | HMPREF0104_02101 | 52.092 | 100 | 0 | 785 | dipeptidyl-peptidase IV [Bacteroides sp. 3_1_19] |
| Q8A028 | HMPREF0103_2768 | 52.092 | 100 | 0 | 783 | peptidase, S9A/B/C family, catalytic domain protein [Bacteroides sp. 2_1_33B] |
| Q8A028 | HMPREF0619_02972 | 51.957 | 100 | 0 | 783 | peptidase, S9A/B/C family, catalytic domain protein [Parabacteroides sp. D13] |
| Q8A028 | HMPREF9448_00433 | 50.136 | 100 | 0 | 776 | hypothetical protein [Barnesiella intestinihominis YIT 11860] |
| Q8A028 | HMPREF1475_00504 | 51.796 | 97 | 0 | 769 | hypothetical protein [Prevotella oralis HGA0225] |
| Q8A028 | HMPREF0673_01202 | 50.067 | 100 | 0 | 766 | peptidase, S9A/B/C family, catalytic domain protein [Prevotella stercorea DSM 18206] |
| Q8A028 | HMPREF1033_00581 | 50.407 | 100 | 0 | 748 | hypothetical protein [Tannerella sp. 6_1_58FAA_CT1] |
| Q8A028 | PREVCOP_04755 | 49.185 | 99 | 0 | 728 | peptidase, S9A/B/C family, catalytic domain protein [Prevotella copri DSM 18205] |
| Q8A028 | HMPREF9456_01479 | 46.883 | 100 | 0 | 721 | hypothetical protein [Dysgonomonas mossii DSM 22836] |
| Q8A028 | HMPREF9455_01995 | 46.944 | 98 | 0 | 701 | hypothetical protein [Dysgonomonas gadei ATCC BAA-286] |
| Q8A028 | HMPREF9449_00338 | 47.684 | 99 | 0 | 693 | hypothetical protein [Odoribacter laneus YIT 12061] |
| Q8A028 | HMPREF9450_01296 | 43.333 | 99 | 0 | 590 | hypothetical protein [Alistipes indistinctus YIT 12060] |
| Q8A028 | ALIPUT_02419 | 38.827 | 96 | 9.26E-159 | 480 | peptidase, S9A/B/C family, catalytic domain protein [Alistipes putredinis DSM 17216] |
| Q8A028 | BACDOR_00958 | 32.94 | 99 | 1.45E-115 | 369 | peptidase, S9A/B/C family, catalytic domain protein [Bacteroides dorei DSM 17855] |
| Q8A028 | HMPREF0105_1254 | 32.983 | 99 | 2.20E-115 | 369 | peptidase, S9A/B/C family, catalytic domain protein [Bacteroides sp. 3_1_33FAA] |

| | | | | | | |
|---|---|---|---|---|---|---|
| Q8A028 | BSEG_00993 | 32.94 | 99 | 2.79E-115 | 368 | peptidase, S9A/B/C family, catalytic domain protein [Bacteroides dorei 5_1_36/D4] |
| Q8A028 | BSBG_00102 | 32.808 | 99 | 1.22E-114 | 367 | peptidase, S9A/B/C family, catalytic domain protein [Bacteroides sp. 9_1_42FAA] |
| Q8A028 | BSFG_00985 | 32.546 | 99 | 1.41E-114 | 366 | peptidase, S9A/B/C family, catalytic domain protein [Bacteroides sp. 4_3_47FAA] |
| Q8A028 | CUU_2370 | 32.546 | 99 | 1.60E-114 | 366 | dipeptidyl peptidase IV N-terminal domain protein [Bacteroides vulgatus PC510] |
| Q8A028 | PRABACTJOHN_03838 | 33.512 | 98 | 2.31E-112 | 361 | peptidase, S9A/B/C family, catalytic domain protein [Parabacteroides johnsonii DSM 18315] |
| Q8A028 | HMPREF9008_02521 | 31.932 | 99 | 1.70E-111 | 358 | peptidase, S9A/B/C family, catalytic domain protein [Parabacteroides sp. 20_3] |
| Q8A028 | PARMER_01283 | 33.469 | 97 | 1.98E-110 | 356 | peptidase, S9A/B/C family, catalytic domain protein [Parabacteroides merdae ATCC 43184] |
| Q8A028 | HMPREF0619_02963 | 31.537 | 99 | 3.52E-110 | 355 | peptidase, S9A/B/C family, catalytic domain protein [Parabacteroides sp. D13] |
| Q8A028 | HMPREF0103_2777 | 31.669 | 99 | 5.37E-110 | 354 | peptidase, S9A/B/C family, catalytic domain protein [Bacteroides sp. 2_1_33B] |
| Q8A028 | HMPREF0104_02092 | 31.537 | 99 | 7.69E-110 | 354 | dipeptidyl peptidase IV [Bacteroides sp. 3_1_19] |
| Q8A028 | HMPREF9011_03920 | 35.058 | 79 | 1.74E-104 | 335 | dipeptidyl aminopeptidase [Bacteroides sp. 3_1_40A] |
| Q8A028 | HMPREF9455_02807 | 32.353 | 99 | 2.63E-101 | 331 | hypothetical protein [Dysgonomonas gadei ATCC BAA-286] |
| Q8A028 | HMPREF9456_00374 | 34.423 | 78 | 5.58E-98 | 322 | hypothetical protein [Dysgonomonas mossii DSM 22836] |
| Q8A028 | HMPREF1033_01873 | 30 | 99 | 1.01E-96 | 319 | hypothetical protein [Tannerella sp. 6_1_58FAA_CT1] |
| Q8A028 | HMPREF1475_02283 | 33.221 | 80 | 1.61E-94 | 312 | hypothetical protein [Prevotella oralis HGA0225] |
| Q8A028 | BACPLE_01539 | 30.9 | 100 | 1.08E-93 | 311 | peptidase, S9A/B/C family, catalytic domain protein [Bacteroides plebeius DSM 17135] |
| Q8A028 | PARMER_02747 | 30.092 | 99 | 8.04E-93 | 309 | peptidase, S9A/B/C family, catalytic domain protein [Parabacteroides merdae ATCC 43184] |
| Q8A028 | BACCOPRO_02325 | 30.749 | 99 | 1.03E-91 | 305 | peptidase, S9A/B/C family, catalytic domain protein [Bacteroides coprophilus DSM 18228 = JCM 13818] |
| Q8A028 | PRABACTJOHN_00695 | 30.223 | 99 | 2.22E-90 | 302 | peptidase, S9A/B/C family, catalytic domain protein [Parabacteroides johnsonii DSM 18315] |
| Q8A028 | HMPREF9420_0906 | 28.892 | 100 | 9.31E-90 | 300 | peptidase, S9A/B/C family, catalytic domain protein [Prevotella salivae DSM 15606] |
| Q8A028 | HMPREF9449_01075 | 30.936 | 98 | 2.40E-89 | 299 | hypothetical protein [Odoribacter laneus YIT 12061] |
| Q8A028 | HMPREF0673_01428 | 31.848 | 79 | 1.68E-88 | 297 | peptidase, S9A/B/C family, catalytic domain protein [Prevotella stercorea DSM 18206] |
| Q8A028 | BFAG_02901 | 31.07 | 79 | 1.53E-87 | 294 | peptidase, S9A/B/C family, catalytic domain protein [Bacteroides fragilis 3_1_12] |
| Q8A028 | HMPREF9441_01209 | 29.517 | 99 | 1.72E-87 | 295 | peptidase, S9A/B/C family, catalytic domain protein [Paraprevotella clara YIT 11840] |
| Q8A028 | HMPREF9442_02116 | 29.262 | 100 | 5.54E-87 | 293 | peptidase, S9A/B/C family, catalytic domain protein [Paraprevotella xylaniphila YIT 11841] |
| Q8A028 | PREVCOP_05027 | 31.624 | 76 | 6.35E-87 | 294 | peptidase, S9A/B/C family, catalytic domain protein [Prevotella copri DSM 18205] |
| Q8A028 | HMPREF9445_01935 | 31.122 | 79 | 9.10E-87 | 293 | peptidase, S9A/B/C family, catalytic domain protein [Bacteroides clarus YIT 12056] |
| Q8A028 | BACCELL_02178 | 32.605 | 73 | 1.86E-86 | 291 | peptidase, S9A/B/C family, catalytic domain protein [Bacteroides cellulosilyticus DSM 14838] |
| Q8A028 | BACSTE_02313 | 30.86 | 79 | 2.62E-86 | 291 | peptidase, S9A/B/C family, catalytic domain protein [Bacteroides stercoris ATCC 43183] |
| Q8A028 | HMPREF0101_02096 | 32.477 | 73 | 1.75E-84 | 286 | peptidase, S9A/B/C family, catalytic domain protein [Bacteroides sp. 2_1_16] |
| Q8A028 | HMPREF1018_00139 | 32.477 | 73 | 2.15E-84 | 286 | hypothetical protein [Bacteroides sp. 2_1_56FAA] |
| Q8A028 | HMPREF9447_02557 | 31.81 | 75 | 3.72E-84 | 286 | hypothetical protein [Bacteroides oleiciplenus YIT 12058] |
| Q8A028 | BSHG_1495 | 32.477 | 73 | 4.16E-84 | 285 | hypothetical protein [Bacteroides sp. 3_2_5] |
| Q8A028 | HMPREF1016_00991 | 31.145 | 79 | 5.65E-83 | 282 | dipeptidyl peptidase IV [Bacteroides eggerthii 1_2_48FAA] |
| Q8A028 | BACEGG_03258 | 30.457 | 79 | 2.35E-82 | 281 | peptidase, S9A/B/C family, catalytic domain protein [Bacteroides eggerthii DSM 20697] |
| Q8A028 | BACINT_01011 | 32.065 | 73 | 2.54E-82 | 281 | peptidase, S9A/B/C family, catalytic domain protein [Bacteroides intestinalis DSM 17393] |
| Q8A028 | HMPREF0969_01715 | 31.652 | 76 | 8.31E-82 | 280 | peptidase, S9A/B/C family, catalytic domain protein [Bacteroides sp. D20] |
| Q8A028 | BACUNI_04166 | 31.652 | 76 | 1.17E-81 | 279 | peptidase, S9A/B/C family, catalytic domain protein [Bacteroides uniformis ATCC 8492] |
| Q8A028 | HMPREF1007_00511 | 31.652 | 76 | 1.17E-81 | 279 | dipeptidyl peptidase IV [Bacteroides sp. 4_1_36] |
| Q8A028 | BACPLE_00583 | 29.15 | 99 | 2.17E-81 | 278 | peptidase, S9A/B/C family, catalytic domain protein [Bacteroides plebeius DSM 17135] |
| Q8A028 | BACCOP_02363 | 32.686 | 73 | 9.26E-81 | 277 | peptidase, S9A/B/C family, catalytic domain protein [Bacteroides coprocola DSM 17136] |
| Q8A028 | HMPREF1532_01808 | 31.103 | 73 | 1.74E-80 | 275 | hypothetical protein [Bacteroides salyersiae WAL 10018 = DSM 18765 = JCM 12988] |
| Q8A028 | HMPREF9446_02242 | 30.796 | 73 | 6.71E-80 | 275 | peptidase, S9A/B/C family, catalytic domain protein [Bacteroides fluxus YIT 12057] |
| Q8A028 | BXY_23820 | 31.404 | 73 | 1.11E-79 | 273 | prolyl tripeptidyl peptidase. Serine peptidase. MEROPS family S09B [Bacteroides xylanisolvens XB1A] |
| Q8A028 | HMPREF0127_01576 | 31.404 | 73 | 1.16E-79 | 273 | hypothetical protein [Bacteroides sp. 1_1_30] |
| Q8A028 | HMPREF0106_04714 | 31.404 | 73 | 1.16E-79 | 273 | dipeptidyl aminopeptidase IV [Bacteroides sp. D22] |
| Q8A028 | HMPREF0102_00604 | 31.404 | 73 | 1.27E-79 | 273 | peptidase, S9A/B/C family, catalytic domain protein [Bacteroides sp. 2_1_22] |
| Q8A028 | BSAG_00356 | 31.404 | 73 | 1.27E-79 | 273 | peptidase, S9A/B/C family, catalytic domain protein [Bacteroides sp. D1] |
| Q8A028 | CW3_2034 | 31.404 | 73 | 1.27E-79 | 273 | dipeptidyl peptidase IV N-terminal domain protein [Bacteroides xylanisolvens SD CC 1b] |
| Q8A028 | CW1_0738 | 31.404 | 73 | 1.27E-79 | 273 | dipeptidyl peptidase IV N-terminal domain protein [Bacteroides xylanisolvens SD CC 2a] |
| Q8A028 | BACOVA_01349 | 30.706 | 79 | 3.37E-79 | 272 | peptidase, S9A/B/C family, catalytic domain protein [Bacteroides ovatus ATCC 8483] |
| Q8A028 | HMPREF1017_00530 | 30.428 | 79 | 8.40E-79 | 271 | hypothetical protein [Bacteroides ovatus 3_8_47FAA] |
| Q8A028 | HMPREF9010_00648 | 30.542 | 79 | 1.95E-78 | 270 | putative dipeptidyl aminopeptidase IV [Bacteroides sp. 3_1_23] |
| Q8A028 | BACCAC_01453 | 30.81 | 73 | 2.79E-78 | 270 | peptidase, S9A/B/C family, catalytic domain protein [Bacteroides caccae ATCC 43185] |
| Q8A028 | CUY_3404 | 30.769 | 79 | 3.15E-78 | 270 | dipeptidyl peptidase IV N-terminal domain protein [Bacteroides ovatus SD CMC 3f] |
| Q8A028 | BSEG_02165 | 27.621 | 100 | 6.04E-78 | 269 | peptidase, S9A/B/C family, catalytic domain protein [Bacteroides dorei 5_1_36/D4] |
| Q8A028 | CUU_0154 | 27.949 | 100 | 8.22E-78 | 269 | dipeptidyl peptidase IV N-terminal domain protein [Bacteroides vulgatus PC510] |
| Q8A028 | BSGG_2370 | 30.606 | 79 | 1.13E-77 | 268 | hypothetical protein [Bacteroides sp. D2] |
| Q8A028 | BACFIN_08470 | 30.81 | 73 | 1.23E-77 | 268 | peptidase, S9A/B/C family, catalytic domain protein [Bacteroides finegoldii DSM 17565] |
| Q8A028 | BSFG_00357 | 27.635 | 100 | 1.40E-77 | 268 | peptidase, S9A/B/C family, catalytic domain protein [Bacteroides sp. 4_3_47FAA] |

| | | | | | | |
|---|---|---|---|---|---|---|
| Q8A028 | HMPREF9011_00439 | 27.635 | 100 | 1.40E-77 | 268 | dipeptidyl peptidase IV [Bacteroides sp. 3_1_40A] |
| Q8A028 | HMPREF0105_2037 | 27.821 | 100 | 7.80E-77 | 266 | peptidase, S9A/B/C family, catalytic domain protein [Bacteroides sp. 3_1_33FAA] |
| Q8A028 | BACDOR_01902 | 27.564 | 100 | 2.19E-76 | 265 | peptidase, S9A/B/C family, catalytic domain protein [Bacteroides dorei DSM 17855] |
| Q8A028 | BSBG_00781 | 27.564 | 100 | 2.19E-76 | 265 | peptidase, S9A/B/C family, catalytic domain protein [Bacteroides sp. 9_1_42FAA] |
| Q8A028 | HMPREF1214_00573 | 29.611 | 79 | 3.74E-76 | 263 | dipeptidyl-peptidase 4 [Bacteroides sp. HPS0048] |
| Q8A028 | HMPREF0104_00231 | 27.953 | 99 | 1.00E-74 | 260 | dipeptidyl aminopeptidase IV [Bacteroides sp. 3_1_19] |
| Q8A028 | HMPREF9008_03751 | 28.571 | 99 | 1.52E-74 | 259 | peptidase, S9A/B/C family, catalytic domain protein [Parabacteroides sp. 20_3] |
| Q8A028 | HMPREF9007_03794 | 30.656 | 79 | 2.72E-74 | 259 | dipeptidyl aminopeptidase IV [Bacteroides sp. 1_1_14] |
| Q8A028 | BSIG_1417 | 30.492 | 79 | 7.84E-74 | 258 | hypothetical protein [Bacteroides sp. 1_1_6] |
| Q8A028 | HMPREF0619_00460 | 28.442 | 99 | 1.84E-73 | 257 | peptidase, S9A/B/C family, catalytic domain protein [Parabacteroides sp. D13] |
| Q8A028 | HMPREF0103_2069 | 27.69 | 99 | 3.22E-73 | 256 | peptidase, S9A/B/C family, catalytic domain protein [Bacteroides sp. 2_1_33B] |
| Q8A028 | BSCG_03097 | 35.501 | 49 | 3.34E-66 | 228 | peptidase, S9A/B/C family, catalytic domain protein [Bacteroides sp. 2_2_4] |
| Q8A028 | HMPREF9007_04016 | 28.176 | 81 | 3.33E-44 | 174 | Xaa-Pro dipeptidyl-peptidase [Bacteroides sp. 1_1_14] |
| Q8A028 | BSIG_1145 | 28.176 | 81 | 3.90E-44 | 174 | hypothetical protein [Bacteroides sp. 1_1_6] |
| Q8A028 | BSCG_03701 | 26.81 | 84 | 4.48E-43 | 171 | peptidase, S9A/B/C family, catalytic domain protein [Bacteroides sp. 2_2_4] |
| Q8A028 | BACINT_00527 | 24.874 | 99 | 9.73E-42 | 167 | peptidase, S9A/B/C family, catalytic domain protein [Bacteroides intestinalis DSM 17393] |
| Q8A028 | HMPREF9449_02863 | 26.187 | 78 | 1.21E-41 | 167 | hypothetical protein [Odoribacter laneus YIT 12061] |
| Q8A028 | BACCOPRO_01823 | 25.118 | 79 | 1.33E-40 | 164 | peptidase, S9A/B/C family, catalytic domain protein [Bacteroides coprophilus DSM 18228 = JCM 13818] |
| Q8A028 | BACCOP_03698 | 29.602 | 52 | 2.77E-40 | 163 | peptidase, S9A/B/C family, catalytic domain protein [Bacteroides coprocola DSM 17136] |
| Q8A028 | HMPREF9441_02554 | 25.78 | 79 | 4.62E-40 | 162 | peptidase, S9A/B/C family, catalytic domain protein [Paraprevotella clara YIT 11840] |
| Q8A028 | HMPREF0106_04740 | 26.935 | 79 | 4.78E-40 | 162 | Xaa-Pro dipeptidyl-peptidase [Bacteroides sp. D22] |
| Q8A028 | HMPREF0127_01607 | 26.935 | 79 | 4.96E-40 | 162 | hypothetical protein [Bacteroides sp. 1_1_30] |
| Q8A028 | BACCAC_01475 | 26.206 | 79 | 6.56E-40 | 162 | peptidase, S9A/B/C family, catalytic domain protein [Bacteroides caccae ATCC 43185] |
| Q8A028 | HMPREF9455_01475 | 23.307 | 99 | 2.47E-39 | 160 | hypothetical protein [Dysgonomonas gadei ATCC BAA-286] |
| Q8A028 | BSCG_03068 | 26.731 | 79 | 9.97E-39 | 158 | peptidase, S9A/B/C family, catalytic domain protein [Bacteroides sp. 2_2_4] |
| Q8A028 | BACPLE_03825 | 25.736 | 79 | 1.21E-38 | 158 | peptidase, S9A/B/C family, catalytic domain protein [Bacteroides plebeius DSM 17135] |
| Q8A028 | HMPREF9010_00743 | 26.731 | 79 | 1.26E-38 | 158 | Xaa-Pro dipeptidyl-peptidase [Bacteroides sp. 3_1_23] |
| Q8A028 | CUY_4412 | 26.452 | 79 | 1.52E-38 | 157 | peptidase, S9A/B/C familie, catalytic domain protein [Bacteroides ovatus SD CMC 3f] |
| Q8A028 | PARMER_02220 | 24.883 | 79 | 2.70E-38 | 157 | peptidase, S9A/B/C family, catalytic domain protein [Parabacteroides merdae ATCC 43184] |
| Q8A028 | HMPREF1486_02696 | 24.519 | 80 | 2.80E-38 | 156 | hypothetical protein [Streptomyces sp. HPH0547] |
| Q8A028 | BACCELL_01959 | 31.429 | 50 | 8.73E-38 | 155 | peptidase, S9A/B/C family, catalytic domain protein [Bacteroides cellulosilyticus DSM 14838] |
| Q8A028 | HMPREF0105_2009 | 31.008 | 50 | 1.52E-37 | 154 | peptidase, S9A/B/C family, catalytic domain protein [Bacteroides sp. 3_1_33FAA] |
| Q8A028 | BSBG_00751 | 31.008 | 50 | 1.84E-37 | 154 | peptidase, S9A/B/C family, catalytic domain protein [Bacteroides sp. 9_1_42FAA] |
| Q8A028 | BACDOR_01931 | 31.008 | 50 | 3.39E-37 | 153 | peptidase, S9A/B/C family, catalytic domain protein [Bacteroides dorei DSM 17855] |
| Q8A028 | BSEG_02195 | 31.266 | 50 | 5.39E-37 | 152 | peptidase, S9A/B/C family, catalytic domain protein [Bacteroides dorei 5_1_36/D4] |
| Q8A028 | CUU_0128 | 31.266 | 50 | 6.06E-37 | 152 | peptidase, S9A/B/C familie, catalytic domain protein [Bacteroides vulgatus PC510] |
| Q8A028 | BSFG_00330 | 31.266 | 50 | 6.58E-37 | 152 | peptidase, S9A/B/C family, catalytic domain protein [Bacteroides sp. 4_3_47FAA] |
| Q8A028 | HMPREF9011_00413 | 31.266 | 50 | 6.58E-37 | 152 | peptidase [Bacteroides sp. 3_1_40A] |
| Q8A028 | HMPREF0104_00820 | 30.73 | 51 | 6.62E-37 | 153 | Xaa-Pro dipeptidyl-peptidase [Bacteroides sp. 3_1_19] |
| Q8A028 | HMPREF0103_1483 | 30.73 | 51 | 6.80E-37 | 153 | peptidase, S9A/B/C family, catalytic domain protein [Bacteroides sp. 2_1_33B] |
| Q8A028 | HMPREF0619_00930 | 30.479 | 51 | 6.93E-37 | 152 | peptidase, S9A/B/C family, catalytic domain protein [Parabacteroides sp. D13] |
| Q8A028 | HMPREF9008_01562 | 30.73 | 51 | 6.99E-37 | 152 | peptidase, S9A/B/C family, catalytic domain protein [Parabacteroides sp. 20_3] |
| Q8A028 | BXY_24090 | 25.04 | 79 | 1.23E-34 | 145 | Dipeptidyl aminopeptidases/acylaminoacyl-peptidases [Bacteroides xylanisolvens XB1A] |
| Q8A028 | BSAG_00328 | 25.04 | 79 | 1.23E-34 | 145 | peptidase, S9A/B/C family, catalytic domain protein [Bacteroides sp. D1] |
| Q8A028 | CW3_2060 | 25.04 | 79 | 1.23E-34 | 145 | peptidase, S9A/B/C familie, catalytic domain protein [Bacteroides xylanisolvens SD CC 1b] |
| Q8A028 | CW1_0762 | 25.04 | 79 | 1.23E-34 | 145 | peptidase, S9A/B/C familie, catalytic domain protein [Bacteroides xylanisolvens SD CC 2a] |
| Q8A028 | HMPREF0102_00578 | 25.04 | 79 | 1.77E-34 | 145 | peptidase, S9A/B/C family, catalytic domain protein [Bacteroides sp. 2_1_22] |
| Q8A028 | HMPREF0102_00577 | 24 | 97 | 8.55E-33 | 140 | peptidase, S9A/B/C family, catalytic domain protein [Bacteroides sp. 2_1_22] |
| Q8A028 | BSAG_00327 | 24 | 97 | 8.55E-33 | 140 | peptidase, S9A/B/C family, catalytic domain protein [Bacteroides sp. D1] |
| Q8A028 | CW3_2061 | 24.252 | 80 | 1.96E-31 | 135 | peptidase, S9A/B/C familie, catalytic domain protein [Bacteroides xylanisolvens SD CC 1b] |
| Q8A028 | CW1_0763 | 24.252 | 80 | 1.96E-31 | 135 | peptidase, S9A/B/C familie, catalytic domain protein [Bacteroides xylanisolvens SD CC 2a] |
| Q8A028 | BACCAC_01476 | 24.71 | 77 | 8.59E-31 | 134 | peptidase, S9A/B/C family, catalytic domain protein [Bacteroides caccae ATCC 43185] |
| Q8A007 | BSIG_2433 | 100 | 100 | 0 | 614 | hypothetical protein [Bacteroides sp. 1_1_6] |
| Q8A007 | HMPREF9007_00446 | 99.655 | 100 | 0 | 612 | conserved hypothetical protein [Bacteroides sp. 1_1_14] |
| Q8A007 | HMPREF0106_02651 | 92.414 | 100 | 0 | 576 | conserved hypothetical protein [Bacteroides sp. D22] |
| Q8A007 | HMPREF0127_04765 | 92.069 | 100 | 0 | 575 | hypothetical protein [Bacteroides sp. 1_1_30] |
| Q8A007 | BXY_33180 | 92.069 | 100 | 0 | 575 | Hydrolases of the alpha/beta superfamily [Bacteroides xylanisolvens XB1A] |
| Q8A007 | BACCAC_01815 | 92.414 | 100 | 0 | 574 | hypothetical protein [Bacteroides caccae ATCC 43185] |
| Q8A007 | BACFIN_08840 | 91.379 | 100 | 0 | 571 | hypothetical protein [Bacteroides finegoldii DSM 17565] |
| Q8A007 | HMPREF0102_03501 | 92.069 | 100 | 0 | 571 | hypothetical protein [Bacteroides sp. 2_1_22] |

| | | | | | | |
|---|---|---|---|---|---|---|
| Q8A007 | CW3_2390 | 92.069 | 100 | 0 | 571 | putative lipoprotein [Bacteroides xylanisolvens SD CC 1b] |
| Q8A007 | BSAG_01744 | 92.069 | 100 | 0 | 571 | hypothetical protein [Bacteroides sp. D1] |
| Q8A007 | CW1_1533 | 92.069 | 100 | 0 | 571 | conserved hypothetical protein [Bacteroides xylanisolvens SD CC 2a] |
| Q8A007 | BSCG_01833 | 92.069 | 100 | 0 | 569 | hypothetical protein [Bacteroides sp. 2_2_4] |
| Q8A007 | BSGG_1362 | 92.069 | 100 | 0 | 569 | hypothetical protein [Bacteroides sp. D2] |
| Q8A007 | BACOVA_05064 | 92.069 | 100 | 0 | 568 | hypothetical protein [Bacteroides ovatus ATCC 8483] |
| Q8A007 | CUY_0760 | 92.069 | 100 | 0 | 568 | conserved hypothetical protein [Bacteroides ovatus SD CMC 3f] |
| Q8A007 | HMPREF9010_02828 | 92.069 | 100 | 0 | 568 | conserved hypothetical protein [Bacteroides sp. 3_1_23] |
| Q8A007 | HMPREF1017_02263 | 91.724 | 100 | 0 | 565 | hypothetical protein [Bacteroides ovatus 3_8_47FAA] |
| Q8A007 | HMPREF1214_01419 | 83.391 | 99 | 0 | 524 | hypothetical protein [Bacteroides sp. HPS0048] |
| Q8A007 | HMPREF1532_00895 | 82.699 | 99 | 0 | 519 | hypothetical protein [Bacteroides salyersiae WAL 10018 = DSM 18765 = JCM 12988] |
| Q8A007 | BSHG_0730 | 81.724 | 100 | 0 | 516 | hypothetical protein [Bacteroides sp. 3_2_5] |
| Q8A007 | HMPREF1018_00932 | 81.724 | 100 | 0 | 516 | hypothetical protein [Bacteroides sp. 2_1_56FAA] |
| Q8A007 | HMPREF0101_02816 | 81.724 | 100 | 0 | 516 | hypothetical protein [Bacteroides sp. 2_1_16] |
| Q8A007 | BFAG_00247 | 81.724 | 100 | 0 | 516 | hypothetical protein [Bacteroides fragilis 3_1_12] |
| Q8A007 | HMPREF9447_04111 | 74.138 | 100 | 1.84E-164 | 462 | hypothetical protein [Bacteroides oleiciplenus YIT 12058] |
| Q8A007 | HMPREF9445_02570 | 73.448 | 100 | 7.23E-164 | 461 | hypothetical protein [Bacteroides clarus YIT 12056] |
| Q8A007 | BACCELL_03210 | 73.103 | 100 | 3.35E-163 | 459 | hypothetical protein [Bacteroides cellulosilyticus DSM 14838] |
| Q8A007 | BACINT_04375 | 73.103 | 100 | 7.96E-163 | 458 | hypothetical protein [Bacteroides intestinalis DSM 17393] |
| Q8A007 | BACEGG_00916 | 73.793 | 100 | 2.20E-161 | 455 | hypothetical protein [Bacteroides eggerthii DSM 20697] |
| Q8A007 | HMPREF1016_01554 | 73.793 | 100 | 2.59E-161 | 454 | hypothetical protein [Bacteroides eggerthii 1_2_48FAA] |
| Q8A007 | HMPREF9449_01038 | 68.75 | 99 | 3.49E-153 | 434 | hypothetical protein [Odoribacter laneus YIT 12061] |
| Q8A007 | HMPREF9011_02830 | 68.276 | 100 | 6.84E-149 | 423 | hypothetical protein [Bacteroides sp. 3_1_40A] |
| Q8A007 | BSFG_02458 | 68.276 | 100 | 1.13E-148 | 422 | hypothetical protein [Bacteroides sp. 4_3_47FAA] |
| Q8A007 | CUU_1898 | 68.276 | 100 | 2.70E-148 | 421 | conserved hypothetical protein [Bacteroides vulgatus PC510] |
| Q8A007 | HMPREF0105_0517 | 67.586 | 100 | 4.10E-147 | 419 | hypothetical protein [Bacteroides sp. 3_1_33FAA] |
| Q8A007 | HMPREF9446_03811 | 65.472 | 100 | 6.73E-147 | 419 | hypothetical protein [Bacteroides fluxus YIT 12057] |
| Q8A007 | BSEG_00635 | 67.586 | 100 | 1.67E-146 | 417 | hypothetical protein [Bacteroides dorei 5_1_36/D4] |
| Q8A007 | BSBG_01732 | 67.586 | 100 | 1.67E-146 | 417 | hypothetical protein [Bacteroides sp. 9_1_42FAA] |
| Q8A007 | BACDOR_03993 | 67.241 | 100 | 5.32E-146 | 416 | hypothetical protein [Bacteroides dorei DSM 17855] |
| Q8A007 | HMPREF0969_00882 | 65.033 | 100 | 5.42E-145 | 414 | hypothetical protein [Bacteroides sp. D20] |
| Q8A007 | BACUNI_03469 | 65.033 | 100 | 5.42E-145 | 414 | hypothetical protein [Bacteroides uniformis ATCC 8492] |
| Q8A007 | HMPREF1007_03165 | 65.033 | 100 | 5.42E-145 | 414 | hypothetical protein [Bacteroides sp. 4_1_36] |
| Q8A007 | BACSTE_03605 | 63.141 | 99 | 1.11E-141 | 406 | hypothetical protein [Bacteroides stercoris ATCC 43183] |
| Q8A007 | BACCOP_02508 | 65.862 | 99 | 4.29E-140 | 401 | hypothetical protein [Bacteroides coprocola DSM 17136] |
| Q8A007 | BACCOPRO_01519 | 64.931 | 99 | 3.73E-136 | 391 | hypothetical protein [Bacteroides coprophilus DSM 18228 = JCM 13818] |
| Q8A007 | BACPLE_02662 | 61.246 | 99 | 5.44E-131 | 378 | hypothetical protein [Bacteroides plebeius DSM 17135] |
| Q8A007 | HMPREF0673_00965 | 62.454 | 91 | 3.29E-126 | 366 | hypothetical protein [Prevotella stercorea DSM 18206] |
| Q8A007 | HMPREF9442_00480 | 63.004 | 93 | 2.38E-125 | 364 | hypothetical protein [Paraprevotella xylaniphila YIT 11841] |
| Q8A007 | HMPREF9441_02529 | 62.637 | 93 | 5.35E-125 | 363 | hypothetical protein [Paraprevotella clara YIT 11840] |
| Q8A007 | PREVCOP_06636 | 54.027 | 99 | 1.09E-107 | 319 | Gram-positive signal peptide protein, YSIRK family [Prevotella copri DSM 18205] |
| Q8A007 | HMPREF1475_01369 | 51.027 | 99 | 1.07E-101 | 303 | hypothetical protein [Prevotella oralis HGA0225] |
| Q8A007 | HMPREF9412_0311 | 48.148 | 84 | 2.41E-80 | 250 | X-Pro dipeptidyl-peptidase (S15 family) [Paenibacillus sp. HGF5] |
| Q8A007 | HMPREF0496_0813 | 49.789 | 81 | 2.44E-75 | 237 | hypothetical protein [Lactobacillus brevis subsp. gravesensis ATCC 27305] |
| Q8A007 | HMPREF0497_0752 | 49.367 | 81 | 1.74E-74 | 234 | hypothetical protein [Lactobacillus buchneri ATCC 11577] |
| Q8A007 | HMPREF1207_04301 | 42.692 | 90 | 4.64E-74 | 233 | hypothetical protein [Paenibacillus sp. HGH0039] |
| Q8A007 | HMPREF9413_4970 | 42.692 | 90 | 4.64E-74 | 233 | hypothetical protein [Paenibacillus sp. HGF7] |
| Q8A007 | HMPREF0519_1547 | 49.367 | 81 | 4.83E-74 | 233 | hypothetical protein [Lactobacillus hilgardii ATCC 8290] |
| Q8A007 | HMPREF0322_00614 | 45.868 | 83 | 1.64E-71 | 228 | hypothetical protein [Desulfitobacterium hafniense DP7] |
| Q8A007 | HMPREF0216_00140 | 44.309 | 85 | 1.73E-70 | 224 | hypothetical protein [Clostridium celatum DSM 1785] |
| Q8A007 | CUO_1816 | 40.493 | 98 | 3.34E-70 | 223 | X-Pro dipeptidyl-peptidase (S15 family) [Enterococcus faecium PC4.1] |
| Q8A007 | HMPREF0352_1292 | 40.493 | 98 | 3.34E-70 | 223 | X-Pro dipeptidyl-peptidase (S15 family) [Enterococcus faecium TX1330] |
| Q8A007 | HMPREF1207_01002 | 42.857 | 84 | 1.37E-69 | 222 | hypothetical protein [Paenibacillus sp. HGH0039] |
| Q8A007 | HMPREF9413_2149 | 42.857 | 84 | 1.37E-69 | 222 | hypothetical protein [Paenibacillus sp. HGF7] |
| Q8A007 | HMPREF9488_01897 | 43.701 | 88 | 2.50E-69 | 221 | peptidase S15 [Coprobacillus sp. 29_1] |
| Q8A007 | HMPREF0531_12659 | 46.414 | 81 | 2.56E-68 | 219 | X-Pro dipeptidyl-peptidase (S15 family) [Lactobacillus plantarum subsp. plantarum ATCC 14917 = JCM 1149 =] |
| Q8A007 | CSBG_02209 | 42.636 | 89 | 5.27E-68 | 218 | hypothetical protein [Clostridium sp. 7_2_43FAA] |
| Q8A007 | HMPREF9024_00971 | 44.304 | 81 | 5.04E-67 | 215 | hypothetical protein [Pediococcus acidilactici 7_4] |
| Q8A007 | HMPREF0557_00531 | 42.194 | 81 | 9.27E-67 | 214 | hypothetical protein [Listeria innocua ATCC 33091] |
| Q8A007 | HOLDEFILI_02222 | 43.952 | 85 | 1.12E-66 | 214 | hypothetical protein [Holdemania filiformis DSM 12042] |

| | | | | | | |
|---|---|---|---|---|---|---|
| Q8A007 | BRYFOR_09865 | 44.13 | 85 | 2.68E-66 | 213 | hypothetical protein [Marvinbryantia formatexigens DSM 14469] |
| Q8A007 | HMPREF0623_1758 | 43.882 | 81 | 3.27E-66 | 213 | hypothetical protein [Pediococcus acidilactici DSM 20284] |
| Q8A007 | HMPREF0348_1710 | 42.975 | 83 | 1.11E-65 | 211 | hypothetical protein [Enterococcus faecalis TX0104] |
| Q8A007 | HMPREF9494_00778 | 42.975 | 83 | 1.11E-65 | 211 | hypothetical protein [Enterococcus faecalis TX2137] |
| Q8A007 | HMPREF0349_1065 | 42.975 | 83 | 1.11E-65 | 211 | hypothetical protein [Enterococcus faecalis TX1322] |
| Q8A007 | CUI_2528 | 42.975 | 83 | 1.11E-65 | 211 | conserved hypothetical protein [Enterococcus faecalis PC1.1] |
| Q8A007 | HMPREF9520_02713 | 42.975 | 83 | 1.11E-65 | 211 | hypothetical protein [Enterococcus faecalis TX1467] |
| Q8A007 | HMPREF9517_00511 | 42.975 | 83 | 1.11E-65 | 211 | hypothetical protein [Enterococcus faecalis TX1341] |
| Q8A007 | HMPREF9516_02291 | 42.975 | 83 | 1.11E-65 | 211 | hypothetical protein [Enterococcus faecalis TX1302] |
| Q8A007 | HMPREF9519_02770 | 42.975 | 83 | 1.11E-65 | 211 | hypothetical protein [Enterococcus faecalis TX1346] |
| Q8A007 | HMPREF9518_02394 | 42.975 | 83 | 1.11E-65 | 211 | hypothetical protein [Enterococcus faecalis TX1342] |
| Q8A007 | HMPREF9497_01466 | 42.975 | 83 | 1.11E-65 | 211 | hypothetical protein [Enterococcus faecalis TX4244] |
| Q8A007 | HMPREF9521_00599 | 42.975 | 83 | 1.11E-65 | 211 | hypothetical protein [Enterococcus faecalis TX2134] |
| Q8A007 | HMPREF0216_01993 | 43.089 | 85 | 2.60E-65 | 211 | hypothetical protein [Clostridium celatum DSM 1785] |
| Q8A007 | STRINF_00059 | 41.224 | 84 | 7.31E-65 | 209 | hypothetical protein [Streptococcus infantarius subsp. infantarius ATCC BAA-102] |
| Q8A007 | HMPREF9478_02915 | 42.437 | 82 | 6.36E-63 | 204 | hypothetical protein [Enterococcus saccharolyticus 30_1] |
| Q8A007 | HMPREF0534_2027 | 45.378 | 81 | 1.90E-62 | 204 | hypothetical protein [Lactobacillus reuteri CF48-3A] |
| Q8A007 | HMPREF0538_20842 | 45.378 | 81 | 1.90E-62 | 204 | hypothetical protein [Lactobacillus reuteri SD2112] |
| Q8A007 | CLOBAR_02672 | 43.75 | 83 | 2.00E-62 | 203 | hypothetical protein [Intestinibacter bartlettii DSM 16795] |
| Q8A007 | LAR_1555 | 44.958 | 81 | 2.63E-62 | 202 | conserved hypothetical protein [Lactobacillus reuteri JCM 1112] |
| Q8A007 | HMPREF0535_0951 | 44.958 | 81 | 2.74E-62 | 202 | hypothetical protein [Lactobacillus reuteri MM2-3] |
| Q8A007 | HMPREF0536_11656 | 44.958 | 81 | 2.74E-62 | 202 | hypothetical protein [Lactobacillus reuteri MM4-1A] |
| Q8A007 | LRHM_0720 | 44.681 | 79 | 3.19E-62 | 202 | putative cell surface hydrolase [Lactobacillus rhamnosus GG] |
| Q8A007 | HMPREF0539_3026 | 44.681 | 79 | 3.52E-62 | 202 | hypothetical protein [Lactobacillus rhamnosus LMS2-1] |
| Q8A007 | LAF_1712 | 45.188 | 81 | 3.61E-62 | 202 | conserved hypothetical protein [Lactobacillus fermentum IFO 3956] |
| Q8A007 | HMPREF5505_1635 | 44.915 | 80 | 4.25E-62 | 202 | alpha/beta hydrolase [Lactobacillus delbrueckii subsp. lactis DSM 20072] |
| Q8A007 | HMPREF0511_1392 | 45.188 | 81 | 4.39E-62 | 203 | hypothetical protein [Lactobacillus fermentum ATCC 14931] |
| Q8A007 | HMPREF0541_02297 | 44.255 | 79 | 2.80E-61 | 200 | hypothetical protein [Lactobacillus rhamnosus ATCC 21052] |
| Q8A007 | HMPREF0993_02154 | 42.51 | 84 | 4.01E-61 | 200 | hypothetical protein [Lachnospiraceae bacterium 5_1_57FAA] |
| Q8A007 | CLOSCI_03946 | 42.51 | 84 | 5.97E-61 | 199 | hypothetical protein [Lachnospiraceae [Clostridium] scindens ATCC 35704] |
| Q8A007 | HMPREF0819_1696 | 39.592 | 84 | 1.02E-60 | 200 | hypothetical protein [Streptococcus equinus ATCC 9812] |
| Q8A007 | HMPREF9469_04155 | 38.462 | 85 | 2.09E-60 | 198 | hypothetical protein [Lachnospiraceae [Clostridium] citroniae WAL-17108] |
| Q8A007 | HMPREF0531_12949 | 38.75 | 83 | 1.18E-59 | 196 | hypothetical protein [Lactobacillus plantarum subsp. plantarum ATCC 14917 = JCM 1149 =] |
| Q8A007 | HMPREF0983_00605 | 42.742 | 85 | 1.21E-59 | 196 | X-Pro dipeptidyl-peptidase (S15 family) [Erysipelotrichaceae bacterium 3_1_53] |
| Q8A007 | CBFG_02158 | 39.271 | 85 | 1.95E-59 | 196 | hypothetical protein [Clostridiales bacterium 1_7_47FAA] |
| Q8A007 | HMPREF1032_03670 | 40.65 | 85 | 5.08E-59 | 195 | hypothetical protein [Subdoligranulum sp. 4_3_54A2FAA] |
| Q8A007 | HMPREF1014_05769 | 44.206 | 79 | 5.12E-59 | 194 | hypothetical protein [Bacillus sp. 7_6_55CFAA_CT2] |
| Q8A007 | HMPREF0548_0665 | 43.443 | 83 | 5.71E-59 | 194 | hypothetical protein [Lactobacillus ultunensis DSM 16047] |
| Q8A007 | HMPREF0542_10204 | 41.975 | 81 | 6.06E-59 | 194 | X-Pro dipeptidyl-peptidase (S15 family) [Lactobacillus ruminis ATCC 25644] |
| Q8A007 | CLOBOL_02399 | 35.798 | 89 | 6.42E-59 | 194 | hypothetical protein [Lachnospiraceae [Clostridium] bolteae ATCC BAA-613] |
| Q8A007 | HMPREF0981_00338 | 41.935 | 85 | 2.11E-58 | 193 | hypothetical protein [Erysipelotrichaceae bacterium 6_1_45] |
| Q8A007 | HMPREF0982_02141 | 41.935 | 85 | 2.11E-58 | 193 | hypothetical protein [Erysipelotrichaceae bacterium 21_3] |
| Q8A007 | HMPREF9406_4244 | 41.935 | 85 | 2.37E-58 | 193 | X-Pro dipeptidyl-peptidase (S15 family) [Clostridium sp. HGF2] |
| Q8A007 | HMPREF9022_01872 | 41.532 | 85 | 3.21E-58 | 192 | hypothetical protein [Erysipelotrichaceae bacterium 2_2_44A] |
| Q8A007 | HMPREF0519_1188 | 38.849 | 95 | 3.81E-58 | 192 | hypothetical protein [Lactobacillus hilgardii ATCC 8290] |
| Q8A007 | HMPREF0530_1739 | 41.339 | 84 | 4.54E-58 | 192 | hypothetical protein [Lactobacillus paracasei subsp. paracasei ATCC 25302] |
| Q8A007 | HMPREF1027_00215 | 39.231 | 89 | 7.80E-58 | 191 | hypothetical protein [Lactobacillus sp. 7_1_47FAA] |
| Q8A007 | HMPREF9467_01811 | 35.271 | 89 | 1.02E-57 | 191 | hypothetical protein [Lachnospiraceae [Clostridium] clostridioforme 2_1_49FAA] |
| Q8A007 | CATMIT_02777 | 40 | 89 | 1.05E-57 | 191 | hypothetical protein [Catenibacterium mitsuokai DSM 15897] |
| Q8A007 | HMPREF0555_1440 | 41.102 | 81 | 1.33E-57 | 191 | hypothetical protein [Leuconostoc mesenteroides subsp. cremoris ATCC 19254] |
| Q8A007 | HMPREF0216_03282 | 39.113 | 85 | 1.61E-57 | 191 | hypothetical protein [Clostridium celatum DSM 1785] |
| Q8A007 | HMPREF0496_0420 | 38.489 | 95 | 2.04E-57 | 190 | hypothetical protein [Lactobacillus brevis subsp. gravesensis ATCC 27305] |
| Q8A007 | HMPREF9488_01857 | 43.852 | 84 | 2.18E-57 | 191 | hypothetical protein [Coprobacillus sp. 29_1] |
| Q8A007 | HMPREF0530_0825 | 41.961 | 87 | 3.24E-57 | 189 | hypothetical protein [Lactobacillus paracasei subsp. paracasei ATCC 25302] |
| Q8A007 | HMPREF9473_03758 | 36.538 | 90 | 5.80E-57 | 189 | hypothetical protein [Lachnospiraceae [Clostridium] hathewayi WAL-18680] |
| Q8A007 | LAF_0029 | 38.129 | 96 | 9.32E-57 | 189 | cell surface hydrolase [Lactobacillus fermentum IFO 3956] |
| Q8A007 | HMPREF0492_0976 | 41.85 | 77 | 1.01E-56 | 188 | hypothetical protein [Lactobacillus acidophilus ATCC 4796] |
| Q8A007 | HMPREF0511_0269 | 36.426 | 100 | 1.13E-56 | 188 | hypothetical protein [Lactobacillus fermentum ATCC 14931] |
| Q8A007 | HMPREF0497_0386 | 38.489 | 95 | 1.56E-56 | 188 | hypothetical protein [Lactobacillus buchneri ATCC 11577] |
| Q8A007 | HMPREF9476_00593 | 38.824 | 87 | 4.72E-56 | 187 | hypothetical protein [Clostridium perfringens WAL-14572] |

| | | | | | | |
|---|---|---|---|---|---|---|
| Q8A007 | HMPREF0518_1269 | 42.324 | 81 | 7.29E-56 | 187 | hypothetical protein [Lactobacillus helveticus DSM 20075] |
| Q8A007 | CATMIT_02406 | 38.832 | 98 | 3.29E-55 | 185 | hypothetical protein [Catenibacterium mitsuokai DSM 15897] |
| Q8A007 | HMPREF0493_1523 | 41.597 | 81 | 6.64E-55 | 184 | hypothetical protein [Lactobacillus amylolyticus DSM 11664] |
| Q8A007 | HMPREF0492_1071 | 41.597 | 81 | 2.81E-54 | 182 | hypothetical protein [Lactobacillus acidophilus ATCC 4796] |
| Q8A007 | HMPREF0548_0756 | 40.506 | 81 | 4.22E-54 | 182 | hypothetical protein [Lactobacillus ultunensis DSM 16047] |
| Q8A007 | HMPREF1021_00790 | 38.306 | 84 | 8.21E-54 | 181 | hypothetical protein [Coprobacillus sp. 3_3_56FAA] |
| Q8A007 | MBAG_02414 | 38.306 | 84 | 8.21E-54 | 181 | hypothetical protein [Coprobacillus sp. D7] |
| Q8A007 | HMPREF0978_01643 | 38.306 | 84 | 8.21E-54 | 181 | hypothetical protein [Coprobacillus sp. 8_2_54BFAA] |
| Q8A007 | HMPREF0494_0603 | 39.916 | 81 | 9.29E-54 | 182 | hypothetical protein [Lactobacillus antri DSM 16041] |
| Q8A007 | CLORAM_00168 | 38.306 | 84 | 1.08E-53 | 181 | hypothetical protein [Erysipelatoclostridium ramosum DSM 1402] |
| Q8A007 | COLAER_01820 | 40.323 | 85 | 1.12E-53 | 181 | hypothetical protein [Collinsella aerofaciens ATCC 25986] |
| Q8A007 | LRHM_2799 | 40.784 | 87 | 1.37E-52 | 177 | putative cell surface hydrolase [Lactobacillus rhamnosus GG] |
| Q8A007 | HMPREF0541_01782 | 40.784 | 87 | 1.37E-52 | 177 | hypothetical protein [Lactobacillus rhamnosus ATCC 21052] |
| Q8A007 | CLOBAR_01854 | 36.694 | 85 | 1.47E-52 | 176 | hypothetical protein [Intestinibacter bartlettii DSM 16795] |
| Q8A007 | CLOBOL_03982 | 41.736 | 83 | 1.55E-52 | 178 | hypothetical protein [Lachnospiraceae [Clostridium] bolteae ATCC BAA-613] |
| Q8A007 | HMPREF0531_11293 | 38.889 | 86 | 6.51E-52 | 176 | hypothetical protein [Lactobacillus plantarum subsp. plantarum ATCC 14917 = JCM 1149 =] |
| Q8A007 | HMPREF0539_0410 | 41.772 | 81 | 9.93E-52 | 173 | hypothetical protein [Lactobacillus rhamnosus LMS2-1] |
| Q8A007 | ERE_20900 | 37.786 | 89 | 1.31E-51 | 176 | Hydrolases of the alpha/beta superfamily [Eubacterium rectale M104/1] |
| Q8A007 | CLOSTASPAR_05412 | 36.596 | 81 | 1.60E-51 | 173 | hypothetical protein [Lachnospiraceae [Clostridium asparagiforme] DSM 15981] |
| Q8A007 | HMPREF0866_00816 | 36.823 | 93 | 1.83E-51 | 175 | hypothetical protein [Ruminococcaceae bacterium D16] |
| Q8A007 | EUR_20570 | 37.405 | 89 | 7.74E-51 | 173 | Hydrolases of the alpha/beta superfamily [Eubacterium rectale DSM 17629] |
| Q8A007 | CLOSS21_03160 | 40 | 83 | 9.11E-51 | 173 | hypothetical protein [Clostridium sp. SS2/1] |
| Q8A007 | CL2_25140 | 40 | 83 | 9.11E-51 | 173 | Prolyl oligopeptidase family. [Clostridiales butyrate-producing bacterium SSC/2] |
| Q8A007 | HMPREF0877_0477 | 39.167 | 82 | 1.92E-50 | 172 | hypothetical protein [Weissella paramesenteroides ATCC 33313] |
| Q8A007 | HMPREF0994_07023 | 36.017 | 81 | 4.70E-50 | 172 | hypothetical protein [Lachnospiraceae bacterium 3_1_57FAA_CT1] |
| Q8A007 | HMPREF0996_01212 | 39.592 | 83 | 4.79E-50 | 171 | hypothetical protein [Lachnospiraceae bacterium 5_1_63FAA] |
| Q8A007 | MITSMUL_03008 | 38.589 | 82 | 3.11E-49 | 169 | X-Pro dipeptidyl-peptidase (S15 family) [Mitsuokella multacida DSM 20544] |
| Q8A007 | HMPREF0623_1395 | 34.979 | 83 | 1.13E-48 | 168 | X-Pro dipeptidyl-peptidase (S15 family) [Pediococcus acidilactici DSM 20284] |
| Q8A007 | HMPREF0369_02016 | 39.184 | 83 | 3.70E-48 | 166 | hypothetical protein [Anaerostipes hadrus DSM 3319] |
| Q8A007 | HMPREF9024_00008 | 34.979 | 83 | 6.83E-48 | 166 | hypothetical protein [Pediococcus acidilactici 7_4] |
| Q8A007 | CBFG_06129 | 38.532 | 75 | 3.89E-47 | 164 | hypothetical protein [Clostridiales bacterium 1_7_47FAA] |
| Q8A007 | HMPREF9469_05716 | 38.8 | 85 | 4.86E-47 | 163 | hypothetical protein [Lachnospiraceae [Clostridium] citroniae WAL-17108] |
| Q8A007 | HOLDEFILI_02011 | 34.363 | 89 | 5.21E-47 | 164 | hypothetical protein [Holdemania filiformis DSM 12042] |
| Q8A007 | HMPREF9494_00812 | 41.395 | 74 | 1.13E-46 | 163 | hypothetical protein [Enterococcus faecalis TX2137] |
| Q8A007 | ENT_08910 | 41.395 | 74 | 1.38E-46 | 163 | hypothetical protein [Enterococcus sp. 7L76] |
| Q8A007 | HMPREF0348_1681 | 41.395 | 74 | 1.47E-46 | 162 | hypothetical protein [Enterococcus faecalis TX0104] |
| Q8A007 | HMPREF9521_00632 | 41.395 | 74 | 1.95E-46 | 162 | hypothetical protein [Enterococcus faecalis TX2134] |
| Q8A007 | HMPREF0531_10501 | 36.052 | 80 | 2.20E-46 | 162 | hypothetical protein [Lactobacillus plantarum subsp. plantarum ATCC 14917 = JCM 1149 =] |
| Q8A007 | HMPREF9517_02618 | 41.395 | 74 | 2.36E-46 | 162 | hypothetical protein [Enterococcus faecalis TX1341] |
| Q8A007 | HMPREF9497_01439 | 41.395 | 74 | 2.36E-46 | 162 | hypothetical protein [Enterococcus faecalis TX4244] |
| Q8A007 | HMPREF0349_1100 | 41.395 | 74 | 2.87E-46 | 162 | hypothetical protein [Enterococcus faecalis TX1322] |
| Q8A007 | HMPREF9516_02258 | 41.395 | 74 | 2.89E-46 | 162 | hypothetical protein [Enterococcus faecalis TX1302] |
| Q8A007 | HMPREF9518_02359 | 40.93 | 74 | 9.02E-46 | 160 | hypothetical protein [Enterococcus faecalis TX1342] |
| Q8A007 | CUI_1761 | 40.93 | 74 | 1.15E-45 | 160 | conserved hypothetical protein [Enterococcus faecalis PC1.1] |
| Q8A007 | HMPREF9520_02753 | 40.93 | 74 | 1.25E-45 | 160 | hypothetical protein [Enterococcus faecalis TX1467] |
| Q8A007 | HMPREF9519_02803 | 40.93 | 74 | 1.84E-45 | 160 | hypothetical protein [Enterococcus faecalis TX1346] |
| Q8A007 | HMPREF9488_00936 | 37.975 | 81 | 1.12E-43 | 154 | hypothetical protein [Coprobacillus sp. 29_1] |
| Q8A007 | HMPREF0982_01652 | 31.923 | 89 | 4.17E-43 | 154 | hypothetical protein [Erysipelotrichaceae bacterium 21_3] |
| Q8A007 | HMPREF9022_04556 | 30.537 | 99 | 1.49E-42 | 152 | hypothetical protein [Erysipelotrichaceae bacterium 2_2_44A] |
| Q8A007 | HMPREF0981_03271 | 30.201 | 99 | 2.98E-42 | 151 | hypothetical protein [Erysipelotrichaceae bacterium 6_1_45] |
| Q8A007 | HMPREF9406_0018 | 30.201 | 99 | 2.98E-42 | 151 | hypothetical protein [Clostridium sp. HGF2] |
| Q8A007 | HMPREF0983_04056 | 36.598 | 67 | 7.99E-41 | 147 | hypothetical protein [Erysipelotrichaceae bacterium 3_1_53] |
| Q8A007 | BBNG_00773 | 33.21 | 88 | 1.48E-40 | 147 | hypothetical protein [Bifidobacterium bifidum NCIMB 41171] |
| Q8A007 | CUO_0104 | 35.294 | 82 | 2.51E-40 | 146 | conserved hypothetical protein [Enterococcus faecium PC4.1] |
| Q8A007 | HMPREF0240_01110 | 32.636 | 82 | 5.86E-40 | 145 | alpha/beta hydrolase [Clostridium sp. D5] |
| Q8A007 | HMPREF0352_1999 | 34.874 | 82 | 2.22E-39 | 144 | hypothetical protein [Enterococcus faecium TX1330] |
| Q8A007 | SUBVAR_07146 | 36.681 | 78 | 5.61E-39 | 143 | hypothetical protein [Subdoligranulum variabile DSM 15176] |
| Q8A007 | HMPREF1014_00754 | 32.489 | 82 | 2.49E-38 | 140 | hypothetical protein [Bacillus sp. 7_6_55CFAA_CT2] |
| Q8A007 | HMPREF9412_1741 | 32.569 | 75 | 8.67E-37 | 138 | X-Pro dipeptidyl-peptidase (S15 family) [Paenibacillus sp. HGF5] |
| Q8A007 | HMPREF1315_1065 | 31.818 | 88 | 1.14E-34 | 132 | alpha/beta hydrolase family protein [Bifidobacterium longum subsp. longum 2-2B] |

| | | | | | | |
|---|---|---|---|---|---|---|
| Q8A007 | HMPREF0175_0327 | 31.439 | 88 | 3.14E-34 | 131 | hypothetical protein [Bifidobacterium longum subsp. longum ATCC 55813] |
| Q8A007 | BIL_10720 | 31.439 | 88 | 3.14E-34 | 131 | hypothetical protein [Bifidobacterium longum subsp. longum F8] |
| Q8A007 | BLLJ_0767 | 31.439 | 88 | 3.14E-34 | 131 | conserved hypothetical protein [Bifidobacterium longum subsp. longum JCM 1217] |
| Q8A007 | BLIF_0801 | 31.439 | 88 | 3.45E-34 | 130 | conserved hypothetical protein [Bifidobacterium longum subsp. infantis 157F] |
| Q8A007 | BIFDEN_02140 | 32.661 | 83 | 8.47E-34 | 128 | hypothetical protein [Bifidobacterium dentium ATCC 27678] |
| Q8A007 | HMPREF1312_2283 | 31.061 | 88 | 4.26E-33 | 128 | alpha/beta hydrolase family protein [Bifidobacterium longum subsp. longum 44B] |
| Q8A007 | BLIG_01056 | 31.061 | 88 | 4.26E-33 | 128 | hypothetical protein [Bifidobacterium longum subsp. infantis CCUG 52486] |
| Q8A007 | HMPREF0177_02024 | 31.061 | 88 | 4.26E-33 | 128 | hypothetical protein [Bifidobacterium sp. 12_1_47BFAA] |
| Q8A007 | CLOHYLEM_04897 | 27.857 | 97 | 6.91E-32 | 124 | hypothetical protein [Lachnospiraceae [Clostridium] hylemonae DSM 15053] |
| Q8A007 | HMPREF1015_00166 | 31.513 | 82 | 1.45E-31 | 123 | hypothetical protein [Bacillus smithii 7_3_47FAA] |
| Q8ABF8 | HMPREF9007_01100 | 100 | 100 | 0 | 561 | esterase [Bacteroides sp. 1_1_14] |
| Q8ABF8 | BSIG_4128 | 99.632 | 100 | 0 | 560 | hypothetical protein [Bacteroides sp. 1_1_6] |
| Q8ABF8 | HMPREF0106_04238 | 85.294 | 100 | 5.39E-179 | 497 | esterase [Bacteroides sp. D22] |
| Q8ABF8 | HMPREF0102_02434 | 84.926 | 100 | 4.47E-178 | 494 | putative esterase [Bacteroides sp. 2_1_22] |
| Q8ABF8 | BSAG_01007 | 84.926 | 100 | 4.47E-178 | 494 | putative esterase [Bacteroides sp. D1] |
| Q8ABF8 | CW3_0980 | 84.926 | 100 | 4.47E-178 | 494 | putative esterase [Bacteroides xylanisolvens SD CC 1b] |
| Q8ABF8 | CW1_0242 | 84.926 | 100 | 4.47E-178 | 494 | putative esterase [Bacteroides xylanisolvens SD CC 2a] |
| Q8ABF8 | HMPREF0127_02851 | 84.191 | 100 | 5.95E-177 | 491 | hypothetical protein [Bacteroides sp. 1_1_30] |
| Q8ABF8 | HMPREF9010_01867 | 85.227 | 97 | 9.13E-174 | 484 | putative esterase [Bacteroides sp. 3_1_23] |
| Q8ABF8 | HMPREF1017_03441 | 85.227 | 97 | 9.13E-174 | 484 | hypothetical protein [Bacteroides ovatus 3_8_47FAA] |
| Q8ABF8 | BSCG_00908 | 85.714 | 95 | 1.05E-171 | 478 | putative esterase [Bacteroides sp. 2_2_4] |
| Q8ABF8 | BACOVA_01285 | 85.714 | 95 | 1.05E-171 | 478 | putative esterase [Bacteroides ovatus ATCC 8483] |
| Q8ABF8 | CUY_4289 | 85.328 | 95 | 3.60E-171 | 477 | putative esterase [Bacteroides ovatus SD CMC 3f] |
| Q8ABF8 | BSGG_0802 | 85.328 | 95 | 7.10E-171 | 476 | hypothetical protein [Bacteroides sp. D2] |
| Q8ABF8 | BACCAC_02168 | 83.088 | 100 | 3.88E-167 | 467 | putative esterase [Bacteroides caccae ATCC 43185] |
| Q8ABF8 | BACFIN_07319 | 80.669 | 99 | 4.42E-158 | 444 | putative esterase [Bacteroides finegoldii DSM 17565] |
| Q8ABF8 | BSHG_0306 | 75.926 | 99 | 1.68E-157 | 442 | hypothetical protein [Bacteroides sp. 3_2_5] |
| Q8ABF8 | HMPREF1018_01506 | 75.926 | 99 | 1.68E-157 | 442 | hypothetical protein [Bacteroides sp. 2_1_56FAA] |
| Q8ABF8 | HMPREF0101_00353 | 75.926 | 99 | 1.68E-157 | 442 | putative esterase [Bacteroides sp. 2_1_16] |
| Q8ABF8 | HMPREF1016_00125 | 72.059 | 100 | 6.71E-154 | 433 | acetyl esterase [Bacteroides eggerthii 1_2_48FAA] |
| Q8ABF8 | BACEGG_01996 | 72.059 | 100 | 1.99E-153 | 432 | putative esterase [Bacteroides eggerthii DSM 20697] |
| Q8ABF8 | BACINT_02983 | 71.691 | 100 | 6.20E-153 | 431 | hypothetical protein [Bacteroides intestinalis DSM 17393] |
| Q8ABF8 | BACCELL_05288 | 71.324 | 100 | 1.25E-152 | 430 | putative esterase [Bacteroides cellulosilyticus DSM 14838] |
| Q8ABF8 | HMPREF0969_00187 | 70.956 | 100 | 1.76E-151 | 427 | putative esterase [Bacteroides sp. D20] |
| Q8ABF8 | BACUNI_01322 | 70.956 | 100 | 1.76E-151 | 427 | putative esterase [Bacteroides uniformis ATCC 8492] |
| Q8ABF8 | HMPREF1007_01620 | 70.956 | 100 | 1.76E-151 | 427 | acetyl esterase [Bacteroides sp. 4_1_36] |
| Q8ABF8 | HMPREF9446_00857 | 70.588 | 100 | 6.65E-150 | 423 | putative esterase [Bacteroides fluxus YIT 12057] |
| Q8ABF8 | HMPREF1532_00361 | 77.075 | 93 | 3.02E-149 | 421 | hypothetical protein [Bacteroides salyersiae WAL 10018 = DSM 18765 = JCM 12988] |
| Q8ABF8 | HMPREF1214_03978 | 76.68 | 93 | 4.36E-148 | 419 | hypothetical protein [Bacteroides sp. HPS0048] |
| Q8ABF8 | HMPREF9447_03836 | 72.059 | 100 | 4.63E-148 | 419 | hypothetical protein [Bacteroides oleiciplenus YIT 12058] |
| Q8ABF8 | BFAG_00626 | 76.8 | 92 | 1.41E-146 | 415 | putative esterase [Bacteroides fragilis 3_1_12] |
| Q8ABF8 | HMPREF9445_00257 | 67.279 | 100 | 2.12E-146 | 414 | putative esterase [Bacteroides clarus YIT 12056] |
| Q8ABF8 | BACSTE_01707 | 66.176 | 100 | 7.41E-143 | 406 | putative esterase [Bacteroides stercoris ATCC 43183] |
| Q8ABF8 | HMPREF0619_00405 | 62.5 | 100 | 4.68E-128 | 368 | putative esterase [Parabacteroides sp. D13] |
| Q8ABF8 | PRABACTJOHN_04554 | 63.258 | 97 | 4.68E-128 | 368 | putative esterase [Parabacteroides johnsonii DSM 18315] |
| Q8ABF8 | HMPREF9008_03800 | 62.5 | 100 | 2.36E-127 | 366 | putative esterase [Parabacteroides sp. 20_3] |
| Q8ABF8 | PARMER_02579 | 64.504 | 96 | 3.90E-127 | 365 | putative esterase [Parabacteroides merdae ATCC 43184] |
| Q8ABF8 | HMPREF0104_00280 | 62.5 | 100 | 9.68E-127 | 365 | esterase [Bacteroides sp. 3_1_19] |
| Q8ABF8 | HMPREF0103_2008 | 62.5 | 100 | 1.38E-126 | 364 | putative esterase [Bacteroides sp. 2_1_33B] |
| Q8ABF8 | BACCOP_04022 | 62.868 | 99 | 5.70E-126 | 363 | putative esterase [Bacteroides coprocola DSM 17136] |
| Q8ABF8 | HMPREF9456_00448 | 60.37 | 99 | 3.85E-124 | 358 | hypothetical protein [Dysgonomonas mossii DSM 22836] |
| Q8ABF8 | BSBG_01789 | 63.971 | 100 | 9.87E-124 | 357 | putative esterase [Bacteroides sp. 9_1_42FAA] |
| Q8ABF8 | HMPREF0105_0634 | 67.46 | 93 | 1.48E-123 | 357 | putative esterase [Bacteroides sp. 3_1_33FAA] |
| Q8ABF8 | BSEG_00689 | 67.46 | 93 | 1.48E-123 | 357 | putative esterase [Bacteroides dorei 5_1_36/D4] |
| Q8ABF8 | BACDOR_03937 | 67.46 | 93 | 1.90E-123 | 356 | putative esterase [Bacteroides dorei DSM 17855] |
| Q8ABF8 | BSFG_04155 | 62.868 | 100 | 6.70E-123 | 355 | putative esterase [Bacteroides sp. 4_3_47FAA] |
| Q8ABF8 | HMPREF9011_02748 | 62.868 | 100 | 6.70E-123 | 355 | acetyl esterase [Bacteroides sp. 3_1_40A] |
| Q8ABF8 | CUU_2570 | 66.667 | 93 | 1.21E-122 | 354 | putative esterase [Bacteroides vulgatus PC510] |
| Q8ABF8 | BACCOPRO_03683 | 60.547 | 94 | 3.13E-116 | 338 | putative esterase [Bacteroides coprophilus DSM 18228 = JCM 13818] |
| Q8ABF8 | HMPREF9442_01591 | 56.777 | 100 | 9.38E-113 | 329 | putative esterase [Paraprevotella xylaniphila YIT 11841] |

| | | | | | | |
|---|---|---|---|---|---|---|
| Q8ABF8 | HMPREF9449_01330 | 56.777 | 99 | 6.14E-111 | 325 | hypothetical protein [Odoribacter laneus YIT 12061] |
| Q8ABF8 | HMPREF9441_00629 | 56.25 | 99 | 1.02E-110 | 324 | putative esterase [Paraprevotella clara YIT 11840] |
| Q8ABF8 | HMPREF1033_02388 | 59.449 | 93 | 2.01E-110 | 323 | hypothetical protein [Tannerella sp. 6_1_58FAA_CT1] |
| Q8ABF8 | HMPREF9455_02698 | 55.678 | 100 | 3.06E-110 | 323 | hypothetical protein [Dysgonomonas gadei ATCC BAA-286] |
| Q8ABF8 | HMPREF9448_00716 | 53.48 | 100 | 4.98E-106 | 312 | hypothetical protein [Barnesiella intestinihominis YIT 11860] |
| Q8ABF8 | BSBG_04718 | 54.023 | 96 | 9.48E-104 | 306 | putative esterase [Bacteroides sp. 9_1_42FAA] |
| Q8ABF8 | BACDOR_04583 | 54.023 | 96 | 1.57E-103 | 307 | putative esterase [Bacteroides dorei DSM 17855] |
| Q8ABF8 | HMPREF9450_00050 | 51.087 | 94 | 4.73E-94 | 283 | hypothetical protein [Alistipes indistinctus YIT 12060] |
| Q8ABF8 | HMPREF9450_00048 | 48.763 | 95 | 1.01E-92 | 283 | hypothetical protein [Alistipes indistinctus YIT 12060] |
| Q8ABF8 | HMPREF9455_02699 | 43.173 | 99 | 2.06E-76 | 237 | hypothetical protein [Dysgonomonas gadei ATCC BAA-286] |
| Q8AB74 | BSIG_0059 | 100 | 100 | 0 | 1431 | hypothetical protein [Bacteroides sp. 1_1_6] |
| Q8AB74 | HMPREF0127_02919 | 92.962 | 100 | 0 | 1347 | hypothetical protein [Bacteroides sp. 1_1_30] |
| Q8AB74 | BXY_42300 | 92.962 | 100 | 0 | 1347 | dipeptidyl-peptidase 7. Serine peptidase. MEROPS family S46 [Bacteroides xylanisolvens XB1A] |
| Q8AB74 | HMPREF0106_04190 | 92.962 | 100 | 0 | 1347 | hypothetical protein [Bacteroides sp. D22] |
| Q8AB74 | HMPREF0102_02503 | 92.815 | 100 | 0 | 1343 | hypothetical protein [Bacteroides sp. 2_1_22] |
| Q8AB74 | BSAG_01079 | 92.815 | 100 | 0 | 1343 | hypothetical protein [Bacteroides sp. D1] |
| Q8AB74 | CW3_4225 | 92.815 | 100 | 0 | 1343 | conserved hypothetical protein [Bacteroides xylanisolvens SD CC 1b] |
| Q8AB74 | CW1_0172 | 92.815 | 100 | 0 | 1343 | conserved hypothetical protein [Bacteroides xylanisolvens SD CC 2a] |
| Q8AB74 | BACFIN_07387 | 92.229 | 100 | 0 | 1338 | hypothetical protein [Bacteroides finegoldii DSM 17565] |
| Q8AB74 | HMPREF9010_01793 | 92.669 | 100 | 0 | 1337 | hypothetical protein [Bacteroides sp. 3_1_23] |
| Q8AB74 | BACOVA_00385 | 92.669 | 100 | 0 | 1337 | hypothetical protein [Bacteroides ovatus ATCC 8483] |
| Q8AB74 | BSGG_0736 | 92.669 | 100 | 0 | 1337 | hypothetical protein [Bacteroides sp. D2] |
| Q8AB74 | BSCG_00819 | 92.669 | 100 | 0 | 1336 | hypothetical protein [Bacteroides sp. 2_2_4] |
| Q8AB74 | HMPREF1017_03508 | 92.522 | 100 | 0 | 1335 | hypothetical protein [Bacteroides ovatus 3_8_47FAA] |
| Q8AB74 | CUY_4107 | 92.522 | 100 | 0 | 1335 | conserved hypothetical protein [Bacteroides ovatus SD CMC 3f] |
| Q8AB74 | BACCAC_02236 | 91.63 | 99 | 0 | 1331 | hypothetical protein [Bacteroides caccae ATCC 43185] |
| Q8AB74 | HMPREF1214_03869 | 83.748 | 100 | 0 | 1235 | hypothetical protein [Bacteroides sp. HPS0048] |
| Q8AB74 | HMPREF1532_00293 | 82.991 | 99 | 0 | 1229 | hypothetical protein [Bacteroides salyersiae WAL 10018 = DSM 18765 = JCM 12988] |
| Q8AB74 | BSHG_1935 | 81.672 | 99 | 0 | 1200 | hypothetical protein [Bacteroides sp. 3_2_5] |
| Q8AB74 | HMPREF0101_03126 | 81.672 | 99 | 0 | 1199 | hypothetical protein [Bacteroides sp. 2_1_16] |
| Q8AB74 | HMPREF1018_03522 | 81.525 | 99 | 0 | 1197 | hypothetical protein [Bacteroides sp. 2_1_56FAA] |
| Q8AB74 | BFAG_03218 | 81.085 | 99 | 0 | 1194 | hypothetical protein [Bacteroides fragilis 3_1_12] |
| Q8AB74 | BACINT_02883 | 79.912 | 99 | 0 | 1185 | hypothetical protein [Bacteroides intestinalis DSM 17393] |
| Q8AB74 | BACCELL_05143 | 79.326 | 99 | 0 | 1176 | hypothetical protein [Bacteroides cellulosilyticus DSM 14838] |
| Q8AB74 | HMPREF9447_03716 | 78.739 | 99 | 0 | 1169 | hypothetical protein [Bacteroides oleiciplenus YIT 12058] |
| Q8AB74 | HMPREF1016_00218 | 79.326 | 99 | 0 | 1168 | peptidase S46 [Bacteroides eggerthii 1_2_48FAA] |
| Q8AB74 | BACEGG_02069 | 79.326 | 99 | 0 | 1168 | hypothetical protein [Bacteroides eggerthii DSM 20697] |
| Q8AB74 | HMPREF9446_00759 | 79.502 | 100 | 0 | 1166 | hypothetical protein [Bacteroides fluxus YIT 12057] |
| Q8AB74 | BACSTE_01783 | 79.179 | 99 | 0 | 1165 | hypothetical protein [Bacteroides stercoris ATCC 43183] |
| Q8AB74 | HMPREF9445_00326 | 78.739 | 99 | 0 | 1165 | hypothetical protein [Bacteroides clarus YIT 12056] |
| Q8AB74 | HMPREF0969_00398 | 78.624 | 100 | 0 | 1163 | hypothetical protein [Bacteroides sp. D20] |
| Q8AB74 | BACUNI_01424 | 78.477 | 100 | 0 | 1162 | hypothetical protein [Bacteroides uniformis ATCC 8492] |
| Q8AB74 | HMPREF1007_01539 | 78.477 | 100 | 0 | 1162 | peptidase S46 [Bacteroides sp. 4_1_36] |
| Q8AB74 | BSFG_01937 | 66.522 | 99 | 0 | 980 | hypothetical protein [Bacteroides sp. 4_3_47FAA] |
| Q8AB74 | HMPREF9011_02237 | 66.522 | 99 | 0 | 980 | hypothetical protein [Bacteroides sp. 3_1_40A] |
| Q8AB74 | CUU_0387 | 66.522 | 99 | 0 | 979 | conserved hypothetical protein [Bacteroides vulgatus PC510] |
| Q8AB74 | BSBG_02477 | 66.232 | 99 | 0 | 971 | hypothetical protein [Bacteroides sp. 9_1_42FAA] |
| Q8AB74 | HMPREF0105_0858 | 66.232 | 99 | 0 | 971 | hypothetical protein [Bacteroides sp. 3_1_33FAA] |
| Q8AB74 | BACDOR_03539 | 66.232 | 99 | 0 | 971 | hypothetical protein [Bacteroides dorei DSM 17855] |
| Q8AB74 | BSEG_00915 | 66.087 | 99 | 0 | 970 | hypothetical protein [Bacteroides dorei 5_1_36/D4] |
| Q8AB74 | BACCOPRO_02782 | 65.982 | 99 | 0 | 964 | hypothetical protein [Bacteroides coprophilus DSM 18228 = JCM 13818] |
| Q8AB74 | BACPLE_01877 | 65.836 | 99 | 0 | 963 | hypothetical protein [Bacteroides plebeius DSM 17135] |
| Q8AB74 | BACCOP_03275 | 65.982 | 99 | 0 | 947 | hypothetical protein [Bacteroides coprocola DSM 17136] |
| Q8AB74 | PARMER_02946 | 52.485 | 100 | 0 | 735 | hypothetical protein [Parabacteroides merdae ATCC 43184] |
| Q8AB74 | PRABACTJOHN_02704 | 52.047 | 100 | 0 | 733 | hypothetical protein [Parabacteroides johnsonii DSM 18315] |
| Q8AB74 | HMPREF1033_02741 | 51.098 | 99 | 0 | 722 | hypothetical protein [Tannerella sp. 6_1_58FAA_CT1] |
| Q8AB74 | HMPREF1033_00928 | 50.292 | 99 | 0 | 718 | hypothetical protein [Tannerella sp. 6_1_58FAA_CT1] |
| Q8AB74 | HMPREF0103_1442 | 51.245 | 99 | 0 | 713 | hypothetical protein [Bacteroides sp. 2_1_33B] |
| Q8AB74 | HMPREF9008_01598 | 51.101 | 99 | 0 | 710 | hypothetical protein [Parabacteroides sp. 20_3] |
| Q8AB74 | HMPREF0619_00990 | 50.954 | 99 | 0 | 709 | hypothetical protein [Parabacteroides sp. D13] |

| | | | | | | |
|---|---|---|---|---|---|---|
| Q8AB74 | HMPREF0104_00878 | 50.954 | 99 | 0 | 709 | hypothetical protein [Bacteroides sp. 3_1_19] |
| Q8AB74 | BACCAC_02237 | 48.538 | 99 | 0 | 669 | hypothetical protein [Bacteroides caccae ATCC 43185] |
| Q8AB74 | HMPREF1214_03868 | 48.326 | 99 | 0 | 669 | hypothetical protein [Bacteroides sp. HPS0048] |
| Q8AB74 | HMPREF0105_0857 | 49.341 | 99 | 0 | 668 | hypothetical protein [Bacteroides sp. 3_1_33FAA] |
| Q8AB74 | BSEG_00914 | 49.341 | 99 | 0 | 668 | hypothetical protein [Bacteroides dorei 5_1_36/D4] |
| Q8AB74 | BSBG_02476 | 49.341 | 99 | 0 | 668 | hypothetical protein [Bacteroides sp. 9_1_42FAA] |
| Q8AB74 | BSFG_01936 | 49.341 | 99 | 0 | 667 | hypothetical protein [Bacteroides sp. 4_3_47FAA] |
| Q8AB74 | HMPREF9011_02238 | 49.341 | 99 | 0 | 667 | hypothetical protein [Bacteroides sp. 3_1_40A] |
| Q8AB74 | BACDOR_03540 | 49.195 | 99 | 0 | 666 | hypothetical protein [Bacteroides dorei DSM 17855] |
| Q8AB74 | CUU_0388 | 49.195 | 99 | 0 | 665 | conserved hypothetical protein [Bacteroides vulgatus PC510] |
| Q8AB74 | HMPREF1532_00292 | 47.889 | 99 | 0 | 665 | hypothetical protein [Bacteroides salyersiae WAL 10018 = DSM 18765 = JCM 12988] |
| Q8AB74 | BACCOP_03276 | 48.463 | 99 | 0 | 661 | hypothetical protein [Bacteroides coprocola DSM 17136] |
| Q8AB74 | BACCOPRO_02783 | 48.035 | 99 | 0 | 657 | hypothetical protein [Bacteroides coprophilus DSM 18228 = JCM 13818] |
| Q8AB74 | BFAG_03217 | 47.584 | 99 | 0 | 656 | hypothetical protein [Bacteroides fragilis 3_1_12] |
| Q8AB74 | BSHG_1934 | 47.877 | 99 | 0 | 656 | hypothetical protein [Bacteroides sp. 3_2_5] |
| Q8AB74 | HMPREF1018_03521 | 47.877 | 99 | 0 | 656 | hypothetical protein [Bacteroides sp. 2_1_56FAA] |
| Q8AB74 | BACFIN_07388 | 47.953 | 99 | 0 | 656 | hypothetical protein [Bacteroides finegoldii DSM 17565] |
| Q8AB74 | HMPREF0101_03127 | 47.877 | 99 | 0 | 656 | hypothetical protein [Bacteroides sp. 2_1_16] |
| Q8AB74 | BSIG_0060 | 47.507 | 99 | 0 | 656 | hypothetical protein [Bacteroides sp. 1_1_6] |
| Q8AB74 | BACPLE_01878 | 47.591 | 99 | 0 | 654 | hypothetical protein [Bacteroides plebeius DSM 17135] |
| Q8AB74 | HMPREF9007_01172 | 47.507 | 99 | 0 | 654 | hypothetical protein [Bacteroides sp. 1_1_14] |
| Q8AB74 | BXY_42310 | 48.099 | 99 | 0 | 654 | dipeptidyl-peptidase 7. Serine peptidase. MEROPS family S46 [Bacteroides xylanisolvens XB1A] |
| Q8AB74 | HMPREF0106_04191 | 47.953 | 99 | 0 | 653 | hypothetical protein [Bacteroides sp. D22] |
| Q8AB74 | HMPREF0127_02920 | 47.953 | 99 | 0 | 652 | hypothetical protein [Bacteroides sp. 1_1_30] |
| Q8AB74 | BSGG_0735 | 47.947 | 99 | 0 | 651 | hypothetical protein [Bacteroides sp. D2] |
| Q8AB74 | HMPREF1017_03509 | 47.947 | 99 | 0 | 651 | hypothetical protein [Bacteroides ovatus 3_8_47FAA] |
| Q8AB74 | HMPREF0102_02504 | 47.807 | 99 | 0 | 651 | hypothetical protein [Bacteroides sp. 2_1_22] |
| Q8AB74 | BSAG_01080 | 47.807 | 99 | 0 | 651 | hypothetical protein [Bacteroides sp. D1] |
| Q8AB74 | CW3_4226 | 47.807 | 99 | 0 | 651 | conserved hypothetical protein [Bacteroides xylanisolvens SD CC 1b] |
| Q8AB74 | CW1_0171 | 47.807 | 99 | 0 | 651 | conserved hypothetical protein [Bacteroides xylanisolvens SD CC 2a] |
| Q8AB74 | CUY_4106 | 47.801 | 99 | 0 | 649 | conserved hypothetical protein [Bacteroides ovatus SD CMC 3f] |
| Q8AB74 | HMPREF9010_01792 | 47.654 | 99 | 0 | 648 | hypothetical protein [Bacteroides sp. 3_1_23] |
| Q8AB74 | BSCG_00818 | 47.654 | 99 | 0 | 646 | hypothetical protein [Bacteroides sp. 2_2_4] |
| Q8AB74 | BACOVA_00386 | 47.507 | 99 | 0 | 645 | hypothetical protein [Bacteroides ovatus ATCC 8483] |
| Q8AB74 | HMPREF9445_00327 | 46.852 | 99 | 0 | 641 | hypothetical protein [Bacteroides clarus YIT 12056] |
| Q8AB74 | HMPREF9447_03715 | 46.725 | 99 | 0 | 639 | hypothetical protein [Bacteroides oleiciplenus YIT 12058] |
| Q8AB74 | BACEGG_02070 | 46.999 | 99 | 0 | 639 | hypothetical protein [Bacteroides eggerthii DSM 20697] |
| Q8AB74 | BACINT_02882 | 46.413 | 99 | 0 | 638 | hypothetical protein [Bacteroides intestinalis DSM 17393] |
| Q8AB74 | HMPREF9446_00758 | 46.579 | 99 | 0 | 635 | hypothetical protein [Bacteroides fluxus YIT 12057] |
| Q8AB74 | HMPREF1016_00219 | 46.852 | 99 | 0 | 632 | peptidase S46 [Bacteroides eggerthii 1_2_48FAA] |
| Q8AB74 | HMPREF0969_00399 | 46.559 | 99 | 0 | 632 | hypothetical protein [Bacteroides sp. D20] |
| Q8AB74 | BACUNI_01423 | 46.559 | 99 | 0 | 632 | hypothetical protein [Bacteroides uniformis ATCC 8492] |
| Q8AB74 | HMPREF1007_01538 | 46.559 | 99 | 0 | 632 | peptidase S46 [Bacteroides sp. 4_1_36] |
| Q8AB74 | BACCELL_05142 | 46.413 | 99 | 0 | 631 | hypothetical protein [Bacteroides cellulosilyticus DSM 14838] |
| Q8AB74 | BACSTE_01784 | 46.266 | 99 | 0 | 624 | hypothetical protein [Bacteroides stercoris ATCC 43183] |
| Q8AB74 | HMPREF9450_00755 | 44.331 | 99 | 0 | 600 | hypothetical protein [Alistipes indistinctus YIT 12060] |
| Q8AB74 | ALIPUT_02112 | 45.743 | 99 | 0 | 591 | hypothetical protein [Alistipes putredinis DSM 17216] |
| Q8AB74 | AL1_16030 | 44.023 | 100 | 0 | 574 | Peptidase S46. [Alistipes shahii WAL 8301] |
| Q8AB74 | HMPREF9449_02133 | 40.514 | 99 | 6.07E-180 | 533 | hypothetical protein [Odoribacter laneus YIT 12061] |
| Q8AB74 | HMPREF9450_02265 | 39.971 | 99 | 9.12E-166 | 496 | hypothetical protein [Alistipes indistinctus YIT 12060] |
| Q8AB74 | PARMER_00304 | 37.936 | 99 | 3.63E-165 | 494 | hypothetical protein [Parabacteroides merdae ATCC 43184] |
| Q8AB74 | CUU_0716 | 39.213 | 99 | 3.42E-164 | 492 | conserved hypothetical protein [Bacteroides vulgatus PC510] |
| Q8AB74 | HMPREF9445_01804 | 39.39 | 99 | 5.10E-164 | 492 | hypothetical protein [Bacteroides clarus YIT 12056] |
| Q8AB74 | HMPREF0105_3274 | 39.01 | 100 | 5.15E-164 | 491 | hypothetical protein [Bacteroides sp. 3_1_33FAA] |
| Q8AB74 | BSBG_03122 | 39.01 | 100 | 5.15E-164 | 491 | hypothetical protein [Bacteroides sp. 9_1_42FAA] |
| Q8AB74 | BSEG_03017 | 39.01 | 100 | 5.86E-164 | 491 | hypothetical protein [Bacteroides dorei 5_1_36/D4] |
| Q8AB74 | BACDOR_02152 | 39.01 | 100 | 5.86E-164 | 491 | hypothetical protein [Bacteroides dorei DSM 17855] |
| Q8AB74 | BSFG_00163 | 39.067 | 99 | 4.51E-163 | 489 | hypothetical protein [Bacteroides sp. 4_3_47FAA] |
| Q8AB74 | HMPREF9011_00248 | 39.067 | 99 | 4.51E-163 | 489 | hypothetical protein [Bacteroides sp. 3_1_40A] |
| Q8AB74 | HMPREF1007_00282 | 38.776 | 99 | 5.61E-163 | 489 | peptidase S46 [Bacteroides sp. 4_1_36] |

| | | | | | | |
|---|---|---|---|---|---|---|
| Q8AB74 | HMPREF0969_01517 | 38.776 | 99 | 6.05E-163 | 489 | hypothetical protein [Bacteroides sp. D20] |
| Q8AB74 | HMPREF9446_00563 | 38.808 | 100 | 1.01E-162 | 488 | hypothetical protein [Bacteroides fluxus YIT 12057] |
| Q8AB74 | PRABACTJOHN_00535 | 37.609 | 99 | 1.72E-162 | 488 | hypothetical protein [Parabacteroides johnsonii DSM 18315] |
| Q8AB74 | BACUNI_01062 | 38.63 | 99 | 2.65E-162 | 487 | hypothetical protein [Bacteroides uniformis ATCC 8492] |
| Q8AB74 | BACSTE_02429 | 39.332 | 99 | 3.82E-162 | 487 | hypothetical protein [Bacteroides stercoris ATCC 43183] |
| Q8AB74 | HMPREF0619_00047 | 38.265 | 99 | 7.87E-162 | 486 | hypothetical protein [Parabacteroides sp. D13] |
| Q8AB74 | HMPREF9008_04196 | 38.122 | 99 | 1.51E-160 | 483 | hypothetical protein [Parabacteroides sp. 20_3] |
| Q8AB74 | HMPREF1016_00890 | 38.63 | 99 | 1.81E-160 | 483 | peptidase S46 [Bacteroides eggerthii 1_2_48FAA] |
| Q8AB74 | BACEGG_03368 | 38.63 | 99 | 1.81E-160 | 483 | hypothetical protein [Bacteroides eggerthii DSM 20697] |
| Q8AB74 | BSGG_2332 | 38.428 | 99 | 2.40E-160 | 482 | hypothetical protein [Bacteroides sp. D2] |
| Q8AB74 | BACOVA_00899 | 38.428 | 99 | 2.40E-160 | 482 | hypothetical protein [Bacteroides ovatus ATCC 8483] |
| Q8AB74 | CUY_4404 | 38.428 | 99 | 2.98E-160 | 482 | conserved hypothetical protein [Bacteroides ovatus SD CMC 3f] |
| Q8AB74 | HMPREF9010_00752 | 38.428 | 99 | 2.98E-160 | 482 | hypothetical protein [Bacteroides sp. 3_1_23] |
| Q8AB74 | BSCG_03059 | 38.428 | 99 | 2.98E-160 | 482 | hypothetical protein [Bacteroides sp. 2_2_4] |
| Q8AB74 | HMPREF1532_01775 | 39.424 | 99 | 4.96E-160 | 481 | hypothetical protein [Bacteroides salyersiae WAL 10018 = DSM 18765 = JCM 12988] |
| Q8AB74 | HMPREF0104_00053 | 37.946 | 99 | 6.41E-160 | 481 | hypothetical protein [Bacteroides sp. 3_1_19] |
| Q8AB74 | BXY_25380 | 38.574 | 99 | 7.78E-160 | 480 | dipeptidyl-peptidase 7. Serine peptidase. MEROPS family S46 [Bacteroides xylanisolvens XB1A] |
| Q8AB74 | HMPREF0127_01618 | 38.574 | 99 | 7.78E-160 | 481 | hypothetical protein [Bacteroides sp. 1_1_30] |
| Q8AB74 | HMPREF0106_04280 | 38.574 | 99 | 7.78E-160 | 481 | hypothetical protein [Bacteroides sp. D22] |
| Q8AB74 | BACCAC_01485 | 38.428 | 99 | 1.14E-159 | 481 | hypothetical protein [Bacteroides caccae ATCC 43185] |
| Q8AB74 | HMPREF1017_00565 | 38.282 | 99 | 1.80E-159 | 480 | hypothetical protein [Bacteroides ovatus 3_8_47FAA] |
| Q8AB74 | HMPREF0101_02131 | 39.01 | 100 | 1.95E-159 | 480 | hypothetical protein [Bacteroides sp. 2_1_16] |
| Q8AB74 | BFAG_02873 | 39.01 | 100 | 2.51E-159 | 479 | hypothetical protein [Bacteroides fragilis 3_1_12] |
| Q8AB74 | HMPREF0103_2247 | 37.838 | 99 | 6.65E-159 | 479 | hypothetical protein [Bacteroides sp. 2_1_33B] |
| Q8AB74 | HMPREF1018_00169 | 38.865 | 100 | 7.13E-159 | 479 | hypothetical protein [Bacteroides sp. 2_1_56FAA] |
| Q8AB74 | HMPREF0102_00568 | 38.428 | 99 | 8.16E-159 | 478 | hypothetical protein [Bacteroides sp. 2_1_22] |
| Q8AB74 | BSAG_00318 | 38.428 | 99 | 8.16E-159 | 478 | hypothetical protein [Bacteroides sp. D1] |
| Q8AB74 | CW3_2069 | 38.428 | 99 | 8.16E-159 | 478 | conserved hypothetical protein [Bacteroides xylanisolvens SD CC 1b] |
| Q8AB74 | CW1_0771 | 38.428 | 99 | 8.16E-159 | 478 | conserved hypothetical protein [Bacteroides xylanisolvens SD CC 2a] |
| Q8AB74 | BSHG_1458 | 38.865 | 100 | 1.76E-158 | 478 | hypothetical protein [Bacteroides sp. 3_2_5] |
| Q8AB74 | HMPREF9455_01975 | 38.439 | 99 | 8.67E-158 | 476 | hypothetical protein [Dysgonomonas gadei ATCC BAA-286] |
| Q8AB74 | BACFIN_05173 | 38.574 | 99 | 4.30E-157 | 474 | hypothetical protein [Bacteroides finegoldii DSM 17565] |
| Q8AB74 | BSIG_1382 | 37.7 | 99 | 8.04E-157 | 473 | hypothetical protein [Bacteroides sp. 1_1_6] |
| Q8AB74 | HMPREF9007_03829 | 37.7 | 99 | 1.17E-156 | 473 | hypothetical protein [Bacteroides sp. 1_1_14] |
| Q8AB74 | HMPREF1214_00608 | 38.417 | 99 | 1.22E-156 | 473 | hypothetical protein [Bacteroides sp. HPS0048] |
| Q8AB74 | BACINT_01296 | 38.34 | 99 | 1.26E-156 | 473 | hypothetical protein [Bacteroides intestinalis DSM 17393] |
| Q8AB74 | HMPREF9447_02263 | 37.971 | 99 | 1.34E-155 | 470 | hypothetical protein [Bacteroides oleiciplenus YIT 12058] |
| Q8AB74 | HMPREF9456_01457 | 37.881 | 99 | 4.61E-155 | 469 | hypothetical protein [Dysgonomonas mossii DSM 22836] |
| Q8AB74 | BACCELL_01726 | 38.116 | 99 | 5.22E-155 | 468 | hypothetical protein [Bacteroides cellulosilyticus DSM 14838] |
| Q8AB74 | HMPREF9449_00904 | 37.625 | 99 | 3.35E-149 | 454 | hypothetical protein [Odoribacter laneus YIT 12061] |
| Q8AB74 | ALIPUT_00511 | 37.172 | 99 | 2.30E-146 | 445 | hypothetical protein [Alistipes putredinis DSM 17216] |
| Q8AB74 | HMPREF9442_02956 | 36.403 | 99 | 2.02E-145 | 444 | hypothetical protein [Paraprevotella xylaniphila YIT 11841] |
| Q8AB74 | HMPREF9441_00315 | 36.403 | 99 | 1.17E-143 | 439 | hypothetical protein [Paraprevotella clara YIT 11840] |
| Q8AB74 | AL1_16890 | 37.5 | 100 | 2.86E-142 | 434 | Peptidase S46. [Alistipes shahii WAL 8301] |
| Q8AB74 | HMPREF9449_02174 | 36.219 | 100 | 8.34E-142 | 435 | hypothetical protein [Odoribacter laneus YIT 12061] |
| Q8AB74 | AL1_16630 | 35.694 | 99 | 7.53E-137 | 422 | Peptidase S46. [Alistipes shahii WAL 8301] |
| Q8AB74 | PREVCOP_06589 | 34.857 | 99 | 3.44E-127 | 397 | hypothetical protein [Prevotella copri DSM 18205] |
| Q8AB74 | HMPREF1475_01683 | 35.613 | 99 | 4.07E-127 | 396 | hypothetical protein [Prevotella oralis HGA0225] |
| Q8AB74 | HMPREF9420_1755 | 34.343 | 99 | 2.16E-122 | 384 | hypothetical protein [Prevotella salivae DSM 15606] |
| Q8AB74 | HMPREF0673_00993 | 34.052 | 100 | 1.61E-120 | 379 | hypothetical protein [Prevotella stercorea DSM 18206] |
| Q8AB74 | BACOVA_04406 | 27.514 | 99 | 1.52E-83 | 283 | hypothetical protein [Bacteroides ovatus ATCC 8483] |
| Q8AB74 | BSGG_3154 | 27.235 | 99 | 7.00E-82 | 278 | hypothetical protein [Bacteroides sp. D2] |
| Q8A8T0 | BSIG_0525 | 98.835 | 100 | 0 | 1063 | hypothetical protein [Bacteroides sp. 1_1_6] |
| Q8A8T0 | HMPREF9007_04520 | 98.81 | 98 | 0 | 1039 | xylanase [Bacteroides sp. 1_1_14] |
| Q8A8T0 | BSAG_01480 | 76.654 | 99 | 0 | 847 | hypothetical protein [Bacteroides sp. D1] |
| Q8A8T0 | CW3_0320 | 76.654 | 99 | 0 | 847 | conserved hypothetical protein [Bacteroides xylanisolvens SD CC 1b] |
| Q8A8T0 | CW1_0848 | 76.654 | 99 | 0 | 847 | conserved hypothetical protein [Bacteroides xylanisolvens SD CC 2a] |
| Q8A8T0 | CUY_0931 | 76.654 | 99 | 0 | 846 | conserved hypothetical protein [Bacteroides ovatus SD CMC 3f] |
| Q8A8T0 | HMPREF0127_00631 | 76.459 | 99 | 0 | 845 | hypothetical protein [Bacteroides sp. 1_1_30] |
| Q8A8T0 | HMPREF1017_03770 | 76.654 | 99 | 0 | 844 | hypothetical protein [Bacteroides ovatus 3_8_47FAA] |

| | | | | | | |
|---|---|---|---|---|---|---|
| Q8A8T0 | HMPREF0106_04667 | 76.459 | 99 | 0 | 843 | xylanase [Bacteroides sp. D22] |
| Q8A8T0 | HMPREF9010_00870 | 76.459 | 99 | 0 | 843 | putative xylanase [Bacteroides sp. 3_1_23] |
| Q8A8T0 | BSGG_0258 | 76.265 | 99 | 0 | 843 | hypothetical protein [Bacteroides sp. D2] |
| Q8A8T0 | BACOVA_04052 | 76.265 | 99 | 0 | 841 | hypothetical protein [Bacteroides ovatus ATCC 8483] |
| Q8A8T0 | HMPREF0102_02896 | 76.278 | 95 | 0 | 803 | hypothetical protein [Bacteroides sp. 2_1_22] |
| Q8A8T0 | BSCG_02541 | 76.278 | 95 | 0 | 802 | hypothetical protein [Bacteroides sp. 2_2_4] |
| Q8A8T0 | HMPREF1018_02143 | 70.979 | 55 | 1.18E-145 | 423 | hypothetical protein [Bacteroides sp. 2_1_56FAA] |
| Q8A8T0 | HMPREF0101_00969 | 70.629 | 55 | 1.59E-145 | 422 | hypothetical protein [Bacteroides sp. 2_1_16] |
| Q8A8T0 | HMPREF1016_03185 | 66.436 | 56 | 5.76E-141 | 411 | hypothetical protein [Bacteroides eggerthii 1_2_48FAA] |
| Q8A8T0 | BACEGG_02772 | 66.436 | 56 | 5.76E-141 | 411 | hypothetical protein [Bacteroides eggerthii DSM 20697] |
| Q8A8T0 | BACINT_02372 | 64.828 | 56 | 8.36E-141 | 410 | hypothetical protein [Bacteroides intestinalis DSM 17393] |
| Q8A8T0 | BACSTE_00652 | 66.782 | 56 | 1.29E-140 | 410 | hypothetical protein [Bacteroides stercoris ATCC 43183] |
| Q8A8T0 | BACCELL_04558 | 64.706 | 56 | 1.75E-140 | 410 | hypothetical protein [Bacteroides cellulosilyticus DSM 14838] |
| Q8A8T0 | HMPREF9447_00465 | 65.052 | 56 | 4.14E-140 | 409 | hypothetical protein [Bacteroides oleiciplenus YIT 12058] |
| Q8A8T0 | HMPREF9445_00632 | 66.667 | 56 | 3.38E-139 | 407 | pectin acetylesterase family protein [Bacteroides clarus YIT 12056] |
| Q8A8T0 | HMPREF0969_03517 | 66.899 | 55 | 3.86E-138 | 404 | hypothetical protein [Bacteroides sp. D20] |
| Q8A8T0 | HMPREF1007_01252 | 66.899 | 55 | 3.86E-138 | 404 | hypothetical protein [Bacteroides sp. 4_1_36] |
| Q8A8T0 | HMPREF0105_0225 | 68.929 | 54 | 1.08E-136 | 400 | hypothetical protein [Bacteroides sp. 3_1_33FAA] |
| Q8A8T0 | BSEG_00286 | 68.929 | 54 | 1.08E-136 | 400 | hypothetical protein [Bacteroides dorei 5_1_36/D4] |
| Q8A8T0 | CUU_1097 | 68.571 | 54 | 1.74E-136 | 399 | conserved hypothetical protein [Bacteroides vulgatus PC510] |
| Q8A8T0 | BACUNI_02945 | 66.551 | 55 | 2.97E-136 | 399 | hypothetical protein [Bacteroides uniformis ATCC 8492] |
| Q8A8T0 | BACDOR_04358 | 68.571 | 54 | 4.15E-136 | 398 | hypothetical protein [Bacteroides dorei DSM 17855] |
| Q8A8T0 | BSBG_01438 | 68.571 | 54 | 4.15E-136 | 398 | hypothetical protein [Bacteroides sp. 9_1_42FAA] |
| Q8A8T0 | HMPREF9446_02343 | 67.368 | 55 | 4.53E-136 | 398 | hypothetical protein [Bacteroides fluxus YIT 12057] |
| Q8A8T0 | BSFG_03001 | 68.571 | 54 | 4.58E-136 | 398 | hypothetical protein [Bacteroides sp. 4_3_47FAA] |
| Q8A8T0 | HMPREF9011_03700 | 68.571 | 54 | 4.58E-136 | 398 | xylanase [Bacteroides sp. 3_1_40A] |
| Q8A8T0 | BFAG_01517 | 65.625 | 56 | 5.14E-134 | 393 | hypothetical protein [Bacteroides fragilis 3_1_12] |
| Q8A8T0 | BACCAC_00970 | 78.07 | 44 | 4.60E-131 | 384 | hypothetical protein [Bacteroides caccae ATCC 43185] |
| Q8A8T0 | BACFIN_06874 | 76.957 | 45 | 2.73E-129 | 379 | hypothetical protein [Bacteroides finegoldii DSM 17565] |
| Q8A8T0 | BACCOP_02646 | 62.411 | 54 | 6.12E-121 | 360 | hypothetical protein [Bacteroides coprocola DSM 17136] |
| Q8A8T0 | BACPLE_03269 | 63.158 | 51 | 1.96E-114 | 343 | hypothetical protein [Bacteroides plebeius DSM 17135] |
| Q8A8T0 | BACCOPRO_01262 | 60.985 | 51 | 6.80E-109 | 329 | hypothetical protein [Bacteroides coprophilus DSM 18228 = JCM 13818] |
| Q8A8T0 | HMPREF0103_3520 | 60.63 | 49 | 1.89E-105 | 320 | hypothetical protein [Bacteroides sp. 2_1_33B] |
| Q8A8T0 | HMPREF9008_03075 | 60.236 | 49 | 2.67E-105 | 319 | hypothetical protein [Parabacteroides sp. 20_3] |
| Q8A8T0 | HMPREF0104_01154 | 60.236 | 49 | 2.05E-104 | 317 | pectin acetylesterase [Bacteroides sp. 3_1_19] |
| Q8A8T0 | HMPREF0619_01136 | 59.843 | 49 | 5.49E-104 | 316 | hypothetical protein [Parabacteroides sp. D13] |
| Q8A8T0 | HMPREF9446_02342 | 62.447 | 44 | 2.31E-98 | 301 | hypothetical protein [Bacteroides fluxus YIT 12057] |
| Q8A8T0 | HMPREF1532_03457 | 60.593 | 44 | 4.13E-95 | 293 | hypothetical protein [Bacteroides salyersiae WAL 10018 = DSM 18765 = JCM 12988] |
| Q8A8T0 | HMPREF0969_03519 | 62.931 | 43 | 6.15E-95 | 292 | hypothetical protein [Bacteroides sp. D20] |
| Q8A8T0 | BACUNI_02947 | 62.931 | 43 | 7.58E-95 | 292 | hypothetical protein [Bacteroides uniformis ATCC 8492] |
| Q8A8T0 | HMPREF1007_01250 | 62.931 | 43 | 7.83E-95 | 292 | hypothetical protein [Bacteroides sp. 4_1_36] |
| Q8A8T0 | HMPREF1214_02158 | 56.554 | 51 | 2.43E-94 | 299 | hypothetical protein [Bacteroides sp. HPS0048] |
| Q8A8T0 | HMPREF1214_02572 | 59.167 | 45 | 1.35E-93 | 289 | hypothetical protein [Bacteroides sp. HPS0048] |
| Q8A8T0 | BACCAC_01030 | 57.371 | 48 | 4.95E-93 | 293 | hypothetical protein [Bacteroides caccae ATCC 43185] |
| Q8A8T0 | HMPREF1532_03226 | 55.094 | 51 | 1.61E-91 | 285 | hypothetical protein [Bacteroides salyersiae WAL 10018 = DSM 18765 = JCM 12988] |
| Q8A8T0 | BACFIN_07184 | 56.972 | 48 | 4.03E-91 | 290 | GDSL-like protein [Bacteroides finegoldii DSM 17565] |
| Q8A8T0 | HMPREF1018_02142 | 59.836 | 46 | 4.80E-91 | 282 | hypothetical protein [Bacteroides sp. 2_1_56FAA] |
| Q8A8T0 | BSCG_02598 | 56.4 | 48 | 1.19E-90 | 288 | GDSL-like protein [Bacteroides sp. 2_2_4] |
| Q8A8T0 | HMPREF0102_02833 | 56.4 | 48 | 1.19E-90 | 288 | GDSL-like protein [Bacteroides sp. 2_1_22] |
| Q8A8T0 | BSAG_01415 | 56.4 | 48 | 1.19E-90 | 288 | GDSL-like protein [Bacteroides sp. D1] |
| Q8A8T0 | CW3_0258 | 56.4 | 48 | 1.19E-90 | 288 | GDSL-like protein [Bacteroides xylanisolvens SD CC 1b] |
| Q8A8T0 | CW1_0908 | 56.4 | 48 | 1.19E-90 | 288 | GDSL-like protein [Bacteroides xylanisolvens SD CC 2a] |
| Q8A8T0 | HMPREF1016_03183 | 58.874 | 43 | 1.28E-90 | 281 | hypothetical protein [Bacteroides eggerthii 1_2_48FAA] |
| Q8A8T0 | HMPREF0127_03401 | 56.4 | 48 | 1.31E-90 | 288 | hypothetical protein [Bacteroides sp. 1_1_30] |
| Q8A8T0 | BXY_07760 | 56.4 | 48 | 1.31E-90 | 288 | Esterase/lipase [Bacteroides xylanisolvens XB1A] |
| Q8A8T0 | HMPREF0106_01451 | 56.4 | 48 | 1.31E-90 | 288 | xylanase [Bacteroides sp. D22] |
| Q8A8T0 | CUY_0871 | 56.4 | 48 | 1.55E-90 | 288 | GDSL-like protein [Bacteroides ovatus SD CMC 3f] |
| Q8A8T0 | BACOVA_03983 | 56.175 | 48 | 1.90E-90 | 288 | GDSL-like protein [Bacteroides ovatus ATCC 8483] |
| Q8A8T0 | HMPREF9010_00809 | 56.4 | 48 | 3.12E-90 | 287 | xylanase [Bacteroides sp. 3_1_23] |
| Q8A8T0 | HMPREF1017_03830 | 56.4 | 48 | 3.43E-90 | 287 | hypothetical protein [Bacteroides ovatus 3_8_47FAA] |

| | | | | | | |
|---|---|---|---|---|---|---|
| Q8A8T0 | BACEGG_02774 | 58.442 | 43 | 6.36E-90 | 279 | hypothetical protein [Bacteroides eggerthii DSM 20697] |
| Q8A8T0 | BSGG_0323 | 56 | 48 | 7.03E-90 | 286 | hypothetical protein [Bacteroides sp. D2] |
| Q8A8T0 | HMPREF1017_02201 | 53.257 | 50 | 9.47E-90 | 280 | hypothetical protein [Bacteroides ovatus 3_8_47FAA] |
| Q8A8T0 | HMPREF9455_04127 | 49.123 | 54 | 1.28E-89 | 280 | hypothetical protein [Dysgonomonas gadei ATCC BAA-286] |
| Q8A8T0 | BACSTE_00654 | 58.009 | 43 | 2.84E-89 | 277 | hypothetical protein [Bacteroides stercoris ATCC 43183] |
| Q8A8T0 | HMPREF9445_00629 | 57.576 | 43 | 2.26E-88 | 276 | hypothetical protein [Bacteroides clarus YIT 12056] |
| Q8A8T0 | BSIG_0461 | 55.2 | 48 | 9.40E-88 | 281 | hypothetical protein [Bacteroides sp. 1_1_6] |
| Q8A8T0 | HMPREF9007_04460 | 54.8 | 48 | 4.48E-87 | 279 | xylanase [Bacteroides sp. 1_1_14] |
| Q8A8T0 | HMPREF9448_02673 | 47.842 | 54 | 1.70E-86 | 271 | hypothetical protein [Barnesiella intestinihominis YIT 11860] |
| Q8A8T0 | HMPREF1214_02379 | 48.754 | 53 | 1.92E-86 | 272 | hypothetical protein [Bacteroides sp. HPS0048] |
| Q8A8T0 | HMPREF9450_01060 | 47.687 | 54 | 2.31E-86 | 272 | hypothetical protein [Alistipes indistinctus YIT 12060] |
| Q8A8T0 | HMPREF9456_02498 | 48.387 | 53 | 3.40E-86 | 271 | hypothetical protein [Dysgonomonas mossii DSM 22836] |
| Q8A8T0 | BACINT_02374 | 61.207 | 43 | 1.82E-85 | 268 | hypothetical protein [Bacteroides intestinalis DSM 17393] |
| Q8A8T0 | BACCELL_04556 | 60.776 | 43 | 2.14E-85 | 268 | hypothetical protein [Bacteroides cellulosilyticus DSM 14838] |
| Q8A8T0 | HMPREF9420_1566 | 48.201 | 54 | 1.03E-84 | 268 | hypothetical protein [Prevotella salivae DSM 15606] |
| Q8A8T0 | HMPREF0105_3767 | 56.034 | 44 | 1.29E-84 | 273 | GDSL-like protein [Bacteroides sp. 3_1_33FAA] |
| Q8A8T0 | BSEG_03737 | 56.034 | 44 | 1.29E-84 | 273 | GDSL-like protein [Bacteroides dorei 5_1_36/D4] |
| Q8A8T0 | BSBG_04066 | 56.034 | 44 | 1.29E-84 | 273 | GDSL-like protein [Bacteroides sp. 9_1_42FAA] |
| Q8A8T0 | CUU_3543 | 56.034 | 44 | 1.77E-84 | 273 | GDSL-like protein [Bacteroides vulgatus PC510] |
| Q8A8T0 | HMPREF9011_00854 | 56.034 | 44 | 1.77E-84 | 272 | xylanase [Bacteroides sp. 3_1_40A] |
| Q8A8T0 | BSFG_01659 | 56.034 | 44 | 1.83E-84 | 272 | GDSL-like protein [Bacteroides sp. 4_3_47FAA] |
| Q8A8T0 | BACDOR_01226 | 56.034 | 44 | 2.01E-84 | 273 | GDSL-like protein [Bacteroides dorei DSM 17855] |
| Q8A8T0 | BSEG_03287 | 49.638 | 53 | 4.79E-84 | 275 | hypothetical protein [Bacteroides dorei 5_1_36/D4] |
| Q8A8T0 | HMPREF9447_00467 | 59.483 | 43 | 2.05E-83 | 263 | hypothetical protein [Bacteroides oleiciplenus YIT 12058] |
| Q8A8T0 | HMPREF9450_00844 | 46.503 | 54 | 4.99E-83 | 263 | hypothetical protein [Alistipes indistinctus YIT 12060] |
| Q8A8T0 | HMPREF0101_01286 | 52.838 | 44 | 2.97E-81 | 257 | hypothetical protein [Bacteroides sp. 2_1_16] |
| Q8A8T0 | BSHG_2414 | 52.838 | 44 | 3.55E-81 | 256 | hypothetical protein [Bacteroides sp. 3_2_5] |
| Q8A8T0 | HMPREF1033_01171 | 45.035 | 54 | 3.72E-81 | 258 | hypothetical protein [Tannerella sp. 6_1_58FAA_CT1] |
| Q8A8T0 | HMPREF0673_01369 | 47.653 | 54 | 9.57E-81 | 257 | hypothetical protein [Prevotella stercorea DSM 18206] |
| Q8A8T0 | HMPREF1018_02313 | 52.402 | 44 | 1.90E-80 | 254 | hypothetical protein [Bacteroides sp. 2_1_56FAA] |
| Q8A8T0 | HMPREF1475_01707 | 49.43 | 51 | 4.26E-80 | 255 | hypothetical protein [Prevotella oralis HGA0225] |
| Q8A8T0 | BFAG_01779 | 53.478 | 44 | 5.93E-79 | 251 | hypothetical protein [Bacteroides fragilis 3_1_12] |
| Q8A8T0 | AL1_17710 | 48.551 | 52 | 1.62E-78 | 251 | Esterase/lipase [Alistipes shahii WAL 8301] |
| Q8A8T0 | HMPREF9441_02836 | 36.364 | 82 | 9.62E-78 | 256 | GDSL-like protein [Paraprevotella clara YIT 11840] |
| Q8A8T0 | HMPREF9450_02181 | 54.667 | 43 | 2.36E-77 | 246 | hypothetical protein [Alistipes indistinctus YIT 12060] |
| Q8A8T0 | PARMER_02724 | 44.643 | 54 | 7.64E-77 | 246 | hypothetical protein [Parabacteroides merdae ATCC 43184] |
| Q8A8T0 | BSGG_2818 | 49.242 | 51 | 2.83E-75 | 251 | hypothetical protein [Bacteroides sp. D2] |
| Q8A8T0 | HMPREF9442_00054 | 44.056 | 55 | 4.51E-75 | 249 | GDSL-like protein [Paraprevotella xylaniphila YIT 11841] |
| Q8A8T0 | PRABACTJOHN_00674 | 43.816 | 54 | 7.12E-73 | 236 | hypothetical protein [Parabacteroides johnsonii DSM 18315] |
| Q8A8T0 | PREVCOP_03544 | 50 | 43 | 1.20E-69 | 228 | hypothetical protein [Prevotella copri DSM 18205] |
| Q8A8T0 | HMPREF9450_01136 | 46.992 | 50 | 3.69E-69 | 228 | hypothetical protein [Alistipes indistinctus YIT 12060] |
| Q8A8T0 | HMPREF9442_02977 | 49.778 | 43 | 2.44E-68 | 233 | hypothetical protein [Paraprevotella xylaniphila YIT 11841] |
| Q8A8T0 | HMPREF1017_03697 | 45.064 | 45 | 3.03E-68 | 224 | hypothetical protein [Bacteroides ovatus 3_8_47FAA] |
| Q8A8T0 | HMPREF9010_00954 | 45.064 | 45 | 5.05E-68 | 223 | xylanase [Bacteroides sp. 3_1_23] |
| Q8A8T0 | FMAG_01345 | 38.413 | 56 | 8.05E-68 | 224 | pectin acetylesterase [Fusobacterium mortiferum ATCC 9817] |
| Q8A8T0 | HMPREF1475_01918 | 42.254 | 54 | 3.42E-67 | 228 | hypothetical protein [Prevotella oralis HGA0225] |
| Q8A8T0 | HMPREF9007_02455 | 45.887 | 45 | 5.11E-67 | 220 | xylanase [Bacteroides sp. 1_1_14] |
| Q8A8T0 | BSIG_0607 | 45.455 | 45 | 2.07E-66 | 219 | hypothetical protein [Bacteroides sp. 1_1_6] |
| Q8A8T0 | BACCELL_02533 | 39.223 | 54 | 3.92E-62 | 208 | hypothetical protein [Bacteroides cellulosilyticus DSM 14838] |
| Q8A8T0 | HMPREF1033_02075 | 41.6 | 47 | 3.01E-58 | 199 | hypothetical protein [Tannerella sp. 6_1_58FAA_CT1] |
| Q8A8T0 | HMPREF1532_00940 | 37.838 | 54 | 3.59E-57 | 196 | hypothetical protein [Bacteroides salyersiae WAL 10018 = DSM 18765 = JCM 12988] |
| Q8A8T0 | HMPREF1214_01385 | 37.415 | 53 | 1.39E-54 | 189 | hypothetical protein [Bacteroides sp. HPS0048] |
| Q8A8T0 | HMPREF9465_00834 | 37.786 | 50 | 4.12E-52 | 182 | hypothetical protein [Sutterella wadsworthensis 2_1_59BFAA] |
| Q8A8T0 | PARMER_04255 | 38.264 | 55 | 4.61E-52 | 182 | hypothetical protein [Parabacteroides merdae ATCC 43184] |
| Q8A8T0 | HMPREF9428_01428 | 38.023 | 46 | 4.19E-51 | 180 | hypothetical protein [Citrobacter freundii 4_7_47CFAA] |
| Q8A8T0 | HMPREF0105_2554 | 39.044 | 47 | 8.19E-51 | 178 | hypothetical protein [Bacteroides sp. 3_1_33FAA] |
| Q8A8T0 | PRABACTJOHN_03370 | 37.621 | 55 | 1.25E-50 | 179 | hypothetical protein [Parabacteroides johnsonii DSM 18315] |
| Q8A8T0 | BSBG_02666 | 39.044 | 47 | 2.12E-50 | 177 | hypothetical protein [Bacteroides sp. 9_1_42FAA] |
| Q8A8T0 | BSFG_01217 | 38.645 | 47 | 2.81E-50 | 177 | hypothetical protein [Bacteroides sp. 4_3_47FAA] |
| Q8A8T0 | CSAG_01440 | 39.827 | 42 | 3.42E-50 | 178 | hypothetical protein [Citrobacter sp. 30_2] |

| | | | | | | |
|---|---|---|---|---|---|---|
| Q8A8T0 | BSEG_02355 | 39.044 | 47 | 4.24E-50 | 177 | hypothetical protein [Bacteroides dorei 5_1_36/D4] |
| Q8A8T0 | BACDOR_02642 | 39.044 | 47 | 4.24E-50 | 177 | hypothetical protein [Bacteroides dorei DSM 17855] |
| Q8A8T0 | CUU_3759 | 38.645 | 47 | 5.01E-50 | 177 | conserved hypothetical protein [Bacteroides vulgatus PC510] |
| Q8A8T0 | HMPREF9011_01328 | 38.645 | 47 | 5.01E-50 | 177 | xylanase [Bacteroides sp. 3_1_40A] |
| Q8A8T0 | HMPREF0103_0001 | 36.928 | 54 | 2.71E-49 | 175 | hypothetical protein [Bacteroides sp. 2_1_33B] |
| Q8A8T0 | HMPREF9412_3022 | 40.833 | 44 | 3.36E-49 | 174 | hypothetical protein [Paenibacillus sp. HGF5] |
| Q8A8T0 | HMPREF0106_03199 | 37.5 | 54 | 4.33E-49 | 175 | pectin acetylesterase [Bacteroides sp. D22] |
| Q8A8T0 | HMPREF0619_01988 | 36.928 | 54 | 4.55E-49 | 175 | hypothetical protein [Parabacteroides sp. D13] |
| Q8A8T0 | HMPREF9008_03993 | 36.928 | 54 | 4.55E-49 | 175 | hypothetical protein [Parabacteroides sp. 20_3] |
| Q8A8T0 | HMPREF0104_03640 | 36.928 | 54 | 4.55E-49 | 175 | pectin acetylesterase [Bacteroides sp. 3_1_19] |
| Q8A8T0 | BSEG_01983 | 36.333 | 53 | 4.62E-47 | 169 | pectin acetylesterase family protein [Bacteroides dorei 5_1_36/D4] |
| Q8A8T0 | HMPREF9449_02785 | 40 | 43 | 1.44E-45 | 165 | hypothetical protein [Odoribacter laneus YIT 12061] |
| Q8A8T0 | HMPREF0103_3361 | 37.238 | 44 | 2.42E-45 | 164 | hypothetical protein [Bacteroides sp. 2_1_33B] |
| Q8A8T0 | HMPREF0619_03864 | 37.657 | 44 | 4.81E-45 | 164 | hypothetical protein [Parabacteroides sp. D13] |
| Q8A8T0 | PRABACTJOHN_02814 | 39.095 | 45 | 7.48E-45 | 163 | hypothetical protein [Parabacteroides johnsonii DSM 18315] |
| Q8A8T0 | HMPREF0104_03562 | 37.657 | 44 | 7.75E-45 | 163 | xylanase [Bacteroides sp. 3_1_19] |
| Q8A8T0 | HMPREF9008_03396 | 36.82 | 44 | 1.58E-44 | 162 | hypothetical protein [Parabacteroides sp. 20_3] |
| Q8A8T0 | HMPREF1207_04297 | 35.424 | 50 | 9.66E-44 | 162 | hypothetical protein [Paenibacillus sp. HGH0039] |
| Q8A8T0 | HMPREF9413_4974 | 35.424 | 50 | 9.66E-44 | 162 | hypothetical protein [Paenibacillus sp. HGF7] |
| Q8A8T0 | HMPREF9456_00347 | 36.638 | 43 | 1.30E-43 | 160 | hypothetical protein [Dysgonomonas mossii DSM 22836] |
| Q8A8T0 | HMPREF9447_02322 | 37.919 | 54 | 4.00E-43 | 159 | hypothetical protein [Bacteroides oleiciplenus YIT 12058] |
| Q8A8T0 | HMPREF9544_00250 | 34.749 | 47 | 1.36E-42 | 158 | hypothetical protein [Escherichia coli MS 153-1] |
| Q8A8T0 | HMPREF9531_00350 | 34.749 | 47 | 1.36E-42 | 158 | hypothetical protein [Escherichia coli MS 45-1] |
| Q8A8T0 | ESCG_00402 | 35.317 | 46 | 1.60E-42 | 157 | hypothetical protein [Escherichia sp. 1_1_43] |
| Q8A8T0 | HMPREF9534_01084 | 35.317 | 46 | 1.60E-42 | 157 | hypothetical protein [Escherichia coli MS 69-1] |
| Q8A8T0 | HMPREF9552_00013 | 35.317 | 46 | 2.03E-42 | 157 | hypothetical protein [Escherichia coli MS 198-1] |
| Q8A8T0 | HMPREF9455_03230 | 34.599 | 44 | 4.54E-42 | 155 | hypothetical protein [Dysgonomonas gadei ATCC BAA-286] |
| Q8A8T0 | HMPREF9442_00780 | 35.985 | 48 | 5.75E-42 | 155 | hypothetical protein [Paraprevotella xylaniphila YIT 11841] |
| Q8A8T0 | BACPEC_02489 | 32.787 | 51 | 8.45E-42 | 155 | hypothetical protein [Clostridiales [Bacteroides] pectinophilus ATCC 43243] |
| Q8A8T0 | PARMER_03974 | 37.931 | 43 | 1.32E-40 | 151 | hypothetical protein [Parabacteroides merdae ATCC 43184] |
| Q8A8T0 | HMPREF1475_00142 | 35.897 | 43 | 2.20E-39 | 148 | hypothetical protein [Prevotella oralis HGA0225] |
| Q8A8T0 | HMPREF0673_01326 | 38.397 | 43 | 3.46E-39 | 148 | hypothetical protein [Prevotella stercorea DSM 18206] |
| Q8A8T0 | HMPREF9441_01409 | 36.975 | 43 | 1.42E-38 | 146 | hypothetical protein [Paraprevotella clara YIT 11840] |
| Q8A8T0 | RUM_12320 | 37.559 | 39 | 3.40E-36 | 140 | Esterase/lipase [Ruminococcus champanellensis 18P13 = JCM 17042] |
| Q8A8T0 | ESCG_00405 | 36.207 | 44 | 1.01E-35 | 138 | hypothetical protein [Escherichia sp. 1_1_43] |
| Q8A8T0 | HMPREF9534_01087 | 36.207 | 44 | 1.09E-35 | 138 | hypothetical protein [Escherichia coli MS 69-1] |
| Q8A8T0 | HMPREF9552_00010 | 36.207 | 44 | 1.11E-35 | 138 | hypothetical protein [Escherichia coli MS 198-1] |
| Q8A8T0 | HMPREF9544_00247 | 36.207 | 44 | 1.11E-35 | 138 | hypothetical protein [Escherichia coli MS 153-1] |
| Q8A8T0 | HMPREF9531_00353 | 36.207 | 44 | 1.11E-35 | 138 | hypothetical protein [Escherichia coli MS 45-1] |
| Q8A8T0 | HMPREF9420_0224 | 34 | 46 | 1.56E-34 | 135 | hypothetical protein [Prevotella salivae DSM 15606] |
| Q8A8T0 | PREVCOP_03849 | 33.083 | 49 | 2.94E-34 | 134 | hypothetical protein [Prevotella copri DSM 18205] |
| Q8A8T0 | HMPREF9420_2725 | 35.391 | 43 | 3.66E-33 | 131 | hypothetical protein [Prevotella salivae DSM 15606] |
| Q8A8T0 | BACPEC_00823 | 32.453 | 45 | 1.18E-32 | 130 | hypothetical protein [Clostridiales [Bacteroides] pectinophilus ATCC 43243] |
| Q8A8T0 | HMPREF0201_02268 | 32.932 | 45 | 4.41E-32 | 129 | Tat pathway signal sequence domain protein [Cedecea davisae DSM 4568] |
| Q8A8T0 | HMPREF9473_01818 | 33.469 | 43 | 1.60E-31 | 126 | hypothetical protein [Lachnospiraceae [Clostridium] hathewayi WAL-18680] |
| Q8A8H5 | BSIG_0461 | 100 | 100 | 0 | 959 | hypothetical protein [Bacteroides sp. 1_1_6] |
| Q8A8H5 | HMPREF9007_04460 | 99.568 | 100 | 0 | 956 | xylanase [Bacteroides sp. 1_1_14] |
| Q8A8H5 | BSGG_0323 | 87.364 | 99 | 0 | 838 | hypothetical protein [Bacteroides sp. D2] |
| Q8A8H5 | CUY_0871 | 85.839 | 99 | 0 | 836 | GDSL-like protein [Bacteroides ovatus SD CMC 3f] |
| Q8A8H5 | BSCG_02598 | 85.621 | 99 | 0 | 834 | GDSL-like protein [Bacteroides sp. 2_2_4] |
| Q8A8H5 | HMPREF1017_03830 | 87.364 | 99 | 0 | 834 | hypothetical protein [Bacteroides ovatus 3_8_47FAA] |
| Q8A8H5 | HMPREF0127_03401 | 85.621 | 99 | 0 | 834 | hypothetical protein [Bacteroides sp. 1_1_30] |
| Q8A8H5 | BXY_07760 | 85.621 | 99 | 0 | 834 | Esterase/lipase [Bacteroides xylanisolvens XB1A] |
| Q8A8H5 | HMPREF0102_02833 | 85.621 | 99 | 0 | 834 | GDSL-like protein [Bacteroides sp. 2_1_22] |
| Q8A8H5 | BSAG_01415 | 85.621 | 99 | 0 | 834 | GDSL-like protein [Bacteroides sp. D1] |
| Q8A8H5 | HMPREF0106_01451 | 85.621 | 99 | 0 | 834 | xylanase [Bacteroides sp. D22] |
| Q8A8H5 | CW3_0258 | 85.621 | 99 | 0 | 834 | GDSL-like protein [Bacteroides xylanisolvens SD CC 1b] |
| Q8A8H5 | CW1_0908 | 85.621 | 99 | 0 | 834 | GDSL-like protein [Bacteroides xylanisolvens SD CC 2a] |
| Q8A8H5 | HMPREF9010_00809 | 86.71 | 99 | 0 | 831 | xylanase [Bacteroides sp. 3_1_23] |
| Q8A8H5 | BACOVA_03983 | 86.057 | 99 | 0 | 823 | GDSL-like protein [Bacteroides ovatus ATCC 8483] |

| | | | | | | |
|---|---|---|---|---|---|---|
| Q8A8H5 | BACFIN_07184 | 85.403 | 99 | 0 | 816 | GDSL-like protein [Bacteroides finegoldii DSM 17565] |
| Q8A8H5 | BACDOR_01226 | 83.224 | 99 | 0 | 810 | GDSL-like protein [Bacteroides dorei DSM 17855] |
| Q8A8H5 | CUU_3543 | 83.261 | 99 | 0 | 809 | GDSL-like protein [Bacteroides vulgatus PC510] |
| Q8A8H5 | HMPREF0105_3767 | 83.224 | 99 | 0 | 808 | GDSL-like protein [Bacteroides sp. 3_1_33FAA] |
| Q8A8H5 | BSEG_03737 | 83.224 | 99 | 0 | 808 | GDSL-like protein [Bacteroides dorei 5_1_36/D4] |
| Q8A8H5 | BSBG_04066 | 83.224 | 99 | 0 | 808 | GDSL-like protein [Bacteroides sp. 9_1_42FAA] |
| Q8A8H5 | BSFG_01659 | 82.826 | 99 | 0 | 806 | GDSL-like protein [Bacteroides sp. 4_3_47FAA] |
| Q8A8H5 | HMPREF9011_00854 | 84.773 | 95 | 0 | 793 | xylanase [Bacteroides sp. 3_1_40A] |
| Q8A8H5 | HMPREF1214_02158 | 71.983 | 95 | 0 | 713 | hypothetical protein [Bacteroides sp. HPS0048] |
| Q8A8H5 | BACCAC_01030 | 86.822 | 84 | 0 | 699 | hypothetical protein [Bacteroides caccae ATCC 43185] |
| Q8A8H5 | HMPREF9442_00054 | 54.884 | 92 | 7.17E-167 | 483 | GDSL-like protein [Paraprevotella xylaniphila YIT 11841] |
| Q8A8H5 | HMPREF9441_02836 | 54.419 | 92 | 5.87E-166 | 480 | GDSL-like protein [Paraprevotella clara YIT 11840] |
| Q8A8H5 | HMPREF1475_01918 | 46.07 | 98 | 3.11E-139 | 412 | hypothetical protein [Prevotella oralis HGA0225] |
| Q8A8H5 | HMPREF1532_03226 | 66.327 | 58 | 1.04E-136 | 399 | hypothetical protein [Bacteroides salyersiae WAL 10018 = DSM 18765 = JCM 12988] |
| Q8A8H5 | HMPREF1214_02379 | 56.643 | 57 | 1.29E-113 | 340 | hypothetical protein [Bacteroides sp. HPS0048] |
| Q8A8H5 | HMPREF1475_01707 | 61.277 | 51 | 8.02E-101 | 306 | hypothetical protein [Prevotella oralis HGA0225] |
| Q8A8H5 | HMPREF9420_1566 | 60.944 | 50 | 9.44E-100 | 305 | hypothetical protein [Prevotella salivae DSM 15606] |
| Q8A8H5 | BFAG_01517 | 59.829 | 50 | 5.28E-99 | 301 | hypothetical protein [Bacteroides fragilis 3_1_12] |
| Q8A8H5 | HMPREF9420_0820 | 42.564 | 83 | 1.12E-97 | 305 | GDSL-like protein [Prevotella salivae DSM 15606] |
| Q8A8H5 | HMPREF1017_02201 | 58.009 | 50 | 3.85E-97 | 298 | hypothetical protein [Bacteroides ovatus 3_8_47FAA] |
| Q8A8H5 | HMPREF9448_02673 | 51.748 | 56 | 7.14E-95 | 291 | hypothetical protein [Barnesiella intestinihominis YIT 11860] |
| Q8A8H5 | BACCOP_02646 | 57.143 | 53 | 1.93E-93 | 288 | hypothetical protein [Bacteroides coprocola DSM 17136] |
| Q8A8H5 | PREVCOP_03544 | 54.167 | 54 | 2.03E-91 | 282 | hypothetical protein [Prevotella copri DSM 18205] |
| Q8A8H5 | HMPREF1018_02143 | 56.838 | 50 | 2.82E-91 | 282 | hypothetical protein [Bacteroides sp. 2_1_56FAA] |
| Q8A8H5 | HMPREF0673_01369 | 56.14 | 49 | 1.03E-90 | 281 | hypothetical protein [Prevotella stercorea DSM 18206] |
| Q8A8H5 | HMPREF0101_00969 | 56.652 | 50 | 1.97E-90 | 280 | hypothetical protein [Bacteroides sp. 2_1_16] |
| Q8A8H5 | HMPREF0969_03517 | 54.936 | 50 | 1.63E-88 | 275 | hypothetical protein [Bacteroides sp. D20] |
| Q8A8H5 | HMPREF1007_01252 | 54.936 | 50 | 1.63E-88 | 275 | hypothetical protein [Bacteroides sp. 4_1_36] |
| Q8A8H5 | HMPREF9007_04520 | 55.2 | 54 | 1.89E-88 | 282 | xylanase [Bacteroides sp. 1_1_14] |
| Q8A8H5 | BSIG_0525 | 55.2 | 54 | 2.36E-88 | 282 | hypothetical protein [Bacteroides sp. 1_1_6] |
| Q8A8H5 | BACSTE_00652 | 54.936 | 50 | 6.77E-88 | 274 | hypothetical protein [Bacteroides stercoris ATCC 43183] |
| Q8A8H5 | HMPREF9445_00632 | 54.701 | 50 | 2.27E-87 | 272 | pectin acetylesterase family protein [Bacteroides clarus YIT 12056] |
| Q8A8H5 | BACUNI_02945 | 54.506 | 50 | 4.73E-87 | 272 | hypothetical protein [Bacteroides uniformis ATCC 8492] |
| Q8A8H5 | HMPREF9446_02343 | 54.585 | 49 | 5.60E-87 | 271 | hypothetical protein [Bacteroides fluxus YIT 12057] |
| Q8A8H5 | HMPREF1016_03185 | 54.077 | 50 | 8.39E-87 | 271 | hypothetical protein [Bacteroides eggerthii 1_2_48FAA] |
| Q8A8H5 | BACEGG_02772 | 54.077 | 50 | 8.39E-87 | 271 | hypothetical protein [Bacteroides eggerthii DSM 20697] |
| Q8A8H5 | BACPLE_03269 | 59.908 | 47 | 1.16E-86 | 270 | hypothetical protein [Bacteroides plebeius DSM 17135] |
| Q8A8H5 | BSFG_03001 | 55.983 | 51 | 4.76E-85 | 266 | hypothetical protein [Bacteroides sp. 4_3_47FAA] |
| Q8A8H5 | HMPREF9011_03700 | 55.983 | 51 | 4.76E-85 | 266 | xylanase [Bacteroides sp. 3_1_40A] |
| Q8A8H5 | BACDOR_04358 | 55.983 | 51 | 5.02E-85 | 266 | hypothetical protein [Bacteroides dorei DSM 17855] |
| Q8A8H5 | BSBG_01438 | 55.983 | 51 | 5.02E-85 | 266 | hypothetical protein [Bacteroides sp. 9_1_42FAA] |
| Q8A8H5 | HMPREF0105_0225 | 55.983 | 51 | 5.30E-85 | 266 | hypothetical protein [Bacteroides sp. 3_1_33FAA] |
| Q8A8H5 | BSEG_00286 | 55.983 | 51 | 5.30E-85 | 266 | hypothetical protein [Bacteroides dorei 5_1_36/D4] |
| Q8A8H5 | CUU_1097 | 55.556 | 51 | 7.49E-85 | 266 | conserved hypothetical protein [Bacteroides vulgatus PC510] |
| Q8A8H5 | HMPREF9447_00465 | 50.644 | 50 | 1.66E-83 | 263 | hypothetical protein [Bacteroides oleiciplenus YIT 12058] |
| Q8A8H5 | BACCELL_04558 | 50.644 | 50 | 3.41E-83 | 261 | hypothetical protein [Bacteroides cellulosilyticus DSM 14838] |
| Q8A8H5 | BACINT_02372 | 50.644 | 50 | 4.01E-83 | 261 | hypothetical protein [Bacteroides intestinalis DSM 17393] |
| Q8A8H5 | BSGG_0258 | 52.4 | 54 | 6.94E-83 | 268 | hypothetical protein [Bacteroides sp. D2] |
| Q8A8H5 | BSGG_2818 | 52.344 | 54 | 4.19E-82 | 268 | hypothetical protein [Bacteroides sp. D2] |
| Q8A8H5 | HMPREF0127_00631 | 51.21 | 53 | 5.42E-80 | 261 | hypothetical protein [Bacteroides sp. 1_1_30] |
| Q8A8H5 | HMPREF0102_02896 | 51.21 | 53 | 7.23E-80 | 260 | hypothetical protein [Bacteroides sp. 2_1_22] |
| Q8A8H5 | BSAG_01480 | 51.21 | 53 | 1.08E-79 | 260 | hypothetical protein [Bacteroides sp. D1] |
| Q8A8H5 | CW3_0320 | 51.21 | 53 | 1.08E-79 | 260 | conserved hypothetical protein [Bacteroides xylanisolvens SD CC 1b] |
| Q8A8H5 | CW1_0848 | 51.21 | 53 | 1.08E-79 | 260 | conserved hypothetical protein [Bacteroides xylanisolvens SD CC 2a] |
| Q8A8H5 | BSCG_02541 | 50.806 | 53 | 2.77E-79 | 258 | hypothetical protein [Bacteroides sp. 2_2_4] |
| Q8A8H5 | CUY_0931 | 50.806 | 53 | 7.92E-79 | 258 | conserved hypothetical protein [Bacteroides ovatus SD CMC 3f] |
| Q8A8H5 | HMPREF0106_04667 | 52.155 | 50 | 1.44E-78 | 257 | xylanase [Bacteroides sp. D22] |
| Q8A8H5 | HMPREF1017_03770 | 50.403 | 53 | 2.61E-78 | 256 | hypothetical protein [Bacteroides ovatus 3_8_47FAA] |
| Q8A8H5 | BACOVA_04052 | 50.403 | 53 | 2.93E-78 | 256 | hypothetical protein [Bacteroides ovatus ATCC 8483] |
| Q8A8H5 | HMPREF9010_00870 | 50.403 | 53 | 3.74E-78 | 256 | putative xylanase [Bacteroides sp. 3_1_23] |

| | | | | | | |
|---|---|---|---|---|---|---|
| Q8A8H5 | BACCOPRO_01262 | 54.299 | 48 | 1.16E-76 | 244 | hypothetical protein [Bacteroides coprophilus DSM 18228 = JCM 13818] |
| Q8A8H5 | PRABACTJOHN_00674 | 48.498 | 50 | 1.38E-76 | 244 | hypothetical protein [Parabacteroides johnsonii DSM 18315] |
| Q8A8H5 | PARMER_02724 | 44.944 | 57 | 2.92E-76 | 243 | hypothetical protein [Parabacteroides merdae ATCC 43184] |
| Q8A8H5 | HMPREF9450_01060 | 46.352 | 50 | 3.26E-73 | 236 | hypothetical protein [Alistipes indistinctus YIT 12060] |
| Q8A8H5 | HMPREF1033_01171 | 50.218 | 49 | 1.05E-72 | 234 | hypothetical protein [Tannerella sp. 6_1_58FAA_CT1] |
| Q8A8H5 | BSHG_3584 | 54.46 | 45 | 1.18E-72 | 249 | hypothetical protein [Bacteroides sp. 3_2_5] |
| Q8A8H5 | HMPREF0101_04163 | 54.93 | 45 | 4.47E-72 | 247 | GDSL-like protein [Bacteroides sp. 2_1_16] |
| Q8A8H5 | HMPREF1018_03978 | 54.46 | 45 | 4.90E-72 | 247 | hypothetical protein [Bacteroides sp. 2_1_56FAA] |
| Q8A8H5 | HMPREF9456_02498 | 49.372 | 51 | 9.15E-72 | 232 | hypothetical protein [Dysgonomonas mossii DSM 22836] |
| Q8A8H5 | HMPREF9455_04127 | 47.699 | 51 | 1.45E-71 | 232 | hypothetical protein [Dysgonomonas gadei ATCC BAA-286] |
| Q8A8H5 | HMPREF1033_01363 | 61.458 | 41 | 1.51E-70 | 234 | hypothetical protein [Tannerella sp. 6_1_58FAA_CT1] |
| Q8A8H5 | HMPREF9450_01136 | 48.927 | 50 | 1.76E-65 | 216 | hypothetical protein [Alistipes indistinctus YIT 12060] |
| Q8A8H5 | BACCELL_02534 | 48.804 | 44 | 7.41E-65 | 220 | GDSL-like protein [Bacteroides cellulosilyticus DSM 14838] |
| Q8A8H5 | BACINT_01713 | 51.795 | 42 | 4.26E-64 | 225 | hypothetical protein [Bacteroides intestinalis DSM 17393] |
| Q8A8H5 | HMPREF9450_00844 | 41.463 | 53 | 2.56E-63 | 211 | hypothetical protein [Alistipes indistinctus YIT 12060] |
| Q8A8H5 | BACCELL_02624 | 51.282 | 42 | 3.67E-63 | 223 | GDSL-like protein [Bacteroides cellulosilyticus DSM 14838] |
| Q8A8H5 | HMPREF0103_3520 | 43.83 | 51 | 5.35E-63 | 209 | hypothetical protein [Bacteroides sp. 2_1_33B] |
| Q8A8H5 | HMPREF9008_03075 | 43.404 | 51 | 6.15E-63 | 209 | hypothetical protein [Parabacteroides sp. 20_3] |
| Q8A8H5 | HMPREF0104_01154 | 43.404 | 51 | 4.07E-62 | 206 | pectin acetylesterase [Bacteroides sp. 3_1_19] |
| Q8A8H5 | HMPREF0619_01136 | 43.404 | 51 | 4.48E-62 | 206 | hypothetical protein [Parabacteroides sp. D13] |
| Q8A8H5 | BACCELL_02533 | 40.52 | 55 | 1.11E-61 | 206 | hypothetical protein [Bacteroides cellulosilyticus DSM 14838] |
| Q8A8H5 | BSHG_2414 | 44.444 | 47 | 1.12E-61 | 204 | hypothetical protein [Bacteroides sp. 3_2_5] |
| Q8A8H5 | HMPREF9447_05160 | 50.256 | 42 | 1.20E-61 | 219 | hypothetical protein [Bacteroides oleiciplenus YIT 12058] |
| Q8A8H5 | HMPREF0101_01286 | 44.444 | 47 | 2.60E-61 | 204 | hypothetical protein [Bacteroides sp. 2_1_16] |
| Q8A8H5 | HMPREF1018_02313 | 44.444 | 47 | 4.14E-61 | 203 | hypothetical protein [Bacteroides sp. 2_1_56FAA] |
| Q8A8H5 | BFAG_01779 | 43.981 | 47 | 1.09E-60 | 202 | hypothetical protein [Bacteroides fragilis 3_1_12] |
| Q8A8H5 | AL1_17710 | 44.206 | 50 | 2.35E-60 | 202 | Esterase/lipase [Alistipes shahii WAL 8301] |
| Q8A8H5 | HMPREF9442_02977 | 44.344 | 48 | 6.05E-59 | 206 | hypothetical protein [Paraprevotella xylaniphila YIT 11841] |
| Q8A8H5 | HMPREF9450_02181 | 43.925 | 46 | 1.75E-57 | 194 | hypothetical protein [Alistipes indistinctus YIT 12060] |
| Q8A8H5 | FMAG_01345 | 40.693 | 46 | 7.34E-55 | 189 | pectin acetylesterase [Fusobacterium mortiferum ATCC 9817] |
| Q8A8H5 | HMPREF1017_03697 | 39.474 | 49 | 9.28E-55 | 187 | hypothetical protein [Bacteroides ovatus 3_8_47FAA] |
| Q8A8H5 | HMPREF9448_02731 | 41.667 | 48 | 1.43E-54 | 186 | hypothetical protein [Barnesiella intestinihominis YIT 11860] |
| Q8A8H5 | HMPREF9010_00954 | 39.474 | 49 | 1.93E-54 | 186 | xylanase [Bacteroides sp. 3_1_23] |
| Q8A8H5 | BSIG_0607 | 41.256 | 48 | 3.70E-54 | 186 | hypothetical protein [Bacteroides sp. 1_1_6] |
| Q8A8H5 | HMPREF9007_02455 | 41.256 | 48 | 6.63E-54 | 184 | xylanase [Bacteroides sp. 1_1_14] |
| Q8A8H5 | BSEG_03287 | 45.299 | 50 | 8.55E-52 | 187 | hypothetical protein [Bacteroides dorei 5_1_36/D4] |
| Q8A8H5 | BACCOP_01058 | 46.734 | 43 | 6.65E-51 | 190 | GDSL-like protein [Bacteroides coprocola DSM 17136] |
| Q8A8H5 | BSCG_01719 | 45.098 | 43 | 3.69E-50 | 183 | GDSL-like protein [Bacteroides sp. 2_2_4] |
| Q8A8H5 | HMPREF1017_02929 | 45.098 | 43 | 2.00E-49 | 183 | hypothetical protein [Bacteroides ovatus 3_8_47FAA] |
| Q8A8H5 | BACOVA_04937 | 45.098 | 43 | 2.08E-49 | 183 | GDSL-like protein [Bacteroides ovatus ATCC 8483] |
| Q8A8H5 | HMPREF9010_02721 | 45.098 | 43 | 3.23E-49 | 183 | sialate O-acetylesterase [Bacteroides sp. 3_1_23] |
| Q8A8H5 | BSGG_1462 | 45.098 | 43 | 5.92E-49 | 182 | hypothetical protein [Bacteroides sp. D2] |
| Q8A8H5 | HMPREF1033_00964 | 45.989 | 40 | 1.44E-48 | 170 | hypothetical protein [Tannerella sp. 6_1_58FAA_CT1] |
| Q8A8H5 | HMPREF9007_01631 | 45.128 | 41 | 3.94E-48 | 169 | conserved hypothetical protein [Bacteroides sp. 1_1_14] |
| Q8A8H5 | CUY_2930 | 44.118 | 43 | 5.34E-48 | 179 | GDSL-like protein [Bacteroides ovatus SD CMC 3f] |
| Q8A8H5 | HMPREF9447_01915 | 46.774 | 40 | 6.92E-45 | 172 | hypothetical protein [Bacteroides oleiciplenus YIT 12058] |
| Q8A8H5 | HMPREF1214_01385 | 36.864 | 47 | 1.98E-44 | 161 | hypothetical protein [Bacteroides sp. HPS0048] |
| Q8A8H5 | HMPREF1532_00940 | 38.136 | 47 | 1.12E-43 | 159 | hypothetical protein [Bacteroides salyersiae WAL 10018 = DSM 18765 = JCM 12988] |
| Q8A8H5 | HMPREF1214_04453 | 43.434 | 41 | 1.57E-43 | 166 | hypothetical protein [Bacteroides sp. HPS0048] |
| Q8A8H5 | HMPREF9420_2498 | 42.132 | 41 | 2.66E-43 | 166 | GDSL-like protein [Prevotella salivae DSM 15606] |
| Q8A8H5 | HMPREF1532_04071 | 44.444 | 41 | 1.96E-42 | 162 | hypothetical protein [Bacteroides salyersiae WAL 10018 = DSM 18765 = JCM 12988] |
| Q8A8H5 | HMPREF9447_02322 | 38.889 | 51 | 3.03E-42 | 155 | hypothetical protein [Bacteroides oleiciplenus YIT 12058] |
| Q8A8H5 | HMPREF9455_01371 | 42.553 | 40 | 1.37E-41 | 158 | hypothetical protein [Dysgonomonas gadei ATCC BAA-286] |
| Q8A8H5 | HMPREF9412_3022 | 36.68 | 53 | 3.16E-41 | 152 | hypothetical protein [Paenibacillus sp. HGF5] |
| Q8A8H5 | HMPREF9428_01428 | 35.498 | 46 | 4.31E-40 | 150 | hypothetical protein [Citrobacter freundii 4_7_47CFAA] |
| Q8A8H5 | HMPREF9465_00834 | 35.417 | 51 | 4.74E-40 | 149 | hypothetical protein [Sutterella wadsworthensis 2_1_59BFAA] |
| Q8A8H5 | CSAG_01440 | 35.498 | 46 | 8.45E-40 | 149 | hypothetical protein [Citrobacter sp. 30_2] |
| Q8A8H5 | HMPREF0106_03199 | 35.965 | 45 | 1.73E-39 | 149 | pectin acetylesterase [Bacteroides sp. D22] |
| Q8A8H5 | BSEG_01983 | 36.404 | 45 | 1.20E-38 | 146 | pectin acetylesterase family protein [Bacteroides dorei 5_1_36/D4] |
| Q8A8H5 | HMPREF9552_00013 | 36.957 | 46 | 1.38E-37 | 143 | hypothetical protein [Escherichia coli MS 198-1] |

| | | | | | | |
|---|---|---|---|---|---|---|
| Q8A8H5 | BACCOPRO_02259 | 38.421 | 40 | 1.38E-37 | 150 | GDSL-like protein [Bacteroides coprophilus DSM 18228 = JCM 13818] |
| Q8A8H5 | ESCG_00402 | 36.957 | 46 | 1.83E-37 | 143 | hypothetical protein [Escherichia sp. 1_1_43] |
| Q8A8H5 | HMPREF9534_01084 | 36.957 | 46 | 1.83E-37 | 143 | hypothetical protein [Escherichia coli MS 69-1] |
| Q8A8H5 | HMPREF9544_00250 | 36.957 | 46 | 2.03E-37 | 143 | hypothetical protein [Escherichia coli MS 153-1] |
| Q8A8H5 | HMPREF9531_00350 | 36.957 | 46 | 2.03E-37 | 143 | hypothetical protein [Escherichia coli MS 45-1] |
| Q8A8H5 | BACPLE_00585 | 38.863 | 43 | 2.97E-37 | 149 | GDSL-like protein [Bacteroides plebeius DSM 17135] |
| Q8A8H5 | HMPREF0619_00082 | 37.113 | 41 | 2.27E-36 | 146 | GDSL-like protein [Parabacteroides sp. D13] |
| Q8A8H5 | HMPREF9008_04226 | 36.598 | 41 | 4.27E-36 | 145 | GDSL-like protein [Parabacteroides sp. 20_3] |
| Q8A8H5 | HMPREF1214_02929 | 39.286 | 41 | 1.23E-35 | 144 | sialate O-acetylesterase [Bacteroides sp. HPS0048] |
| Q8A8H5 | PARMER_00328 | 38.679 | 45 | 1.30E-35 | 145 | cyclically-permuted mutarotase family protein [Parabacteroides merdae ATCC 43184] |
| Q8A8H5 | HMPREF0104_00088 | 36.082 | 41 | 1.40E-35 | 144 | sialate O-acetylesterase [Bacteroides sp. 3_1_19] |
| Q8A8H5 | HMPREF9449_01521 | 38 | 41 | 1.49E-35 | 144 | hypothetical protein [Odoribacter laneus YIT 12061] |
| Q8A8H5 | HMPREF1532_00169 | 39.614 | 43 | 1.95E-35 | 144 | hypothetical protein [Bacteroides salyersiae WAL 10018 = DSM 18765 = JCM 12988] |
| Q8A8H5 | HMPREF0103_2214 | 36.082 | 41 | 2.65E-35 | 143 | GDSL-like protein [Bacteroides sp. 2_1_33B] |
| Q8A8H5 | PARMER_04255 | 30.682 | 53 | 7.70E-35 | 135 | hypothetical protein [Parabacteroides merdae ATCC 43184] |
| Q8A8H5 | BACCOPRO_02673 | 36.041 | 41 | 1.93E-34 | 140 | GDSL-like protein [Bacteroides coprophilus DSM 18228 = JCM 13818] |
| Q8A8H5 | HMPREF0103_0001 | 28.517 | 53 | 3.46E-34 | 134 | hypothetical protein [Bacteroides sp. 2_1_33B] |
| Q8A8H5 | HMPREF0619_01988 | 28.517 | 53 | 4.65E-34 | 134 | hypothetical protein [Parabacteroides sp. D13] |
| Q8A8H5 | HMPREF9008_03993 | 28.517 | 53 | 4.65E-34 | 134 | hypothetical protein [Parabacteroides sp. 20_3] |
| Q8A8H5 | HMPREF0104_03640 | 28.517 | 53 | 4.65E-34 | 134 | pectin acetylesterase [Bacteroides sp. 3_1_19] |
| Q8A8H5 | PRABACTJOHN_03370 | 30.682 | 53 | 4.79E-34 | 133 | hypothetical protein [Parabacteroides johnsonii DSM 18315] |
| Q8A8H5 | HMPREF9007_01395 | 37.5 | 42 | 9.55E-34 | 139 | sialate O-acetylesterase [Bacteroides sp. 1_1_14] |
| Q8A8H5 | BSIG_5696 | 37.129 | 42 | 1.94E-33 | 137 | hypothetical protein [Bacteroides sp. 1_1_6] |
| Q8A8H5 | HMPREF0105_1543 | 36.318 | 41 | 3.98E-33 | 137 | GDSL-like protein [Bacteroides sp. 3_1_33FAA] |
| Q8A8H5 | BSBG_00384 | 36.318 | 41 | 3.98E-33 | 137 | GDSL-like protein [Bacteroides sp. 9_1_42FAA] |
| Q8A8H5 | BSFG_00731 | 36.318 | 41 | 4.13E-33 | 137 | GDSL-like protein [Bacteroides sp. 4_3_47FAA] |
| Q8A8H5 | HMPREF9011_03104 | 36.318 | 41 | 4.13E-33 | 137 | sialate O-acetylesterase [Bacteroides sp. 3_1_40A] |
| Q8A8H5 | CUU_3079 | 36.318 | 41 | 4.21E-33 | 137 | GDSL-like protein [Bacteroides vulgatus PC510] |
| Q8A8H5 | BACDOR_00666 | 36.318 | 41 | 4.54E-33 | 137 | GDSL-like protein [Bacteroides dorei DSM 17855] |
| Q8A8H5 | PRABACTJOHN_00552 | 38.021 | 41 | 5.99E-33 | 137 | cyclically-permuted mutarotase family protein [Parabacteroides johnsonii DSM 18315] |
| Q8A8H5 | BFAG_01068 | 42.211 | 41 | 1.31E-32 | 135 | GDSL-like protein [Bacteroides fragilis 3_1_12] |
| Q8A8H5 | BSEG_01307 | 35.821 | 41 | 5.74E-32 | 133 | GDSL-like protein [Bacteroides dorei 5_1_36/D4] |
| Q8A8H5 | HMPREF0984_01372 | 35.644 | 42 | 6.46E-32 | 127 | hypothetical protein [Eubacterium sp. 3_1_31] |
| Q8A8H5 | HMPREF0863_01666 | 35.644 | 42 | 6.46E-32 | 127 | hypothetical protein [Erysipelotrichaceae bacterium 5_2_54FAA] |
| Q8A8H5 | HMPREF0101_00674 | 41.206 | 41 | 8.22E-32 | 133 | GDSL-like protein [Bacteroides sp. 2_1_16] |
| Q8A8H5 | BSHG_4430 | 41.206 | 41 | 9.55E-32 | 133 | hypothetical protein [Bacteroides sp. 3_2_5] |
| Q8A8H5 | HMPREF1018_01836 | 41.206 | 41 | 9.55E-32 | 133 | hypothetical protein [Bacteroides sp. 2_1_56FAA] |
| Q8A8H5 | BLAHAN_06726 | 35.323 | 41 | 5.96E-31 | 124 | hypothetical protein [Blautia hansenii DSM 20583] |
| Q8A8H5 | ESCG_00405 | 34.513 | 48 | 9.90E-31 | 124 | hypothetical protein [Escherichia sp. 1_1_43] |
| Q8A7W6 | BACCAC_02472 | 90.884 | 100 | 0 | 2096 | WD40-like protein [Bacteroides caccae ATCC 43185] |
| Q8A7W6 | BSCG_00162 | 89.544 | 100 | 0 | 2070 | WD40-like protein [Bacteroides sp. 2_2_4] |
| Q8A7W6 | HMPREF0102_03876 | 88.114 | 100 | 0 | 2069 | WD40-like protein [Bacteroides sp. 2_1_22] |
| Q8A7W6 | BSAG_02467 | 88.114 | 100 | 0 | 2069 | WD40-like protein [Bacteroides sp. D1] |
| Q8A7W6 | CW3_0633 | 88.114 | 100 | 0 | 2069 | dipeptidyl peptidase IV N-terminal domain protein [Bacteroides xylanisolvens SD CC 1b] |
| Q8A7W6 | CW1_2522 | 88.114 | 100 | 0 | 2069 | dipeptidyl peptidase IV N-terminal domain protein [Bacteroides xylanisolvens SD CC 2a] |
| Q8A7W6 | BACOVA_00815 | 89.455 | 100 | 0 | 2067 | WD40-like protein [Bacteroides ovatus ATCC 8483] |
| Q8A7W6 | HMPREF1017_01769 | 89.187 | 100 | 0 | 2062 | hypothetical protein [Bacteroides ovatus 3_8_47FAA] |
| Q8A7W6 | HMPREF9010_02092 | 89.366 | 100 | 0 | 2061 | MdsD protein [Bacteroides sp. 3_1_23] |
| Q8A7W6 | HMPREF0127_03060 | 90.472 | 100 | 0 | 2054 | hypothetical protein [Bacteroides sp. 1_1_30] |
| Q8A7W6 | CUY_4364 | 90.381 | 100 | 0 | 2045 | WD40-like protein [Bacteroides ovatus SD CMC 3f] |
| Q8A7W6 | BSGG_3438 | 90.381 | 100 | 0 | 2043 | hypothetical protein [Bacteroides sp. D2] |
| Q8A7W6 | HMPREF0106_02235 | 89.927 | 100 | 0 | 2031 | MdsD protein [Bacteroides sp. D22] |
| Q8A7W6 | BXY_43820 | 90.353 | 98 | 0 | 2001 | Periplasmic protease [Bacteroides xylanisolvens XB1A] |
| Q8A7W6 | BACFIN_05030 | 86.322 | 100 | 0 | 1946 | WD40-like protein [Bacteroides finegoldii DSM 17565] |
| Q8A7W6 | BACDOR_02023 | 77.566 | 99 | 0 | 1805 | WD40-like protein [Bacteroides dorei DSM 17855] |
| Q8A7W6 | BSBG_03541 | 77.475 | 99 | 0 | 1802 | WD40-like protein [Bacteroides sp. 9_1_42FAA] |
| Q8A7W6 | HMPREF0105_1915 | 77.384 | 99 | 0 | 1802 | WD40-like protein [Bacteroides sp. 3_1_33FAA] |
| Q8A7W6 | CUU_0059 | 77.475 | 99 | 0 | 1800 | WD40-like protein [Bacteroides vulgatus PC510] |
| Q8A7W6 | BSFG_00259 | 77.384 | 99 | 0 | 1800 | WD40-like protein [Bacteroides sp. 4_3_47FAA] |
| Q8A7W6 | HMPREF9011_00341 | 77.384 | 99 | 0 | 1800 | peptidase/protease [Bacteroides sp. 3_1_40A] |

| | | | | | | |
|---|---|---|---|---|---|---|
| Q8A7W6 | HMPREF9445_02888 | 76.294 | 99 | 0 | 1793 | WD40-like protein [Bacteroides clarus YIT 12056] |
| Q8A7W6 | HMPREF0969_01045 | 77.252 | 99 | 0 | 1785 | WD40-like protein [Bacteroides sp. D20] |
| Q8A7W6 | BSHG_1511 | 76.337 | 99 | 0 | 1774 | hypothetical protein [Bacteroides sp. 3_2_5] |
| Q8A7W6 | HMPREF1018_00121 | 76.337 | 99 | 0 | 1774 | hypothetical protein [Bacteroides sp. 2_1_56FAA] |
| Q8A7W6 | HMPREF0619_01939 | 76.385 | 99 | 0 | 1772 | WD40-like protein [Parabacteroides sp. D13] |
| Q8A7W6 | HMPREF0101_02080 | 76.247 | 99 | 0 | 1772 | WD40-like protein [Bacteroides sp. 2_1_16] |
| Q8A7W6 | HMPREF0104_02956 | 76.385 | 99 | 0 | 1771 | MdsD protein [Bacteroides sp. 3_1_19] |
| Q8A7W6 | HMPREF9008_00455 | 76.294 | 99 | 0 | 1771 | WD40-like protein [Parabacteroides sp. 20_3] |
| Q8A7W6 | HMPREF0103_0682 | 76.203 | 99 | 0 | 1769 | WD40-like protein [Bacteroides sp. 2_1_33B] |
| Q8A7W6 | BACUNI_03723 | 76.545 | 99 | 0 | 1769 | WD40-like protein [Bacteroides uniformis ATCC 8492] |
| Q8A7W6 | HMPREF1007_03343 | 76.545 | 99 | 0 | 1769 | hypothetical protein [Bacteroides sp. 4_1_36] |
| Q8A7W6 | HMPREF9446_01471 | 76.273 | 99 | 0 | 1768 | WD40-like protein [Bacteroides fluxus YIT 12057] |
| Q8A7W6 | BFAG_02921 | 76.282 | 99 | 0 | 1764 | WD40-like protein [Bacteroides fragilis 3_1_12] |
| Q8A7W6 | BACSTE_03395 | 75.545 | 99 | 0 | 1751 | WD40-like protein [Bacteroides stercoris ATCC 43183] |
| Q8A7W6 | HMPREF1016_01431 | 74.478 | 99 | 0 | 1743 | hypothetical protein [Bacteroides eggerthii 1_2_48FAA] |
| Q8A7W6 | BACEGG_00785 | 74.478 | 99 | 0 | 1743 | WD40-like protein [Bacteroides eggerthii DSM 20697] |
| Q8A7W6 | BACINT_04582 | 73.43 | 99 | 0 | 1726 | WD40-like protein [Bacteroides intestinalis DSM 17393] |
| Q8A7W6 | BACCELL_02934 | 73.564 | 99 | 0 | 1719 | WD40-like protein [Bacteroides cellulosilyticus DSM 14838] |
| Q8A7W6 | HMPREF9447_04309 | 71.793 | 99 | 0 | 1681 | hypothetical protein [Bacteroides oleiciplenus YIT 12058] |
| Q8A7W6 | BACCOPRO_00170 | 73.545 | 99 | 0 | 1664 | WD40-like protein [Bacteroides coprophilus DSM 18228 = JCM 13818] |
| Q8A7W6 | BACCOP_00819 | 71.416 | 99 | 0 | 1655 | WD40-like protein [Bacteroides coprocola DSM 17136] |
| Q8A7W6 | BACPLE_00276 | 70.327 | 99 | 0 | 1615 | WD40-like protein [Bacteroides plebeius DSM 17135] |
| Q8A7W6 | HMPREF9449_01222 | 62.285 | 99 | 0 | 1394 | hypothetical protein [Odoribacter laneus YIT 12061] |
| Q8A7W6 | PRABACTJOHN_04417 | 48.367 | 99 | 0 | 1026 | WD40-like protein [Parabacteroides johnsonii DSM 18315] |
| Q8A7W6 | HMPREF9449_02001 | 46.801 | 99 | 0 | 1018 | hypothetical protein [Odoribacter laneus YIT 12061] |
| Q8A7W6 | BSGG_3550 | 48.783 | 99 | 0 | 1007 | hypothetical protein [Bacteroides sp. D2] |
| Q8A7W6 | BACOVA_00608 | 48.783 | 99 | 0 | 1007 | WD40-like protein [Bacteroides ovatus ATCC 8483] |
| Q8A7W6 | CUY_3536 | 48.693 | 99 | 0 | 1006 | WD40-like protein [Bacteroides ovatus SD CMC 3f] |
| Q8A7W6 | BSCG_05082 | 48.693 | 99 | 0 | 1006 | WD40-like protein [Bacteroides sp. 2_2_4] |
| Q8A7W6 | HMPREF9010_03364 | 48.693 | 99 | 0 | 1006 | MdsD protein [Bacteroides sp. 3_1_23] |
| Q8A7W6 | PARMER_03129 | 48 | 99 | 0 | 1004 | WD40-like protein [Parabacteroides merdae ATCC 43184] |
| Q8A7W6 | BSHG_4122 | 50.225 | 99 | 0 | 1003 | hypothetical protein [Bacteroides sp. 3_2_5] |
| Q8A7W6 | HMPREF0101_04280 | 50.135 | 99 | 0 | 1002 | WD40-like protein [Bacteroides sp. 2_1_16] |
| Q8A7W6 | HMPREF1018_04474 | 50.181 | 99 | 0 | 999 | hypothetical protein [Bacteroides sp. 2_1_56FAA] |
| Q8A7W6 | BACEGG_00077 | 48.378 | 99 | 0 | 998 | WD40-like protein [Bacteroides eggerthii DSM 20697] |
| Q8A7W6 | HMPREF1016_01913 | 48.378 | 99 | 0 | 996 | hypothetical protein [Bacteroides eggerthii 1_2_48FAA] |
| Q8A7W6 | BFAG_04470 | 50 | 98 | 0 | 993 | WD40-like protein [Bacteroides fragilis 3_1_12] |
| Q8A7W6 | BACCAC_02584 | 48.422 | 99 | 0 | 993 | WD40-like protein [Bacteroides caccae ATCC 43185] |
| Q8A7W6 | HMPREF9445_02665 | 48.602 | 99 | 0 | 988 | WD40-like protein [Bacteroides clarus YIT 12056] |
| Q8A7W6 | HMPREF9446_02487 | 48.294 | 99 | 0 | 987 | WD40-like protein [Bacteroides fluxus YIT 12057] |
| Q8A7W6 | BACPLE_02804 | 47.702 | 98 | 0 | 984 | WD40-like protein [Bacteroides plebeius DSM 17135] |
| Q8A7W6 | BSFG_00797 | 46.883 | 99 | 0 | 983 | WD40-like protein [Bacteroides sp. 4_3_47FAA] |
| Q8A7W6 | HMPREF9011_03169 | 46.883 | 99 | 0 | 983 | exported peptidase [Bacteroides sp. 3_1_40A] |
| Q8A7W6 | BACUNI_03133 | 47.705 | 99 | 0 | 979 | WD40-like protein [Bacteroides uniformis ATCC 8492] |
| Q8A7W6 | CUU_2511 | 46.703 | 99 | 0 | 979 | WD40-like protein [Bacteroides vulgatus PC510] |
| Q8A7W6 | HMPREF1007_02020 | 47.705 | 99 | 0 | 979 | hypothetical protein [Bacteroides sp. 4_1_36] |
| Q8A7W6 | HMPREF0104_03074 | 46.494 | 98 | 0 | 979 | MdsD protein [Bacteroides sp. 3_1_19] |
| Q8A7W6 | HMPREF0619_03703 | 46.679 | 98 | 0 | 978 | WD40-like protein [Parabacteroides sp. D13] |
| Q8A7W6 | HMPREF9008_02096 | 46.402 | 98 | 0 | 978 | WD40-like protein [Parabacteroides sp. 20_3] |
| Q8A7W6 | HMPREF0969_02303 | 47.795 | 99 | 0 | 978 | WD40-like protein [Bacteroides sp. D20] |
| Q8A7W6 | HMPREF0103_3260 | 46.402 | 98 | 0 | 974 | WD40-like protein [Bacteroides sp. 2_1_33B] |
| Q8A7W6 | PREVCOP_03747 | 47.005 | 99 | 0 | 960 | WD40-like protein [Prevotella copri DSM 18205] |
| Q8A7W6 | BSBG_00310 | 45.89 | 99 | 0 | 959 | WD40-like protein [Bacteroides sp. 9_1_42FAA] |
| Q8A7W6 | BACDOR_00760 | 45.89 | 99 | 0 | 958 | WD40-like protein [Bacteroides dorei DSM 17855] |
| Q8A7W6 | BSEG_01184 | 45.89 | 99 | 0 | 958 | WD40-like protein [Bacteroides dorei 5_1_36/D4] |
| Q8A7W6 | HMPREF0105_1469 | 45.89 | 99 | 0 | 957 | WD40-like protein [Bacteroides sp. 3_1_33FAA] |
| Q8A7W6 | BACCOPRO_00458 | 45.946 | 100 | 0 | 956 | WD40-like protein [Bacteroides coprophilus DSM 18228 = JCM 13818] |
| Q8A7W6 | HMPREF9455_01942 | 46.432 | 97 | 0 | 941 | hypothetical protein [Dysgonomonas gadei ATCC BAA-286] |
| Q8A7W6 | HMPREF1475_01715 | 44.11 | 99 | 0 | 907 | hypothetical protein [Prevotella oralis HGA0225] |
| Q8A7W6 | HMPREF0619_03697 | 27.336 | 99 | 7.10E-96 | 333 | WD40-like protein [Parabacteroides sp. D13] |

| | | | | | | |
|---|---|---|---|---|---|---|
| Q8A7W6 | HMPREF9008_02090 | 27.48 | 99 | 7.38E-96 | 333 | WD40-like protein [Parabacteroides sp. 20_3] |
| Q8A7W6 | HMPREF0104_03080 | 27.48 | 99 | 1.21E-95 | 333 | exported tricorn protease [Bacteroides sp. 3_1_19] |
| Q8A7W6 | PARMER_03138 | 27.817 | 99 | 1.33E-95 | 333 | WD40-like protein [Parabacteroides merdae ATCC 43184] |
| Q8A7W6 | BACINT_00511 | 27.336 | 99 | 4.20E-95 | 331 | WD40-like protein [Bacteroides intestinalis DSM 17393] |
| Q8A7W6 | BACCELL_01948 | 27.424 | 99 | 4.46E-95 | 331 | WD40-like protein [Bacteroides cellulosilyticus DSM 14838] |
| Q8A7W6 | HMPREF0103_3254 | 27.392 | 99 | 4.52E-95 | 331 | WD40-like protein [Bacteroides sp. 2_1_33B] |
| Q8A7W6 | HMPREF0101_01621 | 27.036 | 99 | 3.06E-91 | 320 | WD40-like protein [Bacteroides sp. 2_1_16] |
| Q8A7W6 | PRABACTJOHN_04411 | 26.984 | 99 | 4.81E-91 | 320 | WD40-like protein [Parabacteroides johnsonii DSM 18315] |
| Q8A7W6 | BFAG_02007 | 26.91 | 99 | 1.47E-90 | 318 | WD40-like protein [Bacteroides fragilis 3_1_12] |
| Q8A7W6 | BSHG_2651 | 26.95 | 99 | 3.32E-90 | 317 | hypothetical protein [Bacteroides sp. 3_2_5] |
| Q8A7W6 | HMPREF1018_02587 | 26.863 | 99 | 5.42E-90 | 317 | hypothetical protein [Bacteroides sp. 2_1_56FAA] |
| Q8A7W6 | HMPREF9447_01774 | 26.136 | 99 | 6.57E-86 | 305 | hypothetical protein [Bacteroides oleiciplenus YIT 12058] |
| Q8A7W6 | CUU_0099 | 25.697 | 99 | 1.67E-83 | 298 | dipeptidyl peptidase IV N-terminal domain protein [Bacteroides vulgatus PC510] |
| Q8A7W6 | BSBG_03582 | 25.849 | 99 | 7.59E-83 | 296 | WD40-like protein [Bacteroides sp. 9_1_42FAA] |
| Q8A7W6 | BACDOR_01984 | 25.849 | 99 | 7.81E-83 | 296 | WD40-like protein [Bacteroides dorei DSM 17855] |
| Q8A7W6 | HMPREF0105_1954 | 25.762 | 99 | 2.53E-82 | 295 | WD40-like protein [Bacteroides sp. 3_1_33FAA] |
| Q8A7W6 | BSEG_03417 | 25.587 | 99 | 3.72E-82 | 294 | WD40-like protein [Bacteroides dorei 5_1_36/D4] |
| Q8A7W6 | BSFG_00296 | 25.348 | 99 | 6.54E-81 | 291 | WD40-like protein [Bacteroides sp. 4_3_47FAA] |
| Q8A7W6 | HMPREF9011_00378 | 25.348 | 99 | 6.54E-81 | 291 | exported protease [Bacteroides sp. 3_1_40A] |
| Q8A7W6 | HMPREF9449_02293 | 24.563 | 99 | 8.60E-74 | 270 | hypothetical protein [Odoribacter laneus YIT 12061] |
| Q8A7W6 | HMPREF1017_01488 | 52.79 | 21 | 2.24E-71 | 241 | hypothetical protein [Bacteroides ovatus 3_8_47FAA] |
| Q8A6Z2 | BSIG_4300 | 100 | 100 | 0 | 2568 | phosphoribosylformylglycinamidine synthase [Bacteroides sp. 1_1_6] |
| Q8A6Z2 | HMPREF9007_04316 | 99.755 | 99 | 0 | 2536 | phosphoribosylformylglycinamidine synthase [Bacteroides sp. 1_1_14] |
| Q8A6Z2 | BACCAC_02754 | 97.812 | 100 | 0 | 2527 | putative phosphoribosylformylglycinamidine synthase [Bacteroides caccae ATCC 43185] |
| Q8A6Z2 | BACOVA_04462 | 96.677 | 100 | 0 | 2498 | putative phosphoribosylformylglycinamidine synthase [Bacteroides ovatus ATCC 8483] |
| Q8A6Z2 | BACFIN_09028 | 96.272 | 100 | 0 | 2496 | putative phosphoribosylformylglycinamidine synthase [Bacteroides finegoldii DSM 17565] |
| Q8A6Z2 | BXY_47110 | 96.515 | 100 | 0 | 2493 | phosphoribosylformylglycinamidine synthase [Bacteroides xylanisolvens XB1A] |
| Q8A6Z2 | HMPREF0106_03358 | 96.515 | 100 | 0 | 2493 | phosphoribosylformylglycinamidine synthase [Bacteroides sp. D22] |
| Q8A6Z2 | CW3_2629 | 96.434 | 100 | 0 | 2492 | putative phosphoribosylformylglycinamidine synthase [Bacteroides xylanisolvens SD CC 1b] |
| Q8A6Z2 | CW1_1264 | 96.434 | 100 | 0 | 2492 | putative phosphoribosylformylglycinamidine synthase [Bacteroides xylanisolvens SD CC 2a] |
| Q8A6Z2 | BSCG_00549 | 96.434 | 100 | 0 | 2492 | putative phosphoribosylformylglycinamidine synthase [Bacteroides sp. 2_2_4] |
| Q8A6Z2 | HMPREF9010_04478 | 96.353 | 100 | 0 | 2491 | phosphoribosylformylglycinamidine synthase [Bacteroides sp. 3_1_23] |
| Q8A6Z2 | HMPREF0127_04209 | 96.434 | 100 | 0 | 2491 | hypothetical protein [Bacteroides sp. 1_1_30] |
| Q8A6Z2 | HMPREF0102_04318 | 96.353 | 100 | 0 | 2491 | putative phosphoribosylformylglycinamidine synthase [Bacteroides sp. 2_1_22] |
| Q8A6Z2 | BSAG_03685 | 96.353 | 100 | 0 | 2491 | putative phosphoribosylformylglycinamidine synthase [Bacteroides sp. D1] |
| Q8A6Z2 | CUY_4914 | 96.11 | 100 | 0 | 2489 | putative phosphoribosylformylglycinamidine synthase [Bacteroides ovatus SD CMC 3f] |
| Q8A6Z2 | BSGG_3111 | 96.272 | 100 | 0 | 2489 | phosphoribosylformylglycinamidine synthase [Bacteroides sp. D2] |
| Q8A6Z2 | HMPREF1017_02173 | 96.402 | 99 | 0 | 2468 | hypothetical protein [Bacteroides ovatus 3_8_47FAA] |
| Q8A6Z2 | HMPREF1532_02473 | 95.138 | 100 | 0 | 2466 | phosphoribosylformylglycinamidine synthase [Bacteroides salyersiae WAL 10018 = DSM 18765 = JCM 12988] |
| Q8A6Z2 | BFAG_03435 | 94.571 | 100 | 0 | 2459 | putative phosphoribosylformylglycinamidine synthase [Bacteroides fragilis 3_1_12] |
| Q8A6Z2 | HMPREF1018_03750 | 94.652 | 100 | 0 | 2458 | hypothetical protein [Bacteroides sp. 2_1_56FAA] |
| Q8A6Z2 | BSHG_2082 | 94.489 | 100 | 0 | 2457 | phosphoribosylformylglycinamidine synthase [Bacteroides sp. 3_2_5] |
| Q8A6Z2 | HMPREF1214_03144 | 94.814 | 100 | 0 | 2457 | phosphoribosylformylglycinamidine synthase [Bacteroides sp. HPS0048] |
| Q8A6Z2 | HMPREF0101_02976 | 94.571 | 100 | 0 | 2454 | putative phosphoribosylformylglycinamidine synthase [Bacteroides sp. 2_1_16] |
| Q8A6Z2 | BACUNI_01183 | 93.031 | 100 | 0 | 2423 | putative phosphoribosylformylglycinamidine synthase [Bacteroides uniformis ATCC 8492] |
| Q8A6Z2 | HMPREF1007_01738 | 93.112 | 100 | 0 | 2422 | phosphoribosylformylglycinamidine synthase [Bacteroides sp. 4_1_36] |
| Q8A6Z2 | HMPREF9446_00149 | 92.869 | 100 | 0 | 2422 | putative phosphoribosylformylglycinamidine synthase [Bacteroides fluxus YIT 12057] |
| Q8A6Z2 | HMPREF0969_00061 | 93.031 | 100 | 0 | 2420 | putative phosphoribosylformylglycinamidine synthase [Bacteroides sp. D20] |
| Q8A6Z2 | BACINT_03163 | 93.031 | 100 | 0 | 2419 | phosphoribosylformylglycinamidine synthase [Bacteroides intestinalis DSM 17393] |
| Q8A6Z2 | BACCELL_05479 | 92.869 | 100 | 0 | 2417 | phosphoribosylformylglycinamidine synthase [Bacteroides cellulosilyticus DSM 14838] |
| Q8A6Z2 | HMPREF9445_02351 | 92.382 | 100 | 0 | 2416 | putative phosphoribosylformylglycinamidine synthase [Bacteroides clarus YIT 12056] |
| Q8A6Z2 | HMPREF9447_05349 | 92.301 | 100 | 0 | 2415 | phosphoribosylformylglycinamidine synthase [Bacteroides oleiciplenus YIT 12058] |
| Q8A6Z2 | BACSTE_03451 | 92.539 | 99 | 0 | 2415 | putative phosphoribosylformylglycinamidine synthase [Bacteroides stercoris ATCC 43183] |
| Q8A6Z2 | BACEGG_01868 | 92.376 | 99 | 0 | 2405 | putative phosphoribosylformylglycinamidine synthase [Bacteroides eggerthii DSM 20697] |
| Q8A6Z2 | HMPREF1016_00027 | 92.226 | 99 | 0 | 2380 | phosphoribosylformylglycinamidine synthase [Bacteroides eggerthii 1_2_48FAA] |
| Q8A6Z2 | BSFG_01436 | 86.386 | 100 | 0 | 2268 | putative phosphoribosylformylglycinamidine synthase [Bacteroides sp. 4_3_47FAA] |
| Q8A6Z2 | HMPREF9011_03482 | 86.386 | 100 | 0 | 2268 | phosphoribosylformylglycinamidine synthase [Bacteroides sp. 3_1_40A] |
| Q8A6Z2 | CUU_1004 | 86.305 | 100 | 0 | 2266 | putative phosphoribosylformylglycinamidine synthase [Bacteroides vulgatus PC510] |
| Q8A6Z2 | HMPREF0105_1666 | 86.305 | 100 | 0 | 2266 | putative phosphoribosylformylglycinamidine synthase [Bacteroides sp. 3_1_33FAA] |
| Q8A6Z2 | BSEG_01433 | 86.305 | 100 | 0 | 2266 | putative phosphoribosylformylglycinamidine synthase [Bacteroides dorei 5_1_36/D4] |

| | | | | | | |
|---|---|---|---|---|---|---|
| Q8A6Z2 | BSBG_00522 | 86.143 | 100 | 0 | 2263 | putative phosphoribosylformylglycinamidine synthase [Bacteroides sp. 9_1_42FAA] |
| Q8A6Z2 | BACDOR_00488 | 86.143 | 100 | 0 | 2261 | putative phosphoribosylformylglycinamidine synthase [Bacteroides dorei DSM 17855] |
| Q8A6Z2 | BACPLE_03254 | 86.143 | 100 | 0 | 2259 | putative phosphoribosylformylglycinamidine synthase [Bacteroides plebeius DSM 17135] |
| Q8A6Z2 | BACCOP_01508 | 85.981 | 100 | 0 | 2258 | putative phosphoribosylformylglycinamidine synthase [Bacteroides coprocola DSM 17136] |
| Q8A6Z2 | HMPREF9442_01312 | 83.617 | 99 | 0 | 2179 | putative phosphoribosylformylglycinamidine synthase [Paraprevotella xylaniphila YIT 11841] |
| Q8A6Z2 | HMPREF9441_02147 | 83.617 | 99 | 0 | 2176 | putative phosphoribosylformylglycinamidine synthase [Paraprevotella clara YIT 11840] |
| Q8A6Z2 | HMPREF0673_00982 | 76.886 | 99 | 0 | 2034 | putative phosphoribosylformylglycinamidine synthase [Prevotella stercorea DSM 18206] |
| Q8A6Z2 | PREVCOP_05841 | 76.632 | 99 | 0 | 2011 | putative phosphoribosylformylglycinamidine synthase [Prevotella copri DSM 18205] |
| Q8A6Z2 | HMPREF9420_2567 | 74.859 | 99 | 0 | 1972 | putative phosphoribosylformylglycinamidine synthase [Prevotella salivae DSM 15606] |
| Q8A6Z2 | HMPREF1475_01150 | 73.739 | 99 | 0 | 1964 | phosphoribosylformylglycinamidine synthase [Prevotella oralis HGA0225] |
| Q8A6Z2 | HMPREF1033_00479 | 73.684 | 100 | 0 | 1949 | phosphoribosylformylglycinamidine synthase [Tannerella sp. 6_1_58FAA_CT1] |
| Q8A6Z2 | HMPREF0103_3073 | 72.764 | 99 | 0 | 1919 | putative phosphoribosylformylglycinamidine synthase [Bacteroides sp. 2_1_33B] |
| Q8A6Z2 | HMPREF0104_03310 | 72.764 | 99 | 0 | 1918 | phosphoribosylformylglycinamidine synthase [Bacteroides sp. 3_1_19] |
| Q8A6Z2 | HMPREF9008_01872 | 72.683 | 99 | 0 | 1917 | putative phosphoribosylformylglycinamidine synthase [Parabacteroides sp. 20_3] |
| Q8A6Z2 | HMPREF0619_03494 | 72.764 | 99 | 0 | 1916 | putative phosphoribosylformylglycinamidine synthase [Parabacteroides sp. D13] |
| Q8A6Z2 | HMPREF9448_01782 | 72.389 | 100 | 0 | 1906 | phosphoribosylformylglycinamidine synthase [Barnesiella intestinihominis YIT 11860] |
| Q8A6Z2 | PARMER_03288 | 72.204 | 100 | 0 | 1901 | putative phosphoribosylformylglycinamidine synthase [Parabacteroides merdae ATCC 43184] |
| Q8A6Z2 | PRABACTJOHN_00145 | 72.123 | 100 | 0 | 1895 | putative phosphoribosylformylglycinamidine synthase [Parabacteroides johnsonii DSM 18315] |
| Q8A6Z2 | HMPREF9456_00888 | 68.618 | 99 | 0 | 1817 | hypothetical protein [Dysgonomonas mossii DSM 22836] |
| Q8A6Z2 | HMPREF9455_00911 | 68.318 | 99 | 0 | 1811 | hypothetical protein [Dysgonomonas gadei ATCC BAA-286] |
| Q8A6Z2 | BACCOPRO_03260 | 87.318 | 78 | 0 | 1787 | putative phosphoribosylformylglycinamidine synthase [Bacteroides coprophilus DSM 18228 = JCM 13818] |
| Q8A6Z2 | HMPREF9450_01192 | 63.098 | 99 | 0 | 1659 | phosphoribosylformylglycinamidine synthase [Alistipes indistinctus YIT 12060] |
| Q8A6Z2 | HMPREF9449_00369 | 62.804 | 99 | 0 | 1634 | phosphoribosylformylglycinamidine synthase [Odoribacter laneus YIT 12061] |
| Q8A6Z2 | OFAG_00255 | 35.217 | 98 | 0 | 697 | phosphoribosylformylglycinamidine synthase [Oxalobacter formigenes HOxBLS] |
| Q8A6Z2 | PROVRUST_06703 | 35.731 | 98 | 0 | 695 | phosphoribosylformylglycinamidine synthase [Providencia rustigianii DSM 4541] |
| Q8A6Z2 | PROSTU_03273 | 36.206 | 98 | 0 | 695 | phosphoribosylformylglycinamidine synthase [Providencia stuartii ATCC 25827] |
| Q8A6Z2 | PROVRETT_09715 | 36.22 | 98 | 0 | 694 | phosphoribosylformylglycinamidine synthase [Providencia rettgeri DSM 1131] |
| Q8A6Z2 | HMPREF9465_01842 | 35.552 | 98 | 0 | 694 | phosphoribosylformylglycinamidine synthase [Sutterella wadsworthensis 2_1_59BFAA] |
| Q8A6Z2 | HMPREF1030_00465 | 36.492 | 98 | 0 | 692 | phosphoribosylformylglycinamidine synthase [Pseudomonas sp. 2_1_26] |
| Q8A6Z2 | PROVALCAL_01718 | 36.285 | 98 | 0 | 692 | phosphoribosylformylglycinamidine synthase [Providencia alcalifaciens DSM 30120] |
| Q8A6Z2 | HMPREF9444_00197 | 36.715 | 96 | 0 | 691 | phosphoribosylformylglycinamidine synthase [Succinatimonas hippei YIT 12066] |
| Q8A6Z2 | HMPREF1311_01848 | 35.765 | 98 | 0 | 687 | phosphoribosylformylglycinamidine synthase [Proteus mirabilis WGLW6] |
| Q8A6Z2 | OFBG_01185 | 34.743 | 98 | 0 | 680 | phosphoribosylformylglycinamidine synthase [Oxalobacter formigenes OXCC13] |
| Q8A6Z2 | HMPREF0864_02337 | 35.06 | 97 | 0 | 676 | phosphoribosylformylglycinamidine synthase [Enterobacteriaceae bacterium 9_2_54FAA] |
| Q8A6Z2 | HMPREF0026_01499 | 36.032 | 96 | 0 | 674 | phosphoribosylformylglycinamidine synthase [Acinetobacter junii SH205] |
| Q8A6Z2 | HMPREF0454_03258 | 34.953 | 98 | 0 | 673 | phosphoribosylformylglycinamidine synthase [Hafnia alvei ATCC 51873] |
| Q8A6Z2 | HMPREF0018_01359 | 35.864 | 96 | 0 | 672 | phosphoribosylformylglycinamidine synthase [Acinetobacter radioresistens SH164] |
| Q8A6Z2 | EDWATA_00951 | 35.518 | 96 | 0 | 668 | phosphoribosylformylglycinamidine synthase [Edwardsiella tarda ATCC 23685] |
| Q8A6Z2 | HMPREF0880_04767 | 35.41 | 98 | 0 | 663 | phosphoribosylformylglycinamidine synthase [Yokenella regensburgei ATCC 43003] |
| Q8A6Z2 | HMPREF1024_04245 | 35.012 | 98 | 0 | 662 | phosphoribosylformylglycinamidine synthase [Klebsiella sp. 4_1_44FAA] |
| Q8A6Z2 | HMPREF0485_00029 | 34.961 | 98 | 0 | 662 | phosphoribosylformylglycinamidine synthase [Klebsiella sp. 1_1_55] |
| Q8A6Z2 | HMPREF9538_03727 | 35.012 | 98 | 0 | 661 | phosphoribosylformylglycinamidine synthase [Klebsiella sp. MS 92-3] |
| Q8A6Z2 | HMPREF1308_03080 | 35.012 | 98 | 0 | 661 | phosphoribosylformylglycinamidine synthase [Klebsiella pneumoniae subsp. pneumoniae WGLW5] |
| Q8A6Z2 | HMPREF9530_00726 | 34.961 | 99 | 0 | 660 | phosphoribosylformylglycinamidine synthase [Escherichia coli MS 21-1] |
| Q8A6Z2 | HMPREF1307_01501 | 35.012 | 98 | 0 | 660 | phosphoribosylformylglycinamidine synthase [Klebsiella pneumoniae subsp. pneumoniae WGLW3] |
| Q8A6Z2 | HMPREF9418_1969 | 34.833 | 96 | 0 | 659 | phosphoribosylformylglycinamidine synthase II [Neisseria macacae ATCC 33926] |
| Q8A6Z2 | HMPREF9551_01380 | 35.09 | 98 | 0 | 659 | phosphoribosylformylglycinamidine synthase [Escherichia coli MS 196-1] |
| Q8A6Z2 | HMPREF9541_05032 | 35.09 | 98 | 0 | 659 | phosphoribosylformylglycinamidine synthase [Escherichia coli MS 116-1] |
| Q8A6Z2 | HMPREF9547_04496 | 35.09 | 98 | 0 | 659 | phosphoribosylformylglycinamidine synthase [Escherichia coli MS 175-1] |
| Q8A6Z2 | HMPREF9543_04802 | 35.09 | 98 | 0 | 659 | phosphoribosylformylglycinamidine synthase [Escherichia coli MS 146-1] |
| Q8A6Z2 | ESBG_00033 | 35.09 | 98 | 0 | 659 | phosphoribosylformylglycinamidine synthase [Escherichia sp. 4_1_40B] |
| Q8A6Z2 | HMPREF9347_01502 | 35.248 | 98 | 0 | 658 | phosphoribosylformylglycinamidine synthase [Escherichia coli MS 124-1] |
| Q8A6Z2 | HMPREF9350_01721 | 35.248 | 98 | 0 | 658 | phosphoribosylformylglycinamidine synthase [Escherichia coli MS 85-1] |
| Q8A6Z2 | HMPREF9536_02346 | 35.248 | 98 | 0 | 658 | phosphoribosylformylglycinamidine synthase [Escherichia coli MS 84-1] |
| Q8A6Z2 | ESCG_04546 | 35.169 | 98 | 0 | 658 | phosphoribosylformylglycinamidine synthase [Escherichia sp. 1_1_43] |
| Q8A6Z2 | HMPREF9540_04374 | 35.09 | 98 | 0 | 657 | phosphoribosylformylglycinamidine synthase [Escherichia coli MS 115-1] |
| Q8A6Z2 | HMPREF9349_03660 | 35.09 | 98 | 0 | 657 | phosphoribosylformylglycinamidine synthase [Escherichia coli MS 79-10] |
| Q8A6Z2 | HMPREF9345_02738 | 35.09 | 98 | 0 | 657 | phosphoribosylformylglycinamidine synthase [Escherichia coli MS 107-1] |
| Q8A6Z2 | ECSE_2845 | 35.09 | 98 | 0 | 657 | phosphoribosylformyl-glycineamide synthase [Escherichia coli SE11] |
| Q8A6Z2 | HMPREF9550_02495 | 35.09 | 98 | 0 | 657 | phosphoribosylformylglycinamidine synthase [Escherichia coli MS 187-1] |

| | | | | | | |
|---|---|---|---|---|---|---|
| Q8A6Z2 | HMPREF9542_01288 | 35.09 | 98 | 0 | 656 | phosphoribosylformylglycinamidine synthase [Escherichia coli MS 117-3] |
| Q8A6Z2 | HMPREF9544_03086 | 35.09 | 98 | 0 | 656 | phosphoribosylformylglycinamidine synthase [Escherichia coli MS 153-1] |
| Q8A6Z2 | HMPREF9346_00577 | 35.09 | 98 | 0 | 656 | phosphoribosylformylglycinamidine synthase [Escherichia coli MS 119-7] |
| Q8A6Z2 | HMPREF9549_01266 | 35.09 | 98 | 0 | 656 | phosphoribosylformylglycinamidine synthase [Escherichia coli MS 185-1] |
| Q8A6Z2 | HMPREF9531_02552 | 35.09 | 98 | 0 | 656 | phosphoribosylformylglycinamidine synthase [Escherichia coli MS 45-1] |
| Q8A6Z2 | HMPREF9553_00437 | 34.783 | 99 | 0 | 656 | phosphoribosylformylglycinamidine synthase [Escherichia coli MS 200-1] |
| Q8A6Z2 | HMPREF0986_03095 | 34.938 | 99 | 0 | 656 | phosphoribosylformylglycinamidine synthase [Escherichia coli 4_1_47FAA] |
| Q8A6Z2 | HMPREF9533_02378 | 34.783 | 99 | 0 | 656 | phosphoribosylformylglycinamidine synthase [Escherichia coli MS 60-1] |
| Q8A6Z2 | HMPREF9552_02508 | 34.938 | 99 | 0 | 655 | phosphoribosylformylglycinamidine synthase [Escherichia coli MS 198-1] |
| Q8A6Z2 | HMPREF9532_01834 | 35.09 | 98 | 0 | 655 | phosphoribosylformylglycinamidine synthase [Escherichia coli MS 57-2] |
| Q8A6Z2 | HMPREF9535_04675 | 35.012 | 98 | 0 | 655 | phosphoribosylformylglycinamidine synthase [Escherichia coli MS 78-1] |
| Q8A6Z2 | CSAG_02352 | 34.646 | 98 | 0 | 655 | phosphoribosylformylglycinamidine synthase [Citrobacter sp. 30_2] |
| Q8A6Z2 | HMPREF9545_00413 | 34.961 | 98 | 0 | 655 | phosphoribosylformylglycinamidine synthase [Escherichia coli MS 16-3] |
| Q8A6Z2 | HMPREF9534_04086 | 34.86 | 99 | 0 | 655 | phosphoribosylformylglycinamidine synthase [Escherichia coli MS 69-1] |
| Q8A6Z2 | ENTCAN_07231 | 34.803 | 98 | 0 | 654 | phosphoribosylformylglycinamidine synthase [Enterobacter cancerogenus ATCC 35316] |
| Q8A6Z2 | ECSF_2396 | 34.783 | 99 | 0 | 654 | phosphoribosylformyl-glycineamide synthetase [Escherichia coli SE15] |
| Q8A6Z2 | SSJG_00034 | 35.09 | 98 | 0 | 654 | phosphoribosylformylglycinamidine synthase [Escherichia coli D9] |
| Q8A6Z2 | HMPREF9539_02029 | 35.012 | 98 | 0 | 654 | phosphoribosylformylglycinamidine synthase [Escherichia coli MS 110-3] |
| Q8A6Z2 | ESAG_03402 | 35.012 | 98 | 0 | 654 | phosphoribosylformylglycinamidine synthase [Escherichia sp. 3_2_53FAA] |
| Q8A6Z2 | CIT292_05939 | 34.409 | 98 | 0 | 653 | phosphoribosylformylglycinamidine synthase [Citrobacter youngae ATCC 29220] |
| Q8A6Z2 | HMPREF9348_01580 | 35.008 | 98 | 0 | 652 | phosphoribosylformylglycinamidine synthase [Escherichia coli MS 145-7] |
| Q8A6Z2 | HMPREF0201_03523 | 34.964 | 98 | 0 | 652 | phosphoribosylformylglycinamidine synthase [Cedecea davisae DSM 4568] |
| Q8A6Z2 | HMPREF9440_00567 | 34.292 | 98 | 0 | 652 | phosphoribosylformylglycinamidine synthase [Sutterella parvirubra YIT 11816] |
| Q8A6Z2 | HMPREF1004_03311 | 34.228 | 98 | 0 | 650 | phosphoribosylformylglycinamidine synthase [Ralstonia sp. 5_7_47FAA] |
| Q8A6Z2 | HMPREF0989_04307 | 34.228 | 98 | 0 | 650 | phosphoribosylformylglycinamidine synthase [Ralstonia sp. 5_2_56FAA] |
| Q8A6Z2 | HMPREF9428_02491 | 34.488 | 98 | 0 | 650 | phosphoribosylformylglycinamidine synthase [Citrobacter freundii 4_7_47CFAA] |
| Q8A6Z2 | HMPREF9464_00699 | 33.591 | 99 | 0 | 648 | phosphoribosylformylglycinamidine synthase [Sutterella wadsworthensis 3_1_45B] |
| Q8A6Z2 | HMPREF1476_01408 | 33.514 | 99 | 0 | 646 | phosphoribosylformylglycinamidine synthase [Sutterella wadsworthensis HGA0223] |
| Q8A6Z2 | HMPREF9548_05055 | 36.084 | 89 | 0 | 626 | phosphoribosylformylglycinamidine synthase [Escherichia coli MS 182-1] |
| Q8A6Z2 | HMPREF0189_01390 | 34.123 | 97 | 0 | 611 | phosphoribosylformylglycinamidine synthase [Burkholderiales bacterium 1_1_47] |
| Q8A6Z2 | HMPREF9439_02567 | 34.123 | 97 | 0 | 611 | phosphoribosylformylglycinamidine synthase [Parasutterella excrementihominis YIT 11859] |
| Q8A6Z2 | PROPEN_01937 | 35.559 | 50 | 4.14E-104 | 347 | putative phosphoribosylformylglycinamidine synthase [Proteus penneri ATCC 35198] |
| Q8A6Z2 | BACCOPRO_03261 | 77.387 | 16 | 5.63E-103 | 328 | phosphoribosylformylglycinamidine synthase [Bacteroides coprophilus DSM 18228 = JCM 13818] |
| Q8A6Z2 | HMPREF0178_03775 | 27.273 | 65 | 1.25E-47 | 190 | hypothetical protein [Bilophila sp. 4_1_30] |
| Q8A6Z2 | HMPREF0179_03053 | 27.156 | 65 | 1.60E-47 | 190 | phosphoribosylformylglycinamidine synthase II [Bilophila wadsworthia 3_1_6] |
| Q8A6Z2 | CLOSPO_02429 | 25.214 | 87 | 4.89E-47 | 189 | phosphoribosylformylglycinamidine synthase [Clostridium sporogenes ATCC 15579] |
| Q8A6Z2 | HMPREF0219_3458 | 24.108 | 87 | 2.05E-45 | 184 | phosphoribosylformylglycinamidine synthase [Peptoclostridium difficile NAP07] |
| Q8A6Z2 | HMPREF0220_3540 | 24.108 | 87 | 2.05E-45 | 184 | phosphoribosylformylglycinamidine synthase [Peptoclostridium difficile NAP08] |
| Q8A6Z2 | HMPREF9945_01249 | 23.888 | 87 | 2.30E-44 | 181 | phosphoribosylformylglycinamidine synthase [Peptoclostridium difficile 70-100-2010] |
| Q8A6Z2 | HMPREF9476_00105 | 24.048 | 87 | 3.68E-44 | 180 | phosphoribosylformylglycinamidine synthase [Clostridium perfringens WAL-14572] |
| Q8A6Z2 | HMPREF1481_01511 | 24.162 | 89 | 4.15E-44 | 179 | phosphoribosylformylglycinamidine synthase [Streptococcus sp. HPH0090] |
| Q8A6Z2 | DESPIG_02920 | 27.66 | 62 | 4.92E-44 | 179 | putative phosphoribosylformylglycinamidine synthase II [Desulfovibrio piger ATCC 29098] |
| Q8A6Z2 | COLAER_00518 | 23.27 | 89 | 2.24E-43 | 177 | phosphoribosylformylglycinamidine synthase [Collinsella aerofaciens ATCC 25986] |
| Q8A6Z2 | HMPREF0819_1761 | 23.641 | 88 | 5.86E-43 | 176 | phosphoribosylformylglycinamidine synthase [Streptococcus equinus ATCC 9812] |
| Q8A6Z2 | CC1_33390 | 24.754 | 77 | 5.85E-40 | 166 | phosphoribosylformylglycinamidine synthase [Coprococcus catus GD/7] |
| Q8A6Z2 | HMPREF0542_10428 | 26.816 | 56 | 4.35E-38 | 158 | phosphoribosylformylglycinamidine synthase II [Lactobacillus ruminis ATCC 25644] |
| Q8A6Z2 | HMPREF0216_02029 | 22.82 | 92 | 4.68E-37 | 157 | phosphoribosylformylglycinamidine synthase [Clostridium celatum DSM 1785] |
| Q8A6Z2 | HMPREF1485_00657 | 23.345 | 65 | 3.29E-36 | 152 | phosphoribosylformylglycinamidine synthase 2 [Propionibacterium sp. HGH0353] |
| Q8A6Z2 | HMPREF9412_5785 | 25.983 | 56 | 8.08E-36 | 151 | phosphoribosylformylglycinamidine synthase II [Paenibacillus sp. HGF5] |
| Q8A6Z2 | HMPREF0492_1934 | 24.644 | 56 | 2.03E-34 | 147 | phosphoribosylformylglycinamidine synthase II [Lactobacillus acidophilus ATCC 4796] |
| Q8A6Z2 | HMPREF0519_1395 | 24.148 | 56 | 1.46E-32 | 141 | phosphoribosylformylglycinamidine synthase II [Lactobacillus hilgardii ATCC 8290] |
| Q8A6Z2 | HMPREF0497_0606 | 24.148 | 56 | 1.46E-32 | 141 | phosphoribosylformylglycinamidine synthase II [Lactobacillus buchneri ATCC 11577] |
| Q8A6Z2 | HMPREF0496_0646 | 24.006 | 56 | 1.22E-31 | 138 | phosphoribosylformylglycinamidine synthase II [Lactobacillus brevis subsp. gravesensis ATCC 27305] |
| Q8A6Z2 | HMPREF1207_01882 | 25.384 | 56 | 2.37E-31 | 137 | phosphoribosylformylglycinamidine synthase II [Paenibacillus sp. HGH0039] |
| Q8A6Z2 | HMPREF9413_2173 | 25.384 | 56 | 2.37E-31 | 137 | phosphoribosylformylglycinamidine synthase II [Paenibacillus sp. HGF7] |
| Q8A4P8 | BSIG_4968 | 99.669 | 100 | 0 | 622 | hypothetical protein [Bacteroides sp. 1_1_6] |
| Q8A4P8 | HMPREF9007_04599 | 98.675 | 100 | 0 | 618 | muramoyltetrapeptide carboxypeptidase [Bacteroides sp. 1_1_14] |
| Q8A4P8 | BXY_16870 | 90.397 | 100 | 0 | 577 | Uncharacterized proteins, homologs of microcin C7 resistance protein MccF [Bacteroides xylanisolvens XB1A] |
| Q8A4P8 | HMPREF0102_04106 | 89.735 | 100 | 0 | 573 | LD-carboxypeptidase [Bacteroides sp. 2_1_22] |
| Q8A4P8 | BSAG_03604 | 89.735 | 100 | 0 | 573 | LD-carboxypeptidase [Bacteroides sp. D1] |

| | | | | | |
|---|---|---|---|---|---|
| Q8A4P8 | HMPREF0106_02702 | 89.735 | 100 | 0 | 573 muramoyltetrapeptide carboxypeptidase [Bacteroides sp. D22] |
| Q8A4P8 | CW3_4238 | 89.735 | 100 | 0 | 573 LD-carboxypeptidase [Bacteroides xylanisolvens SD CC 1b] |
| Q8A4P8 | CW1_3273 | 89.735 | 100 | 0 | 573 LD-carboxypeptidase [Bacteroides xylanisolvens SD CC 2a] |
| Q8A4P8 | BACCAC_00303 | 89.404 | 100 | 0 | 573 LD-carboxypeptidase [Bacteroides caccae ATCC 43185] |
| Q8A4P8 | HMPREF0127_04309 | 89.735 | 100 | 0 | 573 hypothetical protein [Bacteroides sp. 1_1_30] |
| Q8A4P8 | BSGG_4838 | 89.404 | 100 | 0 | 572 hypothetical protein [Bacteroides sp. D2] |
| Q8A4P8 | HMPREF9010_04622 | 89.073 | 100 | 0 | 568 putative muramoyltetrapeptide carboxypeptidase [Bacteroides sp. 3_1_23] |
| Q8A4P8 | BACOVA_01632 | 89.073 | 100 | 0 | 568 LD-carboxypeptidase [Bacteroides ovatus ATCC 8483] |
| Q8A4P8 | CUY_2716 | 88.742 | 100 | 0 | 567 LD-carboxypeptidase [Bacteroides ovatus SD CMC 3f] |
| Q8A4P8 | BSCG_00120 | 88.742 | 100 | 0 | 567 LD-carboxypeptidase [Bacteroides sp. 2_2_4] |
| Q8A4P8 | HMPREF1017_04372 | 88.742 | 100 | 0 | 567 hypothetical protein [Bacteroides ovatus 3_8_47FAA] |
| Q8A4P8 | BACFIN_08264 | 89.404 | 100 | 0 | 561 LD-carboxypeptidase [Bacteroides finegoldii DSM 17565] |
| Q8A4P8 | HMPREF1532_02182 | 81.544 | 99 | 0 | 509 hypothetical protein [Bacteroides salyersiae WAL 10018 = DSM 18765 = JCM 12988] |
| Q8A4P8 | HMPREF1214_00145 | 79.53 | 99 | 0 | 504 muramoyltetrapeptide carboxypeptidase [Bacteroides sp. HPS0048] |
| Q8A4P8 | BSGG_3797 | 76.923 | 99 | 9.52E-175 | 490 hypothetical protein [Bacteroides sp. D2] |
| Q8A4P8 | HMPREF9446_00340 | 73.841 | 98 | 1.01E-166 | 469 LD-carboxypeptidase [Bacteroides fluxus YIT 12057] |
| Q8A4P8 | BACSTE_02094 | 72.185 | 98 | 7.34E-165 | 464 LD-carboxypeptidase [Bacteroides stercoris ATCC 43183] |
| Q8A4P8 | BFAG_04203 | 73.49 | 99 | 3.74E-164 | 462 LD-carboxypeptidase [Bacteroides fragilis 3_1_12] |
| Q8A4P8 | HMPREF9445_02085 | 71.523 | 98 | 4.81E-163 | 459 LD-carboxypeptidase [Bacteroides clarus YIT 12056] |
| Q8A4P8 | BACCELL_02563 | 70.323 | 98 | 6.48E-163 | 459 LD-carboxypeptidase [Bacteroides cellulosilyticus DSM 14838] |
| Q8A4P8 | HMPREF1016_02641 | 71.854 | 98 | 1.80E-162 | 459 LD-carboxypeptidase [Bacteroides eggerthii 1_2_48FAA] |
| Q8A4P8 | BACEGG_03014 | 71.854 | 98 | 2.63E-162 | 461 LD-carboxypeptidase [Bacteroides eggerthii DSM 20697] |
| Q8A4P8 | BACUNI_03919 | 71.523 | 98 | 4.26E-162 | 457 LD-carboxypeptidase [Bacteroides uniformis ATCC 8492] |
| Q8A4P8 | BSHG_3542 | 73.49 | 99 | 6.88E-162 | 456 hypothetical protein [Bacteroides sp. 3_2_5] |
| Q8A4P8 | HMPREF1007_00699 | 71.192 | 98 | 7.36E-162 | 456 LD-carboxypeptidase [Bacteroides sp. 4_1_36] |
| Q8A4P8 | HMPREF0969_01945 | 71.192 | 98 | 1.45E-161 | 456 LD-carboxypeptidase [Bacteroides sp. D20] |
| Q8A4P8 | HMPREF0101_04104 | 73.49 | 99 | 2.20E-161 | 455 LD-carboxypeptidase [Bacteroides sp. 2_1_16] |
| Q8A4P8 | HMPREF9447_05145 | 70.323 | 98 | 5.35E-161 | 454 hypothetical protein [Bacteroides oleiciplenus YIT 12058] |
| Q8A4P8 | HMPREF1018_04019 | 73.154 | 99 | 8.00E-161 | 453 hypothetical protein [Bacteroides sp. 2_1_56FAA] |
| Q8A4P8 | BACINT_01681 | 69.481 | 97 | 1.58E-159 | 451 hypothetical protein [Bacteroides intestinalis DSM 17393] |
| Q8A4P8 | HMPREF9448_00941 | 53.898 | 98 | 1.42E-109 | 323 hypothetical protein [Barnesiella intestinihominis YIT 11860] |
| Q8A4P8 | HMPREF9450_01386 | 38.127 | 98 | 3.72E-65 | 211 hypothetical protein [Alistipes indistinctus YIT 12060] |
| Q8A4P8 | FUAG_00779 | 34.459 | 96 | 1.20E-49 | 171 hypothetical protein [Fusobacterium ulcerans ATCC 49185] |
| Q8A4P8 | HMPREF0402_00398 | 34.122 | 96 | 1.33E-48 | 168 hypothetical protein [Fusobacterium ulcerans 12-1B] |
| Q8A4P8 | CLOHYLEM_04225 | 34.694 | 94 | 7.26E-45 | 158 LD-carboxypeptidase [Lachnospiraceae [Clostridium] hylemonae DSM 15053] |
| Q8A4P8 | MHY_24330 | 32.776 | 95 | 2.82E-43 | 154 Uncharacterized proteins, homologs of microcin C7 resistance protein MccF [Megamonas hypermegale ART12/1] |
| Q8A4P8 | CLOSPO_00911 | 32.89 | 96 | 3.93E-43 | 153 LD-carboxypeptidase [Clostridium sporogenes ATCC 15579] |
| Q8A4P8 | HMPREF9454_00143 | 32.441 | 95 | 9.56E-43 | 153 hypothetical protein [Megamonas funiformis YIT 11815] |
| Q8A4P8 | HMPREF0993_01236 | 33.442 | 96 | 1.12E-42 | 152 hypothetical protein [Lachnospiraceae bacterium 5_1_57FAA] |
| Q8A4P8 | CLOSCI_00595 | 33.442 | 96 | 3.48E-42 | 152 LD-carboxypeptidase [Lachnospiraceae [Clostridium] scindens ATCC 35704] |
| Q8A4P8 | HMPREF1014_03755 | 32.333 | 97 | 3.83E-42 | 151 hypothetical protein [Bacillus sp. 7_6_55CFAA_CT2] |
| Q8A4P8 | HMPREF9453_01086 | 34.256 | 93 | 5.48E-41 | 149 hypothetical protein [Dialister succinatiphilus YIT 11850] |
| Q8A4P8 | CSBG_00001 | 32.886 | 93 | 2.28E-40 | 146 hypothetical protein [Clostridium sp. 7_2_43FAA] |
| Q8A4P8 | HMPREF1032_01291 | 29.739 | 97 | 2.45E-40 | 146 hypothetical protein [Subdoligranulum sp. 4_3_54A2FAA] |
| Q8A4P8 | FVAG_01986 | 33.446 | 96 | 2.91E-40 | 146 LD-carboxypeptidase [Fusobacterium varium ATCC 27725] |
| Q8A4P8 | HMPREF1705_00197 | 30.662 | 93 | 3.92E-40 | 145 LD-carboxypeptidase [Anaerobaculum hydrogeniformans ATCC BAA-1850] |
| Q8A4P8 | HMPREF9454_00157 | 31.683 | 97 | 6.06E-40 | 146 hypothetical protein [Megamonas funiformis YIT 11815] |
| Q8A4P8 | HMPREF1210_02249 | 36.082 | 89 | 3.03E-39 | 143 hypothetical protein [Paenisporosarcina sp. HGH0030] |
| Q8A4P8 | HMPREF9476_02507 | 31.525 | 97 | 3.24E-39 | 143 hypothetical protein [Clostridium perfringens WAL-14572] |
| Q8A4P8 | HMPREF9449_03029 | 30.651 | 85 | 4.38E-39 | 141 hypothetical protein [Odoribacter laneus YIT 12061] |
| Q8A4P8 | FMAG_00848 | 31.359 | 93 | 7.35E-39 | 142 LD-carboxypeptidase [Fusobacterium mortiferum ATCC 9817] |
| Q8A4P8 | HMPREF0219_2738 | 33.673 | 94 | 5.16E-38 | 140 LD-carboxypeptidase [Peptoclostridium difficile NAP07] |
| Q8A4P8 | HMPREF0220_2384 | 33.673 | 94 | 5.16E-38 | 140 LD-carboxypeptidase [Peptoclostridium difficile NAP08] |
| Q8A4P8 | CLOBAR_02417 | 30.99 | 98 | 1.69E-36 | 136 LD-carboxypeptidase [Intestinibacter bartlettii DSM 16795] |
| Q8A4P8 | CLOSTASPAR_05380 | 31.09 | 97 | 2.86E-36 | 135 LD-carboxypeptidase [Lachnospiraceae [Clostridium asparagiforme] DSM 15981] |
| Q8A4P8 | HMPREF9456_00356 | 27.673 | 97 | 2.19E-35 | 134 hypothetical protein [Dysgonomonas mossii DSM 22836] |
| Q8A4P8 | HMPREF1211_01754 | 31.683 | 98 | 5.31E-35 | 132 hypothetical protein [Streptomyces sp. HGB0020] |
| Q8A4P8 | HMPREF9467_01949 | 31.56 | 89 | 9.30E-34 | 129 hypothetical protein [Lachnospiraceae [Clostridium] clostridioforme 2_1_49FAA] |
| Q8A4P8 | CLOBOL_06414 | 31.69 | 89 | 1.82E-33 | 128 hypothetical protein [Lachnospiraceae [Clostridium] bolteae ATCC BAA-613] |
| Q8A4P8 | HMPREF9412_4142 | 30.952 | 94 | 2.17E-33 | 128 LD-carboxypeptidase [Paenibacillus sp. HGF5] |

| | | | | | | |
|---|---|---|---|---|---|---|
| Q8A4P8 | OFAG_02310 | 31.034 | 94 | 1.65E-31 | 124 | hypothetical protein [Oxalobacter formigenes HOxBLS] |
| Q8A4P8 | HMPREF1207_04435 | 29.01 | 94 | 1.85E-31 | 123 | hypothetical protein [Paenibacillus sp. HGH0039] |
| Q8A4P8 | HMPREF9413_3403 | 29.01 | 94 | 1.85E-31 | 123 | LD-carboxypeptidase [Paenibacillus sp. HGF7] |
| Q8A4P8 | CUW_2297 | 31.229 | 99 | 3.71E-31 | 122 | LD-carboxypeptidase [Turicibacter sanguinis PC909] |
| Q8A4P8 | HMPREF9402_2783 | 31.229 | 99 | 3.71E-31 | 122 | LD-carboxypeptidase [Turicibacter sp. HGF1] |
| Q8A3J3 | HMPREF9007_02678 | 100 | 100 | 0 | 598 | conserved hypothetical protein [Bacteroides sp. 1_1_14] |
| Q8A3J3 | BSIG_1684 | 100 | 100 | 0 | 598 | hypothetical protein [Bacteroides sp. 1_1_6] |
| Q8A3J3 | HMPREF0127_02155 | 76.491 | 100 | 1.56E-173 | 484 | hypothetical protein [Bacteroides sp. 1_1_30] |
| Q8A3J3 | HMPREF9010_00272 | 76.491 | 100 | 2.00E-173 | 484 | conserved hypothetical protein [Bacteroides sp. 3_1_23] |
| Q8A3J3 | BSGG_2718 | 76.491 | 100 | 2.00E-173 | 484 | hypothetical protein [Bacteroides sp. D2] |
| Q8A3J3 | CUY_0456 | 76.491 | 100 | 2.16E-173 | 484 | conserved hypothetical protein [Bacteroides ovatus SD CMC 3f] |
| Q8A3J3 | BXY_20840 | 76.491 | 100 | 2.72E-173 | 483 | GDSL-like Lipase/Acylhydrolase. [Bacteroides xylanisolvens XB1A] |
| Q8A3J3 | HMPREF0102_00897 | 76.491 | 100 | 2.97E-173 | 483 | hypothetical protein [Bacteroides sp. 2_1_22] |
| Q8A3J3 | BACOVA_02193 | 76.491 | 100 | 2.97E-173 | 483 | hypothetical protein [Bacteroides ovatus ATCC 8483] |
| Q8A3J3 | BSAG_00647 | 76.491 | 100 | 2.97E-173 | 483 | hypothetical protein [Bacteroides sp. D1] |
| Q8A3J3 | HMPREF0106_00356 | 76.491 | 100 | 2.97E-173 | 483 | conserved hypothetical protein [Bacteroides sp. D22] |
| Q8A3J3 | CW3_0098 | 76.491 | 100 | 2.97E-173 | 483 | conserved hypothetical protein [Bacteroides xylanisolvens SD CC 1b] |
| Q8A3J3 | CW1_0097 | 76.491 | 100 | 2.97E-173 | 483 | conserved hypothetical protein [Bacteroides xylanisolvens SD CC 2a] |
| Q8A3J3 | BACCAC_01236 | 76.761 | 99 | 4.39E-173 | 483 | GDSL-like protein [Bacteroides caccae ATCC 43185] |
| Q8A3J3 | BSCG_01431 | 76.14 | 100 | 8.60E-173 | 482 | hypothetical protein [Bacteroides sp. 2_2_4] |
| Q8A3J3 | HMPREF1017_00194 | 75.789 | 100 | 1.55E-171 | 479 | hypothetical protein [Bacteroides ovatus 3_8_47FAA] |
| Q8A3J3 | BSFG_02291 | 75.789 | 100 | 2.09E-165 | 463 | GDSL-like protein [Bacteroides sp. 4_3_47FAA] |
| Q8A3J3 | HMPREF9011_03781 | 75.789 | 100 | 2.09E-165 | 463 | hypothetical protein [Bacteroides sp. 3_1_40A] |
| Q8A3J3 | HMPREF0105_1087 | 76.491 | 100 | 1.15E-164 | 462 | GDSL-like protein [Bacteroides sp. 3_1_33FAA] |
| Q8A3J3 | BSEG_03837 | 76.491 | 100 | 1.29E-164 | 461 | GDSL-like protein [Bacteroides dorei 5_1_36/D4] |
| Q8A3J3 | BACDOR_03379 | 76.491 | 100 | 1.29E-164 | 461 | GDSL-like protein [Bacteroides dorei DSM 17855] |
| Q8A3J3 | BSBG_04104 | 76.491 | 100 | 1.29E-164 | 461 | GDSL-like protein [Bacteroides sp. 9_1_42FAA] |
| Q8A3J3 | CUU_2039 | 76.491 | 100 | 1.02E-163 | 459 | putative lipoprotein [Bacteroides vulgatus PC510] |
| Q8A3J3 | PRABACTJOHN_03923 | 51.21 | 87 | 8.04E-90 | 270 | GDSL-like protein [Parabacteroides johnsonii DSM 18315] |
| Q8A3J3 | BACCOP_00756 | 50.82 | 85 | 1.96E-89 | 270 | GDSL-like protein [Bacteroides coprocola DSM 17136] |
| Q8A3J3 | HMPREF9446_02249 | 48.438 | 90 | 1.06E-88 | 270 | GDSL-like protein [Bacteroides fluxus YIT 12057] |
| Q8A3J3 | BACPLE_00396 | 50.204 | 86 | 2.10E-81 | 250 | GDSL-like protein [Bacteroides plebeius DSM 17135] |
| Q8A371 | BSIG_1565 | 100 | 100 | 0 | 566 | hypothetical protein [Bacteroides sp. 1_1_6] |
| Q8A371 | HMPREF1017_00341 | 85.185 | 99 | 4.34E-178 | 495 | hypothetical protein [Bacteroides ovatus 3_8_47FAA] |
| Q8A371 | BACFIN_08783 | 85.556 | 99 | 7.57E-178 | 494 | hypothetical protein [Bacteroides finegoldii DSM 17565] |
| Q8A371 | HMPREF0106_03039 | 84.815 | 99 | 7.51E-177 | 492 | lipase [Bacteroides sp. D22] |
| Q8A371 | HMPREF0127_01336 | 84.528 | 97 | 7.00E-174 | 484 | hypothetical protein [Bacteroides sp. 1_1_30] |
| Q8A371 | BSCG_01536 | 83.774 | 97 | 8.07E-172 | 478 | hypothetical protein [Bacteroides sp. 2_2_4] |
| Q8A371 | BACOVA_02773 | 86.111 | 92 | 3.87E-169 | 472 | hypothetical protein [Bacteroides ovatus ATCC 8483] |
| Q8A371 | HMPREF0102_00791 | 85.714 | 92 | 2.47E-168 | 469 | hypothetical protein [Bacteroides sp. 2_1_22] |
| Q8A371 | BSAG_00543 | 85.185 | 99 | 2.73E-168 | 470 | hypothetical protein [Bacteroides sp. D1] |
| Q8A371 | CW3_4721 | 85.185 | 99 | 2.73E-168 | 470 | conserved hypothetical protein [Bacteroides xylanisolvens SD CC 1b] |
| Q8A371 | CW1_0448 | 85.185 | 99 | 2.73E-168 | 470 | conserved hypothetical protein [Bacteroides xylanisolvens SD CC 2a] |
| Q8A371 | CUY_1628 | 85.714 | 92 | 3.55E-168 | 469 | conserved hypothetical protein [Bacteroides ovatus SD CMC 3f] |
| Q8A371 | BSGG_2580 | 85.768 | 98 | 1.95E-167 | 468 | hypothetical protein [Bacteroides sp. D2] |
| Q8A371 | BXY_21900 | 84.815 | 99 | 2.21E-167 | 468 | Esterase/lipase [Bacteroides xylanisolvens XB1A] |
| Q8A371 | HMPREF9007_02511 | 99.548 | 81 | 4.51E-164 | 457 | lipase [Bacteroides sp. 1_1_14] |
| Q8A371 | HMPREF9447_02121 | 75.188 | 97 | 2.48E-154 | 435 | hypothetical protein [Bacteroides oleiciplenus YIT 12058] |
| Q8A371 | BACINT_01421 | 75.94 | 97 | 8.98E-150 | 423 | hypothetical protein [Bacteroides intestinalis DSM 17393] |
| Q8A371 | BACCELL_01548 | 80.162 | 90 | 6.47E-145 | 410 | hydrolase, alpha/beta domain protein [Bacteroides cellulosilyticus DSM 14838] |
| Q8A371 | BFAG_04647 | 72.414 | 96 | 1.43E-136 | 389 | hypothetical protein [Bacteroides fragilis 3_1_12] |
| Q8A371 | BSHG_1590 | 72.201 | 95 | 3.12E-131 | 376 | hypothetical protein [Bacteroides sp. 3_2_5] |
| Q8A371 | HMPREF1018_00045 | 72.201 | 95 | 9.96E-131 | 375 | hypothetical protein [Bacteroides sp. 2_1_56FAA] |
| Q8A371 | HMPREF0101_01998 | 71.815 | 95 | 1.50E-129 | 372 | hypothetical protein [Bacteroides sp. 2_1_16] |
| Q8A371 | HMPREF9455_02060 | 59.091 | 97 | 2.69E-125 | 361 | hypothetical protein [Dysgonomonas gadei ATCC BAA-286] |
| Q8A371 | HMPREF9456_03136 | 62.222 | 97 | 4.60E-119 | 345 | hypothetical protein [Dysgonomonas mossii DSM 22836] |
| Q8A371 | HMPREF0969_02650 | 57.955 | 96 | 4.89E-118 | 343 | hydrolase, alpha/beta domain protein [Bacteroides sp. D20] |
| Q8A371 | HMPREF1007_02910 | 57.955 | 96 | 1.43E-117 | 342 | alpha/beta hydrolase fold protein [Bacteroides sp. 4_1_36] |
| Q8A371 | BACUNI_04717 | 57.955 | 96 | 7.45E-117 | 340 | hypothetical protein [Bacteroides uniformis ATCC 8492] |
| Q8A371 | HMPREF9456_00758 | 56.929 | 97 | 9.58E-107 | 314 | hypothetical protein [Dysgonomonas mossii DSM 22836] |

| | | | | | | |
|---|---|---|---|---|---|---|
| Q8A371 | HMPREF1475_00688 | 56.225 | 91 | 2.59E-101 | 300 | hypothetical protein [Prevotella oralis HGA0225] |
| Q8A371 | HMPREF0104_01033 | 51.136 | 96 | 6.63E-96 | 286 | lipase [Bacteroides sp. 3_1_19] |
| Q8A371 | HMPREF0619_01109 | 50.187 | 97 | 1.69E-95 | 286 | hydrolase, alpha/beta domain protein [Parabacteroides sp. D13] |
| Q8A371 | HMPREF0103_3491 | 50.951 | 96 | 3.16E-95 | 285 | hydrolase, alpha/beta domain protein [Bacteroides sp. 2_1_33B] |
| Q8A371 | HMPREF9008_03047 | 50.951 | 96 | 3.16E-95 | 285 | hydrolase, alpha/beta domain protein [Parabacteroides sp. 20_3] |
| Q8A371 | HMPREF9010_00520 | 50.376 | 97 | 5.10E-95 | 284 | probable lipase [Bacteroides sp. 3_1_23] |
| Q8A371 | BSGG_2495 | 49.805 | 94 | 8.95E-91 | 273 | hypothetical protein [Bacteroides sp. D2] |
| Q8A371 | BACCOP_00577 | 44.891 | 97 | 3.41E-82 | 252 | hypothetical protein [Bacteroides coprocola DSM 17136] |
| Q8A371 | PREVCOP_05110 | 44.074 | 97 | 4.51E-81 | 249 | esterase/lipase/thioesterase family protein [Prevotella copri DSM 18205] |
| Q8A371 | PARMER_00333 | 45.349 | 94 | 1.19E-69 | 219 | hypothetical protein [Parabacteroides merdae ATCC 43184] |
| Q8A371 | AL1_16970 | 42.366 | 96 | 2.61E-69 | 219 | Esterase/lipase [Alistipes shahii WAL 8301] |
| Q8A371 | PRABACTJOHN_00567 | 43.182 | 97 | 6.38E-67 | 213 | hypothetical protein [Parabacteroides johnsonii DSM 18315] |
| Q8A2Q1 | HMPREF9007_03794 | 99.18 | 100 | 0 | 1510 | dipeptidyl aminopeptidase IV [Bacteroides sp. 1_1_14] |
| Q8A2Q1 | BSIG_1417 | 99.18 | 100 | 0 | 1510 | hypothetical protein [Bacteroides sp. 1_1_6] |
| Q8A2Q1 | BACCAC_01453 | 81.446 | 100 | 0 | 1265 | peptidase, S9A/B/C family, catalytic domain protein [Bacteroides caccae ATCC 43185] |
| Q8A2Q1 | HMPREF0102_00604 | 80.245 | 100 | 0 | 1249 | peptidase, S9A/B/C family, catalytic domain protein [Bacteroides sp. 2_1_22] |
| Q8A2Q1 | BSAG_00356 | 80.245 | 100 | 0 | 1249 | peptidase, S9A/B/C family, catalytic domain protein [Bacteroides sp. D1] |
| Q8A2Q1 | CW3_2034 | 80.245 | 100 | 0 | 1249 | dipeptidyl peptidase IV N-terminal domain protein [Bacteroides xylanisolvens SD CC 1b] |
| Q8A2Q1 | CW1_0738 | 80.245 | 100 | 0 | 1249 | dipeptidyl peptidase IV N-terminal domain protein [Bacteroides xylanisolvens SD CC 2a] |
| Q8A2Q1 | HMPREF0127_01576 | 80.381 | 100 | 0 | 1249 | hypothetical protein [Bacteroides sp. 1_1_30] |
| Q8A2Q1 | BXY_23820 | 80.109 | 100 | 0 | 1249 | prolyl tripeptidyl peptidase. Serine peptidase. MEROPS family S09B [Bacteroides xylanisolvens XB1A] |
| Q8A2Q1 | HMPREF0106_04714 | 80.381 | 100 | 0 | 1249 | dipeptidyl aminopeptidase IV [Bacteroides sp. D22] |
| Q8A2Q1 | HMPREF1017_00530 | 80.109 | 100 | 0 | 1237 | hypothetical protein [Bacteroides ovatus 3_8_47FAA] |
| Q8A2Q1 | BSGG_2370 | 79.837 | 100 | 0 | 1233 | hypothetical protein [Bacteroides sp. D2] |
| Q8A2Q1 | CUY_3404 | 79.7 | 100 | 0 | 1230 | dipeptidyl peptidase IV N-terminal domain protein [Bacteroides ovatus SD CMC 3f] |
| Q8A2Q1 | BACOVA_01349 | 79.564 | 100 | 0 | 1228 | peptidase, S9A/B/C family, catalytic domain protein [Bacteroides ovatus ATCC 8483] |
| Q8A2Q1 | HMPREF9010_00648 | 79.292 | 100 | 0 | 1227 | putative dipeptidyl aminopeptidase IV [Bacteroides sp. 3_1_23] |
| Q8A2Q1 | HMPREF1214_00573 | 76.157 | 97 | 0 | 1173 | dipeptidyl-peptidase 4 [Bacteroides sp. HPS0048] |
| Q8A2Q1 | HMPREF1532_01808 | 74.863 | 100 | 0 | 1165 | hypothetical protein [Bacteroides salyersiae WAL 10018 = DSM 18765 = JCM 12988] |
| Q8A2Q1 | BACFIN_08470 | 75.886 | 100 | 0 | 1154 | peptidase, S9A/B/C family, catalytic domain protein [Bacteroides finegoldii DSM 17565] |
| Q8A2Q1 | BACCELL_02178 | 72.418 | 100 | 0 | 1113 | peptidase, S9A/B/C family, catalytic domain protein [Bacteroides cellulosilyticus DSM 14838] |
| Q8A2Q1 | HMPREF9447_02557 | 70.516 | 100 | 0 | 1087 | hypothetical protein [Bacteroides oleiciplenus YIT 12058] |
| Q8A2Q1 | BFAG_02901 | 71.049 | 97 | 0 | 1071 | peptidase, S9A/B/C family, catalytic domain protein [Bacteroides fragilis 3_1_12] |
| Q8A2Q1 | HMPREF0101_02096 | 71.189 | 97 | 0 | 1068 | peptidase, S9A/B/C family, catalytic domain protein [Bacteroides sp. 2_1_16] |
| Q8A2Q1 | HMPREF1018_00139 | 70.909 | 97 | 0 | 1065 | hypothetical protein [Bacteroides sp. 2_1_56FAA] |
| Q8A2Q1 | HMPREF9445_01935 | 67.867 | 100 | 0 | 1064 | peptidase, S9A/B/C family, catalytic domain protein [Bacteroides clarus YIT 12056] |
| Q8A2Q1 | BSHG_1495 | 70.909 | 97 | 0 | 1063 | hypothetical protein [Bacteroides sp. 3_2_5] |
| Q8A2Q1 | HMPREF9446_02242 | 69.536 | 98 | 0 | 1057 | peptidase, S9A/B/C family, catalytic domain protein [Bacteroides fluxus YIT 12057] |
| Q8A2Q1 | BACSTE_02313 | 68 | 100 | 0 | 1057 | peptidase, S9A/B/C family, catalytic domain protein [Bacteroides stercoris ATCC 43183] |
| Q8A2Q1 | HMPREF1016_00991 | 68.45 | 97 | 0 | 1036 | dipeptidyl peptidase IV [Bacteroides eggerthii 1_2_48FAA] |
| Q8A2Q1 | BACEGG_03258 | 68.176 | 97 | 0 | 1035 | peptidase, S9A/B/C family, catalytic domain protein [Bacteroides eggerthii DSM 20697] |
| Q8A2Q1 | BACINT_01011 | 64.829 | 100 | 0 | 976 | peptidase, S9A/B/C family, catalytic domain protein [Bacteroides intestinalis DSM 17393] |
| Q8A2Q1 | HMPREF0969_01715 | 62.435 | 99 | 0 | 941 | peptidase, S9A/B/C family, catalytic domain protein [Bacteroides sp. D20] |
| Q8A2Q1 | BACUNI_04166 | 62.304 | 99 | 0 | 938 | peptidase, S9A/B/C family, catalytic domain protein [Bacteroides uniformis ATCC 8492] |
| Q8A2Q1 | HMPREF1007_00511 | 62.304 | 99 | 0 | 938 | dipeptidyl peptidase IV [Bacteroides sp. 4_1_36] |
| Q8A2Q1 | HMPREF0105_2037 | 60.676 | 99 | 0 | 900 | peptidase, S9A/B/C family, catalytic domain protein [Bacteroides sp. 3_1_33FAA] |
| Q8A2Q1 | BSEG_02165 | 60.405 | 99 | 0 | 898 | peptidase, S9A/B/C family, catalytic domain protein [Bacteroides dorei 5_1_36/D4] |
| Q8A2Q1 | CUU_0154 | 60.541 | 99 | 0 | 898 | dipeptidyl peptidase IV N-terminal domain protein [Bacteroides vulgatus PC510] |
| Q8A2Q1 | BSFG_00357 | 60.405 | 99 | 0 | 897 | peptidase, S9A/B/C family, catalytic domain protein [Bacteroides sp. 4_3_47FAA] |
| Q8A2Q1 | BACDOR_01902 | 60.541 | 99 | 0 | 897 | peptidase, S9A/B/C family, catalytic domain protein [Bacteroides dorei DSM 17855] |
| Q8A2Q1 | HMPREF9011_00439 | 60.405 | 99 | 0 | 897 | dipeptidyl peptidase IV [Bacteroides sp. 3_1_40A] |
| Q8A2Q1 | BSBG_00781 | 60.541 | 99 | 0 | 897 | peptidase, S9A/B/C family, catalytic domain protein [Bacteroides sp. 9_1_42FAA] |
| Q8A2Q1 | BACCOPRO_02325 | 57.084 | 97 | 0 | 837 | peptidase, S9A/B/C family, catalytic domain protein [Bacteroides coprophilus DSM 18228 = JCM 13818] |
| Q8A2Q1 | BACPLE_00583 | 56.463 | 98 | 0 | 834 | peptidase, S9A/B/C family, catalytic domain protein [Bacteroides plebeius DSM 17135] |
| Q8A2Q1 | BACCOP_02363 | 54.605 | 99 | 0 | 830 | peptidase, S9A/B/C family, catalytic domain protein [Bacteroides coprocola DSM 17136] |
| Q8A2Q1 | HMPREF9441_01209 | 52.005 | 99 | 0 | 769 | peptidase, S9A/B/C family, catalytic domain protein [Paraprevotella clara YIT 11840] |
| Q8A2Q1 | HMPREF9442_02116 | 51.793 | 100 | 0 | 768 | peptidase, S9A/B/C family, catalytic domain protein [Paraprevotella xylaniphila YIT 11841] |
| Q8A2Q1 | PARMER_02747 | 53.61 | 99 | 0 | 754 | peptidase, S9A/B/C family, catalytic domain protein [Parabacteroides merdae ATCC 43184] |
| Q8A2Q1 | PRABACTJOHN_00695 | 53.36 | 99 | 0 | 753 | peptidase, S9A/B/C family, catalytic domain protein [Parabacteroides johnsonii DSM 18315] |
| Q8A2Q1 | HMPREF0673_01428 | 51.075 | 99 | 0 | 744 | peptidase, S9A/B/C family, catalytic domain protein [Prevotella stercorea DSM 18206] |

| | | | | | |
|---|---|---|---|---|---|
| Q8A2Q1 | HMPREF9456_00374 | 51.15 | 99 | 0 | 738 hypothetical protein [Dysgonomonas mossii DSM 22836] |
| Q8A2Q1 | HMPREF9455_02807 | 50.882 | 99 | 0 | 734 hypothetical protein [Dysgonomonas gadei ATCC BAA-286] |
| Q8A2Q1 | HMPREF9008_03751 | 51.2 | 99 | 0 | 722 peptidase, S9A/B/C family, catalytic domain protein [Parabacteroides sp. 20_3] |
| Q8A2Q1 | HMPREF0104_00231 | 51.265 | 99 | 0 | 721 dipeptidyl aminopeptidase IV [Bacteroides sp. 3_1_19] |
| Q8A2Q1 | HMPREF0619_00460 | 50.933 | 99 | 0 | 720 peptidase, S9A/B/C family, catalytic domain protein [Parabacteroides sp. D13] |
| Q8A2Q1 | HMPREF0103_2069 | 51.132 | 99 | 0 | 719 peptidase, S9A/B/C family, catalytic domain protein [Bacteroides sp. 2_1_33B] |
| Q8A2Q1 | PREVCOP_05027 | 50.477 | 96 | 0 | 711 peptidase, S9A/B/C family, catalytic domain protein [Prevotella copri DSM 18205] |
| Q8A2Q1 | HMPREF1475_02283 | 50.761 | 97 | 0 | 698 hypothetical protein [Prevotella oralis HGA0225] |
| Q8A2Q1 | HMPREF9420_0906 | 46.505 | 99 | 0 | 673 peptidase, S9A/B/C family, catalytic domain protein [Prevotella salivae DSM 15606] |
| Q8A2Q1 | HMPREF1033_01873 | 48.735 | 99 | 0 | 669 hypothetical protein [Tannerella sp. 6_1_58FAA_CT1] |
| Q8A2Q1 | BSCG_03097 | 81.201 | 52 | 0 | 654 peptidase, S9A/B/C family, catalytic domain protein [Bacteroides sp. 2_2_4] |
| Q8A2Q1 | HMPREF9449_01075 | 42.72 | 97 | 0 | 563 hypothetical protein [Odoribacter laneus YIT 12061] |
| Q8A2Q1 | BSCG_03096 | 80.938 | 44 | 0 | 545 dipeptidyl peptidase IV N-terminal domain protein [Bacteroides sp. 2_2_4] |
| Q8A2Q1 | BSFG_01970 | 30.519 | 99 | 4.02E-86 | 291 peptidase, S9A/B/C family, catalytic domain protein [Bacteroides sp. 4_3_47FAA] |
| Q8A2Q1 | HMPREF9011_04151 | 30.519 | 99 | 4.02E-86 | 291 dipeptidyl peptidase IV [Bacteroides sp. 3_1_40A] |
| Q8A2Q1 | CUU_1533 | 30.519 | 99 | 4.15E-86 | 291 putative prolyl dipeptidyl peptidase [Bacteroides vulgatus PC510] |
| Q8A2Q1 | BSEG_03560 | 30.452 | 100 | 1.47E-85 | 289 peptidase, S9A/B/C family, catalytic domain protein [Bacteroides dorei 5_1_36/D4] |
| Q8A2Q1 | BACDOR_03064 | 30.452 | 100 | 1.47E-85 | 289 peptidase, S9A/B/C family, catalytic domain protein [Bacteroides dorei DSM 17855] |
| Q8A2Q1 | BSBG_03825 | 30.452 | 100 | 1.47E-85 | 289 peptidase, S9A/B/C family, catalytic domain protein [Bacteroides sp. 9_1_42FAA] |
| Q8A2Q1 | HMPREF0105_4156 | 30.233 | 100 | 2.34E-85 | 289 peptidase, S9A/B/C family, catalytic domain protein [Bacteroides sp. 3_1_33FAA] |
| Q8A2Q1 | HMPREF9449_00338 | 28.571 | 98 | 1.88E-83 | 283 hypothetical protein [Odoribacter laneus YIT 12061] |
| Q8A2Q1 | HMPREF9456_01479 | 29.384 | 84 | 1.27E-80 | 276 hypothetical protein [Dysgonomonas mossii DSM 22836] |
| Q8A2Q1 | BACCOP_03926 | 28.923 | 99 | 2.53E-80 | 275 peptidase, S9A/B/C family, catalytic domain protein [Bacteroides coprocola DSM 17136] |
| Q8A2Q1 | HMPREF9455_01995 | 29.261 | 88 | 1.17E-78 | 270 hypothetical protein [Dysgonomonas gadei ATCC BAA-286] |
| Q8A2Q1 | HMPREF9446_02000 | 30.882 | 80 | 2.61E-78 | 270 peptidase, S9A/B/C family, catalytic domain protein [Bacteroides fluxus YIT 12057] |
| Q8A2Q1 | HMPREF0969_00958 | 31.596 | 80 | 1.07E-77 | 268 peptidase, S9A/B/C family, catalytic domain protein [Bacteroides sp. D20] |
| Q8A2Q1 | BACUNI_03587 | 31.433 | 80 | 1.63E-77 | 268 peptidase, S9A/B/C family, catalytic domain protein [Bacteroides uniformis ATCC 8492] |
| Q8A2Q1 | HMPREF1007_03240 | 31.433 | 80 | 1.63E-77 | 268 dipeptidyl peptidase IV [Bacteroides sp. 4_1_36] |
| Q8A2Q1 | HMPREF9450_01296 | 28.799 | 89 | 5.87E-77 | 266 hypothetical protein [Alistipes indistinctus YIT 12060] |
| Q8A2Q1 | BACSTE_03271 | 30.998 | 76 | 9.95E-77 | 266 peptidase, S9A/B/C family, catalytic domain protein [Bacteroides stercoris ATCC 43183] |
| Q8A2Q1 | BACPLE_03112 | 28.982 | 99 | 1.22E-76 | 265 peptidase, S9A/B/C family, catalytic domain protein [Bacteroides plebeius DSM 17135] |
| Q8A2Q1 | BACCELL_03148 | 28.331 | 99 | 3.51E-76 | 264 peptidase, S9A/B/C family, catalytic domain protein [Bacteroides cellulosilyticus DSM 14838] |
| Q8A2Q1 | BACINT_04434 | 30.213 | 79 | 5.74E-75 | 262 peptidase, S9A/B/C family, catalytic domain protein [Bacteroides intestinalis DSM 17393] |
| Q8A2Q1 | HMPREF9445_02996 | 31.338 | 76 | 6.21E-75 | 261 peptidase, S9A/B/C family, catalytic domain protein [Bacteroides clarus YIT 12056] |
| Q8A2Q1 | HMPREF0103_2777 | 27.332 | 99 | 3.92E-74 | 259 peptidase, S9A/B/C family, catalytic domain protein [Bacteroides sp. 2_1_33B] |
| Q8A2Q1 | PARMER_03777 | 29.868 | 79 | 1.01E-73 | 257 peptidase, S9A/B/C family, catalytic domain protein [Parabacteroides merdae ATCC 43184] |
| Q8A2Q1 | HMPREF0104_02092 | 27.599 | 99 | 1.24E-73 | 257 dipeptidyl peptidase IV [Bacteroides sp. 3_1_19] |
| Q8A2Q1 | BACEGG_00881 | 30.986 | 76 | 1.49E-73 | 257 peptidase, S9A/B/C family, catalytic domain protein [Bacteroides eggerthii DSM 20697] |
| Q8A2Q1 | HMPREF1016_01518 | 30.986 | 76 | 1.54E-73 | 257 dipeptidyl peptidase IV [Bacteroides eggerthii 1_2_48FAA] |
| Q8A2Q1 | HMPREF0619_02963 | 27.471 | 99 | 2.57E-73 | 256 peptidase, S9A/B/C family, catalytic domain protein [Parabacteroides sp. D13] |
| Q8A2Q1 | HMPREF0103_2768 | 29.511 | 86 | 2.90E-73 | 256 peptidase, S9A/B/C family, catalytic domain protein [Bacteroides sp. 2_1_33B] |
| Q8A2Q1 | HMPREF9448_00433 | 28.665 | 99 | 3.34E-73 | 256 hypothetical protein [Barnesiella intestinihominis YIT 11860] |
| Q8A2Q1 | BACCOPRO_03615 | 29.079 | 99 | 3.43E-73 | 256 peptidase, S9A/B/C family, catalytic domain protein [Bacteroides coprophilus DSM 18228 = JCM 13818] |
| Q8A2Q1 | HMPREF0104_02101 | 29.358 | 86 | 3.67E-73 | 256 dipeptidyl-peptidase IV [Bacteroides sp. 3_1_19] |
| Q8A2Q1 | HMPREF9447_04168 | 28.719 | 99 | 3.73E-73 | 256 hypothetical protein [Bacteroides oleiciplenus YIT 12058] |
| Q8A2Q1 | BACPLE_01539 | 30.897 | 79 | 6.67E-73 | 255 peptidase, S9A/B/C family, catalytic domain protein [Bacteroides plebeius DSM 17135] |
| Q8A2Q1 | BSIG_2455 | 30.656 | 79 | 7.35E-73 | 256 hypothetical protein [Bacteroides sp. 1_1_6] |
| Q8A2Q1 | HMPREF0619_02972 | 29.358 | 86 | 9.75E-73 | 254 peptidase, S9A/B/C family, catalytic domain protein [Parabacteroides sp. D13] |
| Q8A2Q1 | HMPREF9008_02510 | 29.358 | 86 | 1.30E-72 | 254 peptidase, S9A/B/C family, catalytic domain protein [Parabacteroides sp. 20_3] |
| Q8A2Q1 | HMPREF1532_00929 | 30.428 | 79 | 1.47E-72 | 254 hypothetical protein [Bacteroides salyersiae WAL 10018 = DSM 18765 = JCM 12988] |
| Q8A2Q1 | BFAG_00311 | 29.441 | 79 | 2.26E-72 | 253 peptidase, S9A/B/C family, catalytic domain protein [Bacteroides fragilis 3_1_12] |
| Q8A2Q1 | HMPREF9008_02521 | 27.051 | 99 | 2.80E-72 | 254 peptidase, S9A/B/C family, catalytic domain protein [Parabacteroides sp. 20_3] |
| Q8A2Q1 | HMPREF9007_00426 | 30.492 | 79 | 5.40E-72 | 253 dipeptidyl-peptidase IV [Bacteroides sp. 1_1_14] |
| Q8A2Q1 | BACCAC_01795 | 30.428 | 79 | 9.30E-72 | 252 peptidase, S9A/B/C family, catalytic domain protein [Bacteroides caccae ATCC 43185] |
| Q8A2Q1 | BACOVA_05046 | 30.492 | 79 | 1.08E-71 | 251 peptidase, S9A/B/C family, catalytic domain protein [Bacteroides ovatus ATCC 8483] |
| Q8A2Q1 | HMPREF0127_04782 | 30.492 | 79 | 2.28E-71 | 251 hypothetical protein [Bacteroides sp. 1_1_30] |
| Q8A2Q1 | BXY_33000 | 30.492 | 79 | 2.28E-71 | 251 dipeptidyl-peptidase IV . Serine peptidase. MEROPS family S09B [Bacteroides xylanisolvens XB1A] |
| Q8A2Q1 | HMPREF0101_00028 | 29.739 | 80 | 2.55E-71 | 251 peptidase, S9A/B/C family, catalytic domain protein [Bacteroides sp. 2_1_16] |
| Q8A2Q1 | PREVCOP_04755 | 30.556 | 84 | 3.08E-71 | 250 peptidase, S9A/B/C family, catalytic domain protein [Prevotella copri DSM 18205] |
| Q8A2Q1 | BSGG_1381 | 29.819 | 79 | 3.29E-71 | 251 hypothetical protein [Bacteroides sp. D2] |

| | | | | | | |
|---|---|---|---|---|---|---|
| Q8A2Q1 | BSHG_0669 | 29.739 | 80 | 3.56E-71 | 250 | hypothetical protein [Bacteroides sp. 3_2_5] |
| Q8A2Q1 | HMPREF1018_00999 | 29.739 | 80 | 3.75E-71 | 250 | hypothetical protein [Bacteroides sp. 2_1_56FAA] |
| Q8A2Q1 | HMPREF9010_02811 | 29.819 | 79 | 4.73E-71 | 250 | dipeptidyl-peptidase IV [Bacteroides sp. 3_1_23] |
| Q8A2Q1 | HMPREF1017_02246 | 29.819 | 79 | 7.17E-71 | 249 | hypothetical protein [Bacteroides ovatus 3_8_47FAA] |
| Q8A2Q1 | BSCG_01817 | 29.819 | 79 | 7.31E-71 | 249 | peptidase, S9A/B/C family, catalytic domain protein [Bacteroides sp. 2_2_4] |
| Q8A2Q1 | HMPREF0106_02634 | 29.885 | 79 | 1.49E-70 | 249 | dipeptidyl-peptidase IV [Bacteroides sp. D22] |
| Q8A2Q1 | PRABACTJOHN_02440 | 29.043 | 79 | 2.65E-70 | 248 | peptidase, S9A/B/C family, catalytic domain protein [Parabacteroides johnsonii DSM 18315] |
| Q8A2Q1 | HMPREF0102_03488 | 30.328 | 79 | 3.34E-70 | 248 | peptidase, S9A/B/C family, catalytic domain protein [Bacteroides sp. 2_1_22] |
| Q8A2Q1 | BSAG_01729 | 30.328 | 79 | 3.34E-70 | 248 | peptidase, S9A/B/C family, catalytic domain protein [Bacteroides sp. D1] |
| Q8A2Q1 | CW3_2404 | 30.328 | 79 | 3.34E-70 | 248 | dipeptidyl peptidase IV N-terminal domain protein [Bacteroides xylanisolvens SD CC 1b] |
| Q8A2Q1 | CW1_1547 | 30.328 | 79 | 3.34E-70 | 248 | dipeptidyl peptidase IV N-terminal domain protein [Bacteroides xylanisolvens SD CC 2a] |
| Q8A2Q1 | CUY_0743 | 29.885 | 79 | 4.13E-70 | 248 | dipeptidyl peptidase IV N-terminal domain protein [Bacteroides ovatus SD CMC 3f] |
| Q8A2Q1 | BACFIN_08852 | 29.934 | 79 | 4.65E-69 | 245 | peptidase, S9A/B/C family, catalytic domain protein [Bacteroides finegoldii DSM 17565] |
| Q8A2Q1 | HMPREF9442_00881 | 28.113 | 100 | 1.76E-67 | 240 | peptidase, S9A/B/C family, catalytic domain protein [Paraprevotella xylaniphila YIT 11841] |
| Q8A2Q1 | PRABACTJOHN_03838 | 28.956 | 78 | 1.06E-66 | 239 | peptidase, S9A/B/C family, catalytic domain protein [Parabacteroides johnsonii DSM 18315] |
| Q8A2Q1 | HMPREF1214_01399 | 29.934 | 79 | 1.11E-66 | 238 | dipeptidyl-peptidase 4 [Bacteroides sp. HPS0048] |
| Q8A2Q1 | PARMER_01283 | 28.836 | 78 | 2.58E-66 | 238 | peptidase, S9A/B/C family, catalytic domain protein [Parabacteroides merdae ATCC 43184] |
| Q8A2Q1 | HMPREF9011_03920 | 28.904 | 79 | 2.69E-64 | 228 | dipeptidyl aminopeptidase [Bacteroides sp. 3_1_40A] |
| Q8A2Q1 | HMPREF1475_00504 | 29.984 | 80 | 4.92E-64 | 231 | hypothetical protein [Prevotella oralis HGA0225] |
| Q8A2Q1 | BACDOR_00958 | 27.297 | 99 | 9.44E-64 | 230 | peptidase, S9A/B/C family, catalytic domain protein [Bacteroides dorei DSM 17855] |
| Q8A2Q1 | BSFG_00985 | 27.034 | 99 | 9.83E-64 | 230 | peptidase, S9A/B/C family, catalytic domain protein [Bacteroides sp. 4_3_47FAA] |
| Q8A2Q1 | BSEG_00993 | 27.297 | 99 | 1.03E-63 | 230 | peptidase, S9A/B/C family, catalytic domain protein [Bacteroides dorei 5_1_36/D4] |
| Q8A2Q1 | HMPREF0105_1254 | 27.297 | 99 | 1.18E-63 | 230 | peptidase, S9A/B/C family, catalytic domain protein [Bacteroides sp. 3_1_33FAA] |
| Q8A2Q1 | CUU_2370 | 27.034 | 99 | 1.21E-63 | 230 | dipeptidyl peptidase IV N-terminal domain protein [Bacteroides vulgatus PC510] |
| Q8A2Q1 | BSBG_00102 | 27.297 | 99 | 1.26E-63 | 230 | peptidase, S9A/B/C family, catalytic domain protein [Bacteroides sp. 9_1_42FAA] |
| Q8A2Q1 | HMPREF9441_02325 | 27.344 | 99 | 1.82E-63 | 229 | peptidase, S9A/B/C family, catalytic domain protein [Paraprevotella clara YIT 11840] |
| Q8A2Q1 | HMPREF9420_0548 | 30.42 | 76 | 1.88E-63 | 229 | peptidase, S9A/B/C family, catalytic domain protein [Prevotella salivae DSM 15606] |
| Q8A2Q1 | ALIPUT_02419 | 28.286 | 79 | 3.13E-62 | 225 | peptidase, S9A/B/C family, catalytic domain protein [Alistipes putredinis DSM 17216] |
| Q8A2Q1 | HMPREF0673_01202 | 25.969 | 99 | 2.11E-61 | 223 | peptidase, S9A/B/C family, catalytic domain protein [Prevotella stercorea DSM 18206] |
| Q8A2Q1 | HMPREF1033_00581 | 27.72 | 86 | 4.15E-60 | 219 | hypothetical protein [Tannerella sp. 6_1_58FAA_CT1] |
| Q8A2Q1 | HMPREF9449_02863 | 25.795 | 74 | 3.95E-41 | 165 | hypothetical protein [Odoribacter laneus YIT 12061] |
| Q8A2Q1 | HMPREF9455_01475 | 25.856 | 77 | 5.28E-41 | 165 | hypothetical protein [Dysgonomonas gadei ATCC BAA-286] |
| Q8A2Q1 | BSIG_1145 | 27.235 | 64 | 3.33E-40 | 162 | hypothetical protein [Bacteroides sp. 1_1_6] |
| Q8A2Q1 | BXY_24100 | 25.85 | 76 | 3.82E-40 | 162 | Dipeptidyl peptidase IV (DPP IV) N-terminal region./Prolyl oligopeptidase family. [Bacteroides xylanisolvens XB1A] |
| Q8A2Q1 | HMPREF0102_00577 | 25.85 | 76 | 1.34E-39 | 161 | peptidase, S9A/B/C family, catalytic domain protein [Bacteroides sp. 2_1_22] |
| Q8A2Q1 | BSAG_00327 | 25.85 | 76 | 1.34E-39 | 161 | peptidase, S9A/B/C family, catalytic domain protein [Bacteroides sp. D1] |
| Q8A2Q1 | CW3_2061 | 25.85 | 76 | 1.37E-39 | 160 | peptidase, S9A/B/C familie, catalytic domain protein [Bacteroides xylanisolvens SD CC 1b] |
| Q8A2Q1 | CW1_0763 | 25.85 | 76 | 1.37E-39 | 160 | peptidase, S9A/B/C familie, catalytic domain protein [Bacteroides xylanisolvens SD CC 2a] |
| Q8A2Q1 | BACCAC_01476 | 26.576 | 76 | 2.02E-39 | 160 | peptidase, S9A/B/C family, catalytic domain protein [Bacteroides caccae ATCC 43185] |
| Q8A2Q1 | BACCELL_01959 | 26.819 | 64 | 4.86E-38 | 156 | peptidase, S9A/B/C family, catalytic domain protein [Bacteroides cellulosilyticus DSM 14838] |
| Q8A2Q1 | BACFIN_08494 | 28.407 | 68 | 1.16E-37 | 155 | peptidase, S9A/B/C family, catalytic domain protein [Bacteroides finegoldii DSM 17565] |
| Q8A2Q1 | BSCG_03701 | 26.884 | 64 | 1.41E-37 | 154 | peptidase, S9A/B/C family, catalytic domain protein [Bacteroides sp. 2_2_4] |
| Q8A2Q1 | HMPREF0104_00820 | 26.783 | 74 | 1.97E-37 | 154 | Xaa-Pro dipeptidyl-peptidase [Bacteroides sp. 3_1_19] |
| Q8A2Q1 | PRABACTJOHN_04237 | 24.602 | 74 | 2.98E-37 | 153 | peptidase, S9A/B/C family, catalytic domain protein [Parabacteroides johnsonii DSM 18315] |
| Q8A2Q1 | BSGG_2341 | 26.61 | 77 | 3.54E-37 | 153 | hypothetical protein [Bacteroides sp. D2] |
| Q8A2Q1 | HMPREF1017_00556 | 26.61 | 77 | 4.01E-37 | 153 | hypothetical protein [Bacteroides ovatus 3_8_47FAA] |
| Q8A2Q1 | BACCOP_03698 | 25.621 | 69 | 4.10E-37 | 153 | peptidase, S9A/B/C family, catalytic domain protein [Bacteroides coprocola DSM 17136] |
| Q8A2Q1 | HMPREF0105_2009 | 26.872 | 61 | 4.21E-37 | 153 | peptidase, S9A/B/C family, catalytic domain protein [Bacteroides sp. 3_1_33FAA] |
| Q8A2Q1 | BACINT_00527 | 26.611 | 64 | 4.45E-37 | 153 | peptidase, S9A/B/C family, catalytic domain protein [Bacteroides intestinalis DSM 17393] |
| Q8A2Q1 | PARMER_01642 | 24.779 | 74 | 4.87E-37 | 152 | peptidase, S9A/B/C family, catalytic domain protein [Parabacteroides merdae ATCC 43184] |
| Q8A2Q1 | HMPREF0103_1483 | 26.609 | 74 | 6.15E-37 | 153 | peptidase, S9A/B/C family, catalytic domain protein [Bacteroides sp. 2_1_33B] |
| Q8A2Q1 | PARMER_02220 | 26.736 | 74 | 7.79E-37 | 152 | peptidase, S9A/B/C family, catalytic domain protein [Parabacteroides merdae ATCC 43184] |
| Q8A2Q1 | HMPREF9008_01562 | 28.099 | 64 | 1.08E-36 | 152 | peptidase, S9A/B/C family, catalytic domain protein [Parabacteroides sp. 20_3] |
| Q8A2Q1 | HMPREF0619_00930 | 27.893 | 64 | 2.33E-36 | 151 | peptidase, S9A/B/C family, catalytic domain protein [Parabacteroides sp. D13] |
| Q8A2Q1 | BACOVA_00890 | 26.644 | 77 | 3.47E-36 | 150 | peptidase, S9A/B/C family, catalytic domain protein [Bacteroides ovatus ATCC 8483] |
| Q8A2Q1 | HMPREF9442_00872 | 25.625 | 64 | 3.54E-36 | 150 | peptidase, S9A/B/C family, catalytic domain protein [Paraprevotella xylaniphila YIT 11841] |
| Q8A2Q1 | BSEG_02195 | 25.988 | 64 | 7.59E-36 | 149 | peptidase, S9A/B/C family, catalytic domain protein [Bacteroides dorei 5_1_36/D4] |
| Q8A2Q1 | HMPREF9441_02554 | 25.311 | 64 | 1.73E-35 | 148 | peptidase, S9A/B/C family, catalytic domain protein [Paraprevotella clara YIT 11840] |
| Q8A2Q1 | BACFIN_08493 | 25.487 | 74 | 3.20E-35 | 147 | peptidase, S9A/B/C family, catalytic domain protein [Bacteroides finegoldii DSM 17565] |
| Q8A2Q1 | CUU_0128 | 25.78 | 64 | 3.36E-35 | 147 | peptidase, S9A/B/C familie, catalytic domain protein [Bacteroides vulgatus PC510] |

| | | | | | | |
|---|---|---|---|---|---|---|
| Q8A2Q1 | BACDOR_01931 | 25.572 | 64 | 3.72E-35 | 147 | peptidase, S9A/B/C family, catalytic domain protein [Bacteroides dorei DSM 17855] |
| Q8A2Q1 | BSBG_00751 | 25.572 | 64 | 4.03E-35 | 147 | peptidase, S9A/B/C family, catalytic domain protein [Bacteroides sp. 9_1_42FAA] |
| Q8A2Q1 | BSFG_00330 | 25.78 | 64 | 4.53E-35 | 147 | peptidase, S9A/B/C family, catalytic domain protein [Bacteroides sp. 4_3_47FAA] |
| Q8A2Q1 | HMPREF9011_00413 | 25.78 | 64 | 4.53E-35 | 147 | peptidase [Bacteroides sp. 3_1_40A] |
| Q8A2Q1 | BSGG_2342 | 23.57 | 91 | 7.20E-34 | 143 | hypothetical protein [Bacteroides sp. D2] |
| Q8A2Q1 | HMPREF1017_00555 | 23.57 | 91 | 7.20E-34 | 143 | hypothetical protein [Bacteroides ovatus 3_8_47FAA] |
| Q8A2Q1 | BACOVA_00889 | 23.57 | 91 | 7.20E-34 | 143 | peptidase, S9A/B/C family, catalytic domain protein [Bacteroides ovatus ATCC 8483] |
| Q8A2Q1 | HMPREF9010_00743 | 24.559 | 79 | 1.02E-33 | 143 | Xaa-Pro dipeptidyl-peptidase [Bacteroides sp. 3_1_23] |
| Q8A2Q1 | BSCG_03068 | 24.559 | 79 | 1.28E-33 | 142 | peptidase, S9A/B/C family, catalytic domain protein [Bacteroides sp. 2_2_4] |
| Q8A2Q1 | HMPREF0106_04740 | 24.559 | 79 | 1.35E-33 | 142 | Xaa-Pro dipeptidyl-peptidase [Bacteroides sp. D22] |
| Q8A2Q1 | CUY_4412 | 24.559 | 79 | 1.40E-33 | 142 | peptidase, S9A/B/C familie, catalytic domain protein [Bacteroides ovatus SD CMC 3f] |
| Q8A2Q1 | HMPREF0127_01607 | 24.559 | 79 | 1.74E-33 | 142 | hypothetical protein [Bacteroides sp. 1_1_30] |
| Q8A2Q1 | BACCAC_01475 | 26.316 | 64 | 8.28E-33 | 140 | peptidase, S9A/B/C family, catalytic domain protein [Bacteroides caccae ATCC 43185] |
| Q8A2Q1 | HMPREF1033_02114 | 24.359 | 81 | 3.66E-32 | 138 | hypothetical protein [Tannerella sp. 6_1_58FAA_CT1] |
| Q8A2Q1 | BACFIN_08525 | 24.826 | 91 | 9.49E-32 | 137 | peptidase, S9A/B/C family, catalytic domain protein [Bacteroides finegoldii DSM 17565] |
| Q8A2L6 | BSIG_1382 | 99.719 | 100 | 0 | 1481 | hypothetical protein [Bacteroides sp. 1_1_6] |
| Q8A2L6 | HMPREF9007_03829 | 99.579 | 100 | 0 | 1478 | hypothetical protein [Bacteroides sp. 1_1_14] |
| Q8A2L6 | HMPREF0127_01618 | 91.585 | 100 | 0 | 1384 | hypothetical protein [Bacteroides sp. 1_1_30] |
| Q8A2L6 | HMPREF0106_04280 | 91.585 | 100 | 0 | 1384 | hypothetical protein [Bacteroides sp. D22] |
| Q8A2L6 | HMPREF1017_00565 | 91.725 | 100 | 0 | 1382 | hypothetical protein [Bacteroides ovatus 3_8_47FAA] |
| Q8A2L6 | BSCG_03059 | 91.725 | 100 | 0 | 1380 | hypothetical protein [Bacteroides sp. 2_2_4] |
| Q8A2L6 | BSGG_2332 | 91.585 | 100 | 0 | 1380 | hypothetical protein [Bacteroides sp. D2] |
| Q8A2L6 | BACOVA_00899 | 91.585 | 100 | 0 | 1380 | hypothetical protein [Bacteroides ovatus ATCC 8483] |
| Q8A2L6 | CUY_4404 | 91.585 | 100 | 0 | 1379 | conserved hypothetical protein [Bacteroides ovatus SD CMC 3f] |
| Q8A2L6 | HMPREF9010_00752 | 91.585 | 100 | 0 | 1379 | hypothetical protein [Bacteroides sp. 3_1_23] |
| Q8A2L6 | HMPREF0102_00568 | 91.164 | 100 | 0 | 1379 | hypothetical protein [Bacteroides sp. 2_1_22] |
| Q8A2L6 | BSAG_00318 | 91.164 | 100 | 0 | 1379 | hypothetical protein [Bacteroides sp. D1] |
| Q8A2L6 | CW3_2069 | 91.164 | 100 | 0 | 1379 | conserved hypothetical protein [Bacteroides xylanisolvens SD CC 1b] |
| Q8A2L6 | CW1_0771 | 91.164 | 100 | 0 | 1379 | conserved hypothetical protein [Bacteroides xylanisolvens SD CC 2a] |
| Q8A2L6 | BACCAC_01485 | 91.842 | 99 | 0 | 1375 | hypothetical protein [Bacteroides caccae ATCC 43185] |
| Q8A2L6 | BACFIN_05173 | 89.621 | 100 | 0 | 1355 | hypothetical protein [Bacteroides finegoldii DSM 17565] |
| Q8A2L6 | BXY_25380 | 92.464 | 97 | 0 | 1350 | dipeptidyl-peptidase 7. Serine peptidase. MEROPS family S46 [Bacteroides xylanisolvens XB1A] |
| Q8A2L6 | HMPREF1532_01775 | 80.591 | 99 | 0 | 1206 | hypothetical protein [Bacteroides salyersiae WAL 10018 = DSM 18765 = JCM 12988] |
| Q8A2L6 | HMPREF1214_00608 | 79.044 | 99 | 0 | 1190 | hypothetical protein [Bacteroides sp. HPS0048] |
| Q8A2L6 | BFAG_02873 | 78.298 | 99 | 0 | 1179 | hypothetical protein [Bacteroides fragilis 3_1_12] |
| Q8A2L6 | HMPREF0101_02131 | 79.046 | 97 | 0 | 1169 | hypothetical protein [Bacteroides sp. 2_1_16] |
| Q8A2L6 | HMPREF1018_00169 | 78.902 | 97 | 0 | 1167 | hypothetical protein [Bacteroides sp. 2_1_56FAA] |
| Q8A2L6 | BSHG_1458 | 78.902 | 97 | 0 | 1167 | hypothetical protein [Bacteroides sp. 3_2_5] |
| Q8A2L6 | HMPREF0969_01517 | 76.611 | 99 | 0 | 1148 | hypothetical protein [Bacteroides sp. D20] |
| Q8A2L6 | HMPREF1007_00282 | 76.471 | 99 | 0 | 1146 | peptidase S46 [Bacteroides sp. 4_1_36] |
| Q8A2L6 | HMPREF9445_01804 | 76.718 | 99 | 0 | 1145 | hypothetical protein [Bacteroides clarus YIT 12056] |
| Q8A2L6 | BACUNI_01062 | 76.331 | 99 | 0 | 1143 | hypothetical protein [Bacteroides uniformis ATCC 8492] |
| Q8A2L6 | HMPREF9446_00563 | 75.387 | 99 | 0 | 1139 | hypothetical protein [Bacteroides fluxus YIT 12057] |
| Q8A2L6 | BACSTE_02429 | 75.562 | 99 | 0 | 1132 | hypothetical protein [Bacteroides stercoris ATCC 43183] |
| Q8A2L6 | HMPREF9447_02263 | 74.438 | 99 | 0 | 1127 | hypothetical protein [Bacteroides oleiciplenus YIT 12058] |
| Q8A2L6 | BACCELL_01726 | 73.418 | 99 | 0 | 1123 | hypothetical protein [Bacteroides cellulosilyticus DSM 14838] |
| Q8A2L6 | BACINT_01296 | 73.558 | 99 | 0 | 1121 | hypothetical protein [Bacteroides intestinalis DSM 17393] |
| Q8A2L6 | HMPREF1016_00890 | 73.315 | 99 | 0 | 1115 | peptidase S46 [Bacteroides eggerthii 1_2_48FAA] |
| Q8A2L6 | BACEGG_03368 | 73.315 | 99 | 0 | 1115 | hypothetical protein [Bacteroides eggerthii DSM 20697] |
| Q8A2L6 | CUU_0716 | 63.256 | 97 | 0 | 944 | conserved hypothetical protein [Bacteroides vulgatus PC510] |
| Q8A2L6 | BSFG_00163 | 63.112 | 97 | 0 | 942 | hypothetical protein [Bacteroides sp. 4_3_47FAA] |
| Q8A2L6 | HMPREF9011_00248 | 63.112 | 97 | 0 | 942 | hypothetical protein [Bacteroides sp. 3_1_40A] |
| Q8A2L6 | HMPREF0105_3274 | 63.203 | 97 | 0 | 940 | hypothetical protein [Bacteroides sp. 3_1_33FAA] |
| Q8A2L6 | BSBG_03122 | 63.203 | 97 | 0 | 940 | hypothetical protein [Bacteroides sp. 9_1_42FAA] |
| Q8A2L6 | BSEG_03017 | 63.203 | 97 | 0 | 940 | hypothetical protein [Bacteroides dorei 5_1_36/D4] |
| Q8A2L6 | BACDOR_02152 | 63.203 | 97 | 0 | 940 | hypothetical protein [Bacteroides dorei DSM 17855] |
| Q8A2L6 | HMPREF0103_2247 | 48 | 98 | 0 | 677 | hypothetical protein [Bacteroides sp. 2_1_33B] |
| Q8A2L6 | HMPREF0619_00047 | 47.857 | 98 | 0 | 676 | hypothetical protein [Parabacteroides sp. D13] |
| Q8A2L6 | HMPREF9008_04196 | 47.714 | 98 | 0 | 675 | hypothetical protein [Parabacteroides sp. 20_3] |
| Q8A2L6 | PARMER_00304 | 47.714 | 98 | 0 | 675 | hypothetical protein [Parabacteroides merdae ATCC 43184] |

| | | | | | | |
|---|---|---|---|---|---|---|
| Q8A2L6 | HMPREF0104_00053 | 47.571 | 98 | 0 | 673 | hypothetical protein [Bacteroides sp. 3_1_19] |
| Q8A2L6 | PRABACTJOHN_00535 | 47.714 | 98 | 0 | 672 | hypothetical protein [Parabacteroides johnsonii DSM 18315] |
| Q8A2L6 | HMPREF9441_00315 | 46.961 | 99 | 0 | 665 | hypothetical protein [Paraprevotella clara YIT 11840] |
| Q8A2L6 | HMPREF9442_02956 | 46.703 | 99 | 0 | 659 | hypothetical protein [Paraprevotella xylaniphila YIT 11841] |
| Q8A2L6 | HMPREF9456_01457 | 44.183 | 99 | 0 | 617 | hypothetical protein [Dysgonomonas mossii DSM 22836] |
| Q8A2L6 | HMPREF9455_01975 | 44.615 | 99 | 0 | 616 | hypothetical protein [Dysgonomonas gadei ATCC BAA-286] |
| Q8A2L6 | HMPREF9420_1755 | 45.969 | 98 | 0 | 608 | hypothetical protein [Prevotella salivae DSM 15606] |
| Q8A2L6 | PREVCOP_06589 | 44.847 | 99 | 0 | 602 | hypothetical protein [Prevotella copri DSM 18205] |
| Q8A2L6 | HMPREF1475_01683 | 45.455 | 98 | 0 | 600 | hypothetical protein [Prevotella oralis HGA0225] |
| Q8A2L6 | HMPREF9450_00755 | 42.6 | 100 | 0 | 568 | hypothetical protein [Alistipes indistinctus YIT 12060] |
| Q8A2L6 | HMPREF0969_00399 | 42.996 | 99 | 0 | 558 | hypothetical protein [Bacteroides sp. D20] |
| Q8A2L6 | BACUNI_01423 | 42.996 | 99 | 0 | 558 | hypothetical protein [Bacteroides uniformis ATCC 8492] |
| Q8A2L6 | HMPREF1007_01538 | 42.996 | 99 | 0 | 558 | peptidase S46 [Bacteroides sp. 4_1_36] |
| Q8A2L6 | ALIPUT_02112 | 42.698 | 99 | 0 | 555 | hypothetical protein [Alistipes putredinis DSM 17216] |
| Q8A2L6 | HMPREF1033_00928 | 41.46 | 100 | 0 | 551 | hypothetical protein [Tannerella sp. 6_1_58FAA_CT1] |
| Q8A2L6 | HMPREF9446_00758 | 41.77 | 99 | 0 | 551 | hypothetical protein [Bacteroides fluxus YIT 12057] |
| Q8A2L6 | HMPREF0673_00993 | 43.208 | 97 | 0 | 547 | hypothetical protein [Prevotella stercorea DSM 18206] |
| Q8A2L6 | HMPREF9445_00327 | 40.903 | 99 | 0 | 541 | hypothetical protein [Bacteroides clarus YIT 12056] |
| Q8A2L6 | BACCOP_03276 | 40.741 | 99 | 0 | 539 | hypothetical protein [Bacteroides coprocola DSM 17136] |
| Q8A2L6 | BACCOPRO_02783 | 41.632 | 98 | 0 | 538 | hypothetical protein [Bacteroides coprophilus DSM 18228 = JCM 13818] |
| Q8A2L6 | AL1_16030 | 41.433 | 99 | 0 | 538 | Peptidase S46. [Alistipes shahii WAL 8301] |
| Q8A2L6 | BACCELL_05142 | 41.541 | 99 | 0 | 537 | hypothetical protein [Bacteroides cellulosilyticus DSM 14838] |
| Q8A2L6 | BACPLE_01878 | 41.103 | 99 | 3.77E-180 | 535 | hypothetical protein [Bacteroides plebeius DSM 17135] |
| Q8A2L6 | HMPREF9449_02133 | 39.726 | 99 | 5.18E-180 | 534 | hypothetical protein [Odoribacter laneus YIT 12061] |
| Q8A2L6 | HMPREF1016_00219 | 41.803 | 99 | 6.86E-179 | 531 | peptidase S46 [Bacteroides eggerthii 1_2_48FAA] |
| Q8A2L6 | HMPREF0105_0857 | 42.254 | 97 | 2.13E-178 | 530 | hypothetical protein [Bacteroides sp. 3_1_33FAA] |
| Q8A2L6 | BSEG_00914 | 42.254 | 97 | 2.13E-178 | 530 | hypothetical protein [Bacteroides dorei 5_1_36/D4] |
| Q8A2L6 | BSBG_02476 | 42.254 | 97 | 2.13E-178 | 530 | hypothetical protein [Bacteroides sp. 9_1_42FAA] |
| Q8A2L6 | BACDOR_03540 | 42.113 | 97 | 4.03E-178 | 529 | hypothetical protein [Bacteroides dorei DSM 17855] |
| Q8A2L6 | BACEGG_02070 | 41.667 | 99 | 4.75E-178 | 529 | hypothetical protein [Bacteroides eggerthii DSM 20697] |
| Q8A2L6 | BACINT_02882 | 40.771 | 99 | 6.50E-178 | 529 | hypothetical protein [Bacteroides intestinalis DSM 17393] |
| Q8A2L6 | BSFG_01936 | 42.113 | 97 | 9.37E-178 | 528 | hypothetical protein [Bacteroides sp. 4_3_47FAA] |
| Q8A2L6 | HMPREF9011_02238 | 42.113 | 97 | 9.37E-178 | 528 | hypothetical protein [Bacteroides sp. 3_1_40A] |
| Q8A2L6 | CUU_0388 | 42.113 | 97 | 1.16E-177 | 528 | conserved hypothetical protein [Bacteroides vulgatus PC510] |
| Q8A2L6 | BACSTE_01784 | 40.608 | 99 | 8.24E-177 | 526 | hypothetical protein [Bacteroides stercoris ATCC 43183] |
| Q8A2L6 | BFAG_03217 | 40.89 | 98 | 4.29E-176 | 524 | hypothetical protein [Bacteroides fragilis 3_1_12] |
| Q8A2L6 | BACFIN_07388 | 40.463 | 99 | 9.57E-176 | 523 | hypothetical protein [Bacteroides finegoldii DSM 17565] |
| Q8A2L6 | HMPREF9447_03715 | 40.083 | 99 | 6.27E-175 | 521 | hypothetical protein [Bacteroides oleiciplenus YIT 12058] |
| Q8A2L6 | BSIG_0060 | 41.044 | 97 | 6.82E-174 | 518 | hypothetical protein [Bacteroides sp. 1_1_6] |
| Q8A2L6 | HMPREF1532_00292 | 40.405 | 99 | 1.45E-173 | 518 | hypothetical protein [Bacteroides salyersiae WAL 10018 = DSM 18765 = JCM 12988] |
| Q8A2L6 | HMPREF0106_04191 | 40.655 | 99 | 5.14E-173 | 516 | hypothetical protein [Bacteroides sp. D22] |
| Q8A2L6 | BXY_42310 | 40.655 | 99 | 5.20E-173 | 516 | dipeptidyl-peptidase 7. Serine peptidase. MEROPS family S46 [Bacteroides xylanisolvens XB1A] |
| Q8A2L6 | BSHG_1934 | 40.73 | 98 | 7.40E-173 | 516 | hypothetical protein [Bacteroides sp. 3_2_5] |
| Q8A2L6 | HMPREF1018_03521 | 40.73 | 98 | 7.40E-173 | 516 | hypothetical protein [Bacteroides sp. 2_1_56FAA] |
| Q8A2L6 | HMPREF0101_03127 | 40.73 | 98 | 7.40E-173 | 516 | hypothetical protein [Bacteroides sp. 2_1_16] |
| Q8A2L6 | HMPREF1214_03868 | 40.382 | 99 | 1.08E-172 | 515 | hypothetical protein [Bacteroides sp. HPS0048] |
| Q8A2L6 | HMPREF0127_02920 | 40.518 | 99 | 2.03E-172 | 514 | hypothetical protein [Bacteroides sp. 1_1_30] |
| Q8A2L6 | BACCAC_02237 | 39.589 | 99 | 3.67E-172 | 514 | hypothetical protein [Bacteroides caccae ATCC 43185] |
| Q8A2L6 | HMPREF0102_02504 | 40.518 | 99 | 5.41E-172 | 513 | hypothetical protein [Bacteroides sp. 2_1_22] |
| Q8A2L6 | BSAG_01080 | 40.518 | 99 | 5.41E-172 | 513 | hypothetical protein [Bacteroides sp. D1] |
| Q8A2L6 | CW3_4226 | 40.518 | 99 | 5.41E-172 | 513 | conserved hypothetical protein [Bacteroides xylanisolvens SD CC 1b] |
| Q8A2L6 | CW1_0171 | 40.518 | 99 | 5.41E-172 | 513 | conserved hypothetical protein [Bacteroides xylanisolvens SD CC 2a] |
| Q8A2L6 | HMPREF9007_01172 | 40.762 | 97 | 8.61E-172 | 513 | hypothetical protein [Bacteroides sp. 1_1_14] |
| Q8A2L6 | BSGG_0735 | 40.223 | 97 | 2.09E-171 | 512 | hypothetical protein [Bacteroides sp. D2] |
| Q8A2L6 | HMPREF1017_03509 | 40.223 | 97 | 2.09E-171 | 512 | hypothetical protein [Bacteroides ovatus 3_8_47FAA] |
| Q8A2L6 | BACOVA_00386 | 40.084 | 97 | 6.07E-171 | 511 | hypothetical protein [Bacteroides ovatus ATCC 8483] |
| Q8A2L6 | CUY_4106 | 40.084 | 97 | 8.77E-171 | 510 | conserved hypothetical protein [Bacteroides ovatus SD CMC 3f] |
| Q8A2L6 | BSCG_00818 | 40.084 | 97 | 3.85E-170 | 509 | hypothetical protein [Bacteroides sp. 2_2_4] |
| Q8A2L6 | BACCELL_05143 | 39.369 | 99 | 5.89E-169 | 506 | hypothetical protein [Bacteroides cellulosilyticus DSM 14838] |
| Q8A2L6 | HMPREF9447_03716 | 38.981 | 99 | 4.33E-168 | 503 | hypothetical protein [Bacteroides oleiciplenus YIT 12058] |

| | | | | | | |
|---|---|---|---|---|---|---|
| Q8A2L6 | BACINT_02883 | 39.369 | 99 | 4.83E-168 | 503 | hypothetical protein [Bacteroides intestinalis DSM 17393] |
| Q8A2L6 | HMPREF9010_01792 | 39.747 | 97 | 5.57E-168 | 502 | hypothetical protein [Bacteroides sp. 3_1_23] |
| Q8A2L6 | HMPREF1532_00293 | 38.387 | 99 | 4.48E-166 | 498 | hypothetical protein [Bacteroides salyersiae WAL 10018 = DSM 18765 = JCM 12988] |
| Q8A2L6 | HMPREF0969_00398 | 39.32 | 99 | 7.37E-166 | 498 | hypothetical protein [Bacteroides sp. D20] |
| Q8A2L6 | BACUNI_01424 | 39.184 | 99 | 5.30E-165 | 496 | hypothetical protein [Bacteroides uniformis ATCC 8492] |
| Q8A2L6 | HMPREF1007_01539 | 39.184 | 99 | 5.30E-165 | 496 | peptidase S46 [Bacteroides sp. 4_1_36] |
| Q8A2L6 | HMPREF9446_00759 | 39.251 | 99 | 7.09E-165 | 495 | hypothetical protein [Bacteroides fluxus YIT 12057] |
| Q8A2L6 | BACCAC_02236 | 38.633 | 98 | 2.32E-164 | 494 | hypothetical protein [Bacteroides caccae ATCC 43185] |
| Q8A2L6 | BACFIN_07387 | 38.075 | 98 | 4.94E-164 | 493 | hypothetical protein [Bacteroides finegoldii DSM 17565] |
| Q8A2L6 | BACSTE_01783 | 39.286 | 99 | 2.63E-163 | 491 | hypothetical protein [Bacteroides stercoris ATCC 43183] |
| Q8A2L6 | BSIG_0059 | 37.796 | 98 | 3.52E-163 | 491 | hypothetical protein [Bacteroides sp. 1_1_6] |
| Q8A2L6 | HMPREF0103_1442 | 38.43 | 99 | 5.23E-163 | 490 | hypothetical protein [Bacteroides sp. 2_1_33B] |
| Q8A2L6 | HMPREF1214_03869 | 38.719 | 98 | 1.44E-162 | 489 | hypothetical protein [Bacteroides sp. HPS0048] |
| Q8A2L6 | HMPREF0619_00990 | 38.592 | 97 | 1.46E-162 | 489 | hypothetical protein [Parabacteroides sp. D13] |
| Q8A2L6 | HMPREF0104_00878 | 38.592 | 97 | 1.46E-162 | 489 | hypothetical protein [Bacteroides sp. 3_1_19] |
| Q8A2L6 | HMPREF1016_00218 | 39.148 | 99 | 1.47E-162 | 489 | peptidase S46 [Bacteroides eggerthii 1_2_48FAA] |
| Q8A2L6 | BACEGG_02069 | 39.148 | 99 | 1.47E-162 | 489 | hypothetical protein [Bacteroides eggerthii DSM 20697] |
| Q8A2L6 | HMPREF9445_00326 | 38.798 | 99 | 1.74E-162 | 489 | hypothetical protein [Bacteroides clarus YIT 12056] |
| Q8A2L6 | HMPREF0127_02919 | 37.238 | 99 | 3.73E-162 | 488 | hypothetical protein [Bacteroides sp. 1_1_30] |
| Q8A2L6 | HMPREF9008_01598 | 38.292 | 99 | 6.04E-162 | 488 | hypothetical protein [Parabacteroides sp. 20_3] |
| Q8A2L6 | BXY_42300 | 37.099 | 99 | 1.68E-161 | 486 | dipeptidyl-peptidase 7. Serine peptidase. MEROPS family S46 [Bacteroides xylanisolvens XB1A] |
| Q8A2L6 | HMPREF0106_04190 | 37.099 | 99 | 1.68E-161 | 486 | hypothetical protein [Bacteroides sp. D22] |
| Q8A2L6 | HMPREF0102_02503 | 37.099 | 99 | 1.03E-160 | 484 | hypothetical protein [Bacteroides sp. 2_1_22] |
| Q8A2L6 | BSAG_01079 | 37.099 | 99 | 1.03E-160 | 484 | hypothetical protein [Bacteroides sp. D1] |
| Q8A2L6 | CW3_4225 | 37.099 | 99 | 1.03E-160 | 484 | conserved hypothetical protein [Bacteroides xylanisolvens SD CC 1b] |
| Q8A2L6 | CW1_0172 | 37.099 | 99 | 1.03E-160 | 484 | conserved hypothetical protein [Bacteroides xylanisolvens SD CC 2a] |
| Q8A2L6 | HMPREF1017_03508 | 37.378 | 99 | 1.42E-160 | 484 | hypothetical protein [Bacteroides ovatus 3_8_47FAA] |
| Q8A2L6 | BACCOP_03275 | 37.326 | 99 | 1.61E-160 | 484 | hypothetical protein [Bacteroides coprocola DSM 17136] |
| Q8A2L6 | BSGG_0736 | 37.378 | 99 | 1.74E-160 | 484 | hypothetical protein [Bacteroides sp. D2] |
| Q8A2L6 | HMPREF9010_01793 | 37.378 | 99 | 2.41E-160 | 484 | hypothetical protein [Bacteroides sp. 3_1_23] |
| Q8A2L6 | BACOVA_00385 | 37.378 | 99 | 2.41E-160 | 484 | hypothetical protein [Bacteroides ovatus ATCC 8483] |
| Q8A2L6 | CUY_4107 | 37.587 | 99 | 3.29E-160 | 483 | conserved hypothetical protein [Bacteroides ovatus SD CMC 3f] |
| Q8A2L6 | PRABACTJOHN_02704 | 38.056 | 99 | 6.07E-160 | 483 | hypothetical protein [Parabacteroides johnsonii DSM 18315] |
| Q8A2L6 | PARMER_02946 | 38.194 | 99 | 1.74E-159 | 481 | hypothetical protein [Parabacteroides merdae ATCC 43184] |
| Q8A2L6 | HMPREF1018_03522 | 37.689 | 99 | 3.07E-159 | 481 | hypothetical protein [Bacteroides sp. 2_1_56FAA] |
| Q8A2L6 | HMPREF0101_03126 | 37.636 | 99 | 3.21E-159 | 481 | hypothetical protein [Bacteroides sp. 2_1_16] |
| Q8A2L6 | BSHG_1935 | 37.636 | 99 | 3.24E-159 | 481 | hypothetical protein [Bacteroides sp. 3_2_5] |
| Q8A2L6 | BFAG_03218 | 37.568 | 99 | 4.44E-159 | 480 | hypothetical protein [Bacteroides fragilis 3_1_12] |
| Q8A2L6 | BACCOPRO_02782 | 37.723 | 100 | 5.32E-159 | 480 | hypothetical protein [Bacteroides coprophilus DSM 18228 = JCM 13818] |
| Q8A2L6 | BACPLE_01877 | 37.415 | 100 | 6.76E-159 | 480 | hypothetical protein [Bacteroides plebeius DSM 17135] |
| Q8A2L6 | HMPREF1033_02741 | 36.827 | 95 | 3.33E-156 | 472 | hypothetical protein [Tannerella sp. 6_1_58FAA_CT1] |
| Q8A2L6 | HMPREF9449_02174 | 36.842 | 99 | 8.10E-156 | 472 | hypothetical protein [Odoribacter laneus YIT 12061] |
| Q8A2L6 | BSCG_00819 | 37.179 | 96 | 1.17E-155 | 471 | hypothetical protein [Bacteroides sp. 2_2_4] |
| Q8A2L6 | BSFG_01937 | 36.856 | 99 | 4.51E-154 | 468 | hypothetical protein [Bacteroides sp. 4_3_47FAA] |
| Q8A2L6 | HMPREF9011_02237 | 36.856 | 99 | 4.51E-154 | 468 | hypothetical protein [Bacteroides sp. 3_1_40A] |
| Q8A2L6 | BSEG_00915 | 37.222 | 97 | 1.66E-153 | 466 | hypothetical protein [Bacteroides dorei 5_1_36/D4] |
| Q8A2L6 | HMPREF9449_00904 | 37.174 | 99 | 1.85E-153 | 466 | hypothetical protein [Odoribacter laneus YIT 12061] |
| Q8A2L6 | BACDOR_03539 | 37.222 | 97 | 1.97E-153 | 466 | hypothetical protein [Bacteroides dorei DSM 17855] |
| Q8A2L6 | HMPREF0105_0858 | 37.222 | 97 | 2.29E-153 | 466 | hypothetical protein [Bacteroides sp. 3_1_33FAA] |
| Q8A2L6 | BSBG_02477 | 37.222 | 97 | 3.95E-153 | 465 | hypothetical protein [Bacteroides sp. 9_1_42FAA] |
| Q8A2L6 | CUU_0387 | 36.585 | 99 | 9.86E-153 | 464 | conserved hypothetical protein [Bacteroides vulgatus PC510] |
| Q8A2L6 | HMPREF9450_02265 | 37.621 | 99 | 1.75E-149 | 455 | hypothetical protein [Alistipes indistinctus YIT 12060] |
| Q8A2L6 | AL1_16890 | 34.566 | 97 | 3.80E-121 | 380 | Peptidase S46. [Alistipes shahii WAL 8301] |
| Q8A2L6 | AL1_16630 | 33.425 | 99 | 7.57E-121 | 382 | Peptidase S46. [Alistipes shahii WAL 8301] |
| Q8A2L6 | ALIPUT_00511 | 33.713 | 97 | 9.59E-121 | 379 | hypothetical protein [Alistipes putredinis DSM 17216] |
| Q8A2L6 | BSGG_3154 | 28.802 | 99 | 2.10E-98 | 323 | hypothetical protein [Bacteroides sp. D2] |
| Q8A2L6 | BACOVA_04406 | 28.745 | 99 | 6.94E-98 | 322 | hypothetical protein [Bacteroides ovatus ATCC 8483] |
| Q8A0D0 | BSIG_2555 | 100 | 100 | 0 | 1368 | hypothetical protein [Bacteroides sp. 1_1_6] |
| Q8A0D0 | HMPREF9007_01646 | 99.849 | 100 | 0 | 1366 | sialic acid-specific 9-O-acetylesterase [Bacteroides sp. 1_1_14] |
| Q8A0D0 | BXY_31750 | 76.704 | 95 | 0 | 1036 | Domain of unknown function (DUF303). [Bacteroides xylanisolvens XB1A] |

| | | | | | | |
|---|---|---|---|---|---|---|
| Q8A0D0 | BACINT_01283 | 55.471 | 99 | 0 | 773 | glycosyl hydrolase family 2, sugar binding domain protein [Bacteroides intestinalis DSM 17393] |
| Q8A0D0 | HMPREF9447_02283 | 55.86 | 99 | 0 | 771 | hypothetical protein [Bacteroides oleiciplenus YIT 12058] |
| Q8A0D0 | BSHG_2842 | 55.235 | 99 | 0 | 757 | hypothetical protein [Bacteroides sp. 3_2_5] |
| Q8A0D0 | HMPREF1018_03421 | 55.235 | 99 | 0 | 757 | hypothetical protein [Bacteroides sp. 2_1_56FAA] |
| Q8A0D0 | HMPREF0101_03409 | 55.235 | 99 | 0 | 757 | glycosyl hydrolase family 2, sugar binding domain protein [Bacteroides sp. 2_1_16] |
| Q8A0D0 | HMPREF1532_04087 | 50.989 | 98 | 0 | 682 | hypothetical protein [Bacteroides salyersiae WAL 10018 = DSM 18765 = JCM 12988] |
| Q8A0D0 | HMPREF1214_04471 | 50.679 | 100 | 0 | 674 | sialate O-acetylesterase [Bacteroides sp. HPS0048] |
| Q8A0D0 | BACCELL_01764 | 53.177 | 90 | 0 | 668 | glycosyl hydrolase family 2, sugar binding domain protein [Bacteroides cellulosilyticus DSM 14838] |
| Q8A0D0 | BSCG_02827 | 49.691 | 98 | 0 | 650 | glycosyl hydrolase family 2, sugar binding domain protein [Bacteroides sp. 2_2_4] |
| Q8A0D0 | HMPREF0102_00368 | 50 | 98 | 0 | 649 | glycosyl hydrolase family 2, sugar binding domain protein [Bacteroides sp. 2_1_22] |
| Q8A0D0 | BSAG_00113 | 50 | 98 | 0 | 649 | glycosyl hydrolase family 2, sugar binding domain protein [Bacteroides sp. D1] |
| Q8A0D0 | HMPREF0127_02525 | 50.39 | 97 | 0 | 649 | hypothetical protein [Bacteroides sp. 1_1_30] |
| Q8A0D0 | CW3_1482 | 50.39 | 97 | 0 | 649 | glycosyl hydrolase family 2, sugar binding domain protein [Bacteroides xylanisolvens SD CC 1b] |
| Q8A0D0 | CW1_3865 | 50.39 | 97 | 0 | 649 | glycosyl hydrolase family 2, sugar binding domain protein [Bacteroides xylanisolvens SD CC 2a] |
| Q8A0D0 | BXY_27890 | 50.234 | 97 | 0 | 647 | Domain of unknown function (DUF303). [Bacteroides xylanisolvens XB1A] |
| Q8A0D0 | HMPREF9441_00871 | 47.812 | 96 | 0 | 629 | glycosyl hydrolase family 2, sugar binding domain protein [Paraprevotella clara YIT 11840] |
| Q8A0D0 | BSEG_01852 | 46.942 | 99 | 0 | 615 | glycosyl hydrolase family 2, sugar binding domain protein [Bacteroides dorei 5_1_36/D4] |
| Q8A0D0 | BACDOR_04782 | 46.789 | 99 | 0 | 615 | glycosyl hydrolase family 2, sugar binding domain protein [Bacteroides dorei DSM 17855] |
| Q8A0D0 | BSBG_01120 | 46.789 | 99 | 0 | 615 | glycosyl hydrolase family 2, sugar binding domain protein [Bacteroides sp. 9_1_42FAA] |
| Q8A0D0 | HMPREF0105_2389 | 46.636 | 99 | 0 | 613 | glycosyl hydrolase family 2, sugar binding domain protein [Bacteroides sp. 3_1_33FAA] |
| Q8A0D0 | HMPREF9442_01438 | 46.375 | 98 | 0 | 613 | glycosyl hydrolase family 2, sugar binding domain protein [Paraprevotella xylaniphila YIT 11841] |
| Q8A0D0 | HMPREF1033_02166 | 46.26 | 99 | 0 | 593 | hypothetical protein [Tannerella sp. 6_1_58FAA_CT1] |
| Q8A0D0 | HMPREF1017_00401 | 45.344 | 98 | 0 | 580 | hypothetical protein [Bacteroides ovatus 3_8_47FAA] |
| Q8A0D0 | HMPREF9007_03698 | 45.122 | 98 | 0 | 575 | sialate O-acetylesterase [Bacteroides sp. 1_1_14] |
| Q8A0D0 | BSIG_1519 | 44.97 | 98 | 0 | 573 | hypothetical protein [Bacteroides sp. 1_1_6] |
| Q8A0D0 | CUY_2134 | 49.393 | 87 | 0 | 570 | glycosyl hydrolase family 2, sugar binding domain protein [Bacteroides ovatus SD CMC 3f] |
| Q8A0D0 | HMPREF1532_01640 | 42.945 | 98 | 0 | 559 | hypothetical protein [Bacteroides salyersiae WAL 10018 = DSM 18765 = JCM 12988] |
| Q8A0D0 | HMPREF1214_00152 | 44.037 | 98 | 0 | 540 | sialate O-acetylesterase [Bacteroides sp. HPS0048] |
| Q8A0D0 | HMPREF9456_00126 | 46.119 | 98 | 0 | 536 | hypothetical protein [Dysgonomonas mossii DSM 22836] |
| Q8A0D0 | HMPREF9455_01452 | 45.06 | 99 | 1.01E-174 | 516 | hypothetical protein [Dysgonomonas gadei ATCC BAA-286] |
| Q8A0D0 | BACCELL_02606 | 44.158 | 99 | 4.33E-173 | 512 | glycosyl hydrolase family 2, sugar binding domain protein [Bacteroides cellulosilyticus DSM 14838] |
| Q8A0D0 | CUU_3743 | 39.084 | 99 | 6.93E-168 | 498 | glycosyl hydrolase family 2, sugar binding domain protein [Bacteroides vulgatus PC510] |
| Q8A0D0 | BSFG_04400 | 38.923 | 98 | 1.11E-166 | 495 | glycosyl hydrolase family 2, sugar binding domain protein [Bacteroides sp. 4_3_47FAA] |
| Q8A0D0 | BSBG_04549 | 38.897 | 98 | 1.14E-166 | 495 | glycosyl hydrolase family 2, sugar binding domain protein [Bacteroides sp. 9_1_42FAA] |
| Q8A0D0 | HMPREF0105_4170 | 38.591 | 98 | 7.77E-166 | 493 | glycosyl hydrolase family 2, sugar binding domain protein [Bacteroides sp. 3_1_33FAA] |
| Q8A0D0 | BACDOR_01315 | 38.744 | 98 | 1.13E-165 | 493 | glycosyl hydrolase family 2, sugar binding domain protein [Bacteroides dorei DSM 17855] |
| Q8A0D0 | BSGG_2683 | 42.249 | 98 | 1.12E-164 | 490 | hypothetical protein [Bacteroides sp. D2] |
| Q8A0D0 | HMPREF0106_00321 | 41.061 | 98 | 1.09E-160 | 480 | sialate O-acetylesterase [Bacteroides sp. D22] |
| Q8A0D0 | HMPREF0127_02122 | 41.123 | 98 | 1.37E-160 | 479 | hypothetical protein [Bacteroides sp. 1_1_30] |
| Q8A0D0 | HMPREF0102_00861 | 40.971 | 98 | 6.84E-160 | 478 | glycosyl hydrolase family 2, sugar binding domain protein [Bacteroides sp. 2_1_22] |
| Q8A0D0 | BSAG_00610 | 40.971 | 98 | 6.84E-160 | 478 | glycosyl hydrolase family 2, sugar binding domain protein [Bacteroides sp. D1] |
| Q8A0D0 | CW3_0062 | 40.971 | 98 | 6.84E-160 | 478 | glycosyl hydrolase family 2, sugar binding domain protein [Bacteroides xylanisolvens SD CC 1b] |
| Q8A0D0 | CW1_0059 | 40.971 | 98 | 6.84E-160 | 478 | glycosyl hydrolase family 2, sugar binding domain protein [Bacteroides xylanisolvens SD CC 2a] |
| Q8A0D0 | HMPREF1017_00242 | 40.668 | 98 | 1.21E-159 | 477 | hypothetical protein [Bacteroides ovatus 3_8_47FAA] |
| Q8A0D0 | BXY_21220 | 41.562 | 95 | 4.36E-159 | 475 | Domain of unknown function (DUF303). [Bacteroides xylanisolvens XB1A] |
| Q8A0D0 | BACOVA_02638 | 41.562 | 95 | 1.46E-158 | 474 | glycosyl hydrolase family 2, sugar binding domain protein [Bacteroides ovatus ATCC 8483] |
| Q8A0D0 | HMPREF9448_01755 | 41.088 | 98 | 3.08E-153 | 461 | hypothetical protein [Barnesiella intestinihominis YIT 11860] |
| Q8A0D0 | HMPREF1214_04111 | 38.897 | 99 | 2.37E-152 | 459 | hypothetical protein [Bacteroides sp. HPS0048] |
| Q8A0D0 | HMPREF9447_02070 | 40.465 | 95 | 3.41E-145 | 440 | hypothetical protein [Bacteroides oleiciplenus YIT 12058] |
| Q8A0D0 | BACCELL_01466 | 40.409 | 95 | 1.07E-142 | 433 | glycosyl hydrolase family 2, sugar binding domain protein [Bacteroides cellulosilyticus DSM 14838] |
| Q8A0D0 | HMPREF1016_02150 | 37.348 | 98 | 9.63E-132 | 405 | glycosyl hydrolase family 2 [Bacteroides eggerthii 1_2_48FAA] |
| Q8A0D0 | BACEGG_01318 | 37.348 | 98 | 9.63E-132 | 405 | glycosyl hydrolase family 2, sugar binding domain protein [Bacteroides eggerthii DSM 20697] |
| Q8A0D0 | BSAG_02204 | 35.964 | 98 | 2.34E-126 | 391 | glycosyl hydrolase family 2, sugar binding domain protein [Bacteroides sp. D1] |
| Q8A0D0 | CW3_1818 | 35.964 | 98 | 2.34E-126 | 391 | glycosyl hydrolase family 2, sugar binding domain protein [Bacteroides xylanisolvens SD CC 1b] |
| Q8A0D0 | CW1_2189 | 35.964 | 98 | 2.34E-126 | 391 | glycosyl hydrolase family 2, sugar binding domain protein [Bacteroides xylanisolvens SD CC 2a] |
| Q8A0D0 | HMPREF0102_00201 | 35.964 | 98 | 3.33E-126 | 391 | glycosyl hydrolase family 2, sugar binding domain protein [Bacteroides sp. 2_1_22] |
| Q8A0D0 | BACCELL_03403 | 35.479 | 98 | 4.27E-123 | 383 | glycosyl hydrolase family 2, sugar binding domain protein [Bacteroides cellulosilyticus DSM 14838] |
| Q8A0D0 | HMPREF9456_02283 | 36.391 | 98 | 4.56E-122 | 380 | hypothetical protein [Dysgonomonas mossii DSM 22836] |
| Q8A0D0 | HMPREF9441_02136 | 34.816 | 98 | 1.85E-120 | 377 | glycosyl hydrolase family 2, sugar binding domain protein [Paraprevotella clara YIT 11840] |
| Q8A0D0 | HMPREF1475_01699 | 69.835 | 70 | 1.43E-119 | 369 | hypothetical protein [Prevotella oralis HGA0225] |

| | | | | | | |
|---|---|---|---|---|---|---|
| Q8A0D0 | HMPREF1475_01699 | 53.636 | 70 | 2.61E-82 | 272 | hypothetical protein [Prevotella oralis HGA0225] |
| Q8A0D0 | HMPREF9442_01303 | 35.552 | 98 | 4.15E-119 | 373 | glycosyl hydrolase family 2, sugar binding domain protein [Paraprevotella xylaniphila YIT 11841] |
| Q8A0D0 | BSBG_00455 | 34.869 | 97 | 6.15E-119 | 372 | glycosyl hydrolase family 2, sugar binding domain protein [Bacteroides sp. 9_1_42FAA] |
| Q8A0D0 | BACDOR_00544 | 34.869 | 97 | 7.70E-119 | 372 | glycosyl hydrolase family 2, sugar binding domain protein [Bacteroides dorei DSM 17855] |
| Q8A0D0 | HMPREF0105_1607 | 34.869 | 97 | 9.34E-119 | 372 | glycosyl hydrolase family 2, sugar binding domain protein [Bacteroides sp. 3_1_33FAA] |
| Q8A0D0 | BSEG_01381 | 34.869 | 97 | 9.34E-119 | 372 | glycosyl hydrolase family 2, sugar binding domain protein [Bacteroides dorei 5_1_36/D4] |
| Q8A0D0 | CUU_3590 | 34.621 | 96 | 4.79E-118 | 370 | glycosyl hydrolase family 2, sugar binding domain protein [Bacteroides vulgatus PC510] |
| Q8A0D0 | BACINT_04222 | 34.488 | 98 | 2.75E-116 | 365 | glycosyl hydrolase family 2, sugar binding domain protein [Bacteroides intestinalis DSM 17393] |
| Q8A0D0 | BSFG_00657 | 34.621 | 96 | 1.10E-115 | 364 | glycosyl hydrolase family 2, sugar binding domain protein [Bacteroides sp. 4_3_47FAA] |
| Q8A0D0 | HMPREF9011_03033 | 34.621 | 96 | 1.10E-115 | 364 | sialic acid-specific 9-O-acetylesterase [Bacteroides sp. 3_1_40A] |
| Q8A0D0 | BACOVA_03450 | 33.03 | 98 | 9.04E-115 | 362 | glycosyl hydrolase family 2, sugar binding domain protein [Bacteroides ovatus ATCC 8483] |
| Q8A0D0 | HMPREF9447_01475 | 34.796 | 98 | 1.14E-113 | 358 | hypothetical protein [Bacteroides oleiciplenus YIT 12058] |
| Q8A0D0 | HMPREF1017_02790 | 32.879 | 98 | 1.66E-113 | 358 | hypothetical protein [Bacteroides ovatus 3_8_47FAA] |
| Q8A0D0 | HMPREF9455_01549 | 34.656 | 98 | 8.56E-112 | 361 | hypothetical protein [Dysgonomonas gadei ATCC BAA-286] |
| Q8A0D0 | HMPREF9011_01515 | 40.281 | 64 | 9.22E-110 | 341 | sialic acid-specific 9-O-acetylesterase [Bacteroides sp. 3_1_40A] |
| Q8A0D0 | BSGG_3169 | 36.347 | 81 | 2.54E-94 | 305 | hypothetical protein [Bacteroides sp. D2] |
| Q8A0D0 | BACEGG_01316 | 34.12 | 82 | 7.61E-93 | 301 | glycosyl hydrolase family 2, sugar binding domain protein [Bacteroides eggerthii DSM 20697] |
| Q8A0D0 | HMPREF1016_02152 | 33.757 | 82 | 1.84E-91 | 298 | sialate O-acetylesterase [Bacteroides eggerthii 1_2_48FAA] |
| Q8A0D0 | CUY_2421 | 35.443 | 81 | 3.54E-91 | 297 | glycosyl hydrolase family 2, sugar binding domain protein [Bacteroides ovatus SD CMC 3f] |
| Q8A0D0 | HMPREF1017_02107 | 35.443 | 81 | 3.54E-91 | 297 | hypothetical protein [Bacteroides ovatus 3_8_47FAA] |
| Q8A0D0 | HMPREF0106_03415 | 35.443 | 81 | 3.54E-91 | 297 | sialate O-acetylesterase [Bacteroides sp. D22] |
| Q8A0D0 | HMPREF9010_02346 | 35.443 | 81 | 1.79E-90 | 296 | putative sialate O-acetylesterase [Bacteroides sp. 3_1_23] |
| Q8A0D0 | BACOVA_04389 | 35.443 | 81 | 1.79E-90 | 296 | glycosyl hydrolase family 2, sugar binding domain protein [Bacteroides ovatus ATCC 8483] |
| Q8A0D0 | BACCELL_03437 | 32.671 | 81 | 1.67E-88 | 290 | glycosyl hydrolase family 2, sugar binding domain protein [Bacteroides cellulosilyticus DSM 14838] |
| Q8A0D0 | HMPREF0127_04268 | 34.946 | 82 | 1.84E-88 | 290 | hypothetical protein [Bacteroides sp. 1_1_30] |
| Q8A0D0 | BXY_46620 | 34.767 | 82 | 9.54E-88 | 288 | Domain of unknown function (DUF303). [Bacteroides xylanisolvens XB1A] |
| Q8A0D0 | BACPLE_01979 | 33.631 | 82 | 3.60E-87 | 286 | glycosyl hydrolase family 2, sugar binding domain protein [Bacteroides plebeius DSM 17135] |
| Q8A0D0 | BACINT_04200 | 32.194 | 81 | 4.84E-87 | 286 | glycosyl hydrolase family 2, sugar binding domain protein [Bacteroides intestinalis DSM 17393] |
| Q8A0D0 | HMPREF0102_04616 | 34.588 | 82 | 1.15E-86 | 285 | glycosyl hydrolase family 2, sugar binding domain protein [Bacteroides sp. 2_1_22] |
| Q8A0D0 | CW3_4008 | 34.588 | 82 | 1.15E-86 | 285 | glycosyl hydrolase family 2, sugar binding domain protein [Bacteroides xylanisolvens SD CC 1b] |
| Q8A0D0 | CW1_3579 | 34.588 | 82 | 1.15E-86 | 285 | glycosyl hydrolase family 2, sugar binding domain protein [Bacteroides xylanisolvens SD CC 2a] |
| Q8A0D0 | BSCG_00489 | 34.87 | 73 | 6.67E-82 | 271 | glycosyl hydrolase family 2, sugar binding domain protein [Bacteroides sp. 2_2_4] |
| Q8A0D0 | CLOSTHATH_04661 | 29.567 | 95 | 7.88E-82 | 275 | glycosyl hydrolase family 2, sugar binding domain protein [Lachnospiraceae [Clostridium] hathewayi DSM 13479] |
| Q8A0D0 | BIFANG_03777 | 30.735 | 97 | 4.33E-81 | 274 | glycosyl hydrolase family 2, sugar binding domain protein [Bifidobacterium angulatum DSM 20098 = JCM 7096] |
| Q8A0D0 | BIFPSEUDO_02469 | 30.735 | 97 | 5.20E-80 | 271 | glycosyl hydrolase family 2, sugar binding domain protein [Bifidobacterium pseudocatenulatum DSM 20438 = JCM 1200 = LMG 10505] |
| Q8A0D0 | BIFCAT_01346 | 30.339 | 97 | 1.05E-79 | 270 | glycosyl hydrolase family 2, sugar binding domain protein [Bifidobacterium catenulatum DSM 16992 = JCM 1194 = LMG 11043] |
| Q8A0D0 | BIFPSEUDO_02491 | 29.528 | 95 | 2.32E-79 | 268 | glycosyl hydrolase family 2, sugar binding domain protein [Bifidobacterium pseudocatenulatum DSM 20438 = JCM 1200 = LMG 10505] |
| Q8A0D0 | HMPREF1033_00396 | 53.785 | 35 | 2.28E-75 | 256 | hypothetical protein [Tannerella sp. 6_1_58FAA_CT1] |
| Q8A0D0 | BIFADO_00693 | 29.102 | 93 | 9.65E-74 | 254 | glycosyl hydrolase family 2, sugar binding domain protein [Bifidobacterium adolescentis L2-32] |
| Q8A0D0 | BSIG_4347 | 52.174 | 35 | 3.51E-70 | 242 | hypothetical protein [Bacteroides sp. 1_1_6] |
| Q8A0D0 | HMPREF9010_04524 | 52.362 | 35 | 1.60E-69 | 240 | sialate O-acetylesterase [Bacteroides sp. 3_1_23] |
| Q8A0D0 | BSGG_3065 | 52.362 | 35 | 1.63E-69 | 240 | hypothetical protein [Bacteroides sp. D2] |
| Q8A0D0 | HMPREF1017_04523 | 52.362 | 35 | 3.23E-69 | 239 | hypothetical protein [Bacteroides ovatus 3_8_47FAA] |
| Q8A0D0 | HMPREF9447_05050 | 47.234 | 68 | 1.02E-62 | 220 | hypothetical protein [Bacteroides oleiciplenus YIT 12058] |
| Q8A0D0 | HMPREF9447_05050 | 38.326 | 68 | 1.17E-35 | 145 | hypothetical protein [Bacteroides oleiciplenus YIT 12058] |
| Q8A0D0 | BACCELL_05108 | 43.404 | 68 | 3.53E-59 | 210 | hypothetical protein [Bacteroides cellulosilyticus DSM 14838] |
| Q8A0D0 | BACCELL_05108 | 41.364 | 68 | 1.23E-43 | 167 | hypothetical protein [Bacteroides cellulosilyticus DSM 14838] |
| Q8A0D0 | BACINT_02833 | 43.83 | 68 | 1.53E-58 | 208 | hypothetical protein [Bacteroides intestinalis DSM 17393] |
| Q8A0D0 | BACINT_02833 | 41.629 | 68 | 2.58E-43 | 166 | hypothetical protein [Bacteroides intestinalis DSM 17393] |
| Q8A0D0 | HMPREF9447_01873 | 46.667 | 72 | 2.93E-57 | 212 | hypothetical protein [Bacteroides oleiciplenus YIT 12058] |
| Q8A0D0 | HMPREF9447_01873 | 40.486 | 72 | 1.85E-45 | 178 | hypothetical protein [Bacteroides oleiciplenus YIT 12058] |
| Q8A0D0 | BACCELL_01180 | 45.833 | 68 | 9.52E-57 | 211 | glycosyl hydrolase, family 88 [Bacteroides cellulosilyticus DSM 14838] |
| Q8A0D0 | BACCELL_01180 | 41.176 | 68 | 1.81E-42 | 169 | glycosyl hydrolase, family 88 [Bacteroides cellulosilyticus DSM 14838] |
| Q8A0D0 | BACCOP_02540 | 46.154 | 67 | 1.30E-56 | 210 | glycosyl hydrolase, family 88 [Bacteroides coprocola DSM 17136] |
| Q8A0D0 | BACCOP_02540 | 37.443 | 67 | 3.62E-40 | 162 | glycosyl hydrolase, family 88 [Bacteroides coprocola DSM 17136] |
| Q8A0D0 | BACINT_04749 | 44.77 | 67 | 2.82E-56 | 202 | hypothetical protein [Bacteroides intestinalis DSM 17393] |
| Q8A0D0 | BACINT_04749 | 42.523 | 67 | 1.03E-48 | 181 | hypothetical protein [Bacteroides intestinalis DSM 17393] |
| Q8A0D0 | BACCELL_00109 | 45.607 | 68 | 1.12E-55 | 200 | hypothetical protein [Bacteroides cellulosilyticus DSM 14838] |
| Q8A0D0 | BACCELL_00109 | 43.318 | 68 | 3.89E-50 | 185 | hypothetical protein [Bacteroides cellulosilyticus DSM 14838] |
| Q8A0D0 | BACCELL_00130 | 43.673 | 72 | 1.41E-55 | 200 | hypothetical protein [Bacteroides cellulosilyticus DSM 14838] |

| | | | | | | |
|---|---|---|---|---|---|---|
| Q8A0D0 | BACCELL_00130 | 44.681 | 72 | 2.58E-50 | 186 | hypothetical protein [Bacteroides cellulosilyticus DSM 14838] |
| Q8A0D0 | BACEGG_03246 | 46.667 | 36 | 4.23E-55 | 200 | hypothetical protein [Bacteroides eggerthii DSM 20697] |
| Q8A0D0 | HMPREF9447_04446 | 44.77 | 68 | 4.56E-55 | 199 | hypothetical protein [Bacteroides oleiciplenus YIT 12058] |
| Q8A0D0 | HMPREF9447_04446 | 44.24 | 68 | 2.79E-50 | 186 | hypothetical protein [Bacteroides oleiciplenus YIT 12058] |
| Q8A0D0 | HMPREF1033_02018 | 47.059 | 68 | 5.94E-55 | 199 | hypothetical protein [Tannerella sp. 6_1_58FAA_CT1] |
| Q8A0D0 | HMPREF1033_02018 | 44.7 | 68 | 3.20E-49 | 183 | hypothetical protein [Tannerella sp. 6_1_58FAA_CT1] |
| Q8A0D0 | HMPREF9445_01947 | 45.833 | 36 | 7.53E-55 | 199 | hypothetical protein [Bacteroides clarus YIT 12056] |
| Q8A0D0 | BACEGG_02278 | 46.186 | 67 | 2.60E-54 | 196 | hypothetical protein [Bacteroides eggerthii DSM 20697] |
| Q8A0D0 | BACEGG_02278 | 40.909 | 67 | 9.04E-47 | 176 | hypothetical protein [Bacteroides eggerthii DSM 20697] |
| Q8A0D0 | HMPREF1016_00473 | 46.186 | 67 | 2.66E-54 | 196 | hypothetical protein [Bacteroides eggerthii 1_2_48FAA] |
| Q8A0D0 | HMPREF1016_00473 | 40.455 | 67 | 2.71E-46 | 174 | hypothetical protein [Bacteroides eggerthii 1_2_48FAA] |
| Q8A0D0 | BACOVA_05004 | 45.417 | 68 | 2.22E-53 | 201 | glycosyl hydrolase, family 88 [Bacteroides ovatus ATCC 8483] |
| Q8A0D0 | BACOVA_05004 | 37.327 | 68 | 1.71E-37 | 154 | glycosyl hydrolase, family 88 [Bacteroides ovatus ATCC 8483] |
| Q8A0D0 | HMPREF0106_02592 | 45.417 | 68 | 2.75E-53 | 200 | sialic acid-specific 9-O-acetylesterase [Bacteroides sp. D22] |
| Q8A0D0 | HMPREF0106_02592 | 36.866 | 68 | 8.82E-37 | 152 | sialic acid-specific 9-O-acetylesterase [Bacteroides sp. D22] |
| Q8A0D0 | HMPREF1017_02204 | 45.417 | 72 | 3.88E-53 | 200 | hypothetical protein [Bacteroides ovatus 3_8_47FAA] |
| Q8A0D0 | HMPREF1017_02204 | 37.037 | 72 | 3.79E-42 | 168 | hypothetical protein [Bacteroides ovatus 3_8_47FAA] |
| Q8A0D0 | CUY_0705 | 45.417 | 72 | 3.96E-53 | 200 | glycosyl hydrolase, family 88 [Bacteroides ovatus SD CMC 3f] |
| Q8A0D0 | CUY_0705 | 36.885 | 72 | 4.04E-42 | 168 | glycosyl hydrolase, family 88 [Bacteroides ovatus SD CMC 3f] |
| Q8A0D0 | BACFIN_06071 | 45.417 | 68 | 6.51E-53 | 199 | glycosyl hydrolase, family 88 [Bacteroides finegoldii DSM 17565] |
| Q8A0D0 | BACFIN_06071 | 37.327 | 68 | 5.25E-37 | 152 | glycosyl hydrolase, family 88 [Bacteroides finegoldii DSM 17565] |
| Q8A0D0 | BXY_32590 | 45 | 64 | 8.21E-53 | 199 | Predicted unsaturated glucuronyl hydrolase involved in regulation of bacterial surface properties, and related proteins [Bacteroides xylanisolvens XB1A] |
| Q8A0D0 | BXY_32590 | 38.743 | 64 | 2.09E-36 | 150 | Predicted unsaturated glucuronyl hydrolase involved in regulation of bacterial surface properties, and related proteins [Bacteroides xylanisolvens XB1A] |
| Q8A0D0 | HMPREF9010_02773 | 45.417 | 68 | 1.01E-52 | 199 | sialic acid-specific 9-O-acetylesterase [Bacteroides sp. 3_1_23] |
| Q8A0D0 | HMPREF9010_02773 | 37.327 | 68 | 1.71E-37 | 154 | sialic acid-specific 9-O-acetylesterase [Bacteroides sp. 3_1_23] |
| Q8A0D0 | BSCG_01778 | 45.417 | 68 | 1.01E-52 | 199 | glycosyl hydrolase, family 88 [Bacteroides sp. 2_2_4] |
| Q8A0D0 | BSCG_01778 | 37.327 | 68 | 1.71E-37 | 154 | glycosyl hydrolase, family 88 [Bacteroides sp. 2_2_4] |
| Q8A0D0 | HMPREF0127_04827 | 45 | 68 | 1.02E-52 | 199 | hypothetical protein [Bacteroides sp. 1_1_30] |
| Q8A0D0 | HMPREF0127_04827 | 38.073 | 68 | 8.90E-41 | 164 | hypothetical protein [Bacteroides sp. 1_1_30] |
| Q8A0D0 | BSGG_1406 | 37.216 | 86 | 1.64E-52 | 198 | hypothetical protein [Bacteroides sp. D2] |
| Q8A0D0 | BSGG_1406 | 36.475 | 86 | 3.21E-42 | 168 | hypothetical protein [Bacteroides sp. D2] |
| Q8A0D0 | HMPREF9445_01759 | 42.562 | 72 | 2.75E-52 | 197 | glycosyl hydrolase, family 88 [Bacteroides clarus YIT 12056] |
| Q8A0D0 | HMPREF9445_01759 | 39.256 | 72 | 3.10E-45 | 177 | glycosyl hydrolase, family 88 [Bacteroides clarus YIT 12056] |
| Q8A0D0 | BACSTE_00371 | 42.562 | 72 | 2.82E-52 | 197 | glycosyl hydrolase, family 88 [Bacteroides stercoris ATCC 43183] |
| Q8A0D0 | BACSTE_00371 | 38.43 | 72 | 5.92E-45 | 176 | glycosyl hydrolase, family 88 [Bacteroides stercoris ATCC 43183] |
| Q8A0D0 | BACOVA_00163 | 43.636 | 69 | 4.56E-52 | 191 | hypothetical protein [Bacteroides ovatus ATCC 8483] |
| Q8A0D0 | BACOVA_00163 | 40.726 | 69 | 4.98E-44 | 168 | hypothetical protein [Bacteroides ovatus ATCC 8483] |
| Q8A0D0 | HMPREF1532_01373 | 42.692 | 80 | 5.47E-52 | 197 | hypothetical protein [Bacteroides salyersiae WAL 10018 = DSM 18765 = JCM 12988] |
| Q8A0D0 | HMPREF1532_01373 | 34.752 | 80 | 9.74E-44 | 172 | hypothetical protein [Bacteroides salyersiae WAL 10018 = DSM 18765 = JCM 12988] |
| Q8A0D0 | HMPREF9446_03361 | 42.149 | 69 | 1.06E-51 | 196 | glycosyl hydrolase, family 88 [Bacteroides fluxus YIT 12057] |
| Q8A0D0 | HMPREF9446_03361 | 40.265 | 69 | 8.83E-44 | 173 | glycosyl hydrolase, family 88 [Bacteroides fluxus YIT 12057] |
| Q8A0D0 | CUY_4445 | 43.182 | 68 | 1.75E-51 | 189 | conserved hypothetical protein [Bacteroides ovatus SD CMC 3f] |
| Q8A0D0 | CUY_4445 | 41.079 | 68 | 7.17E-43 | 165 | conserved hypothetical protein [Bacteroides ovatus SD CMC 3f] |
| Q8A0D0 | HMPREF9007_00837 | 44.144 | 72 | 1.87E-51 | 189 | sialic acid-specific 9-O-acetylesterase [Bacteroides sp. 1_1_14] |
| Q8A0D0 | HMPREF9007_00837 | 37.367 | 72 | 1.64E-36 | 147 | sialic acid-specific 9-O-acetylesterase [Bacteroides sp. 1_1_14] |
| Q8A0D0 | BXY_04090 | 43.182 | 68 | 2.16E-51 | 189 | Domain of unknown function (DUF303). [Bacteroides xylanisolvens XB1A] |
| Q8A0D0 | BXY_04090 | 41.494 | 68 | 2.10E-43 | 167 | Domain of unknown function (DUF303). [Bacteroides xylanisolvens XB1A] |
| Q8A0D0 | HMPREF1017_03623 | 43.182 | 69 | 2.32E-51 | 189 | hypothetical protein [Bacteroides ovatus 3_8_47FAA] |
| Q8A0D0 | HMPREF1017_03623 | 40.726 | 69 | 7.84E-44 | 168 | hypothetical protein [Bacteroides ovatus 3_8_47FAA] |
| Q8A0D0 | BACCELL_02867 | 42.922 | 64 | 2.55E-51 | 188 | hypothetical protein [Bacteroides cellulosilyticus DSM 14838] |
| Q8A0D0 | BACCELL_02867 | 40.376 | 64 | 3.49E-33 | 137 | hypothetical protein [Bacteroides cellulosilyticus DSM 14838] |
| Q8A0D0 | HMPREF9010_01003 | 43.182 | 69 | 2.62E-51 | 189 | sialic acid-specific 9-O-acetylesterase [Bacteroides sp. 3_1_23] |
| Q8A0D0 | HMPREF9010_01003 | 40.726 | 69 | 1.96E-43 | 167 | sialic acid-specific 9-O-acetylesterase [Bacteroides sp. 3_1_23] |
| Q8A0D0 | BSGG_0067 | 43.182 | 69 | 2.78E-51 | 189 | hypothetical protein [Bacteroides sp. D2] |
| Q8A0D0 | BSGG_0067 | 40.323 | 69 | 9.69E-43 | 165 | hypothetical protein [Bacteroides sp. D2] |
| Q8A0D0 | HMPREF9007_02417 | 42.273 | 69 | 3.07E-51 | 188 | sialic acid-specific 9-O-acetylesterase [Bacteroides sp. 1_1_14] |
| Q8A0D0 | HMPREF9007_02417 | 40.726 | 69 | 6.44E-44 | 168 | sialic acid-specific 9-O-acetylesterase [Bacteroides sp. 1_1_14] |
| Q8A0D0 | HMPREF0106_01109 | 44.144 | 72 | 4.92E-51 | 188 | sialic acid-specific 9-O-acetylesterase [Bacteroides sp. D22] |

| | | | | | | |
|---|---|---|---|---|---|---|
| Q8A0D0 | HMPREF0106_01109 | 37.367 | 72 | 1.77E-36 | 147 | sialic acid-specific 9-O-acetylesterase [Bacteroides sp. D22] |
| Q8A0D0 | BACFIN_06776 | 42.922 | 69 | 9.70E-51 | 187 | hypothetical protein [Bacteroides finegoldii DSM 17565] |
| Q8A0D0 | BACFIN_06776 | 40.323 | 69 | 1.37E-42 | 164 | hypothetical protein [Bacteroides finegoldii DSM 17565] |
| Q8A0D0 | BACINT_04752 | 44.351 | 67 | 1.06E-50 | 187 | hypothetical protein [Bacteroides intestinalis DSM 17393] |
| Q8A0D0 | BACINT_04752 | 39.286 | 67 | 1.82E-41 | 161 | hypothetical protein [Bacteroides intestinalis DSM 17393] |
| Q8A0D0 | CW1_3731 | 42.727 | 69 | 1.45E-50 | 186 | conserved hypothetical protein [Bacteroides xylanisolvens SD CC 2a] |
| Q8A0D0 | CW1_3731 | 40.726 | 69 | 1.27E-43 | 167 | conserved hypothetical protein [Bacteroides xylanisolvens SD CC 2a] |
| Q8A0D0 | HMPREF0102_03014 | 42.727 | 69 | 1.77E-50 | 186 | hypothetical protein [Bacteroides sp. 2_1_22] |
| Q8A0D0 | HMPREF0102_03014 | 40.726 | 69 | 1.11E-43 | 167 | hypothetical protein [Bacteroides sp. 2_1_22] |
| Q8A0D0 | BSAG_01602 | 42.727 | 69 | 1.77E-50 | 186 | hypothetical protein [Bacteroides sp. D1] |
| Q8A0D0 | BSAG_01602 | 40.726 | 69 | 1.11E-43 | 167 | hypothetical protein [Bacteroides sp. D1] |
| Q8A0D0 | HMPREF1016_00623 | 41.224 | 68 | 2.59E-50 | 186 | hypothetical protein [Bacteroides eggerthii 1_2_48FAA] |
| Q8A0D0 | HMPREF1016_00623 | 38.14 | 68 | 2.22E-37 | 149 | hypothetical protein [Bacteroides eggerthii 1_2_48FAA] |
| Q8A0D0 | BSIG_0631 | 42.273 | 69 | 2.99E-50 | 186 | hypothetical protein [Bacteroides sp. 1_1_6] |
| Q8A0D0 | BSIG_0631 | 41.129 | 69 | 4.42E-44 | 169 | hypothetical protein [Bacteroides sp. 1_1_6] |
| Q8A0D0 | HMPREF0127_00342 | 42.727 | 69 | 3.45E-50 | 186 | hypothetical protein [Bacteroides sp. 1_1_30] |
| Q8A0D0 | HMPREF0127_00342 | 40.726 | 69 | 1.85E-43 | 167 | hypothetical protein [Bacteroides sp. 1_1_30] |
| Q8A0D0 | HMPREF0106_01711 | 42.727 | 69 | 3.45E-50 | 186 | sialic acid-specific 9-O-acetylesterase [Bacteroides sp. D22] |
| Q8A0D0 | HMPREF0106_01711 | 40.726 | 69 | 1.85E-43 | 167 | sialic acid-specific 9-O-acetylesterase [Bacteroides sp. D22] |
| Q8A0D0 | BACCELL_00129 | 45.116 | 32 | 4.58E-50 | 185 | hypothetical protein [Bacteroides cellulosilyticus DSM 14838] |
| Q8A0D0 | BSCG_02286 | 42.273 | 69 | 8.87E-50 | 184 | hypothetical protein [Bacteroides sp. 2_2_4] |
| Q8A0D0 | BSCG_02286 | 40.726 | 69 | 1.64E-43 | 167 | hypothetical protein [Bacteroides sp. 2_2_4] |
| Q8A0D0 | BACEGG_02266 | 42.857 | 64 | 1.44E-49 | 183 | hypothetical protein [Bacteroides eggerthii DSM 20697] |
| Q8A0D0 | BACEGG_02266 | 38.647 | 64 | 1.04E-35 | 144 | hypothetical protein [Bacteroides eggerthii DSM 20697] |
| Q8A0D0 | HMPREF9447_04495 | 40.816 | 68 | 1.58E-49 | 184 | hypothetical protein [Bacteroides oleiciplenus YIT 12058] |
| Q8A0D0 | HMPREF9447_04495 | 40 | 68 | 4.99E-37 | 149 | hypothetical protein [Bacteroides oleiciplenus YIT 12058] |
| Q8A0D0 | BACEGG_00642 | 41.532 | 73 | 2.02E-49 | 183 | hypothetical protein [Bacteroides eggerthii DSM 20697] |
| Q8A0D0 | BACEGG_00642 | 41.935 | 73 | 6.35E-44 | 168 | hypothetical protein [Bacteroides eggerthii DSM 20697] |
| Q8A0D0 | HMPREF1016_00633 | 41.762 | 70 | 2.17E-49 | 183 | hypothetical protein [Bacteroides eggerthii 1_2_48FAA] |
| Q8A0D0 | HMPREF1016_00633 | 37.668 | 70 | 4.06E-36 | 146 | hypothetical protein [Bacteroides eggerthii 1_2_48FAA] |
| Q8A0D0 | HMPREF9447_04506 | 42.146 | 70 | 2.28E-49 | 183 | hypothetical protein [Bacteroides oleiciplenus YIT 12058] |
| Q8A0D0 | HMPREF9447_04506 | 38.117 | 70 | 2.16E-36 | 147 | hypothetical protein [Bacteroides oleiciplenus YIT 12058] |
| Q8A0D0 | HMPREF1214_01054 | 43.333 | 72 | 2.50E-49 | 189 | unsaturated rhamnogalacturonyl hydrolase [Bacteroides sp. HPS0048] |
| Q8A0D0 | HMPREF1214_01054 | 37.6 | 72 | 1.43E-43 | 172 | unsaturated rhamnogalacturonyl hydrolase [Bacteroides sp. HPS0048] |
| Q8A0D0 | HMPREF1016_01101 | 42.387 | 72 | 2.51E-49 | 183 | hypothetical protein [Bacteroides eggerthii 1_2_48FAA] |
| Q8A0D0 | HMPREF1016_01101 | 42.339 | 72 | 2.86E-44 | 169 | hypothetical protein [Bacteroides eggerthii 1_2_48FAA] |
| Q8A0D0 | HMPREF1016_00462 | 42.857 | 68 | 2.84E-49 | 183 | sialate O-acetylesterase [Bacteroides eggerthii 1_2_48FAA] |
| Q8A0D0 | HMPREF1016_00462 | 38.136 | 68 | 1.14E-40 | 159 | sialate O-acetylesterase [Bacteroides eggerthii 1_2_48FAA] |
| Q8A0D0 | HMPREF1532_01000 | 42.723 | 68 | 3.28E-49 | 182 | hypothetical protein [Bacteroides salyersiae WAL 10018 = DSM 18765 = JCM 12988] |
| Q8A0D0 | HMPREF1532_01000 | 41.224 | 68 | 1.72E-43 | 167 | hypothetical protein [Bacteroides salyersiae WAL 10018 = DSM 18765 = JCM 12988] |
| Q8A0D0 | BSEG_00470 | 41.333 | 70 | 7.99E-49 | 182 | hypothetical protein [Bacteroides dorei 5_1_36/D4] |
| Q8A0D0 | BSEG_00470 | 36.992 | 70 | 1.58E-37 | 150 | hypothetical protein [Bacteroides dorei 5_1_36/D4] |
| Q8A0D0 | BACDOR_04226 | 41.333 | 70 | 7.99E-49 | 182 | hypothetical protein [Bacteroides dorei DSM 17855] |
| Q8A0D0 | BACDOR_04226 | 36.992 | 70 | 1.58E-37 | 150 | hypothetical protein [Bacteroides dorei DSM 17855] |
| Q8A0D0 | BSBG_01569 | 41.333 | 70 | 7.99E-49 | 182 | hypothetical protein [Bacteroides sp. 9_1_42FAA] |
| Q8A0D0 | BSBG_01569 | 36.992 | 70 | 1.58E-37 | 150 | hypothetical protein [Bacteroides sp. 9_1_42FAA] |
| Q8A0D0 | HMPREF0105_0351 | 41.333 | 70 | 8.66E-49 | 182 | hypothetical protein [Bacteroides sp. 3_1_33FAA] |
| Q8A0D0 | HMPREF0105_0351 | 36.992 | 70 | 1.57E-37 | 150 | hypothetical protein [Bacteroides sp. 3_1_33FAA] |
| Q8A0D0 | BACCAC_02404 | 40.496 | 68 | 1.87E-48 | 181 | hypothetical protein [Bacteroides caccae ATCC 43185] |
| Q8A0D0 | BACCAC_02404 | 42.857 | 68 | 2.68E-44 | 169 | hypothetical protein [Bacteroides caccae ATCC 43185] |
| Q8A0D0 | BSGG_1789 | 43.636 | 71 | 3.73E-48 | 180 | hypothetical protein [Bacteroides sp. D2] |
| Q8A0D0 | BSGG_1789 | 35.556 | 71 | 1.66E-34 | 141 | hypothetical protein [Bacteroides sp. D2] |
| Q8A0D0 | BSFG_04267 | 40.444 | 70 | 9.27E-48 | 179 | hypothetical protein [Bacteroides sp. 4_3_47FAA] |
| Q8A0D0 | BSFG_04267 | 36.992 | 70 | 1.88E-37 | 150 | hypothetical protein [Bacteroides sp. 4_3_47FAA] |
| Q8A0D0 | HMPREF9011_02985 | 40.444 | 70 | 9.27E-48 | 179 | sialic acid-specific acetylesterase II [Bacteroides sp. 3_1_40A] |
| Q8A0D0 | HMPREF9011_02985 | 36.992 | 70 | 1.88E-37 | 150 | sialic acid-specific acetylesterase II [Bacteroides sp. 3_1_40A] |
| Q8A0D0 | BACCELL_00106 | 44.118 | 66 | 1.04E-47 | 179 | hypothetical protein [Bacteroides cellulosilyticus DSM 14838] |
| Q8A0D0 | BACCELL_00106 | 38.914 | 66 | 2.64E-38 | 152 | hypothetical protein [Bacteroides cellulosilyticus DSM 14838] |
| Q8A0D0 | CUU_2736 | 40.444 | 70 | 1.06E-47 | 179 | conserved hypothetical protein [Bacteroides vulgatus PC510] |
| Q8A0D0 | CUU_2736 | 37.398 | 70 | 4.96E-38 | 151 | conserved hypothetical protein [Bacteroides vulgatus PC510] |

| | | | | | | |
|---|---|---|---|---|---|---|
| Q8A0D0 | HMPREF9447_04460 | 42.857 | 68 | 1.70E-47 | 178 | hypothetical protein [Bacteroides oleiciplenus YIT 12058] |
| Q8A0D0 | HMPREF9447_04460 | 40.171 | 68 | 1.21E-43 | 167 | hypothetical protein [Bacteroides oleiciplenus YIT 12058] |
| Q8A0D0 | BACOVA_02748 | 40.31 | 71 | 2.54E-47 | 177 | hypothetical protein [Bacteroides ovatus ATCC 8483] |
| Q8A0D0 | BACOVA_02748 | 39.556 | 71 | 4.36E-39 | 154 | hypothetical protein [Bacteroides ovatus ATCC 8483] |
| Q8A0D0 | BSGG_0510 | 42.222 | 69 | 6.50E-47 | 176 | hypothetical protein [Bacteroides sp. D2] |
| Q8A0D0 | BSGG_0510 | 38.43 | 69 | 3.93E-37 | 149 | hypothetical protein [Bacteroides sp. D2] |
| Q8A0D0 | HMPREF0969_01697 | 41.353 | 71 | 1.87E-46 | 175 | hypothetical protein [Bacteroides sp. D20] |
| Q8A0D0 | HMPREF0969_01697 | 37.22 | 71 | 9.83E-42 | 162 | hypothetical protein [Bacteroides sp. D20] |
| Q8A0D0 | BACUNI_04187 | 41.353 | 71 | 1.87E-46 | 175 | hypothetical protein [Bacteroides uniformis ATCC 8492] |
| Q8A0D0 | BACUNI_04187 | 37.22 | 71 | 9.83E-42 | 162 | hypothetical protein [Bacteroides uniformis ATCC 8492] |
| Q8A0D0 | PREVCOP_06527 | 42.009 | 69 | 1.90E-46 | 175 | hypothetical protein [Prevotella copri DSM 18205] |
| Q8A0D0 | PREVCOP_06527 | 34.51 | 69 | 4.63E-32 | 134 | hypothetical protein [Prevotella copri DSM 18205] |
| Q8A0D0 | HMPREF9449_01521 | 41.102 | 35 | 2.03E-46 | 179 | hypothetical protein [Odoribacter laneus YIT 12061] |
| Q8A0D0 | BSGG_3976 | 41.667 | 68 | 2.11E-46 | 175 | hypothetical protein [Bacteroides sp. D2] |
| Q8A0D0 | BSGG_3976 | 39.749 | 68 | 3.88E-37 | 149 | hypothetical protein [Bacteroides sp. D2] |
| Q8A0D0 | HMPREF9007_01383 | 41.333 | 69 | 2.55E-46 | 175 | sialate O-acetylesterase [Bacteroides sp. 1_1_14] |
| Q8A0D0 | HMPREF9007_01383 | 37.449 | 69 | 1.68E-36 | 147 | sialate O-acetylesterase [Bacteroides sp. 1_1_14] |
| Q8A0D0 | BSIG_0282 | 41.333 | 69 | 2.55E-46 | 175 | hypothetical protein [Bacteroides sp. 1_1_6] |
| Q8A0D0 | BSIG_0282 | 37.449 | 69 | 1.68E-36 | 147 | hypothetical protein [Bacteroides sp. 1_1_6] |
| Q8A0D0 | BACINT_01446 | 44.444 | 36 | 3.82E-46 | 179 | hypothetical protein [Bacteroides intestinalis DSM 17393] |
| Q8A0D0 | BACEGG_03295 | 44.444 | 37 | 3.82E-46 | 179 | hypothetical protein [Bacteroides eggerthii DSM 20697] |
| Q8A0D0 | HMPREF0105_3561 | 43.534 | 66 | 3.95E-46 | 174 | hypothetical protein [Bacteroides sp. 3_1_33FAA] |
| Q8A0D0 | HMPREF0105_3561 | 40.845 | 66 | 2.01E-39 | 155 | hypothetical protein [Bacteroides sp. 3_1_33FAA] |
| Q8A0D0 | BSEG_03318 | 43.534 | 66 | 3.95E-46 | 174 | hypothetical protein [Bacteroides dorei 5_1_36/D4] |
| Q8A0D0 | BSEG_03318 | 40.845 | 66 | 2.01E-39 | 155 | hypothetical protein [Bacteroides dorei 5_1_36/D4] |
| Q8A0D0 | HMPREF1016_00957 | 44.444 | 37 | 6.09E-46 | 178 | hypothetical protein [Bacteroides eggerthii 1_2_48FAA] |
| Q8A0D0 | HMPREF0102_01915 | 41.833 | 37 | 6.90E-46 | 169 | hypothetical protein [Bacteroides sp. 2_1_22] |
| Q8A0D0 | CW3_3165 | 41.833 | 37 | 7.10E-46 | 169 | conserved hypothetical protein [Bacteroides xylanisolvens SD CC 1b] |
| Q8A0D0 | CW1_1877 | 41.833 | 37 | 7.10E-46 | 169 | conserved hypothetical protein [Bacteroides xylanisolvens SD CC 2a] |
| Q8A0D0 | BACPLE_03768 | 40 | 69 | 8.00E-46 | 173 | hypothetical protein [Bacteroides plebeius DSM 17135] |
| Q8A0D0 | BACPLE_03768 | 37.22 | 69 | 8.55E-41 | 159 | hypothetical protein [Bacteroides plebeius DSM 17135] |
| Q8A0D0 | HMPREF1007_00492 | 40.977 | 71 | 9.11E-46 | 173 | hypothetical protein [Bacteroides sp. 4_1_36] |
| Q8A0D0 | HMPREF1007_00492 | 37.22 | 71 | 4.07E-41 | 160 | hypothetical protein [Bacteroides sp. 4_1_36] |
| Q8A0D0 | BSIG_1527 | 42.562 | 69 | 1.19E-45 | 173 | hypothetical protein [Bacteroides sp. 1_1_6] |
| Q8A0D0 | BSIG_1527 | 37.118 | 69 | 6.13E-40 | 157 | hypothetical protein [Bacteroides sp. 1_1_6] |
| Q8A0D0 | HMPREF9447_02697 | 44.033 | 36 | 1.33E-45 | 177 | hypothetical protein [Bacteroides oleiciplenus YIT 12058] |
| Q8A0D0 | HMPREF9445_01896 | 44.048 | 37 | 1.52E-45 | 177 | hypothetical protein [Bacteroides clarus YIT 12056] |
| Q8A0D0 | HMPREF0106_00213 | 40.074 | 40 | 1.82E-45 | 175 | sialic acid-specific 9-O-acetylesterase [Bacteroides sp. D22] |
| Q8A0D0 | HMPREF9007_02491 | 42.562 | 66 | 2.65E-45 | 172 | sialic acid-specific 9-O-acetylesterase [Bacteroides sp. 1_1_14] |
| Q8A0D0 | HMPREF9007_02491 | 38.679 | 66 | 1.47E-38 | 153 | sialic acid-specific 9-O-acetylesterase [Bacteroides sp. 1_1_14] |
| Q8A0D0 | AL1_23340 | 42.017 | 68 | 4.41E-45 | 171 | Domain of unknown function (DUF303). [Alistipes shahii WAL 8301] |
| Q8A0D0 | AL1_23340 | 41.935 | 68 | 1.82E-44 | 169 | Domain of unknown function (DUF303). [Alistipes shahii WAL 8301] |
| Q8A0D0 | HMPREF0127_01122 | 41.833 | 37 | 9.64E-45 | 171 | hypothetical protein [Bacteroides sp. 1_1_30] |
| Q8A0D0 | HMPREF9447_03115 | 40.991 | 69 | 1.11E-44 | 170 | hypothetical protein [Bacteroides oleiciplenus YIT 12058] |
| Q8A0D0 | HMPREF9447_03115 | 38.912 | 69 | 2.57E-38 | 152 | hypothetical protein [Bacteroides oleiciplenus YIT 12058] |
| Q8A0D0 | HMPREF0127_03617 | 42.927 | 66 | 1.55E-44 | 174 | hypothetical protein [Bacteroides sp. 1_1_30] |
| Q8A0D0 | HMPREF0127_03617 | 42.797 | 66 | 1.06E-39 | 160 | hypothetical protein [Bacteroides sp. 1_1_30] |
| Q8A0D0 | BSCG_05726 | 41.176 | 70 | 2.02E-44 | 174 | hypothetical protein [Bacteroides sp. 2_2_4] |
| Q8A0D0 | BSCG_05726 | 41.245 | 70 | 3.57E-39 | 159 | hypothetical protein [Bacteroides sp. 2_2_4] |
| Q8A0D0 | CW3_4787 | 43.299 | 65 | 3.65E-44 | 168 | conserved hypothetical protein [Bacteroides xylanisolvens SD CC 1b] |
| Q8A0D0 | CW3_4787 | 40.726 | 65 | 5.65E-44 | 167 | conserved hypothetical protein [Bacteroides xylanisolvens SD CC 1b] |
| Q8A0D0 | HMPREF0102_02680 | 42.439 | 66 | 4.98E-44 | 173 | hypothetical protein [Bacteroides sp. 2_1_22] |
| Q8A0D0 | HMPREF0102_02680 | 42.797 | 66 | 1.40E-39 | 160 | hypothetical protein [Bacteroides sp. 2_1_22] |
| Q8A0D0 | BSAG_01255 | 42.439 | 66 | 4.98E-44 | 173 | hypothetical protein [Bacteroides sp. D1] |
| Q8A0D0 | BSAG_01255 | 42.797 | 66 | 1.40E-39 | 160 | hypothetical protein [Bacteroides sp. D1] |
| Q8A0D0 | CW3_3765 | 42.439 | 66 | 4.98E-44 | 173 | conserved domain protein [Bacteroides xylanisolvens SD CC 1b] |
| Q8A0D0 | CW3_3765 | 42.797 | 66 | 1.40E-39 | 160 | conserved domain protein [Bacteroides xylanisolvens SD CC 1b] |
| Q8A0D0 | CW1_2830 | 42.439 | 66 | 4.98E-44 | 173 | conserved domain protein [Bacteroides xylanisolvens SD CC 2a] |
| Q8A0D0 | CW1_2830 | 42.797 | 66 | 1.40E-39 | 160 | conserved domain protein [Bacteroides xylanisolvens SD CC 2a] |
| Q8A0D0 | BACOVA_01802 | 43.636 | 69 | 1.10E-43 | 173 | hypothetical protein [Bacteroides ovatus ATCC 8483] |

| | | | | | | |
|---|---|---|---|---|---|---|
| Q8A0D0 | BACOVA_01802 | 39.834 | 69 | 1.88E-38 | 157 | hypothetical protein [Bacteroides ovatus ATCC 8483] |
| Q8A0D0 | CUU_3079 | 41.176 | 35 | 1.95E-43 | 170 | GDSL-like protein [Bacteroides vulgatus PC510] |
| Q8A0D0 | HMPREF9007_03720 | 42.152 | 33 | 2.51E-43 | 166 | sialic acid-specific 9-O-acetylesterase [Bacteroides sp. 1_1_14] |
| Q8A0D0 | HMPREF0619_00082 | 38.645 | 37 | 2.83E-43 | 170 | GDSL-like protein [Parabacteroides sp. D13] |
| Q8A0D0 | BSIG_1497 | 42.152 | 33 | 3.12E-43 | 166 | hypothetical protein [Bacteroides sp. 1_1_6] |
| Q8A0D0 | BFAG_01068 | 39.241 | 35 | 3.91E-43 | 169 | GDSL-like protein [Bacteroides fragilis 3_1_12] |
| Q8A0D0 | BACCOP_00150 | 39.592 | 69 | 5.50E-43 | 166 | hypothetical protein [Bacteroides coprocola DSM 17136] |
| Q8A0D0 | BACCOP_00150 | 38.117 | 69 | 1.57E-40 | 159 | hypothetical protein [Bacteroides coprocola DSM 17136] |
| Q8A0D0 | ES1_08140 | 40.506 | 35 | 5.82E-43 | 166 | Domain of unknown function (DUF303). [Ruminiclostridium [Eubacterium] siraeum V10Sc8a] |
| Q8A0D0 | BXY_42630 | 43.182 | 69 | 5.92E-43 | 171 | Uncharacterized protein conserved in bacteria [Bacteroides xylanisolvens XB1A] |
| Q8A0D0 | BXY_42630 | 39.834 | 69 | 3.19E-39 | 159 | Uncharacterized protein conserved in bacteria [Bacteroides xylanisolvens XB1A] |
| Q8A0D0 | CUY_4423 | 43.182 | 69 | 6.11E-43 | 171 | conserved domain protein [Bacteroides ovatus SD CMC 3f] |
| Q8A0D0 | CUY_4423 | 39.419 | 69 | 1.88E-38 | 157 | conserved domain protein [Bacteroides ovatus SD CMC 3f] |
| Q8A0D0 | HMPREF9010_04843 | 43.182 | 69 | 6.55E-43 | 171 | sialic acid-specific 9-O-acetylesterase [Bacteroides sp. 3_1_23] |
| Q8A0D0 | HMPREF9010_04843 | 39.419 | 69 | 1.91E-38 | 157 | sialic acid-specific 9-O-acetylesterase [Bacteroides sp. 3_1_23] |
| Q8A0D0 | BACOVA_01808 | 40.724 | 70 | 6.85E-43 | 170 | hypothetical protein [Bacteroides ovatus ATCC 8483] |
| Q8A0D0 | BACOVA_01808 | 41.634 | 70 | 3.70E-40 | 162 | hypothetical protein [Bacteroides ovatus ATCC 8483] |
| Q8A0D0 | HMPREF9008_04226 | 38.247 | 37 | 1.01E-42 | 168 | GDSL-like protein [Parabacteroides sp. 20_3] |
| Q8A0D0 | HMPREF0127_03621 | 43.182 | 69 | 1.03E-42 | 170 | hypothetical protein [Bacteroides sp. 1_1_30] |
| Q8A0D0 | HMPREF0127_03621 | 39.419 | 69 | 1.90E-38 | 157 | hypothetical protein [Bacteroides sp. 1_1_30] |
| Q8A0D0 | BSEG_01307 | 39.844 | 37 | 1.11E-42 | 168 | GDSL-like protein [Bacteroides dorei 5_1_36/D4] |
| Q8A0D0 | EUBSIR_02263 | 40.928 | 35 | 1.19E-42 | 165 | hypothetical protein [Ruminiclostridium [Eubacterium] siraeum DSM 15702] |
| Q8A0D0 | HMPREF0105_1543 | 39.844 | 37 | 1.26E-42 | 168 | GDSL-like protein [Bacteroides sp. 3_1_33FAA] |
| Q8A0D0 | BSBG_00384 | 39.844 | 37 | 1.26E-42 | 168 | GDSL-like protein [Bacteroides sp. 9_1_42FAA] |
| Q8A0D0 | BSGG_2603 | 38.528 | 66 | 1.66E-42 | 164 | hypothetical protein [Bacteroides sp. D2] |
| Q8A0D0 | BSGG_2603 | 35.981 | 66 | 4.28E-38 | 152 | hypothetical protein [Bacteroides sp. D2] |
| Q8A0D0 | BSFG_00731 | 40.756 | 35 | 1.95E-42 | 167 | GDSL-like protein [Bacteroides sp. 4_3_47FAA] |
| Q8A0D0 | HMPREF9011_03104 | 40.756 | 35 | 1.95E-42 | 167 | sialate O-acetylesterase [Bacteroides sp. 3_1_40A] |
| Q8A0D0 | EUS_22520 | 40.506 | 35 | 1.96E-42 | 165 | Domain of unknown function (DUF303). [Ruminiclostridium [Eubacterium] siraeum 70/3] |
| Q8A0D0 | HMPREF0103_2214 | 37.849 | 37 | 2.61E-42 | 167 | GDSL-like protein [Bacteroides sp. 2_1_33B] |
| Q8A0D0 | HMPREF0104_00088 | 38.247 | 37 | 2.85E-42 | 167 | sialate O-acetylesterase [Bacteroides sp. 3_1_19] |
| Q8A0D0 | BSIG_0266 | 41.322 | 70 | 3.86E-42 | 168 | hypothetical protein [Bacteroides sp. 1_1_6] |
| Q8A0D0 | BSIG_0266 | 40.833 | 70 | 6.21E-40 | 161 | hypothetical protein [Bacteroides sp. 1_1_6] |
| Q8A0D0 | BACDOR_00666 | 39.453 | 37 | 6.04E-42 | 166 | GDSL-like protein [Bacteroides dorei DSM 17855] |
| Q8A0D0 | BSHG_4430 | 38.819 | 35 | 9.95E-42 | 166 | hypothetical protein [Bacteroides sp. 3_2_5] |
| Q8A0D0 | HMPREF1018_01836 | 38.819 | 35 | 9.95E-42 | 166 | hypothetical protein [Bacteroides sp. 2_1_56FAA] |
| Q8A0D0 | HMPREF0101_00674 | 38.819 | 35 | 1.06E-41 | 165 | GDSL-like protein [Bacteroides sp. 2_1_16] |
| Q8A0D0 | HMPREF9007_01365 | 41.202 | 69 | 1.84E-41 | 166 | sialate O-acetylesterase [Bacteroides sp. 1_1_14] |
| Q8A0D0 | HMPREF9007_01365 | 41.25 | 69 | 2.41E-40 | 162 | sialate O-acetylesterase [Bacteroides sp. 1_1_14] |
| Q8A0D0 | PRABACTJOHN_00552 | 37.549 | 37 | 3.53E-41 | 166 | cyclically-permuted mutarotase family protein [Parabacteroides johnsonii DSM 18315] |
| Q8A0D0 | HMPREF0619_04055 | 35.461 | 74 | 9.92E-41 | 159 | hypothetical protein [Parabacteroides sp. D13] |
| Q8A0D0 | HMPREF0619_04055 | 35.747 | 74 | 6.78E-36 | 145 | hypothetical protein [Parabacteroides sp. D13] |
| Q8A0D0 | HMPREF9011_01628 | 35.945 | 33 | 1.47E-40 | 151 | hypothetical protein [Bacteroides sp. 3_1_40A] |
| Q8A0D0 | BACPLE_00585 | 38.462 | 35 | 2.06E-40 | 162 | GDSL-like protein [Bacteroides plebeius DSM 17135] |
| Q8A0D0 | HMPREF0104_03885 | 35.461 | 74 | 2.08E-40 | 158 | sialic acid-specific 9-O-acetylesterase [Bacteroides sp. 3_1_19] |
| Q8A0D0 | HMPREF0104_03885 | 36.199 | 74 | 4.58E-36 | 145 | sialic acid-specific 9-O-acetylesterase [Bacteroides sp. 3_1_19] |
| Q8A0D0 | HMPREF0103_3892 | 35.816 | 74 | 2.58E-40 | 158 | hypothetical protein [Bacteroides sp. 2_1_33B] |
| Q8A0D0 | HMPREF0103_3892 | 35.747 | 74 | 6.28E-36 | 145 | hypothetical protein [Bacteroides sp. 2_1_33B] |
| Q8A0D0 | HMPREF1016_00121 | 37.091 | 40 | 2.79E-40 | 158 | hypothetical protein [Bacteroides eggerthii 1_2_48FAA] |
| Q8A0D0 | BACEGG_01992 | 37.091 | 40 | 2.79E-40 | 158 | hypothetical protein [Bacteroides eggerthii DSM 20697] |
| Q8A0D0 | PARMER_00328 | 37.154 | 37 | 2.82E-40 | 163 | cyclically-permuted mutarotase family protein [Parabacteroides merdae ATCC 43184] |
| Q8A0D0 | HMPREF9420_0396 | 37.615 | 32 | 4.42E-40 | 157 | hypothetical protein [Prevotella salivae DSM 15606] |
| Q8A0D0 | HMPREF1532_00169 | 39.662 | 35 | 5.25E-40 | 160 | hypothetical protein [Bacteroides salyersiae WAL 10018 = DSM 18765 = JCM 12988] |
| Q8A0D0 | BSGG_2455 | 42.678 | 69 | 7.06E-40 | 157 | hypothetical protein [Bacteroides sp. D2] |
| Q8A0D0 | BSGG_2455 | 39.381 | 69 | 7.37E-38 | 151 | hypothetical protein [Bacteroides sp. D2] |
| Q8A0D0 | HMPREF9008_04368 | 35.461 | 74 | 2.04E-39 | 155 | hypothetical protein [Parabacteroides sp. 20_3] |
| Q8A0D0 | HMPREF9008_04368 | 35.747 | 74 | 5.49E-36 | 145 | hypothetical protein [Parabacteroides sp. 20_3] |
| Q8A0D0 | HMPREF0673_02450 | 35.017 | 42 | 2.66E-39 | 159 | GDSL-like protein [Prevotella stercorea DSM 18206] |
| Q8A0D0 | BXY_22860 | 38.621 | 72 | 3.07E-39 | 155 | Domain of unknown function (DUF303). [Bacteroides xylanisolvens XB1A] |
| Q8A0D0 | BXY_22860 | 41.262 | 72 | 1.51E-37 | 150 | Domain of unknown function (DUF303). [Bacteroides xylanisolvens XB1A] |

| | | | | | | |
|---|---|---|---|---|---|---|
| Q8A0D0 | BACOVA_02858 | 38.966 | 72 | 3.73E-39 | 155 | hypothetical protein [Bacteroides ovatus ATCC 8483] |
| Q8A0D0 | BACOVA_02858 | 41.262 | 72 | 5.68E-38 | 151 | hypothetical protein [Bacteroides ovatus ATCC 8483] |
| Q8A0D0 | HMPREF0127_01481 | 42.857 | 30 | 5.08E-39 | 154 | hypothetical protein [Bacteroides sp. 1_1_30] |
| Q8A0D0 | CUY_1625 | 39.013 | 33 | 7.46E-39 | 153 | conserved hypothetical protein [Bacteroides ovatus SD CMC 3f] |
| Q8A0D0 | HMPREF9010_00552 | 38.621 | 72 | 9.28E-39 | 154 | sialic acid-specific 9-O-acetylesterase [Bacteroides sp. 3_1_23] |
| Q8A0D0 | HMPREF9010_00552 | 40.777 | 72 | 1.49E-37 | 150 | sialic acid-specific 9-O-acetylesterase [Bacteroides sp. 3_1_23] |
| Q8A0D0 | BSIG_3572 | 36.818 | 33 | 1.19E-38 | 153 | hypothetical protein [Bacteroides sp. 1_1_6] |
| Q8A0D0 | HMPREF1017_04633 | 39.419 | 36 | 1.72E-38 | 156 | hypothetical protein [Bacteroides ovatus 3_8_47FAA] |
| Q8A0D0 | HMPREF1016_01035 | 38.819 | 35 | 1.82E-38 | 150 | hypothetical protein [Bacteroides eggerthii 1_2_48FAA] |
| Q8A0D0 | HMPREF0102_00710 | 38.621 | 72 | 1.88E-38 | 152 | hypothetical protein [Bacteroides sp. 2_1_22] |
| Q8A0D0 | HMPREF0102_00710 | 41.262 | 72 | 4.91E-38 | 151 | hypothetical protein [Bacteroides sp. 2_1_22] |
| Q8A0D0 | CW3_1978 | 38.621 | 72 | 1.88E-38 | 152 | conserved hypothetical protein [Bacteroides xylanisolvens SD CC 1b] |
| Q8A0D0 | CW3_1978 | 41.262 | 72 | 4.91E-38 | 151 | conserved hypothetical protein [Bacteroides xylanisolvens SD CC 1b] |
| Q8A0D0 | CW1_2507 | 38.621 | 72 | 1.88E-38 | 152 | conserved hypothetical protein [Bacteroides xylanisolvens SD CC 2a] |
| Q8A0D0 | CW1_2507 | 41.262 | 72 | 4.91E-38 | 151 | conserved hypothetical protein [Bacteroides xylanisolvens SD CC 2a] |
| Q8A0D0 | HMPREF1017_00424 | 38.621 | 72 | 2.60E-38 | 152 | hypothetical protein [Bacteroides ovatus 3_8_47FAA] |
| Q8A0D0 | HMPREF1017_00424 | 41.262 | 72 | 6.02E-38 | 152 | hypothetical protein [Bacteroides ovatus 3_8_47FAA] |
| Q8A0D0 | BSAG_00461 | 38.621 | 72 | 2.60E-38 | 152 | hypothetical protein [Bacteroides sp. D1] |
| Q8A0D0 | BSAG_00461 | 41.262 | 72 | 6.02E-38 | 152 | hypothetical protein [Bacteroides sp. D1] |
| Q8A0D0 | BSCG_01640 | 38.621 | 72 | 2.86E-38 | 152 | hypothetical protein [Bacteroides sp. 2_2_4] |
| Q8A0D0 | BSCG_01640 | 40.777 | 72 | 2.51E-37 | 149 | hypothetical protein [Bacteroides sp. 2_2_4] |
| Q8A0D0 | CUY_3870 | 38.276 | 72 | 3.64E-38 | 152 | conserved hypothetical protein [Bacteroides ovatus SD CMC 3f] |
| Q8A0D0 | CUY_3870 | 40.777 | 72 | 1.90E-37 | 150 | conserved hypothetical protein [Bacteroides ovatus SD CMC 3f] |
| Q8A0D0 | BACEGG_03202 | 38.819 | 35 | 3.80E-38 | 151 | hypothetical protein [Bacteroides eggerthii DSM 20697] |
| Q8A0D0 | HMPREF0106_03704 | 38.276 | 72 | 3.82E-38 | 152 | sialic acid-specific 9-O-acetylesterase [Bacteroides sp. D22] |
| Q8A0D0 | HMPREF0106_03704 | 40.777 | 72 | 2.03E-37 | 150 | sialic acid-specific 9-O-acetylesterase [Bacteroides sp. D22] |
| Q8A0D0 | HMPREF9446_03560 | 36.555 | 35 | 5.34E-38 | 154 | GDSL-like protein [Bacteroides fluxus YIT 12057] |
| Q8A0D0 | BACCOPRO_02673 | 37.037 | 36 | 5.84E-38 | 154 | GDSL-like protein [Bacteroides coprophilus DSM 18228 = JCM 13818] |
| Q8A0D0 | HMPREF1214_02929 | 35.274 | 42 | 2.86E-37 | 152 | sialate O-acetylesterase [Bacteroides sp. HPS0048] |
| Q8A0D0 | BSIG_5696 | 36.017 | 35 | 4.64E-37 | 152 | hypothetical protein [Bacteroides sp. 1_1_6] |
| Q8A0D0 | HMPREF9445_00461 | 35.319 | 35 | 5.65E-37 | 152 | cyclically-permuted mutarotase family protein [Bacteroides clarus YIT 12056] |
| Q8A0D0 | HMPREF9007_01395 | 35.593 | 35 | 8.10E-37 | 151 | sialate O-acetylesterase [Bacteroides sp. 1_1_14] |
| Q8A0D0 | HMPREF1033_01559 | 34.513 | 33 | 1.02E-36 | 147 | hypothetical protein [Tannerella sp. 6_1_58FAA_CT1] |
| Q8A0D0 | BACSTE_01984 | 31.683 | 44 | 1.39E-36 | 151 | cyclically-permuted mutarotase family protein [Bacteroides stercoris ATCC 43183] |
| Q8A0D0 | HMPREF1214_04113 | 36.652 | 33 | 1.54E-36 | 148 | hypothetical protein [Bacteroides sp. HPS0048] |
| Q8A0D0 | BACCELL_00054 | 37.668 | 32 | 1.55E-36 | 145 | hypothetical protein [Bacteroides cellulosilyticus DSM 14838] |
| Q8A0D0 | HMPREF9456_02531 | 38.34 | 37 | 1.79E-36 | 147 | hypothetical protein [Dysgonomonas mossii DSM 22836] |
| Q8A0D0 | BSGG_5000 | 42.211 | 29 | 2.42E-36 | 149 | hypothetical protein [Bacteroides sp. D2] |
| Q8A0D0 | BACUNI_03810 | 38.571 | 67 | 4.39E-36 | 150 | hypothetical protein [Bacteroides uniformis ATCC 8492] |
| Q8A0D0 | BACUNI_03810 | 35.417 | 67 | 5.34E-35 | 146 | hypothetical protein [Bacteroides uniformis ATCC 8492] |
| Q8A0D0 | HMPREF1007_00771 | 38.571 | 67 | 4.51E-36 | 150 | carbohydrate binding module [Bacteroides sp. 4_1_36] |
| Q8A0D0 | HMPREF1007_00771 | 35.833 | 67 | 1.26E-35 | 148 | carbohydrate binding module [Bacteroides sp. 4_1_36] |
| Q8A0D0 | BACDOR_00589 | 35.78 | 32 | 4.95E-36 | 145 | hypothetical protein [Bacteroides dorei DSM 17855] |
| Q8A0D0 | HMPREF0969_02025 | 38.571 | 67 | 5.30E-36 | 149 | hypothetical protein [Bacteroides sp. D20] |
| Q8A0D0 | HMPREF0969_02025 | 35.417 | 67 | 6.11E-35 | 146 | hypothetical protein [Bacteroides sp. D20] |
| Q8A0D0 | BACCAC_01087 | 35.865 | 35 | 6.72E-36 | 149 | cyclically-permuted mutarotase family protein [Bacteroides caccae ATCC 43185] |
| Q8A0D0 | HMPREF9420_2498 | 35.294 | 38 | 1.01E-35 | 147 | GDSL-like protein [Prevotella salivae DSM 15606] |
| Q8A0D0 | RUM_20850 | 36.017 | 35 | 1.71E-35 | 145 | Domain of unknown function (DUF303). [Ruminococcus champanellensis 18P13 = JCM 17042] |
| Q8A0D0 | BSGG_2554 | 40.417 | 64 | 7.35E-35 | 142 | hypothetical protein [Bacteroides sp. D2] |
| Q8A0D0 | BSGG_2554 | 40.722 | 64 | 2.13E-34 | 140 | hypothetical protein [Bacteroides sp. D2] |
| Q8A0D0 | BSIG_3231 | 36.134 | 35 | 1.06E-34 | 144 | hypothetical protein [Bacteroides sp. 1_1_6] |
| Q8A0D0 | HMPREF9007_00774 | 35.294 | 35 | 2.39E-34 | 144 | sialate O-acetylesterase [Bacteroides sp. 1_1_14] |
| Q8A0D0 | HMPREF0106_01306 | 35.772 | 35 | 3.28E-34 | 144 | sialate O-acetylesterase [Bacteroides sp. D22] |
| Q8A0D0 | BRYFOR_09438 | 33.113 | 40 | 3.51E-33 | 140 | glycosyl hydrolase family 2, sugar binding domain protein [Marvinbryantia formatexigens DSM 14469] |
| Q8A0D0 | BACPLE_01375 | 36.15 | 32 | 1.89E-32 | 135 | hypothetical protein [Bacteroides plebeius DSM 17135] |
| Q8A0D0 | HMPREF9447_03525 | 37.705 | 35 | 3.89E-32 | 134 | hypothetical protein [Bacteroides oleiciplenus YIT 12058] |
| Q8A0D0 | HMPREF0969_00863 | 34.583 | 35 | 3.94E-32 | 134 | hypothetical protein [Bacteroides sp. D20] |
| Q8A0D0 | CW3_3166 | 36.946 | 31 | 1.87E-31 | 125 | conserved domain protein [Bacteroides xylanisolvens SD CC 1b] |
| Q8A0D0 | CW1_1878 | 36.946 | 31 | 1.87E-31 | 125 | conserved domain protein [Bacteroides xylanisolvens SD CC 2a] |
| Q8A048 | HMPREF9007_00404 | 99.427 | 100 | 0 | 1081 | rhamnogalacturonan acetylesterase [Bacteroides sp. 1_1_14] |

| | | | | | | |
|---|---|---|---|---|---|---|
| Q8A048 | BSIG_2476 | 100 | 98 | 0 | 1063 | hypothetical protein [Bacteroides sp. 1_1_6] |
| Q8A048 | CUY_0682 | 92.366 | 100 | 0 | 1009 | GDSL-like protein [Bacteroides ovatus SD CMC 3f] |
| Q8A048 | HMPREF9010_02750 | 92.366 | 100 | 0 | 1009 | putative rhamnogalacturonan acetylesterase [Bacteroides sp. 3_1_23] |
| Q8A048 | BSCG_01751 | 92.176 | 100 | 0 | 1008 | GDSL-like protein [Bacteroides sp. 2_2_4] |
| Q8A048 | HMPREF0106_02576 | 92.176 | 100 | 0 | 1006 | rhamnogalacturonan acetylesterase [Bacteroides sp. D22] |
| Q8A048 | BXY_32340 | 91.985 | 100 | 0 | 1006 | hypothetical protein [Bacteroides xylanisolvens XB1A] |
| Q8A048 | HMPREF0102_03382 | 91.794 | 100 | 0 | 1004 | GDSL-like protein [Bacteroides sp. 2_1_22] |
| Q8A048 | CW3_2306 | 91.794 | 100 | 0 | 1003 | GDSL-like protein [Bacteroides xylanisolvens SD CC 1b] |
| Q8A048 | CW1_1607 | 91.794 | 100 | 0 | 1003 | GDSL-like protein [Bacteroides xylanisolvens SD CC 2a] |
| Q8A048 | BACOVA_04972 | 91.412 | 100 | 0 | 996 | GDSL-like protein [Bacteroides ovatus ATCC 8483] |
| Q8A048 | BACFIN_06097 | 91.013 | 99 | 0 | 994 | GDSL-like protein [Bacteroides finegoldii DSM 17565] |
| Q8A048 | HMPREF0127_02083 | 93.191 | 98 | 0 | 994 | hypothetical protein [Bacteroides sp. 1_1_30] |
| Q8A048 | BSGG_1433 | 92.802 | 98 | 0 | 989 | hypothetical protein [Bacteroides sp. D2] |
| Q8A048 | BSAG_03195 | 92.607 | 98 | 0 | 988 | GDSL-like protein [Bacteroides sp. D1] |
| Q8A048 | HMPREF1017_02195 | 92.412 | 98 | 0 | 984 | hypothetical protein [Bacteroides ovatus 3_8_47FAA] |
| Q8A048 | HMPREF9447_04469 | 86.26 | 100 | 0 | 955 | hypothetical protein [Bacteroides oleiciplenus YIT 12058] |
| Q8A048 | BACINT_04767 | 86.233 | 99 | 0 | 953 | GDSL-like protein [Bacteroides intestinalis DSM 17393] |
| Q8A048 | BACCELL_00088 | 86.233 | 99 | 0 | 952 | GDSL-like protein [Bacteroides cellulosilyticus DSM 14838] |
| Q8A048 | HMPREF9445_01758 | 80.492 | 100 | 0 | 919 | GDSL-like protein [Bacteroides clarus YIT 12056] |
| Q8A048 | CUU_3549 | 82.353 | 99 | 0 | 913 | GDSL-like protein [Bacteroides vulgatus PC510] |
| Q8A048 | BSFG_01515 | 82.163 | 99 | 0 | 912 | GDSL-like protein [Bacteroides sp. 4_3_47FAA] |
| Q8A048 | HMPREF9011_03399 | 82.163 | 99 | 0 | 912 | carbohydrate esterase family 12 protein [Bacteroides sp. 3_1_40A] |
| Q8A048 | BACDOR_00395 | 81.973 | 99 | 0 | 912 | GDSL-like protein [Bacteroides dorei DSM 17855] |
| Q8A048 | BSBG_00614 | 81.973 | 99 | 0 | 912 | GDSL-like protein [Bacteroides sp. 9_1_42FAA] |
| Q8A048 | HMPREF0105_1747 | 81.784 | 99 | 0 | 910 | GDSL-like protein [Bacteroides sp. 3_1_33FAA] |
| Q8A048 | BSEG_01662 | 81.594 | 99 | 0 | 908 | GDSL-like protein [Bacteroides dorei 5_1_36/D4] |
| Q8A048 | BACPLE_03132 | 81.369 | 99 | 0 | 902 | GDSL-like protein [Bacteroides plebeius DSM 17135] |
| Q8A048 | BACOVA_02693 | 70.802 | 99 | 0 | 750 | GDSL-like protein [Bacteroides ovatus ATCC 8483] |
| Q8A048 | HMPREF1532_02764 | 65.655 | 100 | 0 | 713 | hypothetical protein [Bacteroides salyersiae WAL 10018 = DSM 18765 = JCM 12988] |
| Q8A048 | HMPREF9442_00977 | 63.776 | 98 | 0 | 673 | GDSL-like protein [Paraprevotella xylaniphila YIT 11841] |
| Q8A048 | HMPREF9441_03355 | 63.269 | 98 | 0 | 669 | GDSL-like protein [Paraprevotella clara YIT 11840] |
| Q8A048 | BACPLE_00256 | 48.374 | 93 | 1.15E-70 | 243 | glycosyl hydrolase, family 88 [Bacteroides plebeius DSM 17135] |
| Q8A048 | BACPLE_00256 | 48.583 | 93 | 2.54E-70 | 242 | glycosyl hydrolase, family 88 [Bacteroides plebeius DSM 17135] |
| Q8A048 | HMPREF9455_02344 | 46.694 | 93 | 1.91E-62 | 221 | hypothetical protein [Dysgonomonas gadei ATCC BAA-286] |
| Q8A048 | HMPREF9455_02344 | 43.478 | 93 | 2.20E-61 | 218 | hypothetical protein [Dysgonomonas gadei ATCC BAA-286] |
