## Supplemental Table S6 for "Chemoproteomic identification of a dipeptidyl peptidase 4 (DPP4) homolog in *Bacteroides thetaiotaomicron* important for envelope integrity and fitness"

TableS6: Protein-BLAST results for human DPP4 in HMP gut reference isolates

| Query sequence | Target sequence | % identity | % coverage | E-value | Bit score | Target sequence description |
|---|---|---|---|---|---|---|
| P27487 | HMPREF9008_02510 | 32.429 | 90 | 1.73E-98 | 324 | peptidase, S9A/B/C family, catalytic domain protein [Parabacteroides sp. 20_3] |
| P27488 | PARMER_03777 | 32.338 | 98 | 1.11E-97 | 322 | peptidase, S9A/B/C family, catalytic domain protein [Parabacteroides merdae ATCC 43184] |
| P27489 | HMPREF0104_02101 | 32.143 | 90 | 3.22E-97 | 321 | dipeptidyl-peptidase IV [Bacteroides sp. 3_1_19] |
| P27490 | HMPREF0619_02972 | 32.143 | 90 | 6.03E-97 | 320 | peptidase, S9A/B/C family, catalytic domain protein [Parabacteroides sp. D13] |
| P27491 | PRABACTJOHN_02440 | 32.208 | 98 | 9.25E-97 | 320 | peptidase, S9A/B/C family, catalytic domain protein [Parabacteroides johnsonii DSM 18315] |
| P27492 | HMPREF0103_2768 | 32 | 90 | 2.55E-96 | 318 | peptidase, S9A/B/C family, catalytic domain protein [Bacteroides sp. 2_1_33B] |
| P27493 | BACSTE_03271 | 33.824 | 86 | 6.63E-93 | 310 | peptidase, S9A/B/C family, catalytic domain protein [Bacteroides stercoris ATCC 43183] |
| P27494 | HMPREF9445_02996 | 33.431 | 86 | 1.01E-91 | 307 | peptidase, S9A/B/C family, catalytic domain protein [Bacteroides clarus YIT 12056] |
| P27495 | HMPREF0969_00958 | 33.333 | 86 | 1.16E-91 | 306 | peptidase, S9A/B/C family, catalytic domain protein [Bacteroides sp. D20] |
| P27496 | HMPREF9456_01479 | 33.184 | 86 | 1.18E-91 | 306 | hypothetical protein [Dysgonomonas mossii DSM 22836] |
| P27497 | BACUNI_03587 | 33.333 | 86 | 1.35E-91 | 306 | peptidase, S9A/B/C family, catalytic domain protein [Bacteroides uniformis ATCC 8492] |
| P27498 | HMPREF1007_03240 | 33.333 | 86 | 1.35E-91 | 306 | dipeptidyl peptidase IV [Bacteroides sp. 4_1_36] |
| P27499 | HMPREF9448_00433 | 30.906 | 86 | 2.26E-91 | 305 | hypothetical protein [Barnesiella intestinihominis YIT 11860] |
| P27500 | HMPREF1532_00929 | 32.362 | 86 | 2.78E-91 | 305 | hypothetical protein [Bacteroides salyersiae WAL 10018 = DSM 18765 = JCM 12988] |
| P27501 | HMPREF1017_02246 | 32.07 | 86 | 9.08E-91 | 304 | hypothetical protein [Bacteroides ovatus 3_8_47FAA] |
| P27502 | BSGG_1381 | 32.07 | 86 | 1.07E-90 | 304 | hypothetical protein [Bacteroides sp. D2] |
| P27503 | HMPREF9010_02811 | 32.07 | 86 | 1.73E-90 | 303 | dipeptidyl-peptidase IV [Bacteroides sp. 3_1_23] |
| P27504 | BSCG_01817 | 32.216 | 86 | 2.20E-90 | 303 | peptidase, S9A/B/C family, catalytic domain protein [Bacteroides sp. 2_2_4] |
| P27505 | HMPREF0106_02634 | 31.76 | 88 | 2.24E-90 | 303 | dipeptidyl-peptidase IV [Bacteroides sp. D22] |
| P27506 | ALIPUT_02419 | 32.625 | 84 | 3.91E-90 | 302 | peptidase, S9A/B/C family, catalytic domain protein [Alistipes putredinis DSM 17216] |
| P27507 | HMPREF9455_01995 | 32.493 | 87 | 4.40E-90 | 302 | hypothetical protein [Dysgonomonas gadei ATCC BAA-286] |
| P27508 | BSIG_2455 | 31.693 | 86 | 5.79E-90 | 303 | hypothetical protein [Bacteroides sp. 1_1_6] |
| P27509 | HMPREF9447_04168 | 33.869 | 86 | 6.39E-90 | 302 | hypothetical protein [Bacteroides oleiciplenus YIT 12058] |
| P27510 | CUY_0743 | 31.617 | 88 | 6.93E-90 | 302 | dipeptidyl peptidase IV N-terminal domain protein [Bacteroides ovatus SD CMC 3f] |
| P27511 | BACOVA_05046 | 31.617 | 88 | 8.49E-90 | 301 | peptidase, S9A/B/C family, catalytic domain protein [Bacteroides ovatus ATCC 8483] |
| P27512 | HMPREF0127_04782 | 31.617 | 88 | 1.03E-89 | 301 | hypothetical protein [Bacteroides sp. 1_1_30] |
| P27513 | BXY_33000 | 31.617 | 88 | 1.03E-89 | 301 | dipeptidyl-peptidase IV . Serine peptidase. MEROPS family S09B [Bacteroides xylanisolvens XB1A] |
| P27514 | HMPREF0102_03488 | 31.617 | 88 | 1.04E-89 | 301 | peptidase, S9A/B/C family, catalytic domain protein [Bacteroides sp. 2_1_22] |
| P27515 | BSAG_01729 | 31.617 | 88 | 1.04E-89 | 301 | peptidase, S9A/B/C family, catalytic domain protein [Bacteroides sp. D1] |
| P27516 | CW3_2404 | 31.617 | 88 | 1.04E-89 | 301 | dipeptidyl peptidase IV N-terminal domain protein [Bacteroides xylanisolvens SD CC 1b] |
| P27517 | CW1_1547 | 31.617 | 88 | 1.04E-89 | 301 | dipeptidyl peptidase IV N-terminal domain protein [Bacteroides xylanisolvens SD CC 2a] |
| P27518 | HMPREF1018_00999 | 32.606 | 86 | 1.64E-89 | 301 | hypothetical protein [Bacteroides sp. 2_1_56FAA] |
| P27519 | BACCOPRO_03615 | 32.181 | 94 | 2.16E-89 | 300 | peptidase, S9A/B/C family, catalytic domain protein [Bacteroides coprophilus DSM 18228 = JCM 13818] |
| P27520 | BFAG_00311 | 32.362 | 86 | 2.57E-89 | 300 | peptidase, S9A/B/C family, catalytic domain protein [Bacteroides fragilis 3_1_12] |
| P27521 | HMPREF9007_00426 | 31.404 | 86 | 6.43E-89 | 299 | dipeptidyl-peptidase IV [Bacteroides sp. 1_1_14] |
| P27522 | HMPREF0101_00028 | 32.46 | 86 | 1.52E-88 | 298 | peptidase, S9A/B/C family, catalytic domain protein [Bacteroides sp. 2_1_16] |
| P27523 | BSHG_0669 | 32.46 | 86 | 2.34E-88 | 298 | hypothetical protein [Bacteroides sp. 3_2_5] |
| P27524 | BACINT_04434 | 33.041 | 86 | 3.89E-88 | 298 | peptidase, S9A/B/C family, catalytic domain protein [Bacteroides intestinalis DSM 17393] |
| P27525 | BACCELL_03148 | 32.945 | 86 | 1.19E-87 | 296 | peptidase, S9A/B/C family, catalytic domain protein [Bacteroides cellulosilyticus DSM 14838] |
| P27526 | HMPREF1214_01399 | 32.996 | 87 | 1.73E-87 | 295 | dipeptidyl-peptidase 4 [Bacteroides sp. HPS0048] |
| P27527 | BACEGG_00881 | 32.701 | 86 | 1.88E-87 | 295 | peptidase, S9A/B/C family, catalytic domain protein [Bacteroides eggerthii DSM 20697] |
| P27528 | BACCAC_01795 | 31.441 | 87 | 6.64E-87 | 293 | peptidase, S9A/B/C family, catalytic domain protein [Bacteroides caccae ATCC 43185] |
| P27529 | HMPREF9450_01296 | 31.102 | 92 | 1.03E-86 | 293 | hypothetical protein [Alistipes indistinctus YIT 12060] |
| P27530 | HMPREF1016_01518 | 32.555 | 86 | 1.15E-86 | 293 | dipeptidyl peptidase IV [Bacteroides eggerthii 1_2_48FAA] |
| P27531 | BACFIN_08852 | 31.655 | 87 | 1.42E-86 | 293 | peptidase, S9A/B/C family, catalytic domain protein [Bacteroides finegoldii DSM 17565] |
| P27532 | BSFG_01970 | 31.242 | 93 | 4.70E-86 | 291 | peptidase, S9A/B/C family, catalytic domain protein [Bacteroides sp. 4_3_47FAA] |
| P27533 | HMPREF9011_04151 | 31.242 | 93 | 4.70E-86 | 291 | dipeptidyl peptidase IV [Bacteroides sp. 3_1_40A] |
| P27534 | HMPREF0105_4156 | 31.108 | 93 | 4.75E-86 | 291 | peptidase, S9A/B/C family, catalytic domain protein [Bacteroides sp. 3_1_33FAA] |
| P27535 | BSEG_03560 | 31.108 | 93 | 5.49E-86 | 291 | peptidase, S9A/B/C family, catalytic domain protein [Bacteroides dorei 5_1_36/D4] |
| P27536 | BACDOR_03064 | 31.108 | 93 | 5.49E-86 | 291 | peptidase, S9A/B/C family, catalytic domain protein [Bacteroides dorei DSM 17855] |
| P27537 | BSBG_03825 | 31.108 | 93 | 5.49E-86 | 291 | peptidase, S9A/B/C family, catalytic domain protein [Bacteroides sp. 9_1_42FAA] |
| P27538 | CUU_1533 | 31.242 | 93 | 6.08E-86 | 291 | putative prolyl dipeptidyl peptidase [Bacteroides vulgatus PC510] |
| P27539 | HMPREF9446_02000 | 31.341 | 86 | 6.09E-86 | 291 | peptidase, S9A/B/C family, catalytic domain protein [Bacteroides fluxus YIT 12057] |

| | | | | | | |
|---|---|---|---|---|---|---|
| P27540 | HMPREF9449_00338 | 29.576 | 95 | 9.33E-86 | 290 | hypothetical protein [Odoribacter laneus YIT 12061] |
| P27541 | BACCOP_03926 | 31.126 | 94 | 2.54E-85 | 290 | peptidase, S9A/B/C family, catalytic domain protein [Bacteroides coprocola DSM 17136] |
| P27542 | BACPLE_03112 | 31.684 | 94 | 1.62E-80 | 276 | peptidase, S9A/B/C family, catalytic domain protein [Bacteroides plebeius DSM 17135] |
| P27543 | HMPREF1033_00581 | 31.571 | 86 | 2.71E-79 | 273 | hypothetical protein [Tannerella sp. 6_1_58FAA_CT1] |
| P27544 | HMPREF9441_02325 | 29.118 | 86 | 4.93E-71 | 251 | peptidase, S9A/B/C family, catalytic domain protein [Paraprevotella clara YIT 11840] |
| P27545 | PREVCOP_04755 | 29.094 | 86 | 1.52E-70 | 249 | peptidase, S9A/B/C family, catalytic domain protein [Prevotella copri DSM 18205] |
| P27546 | HMPREF9442_00881 | 28.655 | 86 | 1.23E-69 | 247 | peptidase, S9A/B/C family, catalytic domain protein [Paraprevotella xylaniphila YIT 11841] |
| P27547 | HMPREF1475_00504 | 28.448 | 88 | 5.94E-68 | 242 | hypothetical protein [Prevotella oralis HGA0225] |
| P27548 | HMPREF0673_01202 | 27.507 | 88 | 4.04E-65 | 234 | peptidase, S9A/B/C family, catalytic domain protein [Prevotella stercorea DSM 18206] |
| P27549 | HMPREF9420_0548 | 28.424 | 85 | 1.21E-60 | 222 | peptidase, S9A/B/C family, catalytic domain protein [Prevotella salivae DSM 15606] |
| P27550 | CUU_2370 | 29.462 | 89 | 7.50E-59 | 217 | dipeptidyl peptidase IV N-terminal domain protein [Bacteroides vulgatus PC510] |
| P27551 | BSFG_00985 | 29.462 | 89 | 1.25E-58 | 216 | peptidase, S9A/B/C family, catalytic domain protein [Bacteroides sp. 4_3_47FAA] |
| P27552 | BSEG_00993 | 30.275 | 84 | 2.48E-58 | 216 | peptidase, S9A/B/C family, catalytic domain protein [Bacteroides dorei 5_1_36/D4] |
| P27553 | BACDOR_00958 | 30.275 | 84 | 3.29E-58 | 215 | peptidase, S9A/B/C family, catalytic domain protein [Bacteroides dorei DSM 17855] |
| P27554 | HMPREF0105_1254 | 30.275 | 84 | 4.96E-58 | 215 | peptidase, S9A/B/C family, catalytic domain protein [Bacteroides sp. 3_1_33FAA] |
| P27555 | BSBG_00102 | 30.122 | 84 | 2.55E-57 | 213 | peptidase, S9A/B/C family, catalytic domain protein [Bacteroides sp. 9_1_42FAA] |
| P27556 | HMPREF9011_03920 | 31.219 | 77 | 5.90E-56 | 206 | dipeptidyl aminopeptidase [Bacteroides sp. 3_1_40A] |
| P27557 | HMPREF9455_02807 | 26.856 | 79 | 1.68E-51 | 196 | hypothetical protein [Dysgonomonas gadei ATCC BAA-286] |
| P27558 | HMPREF1033_01873 | 29.604 | 74 | 1.07E-50 | 193 | hypothetical protein [Tannerella sp. 6_1_58FAA_CT1] |
| P27559 | HMPREF9456_00374 | 25.926 | 78 | 3.30E-49 | 189 | hypothetical protein [Dysgonomonas mossii DSM 22836] |
| P27560 | BACCELL_02178 | 25.523 | 91 | 5.79E-46 | 179 | peptidase, S9A/B/C family, catalytic domain protein [Bacteroides cellulosilyticus DSM 14838] |
| P27561 | BFAG_02901 | 25.623 | 90 | 2.25E-45 | 178 | peptidase, S9A/B/C family, catalytic domain protein [Bacteroides fragilis 3_1_12] |
| P27562 | HMPREF9447_02557 | 24.592 | 99 | 6.05E-45 | 177 | hypothetical protein [Bacteroides oleiciplenus YIT 12058] |
| P27563 | BACINT_01011 | 26.624 | 81 | 1.75E-44 | 176 | peptidase, S9A/B/C family, catalytic domain protein [Bacteroides intestinalis DSM 17393] |
| P27564 | HMPREF0969_01715 | 26.721 | 77 | 2.52E-44 | 175 | peptidase, S9A/B/C family, catalytic domain protein [Bacteroides sp. D20] |
| P27565 | HMPREF9442_02116 | 25.484 | 83 | 2.53E-44 | 175 | peptidase, S9A/B/C family, catalytic domain protein [Paraprevotella xylaniphila YIT 11841] |
| P27566 | HMPREF9441_01209 | 25.714 | 83 | 2.93E-44 | 175 | peptidase, S9A/B/C family, catalytic domain protein [Paraprevotella clara YIT 11840] |
| P27567 | BACUNI_04166 | 26.721 | 77 | 3.52E-44 | 174 | peptidase, S9A/B/C family, catalytic domain protein [Bacteroides uniformis ATCC 8492] |
| P27568 | HMPREF1007_00511 | 26.721 | 77 | 3.52E-44 | 174 | dipeptidyl peptidase IV [Bacteroides sp. 4_1_36] |
| P27569 | HMPREF9008_03751 | 24.766 | 92 | 4.15E-44 | 174 | peptidase, S9A/B/C family, catalytic domain protein [Parabacteroides sp. 20_3] |
| P27570 | HMPREF0104_00231 | 24.357 | 92 | 1.11E-43 | 173 | dipeptidyl aminopeptidase IV [Bacteroides sp. 3_1_19] |
| P27571 | HMPREF0619_00460 | 24.42 | 92 | 2.60E-43 | 172 | peptidase, S9A/B/C family, catalytic domain protein [Parabacteroides sp. D13] |
| P27572 | HMPREF0103_2069 | 24.557 | 92 | 4.26E-43 | 171 | peptidase, S9A/B/C family, catalytic domain protein [Bacteroides sp. 2_1_33B] |
| P27573 | HMPREF9007_03794 | 25.379 | 98 | 5.48E-43 | 171 | dipeptidyl aminopeptidase IV [Bacteroides sp. 1_1_14] |
| P27574 | BSHG_1495 | 26.108 | 78 | 8.94E-43 | 170 | hypothetical protein [Bacteroides sp. 3_2_5] |
| P27575 | HMPREF9445_01935 | 23.964 | 97 | 1.28E-42 | 170 | peptidase, S9A/B/C family, catalytic domain protein [Bacteroides clarus YIT 12056] |
| P27576 | HMPREF1018_00139 | 25.944 | 78 | 2.35E-42 | 169 | hypothetical protein [Bacteroides sp. 2_1_56FAA] |
| P27577 | HMPREF0101_02096 | 26.108 | 78 | 3.09E-42 | 169 | peptidase, S9A/B/C family, catalytic domain protein [Bacteroides sp. 2_1_16] |
| P27578 | BSIG_1417 | 24.461 | 98 | 8.98E-42 | 167 | hypothetical protein [Bacteroides sp. 1_1_6] |
| P27579 | HMPREF1016_00991 | 26.059 | 78 | 1.45E-41 | 167 | dipeptidyl peptidase IV [Bacteroides eggerthii 1_2_48FAA] |
| P27580 | BACSTE_02313 | 23.995 | 96 | 4.02E-41 | 165 | peptidase, S9A/B/C family, catalytic domain protein [Bacteroides stercoris ATCC 43183] |
| P27581 | BACEGG_03258 | 24.487 | 91 | 6.30E-41 | 165 | peptidase, S9A/B/C family, catalytic domain protein [Bacteroides eggerthii DSM 20697] |
| P27582 | HMPREF1532_01808 | 24.255 | 93 | 1.53E-40 | 163 | hypothetical protein [Bacteroides salyersiae WAL 10018 = DSM 18765 = JCM 12988] |
| P27583 | BACCAC_01453 | 23.35 | 98 | 9.46E-40 | 161 | peptidase, S9A/B/C family, catalytic domain protein [Bacteroides caccae ATCC 43185] |
| P27584 | PARMER_02747 | 25.135 | 94 | 2.35E-39 | 160 | peptidase, S9A/B/C family, catalytic domain protein [Parabacteroides merdae ATCC 43184] |
| P27585 | BACCOP_02363 | 25.354 | 78 | 2.56E-39 | 160 | peptidase, S9A/B/C family, catalytic domain protein [Bacteroides coprocola DSM 17136] |
| P27586 | BSGG_2370 | 24.478 | 83 | 5.13E-39 | 159 | hypothetical protein [Bacteroides sp. D2] |
| P27587 | HMPREF9010_00648 | 24.776 | 83 | 5.93E-39 | 159 | putative dipeptidyl aminopeptidase IV [Bacteroides sp. 3_1_23] |
| P27588 | HMPREF0102_00604 | 23.192 | 98 | 7.66E-39 | 158 | peptidase, S9A/B/C family, catalytic domain protein [Bacteroides sp. 2_1_22] |
| P27589 | BSAG_00356 | 23.192 | 98 | 7.66E-39 | 158 | peptidase, S9A/B/C family, catalytic domain protein [Bacteroides sp. D1] |
| P27590 | CW3_2034 | 23.192 | 98 | 7.66E-39 | 158 | dipeptidyl peptidase IV N-terminal domain protein [Bacteroides xylanisolvens SD CC 1b] |
| P27591 | CW1_0738 | 23.192 | 98 | 7.66E-39 | 158 | dipeptidyl peptidase IV N-terminal domain protein [Bacteroides xylanisolvens SD CC 2a] |
| P27592 | HMPREF1017_00530 | 24.627 | 83 | 8.71E-39 | 158 | hypothetical protein [Bacteroides ovatus 3_8_47FAA] |
| P27593 | BACOVA_01349 | 24.478 | 83 | 9.89E-39 | 158 | peptidase, S9A/B/C family, catalytic domain protein [Bacteroides ovatus ATCC 8483] |
| P27594 | BXY_23820 | 23.192 | 98 | 1.08E-38 | 158 | prolyl tripeptidyl peptidase. Serine peptidase. MEROPS family S09B [Bacteroides xylanisolvens XB1A] |
| P27595 | CUY_3404 | 24.814 | 83 | 1.19E-38 | 158 | dipeptidyl peptidase IV N-terminal domain protein [Bacteroides ovatus SD CMC 3f] |

| | | | | | | |
|---|---|---|---|---|---|---|
| P27596 | BSFG_00357 | 23.575 | 95 | 2.10E-38 | 157 | peptidase, S9A/B/C family, catalytic domain protein [Bacteroides sp. 4_3_47FAA] |
| P27597 | HMPREF9011_00439 | 23.575 | 95 | 2.10E-38 | 157 | dipeptidyl peptidase IV [Bacteroides sp. 3_1_40A] |
| P27598 | HMPREF0127_01576 | 22.778 | 98 | 2.15E-38 | 157 | hypothetical protein [Bacteroides sp. 1_1_30] |
| P27599 | HMPREF0106_04714 | 22.778 | 98 | 2.15E-38 | 157 | dipeptidyl aminopeptidase IV [Bacteroides sp. D22] |
| P27600 | CUU_0154 | 23.575 | 95 | 2.66E-38 | 157 | dipeptidyl peptidase IV N-terminal domain protein [Bacteroides vulgatus PC510] |
| P27601 | BACDOR_01902 | 23.705 | 95 | 2.96E-38 | 157 | peptidase, S9A/B/C family, catalytic domain protein [Bacteroides dorei DSM 17855] |
| P27602 | BSBG_00781 | 23.705 | 95 | 2.96E-38 | 157 | peptidase, S9A/B/C family, catalytic domain protein [Bacteroides sp. 9_1_42FAA] |
| P27603 | BACCOPRO_02325 | 25.901 | 73 | 3.60E-38 | 156 | peptidase, S9A/B/C family, catalytic domain protein [Bacteroides coprophilus DSM 18228 = JCM 13818] |
| P27604 | HMPREF0105_2037 | 23.446 | 95 | 3.89E-38 | 156 | peptidase, S9A/B/C family, catalytic domain protein [Bacteroides sp. 3_1_33FAA] |
| P27605 | BSEG_02165 | 23.575 | 95 | 6.02E-38 | 155 | peptidase, S9A/B/C family, catalytic domain protein [Bacteroides dorei 5_1_36/D4] |
| P27606 | HMPREF9420_0906 | 25.244 | 78 | 1.14E-37 | 155 | peptidase, S9A/B/C family, catalytic domain protein [Prevotella salivae DSM 15606] |
| P27607 | HMPREF1214_00573 | 23.673 | 93 | 1.03E-36 | 152 | dipeptidyl-peptidase 4 [Bacteroides sp. HPS0048] |
| P27608 | HMPREF9446_02242 | 24.768 | 76 | 3.85E-36 | 150 | peptidase, S9A/B/C family, catalytic domain protein [Bacteroides fluxus YIT 12057] |
| P27609 | BACPLE_00583 | 24.316 | 82 | 8.95E-36 | 149 | peptidase, S9A/B/C family, catalytic domain protein [Bacteroides plebeius DSM 17135] |
| P27610 | HMPREF9449_01075 | 25.974 | 68 | 1.09E-35 | 149 | hypothetical protein [Odoribacter laneus YIT 12061] |
| P27611 | BSCG_03097 | 31.679 | 34 | 1.93E-31 | 131 | peptidase, S9A/B/C family, catalytic domain protein [Bacteroides sp. 2_2_4] |
