## Supplemental Table S7 for "Chemoproteomic identification of a dipeptidyl peptidase 4 (DPP4) homolog in *Bacteroides thetaiotaomicron* important for envelope integrity and fitness"

Table S7: Bacterial strains, DNA, and primers

| Strain | Name | Source |
|---|---|---|
| *Bacteroides thetaiotaomicron* VPI-5482 | VPI-5482 | Kind gift from the laboratory of Dr. Justin Sonnenburg |
| *Bacteroides thetaiotaomicron* VPI-5482 Δ*tdk* (GenR) | *tdk* | doi: 10.1016/j.str.2008.03.017 |
| *Bacteroides thetaiotaomicron* VPI-5482 Δ*tdk* Δ*BT3254* | RC45 | This study |
| *Bacteroides thetaiotaomicron* VPI-5482 Δ*tdk* Δ*BT4193* | RC56 | This study |
| *Bacteroides thetaiotaomicron* VPI-5482 Δ*tdk* Δ*BT4193* Δ*BT3254* | RC57 | This study |
| *E. coli* S17-1 λ*pir*; *zxx::* RP4 2-(Tetr::Mu) (Kanr::Tn7) λ*pir* | S17-1 λpir | doi: 10.1038/nbt1183-784 |
| *E. coli* DH5α *λpir; F - endA1 hsdR17 (r - m +) supE44 thi-1 recA1 gyrA relA1 Δ(lacZYA-argF)U189 φ80lacZ ΔM15 λpir* | DH5α λpir | doi: 10.1128/JB.187.21.7167-7175.2005 |
| *E. coli* Rosetta 2(DE3) | | Kind gift from the laboratory of Dr. Stefan Sieber |
| *Bacteroides fragilis* | | doi: 10.1016/j.chom.2021.12.008 |
| *Bacteroides uniformis* | | doi: 10.1016/j.chom.2021.12.008 |
| *Blautia producta* | | doi: 10.1016/j.chom.2021.12.008 |
| *Clostridium clostridioforme* | | doi: 10.1016/j.chom.2021.12.008 |
| *Clostridium hathewayi* | | doi: 10.1016/j.chom.2021.12.008 |
| *Clostridium hylemonae* | | doi: 10.1016/j.chom.2021.12.008 |
| *Clostridium scindens* | | doi: 10.1016/j.chom.2021.12.008 |
| *Clostridium symbiosum* | | doi: 10.1016/j.chom.2021.12.008 |
| *Enterococcus faecalis* | | doi: 10.1016/j.chom.2021.12.008 |
| *Enterococcus faecium* | | doi: 10.1016/j.chom.2021.12.008 |
| *Enterococcus hirae* | | doi: 10.1016/j.chom.2021.12.008 |
| *Escherichia fergusonii* | | doi: 10.1016/j.chom.2021.12.008 |
| *Flavonifractor plautii* | | doi: 10.1016/j.chom.2021.12.008 |
| *Parabacteroides distasonis* | | doi: 10.1016/j.chom.2021.12.008 |

| Recombinant DNA | Name | Source |
|---|---|---|
| pKNOCK-*bla*-*ermGb*::*tdk* | pExchange-*tdk* | ATCC |
| Upstream and downstream regions of *B. theta BT3254* in pExchange-*tdk* | pRBC9 | This study |
| Upstream and downstream regions of *B. theta BT4193* in pExchange-*tdk* | pRBC8 | This study |
| pET28a | pET28a | Kind gift from the laboratory of Dr. Stefan Sieber |
| pET28a-BT4193-His | pET28a-BT4193-His | This study |
| pET28a-BT3254-His | pET28a-BT3254-His | This study |

| Primer name | Sequence | Description |
|---|---|---|
| BT4193 upstream_fwd | cgaattcctgcagcccggggcttcctgtggagtttctcccaac | Gibson cloning for BT4193 deletion |
| BT4193 upstream_rev | gatttataaatggctgacagagtgcgca | Gibson cloning for BT4193 deletion |
| BT4193 downstream_fwd | ctgtcagccatttataaatcgttttatattttacttgtttaagggggttaag | Gibson cloning for BT4193 deletion |
| BT4193 downstream_rev | cggccgctctagaactagtggggcacgcaaagagagctttttc | Gibson cloning for BT4193 deletion |
| BT3254 upstream_fwd | cgaattcctgcagcccggggcgatgcggagccgaacatac | Gibson cloning for BT3254 deletion |
| BT3254 upstream_rev | aataaaagcacaatctatagttgacagctcacagc | Gibson cloning for BT3254 deletion |
| BT3254 downstream_fwd | ctatagattgtgcttttatttatctttattggttttaaatcc | Gibson cloning for BT3254 deletion |
| BT3254 downstream_rev | cggccgctctagaactagtggaatgcgagttccgcaaatac | Gibson cloning for BT3254 deletion |
| rBT4193_fwd | actttaagaaggagatatacatgagaaaagtaagtttagcc | Gibson cloning for BT4193 expression |
| rBT4193_rev | tggtggtggtggtggctgccgctgcccaaattgttcaggaagaag | Gibson cloning for BT4193 expression |
| rBT3254_fwd | actttaagaaggagatatacatgagtaaaattggaaaaagag | Gibson cloning for BT3254 expression |
| rBT3254_rev | tggtggtggtggtggctgccgctgcctaggttctgctcaaagtatc | Gibson cloning for BT3254 expression |
| backbone_fwd | ggcagcggcagccaccaccaccaccaccac | Gibson cloning for protein expression |
| backbone_rev | gtatatctccttcttaaagttaaacaaaattatttctagaggggaattgttatccgctc | Gibson cloning for protein expression |
