## Supplemental Table S8 for "Chemoproteomic identification of a dipeptidyl peptidase 4 (DPP4) homolog in *Bacteroides thetaiotaomicron* important for envelope integrity and fitness"

Table S8: Medium recipes

| Medium name | Ingredients | Supplement concentration | Source | Notes |
|---|---|---|---|---|
| BHIS | Brain Heart Infusion | | Millipore 53286 | |
| (for deletions) | Hemin | 5 µg/mL | | |
| | L-Cysteine | 1 mg/mL | | |
| mBHI | BBL Brain Heart Infusion Broth, Modified | | BD 299070 | |
| (for proteomics) | | | | |
| mBHIS | BBL Brain Heart Infusion Broth, Modified | | BD 299070 | |
| (for microbiology) | L-Cysteine | 0.48 µg/mL | | |
| | Hemin | 10 µg/mL | | Add after autoclaving |
| | $NaHCO_3$ | 1.9 µg/mL | | Add after autoclaving |
| Varel-Bryant + 0.5% glucose (VBG) | Mineral 3B Solution (20x) | | | Dilute 1:20 |
| (for microbiology) | $FeSO_4 \cdot 7H_2O$ | 4 µg/mL | | |
| | L-Methionine | 20 µg/mL | | |
| | L-Cysteine | 1 mg/mL | | |
| | Glucose | 5 mg/mL | | Adjust pH to 7.1 prior to autoclaving |
| | Hemin | 10 µg/mL | | Add after autoclaving |
| | $NaHCO_3$ | 2 mg/mL | | Add after autoclaving |
| Mineral 3B Solution (20x) | $KH_2PO_4$ | 18 mg/mL | | |
| | NaCl | 18 mg/mL | | |
| | $MgCl_2 \cdot 6H_2O$ | 400 µg/mL | | |
| | $CaCl_2 \cdot 2H_2O$ | 520 µg/mL | | |
| | $CoCl_2 \cdot 6H_2O$ | 20 µg/mL | | |
| | $MnCl_2 \cdot 4H_2O$ | 200 µg/mL | | |
| | $NH_4Cl$ | 10 mg/mL | | |
| | $Na_2SO_4$ | 5 mg/mL | | Autoclave for long-term storage |
| BHI | Brain Heart Infusion | | BD 237500 | |
| (for communities) | | | | |
| BHIhk | Brain Heart Infusion | | BD 237500 | |
| (for communities) | Hemin | 0.5 µg/mL | | |
| | Vitamin K1 | 0.5 µg/mL | | |
| mGAM | Modified Gifu Anaerobic Medium | | HyServe 05433 | |
| (for communities) | | | | |
